# The organization of visual pathways in the *Drosophila* brain

**DOI:** 10.64898/2025.12.22.696097

**Authors:** Judith Hoeller, Arthur Zhao, Aljoscha Nern, Edward M Rogers, Sandro Romani, Michael B Reiser

**Affiliations:** Janelia Research Campus, Howard Hughes Medical Institute, Ashburn, VA USA

## Abstract

Visual systems across species transform photoreceptor inputs into diverse perceptual representations through hierarchical networks that extract features via parallel pathways. In Drosophila, the optic lobes are layered, retinotopic visual processing centers that contain two-thirds of the brain’s neurons and support diverse visually guided behaviors. Although this architecture has long suggested hierarchical and parallel organization, a system-wide account of how behaviorally relevant visual features are routed and integrated across a complete visual system—in any animal—has remained elusive. The new male fly connectome now provides the synapse-level wiring needed to trace visual information from photoreceptors through the optic lobes and across the central brain. Applying a network-based analysis of information flow, we reveal a multi-layered architecture organized into distinct, functionally interpretable pathways. Using this framework to propagate signals through these pathways predicts receptive-field structure and feature selectivity consistent with physiological data, enabling large-scale functional annotation of thousands of neuron types. We find that distinct visual input channels are broadly distributed throughout the brain, yet converge in focal regions of feature specificity and acute spatial vision. Together, these analyses provide a neuron-level, connectome-based view of how a brain organizes and transforms visual input.

## Introduction

The structure of the visual world shapes the neural architectures that interpret it^1,2^. To preserve spatial relationships while extracting diverse features, visual systems across species converge on organizing principles that link neuroanatomy with computation: retinotopic, parallel, and hierarchical processing, with the canonical model of the primate visual cortex^3–5^. Capturing the richness of visual features further demands diverse neural computations and, consequently, specialized cell types^6^. Importantly, these principles often emerge even in circuits with extensive recurrence, where layered functional organization coexists with dense lateral and feedback connectivity. Understanding how such orderly transformations arise from highly interconnected networks remains a central challenge in systems neuroscience^7–9^.

Vision also serves a broad range of non-spatial, evolutionarily conserved functions^10^, suggesting that visual circuits must process information across distinct spatial, temporal, and spectral domains. In the compact yet remarkably elaborate neural network of the *Drosophila* optic lobes, all of these principles are clearly embodied through a striking diversity of neuron types arranged in precise retinotopic maps across layered circuits^11–13^. However, most fly visual neurons remain functionally uncharacterized, and unlike in mammals, where much of vision lies beyond the retina, the extent of visual processing beyond the fly’s optic lobes is unknown^14^. In particular, the functional depth of the hierarchy across the optic lobes and central brain has remained unclear.

Outside of the well-studied visual motion pathways^15–17^, the logic by which visual information is transformed, integrated across feature channels, and routed through the central brain is largely unresolved. Even the definition of what constitutes a “visual system” remains open: a functional description that identifies where visual information is used and transformed has been missing.

These gaps, together with the need to link anatomical pathways to visual computation, motivate this study. Bridging them requires systematic, connectome-based computational approaches that make explicit how network diversity and structure support the organization, routing, and computation of visual features^18–20^.

In *Drosophila*, recent Electron Microscopy reconstructions of the optic lobes^11,21^, the central brain^22^, and most recently the complete central nervous system of both male^23^ and female^24^ flies now provide detailed neuron-to-neuron wiring diagrams (connectomes) linking photoreceptors to higher brain areas. The new male connectome spans the entire brain, including the optic lobes, central brain, and ventral nerve cord (**Fig. 1a,d**), and extends the extensive proofreading and annotation of the right optic lobe^11^. The right-side optic lobe (OL) contains processes of ∼52k neurons across ∼700 cell types, with a balanced composition of excitatory and inhibitory types (**Fig. 1e**). These neurons can be grouped into visual inputs, visual interneurons (VINs), whose arbors lie entirely within the optic lobe, visual centrifugal neurons (VCNs), which relay feedback from the central brain, and Visual Projection Neurons (VPNs) which primarily send information into the central brain.

**Fig. 1.**
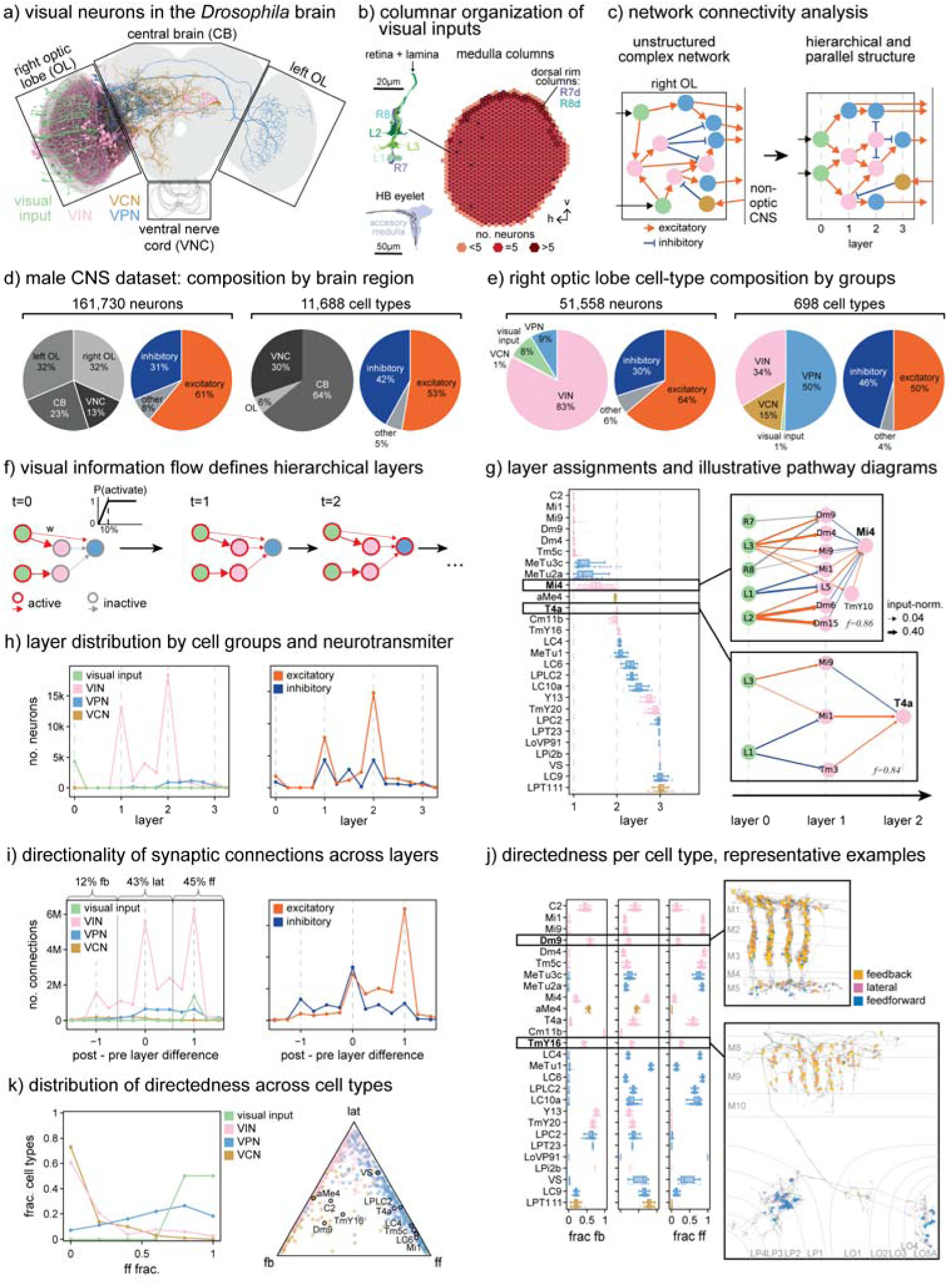
Quantifying hierarchical structure across the optic lobe network. **a)** Random subsample (∼1%) of neurons innervating the right optic lobe, colored by group: visual inputs, visual interneurons (VINs), visual projection neurons (VPNs), and visual centrifugal neurons (VCNs). Boxes indicate brain regions. **b)** All visual input neurons, except HB eyelet, can be assigned to one of the 892 medulla columns, represented as hexagons in a coordinate system where high h corresponds to anterior and high v to dorsal positions in the visual field. Each column typically contains five inputs (L1, L2, L3, R7, and R8), with dorsal rim columns containing R7d and R8d instead of R7 and R8. Shown here is the column map for the right optic lobe^11^, a similar map has been made for the left optic lobe^23^. **c)** The hidden hierarchical and parallel structure of the optic lobe network is revealed with network analysis. This schematic shows representative neurons (dots) colored by their group and connections (arrows) colored by the neurotransmitter of the presynaptic neuron. Acetylcholine was treated as excitatory; GABA and glutamate as inhibitory; others as “other.” Visual processing begins with the visual inputs (black arrows), outputs from the optic lobe are conveyed to other brain regions. **d-e)** Counts of neurons and cell types from the male CNS dataset^23^ included in our analysis, split by brain region and neurotransmitter type (“other” in grey). The optic lobe summary is based on the right-side inventory^11^. **f)** Hierarchy defined by information-flow algorithm^18^. Activation is passed down stepwise from the visual inputs (t = 0) via a nonlinearity (above the first black arrow). Each neuron’s mean activation step defines its hierarchical “layer.” **g)** Representative layer assignment for individual neurons, grouped by cell type. For boxed examples, the dominant activating pathways are shown to the right. Arrow thickness scales with input-normalized connection number and dashed vertical lines mark integer layers. f indicates the fraction of total pathway weights displayed. **h)** Distribution of neuron layers by cell group (left) and neurotransmitter type (right). **i)** Distribution of layer differences across synaptic connections, split by cell group (left) and neurotransmitter type (right). Solid vertical lines separate feedback (fb), lateral (lat), and feedforward (ff) connections; percentages of each (not split by group) are shown. **j)** Representative fractions of fb, lat, and ff connections for individual neurons, grouped by cell type. Boxed examples show spatial distribution of fb (orange), lat (pink), and ff (blue) synapses for one neuron of each type. **k)** Normalized distribution of feedforward fractions across cell types by group (left) and ternary plot of fb, lat, and ff fractions across all right-side optic lobe cell types (right). Points near one edge reflect dominance of two connection types, points near a vertex reflect dominance of one connection type. See also **Fig. S1** and **S2**.

The visual system offers a unique advantage for interpreting structure-function relationships in connectome data: its organization is anchored to the geometry of the eye, so the origin and routing of visual information can be mapped onto a well-defined spatial coordinate system^11,25^. A necessary step in analyzing the flow of visual information is to designate the visual input neurons that seed our analysis and serve as functionally distinct channels (**Fig. 1b**; see Methods). These inputs include the lamina monopolar cells L1, L2, and L3, the inner photoreceptors R7 and R8 and their specialized dorsal rim counterparts, R7d and R8d, and the extra-retinal HB eyelet.

Because neural superposition in the fly eye merges upstream R1-R6 photoreceptor signals from different eye facets in the lamina ^13,26,27^, L1-L3 are the most natural and informative columnar units for seeding the propagation. L1 and L2 convey contrast signals and drive the ON- and OFF-luminance pathways^28–33^; L3 provides luminance-level information^34^, R7 and R8 detect short- and long-wavelength light^35–37;^ R7d and R8d detect perpendicular orientations of polarized UV light^38,39^; and the HB eyelet supplies a non-spatial visual input that photoentrains the circadian clock^40^. Because L1-L3 and R7-R8 map one-to-one onto visual columns (**Fig. 1b**), they preserve the eye’s geometry and allow visual feature transformations to be traced throughout the brain.

A thoroughly annotated connectome is still a dauntingly complex network, so quantitative approaches are essential to reveal global network structure (**Fig. 1c**). To capture how visual signals propagate through the brain, we applied a network-based analysis of information flow^18^ to the new male central nervous system (CNS) connectome. This assigns each synaptic connection a hierarchical depth and direction, classifying it as lateral (within a layer), or as feedforward or feedback (across layers), yielding a data-driven description of hierarchical organization. By deriving a functional hierarchy directly from connectivity, this framework can separate the dominant feedforward pathways from the surrounding recurrent structure, offering an interpretable way to compare sensory circuit organization across systems.

Given the scale of the dataset—over 160k neurons across ∼12k cell types, with the optic lobes contributing ∼64% of the cells **(Fig. 1d**)^23^—we introduce *visual pathways* as an intermediate level of description that captures how information flows through the connectome. To develop and validate this approach, we first apply it to the right optic lobe, and define visual pathways, as strongly connected sets of neurons linking visual inputs and terminating in VPNs (**Fig. 2**). Many VPNs share similar indirect input patterns, which can be grouped into distinct pathway classes. This operational definition of visual pathways, derived directly from the connectome, makes pathways measurable computational units of anatomy, with inferred functional properties. In primate vision, pathways have historically referred to routes of information flow, such as the dorsal and ventral visual streams, which differ in anatomical connectivity and in the functional computations they support^4,41^. Our notion of pathways also parallels the notion of “modules” or “communities” often studied in network science, which are sets of (in our case, overlapping) nodes with dense internal connectivity and likely related function^42^.

**Fig. 2.**
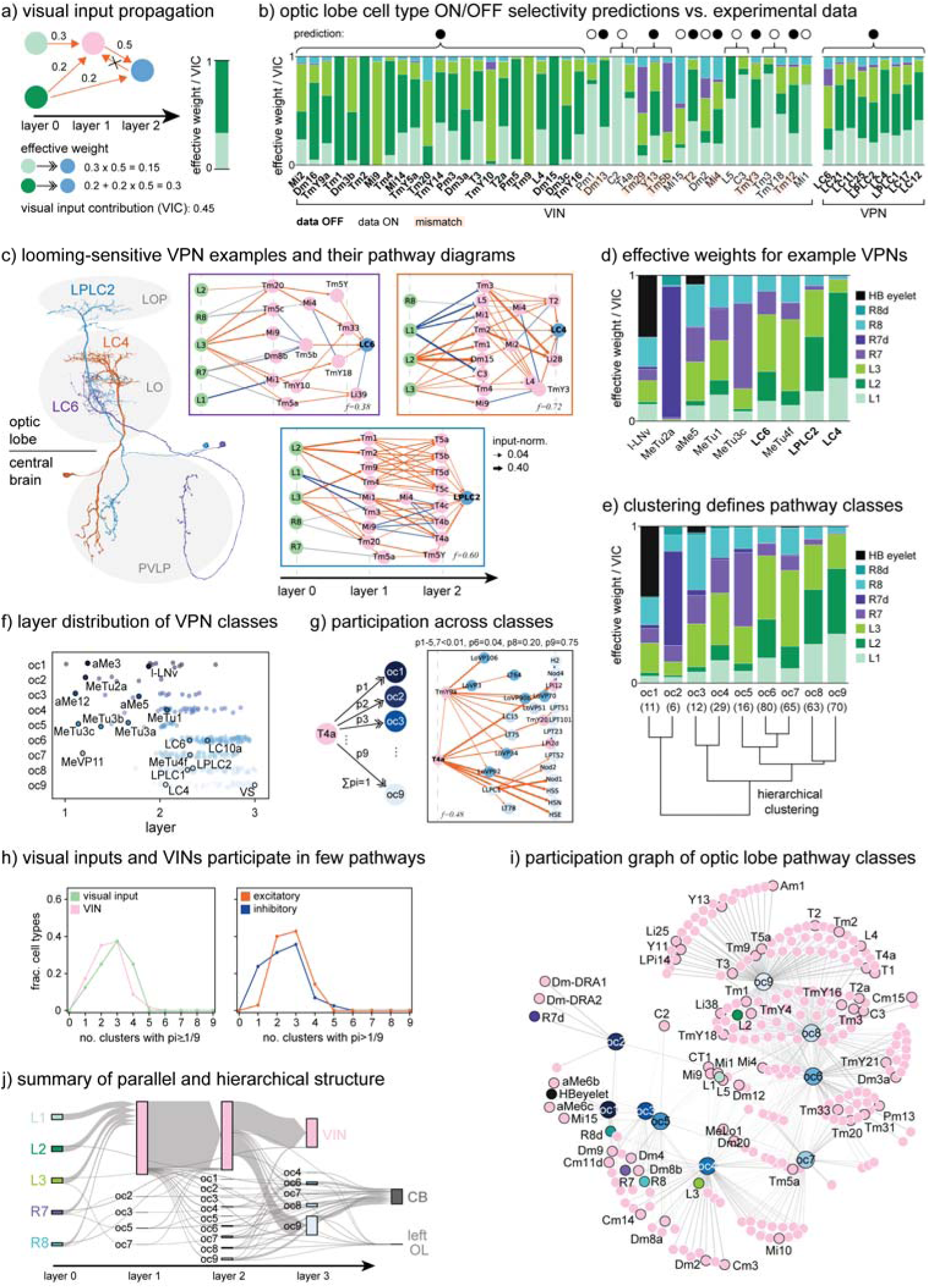
Forward-directed pathways linking visual inputs to optic lobe outputs. **a)** Visual inputs may connect indirectly to VPNs via VINs. The strength of these indirect connections is given by the *effective weight*, computed (bottom) on a trimmed, acyclic graph. The sum of effective weights from all visual inputs defines the *visual input contribution* (VIC). Relative effective weights are visualized as bar plots (right). **b)** Comparison between predicted and measured ON/OFF selectivity. Predictions (top) are based on whether L1 or L2 provides the larger effective weight, indicated by white (ON) or black (OFF) dots, respectively. Experimental calcium imaging data^52,55,57^ are used for experimental validation. OFF (ON) dominance is indicated by a bolded (unbolded) cell type name, mismatches between prediction and data indicated in red. The two blocks correspond to VINs and VPNs. **c)** VPN cell types LC4, LC6, and LPLC2 are known to be looming-sensitive^45,52,55,60^. Despite differing morphologies (left), their pathways (right) reveal shared visual inputs and some overlapping VINs. **d)** Relative effective weights for representative examples of VPN cell types. Each bar represents the normalized contribution from the eight visual-input channels. **e)** Hierarchical clustering of relative effective weights for all VPN cell types defines nine optic lobe pathway classes (oc1-oc9). The dendrogram (bottom) and averaged effective weight profiles (top) summarize the classes; numbers in parentheses indicate the number of cell types per class. Each example in **d** represents one cluster, ordered as in **e**. **f)** VPN types in each pathway class are plotted by their hierarchical layers, colored by class. **g)** Quantification of visual input and VIN contribution to VPN clusters. Effective weights for each cell type were computed as in **a**, then output-normalized to yield a participation fraction vector, pi for i=1-9 (for each class). Example: T4a neurons participate mainly in oc8 and oc9. **h)** Distribution of the number of pathway classes with pi>1/9, for visual inputs and VINs, split by neuron group and neurotransmitter type. A uniform distribution (pi=1/9≈0.11) would peak at 9. **i)** Graph visualization of pathway class participation within the optic lobe. Edges indicate pi>1/9 between cell types (small nodes, colored by cell group) and VPN classes (large nodes, shades of blue). **j)** Sankey diagram summarizing right optic lobe signal flow from visual inputs through VINs and VPN classes (oc1-oc9) to the central brain and left optic lobe. Only connections with positive layer differences are shown. See also **Fig. S3**.

By combining complete connectivity, defined sensory inputs, spatial mapping, and feature propagation, the analysis of the fly visual system is uniquely suited for moving from an anatomical analysis towards functional inferences, including an examination of spatial representation within optic lobe circuits. The retinotopy of the optic lobes constrains the function of many neurons: small neurons engage in mainly local processing, whereas large tangential cells combine information across space. We strengthen this intuition with quantitative signal propagation through successive layers, demonstrating that this predicts receptive fields for many neurons more accurately than direct anatomical measurements (**Fig. 3**).

**Fig. 3.**
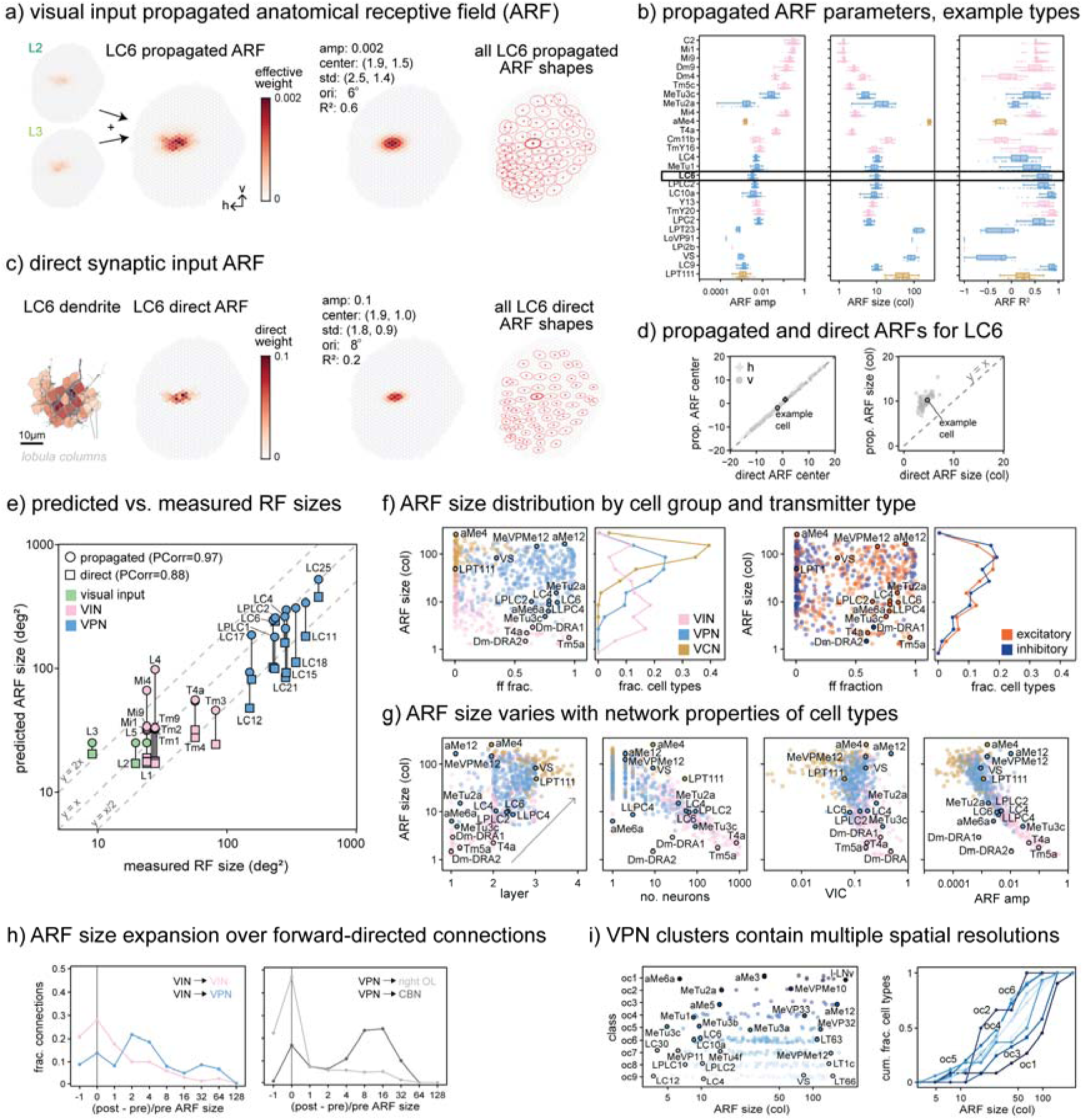
Connectome-based predictions of anatomical receptive fields in the optic lobe. **a)** Propagated visual inputs (Fig. 2a) are summed in each medulla column (Fig. 1b) to yield an anatomical receptive field (ARF). Panels **a**,**c**, and **d** show an example LC6 neuron (body ID 30134). The left panel shows effective weights from L2 and L3, the dominant visual inputs for LC6; the full propagated ARF includes all visual inputs (except for HB eyelet and R7d, R8d). The ARF is fit with a 2D Gaussian to extract its center, size, and shape. Amplitude (*amp*) is the largest effective weight across columns, *std* the standard deviation along the two principal axes, *ori* the orientation of the major axis with respect to -h, and R^2^ the goodness of fit. Right: ARFs of all LC6 neurons displayed tiling the medulla’s columns; crosses mark centers, and ellipses indicate one std (bolded for the example neuron). **b)** Propagated ARF parameters for individual neurons grouped by cell type; size is reported as ellipse area in units of columns (col). **c)** Direct synaptic inputs can also be used to estimate an ARF. The input synapses onto the dendrite of the example LC6 neuron are mapped across lobula columns (left, color represents the fraction of synapses per column). This direct ARF is fit with a 2D Gaussian (right), analogous to **a**. **d)** Scatter plot comparison of propagated and direct ARF parameters (centers and sizes) for all LC6 neurons. The example cell is highlighted; dashed lines indicate equality. **e)** Comparison of predicted and measured receptive field (RF) sizes. Predictions from propagated (circles) and direct (squares) ARFs are connected by lines for each cell type. Experimental calcium imaging data are compiled from multiple studies^53,55,70–72^. Dashed lines indicate equality and two-fold deviations. Propagated ARF predictions correlate more strongly with the measured data (larger Pearson correlation). **f)** Feedforward fraction versus propagated ARF size and normalized ARF size distributions for optic lobe groups (left) and neurotransmitter type (right). **g)** Relationships between propagated ARF size and layer, cell-type abundance, visual input contribution (VIC), and ARF amplitude. **h)** Distribution of relative ARF-size change between connected neurons: from VINs to VINs/VPNs (left), and from VPNs to right optic lobe neurons or central brain neurons (CBNs). The distributions are weighted by connection count. **i)** Each VPN pathway class spans a broad range of ARF sizes (left), indicating multi-resolution visual processing. The right panel shows the cumulative layer distribution for each VPN class. For all panels except **h**, only connections with positive layer difference are included.

Building on the analysis established in the optic lobe, we next extend it to the central brain by mapping and comparing pathways, while detailing the cell types and brain areas involving in routing visual information (**Fig. 4**). This periphery-anchored propagation reveals how the visual input channels (**Fig. 1b**) are preserved and recombined into higher-order visual representations and identifies where this information goes. In essence, our approach uses what is known—both the functional identities of visual inputs and their spatial organization—to inform what remains uncharacterized, providing functional annotations of thousands of neurons across the central nervous system. Finally, we use the propagated spatial information to determine where high-acuity spatial vision is supported and detail the representation of visual space across the fly brain (**Fig. 5**).

**Fig. 4.**
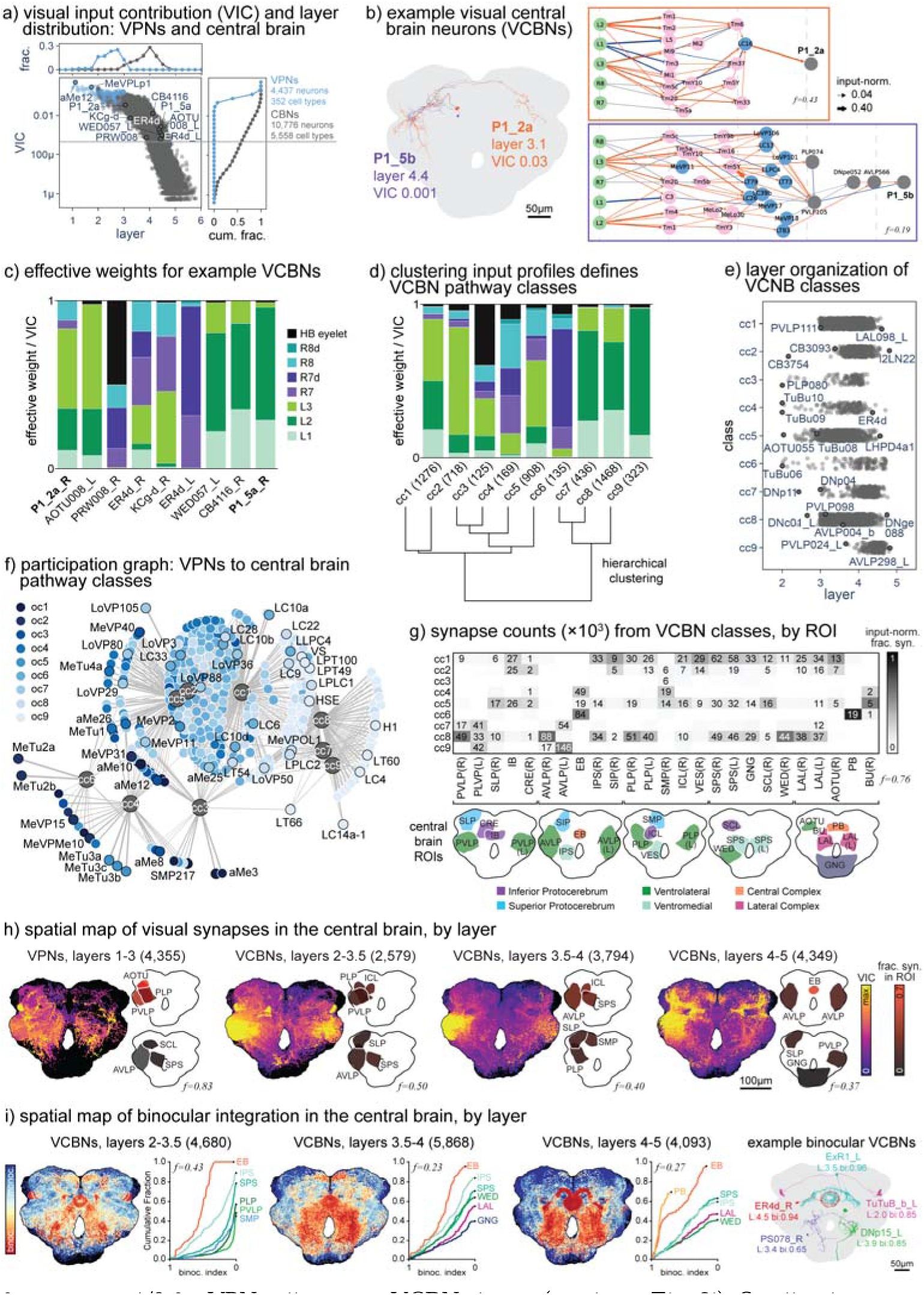
Mapping the flow of visual information into the central brain. **a)** Visual input contribution (VIC, Fig. 2a) of VPN and central brain neuron (CBN) types versus the cell type’s assigned layer based on right-side optic lobe propagation. The marginal histograms show the normalized layer distribution for neurons with VIC > 5X10^-4^ (top) and the cumulative VIC distribution (right). CBN types with VIC > 5X10^-4^ are defined as visual central brain neurons (VCBNs). Numbers above the solid line indicate the counts of neurons and cell types that meet this threshold. **b)** Example VCBNs with their morphologies (left) and pathway diagram (right). Shown are two representative P1 neurons (P1_2a: body ID 15329; P1_5b: body ID 29483), known for their involvement in courtship, but whose specific visual partners and feature selectivity were not generally known. **c)** Relative effective weights from the eight visual inputs for selected VCBN cell types, including the two P1 neurons in **b**. **d)** Hierarchical clustering of the relative effective weights of all VCBN cell types (analogous to Fig. 2e) yields nine central brain clusters, that define pathway classes (cc1-cc9). Bars show the mean relative weights, and the numbers in parentheses denote the number of cell types in each cluster. Each example in **c** represented one cluster, presented in the same order. **e)** Layer distribution of VCBN cell types split by class. **f)** Graph representation of participation fractions pi>1/9 for VPN cell types to VCBN classes (similar to Fig. 2i). Smaller dots correspond to VPN cell types (colored by their VPN cluster oc1-oc9), larger dots correspond to the VCBN clusters. **g)** Number of synapses (in thousands) from VCBNs split by central brain region of interest (ROI) and pathway class. Grayscale intensity indicates the fraction (summing to 1 in each column) of synapses from a class in each ROI. Only fractions > 0.1 are shown; schematics below outline ROI locations. **h)** Projected heatmaps show the spatial distribution of all central brain VPNs and VCBNs presynapses, colored by VIC and separated by layers (as indicated). The brain was collapsed to two dimensions using a maximum-intensity projection, which for each pixel averages the top 10% of VIC values. For each map, the six ROIs with the highest VPN/VCBN synapse counts are listed, and f reports the fraction of all VPN/VCBN presynapses contained within the highlighted ROIs. **i)** Binocular integration across the central brain. As in **h**, projected heatmaps show the 2D spatial distribution of VCBN presynapses based on propagation from both the right and left optic lobes. Color encodes the *binocularity index*, defined for each neuron as the ratio of the smaller to the larger VIC (0=monocular, 1 = fully binocular). Layer assignments shown here use the smaller of the two layer values from left- and right-side propagation. Cumulative plots show the distribution of binocularity indices for the VCBN synapses within the six ROIs with the highest number of binocular synapses (binocular index > 0.5). On the right, five example neurons (body IDs 12656, 520699, 20980, 12069, 31958) show how the layer and binocularity values manifest at the single-neuron level. See also **Fig. S4, S5** and **S6**.

**Fig. 5.**
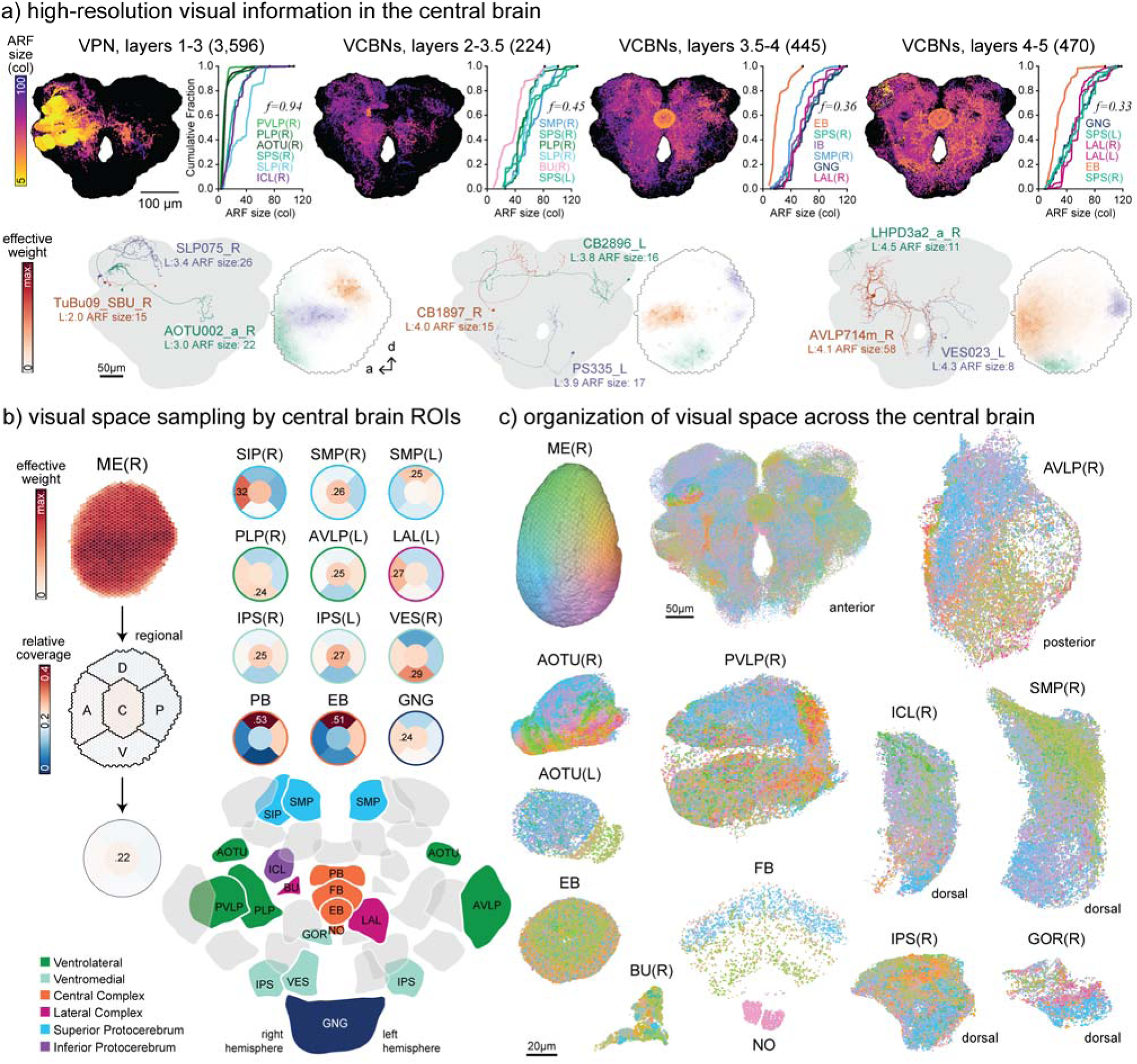
Spatial vision in the central brain. **a)** Extreme-value projections show the 2D spatial distribution of presynaptic sites from VPNs and VCBNs with well-fit anatomical receptive fields (R² > 0.05), colored by ARF size (smaller = higher resolution). Projections as in Fig. 4h, but averaging the 10% smallest ARF values along each projection ray. Neurons are separated by hierarchical layer (as indicated), and the number of neurons in each group is noted in parentheses. Cumulative distributions (right) summarize the ARF sizes for neurons in each group for the six ROIs with the largest numbers of synapses from neurons above the VIC and R² thresholds, and *f* reports the fraction of all above threshold VPN/VCBN presynapses for each grouping, contained within the highlighted ROIs. Below, representative high-resolution VCBNs (Body IDs 17594, 80342, 28227; 53437, 49437, 152122; 87239, 12282, 25289) are shown together with their ARFs and layer and ARF sizes. **b)** To quantify the sampling of visual space, propagated visual inputs from VPNs/VCBNs contributing presynapses to each ROI were grouped into five visual-field sectors—dorsal, ventral, anterior, posterior, and central. The medulla (left) shows a slight increase in the central sector (0.2 corresponds to uniform sampling; maximal value indicated). Sector fractions for the representative ROIs (right; larger set in **Fig. S7e**), highlight consistent biases toward specific portions of the visual field. ROIs are arranged in approximate anatomical order but separated to maximize visual clarity. **c)** To visualize the organization of visual space across the central brain, presynaptic sites of VPNs and VCBNs (ARF R² > -0.5; additional details in **Methods**) are shown in 3D orthographic projection and colored by each neuron’s visual field center. The color map is derived from retinal coordinates and applied to medulla columns (upper left). The full central brain is shown from the anterior view, and selected ROIs are displayed at a common, higher magnification scale (20 µm scale bar) in anterior view unless otherwise indicated. Together, these views illustrate the diversity of spatial organizations across the brain, including dorsal-ventral biases in midline structures, layered structure in the fan-shaped body, and continuous map-like representations in the AOTU. The key in **b** shows the relative spatial positions of the ROIs. See also **Fig. S7**.

This analysis allows long-standing principles of vision to be tested quantitatively and comprehensively:

1. Dominance of feedforward computation: how much of visual processing is carried out by unidirectional transmission versus feedback and lateral refinement?
2. Hierarchical progression: does visual processing advance from local feature detection to neurons encoding larger, more complex receptive fields?
3. Organization into parallel pathways: are distinct feature channels (contrast polarity, luminance, color, polarization, and non-spatial inputs), established at the periphery and propagated within dedicated pathways, or do they instead converge across processing stages?
4. Spatial representation in higher-order visual areas: is retinotopy and high-acuity vision preserved in deeper brain areas?

In the sections that follow, we trace visual information through the fly brain, addressing each of these questions in turn. Together, our analyses provide a quantitative foundation for understanding what constitutes a visual system in fully mapped brain, and how a compact nervous system extracts, combines, and distributes visual information from the retina to the circuits that guide behavior.

## Results

### Quantifying hierarchical structure across the optic lobe network

We first applied this analysis to the right optic lobe, the entry point of visual information into one hemisphere of the brain, to quantify how visual signals propagate from photoreceptors through successive processing layers. Neurons were organized into a hierarchy reflecting the order of signal transmission from visual inputs to downstream targets. Each neuron was assigned a *layer*, defined as the average number of synaptic steps required for activation following a simulated activation of all visual inputs. These activation steps do not represent real time but rather the number of synaptic connections that need to be crossed for activity to reach a neuron through the connectome. Layers capture each neuron’s relative position within the visual processing stream, analogous to layers in artificial feedforward neural networks^43^.

We computed layers by extending an information-flow algorithm originally developed for the fly olfactory system^18^. In this model, each neuron has an activation probability, a bounded activation variable, that accumulates deterministically at each discrete step. This accumulation is computed from the activation of its input neurons through a threshold nonlinearity. The threshold value, the only free parameter in the model, is set (to 10%) so that the motion-sensitive T4 neurons, known to be two synapses downstream of L1 and L3, reach their expected activation at the second step, thereby calibrating the model. Each neuron’s layer is then defined as the average number of steps required to activate it (**Fig. 1f**).

The visual inputs are defined to occupy layer 0. With the calibrated model, all four T4 types fell into layer 2, by design, and well-known downstream partners such as VS neurons^44^ were assigned to layer 3 (T4a and other examples shown in **Fig. 1g**). Most optic lobe interneurons, the VINs, were assigned near-integer layer values (**Fig. 1h**), indicating activation by a single dominant path or multiple paths of the same length. Some neurons fell between integer layers because they are activated along paths of different lengths from the visual inputs. There are some cell types with broad layer distributions across individual cells, which are due to broader distributions in input-normalized synaptic connections, especially for values around the threshold. For example, Mi4 neurons have input-normalized synaptic connections from R8 that range from 0.001 (resulting in assignment to layer 2) to 0.177 (layer 1). Because layer distributions were tight for most cell types, we summarized each by its median layer. The visual cell type groups have distinct layer assignments (**Fig. S1a**) with the more numerous cell types generally assigned to earlier layers (**Fig. S1e**). Layers were treated as continuous for analysis and rounded to integers only for visualization.

To visualize the hierarchical relationships between cell types, we use standardized pathway diagrams. These diagrams show the strongest connections linking visual inputs to specific target neurons (e.g. Mi4 and T4a in **Fig. 1g**), with arrow thickness proportional to the input-normalized connection strength. To provide quantitative context, we introduce the displayed fraction (t), quantifying the fraction of total effective connection strength captured by the displayed paths.

While the optic lobe comprises more than 20 classical anatomic layers (**Fig. S1i-j**)^11,12^, the information-flow analysis revealed a shallow hierarchy within the right optic lobe (**Fig. 1h**): all VINs and VPNs are contained within three processing layers beyond the visual input layer, with only some VCNs extending deeper (**Fig. S1a**). Most VPNs are in layers 2-3 but some are earlier (e.g. MeTu3c, MeTu2a in **Fig. 1g**). This compact, three-layer architecture supports the idea that short synaptic pathways account for the rapid visual processing times observed in electrophysiological recordings, where signals reach VPNs within tens of milliseconds^45–47^. We next examined the distribution of excitatory and inhibitory neurons across layers. This analysis is enabled by accurate neurotransmitter predictions and extensive ground-truth data for the optic lobes^11^. The distributions of excitatory and inhibitory cell types are similar across layers (**Fig. S1a**), although the large population of excitatory motion-sensitive columnar neurons (∼7k T4/T5 cells) skews the ratio of total neurons in layer 2 (**Fig. 1h**). This similarity in excitatory and inhibitory distributions may underlie dynamical stability, often referred to as E/I balance^48^.

To examine the connectivity patterns underlying this hierarchy, we categorized synaptic connections based on the layer difference of their connecting neurons: feedforward (>0.5 layers deeper), lateral (within ±0.5 layers), and feedback (<-0.5; **Fig. 1i**). Across all synaptic connections between optic lobe neurons, 45% are feedforward, 43% lateral and 12% feedback connections. This pattern indicates that recurrent computation is pervasive within the optic lobe. Across the four optic lobe groups, visual inputs make connections that were 97% feedforward, while VINs and VPNs have similar distributions (nearly 50% feedforward and > 40% lateral), whereas VCN connections were mostly feedback (53%), consistent with their expected role in top-down modulation (**Fig. S1b**). As may be expected from canonical neural circuit motifs^49^, feedforward connections are mainly excitatory (81%), lateral synapses are balanced across excitatory and inhibitory neurons, and feedback synapses are predominantly inhibitory (69%; **Fig. 1i, S1b**).

Although these group-level trends define the overall flow of information, individual cell types vary widely in *directedness*, defined as the fractions of feedforward, lateral, and feedback connections that together sum to one and describe each neuron’s directional bias in the network. For example, the interneuron Mi1 is 86% feedforward, while most other VINs are dominated by lateral and feedback connections (**Fig. 1k,j**). Both VCNs and VINs include many minimally feedforward cell types, with VCNs showing higher feedback fractions (**Fig. S1c**). VPNs generally exhibit feedforward connections, though notable exceptions exist (e.g., LPT23 and LPC2, both > 60% feedback). Overall, excitatory neurons tend to favor feedforward transmission, whereas inhibitory neurons more often participate in lateral or feedback interactions (**Fig. S1d**). However, some inhibitory types (Mi9, Tm5c, Dm4, LoVP91) are almost purely feedforward, while some excitatory neurons (Cm11b, TmY20, aMe4) are predominantly feedback (**Fig. 1k,j**). A comprehensive survey of layers and directedness across optic lobe cell types is provided in **Fig. S2**, with the full inventory summarized in **Table S1**. Earlier studies often inferred whether a cell type was primarily feedforward or feedback based on morphology and innervation patterns, but such inferences are difficult to confirm without synaptic connectivity and a systematic notion of directionality. Our connectome-based directedness metrics confirm many of these expectations (e.g., the feedback role of C2 neurons^12,33^) and also reveal distinctions within groups of cells thought to play similar roles. For example, among the well-studied T4 input neurons, Mi1, Mi4, and Mi9 share similar morphologies^12,50^, yet only Mi4 shows an unexpectedly strong mixture of lateral and feedback outputs. Likewise, among Dm cells, whose presynaptic distributions span one or a few adjacent layers, directedness cannot be reliably predicted from anatomy alone, and clear differences emerge even between closely related types, such as Dm9 and Dm4^11,12,51^. Finally, although Tm and TmY neurons are often characterized as feedforward^11,12^, several, including TmY16, exhibit substantial lateral or feedback components. The mapping of directness also shows individual cell types serving complex, region-specific roles: the same neuron may form predominantly lateral and feedback connections in one area and a different mixture of output connections elsewhere (e.g. Dm9 across medulla layers and TmY16 across neuropils). These examples illustrate how connectome-derived directedness can uncover functional diversity that is difficult to anticipate from morphology alone.

Taken together, these analyses reveal an emergent three-layer architecture (**Fig. 1c**, right), with VINs dominating layers 1-2, VPNs populating much of layers 2-3, and VCNs forming the deepest layers (**Fig. S1f**). Although recurrent interactions, especially inhibitory ones, are extensive at every stage (**Fig. S1g**), a large fraction of visual signals flow through a compact set of forward-directed routes. To describe how visual information traverses the fly brain we therefore focus on these forward-directed pathways.

### Forward-directed pathways linking visual inputs to optic lobe outputs

We next ask how visual input signals are transformed within the optic lobe and relayed to visual projection neurons (VPNs). Using the layer hierarchy as a scaffold, we isolate the forward-directed routes carrying features established by the visual inputs (contrast, luminance, color, polarization, and circadian information). To focus on this information flow, we removed all connections between neurons of equal or decreasing layer, thereby eliminating most feedback and some lateral connections. The resulting trimmed network retains ∼65% of the original connections (**Fig. S1h**) and is acyclic, without loops, representing the portion of the network that likely performs most of the direct, often rapid, feedforward visual processing.

Approximately 350 VPN types convey visual information to the central brain. Recent anatomical studies have revealed this full repertoire^11,16,52,53^, although the response properties and behavioral roles of some VPNs have been characterized in detail and intensively studied in recent years^45,52,54–56^, most remain functionally untested. Without principled guidance, discovering their function typically relies on bespoke stimulus protocols and idiosyncratic behavioral inspiration.

To complement this heuristic approach, we use computational tools to quantify the basic visual input features that reach each VPN through both direct and indirect connections. This analysis places strong constraints on the stimulus features to which each neuron can respond, providing testable hypotheses about their responsivity. We then group VPNs with similar predicted feature sensitivity, revealing distinct equivalence classes of visual pathways that we later extend into the central brain (**Fig. 4**).

To quantify how visual input propagates through the trimmed network, we computed the effective weight—the summed product of input-normalized synaptic connections along all paths from visual inputs to a downstream (output) neuron (**Fig. 2a**). The total effective weight from all visual inputs defines an output neuron’s visual input contribution (VIC), quantifying the fraction of its inputs attributable to the eight visual input channels. Relative effective weights were then obtained by dividing effective weights by the VIC, obtaining values that sum to 1. These values represent the relative activation strength, in a linear trimmed network, from each of the eight visual input channels.

How well does propagating visual inputs through the connectome capture known functional properties of optic lobe neurons? We addressed this question using a simple test case, the segregation of ON and OFF contrast polarity, where extensive physiological data are available. By comparing only the relative strengths of L1 (ON) and L2 (OFF) inputs, our approach correctly predicts the response polarity of most cell types with experimental measurements^52,55,57^ (**Fig. 2b)**. Only a few cell types (highlighted in pink) deviated from the expected polarity. While incorporating additional visual inputs and the synaptic signs could further improve these predictions, the success of this minimal linear model underscores that major visual functions are already specified by the propagation of signals from well-characterized visual inputs.

Visual input propagation recovers many other expected properties. For example, large l-LNvs, components of the flies’ circadian clock, receive dominant input from the HB eyelet^58^ and MeTu2a cells from dorsal rim R7d photoreceptors^59^, while also revealing previously unrecognized relationships, such as the distinct input mixtures arriving at three well-studied looming-sensitive neurons (**Fig. 2c,d**). LC4 inherits OFF sensitivity^45,55^ from L2; LPLC2 integrates L1-L3 inputs expected of motion-pathway VPNs downstream of both T4 and T5^44,60^; and LC6 receives substantial R7 and R8 signals, suggesting additional specialization for spectral contrast against the sky^52,61^. These distinctions clarify why multiple looming pathways coexist and demonstrate how this analysis can guide stimulus design and more systematic functional testing.

We performed hierarchical clustering of the relative effective weights across VPNs (**Fig. 2e**). Cutting the dendrogram at nine clusters (Optic lobe Classes, oc1-oc9, **Fig. S3a**) produced a consistent partitioning between the left and right homologous VPNs (**Fig. S3f, h-j**). VPNs make many connections to other VPNs^55^, and the connectivity between VPNs across clusters shows a prominent diagonal structure that persisted even after within-type connections were excluded (**Fig. S3c**). This preferential within-cluster connectivity suggests that the clusters capture VPNs with shared tuning properties, echoing observations from mammalian visual cortex demonstrating like-to-like connectivity among similarly tuned neurons^62–64^.

Each of the nine optic lobe pathway classes integrates distinct combinations of the eight visual input channels. By focusing on the strongest input, a range of functional hypotheses emerges. The pathways in cluster oc1 (dominated by HB eyelet) likely drive circadian circuits. The pathways in oc2 (R7d-R8d) encode polarization, oc3–oc5 (R7-R8) convey color, oc8-9 (L1-L2) process contrast, and oc6-7 (L3) luminance. But the pathway classes feature mixtures of input signals—oc4,6,7 could drive integrated color and contrast circuits, and oc3 color-integrating circadian clock circuits. The pathway classes dominated by HB eyelet, R7d-R8d, or R7-R8 inputs (oc1-3,5) have more cell types in earlier layers (**Fig. 2f**), bypassing the deeper processing of the pathways in oc4,6-9. As anticipated by their differing complements of inputs, the three looming-sensitive VPNs (LPLC2, LC4, LC6) fall into three different classes (**Fig. 2c-f**). Several of these pathway classes are readily interpretable. For example, many visual clock neurons^65^ are found in oc1, and nearly all of the large and small lobula plate motion-sensitive neurons, the LPTs^16^, LPCs, and LLPCs^66^ reside in oc9 (see also **Fig. 2g**), showing that the pathways dominated by the directional-selective neurons are functionally grouped.

The contributions of optic lobe neurons to the pathway classes vary widely. Among the inputs, the HB eyelet and R7d-R8d are the most specialized, mainly supplying oc1,2, whereas L3 is the least. To identify the VINs linking visual inputs to the VPN classes, we introduce the *participation fraction*—the mean effective weight from an optic lobe interneuron to VPNs in each cluster, output-normalized to one (**Fig. 2g**). VINs are markedly more specialized than predicted by uniform connectivity (**Fig. 2h**), each participating in only one to five of the nine VPN classes. Above-uniform participation is visualized in **Fig. 2i**, showing which visual inputs and VINs participate in each VPN cluster. Some VINs only participate strongly (p_i_>1/9) in a single cluster. For instance, the directional-motion-selective T4, T5 neurons only participate strongly in oc8-oc9 (also **Fig. 2g**, right). The major “contrast” classes (oc8-9) share many VINs, some of which also strongly participate in the “luminance” classes (oc6-7), but only a few also participate in the HB eyelet and R7d-R8d classes (oc1-3). These observations indicate that optic lobe pathways are partially segregated at the outputs but overlap within earlier processing stages, supporting both specialization and integration of visual features.

By explicitly revealing the visual pathways of the optic lobe, the network can now be summarized as a three-layer hierarchy with nine output pathway classes (**Fig. 2j**). VPNs are not simple copies of the input channels; rather, they process and transmit distinct mixtures of visual information. As the computations performed by VINs become better characterized, such as the ON/OFF directional selectivity implemented by T4 and T5, the specific visual features represented by each pathway could be predicted in greater detail by applying our attribution procedure to each VIN. Building on this functional mapping of visual pathways, based on the features of the different input channels, we next turn to spatial mapping, based on the columnar arrangement of visual inputs.

### Connectome-based predictions of anatomical receptive fields in the optic lobe

To determine where each neuron “looks” in the fly’s visual field, we estimated anatomical receptive fields (ARFs), the region of visual space from which it receives effective input. Visual inputs were assigned to retinotopic medulla columns within a hexagonal coordinate system, each typically containing five distinct inputs (L1-L3 and R7-R8, **Fig. 1b**). The specialized dorsal rim photoreceptors and the extra-retinal^67^ HB eyelet photoreceptors are excluded from the ARF predictions.

The visual-input-propagated ARF is the column-by-column, summed effective weight from the columnar visual inputs (shown for LC6, **Fig. 3a**). We characterized each ARF by fitting an anisotropic 2D Gaussian to obtain estimates of its amplitude, spatial center, size, and shape; fit quality was measured by the coefficient of determination (R²). Because these measurements were highly consistent across neurons of the same cell type (**Fig. 3b**), we emphasize the type-level medians.

A simpler version of an ARF derives from direct dendritic inputs rather than visual-input-propagated connections (**Fig. 3c**)^11,16,57,68,69^. When we fit Gaussians to these direct ARFs, the estimated centers were nearly identical, but sizes were smaller (LC6 population, **Fig. 3d**), reflecting that direct dendritic inputs typically convey signals integrated over multiple columns.

To test which measure better predicts functionally measured receptive fields, we compared direct and visual-input-propagated ARF sizes with calcium imaging data^53,55,70–72^ (**Fig. 3e, Table S2**). Propagated ARF predictions aligned closely with most measured receptive field sizes (Pearson correlation 0.97), whereas direct ARFs systematically underestimated them (Pearson correlation 0.88). This validation supports the visual-input-propagation framework, and we therefore focus on these estimates, hereafter referred to simply as ARFs.

Having established that ARFs provide realistic estimates of receptive-field size, we next analyzed how size varies with network properties of neuron types. ARF size did not depend on synaptic sign or directedness (fraction of feedforward connections; **Fig. 3f**), but varied systematically with optic lobe group, layer, cell-type abundance, VIC, and ARF amplitude (**Fig. 3f,g**). In early hierarchical layers, most ARFs typically spanned fewer than ten columns; in later layers, they expanded substantially. But among these network properties, ARF size correlated best, in absolute value, with number of neurons per cell types (Pearson correlation with layer: 0.30, number of neurons: -0.36, VIC: -0.26, ARF amp: -0.24). That cell types with many neurons had smaller, higher-resolution ARFs is consistent with their dense space-filling organization^11^. This is a hallmark of hierarchical visual processing, where later stages combine signals from earlier, finer units.

By analyzing the entire optic lobe, we also find prominent deviations away from this trend, including aMe12, a large, wide-field cell (layer 1, oc4) that receives direct photoreceptor input^73^, and projects to the central brain, highlighting a specialized early pathway for global visual cues. Some VPNs in layer 2, such as the three looming-selective cell types (LC4, LC6, LPLC2), span only about 10 columns—much fewer than MeVPMe12 (layer 2, oc7). The ARFs for all ∼350 VPN types are detailed in the supplementary gallery (**Document S2**).

A more direct indicator of hierarchical expansion comes from comparing ARF sizes across connections (**Fig. 3h**). Most connections from VINs to VPNs show at least twofold field expansion, sometimes exceeding 60-fold, whereas most connections among VINs preserve the ARF size. Most, but not all, VPN projections to central-brain targets show substantial expansion, with a modal value of 16-fold, while the majority of connections to other VPNs show minimal ARF size changes. The progressive broadening of receptive fields marks the expansion of local visual information towards more global visual processing. How much local vision remains in the central brain will be explored in **Fig. 5**.

Within each of the visual pathway classes, ARF sizes span more than an order of magnitude, revealing visual processing at multiple spatial resolutions (**Fig. 3i**). The circadian-associated pathway class (oc1) has the largest ARFs, consistent with the integration of wide-field visual inputs to reliably estimate broad features of the environment. Most of the color-associated pathways (oc3–oc5) show intermediate resolutions with few high-resolution VPNs, whereas the contrast- and luminance-dominant pathways (oc6– oc9) contain most of the smallest, highest-resolution VPNs.

In summary, the visual-input-propagated ARF sizes closely match experimentally measured receptive-field sizes and increase through the optic lobe hierarchy, from fine columnar sampling in earlier layers to progressively broader integration in VPNs with further expansion in the central brain. Having established that both functional and spatial predictions agree well with VPN experimental data, we next extend this analysis to the central brain, where experimental data are limited and connectome-based predictions will be critical for guiding future experiments.

### Mapping the flow of visual information into the central brain

Determining which central brain neurons are visually responsive has long been limited by the need for demanding physiological experiments, constraining our ability to map out the fly’s “visual system.” By propagating visual inputs through the connectome, we can now systematically predict these putative “visual” central brain neurons (VCBNs). We extended the feedforward propagation beyond the VPNs to all central brain cell types and quantified their visual input contribution (VIC) score, as a measure of visually driven signaling. We compared the propagated scores for bilateral homologous cell types (across the right and left sides) to determine a threshold above which VIC scores were more consistent (**Fig. S4a,b**). With this criterion, we identify approximately half of all central brain neurons (>5k cell types and nearly 11k individual neurons) as receiving substantial visual input contributions from the right optic lobe (**Fig. 4a**, **Table S3**). This threshold correctly includes all VPNs, as expected, and reveals a large visual contingent across the brain. Most of these VCBNs occupy layers 3-5 of the network hierarchy, several synaptic steps deeper than the VPNs themselves, confirming that visual processing extends broadly and deeply in the central brain.

To illustrate how these propagated visual inputs manifest in individual neurons, we examined representative examples of VCBNs (**Fig. 4b,c**). Among them, the sexually dimorphic P1 neurons provide a recently established example linking vision to behavior^56,74,75^. Our analysis identifies strong visual inputs to P1_2a neurons (layer 3), consistent with the recently noted link between these male courtship neurons and visual cues received via LC16 inputs^76^. In contrast, P1_5b neurons, located more than one hierarchical layer deeper, do not receive direct VPN inputs but indirectly connect to multiple VPN types. P1_2a and P1_5b neurons also receive notably different propagated contributions of each visual input (**Fig. 4c**). Of the 44 annotated P1 cell types, 31 exceed our VIC threshold, underscoring how pervasive visual inputs are to this social behavior circuit.

Using the propagation and clustering framework detailed in Fig. 2, we next examined how visual inputs are organized across all VCBNs (**Fig. 4d**). Clustering all neurons above the VIC threshold and cross-validating across hemispheres revealed nine distinct classes (Central brain Clusters, cc1-cc9) differing in size and composition (**Fig. S4c-i, Fig. S7c**). Each cluster comprises neurons with similar combinations of propagated visual inputs, capturing both the persistence and diversification of visual input features at higher processing levels (also reflected in exemplary neurons of each class, **Fig. 4c**). We find one class, cc6, is dominated by dorsal rim R7d, as a nearly pure polarized light pathway^39,59,73^, but it is striking to find other classes, cc2 and cc9, dominated by L3 (luminance) and L2 (OFF), respectively, showing that narrowly tuned visual pathways can propagate intact through multiple synapses. The other classes show integrated mixtures of visual inputs, consistent with progressive feature integration. The overall input composition of these classes mirrors the major visual pathway classes in the optic lobe (**Fig. 2e**, **Fig. S3i-j**): five (cc1-2, cc7–cc9) are dominated by luminance and contrast pathways, while four (cc3–cc6) are enriched for color-sensitive or circadian inputs from R7/R8 photoreceptors and the HB eyelet. Even six synaptic layers from the periphery, central-brain neurons remain partly sorted by their input channels, revealing how early visual feature streams persist and intermix into deeper central brain circuits.

Across the hierarchy, VCBNs are distributed primarily in layers 3–5, with TuBu neurons being exceptionally early (layer 2). While some classes (e.g., cc9) are restricted to the deeper layers 4-5, most span the full range (**Fig. 4e**). To trace how these signals enter the central brain, we next mapped the participation of individual visual projection neurons across the nine visual pathway classes in the central brain (**Fig. 4f**). As in the related analysis of VINs (**Fig. 2f-i**), each VPN contributes unevenly, most favoring 2-3 classes (**Fig. S4k**), while a few, such as LT66 participate in up to four classes. Together these patterns indicate that visual processing is organized into a combination of distinct and partially overlapping pathways that both preserve and recombine peripheral sensory features.

Our analysis identifies roughly half of all central brain cell types as being visually driven (**Fig. 4a**), but are they concentrated into visual brain areas? To address this, we next examined where the visual pathways project across the major anatomical regions^77^ of the central brain (regions of interest, or ROIs), and find an uneven mapping of the nine visual pathway classes: 76% of the synapses from these neurons target just 25 regions, 16 ipsilateral to the optic lobe source, 5 contralateral mirror-symmetric counterparts, and 4 midline neuropils (**Fig. 4g**). Several smaller classes narrowly target specific ROIs, recapitulating known neuroanatomy: the polarization class (cc6), mainly supplies the Ellipsoid Body (EB)^53,59,78^; the putative circadian class (cc3) primarily targets the Superior Medial Protocerebrum (SMP)^65,79^; and cc4 which combines color, polarization, and circadian inputs, divides its projections between these two regions. The L2-dominated pathways in class cc9 target only a few regions in the protocerebrum, whereas larger classes, including the “color-blind” contrast/luminance pathways (cc1-2, cc7-8) and the color-sensitive pathway cc5, supply many ROIs, establishing several regions as newly relevant centers of visual processing.

By visualizing the flow of vision across the hierarchical layers, we can see how visual the central brain is (**Fig. 4h**). The ∼4350 right-side VPNs with central brain connections already distribute visual signals widely, concentrated in regions where their axons terminate, mainly in the ventrolateral neuropils, such as the Anterior Optic Tubercle (AOTU), and the Ventrolateral Protocerebrum regions (AVLP, PVLP, PLP). Beyond the VPNs, visual pathways progressively expand, with deeper layers engaging increasingly larger sets of VCBNs. In early layers, visual signals are concentrated in the VPN-target regions with a noteworthy expansion into the Inferior Clamp (ICL). In deeper layers, the footprint expands to include superior neuropils (SLP, SMP), and in the deepest layers of our analysis, we encounter midline structures such as the EB, central to navigation^78^, and the Gnathal Ganglion (GNG), housing signals for descending control of behavior^80^. This visualization—quite literally—shows how visual information, originating in specialized sensory circuits, permeates much of the fly’s brain within five-six synaptic steps.

Propagation from the left optic lobe produced nearly mirror-symmetric patterns to those from the right, providing an independent validation for our propagation analysis and demonstrating that the visual organization of the central brain is bilaterally conserved (**Fig. S5a**). Yet, when signals are traced from individual visual inputs (from the right eye), distinct spatial patterns emerge across the brain (**Fig. S5b**), revealing that each channel engages a different subset of VCBNs and ROIs. These differences are further evident at the pathway level: each visual pathway class projects to a characteristic set of neuropils, forming distinct, partially overlapping visual maps across the central brain (**Fig. S6**).

Thus far we have examined the flow of visual information from one eye at a time, but are there specific central brain regions primarily engaged in *binocular* vision—defined here as the integration of inputs from both eyes without regard to binocular overlap (which is limited in flies^25^)? By quantifying each VCBN’s binocularity index (see Methods), we found that across layers, neuropils near the midline feature the highest prevalence of binocular neurons (**Fig. 4i**). The strongest binocularity occurs in the central complex and associated neuropils (EB, PB, LAL), whereas most lateral regions remain strongly monocular. These convergence zones mark the transition from local, eye-specific processing to the global representations required for spatial orientation, navigation, spatial memory, and multimodal integration. Vision in the central brain originates in peripheral regions near the eyes and spreads inward (**Fig. 4h**), while binocular vision is concentrated along the midline, farthest from both eyes (**Fig. 4i**). This organization reveals a striking architectural symmetry between the physical layout of visual processing in the fly brain and its functional visual hierarchy.

### Spatial vision in the central brain

This structural symmetry naturally raises a deeper question: how is visual space represented within the central brain architecture? To assess how spatial organization is preserved as visual signals pass from the retinotopic optic lobes into the central brain, we focused on two complementary aspects of spatial vision: (1) the distribution of high-resolution visual information and (2) regional differences in how visual space is represented.

To look for high-resolution vision in the central brain, we sought neurons that collect visual inputs from only a small region of visual space. We restricted candidates to neurons with an ARF that was well-captured by a 2D Gaussian (R² > 0.05) and mapped their synaptic distribution across hierarchical layers (**Fig. 5a**, top; **Fig. S7a**). Beginning with the VPNs, the highest-resolution projections are in the protocerebrum and AOTU, consistent with the known VPN target zones. As expected, ARF sizes generally increased through the deeper layers of the hierarchy (**Fig. S7b**). Using the stringent criteria for identifying high-resolution neurons, we excluded ∼90% of VCBNs, but hundreds of high-resolution neurons remain at all layers.

Notably, neurons within the EB show some of the smallest ARFs among deep-layer cells. Outside the EB, small, spatially restricted fields are found across many cell types and central brain regions (**Fig. 5a**, bottom). Because our framework is fully traceable, the visual pathways for high-resolution VCBNs can be studied, and these cells can be linked to the specific VPNs that supply them (**Fig. S7c,d**).

It is now well established that localized sampling of specific parts of the visual field occurs in the fly’s central brain, even in regions lacking obvious retinotopic organization^53,68,69^—but how widespread are these spatially biased samples? To ask which parts of the visual field are emphasized, we examined the spatial coverage of all propagated visual inputs, irrespective of resolution, across the most visual central brain ROIs (**Fig. 5b**). For this analysis, the visual field was divided into five sectors—dorsal, ventral, anterior, posterior, and central. In the medulla, coverage is nearly uniform (∼20% per sector), but almost every central ROI deviates significantly from uniform, revealing many spatial biases (**Fig. S7e**). The most consistent trend is an overrepresentation of the central eye sector, but other prominent regional biases are also evident. For example, the EB and PB show extreme dorsal-input dominance, the VES favors ventral inputs, and the SIP, LAL, and AOTU are anterior-biased. None of the “most visual” ROIs are posterior-dominant. These differences show that distinct brain regions differentially sample the visual field, emphasizing specific zones of the fly’s visual world—likely reflecting both sensory specialization (e.g., dorsal polarization vs. ventral ground motion) and the spatial organization of motor and behavioral control.

We next asked whether this organization extends to finer scales—whether central-brain neuron types preserve systematic relationships to visual space as exemplified by (but not limited to) the retinotopy in the optic lobes. To visualize this organization more directly, we mapped the receptive-field centers of VCBNs (with ARF fit R² > -0.5) to presynaptic sites across the central brain, linking eye position to brain locations (**Fig. 5c**). This view reveals an unexpectedly structured landscape of visual space representation. At the global level, different eye regions project to distinct brain territories. Within individual ROIs, the organization is even more precise. The AOTU, for example, contains two distinct zones, each with a continuous visual map, while midline structures such as the EB (dorsal) and NO (ventral) show strong, region-wide spatial biases. The fan-shaped body (FB) exhibits layered organization, with strata emphasizing anterior, dorsal, or ventral visual sectors—patterns likely arising from the spatial biases of a few cell types^78,81^. This data-driven analysis reveals multiple, map-like structures across the central brain, including unexpected spatial patterns in regions such as the ICL and the GOR, raising new questions about how these representations contribute to perception and behavior.

Together, these results show that the central brain remains deeply organized by the structure of the eye. Spatial information is represented in multiple complementary ways: some regions encode map-like retinotopy, others emphasize specific sectors of visual space, and many combine both. This hybrid architecture underscores how the fly brain transforms peripheral spatial sampling into distributed internal maps.

## Discussion

This study provides a connectome-derived account of how visual information is organized across the fly brain. By propagating defined visual inputs through the complete male CNS connectome, we can evaluate all four principles posed in the introduction:

### Feedforward versus recurrent connectivity

Our analysis revealed that despite dense recurrence, with 55% lateral or feedback connections, the optic lobe is organized into a shallow, three-layer hierarchy linking visual inputs to VPNs (**Fig. 1g,i**). Remarkably, the fraction of feedforward, lateral, and feedback connections closely matches recent estimates from the mouse visual cortex (∼0.43, 0.36, 0.21)^82^, indicating that modest feedforward bias can coexist with dense recurrence both within early sensory networks and deeper ones and across species. As in mammalian cortex, where feedforward and feedback streams are functionally segregated^83,84^, we found few neurons with both high feedforward and high feedback fractions (**Fig. 1k**). Trimming the connectome to remove same- or earlier-layer edges isolates the direct pathways likely underlying rapid visual responses while retaining the majority of connections. Within this structure, even deep central-brain regions such as the central complex can be reached within ∼6 synaptic steps from the visual periphery (**Fig. 4**).

### Receptive-field progression across hierarchical layers

To examine spatial integration across the network, we used propagated visual inputs to estimate anatomical receptive fields (ARFs), quantifying each neuron’s effective field of view.

Propagated ARFs showed a clear trend of increasing size with hierarchical depth and matched physiological receptive-field sizes better than estimates based solely on direct dendritic inputs (**Fig. 3e,g-h**). This mirrors classic increases in RF size seen in visual cortex^4,85^ and convolutional neural networks^86^, but the full fly connectome reveals additional structure. Each cell type’s count is the strongest predictor of ARF size, reflecting the tiling of higher-resolution cell types sampling visual space, while a small set of early wide-field neurons form large ARFs consistent with low-resolution or non-spatial roles. For the numerous cell types supporting the fly’s spatial vision, hierarchical progression emerges directly from connectivity, capturing receptive-field structure across neuron classes.

### Organization of parallel feature pathway

By quantifying the inherited visual-input composition of optic lobe and central brain neurons, we traced how distinct visual features propagate through the network. Propagation revealed classes of forward-directed optic-lobe pathways, with neurons in each class sharing characteristic mixtures of peripheral inputs—contrast, luminance, color, polarization, and circadian signals (**Fig. 2d-e**). These mixtures already predict aspects of functional tuning and help explain the functional diversity observed among VPNs (**Fig. 2b,c**). Extending this analysis into the central brain identified additional pathway classes for the nearly half of all neurons that receive substantial visual drive (**Fig. 4**). These classes show that visual information reaches deep brain regions and that certain channels remain largely unmixed, revealing limits to early convergence. Across both the optic lobe and central brain, some pathway classes are strongly biased towards one visual input source, but all classes show at least some cross-channel mixing rather than strict labeled-line isolation.

### Representation of visual space in higher brain areas

By propagating retinally anchored visual information, we found that not only does vision reach deep into the central brain, but that representations of visual space remain richly structured well beyond the optic lobes. Many visual central brain neurons retain small, well-fit ARFs several synapses downstream, including in navigation-related regions such as the ellipsoid body, showing that high-acuity vision persists deep in the hierarchy (**Fig. 5a**). Mapping receptive-field centers across the brain revealed region-specific organization, and in several areas, including the anterior optic tubercle, nearly complete map-like coverage of visual space (**Fig. 5b,c**). These patterns demonstrate that neurons in deeper visual areas preserve fine spatial detail and indicate that retinotopy is transformed rather than lost.

### Limitations of the study

Several methodological choices limit the present analysis. First, to obtain an acyclic graph suitable for propagation, we removed most lateral and feedback connections. This simplification isolates the dominant feedforward pathways but underestimates the contributions of recurrence, particularly inhibitory feedback. Second, we seeded our propagation exclusively with visual inputs. Although this is appropriate for the goal of mapping the visual system, many central-brain neurons integrate signals from other sensory modalities and internal state-dependent signals, which are ignored. Third, the effective-weight model does not incorporate synaptic sign or nonlinear integration. As a result, receptive-field sizes in strongly inhibitory pathways may be overestimated, and ON/OFF polarity (**Fig. 2b**) may be predicted more accurately by explicitly including known neurotransmitter identities. Fourth, our Gaussian ARF fits provide a compact summary of spatial sampling but work best for small, well-localized receptive fields and less well for large or complex ones. Finally, functional validation is strongest in the optic lobe, and most predictions in the central brain, including pathway identities, feature selectivity, and spatial organization, await targeted physiological and behavioral testing. Despite these limitations, the connectome-derived annotations presented here provide a foundation for understanding vision across the fly brain.

### Outlook

The fly is the first animal with spatial vision for which we have complete synapse-level maps of the entire brain, providing the first opportunity to trace all visual pathways from the periphery into the central brain. In doing so, we have mapped out the scope of the visual system, showing that nearly half of all central-brain cell types receive substantial visual drive. This already exceeds long-standing assumptions about the reach of the visual system in *Drosophila* and likely underestimates the full impact of visual processing. Because the propagation framework yields predicted receptive fields and likely visual feature sensitivities, experimental design can now be guided by explicit expectations about what a neuron “sees” and which basic visual features it is most likely to respond to. These predictions provide concrete starting points for identifying relevant neurons or designing targeted stimuli for exploring the functional roles of candidate cells and pathways. At a broader scale, the analysis revealed an architectural principle that becomes apparent only in the full connectome: binocular integration emerges in deep midline neuropils (**Fig. 4i**), reflecting the brain’s physical layout and establishing a spatial correspondence with the structure of its visual hierarchy.

Beyond the specific findings we report, our analysis illustrates how a complete connectome can be used to move from anatomy toward function without explicit simulation of biophysics or nonlinear circuit dynamics. This approach complements recent dynamical simulations of connectome-constrained circuits^19,20^. Simulations are indispensable for capturing timing, nonlinear integration, and adaptation in neural activity, but these features are largely missing from connectome datasets, and models typically require assumptions that are underconstrained even in dense wiring diagrams. In contrast, the propagation method is deliberately transparent, asking what can be directly inferred from connectivity using a small set of interpretable assumptions that can be validated against physiological measurements. This makes it a tractable first step for converting anatomy into functional predictions in large, recurrent networks. The approach is broadly applicable as new connectomes anchored by well-defined sensory inputs—whether in the fly, zebrafish^87,88^, or mammalian retina^89^—become available. While comprehensive cell typing in flies greatly aids interpretability, it is not required. The same propagation framework outlined here can be applied to reveal functional hierarchies, pathway organization, and functional predictions across the nervous system, establishing a foundation for deep connectome-informed comparisons of brain architectures^90^.

## Supporting information

Doc S1

Doc S2

## Resource availability

All analyses are based on the publicly available Male CNS connectome (v0.9), hosted at https://neuprint.janelia.org/.

## Acknowledgements

We thank HHMI and the Janelia Research Campus for supporting our work and that of the FlyEM team, whose production of the underlying connectome data, and early access to it, made this study possible. We thank Marisa Dreher for contributions to graphics design and figure production, Yijie Yin for collaboration on the Connectome Interpreter Toolbox, Stuart Berg for database and software support, and Gerry Rubin for supporting A.N. We also thank Kit Longden, Laura Burnett, and all members of the Reiser and Otopalik laboratories for critical feedback throughout the project. We are grateful to Eyal Gruntman, Pavithraa Seenivasan, Aneesh P B, and Saket Navlakha for early and regular discussions, and to Frank Loesche for assistance with neuron rendering pipeline. This article is subject to HHMI’s Open Access to Publications policy. HHMI lab heads have previously granted a nonexclusive CC BY 4.0 license to the public and a sublicensable license to HHMI in their research articles. Pursuant to those licenses, the author-accepted manuscript of this article can be made freely available under a CC BY 4.0 license immediately upon publication.

## Author contributions

J.H. and A.Z. designed and implemented the connectome analysis, wrote all code, and produced all analysis figures. A.N. provided neurobiological expertise and guidance. E.M.R. contributed to neuron figure production. S.R. and M.B.R. guided the analysis from inception and supervised the research. M.B.R. and J.H. drafted the manuscript with input from all authors. All authors contributed to manuscript revisions.

## STAR Methods

### METHODS DETAILS

#### Analysis software

All analyses were implemented using custom software written in Python (v3.11.8). To access the the data, we used Neuprint (neuprint-cns.janelia.org) for the male-cns:v0.9 dataset^23,91^. To standardize much of our analysis, one of the authors (J. Hoeller) contributed to the development of the Connectome Interpreter Toolkit (github.com/YijieYin/connectome_interpreter)^92^—a python library for efficient computation of indirect paths and effective weights between neurons—which we used throughout this study. For data plotting, we used plotly^93^ and Matplotlib^94^, and for rendering neurons we used the Blender-based visualization pipeline developed for the optic lobe study^11^.

#### Connectomic data selection

Our analysis only considered cell-typed neurons (using “type=NotNull” in neuPrint queries), and the connectivity between them, in terms of the number of synapses—excluding synapses in the partially reconstructed laminae. We did not use a threshold on the number of synapses. Neurons were assigned a neurotransmitter (categories: acetylcholine, glutamate, GABA, dopamine, serotonin, octopamine, histamine, unclear) based on consensus neurotransmitter predictions in Neuprint.

Cell types were divided into regions based on superclass information^23^. Superclasses starting with “cb” as well as “sensory_descending” and “descending_neuron” were classified as “CB”, whereas if they started with “vnc” or were “sensory_ascending”, “efferent_ascending”, “ascending_neuron” we classified them as “VNC”. Right optic lobe cell types were based on the list of instances from the Optic Lobe connectome study^11^, and left optic lobe as the symmetric instances of those same cell types (opposite soma position). For cell types in the right optic lobe list that have cell bodies on both sides, we only classified the cell type with the cell body on the right as right optic lobe and the cell type with the cell body on the left as left optic lobe. optic lobe cell types were further subdivided into four optic lobe groups: visual inputs (L1-L3, R7-R8, R7d-R8d, HB eyelet), VPNs (visual projection neurons), VCNs (visual centrifugal neurons), and the remainder as VINs (visual interneurons). The Optic Lobe connectome study^11^ used 5 groups, including VPNs and VCNs, but the OLINs (optic lobe intrinsic neurons) and the OLCNs (optic lobe connecting neurons) are now split into visual input and VINs, whereas the “other” group was now classified as “CB”.

In all analyses other for **Fig. 1d**, we distinguished cell types on the right and left hemispheres, using the *instance* information from neuprint. To facilitate left-right comparisons, we removed terms in parentheses and the string “_indeterminate” from instance names. For **Fig. 1d**, cell types were defined using the neuPrint “type” excluding neurons whose type names contained “_unclear” and “R1-R6”. Pale and yellow photoreceptors were not counted as separate types, and for the neuron count we used the filled-in R7/R8 dataset (see section **Filling in R7, R8 cells and their connectivity**).

### Columns and hexagonal coordinates

In the optic lobes, the retinal visual inputs can be placed on a hexagonal coordinate system, because the facets on the compound eye form an approximate hexagonal grid, which has been mapped with near one-to-one precision to columnar ROIs in the medulla, the largest neuropil in the optic lobe, and the entry point to the visual system after the lamina^11,25^. This mapping has been extended to the so-called columnar neurons, cell types that have approximately as many neurons as there are lenses. In addition to previously assigned columnar cell types (L1-L2, L5, Mi1, Mi4, Mi9, Tm1-Tm2, Tm9, Tm20, C3, T1, T4)^11,23^, we assigned L3, R7 and R8 neurons to hexagonal coordinates *(hex1_id, hex2_id)* based on the medulla column ROI containing the largest number of synapses.

For the propagation analysis in our study, we focus on eight visual input cell types. The first seven of these provide columnar visual input to the medulla. We use the lamina monopolar cells, L1, L2, and L3 that are direct targets of the R1-6, conveying their signals to the rest of the brain (including the other two LMC types L4, L5)^50,96,97^. We do not use R1-6 photoreceptors as visual inputs in our analysis for several reasons: i) due to neural superposition in the dipteran eye, R1-6 from different ommatidia converge onto the same lamina cartridge, meaning that the columnar sampling of visual space is assembled at the level of the lamina monopolar cells, not the photoreceptors^13,26,27^, ii) by starting with L1-L3 we make use of their known properties, which would not be possible if starting with R1-6, and iii) large parts of both laminae, and as a result many R1-6 photoreceptors are missing in the EM dataset^11^ (we exclude all lamina connectivity data from our analysis as it is less reliable). We include the inner photoreceptors R7 and R8, which project from the retina directly to the medulla and establish columnar, spectrally shifted measurements at each location on the compound eye^73^, but do not distinguish R7 and R8 subtypes (R7p and R7y, R8p and R8y), which have additional, finer differences in their spectral responses. We do include the specialized R7d and R8d from the dorsal rim that provide polarization information^38,39^. Finally, we include HB eyelet inputs, a cluster of specialized extra-retinal photoreceptors that provide a small but important visual contribution to the circadian clock^40^.

As *(hex1_id, hex2_id)* correspond to a non-orthogonal basis, it is convenient to also introduce an orthogonal basic with *(x,y)* coordinates via *x* = *hex*1_id - *hex*2_id and *hex*1_id - *hex*2_id - 22. The coordinates *(x,y)* are integer-valued, but the corresponding orthogonal basis is not normalized. For an orthogonal, normalized basis, which will be important for fitting 2D Gaussians, we used the *(h,v)* coordinates defined by *h* = *x* and 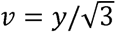.

The hexagonal column maps (**Fig. 1b**, **Fig. 3a,c**, **Fig. 5a,b**) are based on methods previously developed^11,25^. We used the Connectome Interpreter Toolkit^92^ to calculate the effective contributions, by column, to the neurons (or set of neurons) of interest.

### Filling in R7, R8 cells and their connectivity

Some R7 and R8 photoreceptors are missing or are incompletely reconstructed due to sample preparation^11,23^. To enable the propagation of these visual inputs, we generated substitute (“fill-in”) photoreceptors together with their major connectivity. We also introduced substitute weights for selected connections of partially reconstructed cells. The major connections we aimed to fill-in were from R7 to R8, Dm8a, Dm8b, Dm9, Tm5a, Tm5b, MeTu3b, MeTu3c, L3, Dm11 and Dm2, from R8 to R7, Tm20, Tm5c, Dm9, Mi15, Dm2, Mi1, Mi4 and Mi9, and from Dm9 to R7 and R8.

We first identified medulla columns missing an R7 cell (annotated as R7y, R7p, R7d and R7_unclear) or an R8 cell (similarly annotated), excluding columns containing R7d or R8d and those at the medulla edge (less than six neighbors in the hexagonal coordinate system). For each missing R7 and R8 cell, we added a new cell (bodyId) with synthetic connectivity. The bodyIds of these cells have a standardized format, starting with 177777 for a right optic lobe R7, 188888 for a right optic lobe R8, 155555 for a left optic lobe R7 and 166666 for a left optic lobe R8, and ending with the four-digit integer *100 × hex1_id + hex2_id* where *(hex1_id, hex2_id)* is the hexagonal coordinate of the corresponding column.

To identify incompletely reconstructed photoreceptors Rx for *x=7,8*, we split medulla columns (excluding dorsal and edge columns) into two groups: “reference” columns with at least output connections, and “fill-in” columns with less than output connections. We chose *n*_7_ = 75 and *n*_8 =_ 100.

To fill in the connections between photoreceptors, we used fixed values: 12 for R7->R8 and 32 for R8->R7, based on the distribution of the connections between R7 and R8 cells in the reference columns (rounded, median values from the right optic lobe). For all other connections between a photoreceptor and a target or input cell type, we trained linear regression models on ∼ 500 reference columns (selected separately for R7 and R8 per optic lobe). Features included (i) the total input and output connections of the target/input cell in medulla layers M1 to M6 for R7 and M1 to M4 for R8, and (ii) all connections within the fill-in column for each ordered pair among Tm5a, Tm5b, Dm8a and Dm8b (e.g. Dm8a-> Tm5a), which served as a proxy for pale/yellow-related differences between medulla columns. This gave 24 features for R7 and 20 for R8. We trained sklearn.linear_model.Lasso with alpha=0.1 on 80% of reference columns and validated them on the remaining 20% (R^2^ values between 0.5 and 0.9, depending on target/input cell type). The fitted linear models were then applied to fill-in columns to generate synthetic connections.

For the R7 and R8 connectivity data used in our analysis (to be provided in the companion code repository), we renamed Rxp, Rxy, and Rx_unclear as Rx for *x=7,8*, and replaced existing connectivity for Rx in filled-in columns with synthetic values. Note that other existing connections of these Rx cells that were not replaced, i.e., connections to minor synaptic partners, and the connections of photoreceptors in the reference columns were not changed. All connections of cells with the cell type name “R7R8_unclear” were discarded.

### Converting the fly connectome into a network

To reveal global structure in the fly connectome, such as hierarchical and parallel processing, we view neurons as the nodes of a network that are connected by weighted and directed edges for each pair of pre- and post-synaptic neurons. For the weight, we use input-normalized number of synapses, which is the number of synapses (for each pair of connected neurons) divided by the total number of post-synapses a post-synaptic neuron has. One reason to normalize is that columnar neurons are expected to perform the same computation in every column, yet their synapse distributions are not identical across columns^11^. In particular, more synapses are formed near the equator. However, the input-normalized synapse counts are more consistent across columns.

### Information flow algorithm

Numerous algorithms have been proposed to extract hierarchical structure from directed, weighted networks, broadly falling into flow-, spectral-, or community-based (not mutually exclusive). Flow-based methods infer hierarchy from the directional bias of information flow through the network, spectral methods exploit the algebraic properties of network matrices to derive hierarchical coordinates, and community-based methods identify nested groups of nodes with similar connectivity profiles. We employed the information flow algorithm from prior work^18^, a flow-based approach, because it allowed us to initialize the flow from a biologically meaningful set of inputs, the visual inputs. This algorithm is similar to random-walk and diffusion processes on networks^98^, but is input-normalized instead of output-normalized and incorporates nonlinear integration of incoming signals, providing a more biologically plausible model than the linear assumptions underlying spectral methods.

We slightly modified the information flow algorithm, defining a hierarchical “layer” for each neuron based on its activation step under the information-flow dynamics, and using a deterministic, rather than probabilistic, updating of neuron activity. In this model, every neuron has a cumulative activation probability (a bounded activation variable), which is updated (remaining constant or increasing) deterministically at each activation step. Input neurons, by definition, start with an activation probability of 1. Every connection has a traversal probability, which is computed from its input-normalized connection strength via a threshold-linear function. The only parameter in this model is the threshold in this function. Its value is chosen (= 0.1) such that the model assigns T4 neurons, known to be two synapses downstream of L1 and L3 neurons, to be in layer 2. This threshold-linear function is such that (**Fig.1f**, inset) the traversal probability increases linearly from 0 to 1 as the input-normalized connection strength increases from 0 to 0.1, and remains at 1 for connection strengths > 0.1. Note that each connection’s traversal probability is constant throughout all activation steps. At each step, we first compute, for each connection, the product of its traversal probability and its upstream neuron’s cumulative activation probability. This is the connection’s activation probability. Then we computed, for each neuron, the probability that at least one of its input connections is active. This is the update to this neuron’s cumulative activation probability at this step. Despite the use of probabilistic terminology, all updates in this procedure are deterministic, and the algorithm produces the same result on every run.

The layer of a neuron is then given by the average step of activation, representing the average number of synaptic steps required for activation through the connectome under this nonlinear flow model. These layer assignments therefore provide a graph-theoretic estimate of each neuron’s position within the visual hierarchy.We used the function ‘compute_flow_hitting_tim’ from the Connectome Interpreter Toolkit with the parameters *flow_seed_idx* selecting the visual inputs, *flow_steps* equal to *20,* which is large enough to ensure all neurons that can be reached are reached, and *flow_thre* equal to *0.1*. The value of *0.1* corresponds to the saturation of a simple threshold-linear function at 10% active total weighted input, which we chose by testing the algorithm on T4 neurons. For T4 neurons, we expect activation at the second step (considering the most relevant inputs, L1 and L3 are at step 0). If saturation was chosen at 20% or larger, the probability of activating T4 at the second step was small.

### Effective weight (propagation of weights along paths)

To characterize how visual signals propagate from the visual inputs downstream, we focused on the subset of connections that most directly relay information from earlier to later layers. After assigning each neuron to a hierarchical layer, we removed edges that projected to neurons in the same or earlier layer, thereby discarding most feedback and lateral connections. The resulting acyclic, forward-directed graph provides a simplified representation of the likely most relevant routes through which visual information is transmitted to downstream targets.

For an “input” neuron that connects to an “output” neuron via one or more paths, we define the effective weight as a measure of the strength of this indirect influence. To be clear, a *path* is a sequence of nodes in which each consecutive pair is connected by an edge. The effective weight along a single path is computed as the product of the weights along its edges in the trimmed network, and the total effective weight is given by the sum over all paths connecting the two neurons.

This computation is formally equivalent to a linear model, but because we apply it to a trimmed network that retains only edges from neurons in earlier layers to neurons in later layers, it does not correspond to a linear model of the full network. One interpretation of this approximation is that it emphasizes the direct (possibly fast), feedforward components of visual processing. In practice, this approach generates a transparent set of interpretable predictions with explicit assumptions that compare favorably with experimental data (**Fig. 2b** and **Fig.3e**) for optic lobe neurons.

For our implementation, we used the function ‘compress_paths’ from the Connectome Interpreter Toolkit. This function efficiently computes successive powers of the trimmed, input-normalized connectivity matrix. The highest power is set by the parameter *step_number*, which is chosen as *10*, allowing us to capture paths of length up to *10*. This was sufficient, as the latest

CB neurons were in layer 6 (**Fig. 4b**). We further used the parameter output_threshold = 10^-6^, which set all matrix elements below this value to *0* (for each power). This threshold sparsened our matrices and was introduced primarily to reduce memory requirements, but it also limited our ability to make precise predictions for weakly visual neurons in deeper layers (see **Thresholds for central brain analyses**). This is because successive multiplications of weights < 1 gives exponentially small values and the number of forward-directed paths is limited.

### Visual input contribution and relative effective weight

We primarily analyze paths that begin with visual inputs and end at either VPNs or CBNs. The sum of effective weights from all visual inputs defines the visual input contribution (VIC). Since VIC decreases with layers (**Fig. 4a**), even purely visual neurons such as Tm5a have a VIC smaller than *1* (in this case, *0.161*).

A major reason for this decrease is that we did not renormalize the trimmed input-normalized connectivity matrix—thus, the sum over trimmed input weights is <*1*, reflecting the removal of connections. This is particularly relevant in the CB, where renormalizing the trimmed network would artificially inflate VIC values, since trimming might remove non-visual sensory inputs. Hence, the VIC should be viewed as a relative measure rather than an absolute one.

To remove this layer dependance in our pathway analysis and visual response predictions, we normalized effective weights to *1*. For effective weights given at the cell type level, we summed effective weights across neurons of the same input cell type, then took the median across neurons of the same output cell type, and finally normalized these values to *1*, yielding the relative effective weight (**Fig. 2b,d,e** and **Fig. 4c,d**).

### Experimental data for ON/OFF comparison

To validate how well the propagation of visual inputs through the connectome captures known functional properties of optic lobe neurons, we compared the predicted contrast polarity (ON contrast vs. OFF contrast) to existing experimental data. We drew on three calcium imaging studies examining a range of optic lobe cell types (**Fig. 2b**)—one focusing on VINs^57^ and the other on VPNs^55,68^. The data are summarized in **Table S2** and explained below.

Based on the linear spatiotemporal receptive fields (STRF) fit to identified neurons’ responses to ternary (black-gray-white) noise stimuli^57^, neurons were classified as ON if their STRF showed calcium increases to luminance increments and OFF if they showed calcium level reductions to luminance decrements. VPN responses are not well captured by STRFs to noise stimuli; instead, we used the relative responses of dark on bright or bright on dark looming stimul^55,68^. For all reported VPN cell types included in **Fig. 2b**, responses to the black looming stimuli led to stronger calcium responses, consistent with OFF responses. The exceptions are LC15 and LC18, which showed minimal, non-significantly different responses to ON and OFF looming stimuli^55^ and are therefore excluded from our comparison. Our comparative model is the simplest one possible—we predict each cell type as ON if the relative contribution of L1 > L2 and OFF if the contribution of L2 > L1.

### Pathways and pathway diagrams

A pathway is a collection of the strongest paths from a set of input neurons to a set of output neurons, and pathway diagrams are visualizations of pathways. In all pathway diagrams, we used trimmed, input-normalized number of synapses as weights (**Fig. 1g**, **Fig. 2c,g** and **Fig. 4b**).

To identify all neurons connecting a given set of initial neurons to a set of final neurons through a fixed number of intermediate neurons, we used the Connectome Interpreter Toolkit function ‘find_paths_of_length’. The number of intermediate neurons was varied from *0* to *n_diff*, where *n_diff* is the difference between the mean layer of the final and initial neurons, rounded up to the next integer.

To select the strongest paths, we used two parameters, *thre_cumsum* and *thre_step_min* (both numbers between *0* and *1*). For each fixed number of intermediate neurons, we ranked paths by effective weight (rank *1* is the strongest path, i.e., the path with the largest effective weight). We then identified the smallest rank *R* such that effective weights of the paths with ranks *1-R* accounted for at least a fraction *thre_cumsum* of the total effective weight. Among these top-ranked paths, we determined the minimum direct weight *step_min*. If *step_min* was smaller than *thre_step_min*, we set *step_min = thre_step_min*. Finally, we retained only paths for which all direct weights exceeded *step_min*.

For visualization in this manuscript, we manually chose *thre_cumsum* and *thre_step_min* for each panel, such that the number of paths was neither too large nor too small. We used *thre_cumsum=0.8* and *thre_step_min=0.01* for **Fig. 1g**, *thre_cumsum=0.4* and *thre_step_min=0.01* for **Fig. 2c** and **Fig. 4b**, *thre_cumsum=0.3* and *thre_step_min=0.05* for **Fig. 2g**.

The fraction of displayed paths (indicated as *f*) in each pathway diagram was computed as the ratio between the total effective weight along the selected paths and the total effective weight across all paths.

### Hierarchical clustering

We grouped pathways into equivalence classes based on clusters of output cell types—specifically, VPNs in **Fig. 2** and visual CBNs in **Fig. 4**. Output cell types were clustered using the relative effective weight starting from visual inputs (see **Visual input contribution and relative effective weight**). Each output cell type was thus represented by an eight-dimensional feature vector.

Hierarchical clustering was performed on these feature vectors using the scipy.hierarchy^99^ and fastcluster^100^ libraries, with Ward’s linkage method and Euclidean distance as the metric. The number of clusters was chosen manually, guided by the following criteria:

1. Stability: the clustering should remain consistent across a range of dendrogram cut thresholds (*thre_link*, normalized by maximum linkage), such that the number of clusters changes slowly with *thre_link* (**Fig. S3d** and **Fig. S4e**).
2. Consistency: clusters derived from right and left-side output cell types should be relatively consistent at corresponding *thre_link* values (**Fig. S3f,j** and **Fig. S4f,i**).
3. Interpretability: each cluster should contain more than one cell type, and a higher number of clusters is generally preferred for expressiveness.

For criterion 2, relative effective weights of left-side output cell types were computed starting from the left-side visual inputs (**Fig. S3h-i** and **Fig. S4g-h**). Each of the 347 right VPNs has a corresponding left-side type, except for LoVP31, MeVP57, and MeVP64, which exist only on the right. In the CB, 98 cell types have their soma on the right but not the left, and 83 exist only on the left. To compare left and right-side clusters, we only considered those cell types with mirrored left–right homologs. Cluster similarity was quantified using the adjusted Rand index (ARI, **Fig. S3f** and **Fig. S4f**), implemented with the sklearn.metrics^101^ library. The RI measures clustering similarity, and the ARI corrects for chance agreement. An ARI of 1 indicates identical clustering, 0 indicates random similarity, and −0.5 corresponds to clustering more dissimilar than random. We further found an optimal assignment between left and right-side clusters using the Hungarian algorithm on the confusion matrix (**Fig. S3j** and **Fig. S4i**). The (i,j) entry of the confusion matrix equals the number of cell types that are in left-side cluster i and the right-side cluster j. For the implementation, we used the ‘linear_sum_assignment’ function in scipy.optimize^99^.

For criterion 3, we required that each cluster contain at least two output cell types (unlike **Fig. S3e**). Too few clusters would obscure distinctions between groups—for example, cutting the VPN dendrogram at two clusters (**Fig. 2e**) would yield one cluster dominated by cell types responding primarily to HB eyelet and dorsal photoreceptors, and another cluster responding to other visual inputs, which would not be sufficiently expressive.

### Participation computation and participation graph

The clusters of output cell types split pathways into different equivalence classes; in particular, each output cell type only belongs to a unique pathway. On the other hand, earlier cell types (visual inputs and VINs in **Fig. 2** and VPNs in **Fig. 4**) generically participate in a few but not all pathways. This is quantified by the participation fraction *pi* (**Fig. 2g**). For every earlier neuron, we find the effective weight from that neuron to all output neurons. Then we take the mean of this effective weight per output cluster, resulting in one non-negative number per cluster, and finally we normalize these numbers so they sum to 1. If *pi=0* then that earlier neuron has no connection to any neuron in output cluster *i*, while *pi=1* means that this neuron only connects to neurons in output cluster *i*.

The participation graphs in **Fig. 2i** and **Fig. 4f** visualize the major earlier cell types that participate in each output cluster. Edges are given by *pi* and are only displayed if *pi* is larger than what would be expected from uniform participation (the reciprocal of the number of clusters).

We used a force-directed drawing algorithm (Kamada-Kawai^102^) and the igraph^103^ library for plotting.

### Flow diagram

The flow (Sankey) diagram in **Fig. 2j** visualizes the hierarchical and parallel structure of the right optic lobe. We excluded R7d, R8d, and HB eyelet because they form substantially fewer synapses than the other visual inputs, and we excluded VCNs because they primarily provide feedback.

Neuron layers in the right optic lobe were binned to integer values with a maximum layer of 3. For visualization, all CB and left optic lobe neurons were assigned to a nominal “layer” 4. In layer 0, visual inputs were grouped by cell type; in layers 1–3, neurons were grouped either into VINs or the nine VPN clusters; and in “layer” 4, neurons were grouped by brain region (CB or left optic lobe).

For each neuron, we computed the output-normalized number of synapses (the sum of outgoing weights summed to 1). For each layer *k* and each group, we then summed the output-normalized number of synapses over all post-synaptic neurons in layers > *k* and within each group. These values, which scale with the number of pre-synaptic neurons, were chosen as the edge weights for the Sankey diagram. Only edges with weight of *5* or above were displayed, unless this would remove all incoming or outgoing edges for a given node, in which case we retained the edge with the largest weight.

The flow diagram was implemented using the function “go.Sankey” from the plotly library. Node positions were manually arranged: the x-coordinate was determined by layer assignment, and the y-coordinate was adjusted to visually order and space the nodes. The height of a node is determined by the maximum of the incoming and outgoing weights, which for layers 0-3 approximately equals the number of neurons in that node. For CB and the left optic lobe nodes, height reflects the number of neurons projecting to these regions.

### Anatomical receptive fields and 2D Gaussian fits

To compute visual input propagated anatomical receptive field (ARFs), we calculated the effective weight from selected visual inputs (L1-L3, R7-R8) to each downstream neuron in the trimmed network and summed these weights over visual inputs in the same medulla column (example in **Fig. 3a**). HB eyelet and R7d-R8d were excluded because they do not cover 2D portions of medulla columns (**Fig. 1b**).

For direct ARFs, we used the synaptic inputs in the local columnar organization of the medulla, lobula, and lobula plate, depending on which neuropil contained the majority of a neuron’s post-synapses. Each post-synapse was assigned to a local column, also referenced in hexagonal coordinates^11^. The number of post-synapses in each column was counted and normalized to *1*, yielding the direct weight per column (example in **Fig. 3c**).

The ARF amplitude was defined as the maximum value of the ARF distribution across columns. To quantify 2D spatial properties of the ARF, such as center, size, and orientation, we fit it to a 2D Gaussian.

Gaussian fits were performed on the ARF values defined over the hexagonal coordinates (*x,y*), using five parameters: center (*x*_0_*,y*_0_), standard deviations (*a,b*), and orientation *φ*. The fitting procedure consisted of three steps:

1. Upsampling and smoothing: The hexagonal grid was upsampled by a factor of 5, and the distribution was smoothed with a Gaussian filter (σ = 1 in original units).
2. Masking: On the smoothed distribution, values were ranked from highest to lowest, and the threshold value *thre_val* was defined as the point where the cumulative sum reached *0.7*. All values below this threshold were set to *0*, resulting in a masked and smoothed distribution.
3. Parameter estimation: Gaussian parameters were derived using a covariance-based method. The ARF center (*x*_0_,*y*_0_) was computed as the value-weighted mean over the upsampled grid. The weighted covariance matrix of deviations from *x* - *x*_0_ and *y* - *y*_0_ yielded eigenvalues corresponding to the variances *a* and *b*; the larger eigenvalue defined *a*, the smaller *b*. The orientation *φ* was obtained as the phase of the eigenvector associated with *a*.

The ARF size was computed as 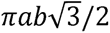, expressed in units of the hexagonal column area, referred to in the texts as “columns” or “col” when abbreviated.

To assess the goodness of fit, we computed the coefficient of determination *R*^2^ between the original (masked) ARF and its Gaussian fit. Both distributions were normalized to sum to *1*, and we calculated the residual sum of squares (*SS_res_*) and the total sum of squares relative to a uniform distribution (*SS_tot_*). The goodness of fit was then *R*^2^ = 1 - *SS_res_*/*SS_tot_*. Large ARFs typically do not have exponentially decaying flanks, resulting in a negative *R*^2^, which means that uniform predictions would perform better as an ARF size estimate than a 2D Gaussian fit. Since *R*^2^ is unbounded from below, bad fits can result in arbitrarily large negative *R*^2^ values. As these large negative *R*^2^ values are hard to interpret, we clip negative values at -1.

### Experimental data for RF size comparison

To evaluate whether our estimated anatomical receptive fields (ARFs) correspond to functionally measured receptive fields (RFs), we compared their sizes for both indirect and direct ARFs.

Functional RF sizes were obtained from published calcium imaging datasets.

From each study, we extracted estimates of the full width at half maximum (FWHM) of neuronal response profiles (in units of degrees squared). Assuming a Gaussian response shape, we converted the FWHM along each axis (*i* = 1, 2) to standard deviations as *σ_i_* = *FWHM_i_*/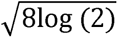, and computed the RF size as *πσ*_1_*σ*_2_.

To compare these functionally measured RF sizes with our predicted ARF sizes, we expressed one column unit as 25 deg^2^, based on a 5° inter-ommatidial angle, a reasonable approximation to measurements of the fly eye^25,104^.

Functional measurements are summarized in **Table S2** based on the following studies:

- Haag, et al.^70^, Fig 2: single ommatidia stimulation, measuring L2 and T4a responses.
- Arenz, et al.^71^, Fig 2: 1D white-noise stimuli with 2.8° resolution used to measure eight medulla columnar cell types (Mi1, Mi4, Mi9, Tm1-Tm3, Tm4, Tm9).
- Drews, et al.^72^, Fig S3: 2D white-noise stimuli with 2.8° resolution used to measure L1-L5 responses.
- Klapoetke, et al.^55^, Table S4: localized drifting gratings used to estimate responses of ten VPNs (LC4, LC11-LC12, LC15, LC17-LC18, LC21, LC25, LPLC1-LPLC2).
- Morimoto, et al.^68^, Fig. 3S3: putative LC6 bouton responses to 9° flashing spots.

### Thresholds for central brain analyses

Identifying visual neurons in the central brain (CB) is challenging because the VIC decreases across layers (see **Visual input contribution and relative effective weight**), making the use of an absolute threshold potentially misleading.

For technical reasons, we nevertheless chose a lower VIC threshold of 5 × 10^-4^, motivated by the fact that the trimmed, input-normalized connectivity matrix was thresholded at 10^-6^). In a worst-case scenario, effective weights from all ∼*4000* visual inputs to a given output neuron each fall just below 10^-6^, yielding a VIC of 0 instead of 4 × 10^-4^. In a more typical scenario, a visual input cell type covering about half the eye (ca. *500* columns) with effective weights just below 10^-6^ would give a VIC of 0 instead of 5 × 10^-4^.

Consequently, we do not interpret analyses for neurons with VIC values at or below 5 × 10^-4^, as these values approach the numerical resolution limit of our computation. We also note that left vs. right-side propagation on the symmetric neurons gave more consistent VICs for VIC > 5 × 10^-4^ (**Fig. S4b**).

To study the high-resolution pathways in the CB, we focused our analysis on CB cell types that additionally had a reasonable 2D Gaussian fit (*R*^2^ > 0.05). We note that the ARF size continued to show an increasing trend across layers (**Fig. S7b**, similar to **Fig. 3g**), a trend not observed without thresholds (the ARF size decreases in deeper layers for VIC<5 × 10^-4^ as the thresholded effective weights would eliminate many of the columns).

### Spatial distributions of quantified metrics across central brain

We use heat-mapped projections to illustrate the spatial distribution of three metrics in the central brain: Visual Input Contribution (VIC, **Figs. 4h, S5, S6**), binocularity (**Fig. 4i**), and ARF size (**Fig. 5a**). The metrics are calculated for each Visual Central Brain Neuron, but since many neurons have large, complex shapes, we instead map the distributions of pre-synapses (T-bars), to capture the locations in the brain where this information is available. To visualize the 3D distribution of these synapses, we performed an *Extreme Value Projection* where we computed the mean of either the 10% highest values (in the case of VIC and binocularity) or 10% lowest values (in the case of ARF size, where smaller values represent higher visual resolution) that are contained in all voxels along a projection ray (along the “z” dimension in the typical orientation at which the brain is viewed). We used an array of 240×200 projection rays to cover the entire central brain region, resulting in a pixel size of 2×2 µm. To enhance the identity and resolvability of patterns in the central brain, we separated visual projection neurons from central brain neurons, and further divided the latter into 3 groups based on their layer assignment. We further provide ROI-level information for the most contributing brain regions to provide more anatomical context.

To produce a map of binocular neurons (**Fig. 4i**), we define a binocularity index as the ratio of the smaller to the larger VIC score from the left and right optic lobes, ranging from 0 (monocular) to 1 (balanced binocular). In the maps of **Fig. 4h,i** and **S5, S6**, we exclude those neuron types whose median VIC is less than 5 × 10^-4^. **Fig.5a** represents the distribution of the smallest ARF size, that is, the synapses representing the highest resolution. Please refer to the section **Anatomical receptive fields and 2D Gaussian fits** for the computation of this value. We applied an additional criterion to filter out neurons with ARF goodness-of-fit R^2^ < 0.05, as most, but not all, neurons with small ARFs are well fit by a 2D Gaussian (**Fig. 3b**).

Accompanying these extreme value projections are additional summary plots that aim to illustrate various aspects of these metrics. For **Fig. 4h** we selected the six ROIs with the highest synapse count from VCBNs for each group, and represented the fraction of synapses from VCBNs in each ROI with a color-mapped fraction. To emphasize the most binocular neurons, in **Fig.4i** we selected the six ROIs with the highest numbers of synapses from VCBNs neurons with a binocularity index > 0.5. The cumulative plots in **Fig.4i, 5a** illustrate the distribution of the corresponding metrics in each ROI, which is not apparent from the extreme value projections themselves. The heatmaps and cumulative plots were generated with custom Python code using matplotlib.

### Visual spatial coverage by ROI

To explore potential spatial sampling bias, we focus on individual ROIs in the central brain and construct their coverage maps, as shown in **Fig. 5b, S7e**. We first collected all visual neurons (defined by the VIC > 5 × 10^-4^ criterion) that project into a given ROI (e.g. that ROI contains pre-synapses from these neurons). We then summed these neurons’ ARFs, weighted by each neuron’s pre-synapse count in that ROI. To facilitate comparison, we grouped the column-resolution ARFs into five sectors corresponding to the dorsal, ventral, anterior, posterior, and central regions of the eye, selected to approximate near uniform column counts per sector, and represented the total regional visual contributions as a fraction, with 0.2 representing uniform coverage (for 5 sectors). In **Fig. 5b** the ROI positions were manually arranged from the original projection view to maximize clarity. The example ARFs in **Fig. 5a**, bottom and the spatial coverage plots in **Fig. 5b** were implemented using the Connectome Interpreter and Plotly.

### Visual RF center brain mapping

To further explore the sampling and representation of visual space in the central brain, we developed a visualization of the 3D pre-synapse locations of VCBNs across the central brain ROIs and colored-coded them based on their parent neuron’s ARF center-of-mass (based on the 2D Gaussian fit) in the retinal coordinates. The visualization was implemented in the powerful, web-browser-based 3D viewer, Neuroglancer^105^. For the views shown in **Fig. 5c**, we used all VCBN synapses (neurons with VIC > 5 × 10^-4^) across all layers but restrict to only ARF goodness-of-fit R^2^ > -0.5, and only show a subset of the points (using the spacing projection parameter of 1.0-3.0 px 988/988, radius = 1.5). The images are generated using screenshots from Neuroglancer in standard brain orientations and with an orthographic projection.

## Declaration of generative AI and AI-assisted technologies in the manuscript preparation process

During the preparation of this work, the authors used ChatGPT to edit and improve the draft’s clarity, and used Copilot to assist with writing analysis and visualization code. After using these tools, the authors reviewed and edited the content and take full responsibility for the content of the published article.

