## Supplementary material for "The organization of visual pathways in the *Drosophila* brain": Doc S1

### Supplementary Figures: S1-S7

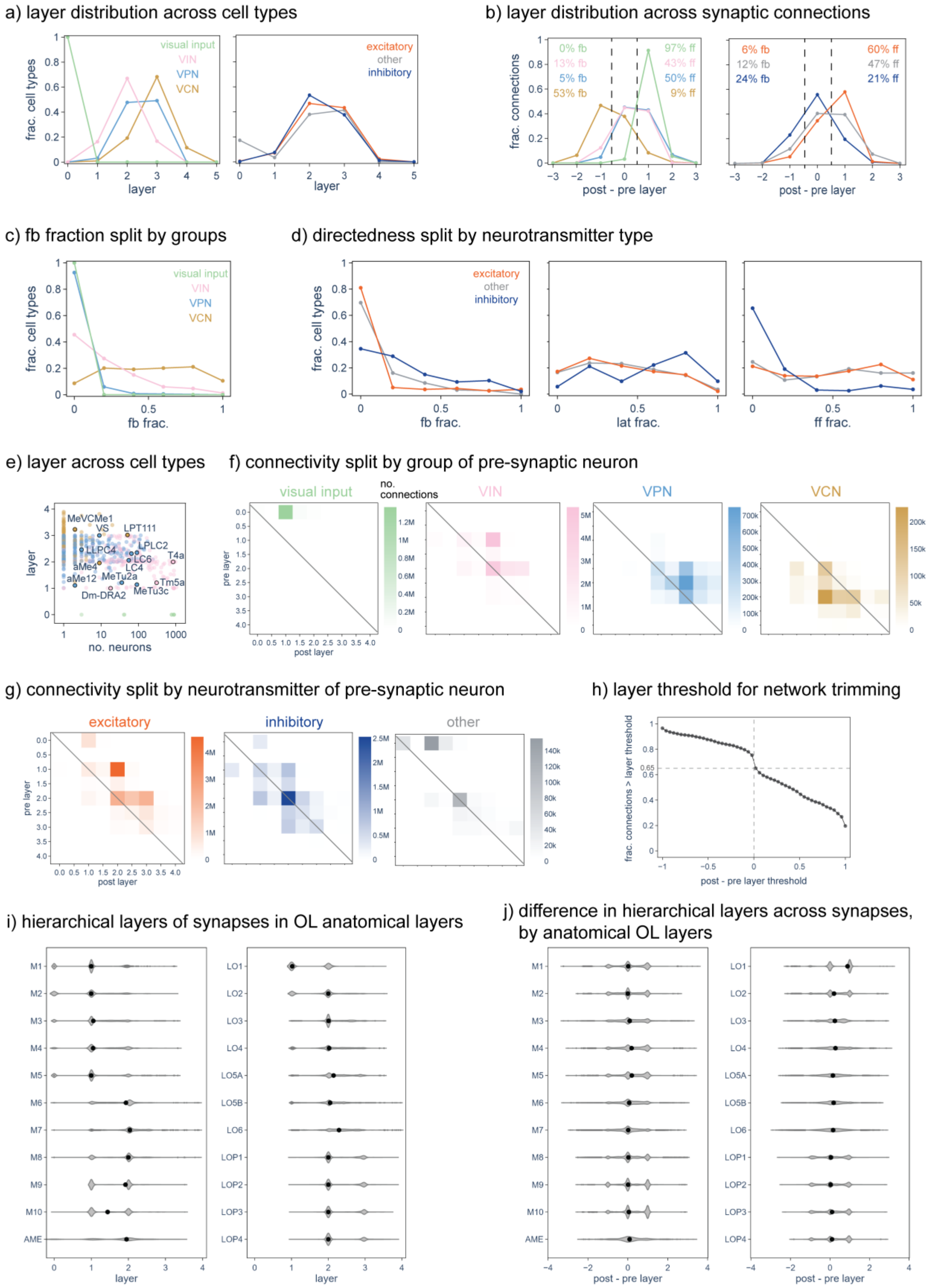

**(previous page)**

**Fig. S1. Layer distributions and connectivity patterns in the right-side optic lobe (related to Fig. 1)**

**a)** Normalized distribution of hierarchical layers across cell types, split by cell group (left) and neurotransmitter type (right). **b)** Normalized distribution of layer differences across synaptic connections (pos-pre), weighted by the number of connections, split by cell group (left) and neurotransmitter type (right). Dashed lines at $\pm 0.5$ separate feedback (fb), lateral (lat), and feedforward (ff) connections, and percentages indicate the fraction of feedback, lateral, and feedforward connections within each group. **c)** Distribution of feedback fractions across cell types, split by cell group. **d)** Distributions of feedback, lateral, and feedforward fractions across cell types, split by neurotransmitter type. **e)** Number of neurons per cell type versus layer, with several example cell types highlighted. **f)** Number of synaptic connections between layers split by pre-synaptic neurons in each cell group; post-synaptic neurons include all neuron types, not restricted to the right optic lobe. Layers were binned at half-integer values. **g)** Similar to **f**, but split by presynaptic neurotransmitter type. **h)** Fraction of connections retained as a function of the layer-difference threshold used for network trimming. A threshold of 0 retains ~65% of all connections. **i)** Hierarchical layers of synapses plotted against anatomical optic lobe layers. M1-M10: medulla, LO1-LO6: lobula, LOP1-LOP4: lobula plate, and AME: accessory medulla, undivided. Black dots indicate medians. **j)** Similar to **i** but for the layer difference (post-pre).

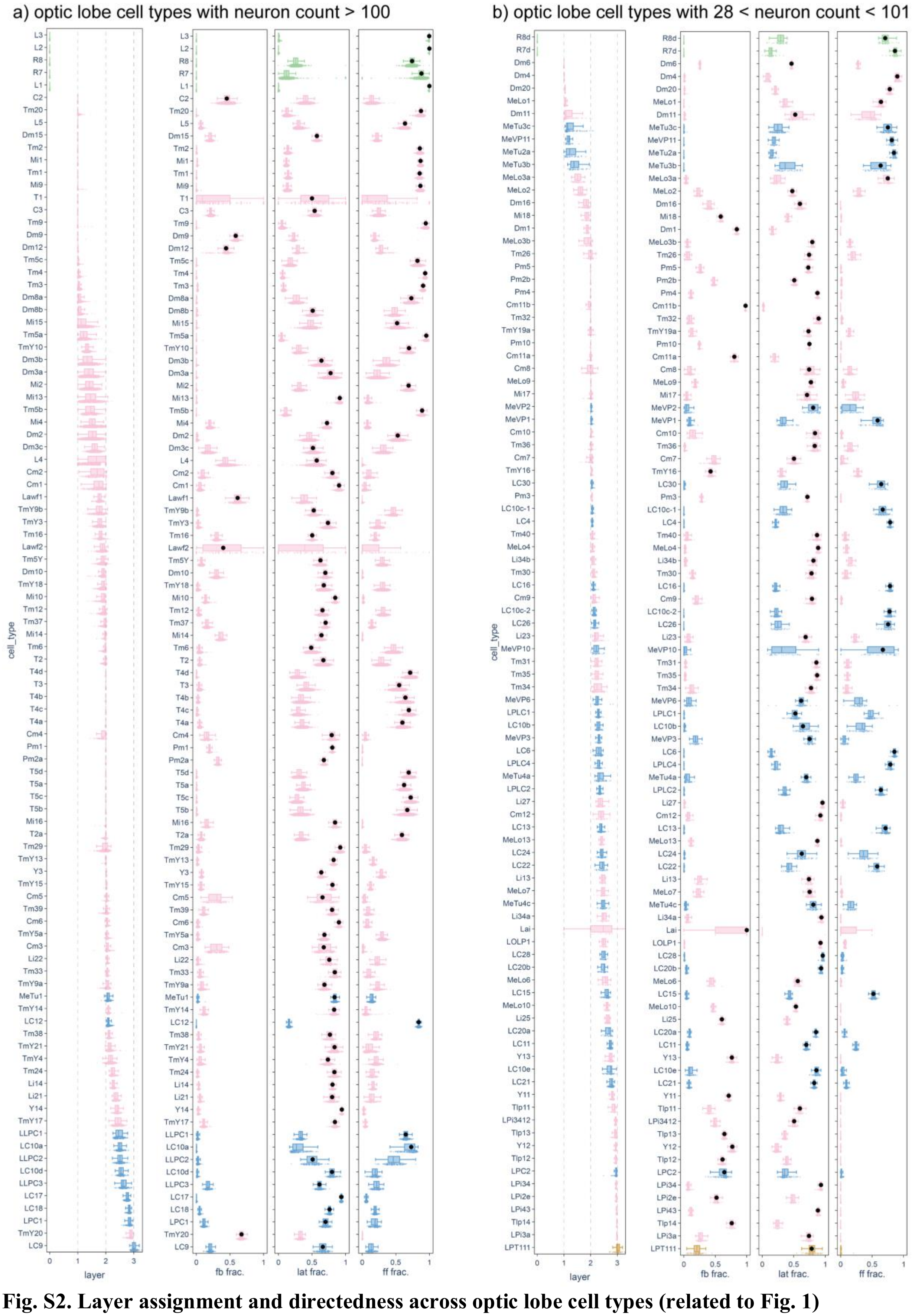


**Fig. S2. Layer assignment and directedness across optic lobe cell types (related to Fig. 1)**
Layer assignment and directedness for individual neurons grouped by cell type in the right optic lobe (as in **Fig. 1g,j**). **a)** Cell types with > 100 neurons. **b)** Cell types with 28-100 neurons. Cell types are ordered by median layer. Black dots indicate medians, but for directedness, the black dot is only shown for the largest median fraction.

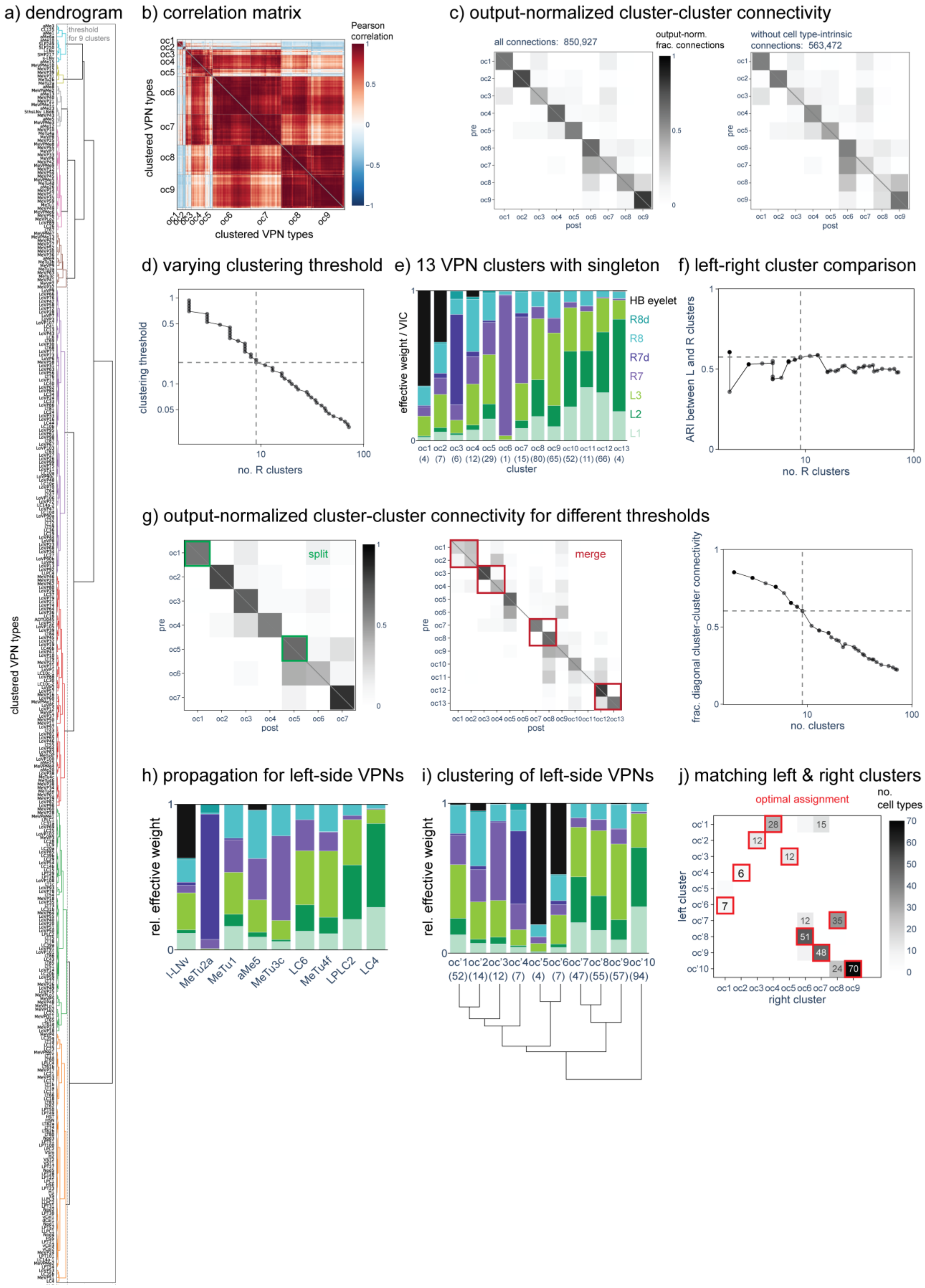

a) dendrogram
threshold for 9 clusters
clustered VPN types
b) correlation matrix
Pearson correlation
oc1
oc2
oc3
oc4
oc5
oc6
oc7
oc8
oc9
clustered VPN types
c) output-normalized cluster-cluster connectivity
all connections: 850,927
output-norm. frac. connections
without cell type-intrinsic connections: 563,472
pre
post
d) varying clustering threshold
clustering threshold
no. R clusters
e) 13 VPN clusters with singleton
effective weight / VIC
HB eyelet
R8d
R8
R7d
R7
L3
L2
L1
oc1 oc2 oc3 oc4 oc5 oc6 oc7 oc8 oc9 oc10 oc11 oc12 oc13
(4) (7) (6) (12) (29) (1) (15) (80) (65) (52) (11) (66) (4)
cluster
f) left-right cluster comparison
ARI between L and R clusters
no. R clusters
g) output-normalized cluster-cluster connectivity for different thresholds
split
merge
pre
post
frac. diagonal cluster-cluster connectivity
no. clusters
h) propagation for left-side VPNs
rel. effective weight
l-LNv
MeTu2a
MeTu1
aMe5
MeTu3c
LC6
MeTu4f
LPLC2
LC4
i) clustering of left-side VPNs
rel. effective weight
oc'1 oc'2 oc'3 oc'4 oc'5 oc'6 oc'7 oc'8 oc'9 oc'10
(52) (14) (12) (7) (4) (7) (47) (55) (57) (94)
j) matching left & right clusters
optimal assignment
no. cell types
left cluster
right cluster
oc'1
oc'2
oc'3
oc'4
oc'5
oc'6
oc'7
oc'8
oc'9
oc'10
oc1 oc2 oc3 oc4 oc5 oc6 oc7 oc8 oc9

**(previous page)**
**Fig. S3. Hierarchical clustering and left-right comparison of VPN pathway classes (related to Fig. 2)**
**a)** Full dendrogram from hierarchical clustering of the 352 VPN cell types in the right optic lobe (**Fig. 2e** shows the portion of the dendrogram to the right of the flat cut at 9 clusters). **b)** Pearson correlation matrix of relative effective weights between VPN cell types, ordered as in panel **a**. Horizontal and vertical gray lines delineate cell types belonging to the same cluster (oc1-oc9). **c)** Output-normalized number of connections between neurons in each VPN cluster (each column sums to 1). The title indicates the total number of connections. The right matrix excludes connections between neurons of the same cell type (and is renormalized so each column again sums to 1). **d)** By varying the clustering threshold (flat cut of the dendrogram in panel **a**, given as a fraction relative to the total height), different numbers of clusters were obtained. The dashed line marks the threshold used in the main text, resulting in 9 clusters. **e)** Mean relative effective weight per cluster for the cut yielding 13 clusters (analogue of **Fig. 2e**). Because cluster oc6 contains only a single cell type (a "singleton" cluster), we did not adopt the 13-cluster assignment. **f)** Comparison of clustering performed on the homologous VPN cell types in the right and left optic lobe, quantified by the adjusted rand index (ARI). For the comparison, we used the same clustering threshold on the left and right. **g)** Similar to panel **c** (left) but for thresholds yielding 7 or 13 clusters. Highlighted blocks indicate clusters that would split or merge to obtain the 9-cluster assignment. The rightmost panel shows the fraction of diagonal entries in the output-normalized cluster-cluster connectivity matrices (relative to the sum of all entries) as a function of the number of clusters. **h)** Example relative effective weights for selected VPN cell types in the left optic lobe (similar to **Fig. 2d** but for the left side). **i)** Using the clustering threshold highlighted in panel **d**, we obtained 10 clusters for left side VPNs, which are denoted (oc'1- oc'9). Bar plots show the mean relative effective weight for each left-side cluster, along with the corresponding dendrogram (left-side analog of **Fig. 2e**). **j)** Confusion matrix between left- and right-side VPN clusters, with optimal matches indicated by red squares.

**(next page)**
**Fig. S4. Hierarchical clustering and left-right comparison of VCBNs (related to Fig. 4).**
**a)** Comparison of layers for homologous VPN and CBN cell types. The histogram shows the marginal distribution of left-side layers. **b)** Comparison of VICs for homologous CBN cell types. CBNs are separated into three groups based on the sum of left and right VICs. The histogram shows the distribution along the diagonal for each group. The groups with higher VIC scores also show greater left-right consistency. **c)** Full dendrogram from hierarchical clustering of the 5,558 VCBN cell types reached from the right optic lobe (cf. **Fig. 4a**; note that **Fig. 4d** only includes part of the dendrogram). Because of their numerosity, cell type names are not shown. **d)** Output-normalized number of connections between neurons in each VCBN cluster (each column sums to 1). Right-side VCBN clusters are denoted cc1-cc9. The title states the total number of connections. **e)** By varying the clustering threshold (flat cut of the dendrogram in panel **c**, normalized by the total height), different numbers of VCBN clusters were obtained. The dashed line marks the threshold used in the main text (9 clusters). **f)** Comparison of clustering performed on homologous VCBN cell types reached from the right and left optic lobes, quantified by the adjusted Rand index (ARI). **g)** Relative effective weights for selected VCBN cell types reached from the left optic lobe visual inputs (similar to **Fig. 4c**). **h)** Using the clustering threshold highlighted in panel **e**, we also obtained 9 VCBN clusters for propagation from left-side visual inputs. Left-side VCBN clusters are denoted as cc'1- cc'9. Bar plots show the mean relative effective weight for each left-side cluster together with the dendrogram (similar to **Fig. 4d**). **i)** Confusion matrix comparing left- and right-side VCBN clusters, with optimal matches indicated by red squares. **j)** Matrix showing the number of connections from VPNs in each oc cluster to VCBNs in each cc cluster. Grayscale values indicate output-normalized fractions (left, each row sums to 1) or input-normalized fractions (right, each column sums to 1). Only entries with fractions (of connections) > 0.05 are displayed. **k)** Normalized distribution of number of VCBN clusters with participation fraction pi≥1/9 across VPN cell types (similar to **Fig. 2h** but for VPN to VCBN clusters). A uniform distribution would have pi=1/9≈0.11, corresponding to equal participation across all 9 cc clusters. **l)** For each central brain neuron (CBN) type , we assign two VIC scores, one for visual input propagation starting in each optic lobe. Unreached cell types are assigned a VIC of $10^{-6}$. Cell types are split into three groups based on the maximum and minimum of their left and right VICs: those with max(VIC) < $5 \times 10^{-4}$, those with min(VIC) < $5 \times 10^{-4}$ and max(VIC) $\geq 5 \times 10^{-4}$, and those with min(VIC) $\geq 5 \times 10^{-4}$. For binocularity in Fig. **4i**, we used only cell types in groups 2 and 3.

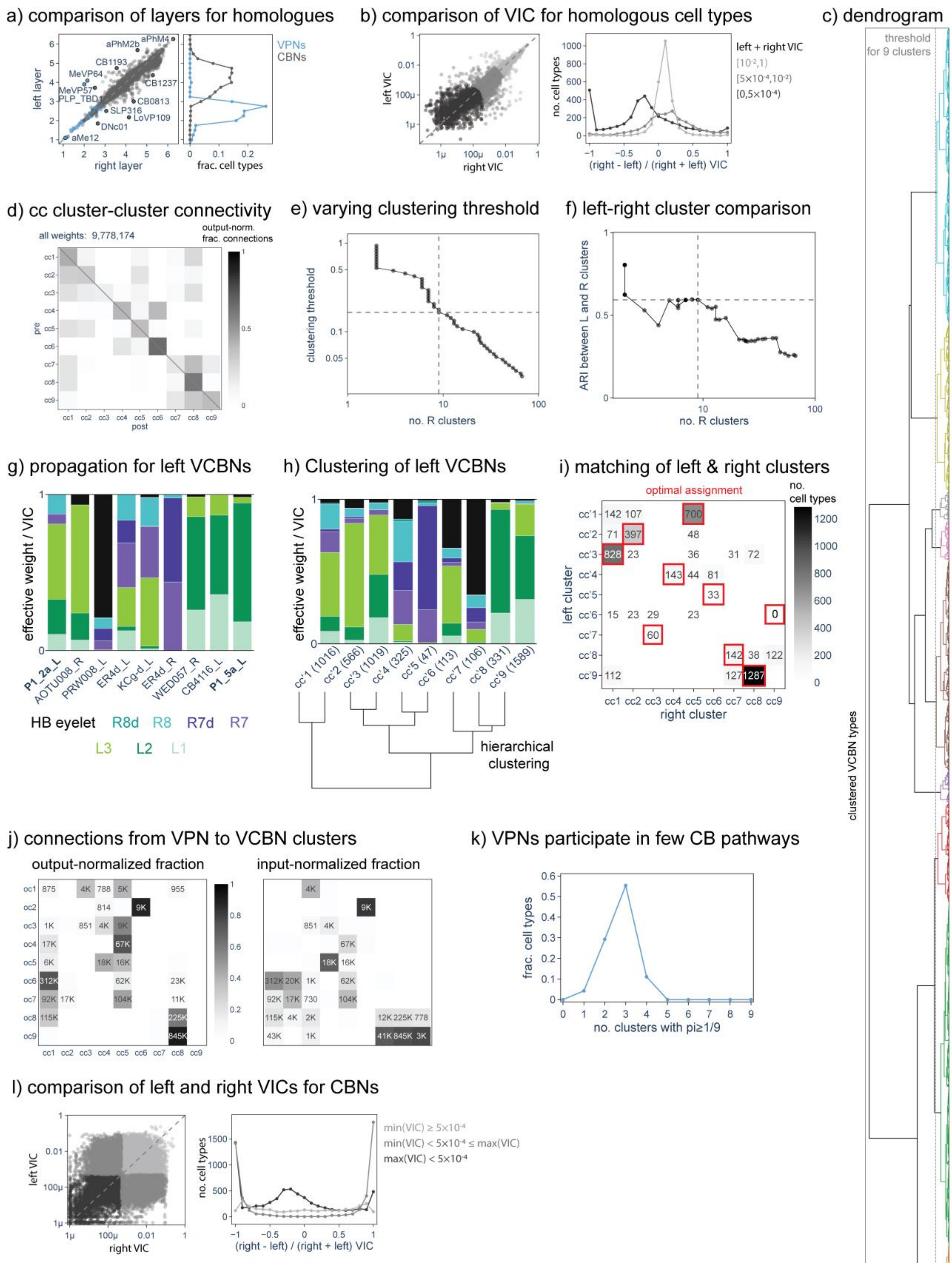
a) comparison of layers for homologues
left layer
right layer
frac. cell types
VPNs
CBNs
aPhM2b
aPhM4
CB1193
MeVP64
MeVP57
PLP_TBD1
CB1237
CB0813
SLP316
LoVP109
DNc01
aMe12
b) comparison of VIC for homologous cell types
left VIC
right VIC
no. cell types
(right - left) / (right + left) VIC
left + right VIC
c) dendrogram
threshold for 9 clusters
clustered VCBN types
d) cc cluster-cluster connectivity
all weights: 9,778,174
output-norm. frac. connections
pre
post
e) varying clustering threshold
clustering threshold
no. R clusters
f) left-right cluster comparison
ARI between L and R clusters
no. R clusters
g) propagation for left VCBNs
effective weight / VIC
P1_2a_L
AOTU008_R
PRW008_L
ER4d_L
KCg-d_L
ER4d_R
WED057_R
CB4116_L
P1_5a_L
HB eyelet
R8d
R8
R7d
R7
L3
L2
L1
h) Clustering of left VCBNs
effective weight / VIC
cc'1 (1016)
cc'2 (566)
cc'3 (1019)
cc'4 (325)
cc'5 (47)
cc'6 (113)
cc'7 (106)
cc'8 (331)
cc'9 (1589)
hierarchical clustering
i) matching of left & right clusters
optimal assignment
no. cell types
left cluster
right cluster
j) connections from VPN to VCBN clusters
output-normalized fraction
input-normalized fraction
k) VPNs participate in few CB pathways
frac. cell types
no. clusters with pi≥1/9
l) comparison of left and right VICs for CBNs
left VIC
right VIC
no. cell types
(right - left) / (right + left) VIC
min(VIC) ≥ 5×10⁻⁴
min(VIC) < 5×10⁻⁴ ≤ max(VIC)
max(VIC) < 5×10⁻⁴

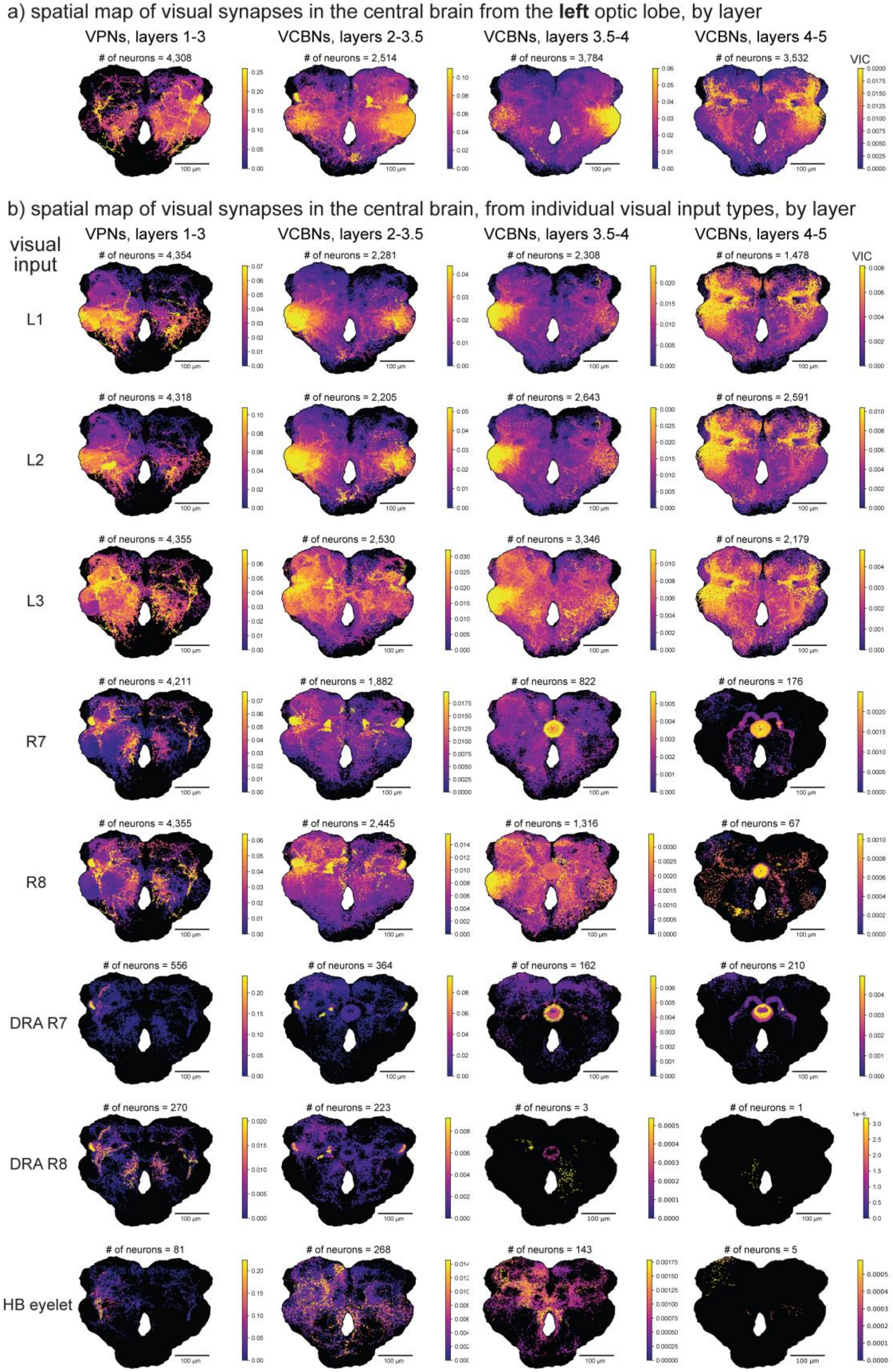
a) spatial map of visual synapses in the central brain from the **left** optic lobe, by layer
VPNs, layers 1-3
VCBNs, layers 2-3.5
VCBNs, layers 3.5-4
VCBNs, layers 4-5
### of neurons = 4,308
### of neurons = 2,514
### of neurons = 3,784
### of neurons = 3,532
VIC
b) spatial map of visual synapses in the central brain, from individual visual input types, by layer
visual input
VPNs, layers 1-3
VCBNs, layers 2-3.5
VCBNs, layers 3.5-4
VCBNs, layers 4-5
VIC
L1
### of neurons = 4,354
### of neurons = 2,281
### of neurons = 2,308
### of neurons = 1,478
L2
### of neurons = 4,318
### of neurons = 2,205
### of neurons = 2,643
### of neurons = 2,591
L3
### of neurons = 4,355
### of neurons = 2,530
### of neurons = 3,346
### of neurons = 2,179
R7
### of neurons = 4,211
### of neurons = 1,882
### of neurons = 822
### of neurons = 176
R8
### of neurons = 4,355
### of neurons = 2,445
### of neurons = 1,316
### of neurons = 67
DRA R7
### of neurons = 556
### of neurons = 364
### of neurons = 162
### of neurons = 210
DRA R8
### of neurons = 270
### of neurons = 223
### of neurons = 3
### of neurons = 1
HB eyelet
### of neurons = 81
### of neurons = 268
### of neurons = 143
### of neurons = 5
100 μm

**(previous page)**

**Fig. S5 Additional spatial maps of visual synapses in the central brain (related to Fig. 4).**
**a)** Spatial maps of presynaptic sites from VPNs/VCBNs reached via *left-side* optic-lobe propagation, separated by hierarchical layer groups (brain collapsed to two dimensions using maximum-intensity projection, as in **Fig. 4h**). Color indicates VIC value for each presynapse; ranges are rescaled per panel. **b)** Projected spatial maps of the visual contributions, represented at presynaptic sites of VPNs and VCBNs, by each visual input type (L1, L2, L3, R7, R8, R7d, 8d, and HB eyelet), with propagation performed from all cells of one input at a time. Panels are grouped by hierarchical layer (brain collapsed to two dimensions using maximum-intensity projection, as in **Fig. 4h**). Color indicates the VIC score for that input, and the color scales differ across panels to capture the dynamic range of the data. Listed numbers indicate the total neurons included in each map (distinct from **Fig. 4d**, which reports cell-type counts).

**(next page)**

**Fig. S6 Spatial maps of visual central brain neurons by central brain pathway class (related to Fig. 4).**
2D projected spatial distribution of VCBN presynapses for each central brain pathway class (cc1-cc9), split by hierarchical layer groups (brain collapsed to two dimensions using maximum-intensity projection, as in **Fig. 4h**). Color indicates the VIC score (across all visual inputs) for the presynapses of the VCBNs within each pathway class. Color scales differ across panels to capture the dynamic range of the data. Total numbers of neurons included in each map are indicated.

a) spatial map of VCBN synapses, by central brain pathway class

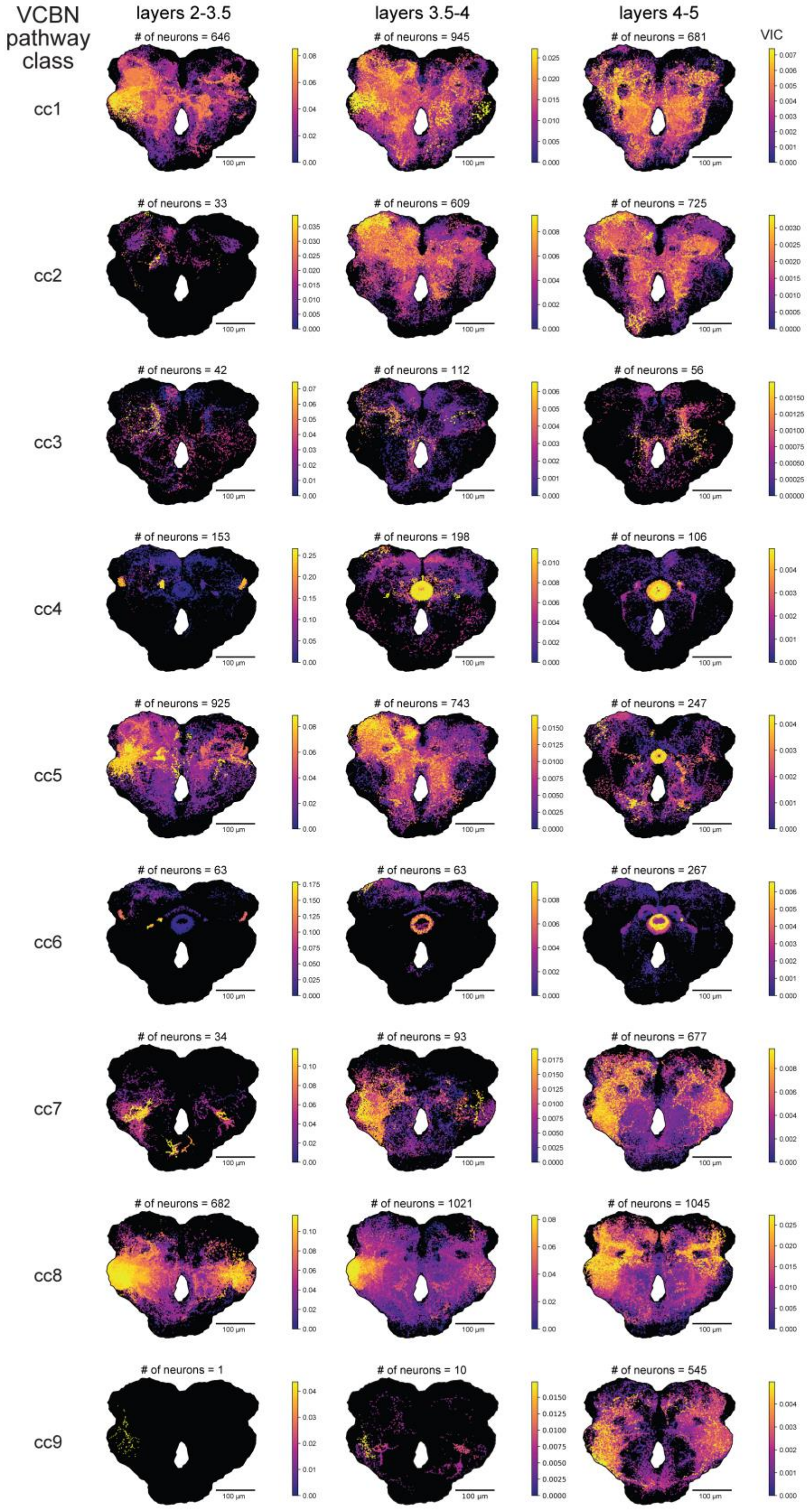

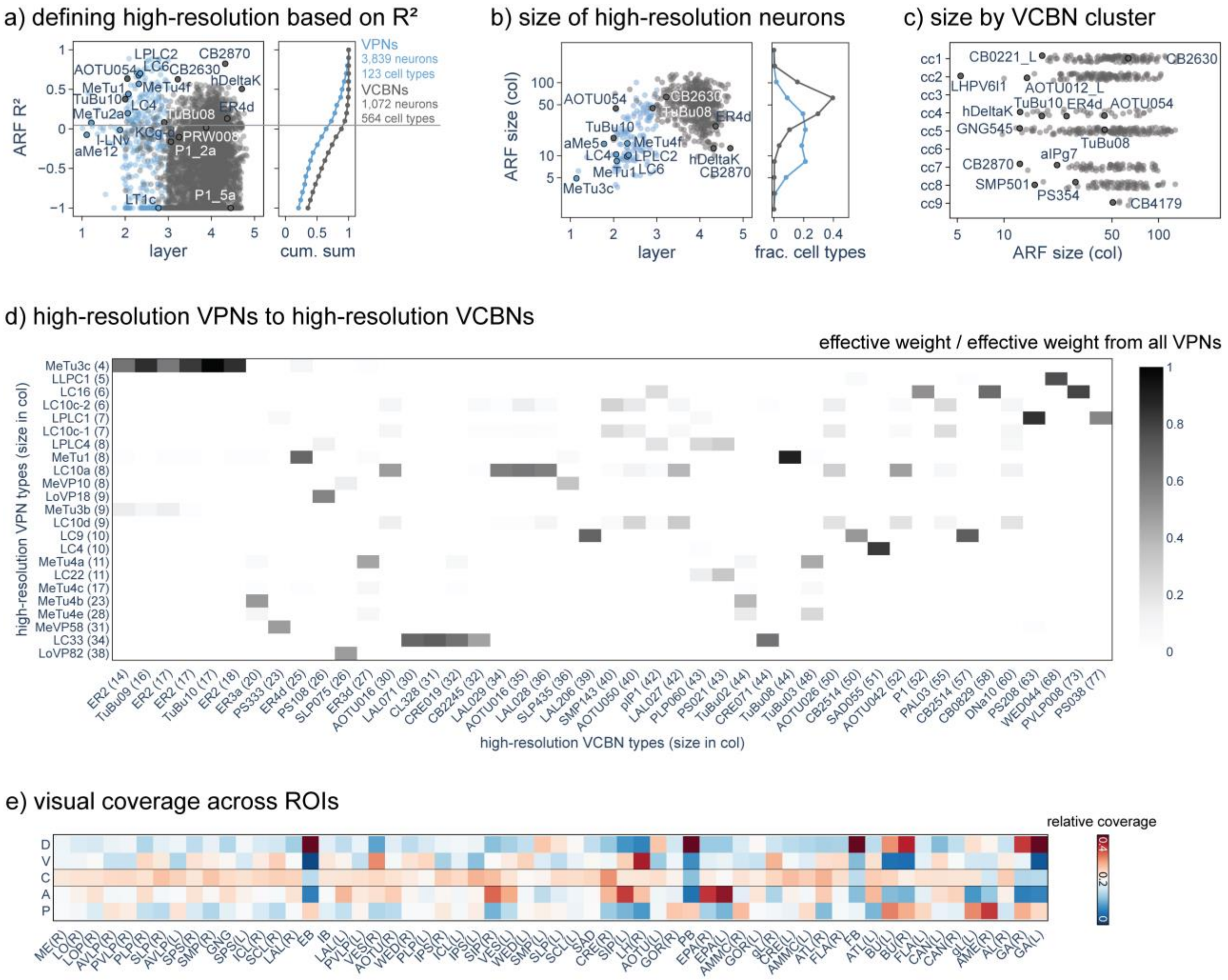


**Fig. S7. Additional analyses of a high-resolution spatial vision (related to Fig. 5).**
**a)** Goodness-of-fit of the ARF Gaussians was used to select high-resolution cell types; for **Fig. 5a** only cell types with ARF $R^2 > 0.05$ (above solid line) were included. The histogram shows the marginal cumulative distribution of the ARF $R^2$ values. Cell types in clusters dominated by HB eyelet or DRA photoreceptors, i.e., oc1-oc2 (**Fig. 2e**) and cc3, cc6 (**Fig. 4d**) we also excluded. The numbers of neurons and cell types satisfying these criteria are shown above the solid line. **b)** ARF size of high-resolution VPN and VCBN cell types as a function of layer, with the marginal distribution of ARF size shown at right. **c)** ARF size of high-resolution cell types split by VCBN cluster. **d)** Strongest indirect connections from high-resolution VPN to high-resolution VCBN types. The heatmap shows the effective weight from each VPN type to each VCBN type, divided by the sum of effective weights from all VPNs—a quantity analogous to the relative effective weight used for visual input propagation, but here computed starting from VPNs. **e)** Relative visual coverage quantifies whether an ROI contains more synapses sampling a particular sector of visual space. Visual space is divided into 5 sectors (D: dorsal, V: ventral, C: central, A: anterior, P: posterior), and compute the relative contribution of propagated visual inputs in each sector to all the presynapses in each brain ROI. These values sum to 1 across all five sectors. Uniform visual coverage would have a value of 1/5=0.2 (for all sectors). The color code emphasizes reductions (blue) or increases (red) from uniform.
