## Supplementary material for "The organization of visual pathways in the *Drosophila* brain": Doc S2

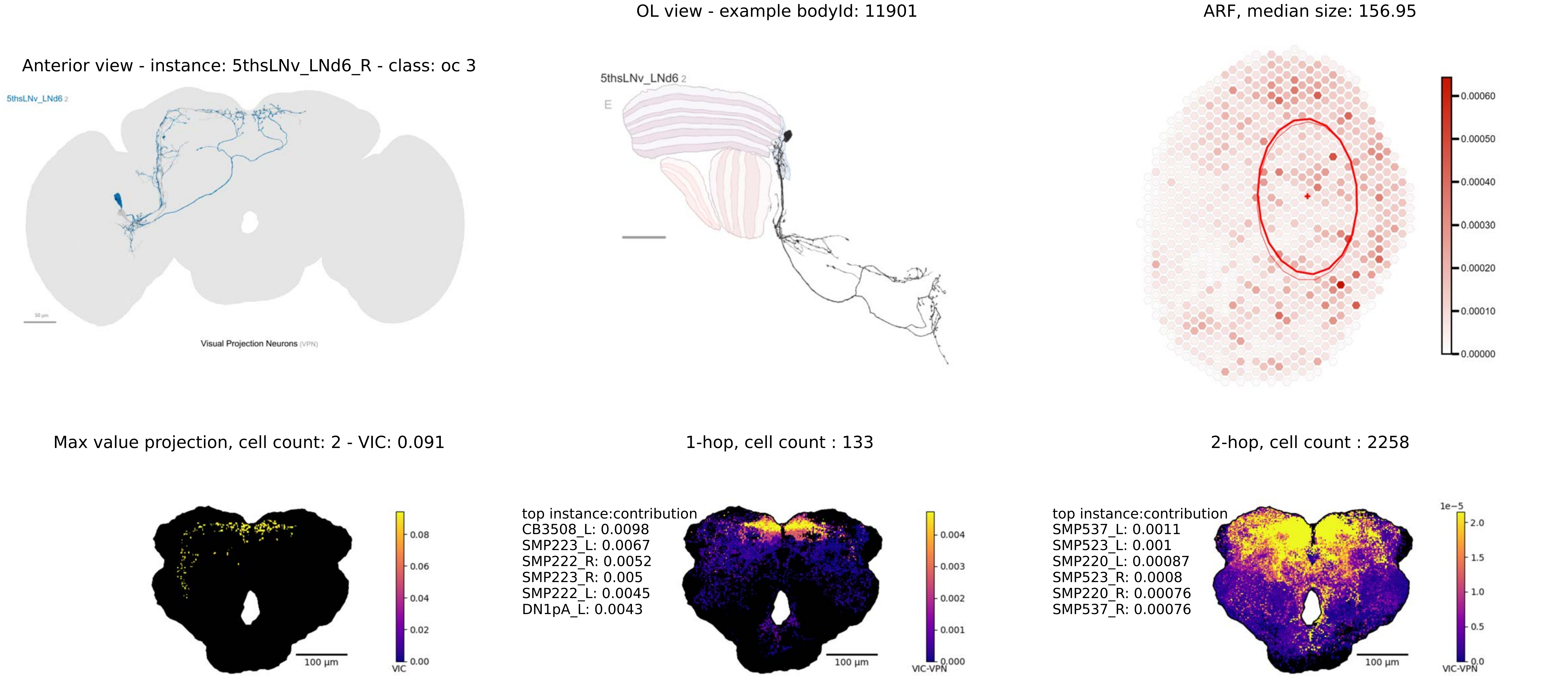
Anterior view - instance: 5thsLNv_LNd6_R - class: oc 3
5thsLNv_LNd6 2
Visual Projection Neurons (VPN)
OL view - example bodyId: 11901
5thsLNv_LNd6 2
E
ARF, median size: 156.95
0.00060
0.00050
0.00040
0.00030
0.00020
0.00010
0.00000
Max value projection, cell count: 2 - VIC: 0.091
100 µm
0.08
0.06
0.04
0.02
0.00
VIC
1-hop, cell count : 133
top instance:contribution
CB3508_L: 0.0098
SMP223_L: 0.0067
SMP222_R: 0.0052
SMP223_R: 0.005
SMP222_L: 0.0045
DN1pA_L: 0.0043
100 µm
0.004
0.003
0.002
0.001
0.000
VIC-VPN
2-hop, cell count : 2258
top instance:contribution
SMP537_L: 0.0011
SMP523_L: 0.001
SMP220_L: 0.00087
SMP523_R: 0.0008
SMP220_R: 0.00076
SMP537_R: 0.00076
100 µm
1e−5
2.0
1.5
1.0
0.5
0.0
VIC-VPN

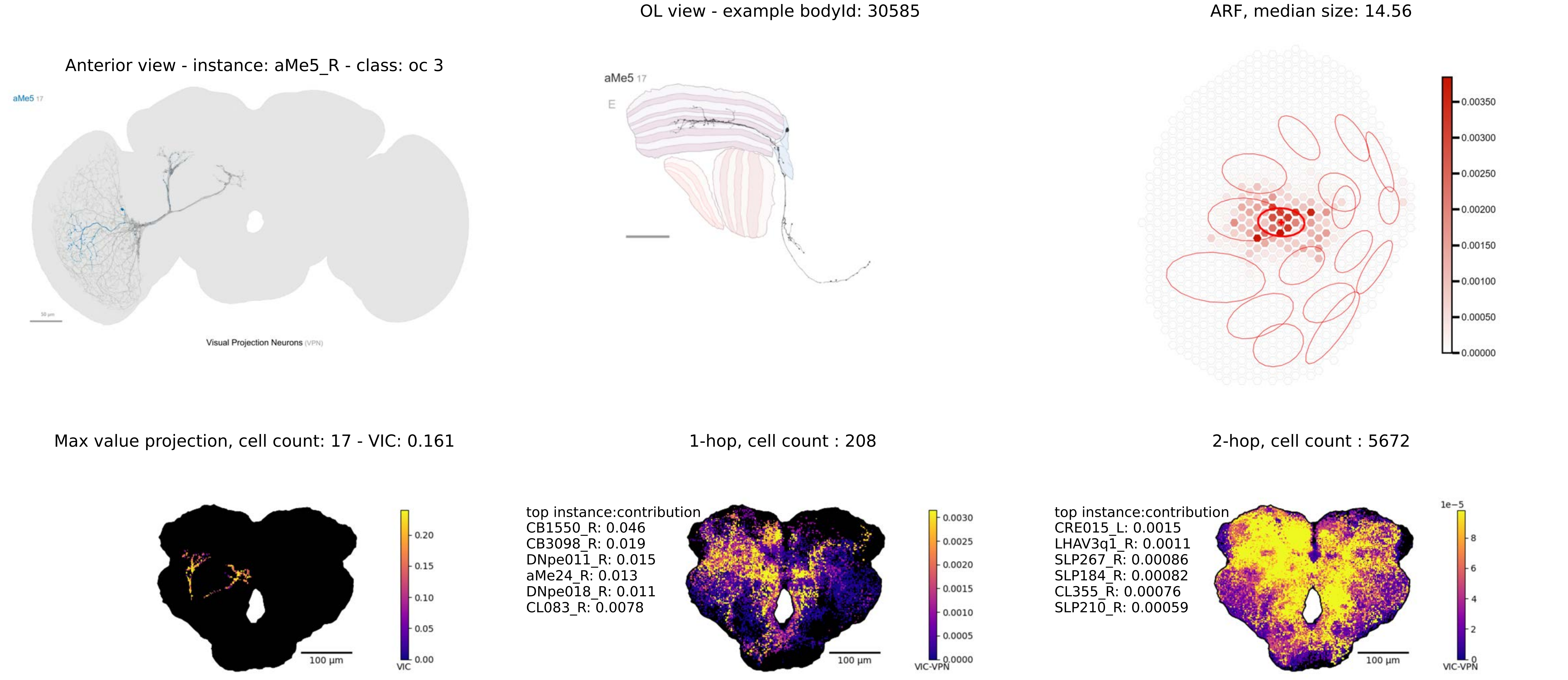
OL view - example bodyId: 30585
ARF, median size: 14.56
Anterior view - instance: aMe5_R - class: oc 3
aMe5 17
Visual Projection Neurons (VPN)
aMe5 17
E
0.00350
0.00300
0.00250
0.00200
0.00150
0.00100
0.00050
0.00000
Max value projection, cell count: 17 - VIC: 0.161
1-hop, cell count : 208
2-hop, cell count : 5672
0.20
0.15
0.10
0.05
0.00
VIC
100 µm
top instance:contribution
CB1550_R: 0.046
CB3098_R: 0.019
DNpe011_R: 0.015
aMe24_R: 0.013
DNpe018_R: 0.011
CL083_R: 0.0078
0.0030
0.0025
0.0020
0.0015
0.0010
0.0005
0.0000
VIC-VPN
100 µm
top instance:contribution
CRE015_L: 0.0015
LHAV3q1_R: 0.0011
SLP267_R: 0.00086
SLP184_R: 0.00082
CL355_R: 0.00076
SLP210_R: 0.00059
1e−5
8
6
4
2
0
VIC-VPN
100 µm

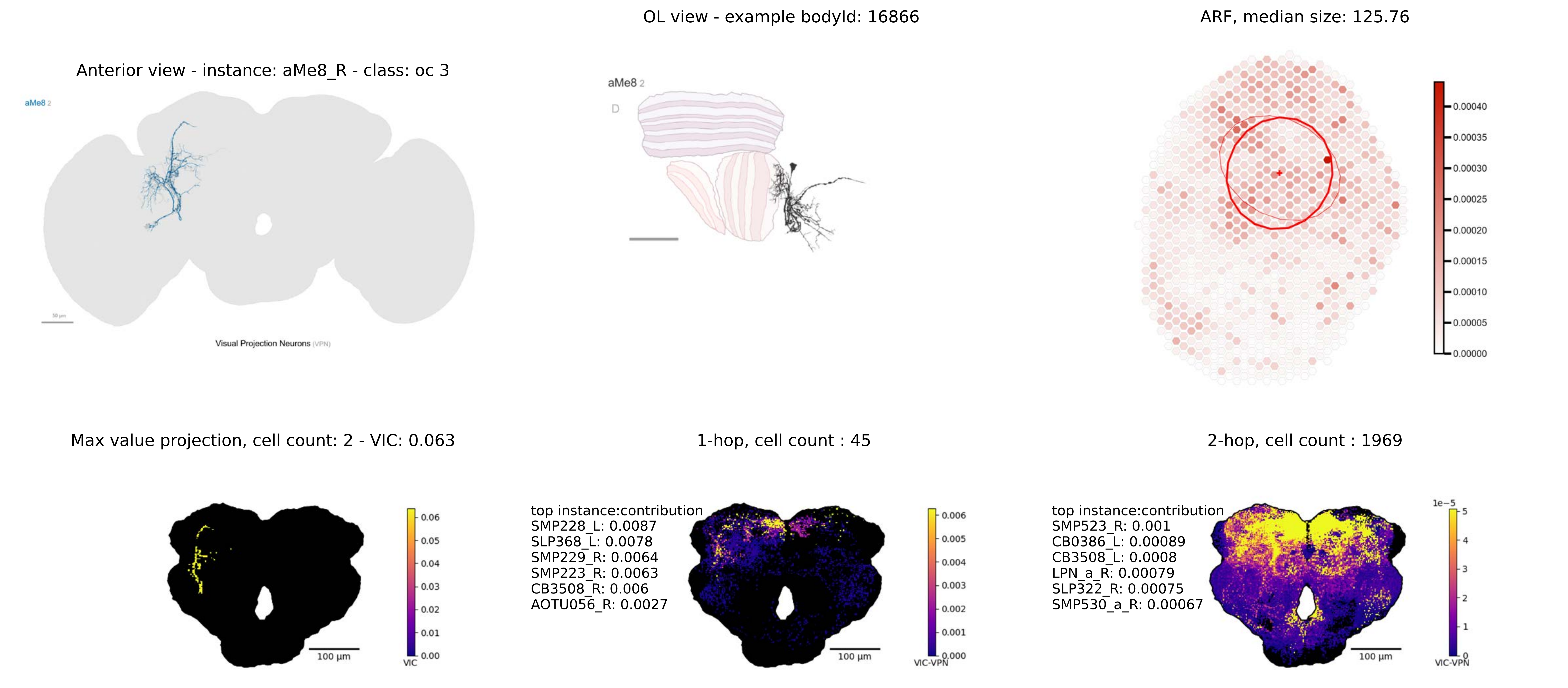

Anterior view - instance: aMe8_R - class: oc 3
aMe8 2
Visual Projection Neurons (VPN)
OL view - example bodyId: 16866
aMe8 2
D
ARF, median size: 125.76
0.00040
0.00035
0.00030
0.00025
0.00020
0.00015
0.00010
0.00005
0.00000
Max value projection, cell count: 2 - VIC: 0.063
100 µm
VIC
1-hop, cell count : 45
top instance:contribution
SMP228_L: 0.0087
SLP368_L: 0.0078
SMP229_R: 0.0064
SMP223_R: 0.0063
CB3508_R: 0.006
AOTU056_R: 0.0027
100 µm
VIC-VPN
2-hop, cell count : 1969
top instance:contribution
SMP523_R: 0.001
CB0386_L: 0.00089
CB3508_L: 0.0008
LPN_a_R: 0.00079
SLP322_R: 0.00075
SMP530_a_R: 0.00067
1e−5
100 µm
VIC-VPN

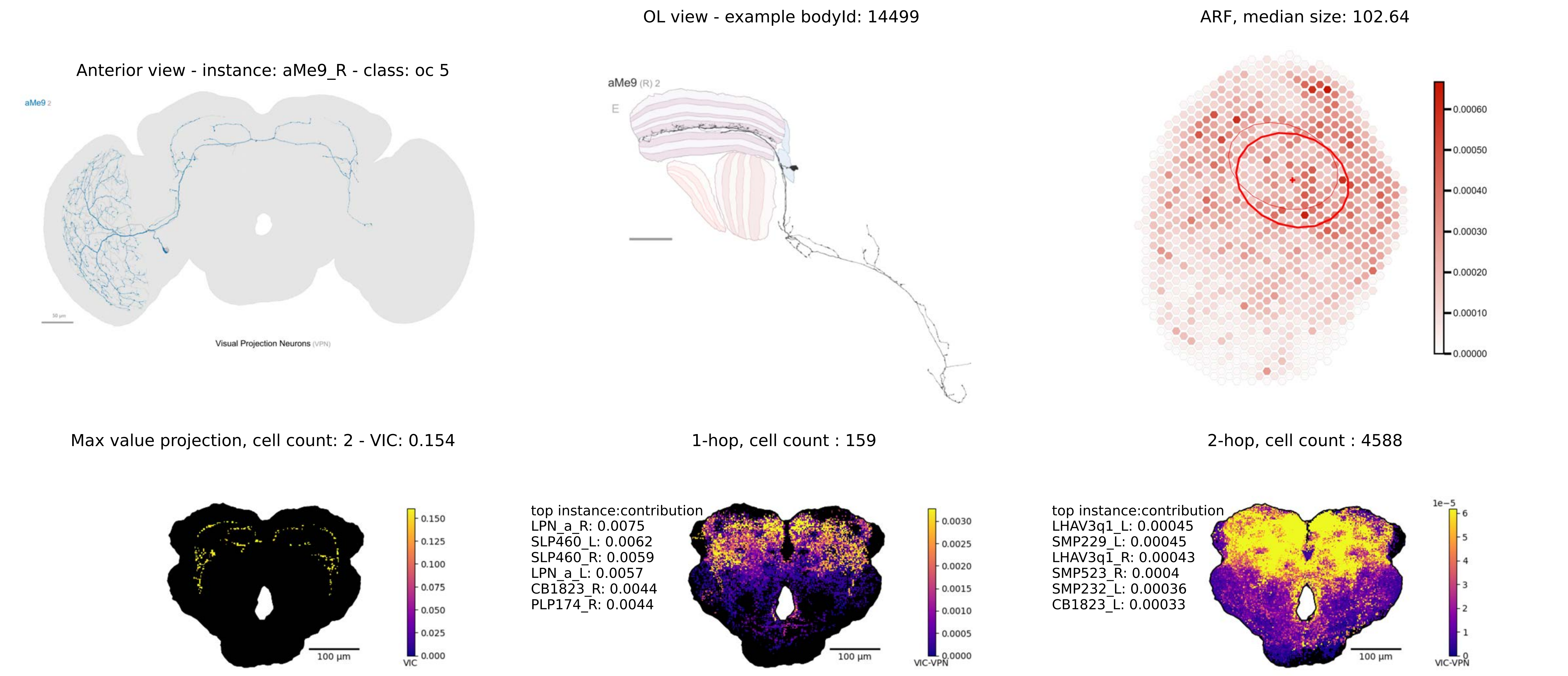
OL view - example bodyId: 14499
ARF, median size: 102.64
Anterior view - instance: aMe9_R - class: oc 5
aMe9 2
Visual Projection Neurons (VPN)
aMe9 (R) 2
E
0.00060
0.00050
0.00040
0.00030
0.00020
0.00010
0.00000
Max value projection, cell count: 2 - VIC: 0.154
1-hop, cell count : 159
2-hop, cell count : 4588
0.150
0.125
0.100
0.075
0.050
0.025
0.000
VIC
100 μm
top instance:contribution
LPN_a_R: 0.0075
SLP460_L: 0.0062
SLP460_R: 0.0059
LPN_a_L: 0.0057
CB1823_R: 0.0044
PLP174_R: 0.0044
0.0030
0.0025
0.0020
0.0015
0.0010
0.0005
0.0000
VIC-VPN
top instance:contribution
LHAV3q1_L: 0.00045
SMP229_L: 0.00045
LHAV3q1_R: 0.00043
SMP523_R: 0.0004
SMP232_L: 0.00036
CB1823_L: 0.00033
1e−5
6
5
4
3
2
1
0
VIC-VPN

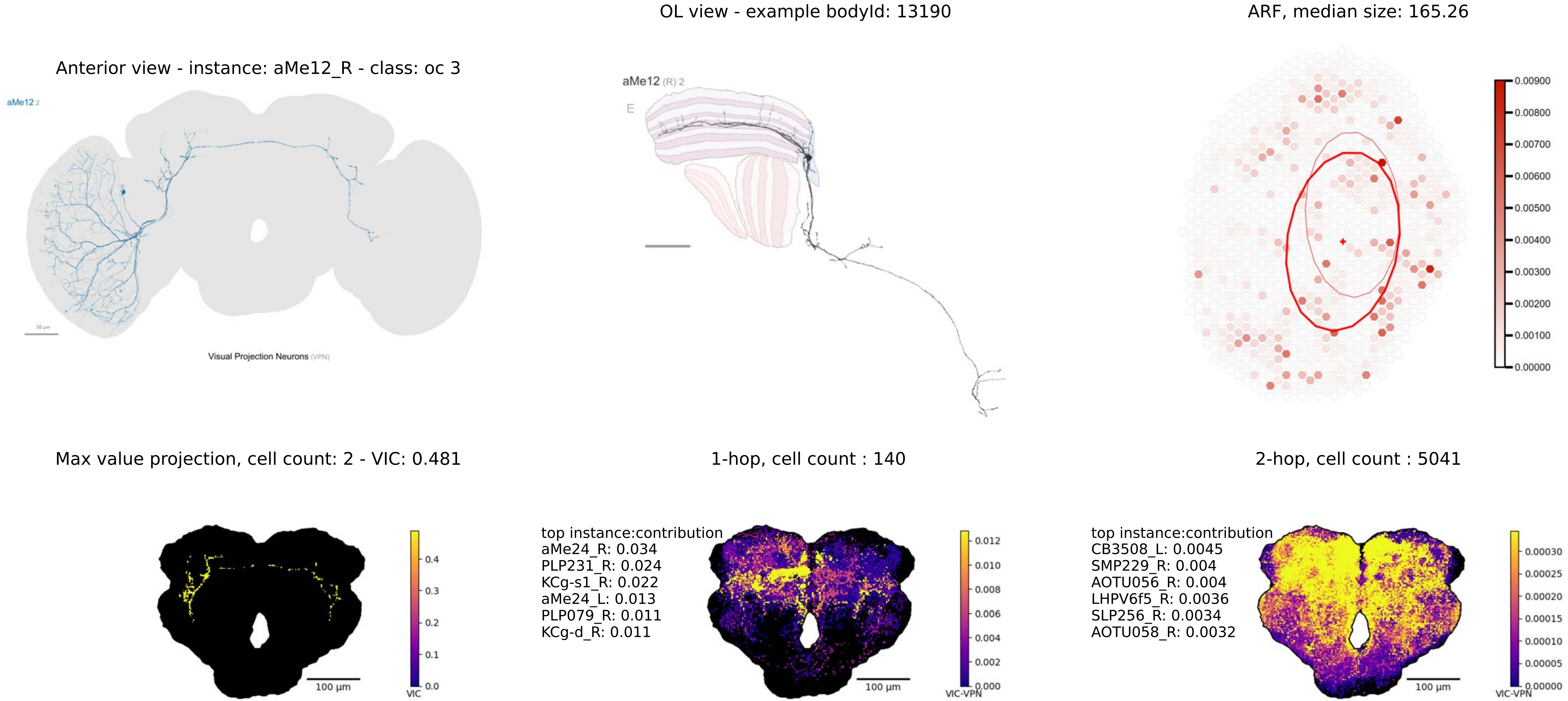

OL view - example bodyId: 13190
ARF, median size: 165.26
Anterior view - instance: aMe12_R - class: oc 3
aMe12 2
Visual Projection Neurons (VPN)
aMe12 (R) 2
E
0.00900
0.00800
0.00700
0.00600
0.00500
0.00400
0.00300
0.00200
0.00100
0.00000
Max value projection, cell count: 2 - VIC: 0.481
1-hop, cell count : 140
2-hop, cell count : 5041
0.4
0.3
0.2
0.1
0.0
VIC
100 µm
top instance:contribution
aMe24_R: 0.034
PLP231_R: 0.024
KCg-s1_R: 0.022
aMe24_L: 0.013
PLP079_R: 0.011
KCg-d_R: 0.011
0.012
0.010
0.008
0.006
0.004
0.002
0.000
VIC-VPN
100 µm
top instance:contribution
CB3508_L: 0.0045
SMP229_R: 0.004
AOTU056_R: 0.004
LHPV6f5_R: 0.0036
SLP256_R: 0.0034
AOTU058_R: 0.0032
0.00030
0.00025
0.00020
0.00015
0.00010
0.00005
0.00000
VIC-VPN
100 µm

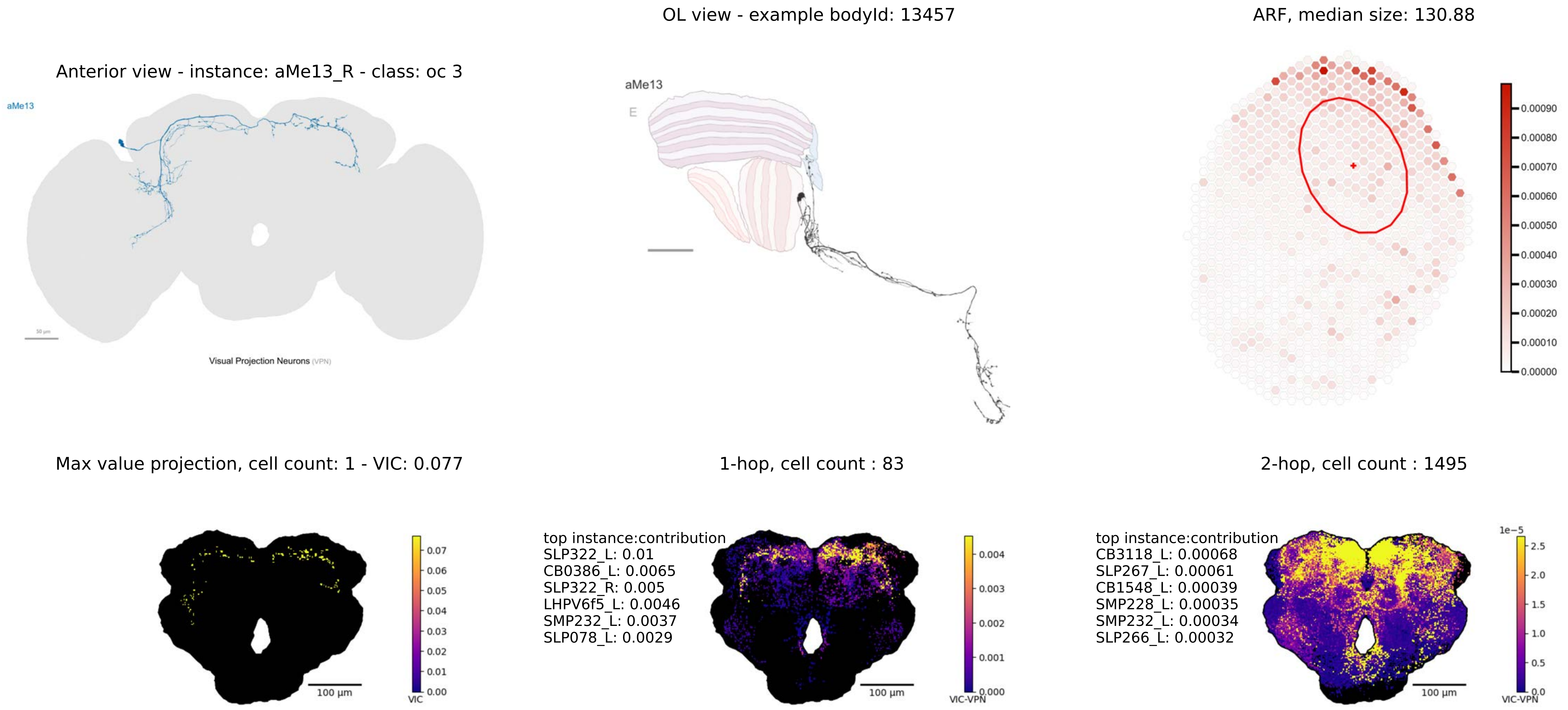
OL view - example bodyId: 13457
ARF, median size: 130.88
Anterior view - instance: aMe13_R - class: oc 3
aMe13
Visual Projection Neurons (VPN)
aMe13
E
0.00090
0.00080
0.00070
0.00060
0.00050
0.00040
0.00030
0.00020
0.00010
0.00000
Max value projection, cell count: 1 - VIC: 0.077
1-hop, cell count : 83
2-hop, cell count : 1495
100 µm
0.07
0.06
0.05
0.04
0.03
0.02
0.01
0.00
VIC
top instance:contribution
SLP322_L: 0.01
CB0386_L: 0.0065
SLP322_R: 0.005
LHPV6f5_L: 0.0046
SMP232_L: 0.0037
SLP078_L: 0.0029
100 µm
0.004
0.003
0.002
0.001
0.000
VIC-VPN
top instance:contribution
CB3118_L: 0.00068
SLP267_L: 0.00061
CB1548_L: 0.00039
SMP228_L: 0.00035
SMP232_L: 0.00034
SLP266_L: 0.00032
100 µm
1e−5
2.5
2.0
1.5
1.0
0.5
0.0
VIC-VPN

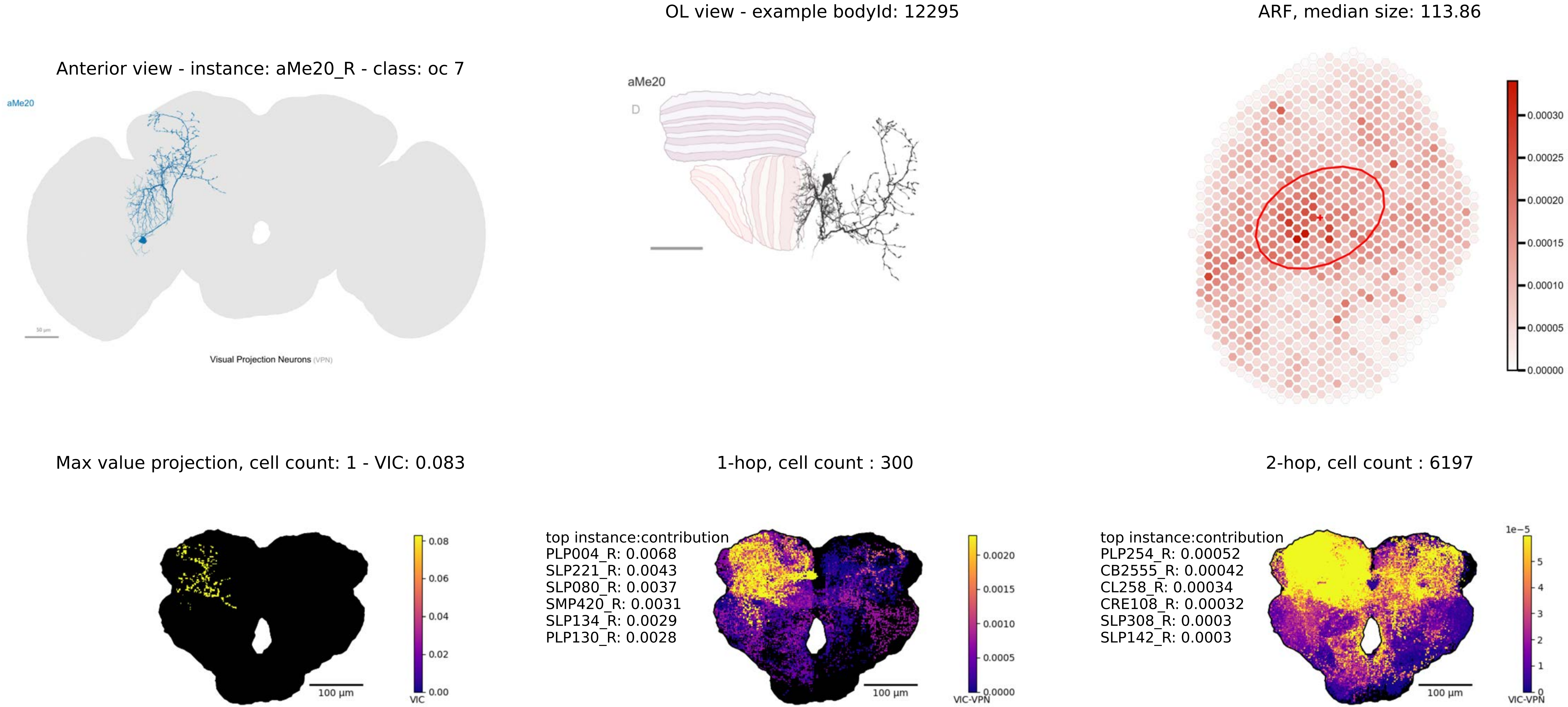
OL view - example bodyId: 12295
ARF, median size: 113.86
Anterior view - instance: aMe20_R - class: oc 7
aMe20
Visual Projection Neurons (VPN)
aMe20
D
0.00030
0.00025
0.00020
0.00015
0.00010
0.00005
0.00000
Max value projection, cell count: 1 - VIC: 0.083
1-hop, cell count : 300
2-hop, cell count : 6197
0.08
0.06
0.04
0.02
0.00
VIC
100 µm
top instance:contribution
PLP004_R: 0.0068
SLP221_R: 0.0043
SLP080_R: 0.0037
SMP420_R: 0.0031
SLP134_R: 0.0029
PLP130_R: 0.0028
0.0020
0.0015
0.0010
0.0005
0.0000
VIC-VPN
100 µm
top instance:contribution
PLP254_R: 0.00052
CB2555_R: 0.00042
CL258_R: 0.00034
CRE108_R: 0.00032
SLP308_R: 0.0003
SLP142_R: 0.0003
1e−5
5
4
3
2
1
0
VIC-VPN
100 µm

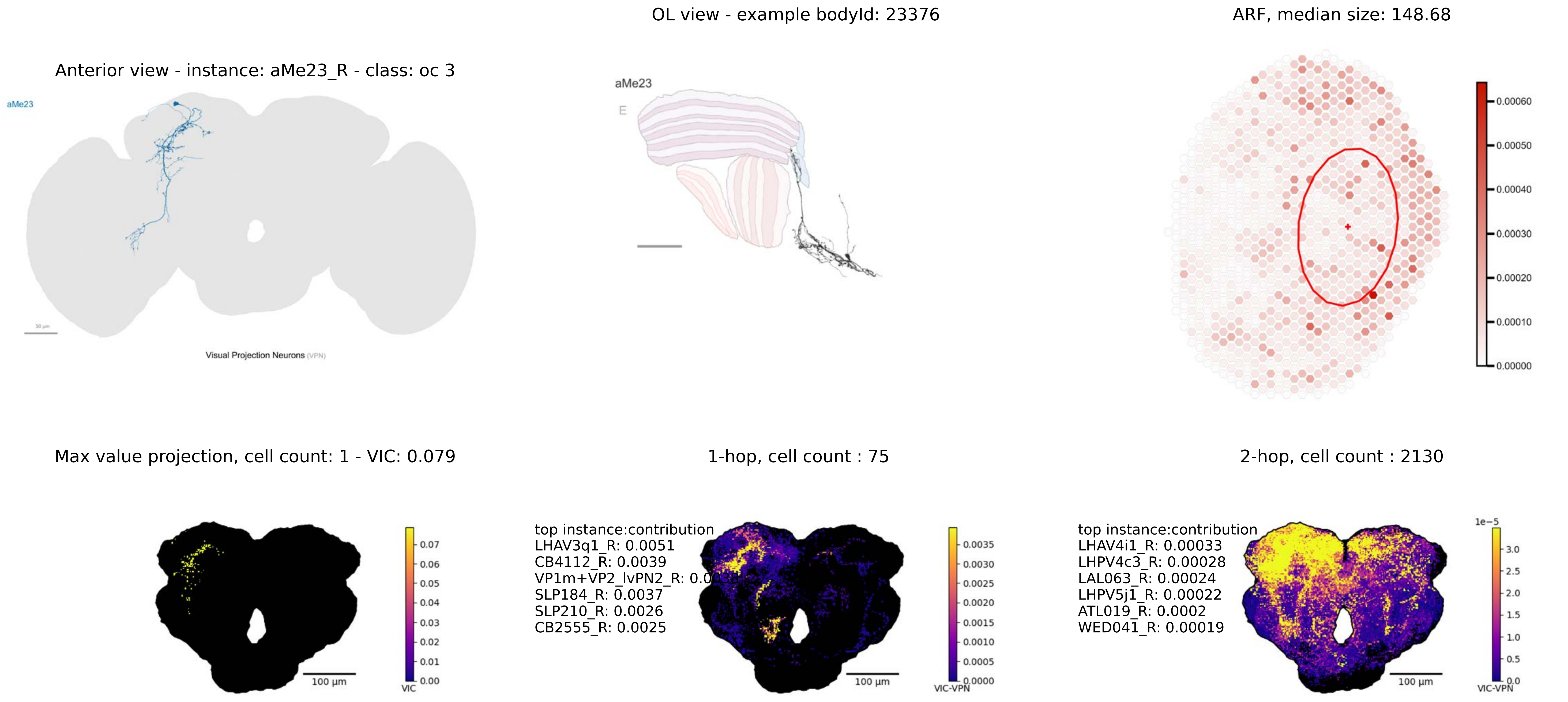

OL view - example bodyId: 23376
ARF, median size: 148.68
Anterior view - instance: aMe23_R - class: oc 3
aMe23
Visual Projection Neurons (VPN)
aMe23
E
0.00060
0.00050
0.00040
0.00030
0.00020
0.00010
0.00000
Max value projection, cell count: 1 - VIC: 0.079
1-hop, cell count : 75
2-hop, cell count : 2130
0.07
0.06
0.05
0.04
0.03
0.02
0.01
0.00
VIC
100 µm
top instance:contribution
LHAV3q1_R: 0.0051
CB4112_R: 0.0039
VP1m+VP2_lvPN2_R: 0.00
SLP184_R: 0.0037
SLP210_R: 0.0026
CB2555_R: 0.0025
0.0035
0.0030
0.0025
0.0020
0.0015
0.0010
0.0005
0.0000
VIC-VPN
100 µm
top instance:contribution
LHAV4i1_R: 0.00033
LHPV4c3_R: 0.00028
LAL063_R: 0.00024
LHPV5j1_R: 0.00022
ATL019_R: 0.0002
WED041_R: 0.00019
1e−5
3.0
2.5
2.0
1.5
1.0
0.5
0.0
VIC-VPN
100 µm

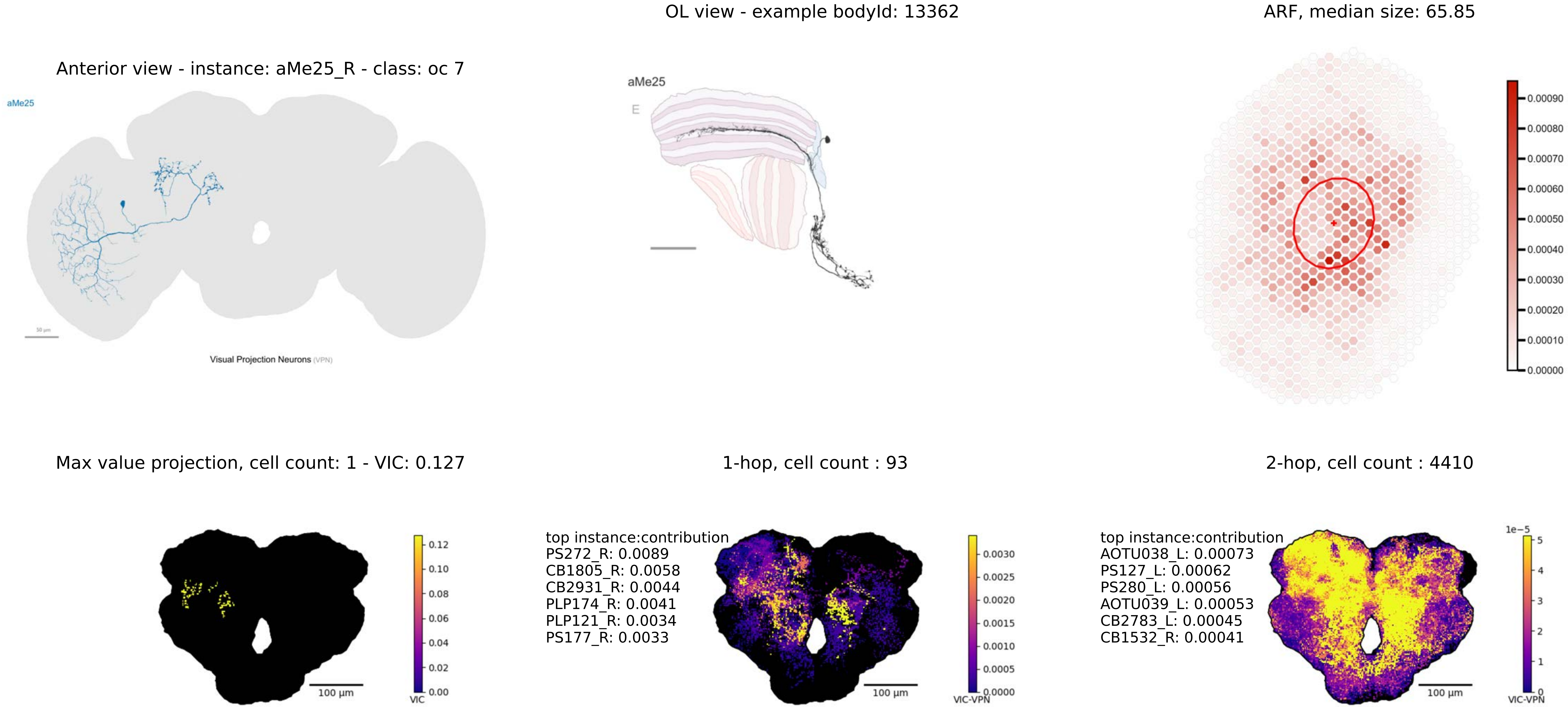

OL view - example bodyId: 13362
ARF, median size: 65.85
Anterior view - instance: aMe25_R - class: oc 7
aMe25
50 µm
Visual Projection Neurons (VPN)
aMe25
E
0.00090
0.00080
0.00070
0.00060
0.00050
0.00040
0.00030
0.00020
0.00010
0.00000
Max value projection, cell count: 1 - VIC: 0.127
1-hop, cell count : 93
2-hop, cell count : 4410
0.12
0.10
0.08
0.06
0.04
0.02
0.00
VIC
100 µm
top instance:contribution
PS272_R: 0.0089
CB1805_R: 0.0058
CB2931_R: 0.0044
PLP174_R: 0.0041
PLP121_R: 0.0034
PS177_R: 0.0033
0.0030
0.0025
0.0020
0.0015
0.0010
0.0005
0.0000
VIC-VPN
100 µm
top instance:contribution
AOTU038_L: 0.00073
PS127_L: 0.00062
PS280_L: 0.00056
AOTU039_L: 0.00053
CB2783_L: 0.00045
CB1532_R: 0.00041
1e−5
5
4
3
2
1
0
VIC-VPN
100 µm

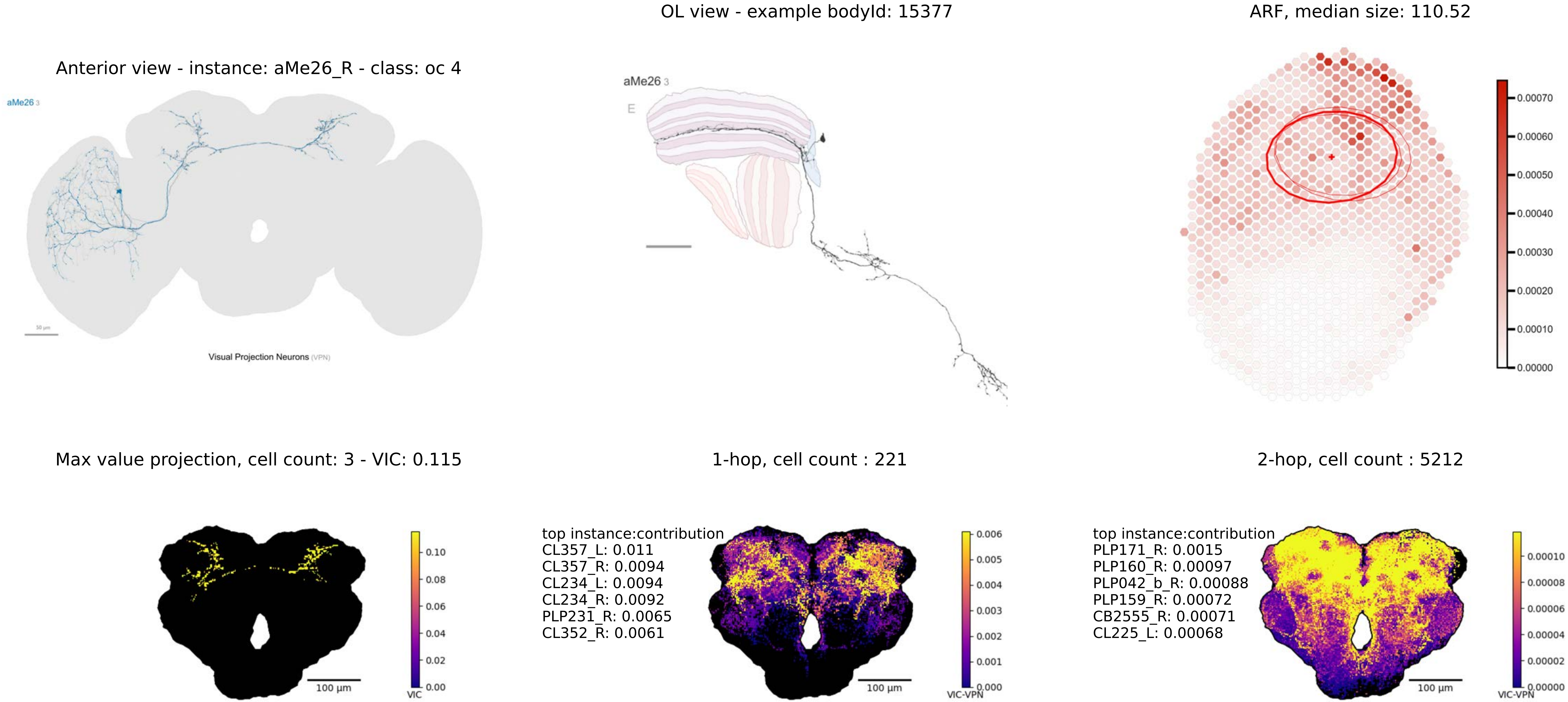

OL view - example bodyId: 15377
ARF, median size: 110.52
Anterior view - instance: aMe26_R - class: oc 4
aMe26 3
Visual Projection Neurons (VPN)
aMe26 3
E
Max value projection, cell count: 3 - VIC: 0.115
1-hop, cell count : 221
2-hop, cell count : 5212
top instance:contribution
CL357_L: 0.011
CL357_R: 0.0094
CL234_L: 0.0094
CL234_R: 0.0092
PLP231_R: 0.0065
CL352_R: 0.0061
top instance:contribution
PLP171_R: 0.0015
PLP160_R: 0.00097
PLP042_b_R: 0.00088
PLP159_R: 0.00072
CB2555_R: 0.00071
CL225_L: 0.00068
100 µm
100 µm
100 µm
VIC
VIC-VPN
VIC-VPN

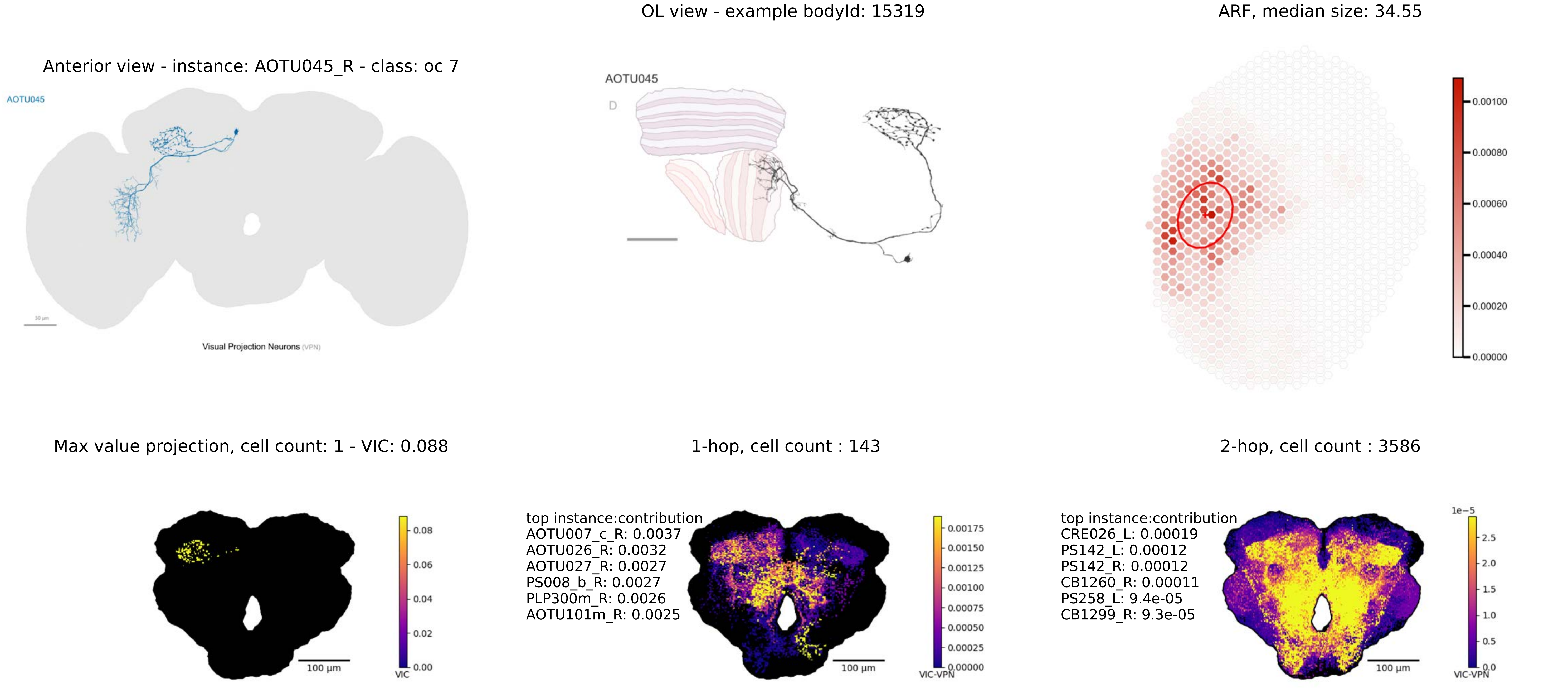
OL view - example bodyId: 15319
ARF, median size: 34.55
Anterior view - instance: AOTU045_R - class: oc 7
AOTU045
Visual Projection Neurons (VPN)
AOTU045
D
0.00100
0.00080
0.00060
0.00040
0.00020
0.00000
Max value projection, cell count: 1 - VIC: 0.088
1-hop, cell count : 143
2-hop, cell count : 3586
0.08
0.06
0.04
0.02
0.00
VIC
100 µm
top instance:contribution
AOTU007_c_R: 0.0037
AOTU026_R: 0.0032
AOTU027_R: 0.0027
PS008_b_R: 0.0027
PLP300m_R: 0.0026
AOTU101m_R: 0.0025
0.00175
0.00150
0.00125
0.00100
0.00075
0.00050
0.00025
0.00000
VIC-VPN
100 µm
top instance:contribution
CRE026_L: 0.00019
PS142_L: 0.00012
PS142_R: 0.00012
CB1260_R: 0.00011
PS258_L: 9.4e-05
CB1299_R: 9.3e-05
1e−5
2.5
2.0
1.5
1.0
0.5
0.0
VIC-VPN
100 µm

Anterior view - instance: dCal1_R - class: oc 9

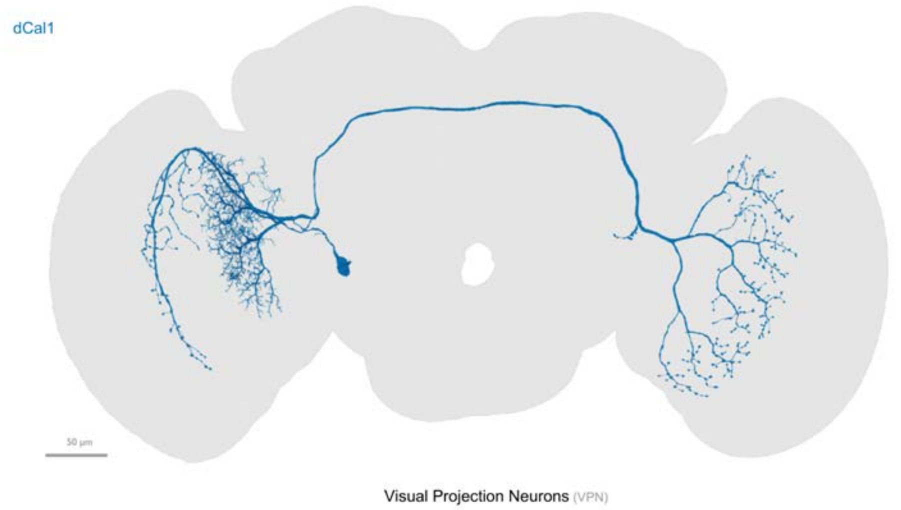


fewer than 100 output connections in central brain

OL view - example bodyId: 10441

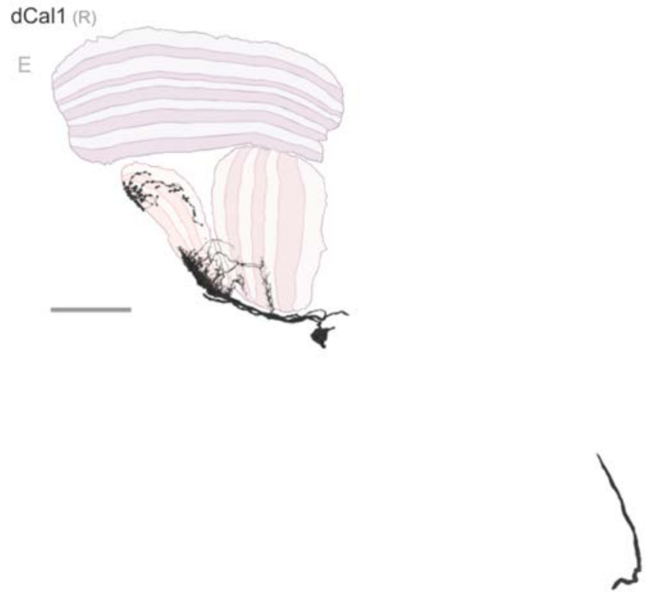


ARF, median size: 28.62

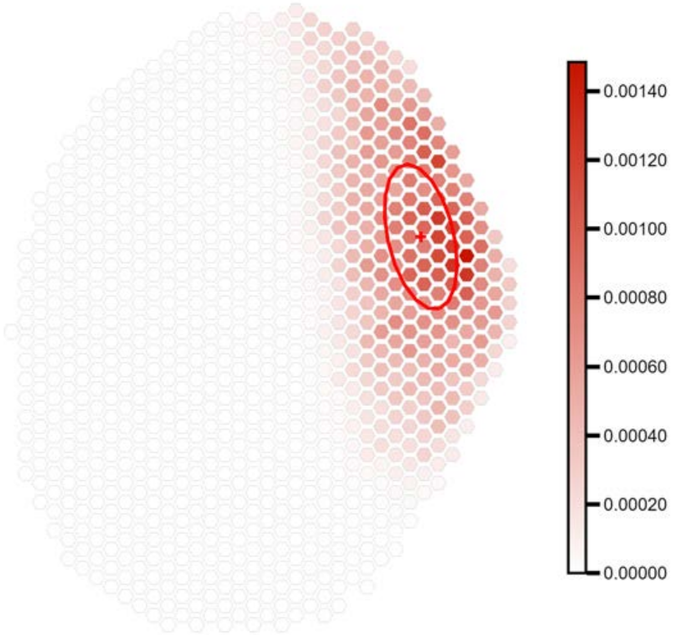

Anterior view - instance: H1_R - class: oc 9

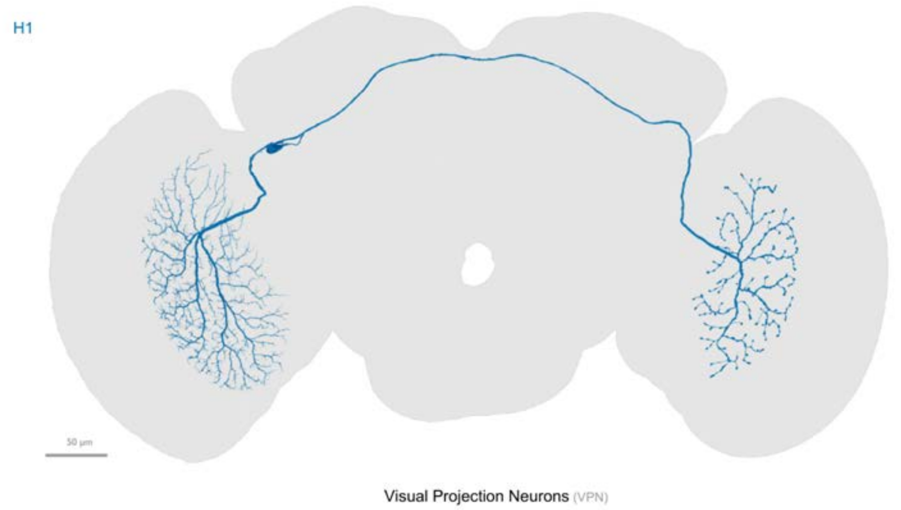


fewer than 100 output connections in central brain

OL view - example bodyId: 10755

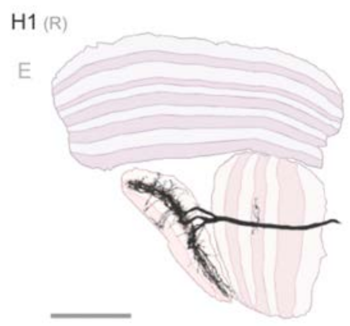


ARF, median size: 98.72

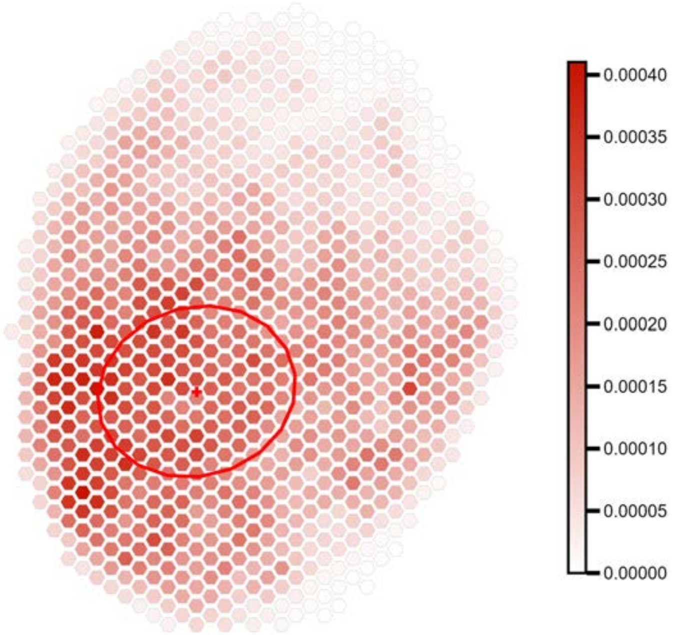

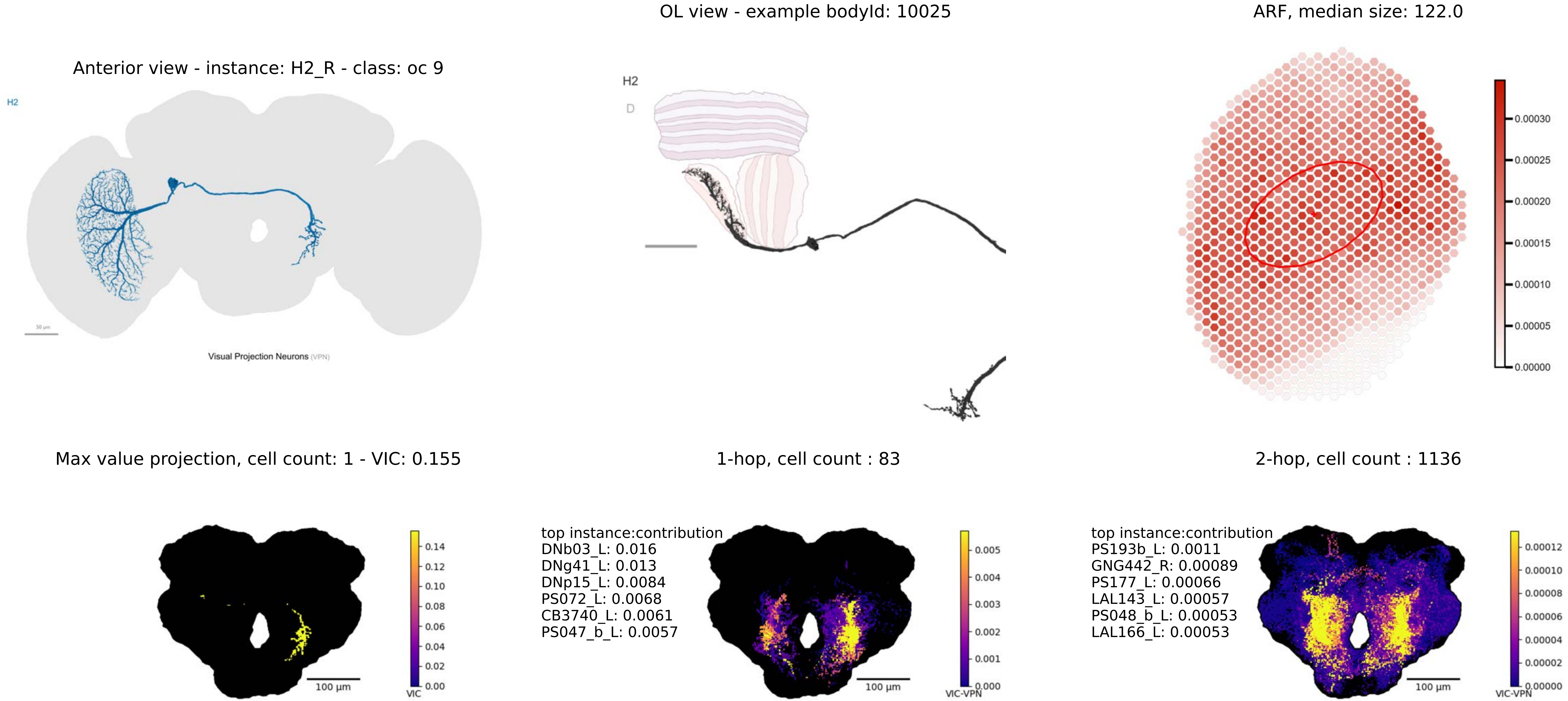

OL view - example bodyId: 10025
ARF, median size: 122.0
Anterior view - instance: H2_R - class: oc 9
H2
Visual Projection Neurons (VPN)
H2
D
0.00030
0.00025
0.00020
0.00015
0.00010
0.00005
0.00000
Max value projection, cell count: 1 - VIC: 0.155
1-hop, cell count : 83
2-hop, cell count : 1136
0.14
0.12
0.10
0.08
0.06
0.04
0.02
0.00
VIC
100 µm
top instance:contribution
DNb03_L: 0.016
DNg41_L: 0.013
DNp15_L: 0.0084
PS072_L: 0.0068
CB3740_L: 0.0061
PS047_b_L: 0.0057
0.005
0.004
0.003
0.002
0.001
0.000
VIC-VPN
100 µm
top instance:contribution
PS193b_L: 0.0011
GNG442_R: 0.00089
PS177_L: 0.00066
LAL143_L: 0.00057
PS048_b_L: 0.00053
LAL166_L: 0.00053
0.00012
0.00010
0.00008
0.00006
0.00004
0.00002
0.00000
VIC-VPN
100 µm

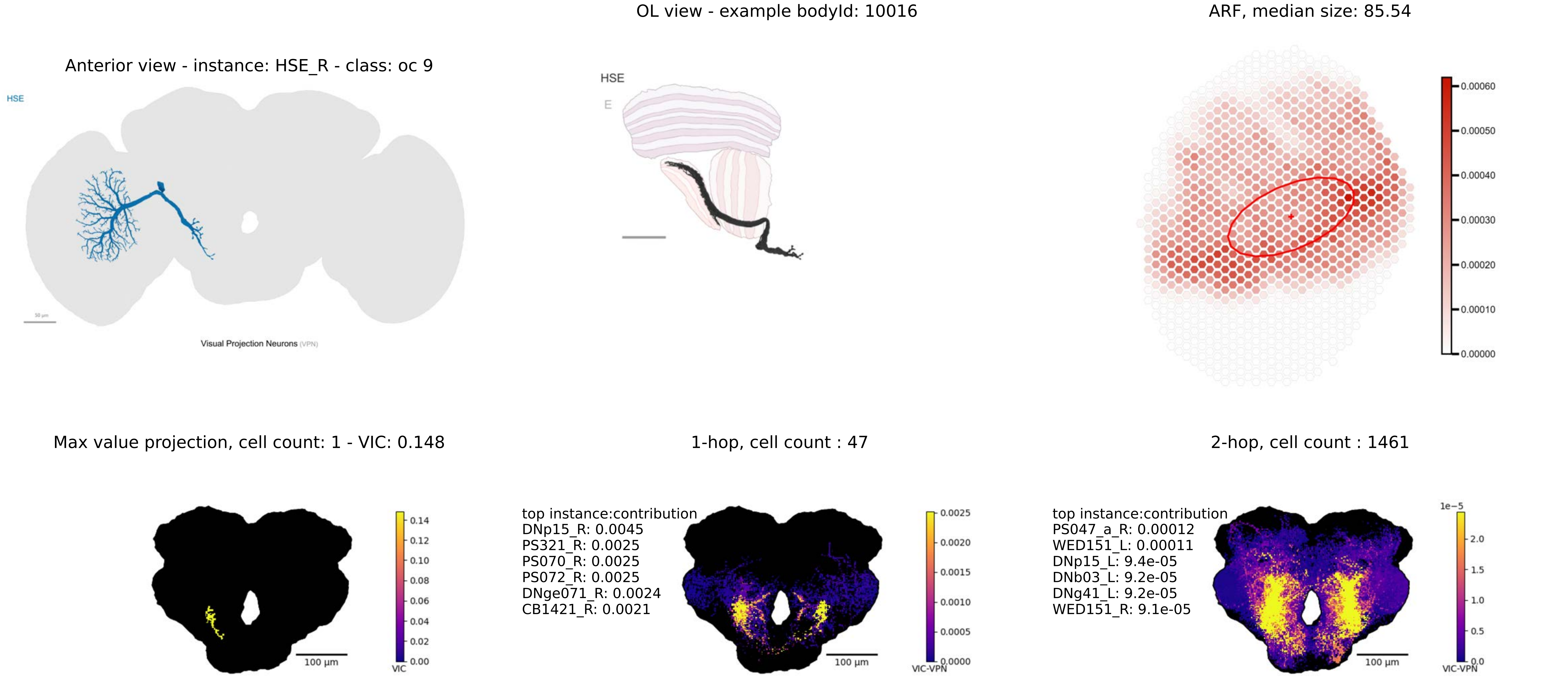

OL view - example bodyId: 10016
ARF, median size: 85.54
Anterior view - instance: HSE_R - class: oc 9
HSE
Visual Projection Neurons (VPN)
HSE
E
0.00060
0.00050
0.00040
0.00030
0.00020
0.00010
0.00000
Max value projection, cell count: 1 - VIC: 0.148
1-hop, cell count : 47
2-hop, cell count : 1461
0.14
0.12
0.10
0.08
0.06
0.04
0.02
0.00
VIC
100 µm
top instance:contribution
DNp15_R: 0.0045
PS321_R: 0.0025
PS070_R: 0.0025
PS072_R: 0.0025
DNge071_R: 0.0024
CB1421_R: 0.0021
0.0025
0.0020
0.0015
0.0010
0.0005
0.0000
VIC-VPN
top instance:contribution
PS047_a_R: 0.00012
WED151_L: 0.00011
DNp15_L: 9.4e-05
DNb03_L: 9.2e-05
DNg41_L: 9.2e-05
WED151_R: 9.1e-05
1e−5
2.0
1.5
1.0
0.5
0.0
VIC-VPN

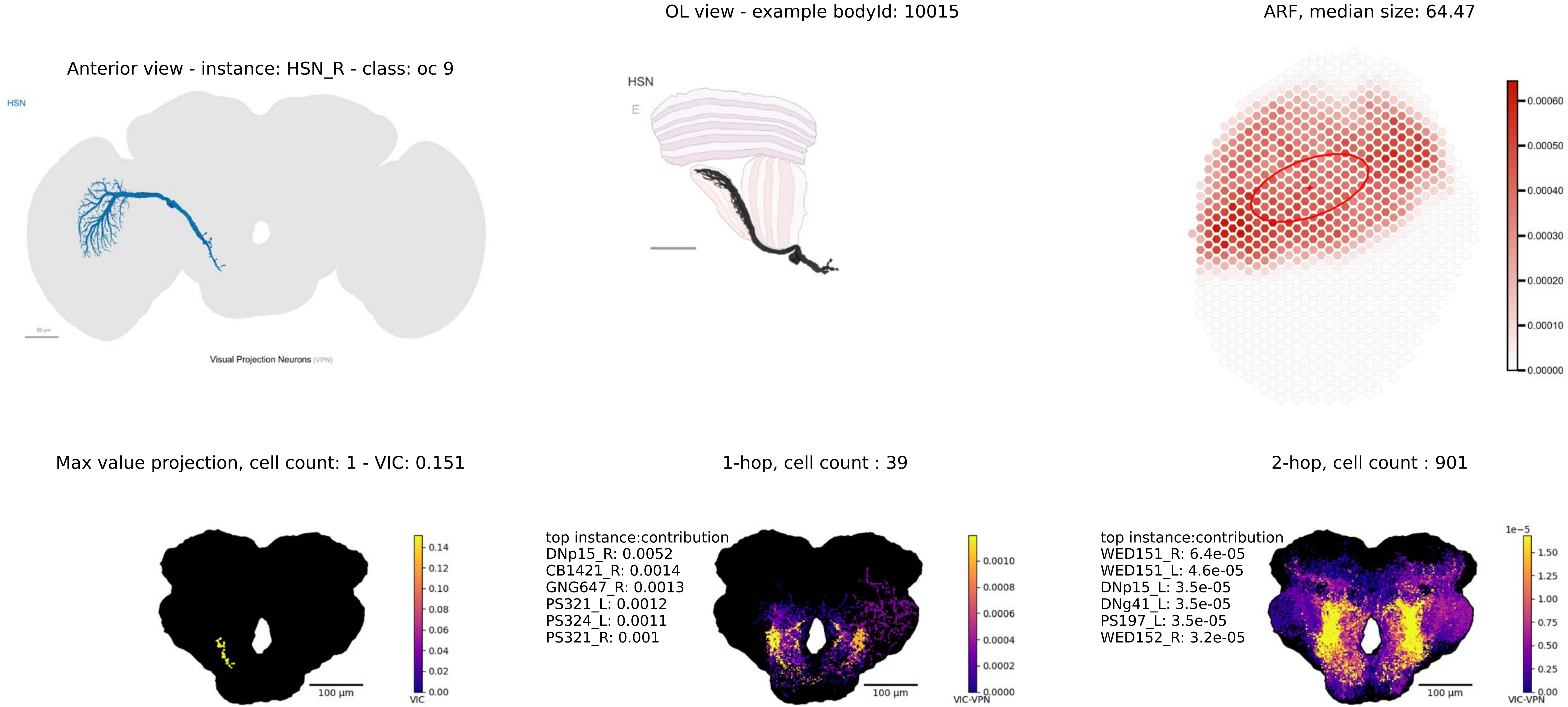
OL view - example bodyId: 10015
ARF, median size: 64.47
Anterior view - instance: HSN_R - class: oc 9
HSN
Visual Projection Neurons (VPN)
HSN
E
0.00060
0.00050
0.00040
0.00030
0.00020
0.00010
0.00000
Max value projection, cell count: 1 - VIC: 0.151
1-hop, cell count : 39
2-hop, cell count : 901
0.14
0.12
0.10
0.08
0.06
0.04
0.02
0.00
VIC
100 µm
top instance:contribution
DNp15_R: 0.0052
CB1421_R: 0.0014
GNG647_R: 0.0013
PS321_L: 0.0012
PS324_L: 0.0011
PS321_R: 0.001
0.0010
0.0008
0.0006
0.0004
0.0002
0.0000
VIC-VPN
100 µm
top instance:contribution
WED151_R: 6.4e-05
WED151_L: 4.6e-05
DNp15_L: 3.5e-05
DNg41_L: 3.5e-05
PS197_L: 3.5e-05
WED152_R: 3.2e-05
1e−5
1.50
1.25
1.00
0.75
0.50
0.25
0.00
VIC-VPN
100 µm

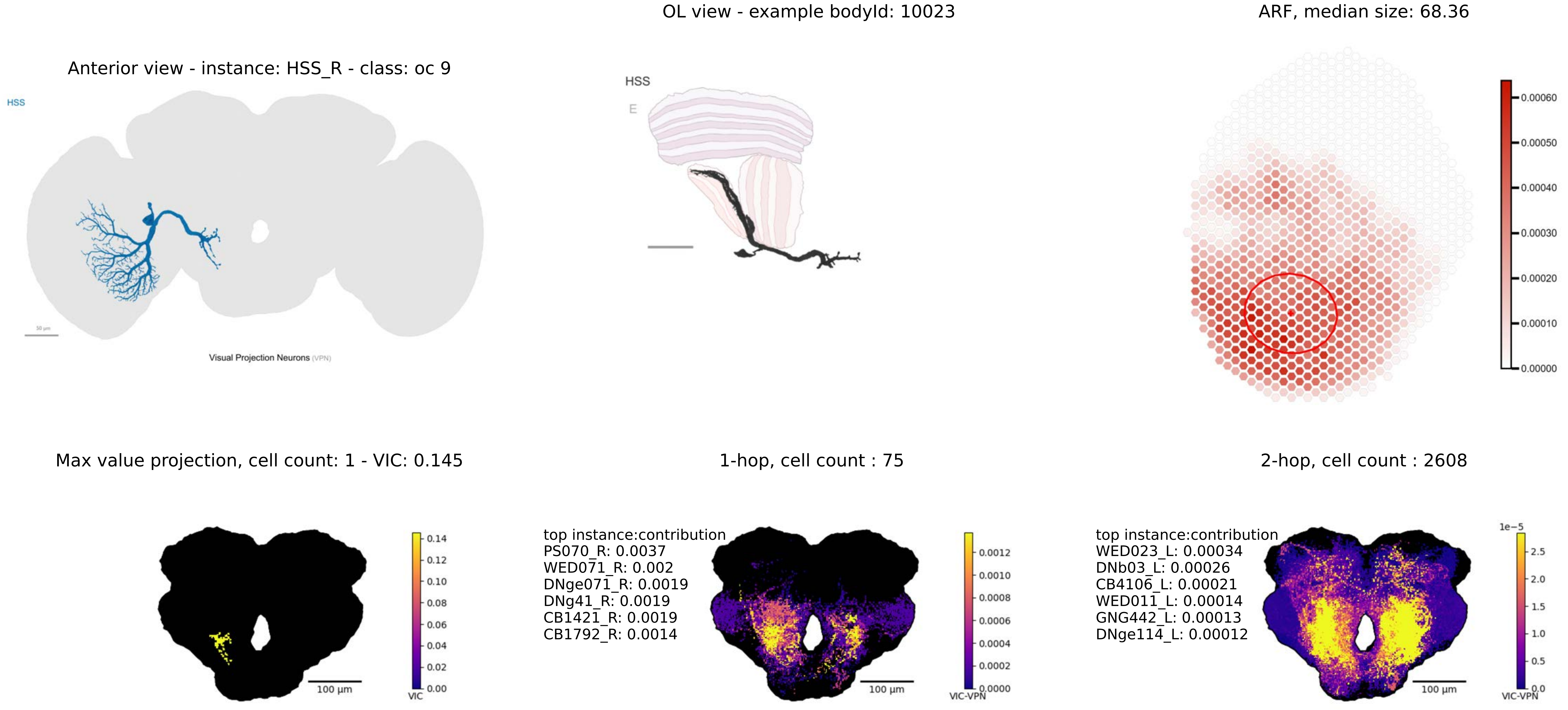
OL view - example bodyId: 10023
ARF, median size: 68.36
Anterior view - instance: HSS_R - class: oc 9
HSS
50 μm
Visual Projection Neurons (VPN)
HSS
E
0.00060
0.00050
0.00040
0.00030
0.00020
0.00010
0.00000
Max value projection, cell count: 1 - VIC: 0.145
1-hop, cell count : 75
2-hop, cell count : 2608
100 μm
0.14
0.12
0.10
0.08
0.06
0.04
0.02
0.00
VIC
top instance:contribution
PS070_R: 0.0037
WED071_R: 0.002
DNge071_R: 0.0019
DNg41_R: 0.0019
CB1421_R: 0.0019
CB1792_R: 0.0014
0.0012
0.0010
0.0008
0.0006
0.0004
0.0002
0.0000
VIC-VPN
top instance:contribution
WED023_L: 0.00034
DNb03_L: 0.00026
CB4106_L: 0.00021
WED011_L: 0.00014
GNG442_L: 0.00013
DNge114_L: 0.00012
1e−5
2.5
2.0
1.5
1.0
0.5
0.0
VIC-VPN

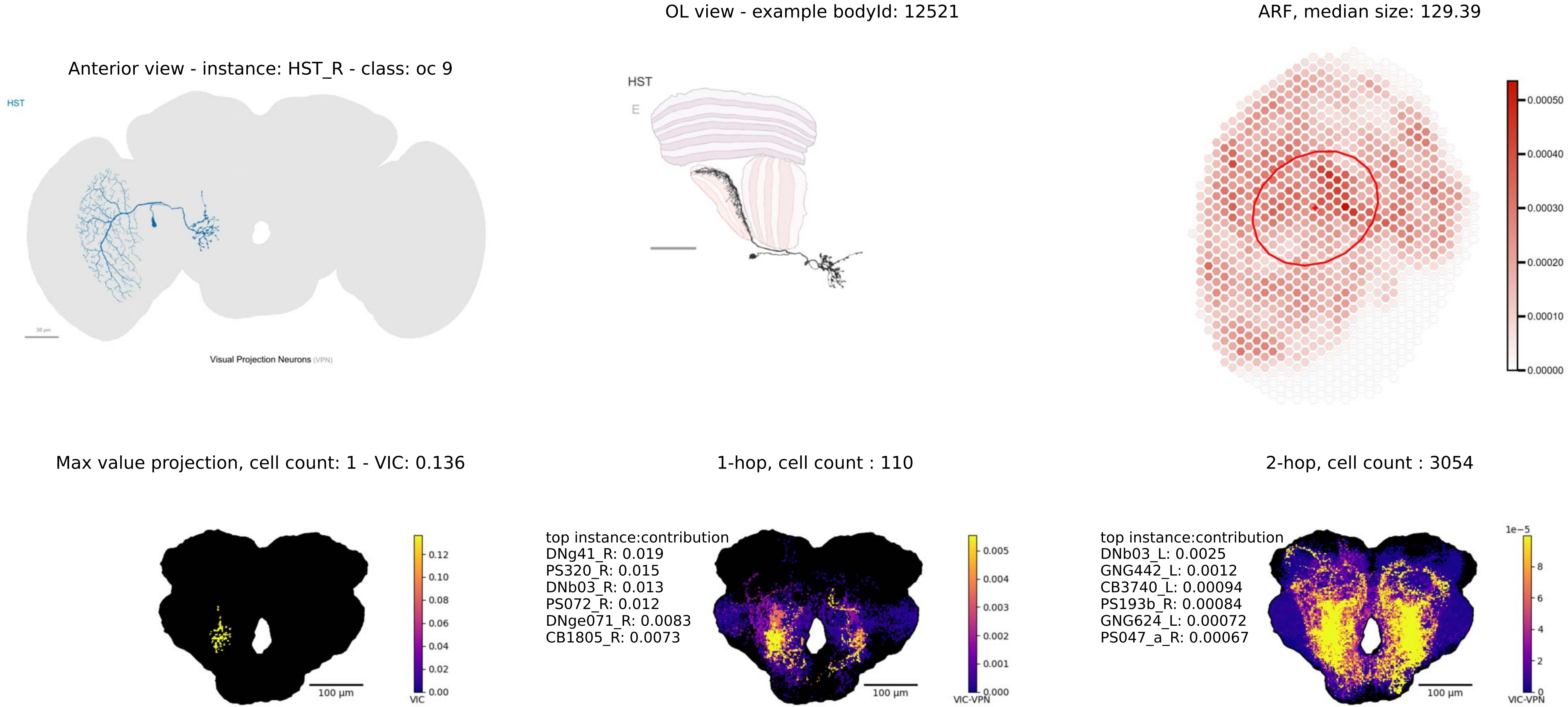
OL view - example bodyId: 12521
ARF, median size: 129.39
Anterior view - instance: HST_R - class: oc 9
HST
Visual Projection Neurons (VPN)
HST
E
0.00050
0.00040
0.00030
0.00020
0.00010
0.00000
Max value projection, cell count: 1 - VIC: 0.136
1-hop, cell count : 110
2-hop, cell count : 3054
0.12
0.10
0.08
0.06
0.04
0.02
0.00
VIC
100 µm
top instance:contribution
DNg41_R: 0.019
PS320_R: 0.015
DNb03_R: 0.013
PS072_R: 0.012
DNge071_R: 0.0083
CB1805_R: 0.0073
0.005
0.004
0.003
0.002
0.001
0.000
VIC-VPN
100 µm
top instance:contribution
DNb03_L: 0.0025
GNG442_L: 0.0012
CB3740_L: 0.00094
PS193b_R: 0.00084
GNG624_L: 0.00072
PS047_a_R: 0.00067
1e−5
8
6
4
2
0
VIC-VPN
100 µm

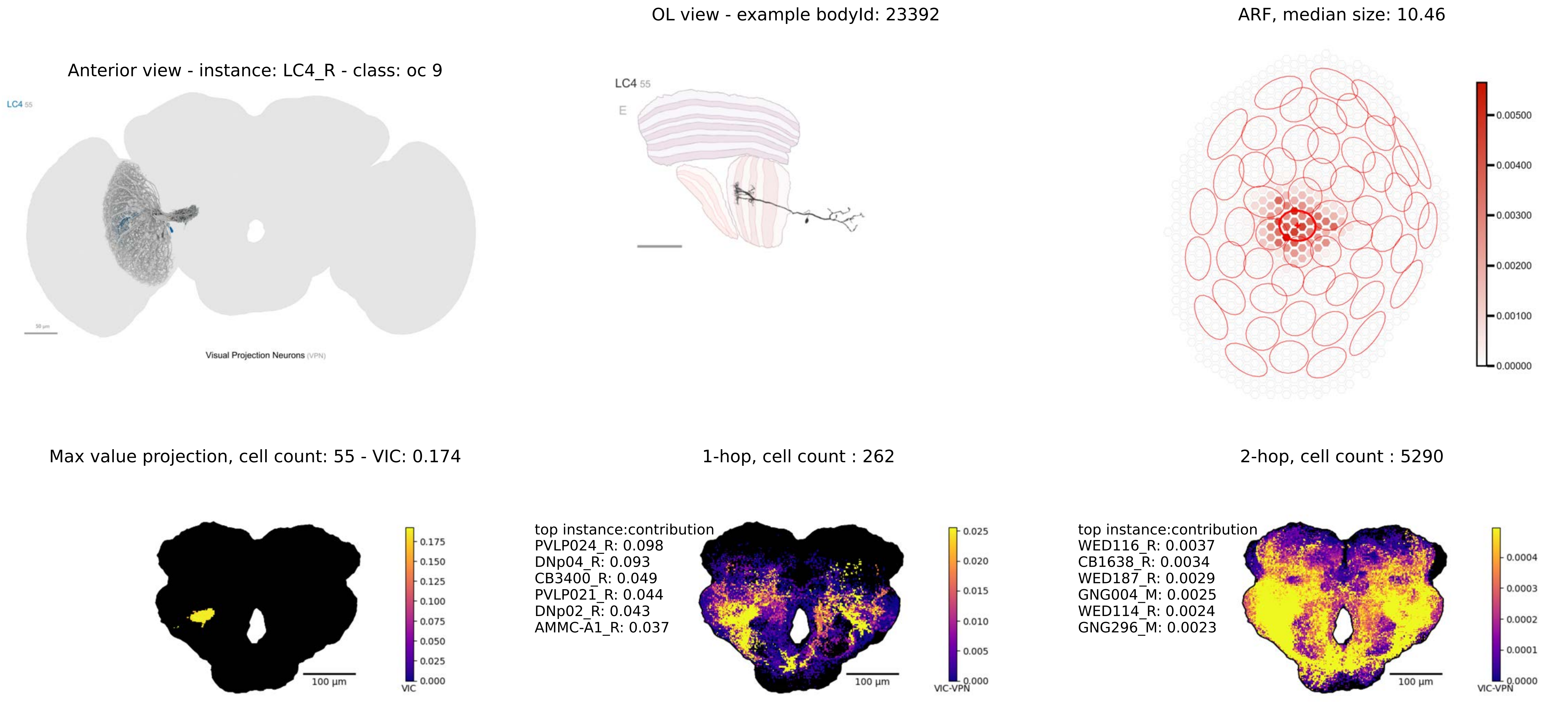

OL view - example bodyId: 23392
ARF, median size: 10.46
Anterior view - instance: LC4_R - class: oc 9
LC4 55
Visual Projection Neurons (VPN)
LC4 55
E
0.00500
0.00400
0.00300
0.00200
0.00100
0.00000
Max value projection, cell count: 55 - VIC: 0.174
1-hop, cell count : 262
2-hop, cell count : 5290
0.175
0.150
0.125
0.100
0.075
0.050
0.025
0.000
VIC
100 µm
top instance:contribution
PVLP024_R: 0.098
DNp04_R: 0.093
CB3400_R: 0.049
PVLP021_R: 0.044
DNp02_R: 0.043
AMMC-A1_R: 0.037
0.025
0.020
0.015
0.010
0.005
0.000
VIC-VPN
100 µm
top instance:contribution
WED116_R: 0.0037
CB1638_R: 0.0034
WED187_R: 0.0029
GNG004_M: 0.0025
WED114_R: 0.0024
GNG296_M: 0.0023
0.0004
0.0003
0.0002
0.0001
0.0000
VIC-VPN
100 µm

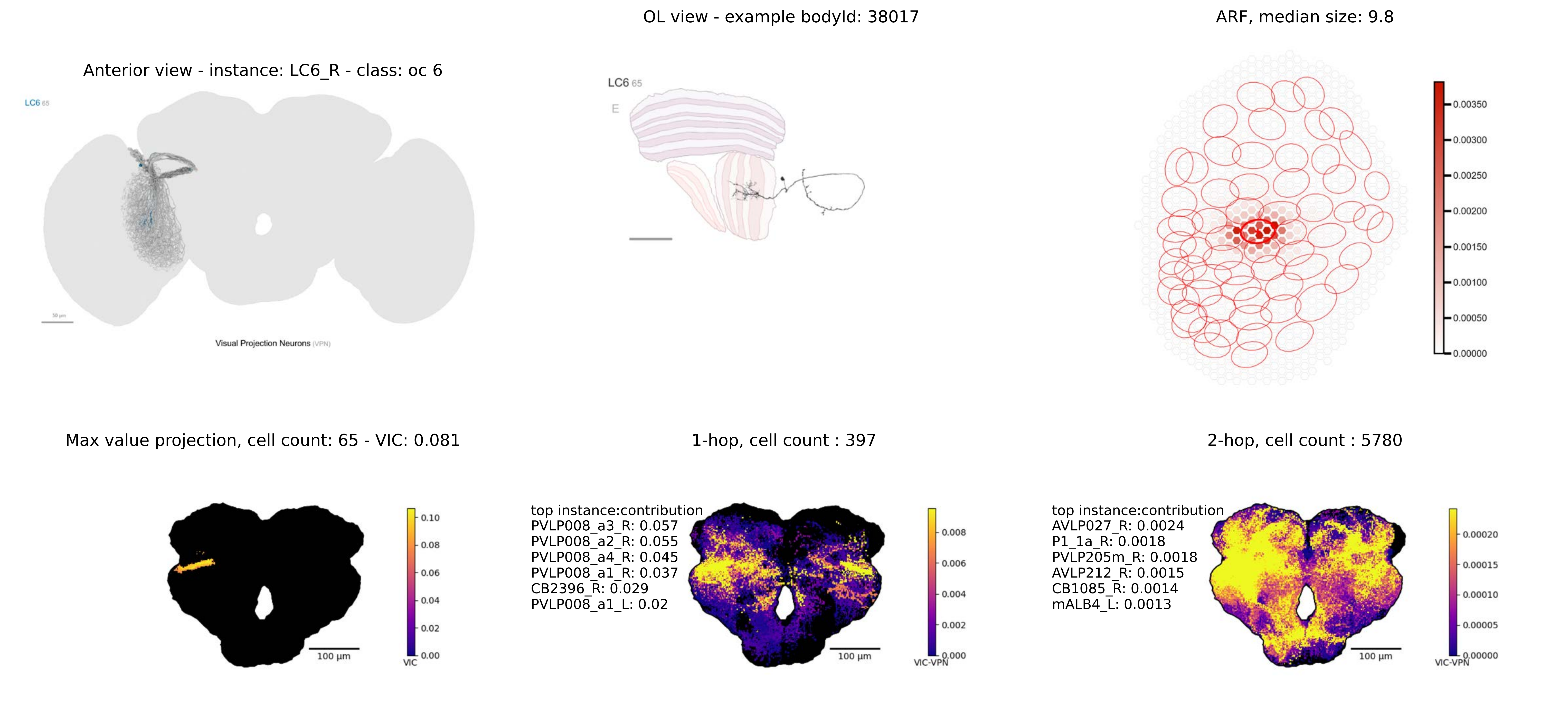
OL view - example bodyId: 38017
ARF, median size: 9.8
Anterior view - instance: LC6_R - class: oc 6
LC6 65
E
Visual Projection Neurons (VPN)
0.00350
0.00300
0.00250
0.00200
0.00150
0.00100
0.00050
0.00000
Max value projection, cell count: 65 - VIC: 0.081
1-hop, cell count : 397
2-hop, cell count : 5780
0.10
0.08
0.06
0.04
0.02
0.00
VIC
100 µm
top instance:contribution
PVLP008_a3_R: 0.057
PVLP008_a2_R: 0.055
PVLP008_a4_R: 0.045
PVLP008_a1_R: 0.037
CB2396_R: 0.029
PVLP008_a1_L: 0.02
0.008
0.006
0.004
0.002
0.000
VIC-VPN
top instance:contribution
AVLP027_R: 0.0024
P1_1a_R: 0.0018
PVLP205m_R: 0.0018
AVLP212_R: 0.0015
CB1085_R: 0.0014
mALB4_L: 0.0013
0.00020
0.00015
0.00010
0.00005
0.00000
VIC-VPN

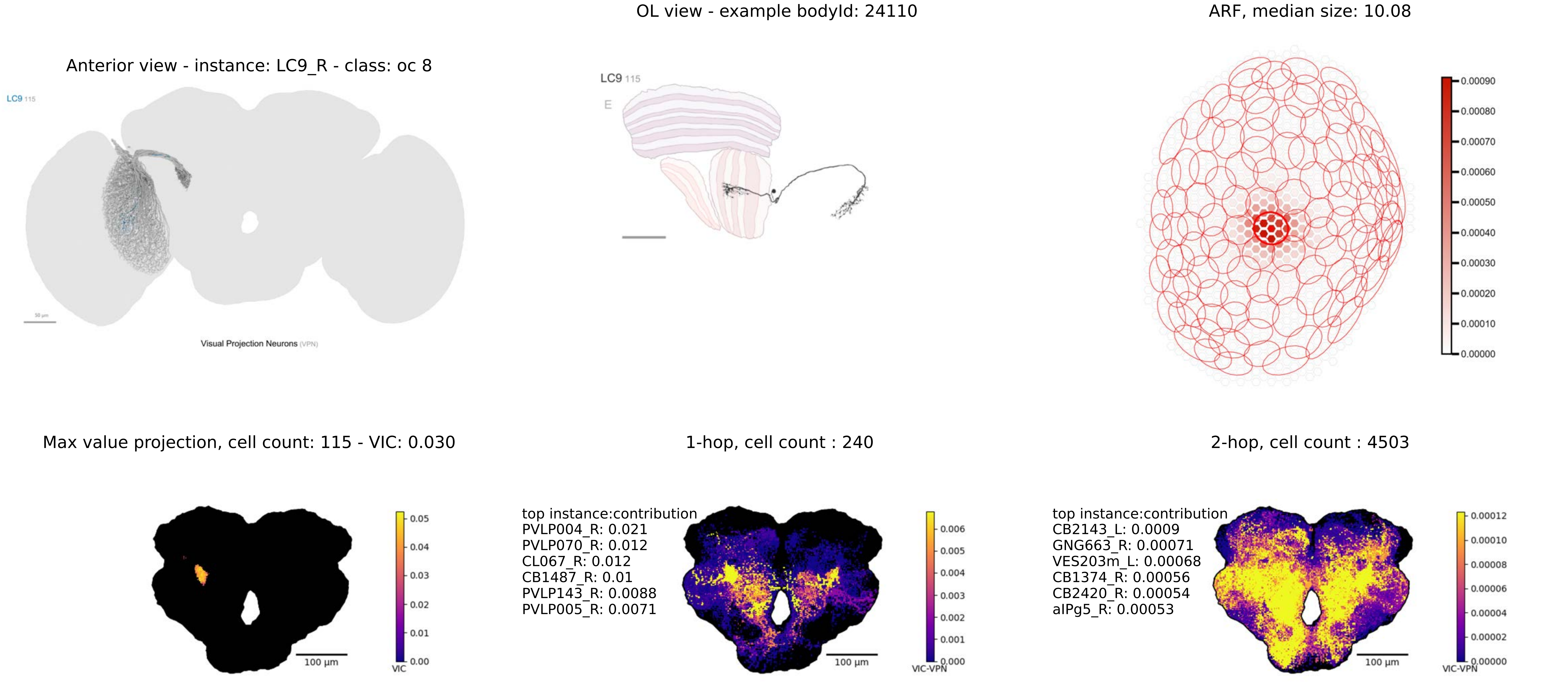
Anterior view - instance: LC9_R - class: oc 8
LC9 115
Visual Projection Neurons (VPN)
OL view - example bodyId: 24110
LC9 115
E
ARF, median size: 10.08
0.00090
0.00080
0.00070
0.00060
0.00050
0.00040
0.00030
0.00020
0.00010
0.00000
Max value projection, cell count: 115 - VIC: 0.030
0.05
0.04
0.03
0.02
0.01
0.00
VIC
100 µm
1-hop, cell count : 240
top instance:contribution
PVLP004_R: 0.021
PVLP070_R: 0.012
CL067_R: 0.012
CB1487_R: 0.01
PVLP143_R: 0.0088
PVLP005_R: 0.0071
0.006
0.005
0.004
0.003
0.002
0.001
0.000
VIC-VPN
100 µm
2-hop, cell count : 4503
top instance:contribution
CB2143_L: 0.0009
GNG663_R: 0.00071
VES203m_L: 0.00068
CB1374_R: 0.00056
CB2420_R: 0.00054
aIPg5_R: 0.00053
0.00012
0.00010
0.00008
0.00006
0.00004
0.00002
0.00000
VIC-VPN
100 µm

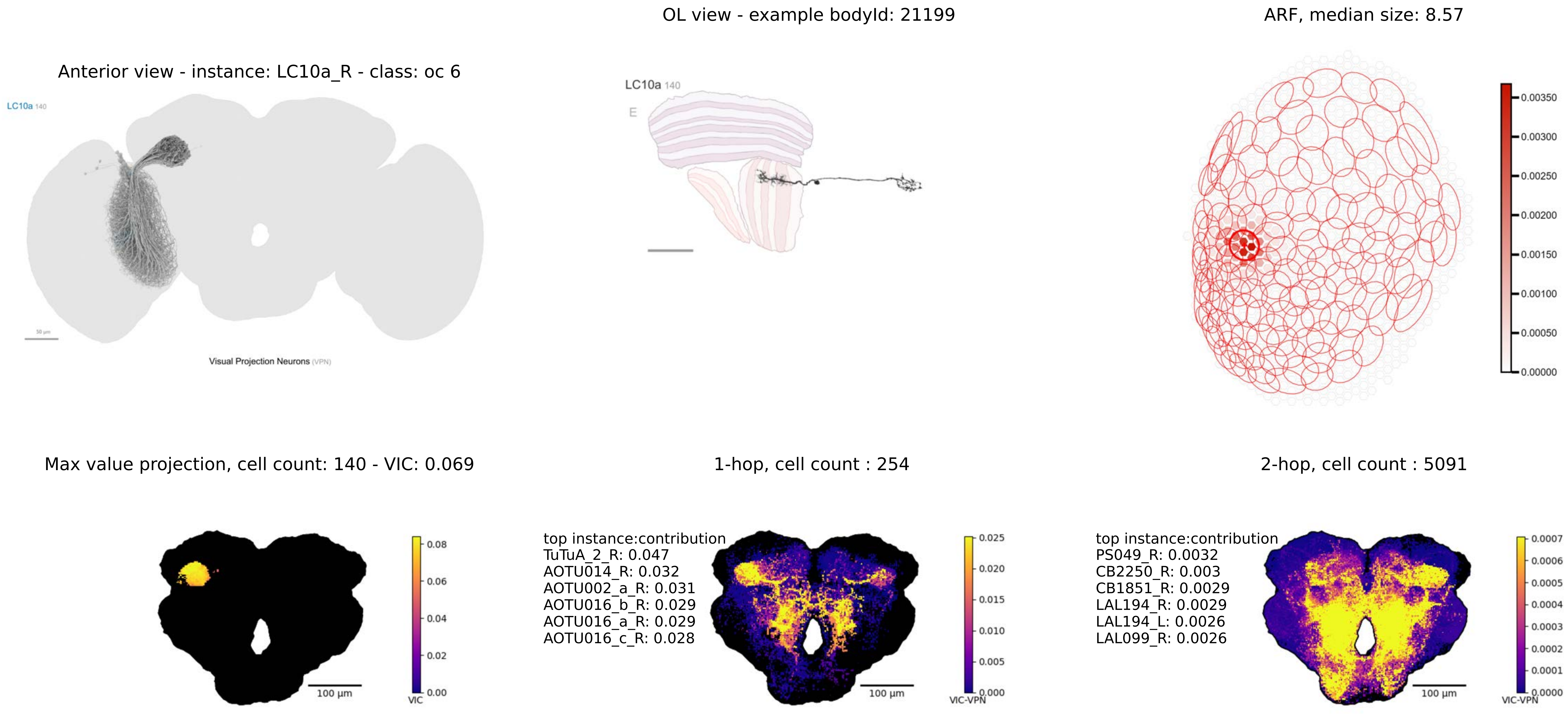

OL view - example bodyId: 21199
ARF, median size: 8.57
Anterior view - instance: LC10a_R - class: oc 6
LC10a 140
Visual Projection Neurons (VPN)
LC10a 140
E
0.00350
0.00300
0.00250
0.00200
0.00150
0.00100
0.00050
0.00000
Max value projection, cell count: 140 - VIC: 0.069
1-hop, cell count : 254
2-hop, cell count : 5091
100 µm
VIC
0.08
0.06
0.04
0.02
0.00
top instance:contribution
TuTuA_2_R: 0.047
AOTU014_R: 0.032
AOTU002_a_R: 0.031
AOTU016_b_R: 0.029
AOTU016_a_R: 0.029
AOTU016_c_R: 0.028
100 µm
VIC-VPN
0.025
0.020
0.015
0.010
0.005
0.000
top instance:contribution
PS049_R: 0.0032
CB2250_R: 0.003
CB1851_R: 0.0029
LAL194_R: 0.0029
LAL194_L: 0.0026
LAL099_R: 0.0026
100 µm
VIC-VPN
0.0007
0.0006
0.0005
0.0004
0.0003
0.0002
0.0001
0.0000

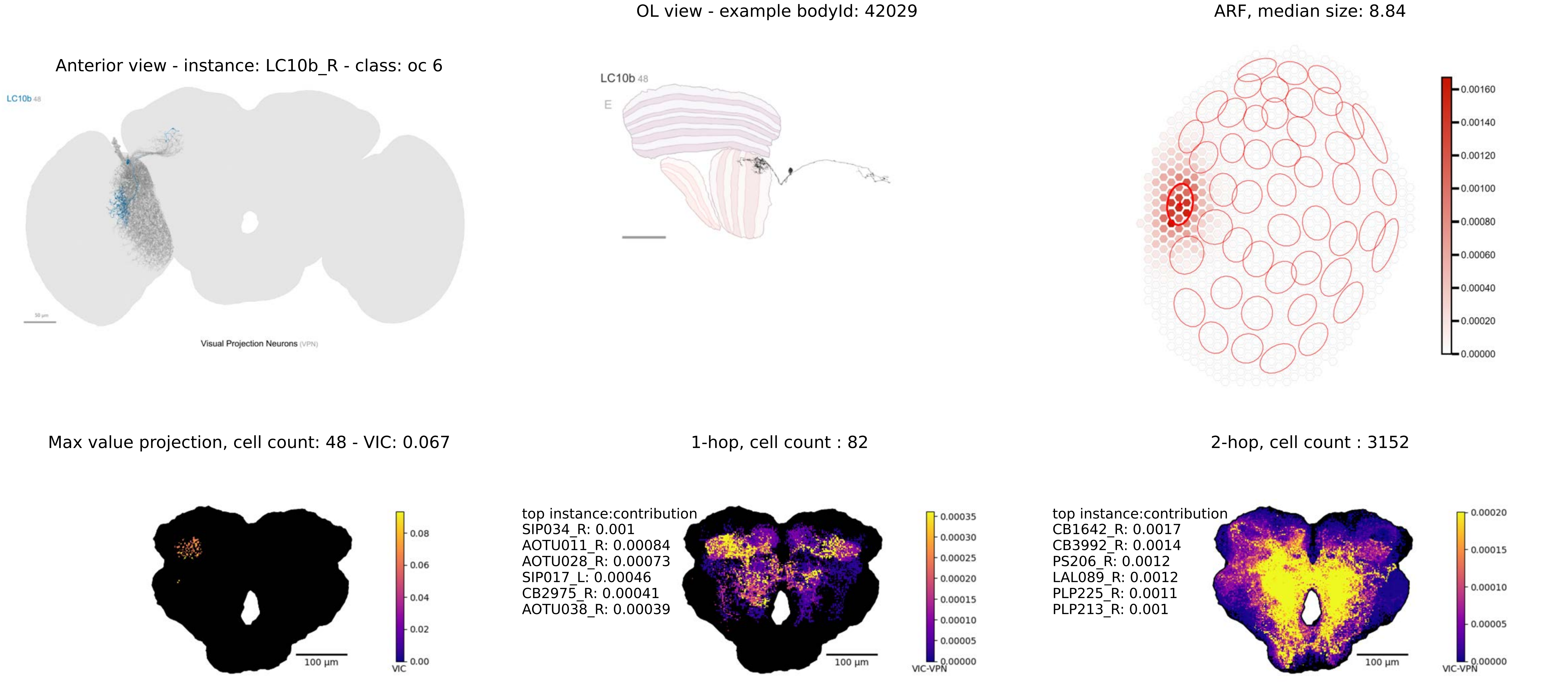

OL view - example bodyId: 42029
ARF, median size: 8.84
Anterior view - instance: LC10b_R - class: oc 6
LC10b 48
Visual Projection Neurons (VPN)
LC10b 48
E
0.00160
0.00140
0.00120
0.00100
0.00080
0.00060
0.00040
0.00020
0.00000
Max value projection, cell count: 48 - VIC: 0.067
1-hop, cell count : 82
2-hop, cell count : 3152
100 µm
VIC
0.08
0.06
0.04
0.02
0.00
top instance:contribution
SIP034_R: 0.001
AOTU011_R: 0.00084
AOTU028_R: 0.00073
SIP017_L: 0.00046
CB2975_R: 0.00041
AOTU038_R: 0.00039
100 µm
VIC-VPN
0.00035
0.00030
0.00025
0.00020
0.00015
0.00010
0.00005
0.00000
top instance:contribution
CB1642_R: 0.0017
CB3992_R: 0.0014
PS206_R: 0.0012
LAL089_R: 0.0012
PLP225_R: 0.0011
PLP213_R: 0.001
100 µm
VIC-VPN
0.00020
0.00015
0.00010
0.00005
0.00000

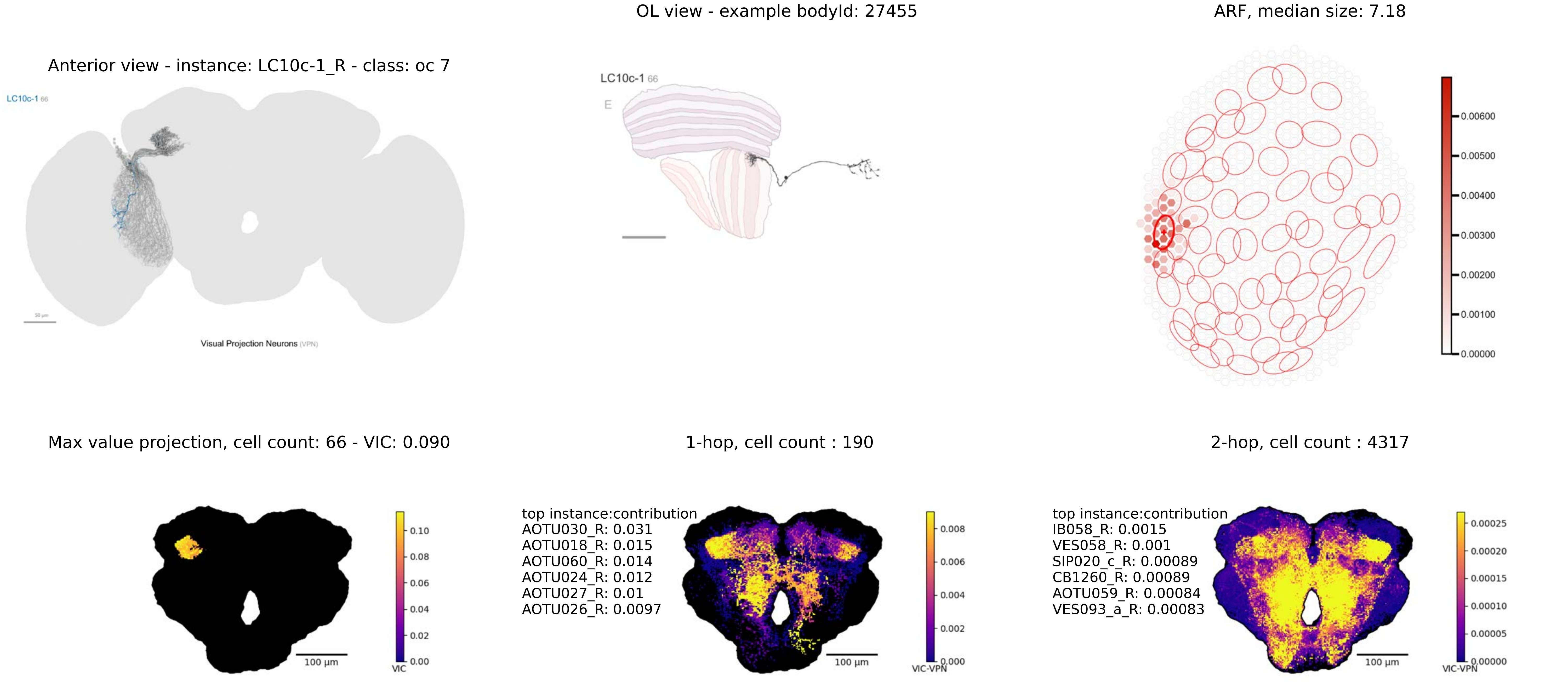
OL view - example bodyId: 27455
ARF, median size: 7.18
Anterior view - instance: LC10c-1_R - class: oc 7
LC10c-1 66
Visual Projection Neurons (VPN)
LC10c-1 66
E
0.00600
0.00500
0.00400
0.00300
0.00200
0.00100
0.00000
Max value projection, cell count: 66 - VIC: 0.090
1-hop, cell count : 190
2-hop, cell count : 4317
0.10
0.08
0.06
0.04
0.02
0.00
VIC
100 µm
top instance:contribution
AOTU030_R: 0.031
AOTU018_R: 0.015
AOTU060_R: 0.014
AOTU024_R: 0.012
AOTU027_R: 0.01
AOTU026_R: 0.0097
0.008
0.006
0.004
0.002
0.000
VIC-VPN
100 µm
top instance:contribution
IB058_R: 0.0015
VES058_R: 0.001
SIP020_c_R: 0.00089
CB1260_R: 0.00089
AOTU059_R: 0.00084
VES093_a_R: 0.00083
0.00025
0.00020
0.00015
0.00010
0.00005
0.00000
VIC-VPN
100 µm

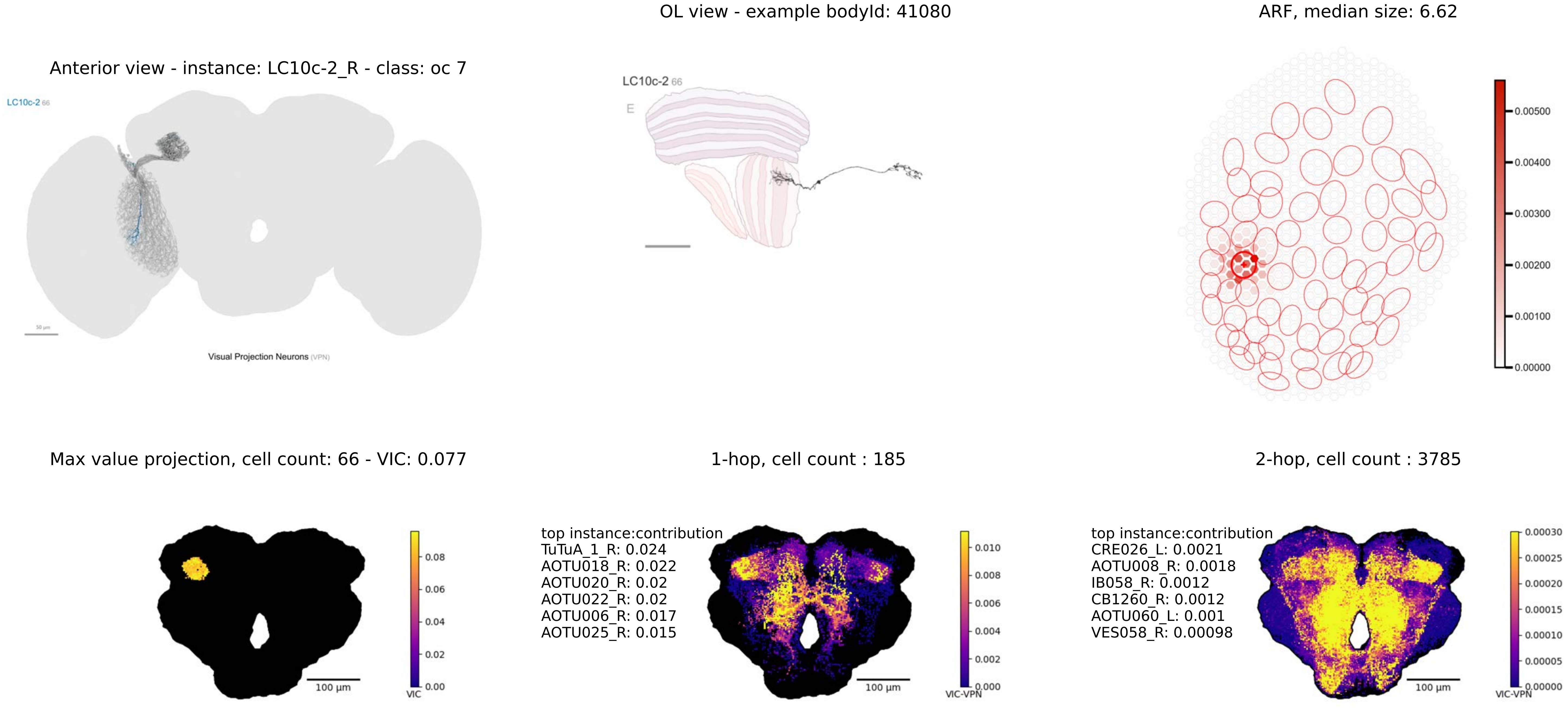
OL view - example bodyId: 41080
ARF, median size: 6.62
Anterior view - instance: LC10c-2_R - class: oc 7
LC10c-2 66
LC10c-2 66
E
Visual Projection Neurons (VPN)
0.00500
0.00400
0.00300
0.00200
0.00100
0.00000
Max value projection, cell count: 66 - VIC: 0.077
1-hop, cell count : 185
2-hop, cell count : 3785
0.08
0.06
0.04
0.02
0.00
VIC
100 µm
top instance:contribution
TuTuA_1_R: 0.024
AOTU018_R: 0.022
AOTU020_R: 0.02
AOTU022_R: 0.02
AOTU006_R: 0.017
AOTU025_R: 0.015
0.010
0.008
0.006
0.004
0.002
0.000
VIC-VPN
100 µm
top instance:contribution
CRE026_L: 0.0021
AOTU008_R: 0.0018
IB058_R: 0.0012
CB1260_R: 0.0012
AOTU060_L: 0.001
VES058_R: 0.00098
0.00030
0.00025
0.00020
0.00015
0.00010
0.00005
0.00000
VIC-VPN
100 µm

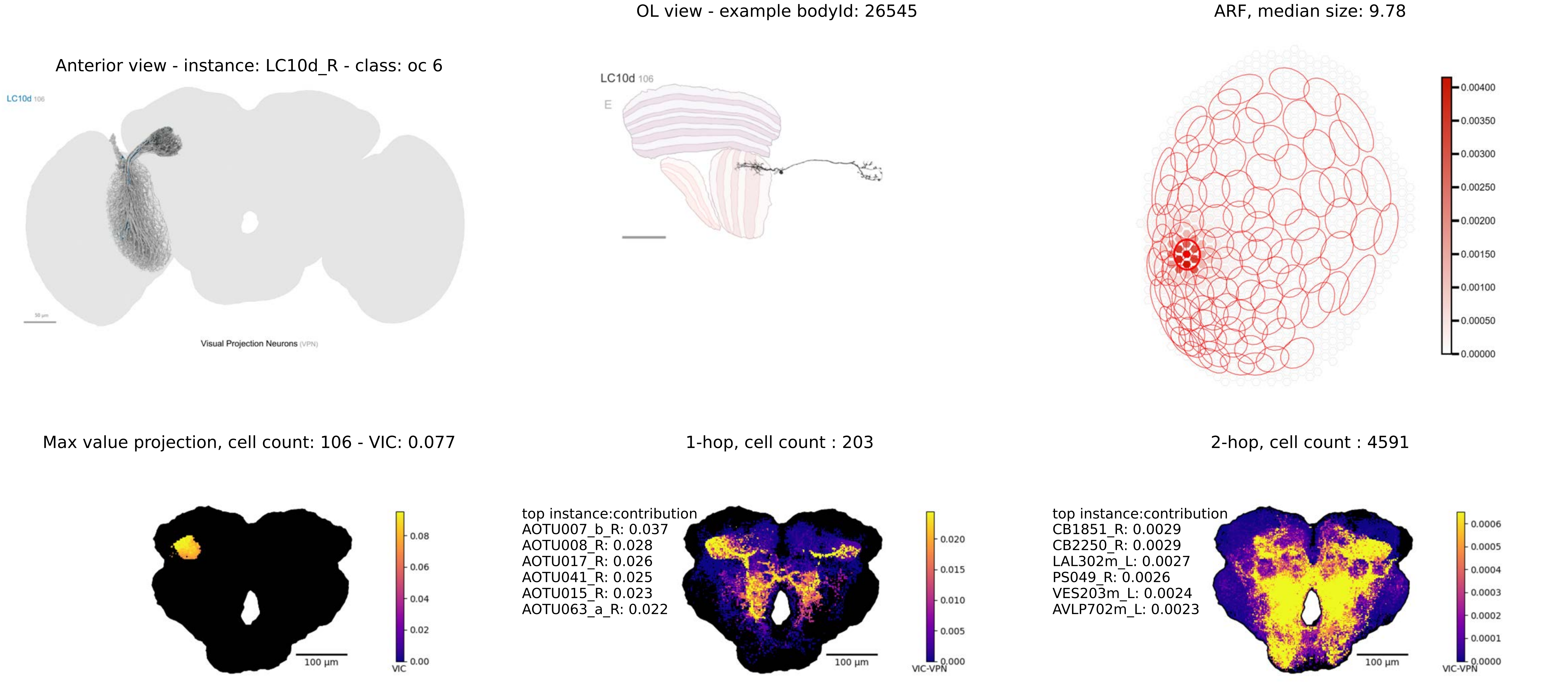

OL view - example bodyId: 26545
ARF, median size: 9.78
Anterior view - instance: LC10d_R - class: oc 6
LC10d 106
E
0.00400
0.00350
0.00300
0.00250
0.00200
0.00150
0.00100
0.00050
0.00000
Visual Projection Neurons (VPN)
Max value projection, cell count: 106 - VIC: 0.077
1-hop, cell count : 203
2-hop, cell count : 4591
top instance:contribution
AOTU007_b_R: 0.037
AOTU008_R: 0.028
AOTU017_R: 0.026
AOTU041_R: 0.025
AOTU015_R: 0.023
AOTU063_a_R: 0.022
top instance:contribution
CB1851_R: 0.0029
CB2250_R: 0.0029
LAL302m_L: 0.0027
PS049_R: 0.0026
VES203m_L: 0.0024
AVLP702m_L: 0.0023
0.08
0.06
0.04
0.02
0.00
VIC
100 µm
0.020
0.015
0.010
0.005
0.000
VIC-VPN
100 µm
0.0006
0.0005
0.0004
0.0003
0.0002
0.0001
0.0000
VIC-VPN
100 µm

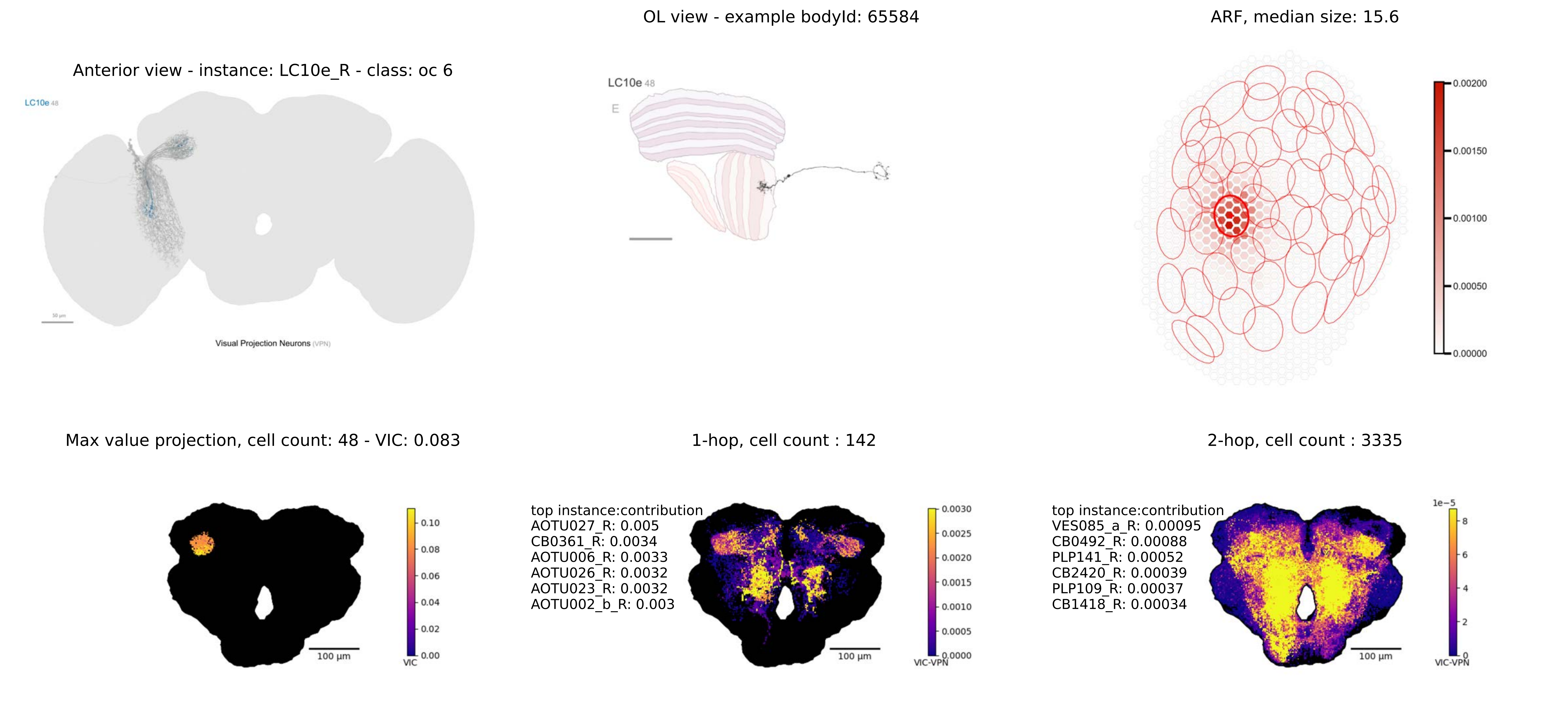
OL view - example bodyId: 65584
ARF, median size: 15.6
Anterior view - instance: LC10e_R - class: oc 6
LC10e 48
Visual Projection Neurons (VPN)
LC10e 48
E
0.00200
0.00150
0.00100
0.00050
0.00000
Max value projection, cell count: 48 - VIC: 0.083
1-hop, cell count : 142
2-hop, cell count : 3335
100 µm
0.10
0.08
0.06
0.04
0.02
0.00
VIC
top instance:contribution
AOTU027_R: 0.005
CB0361_R: 0.0034
AOTU006_R: 0.0033
AOTU026_R: 0.0032
AOTU023_R: 0.0032
AOTU002_b_R: 0.003
100 µm
0.0030
0.0025
0.0020
0.0015
0.0010
0.0005
0.0000
VIC-VPN
top instance:contribution
VES085_a_R: 0.00095
CB0492_R: 0.00088
PLP141_R: 0.00052
CB2420_R: 0.00039
PLP109_R: 0.00037
CB1418_R: 0.00034
100 µm
1e−5
8
6
4
2
0
VIC-VPN

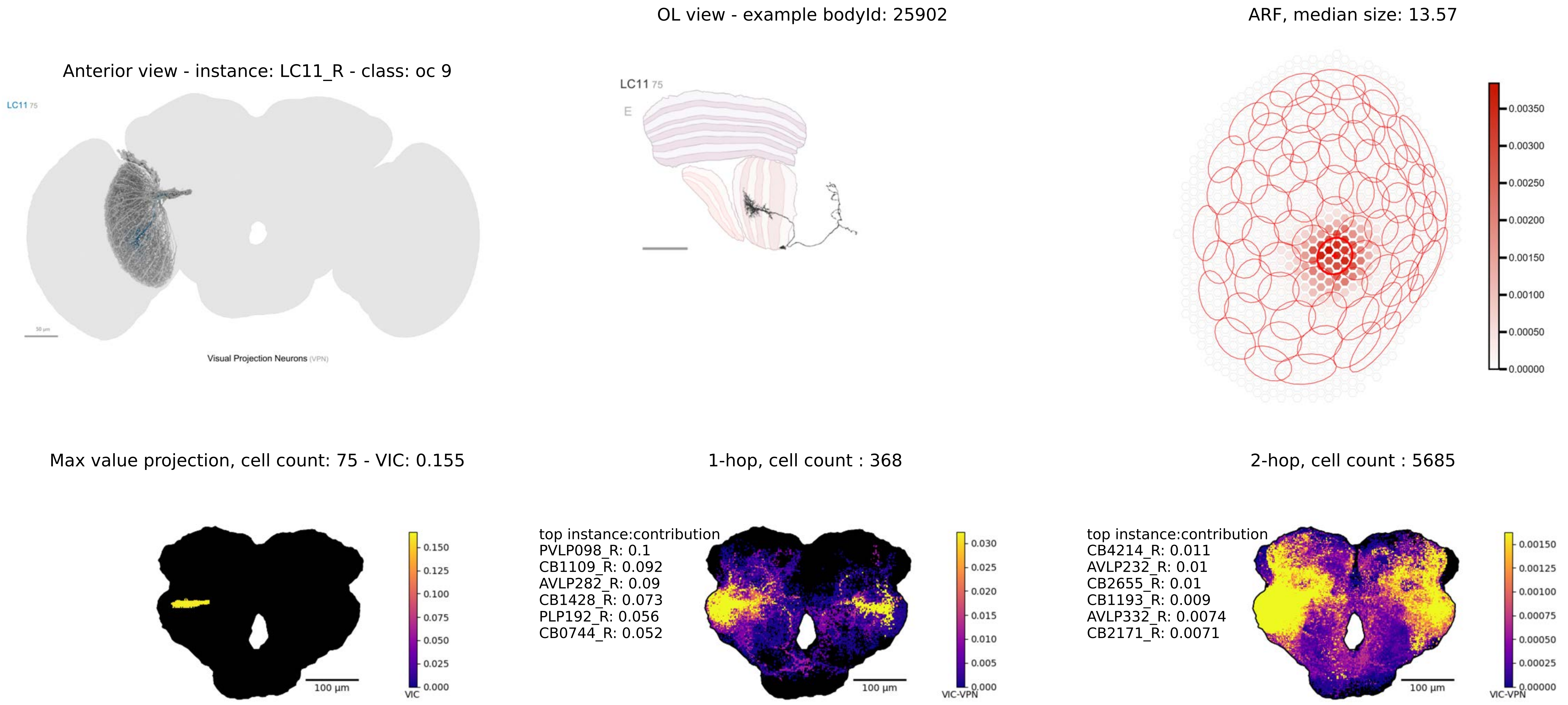
OL view - example bodyId: 25902
ARF, median size: 13.57
Anterior view - instance: LC11_R - class: oc 9
LC11 75
Visual Projection Neurons (VPN)
0.00350
0.00300
0.00250
0.00200
0.00150
0.00100
0.00050
0.00000
Max value projection, cell count: 75 - VIC: 0.155
1-hop, cell count : 368
2-hop, cell count : 5685
top instance:contribution
PVLP098_R: 0.1
CB1109_R: 0.092
AVLP282_R: 0.09
CB1428_R: 0.073
PLP192_R: 0.056
CB0744_R: 0.052
top instance:contribution
CB4214_R: 0.011
AVLP232_R: 0.01
CB2655_R: 0.01
CB1193_R: 0.009
AVLP332_R: 0.0074
CB2171_R: 0.0071
100 µm
VIC
0.150
0.125
0.100
0.075
0.050
0.025
0.000
VIC-VPN
0.030
0.025
0.020
0.015
0.010
0.005
0.000
0.00150
0.00125
0.00100
0.00075
0.00050
0.00025
0.00000

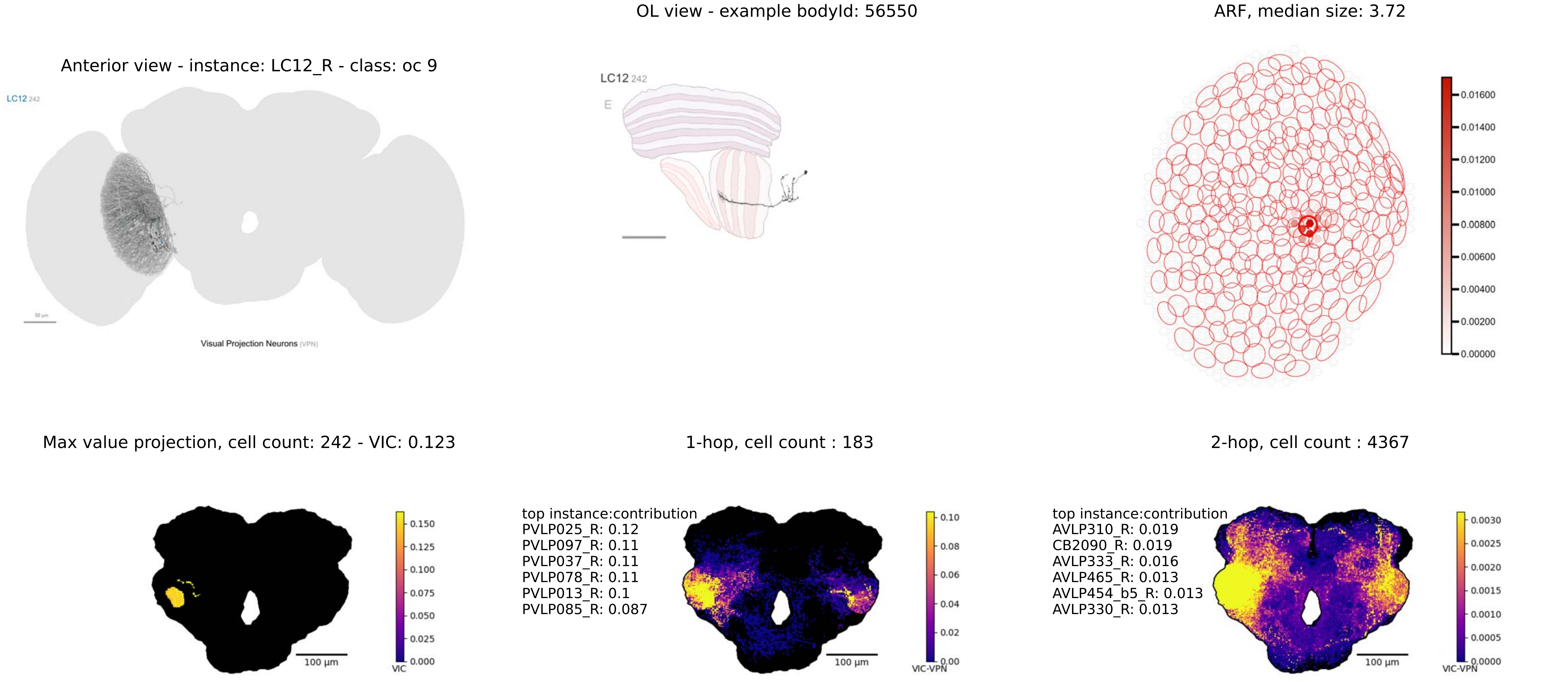
OL view - example bodyId: 56550
ARF, median size: 3.72
Anterior view - instance: LC12_R - class: oc 9
LC12 242
Visual Projection Neurons (VPN)
LC12 242
E
Max value projection, cell count: 242 - VIC: 0.123
1-hop, cell count : 183
2-hop, cell count : 4367
top instance:contribution
PVLP025_R: 0.12
PVLP097_R: 0.11
PVLP037_R: 0.11
PVLP078_R: 0.11
PVLP013_R: 0.1
PVLP085_R: 0.087
top instance:contribution
AVLP310_R: 0.019
CB2090_R: 0.019
AVLP333_R: 0.016
AVLP465_R: 0.013
AVLP454_b5_R: 0.013
AVLP330_R: 0.013
100 µm
100 µm
100 µm
VIC
VIC-VPN
VIC-VPN

OL view - example bodyId: 59045

ARF, median size: 8.82

Anterior view - instance: LC13_R - class: oc 6

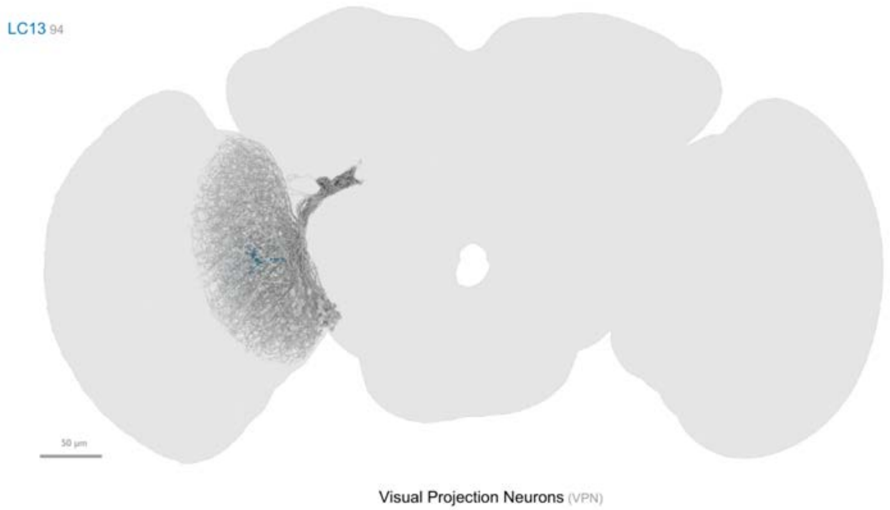


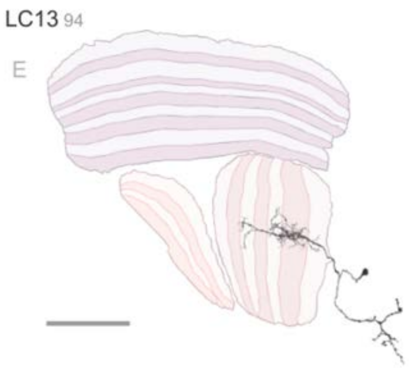


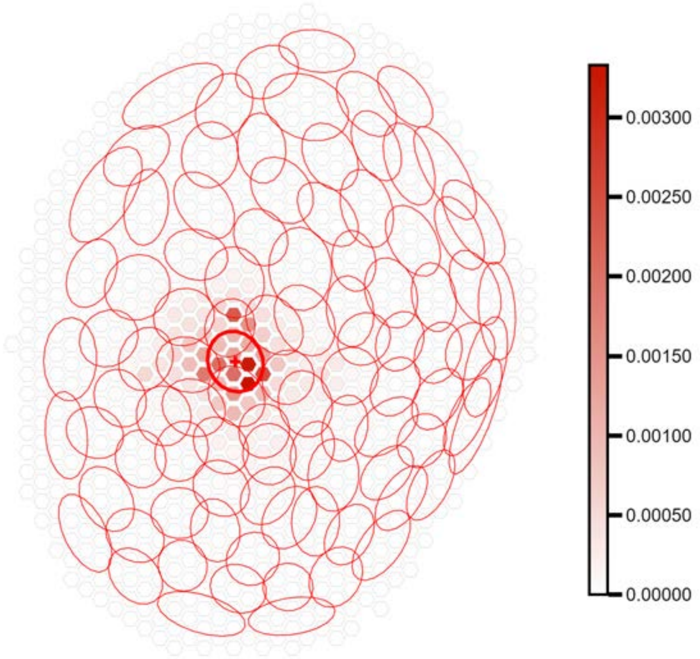


Max value projection, cell count: 94 - VIC: 0.068

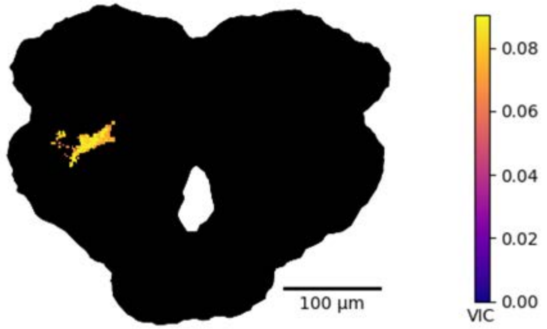


1-hop, cell count : 160

top instance:contribution
PLP008_R: 0.055
PLP016_R: 0.053
AVLP464_R: 0.042
PLP015_R: 0.036
PLP108_R: 0.035
PLP076_R: 0.026

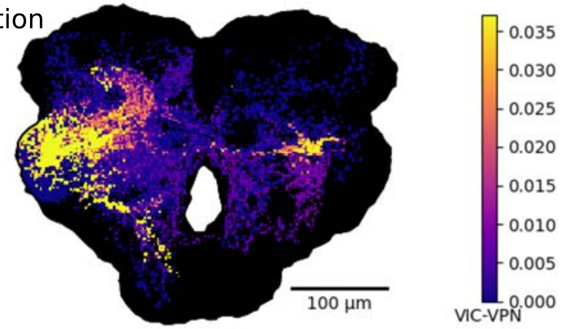


2-hop, cell count : 5740

top instance:contribution
CB4170_R: 0.011
CB4169_R: 0.0058
PVLP134_R: 0.0041
AVLP198_R: 0.0038
CB1269_R: 0.0032
CB0682_R: 0.0032

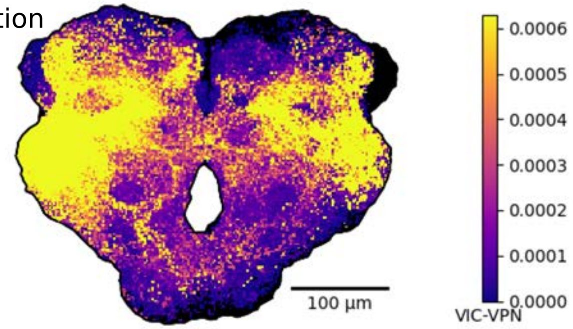

Anterior view - instance: LC14a-1_R - class: oc 9

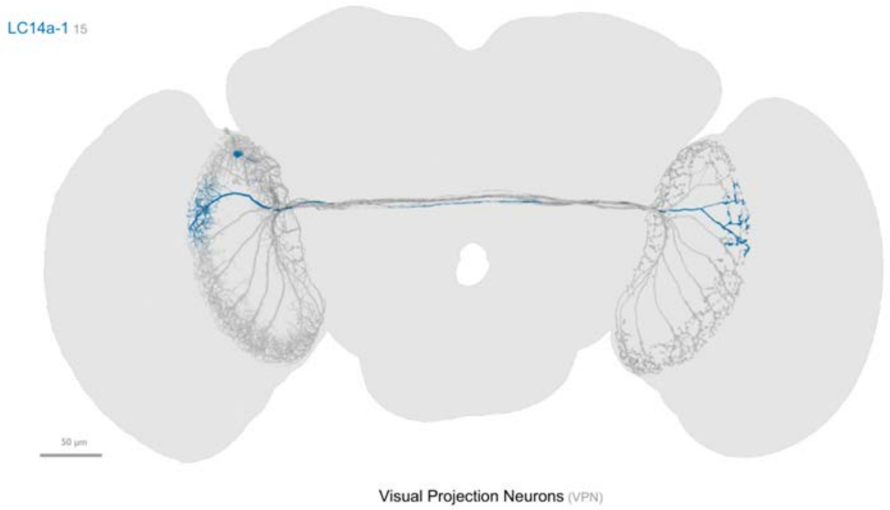


OL view - example bodyId: 24305

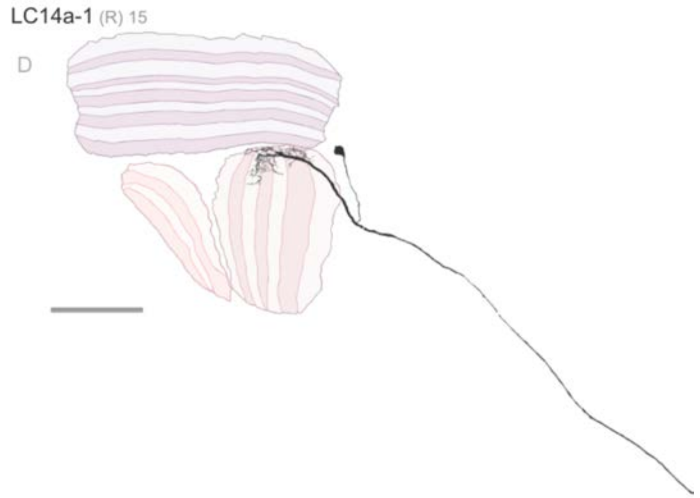


ARF, median size: 13.33

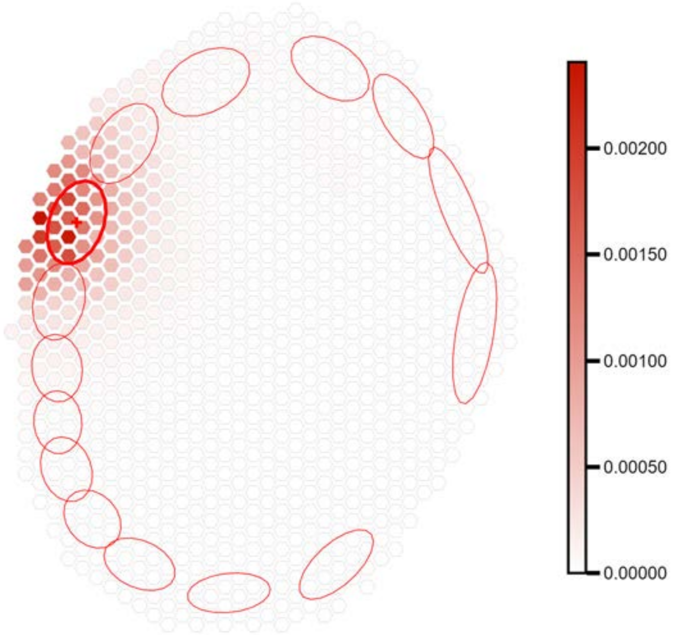


fewer than 100 output connections in central brain

Anterior view - instance: LC14a-2_R - class: oc 6

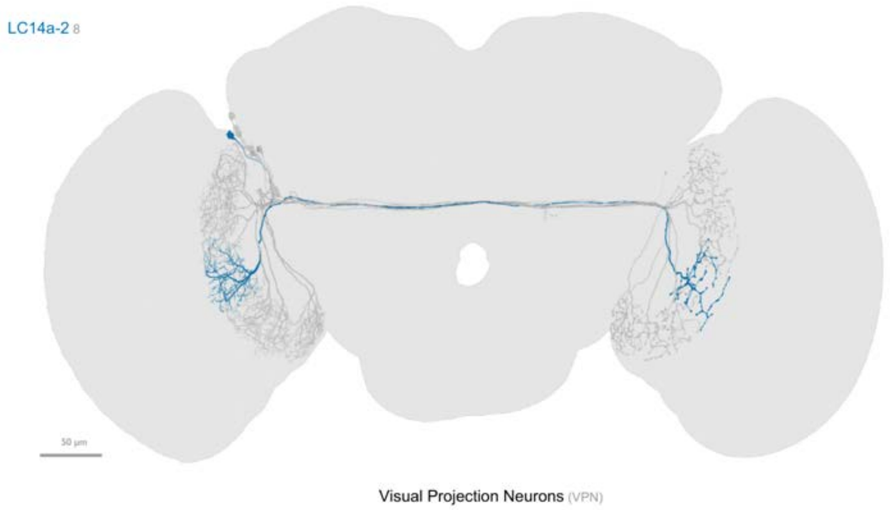


fewer than 100 output connections in central brain

OL view - example bodyId: 22025

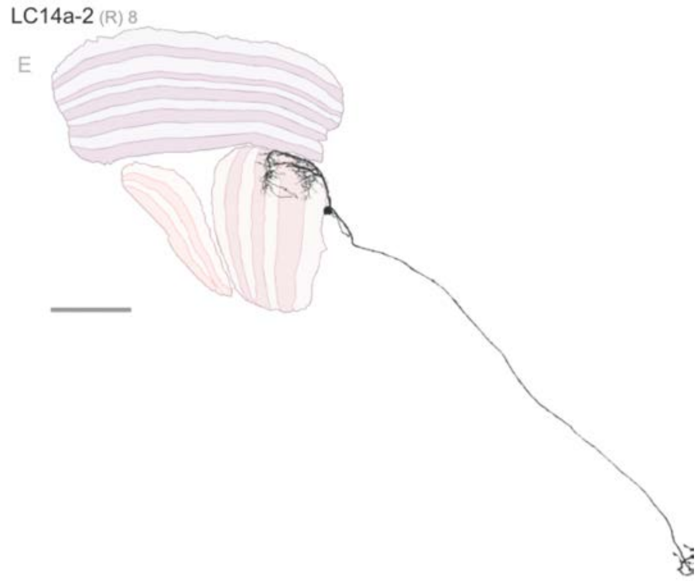


ARF, median size: 17.74

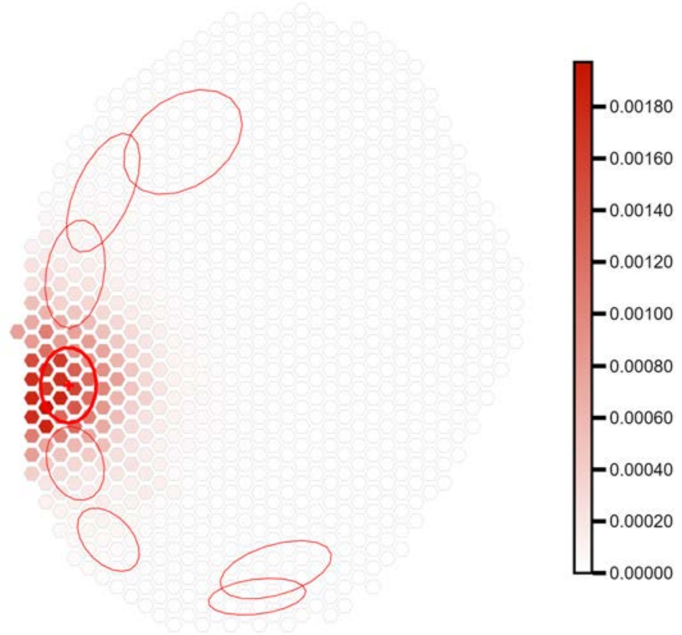

OL view - example bodyId: 21081

ARF, median size: 9.54

Anterior view - instance: LC14b_R - class: oc 8

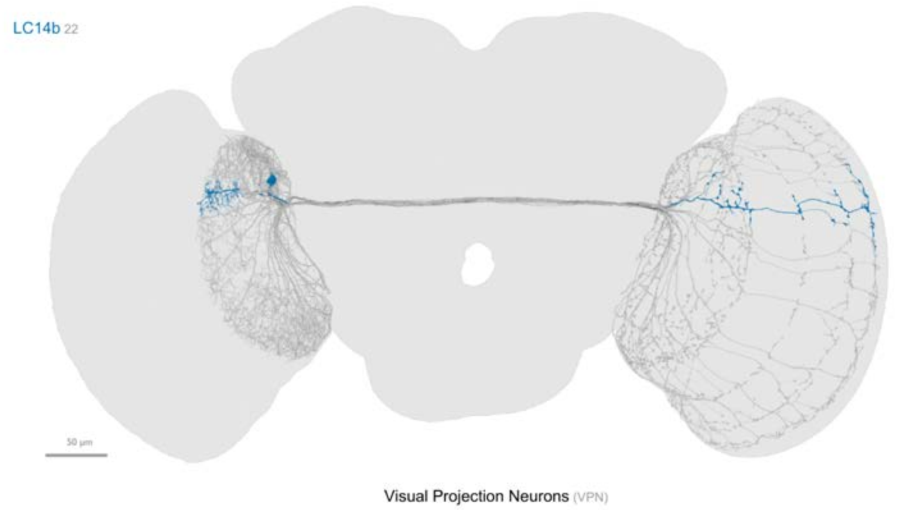


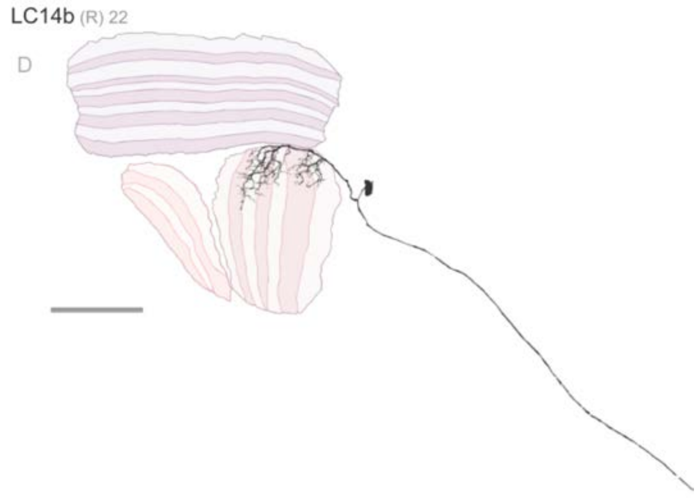


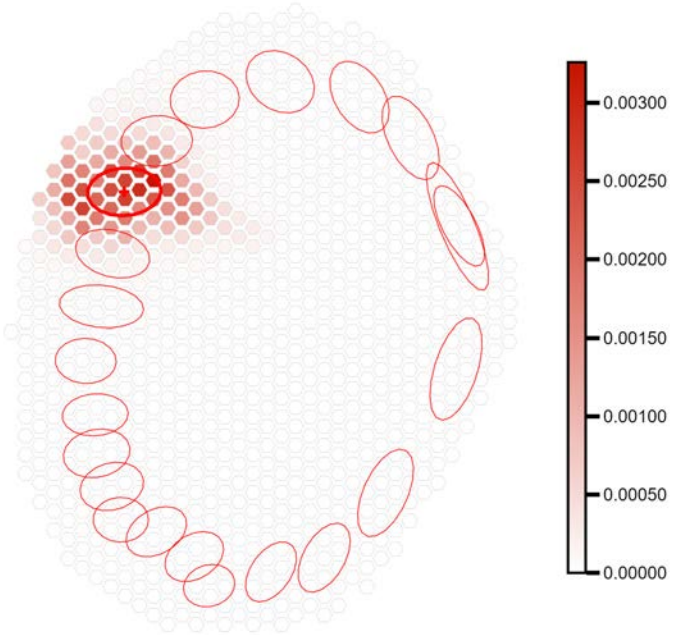


fewer than 100 output connections in central brain

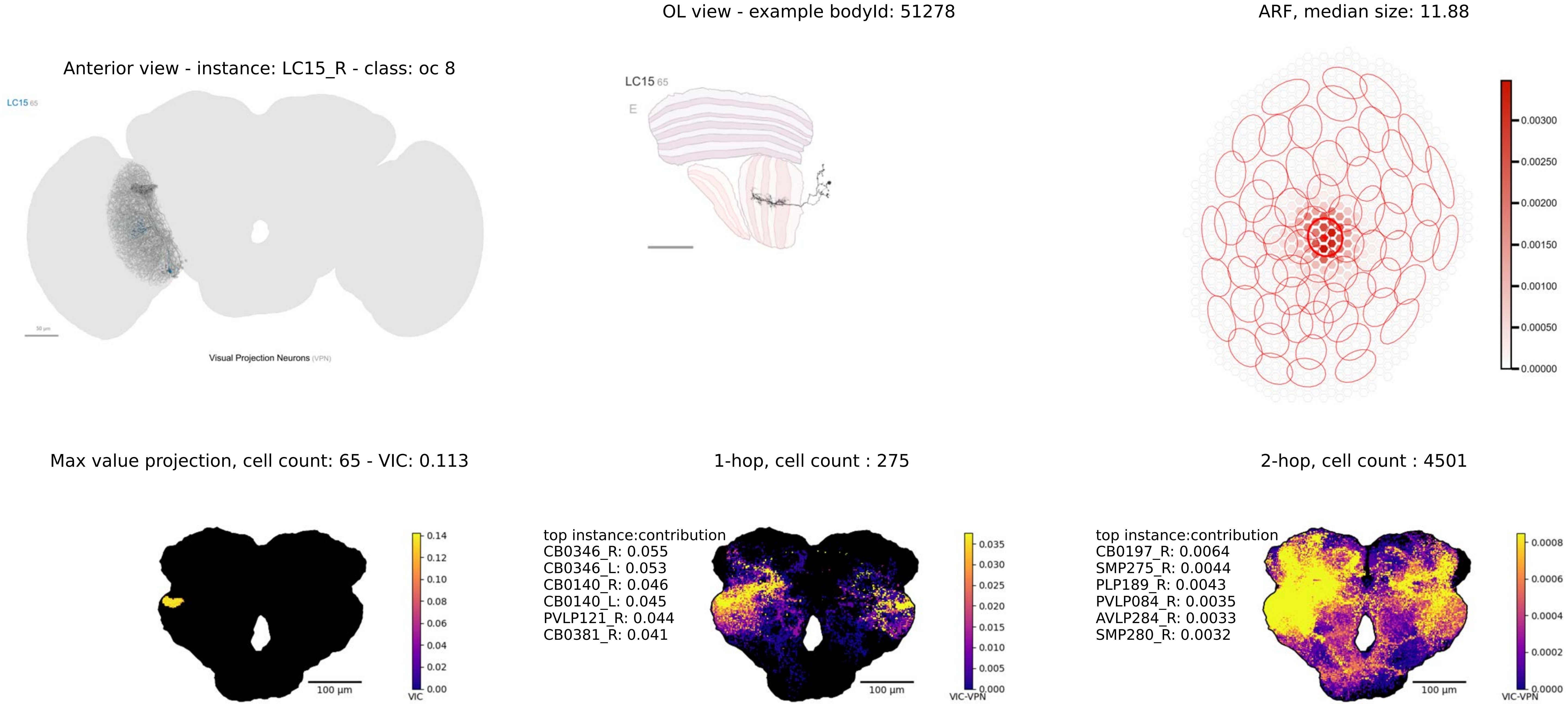
OL view - example bodyId: 51278
ARF, median size: 11.88
Anterior view - instance: LC15_R - class: oc 8
LC15 65
Visual Projection Neurons (VPN)
LC15 65
E
0.00300
0.00250
0.00200
0.00150
0.00100
0.00050
0.00000
Max value projection, cell count: 65 - VIC: 0.113
1-hop, cell count : 275
2-hop, cell count : 4501
0.14
0.12
0.10
0.08
0.06
0.04
0.02
0.00
VIC
100 µm
top instance:contribution
CB0346_R: 0.055
CB0346_L: 0.053
CB0140_R: 0.046
CB0140_L: 0.045
PVLP121_R: 0.044
CB0381_R: 0.041
0.035
0.030
0.025
0.020
0.015
0.010
0.005
0.000
VIC-VPN
100 µm
top instance:contribution
CB0197_R: 0.0064
SMP275_R: 0.0044
PLP189_R: 0.0043
PVLP084_R: 0.0035
AVLP284_R: 0.0033
SMP280_R: 0.0032
0.0008
0.0006
0.0004
0.0002
0.0000
VIC-VPN
100 µm

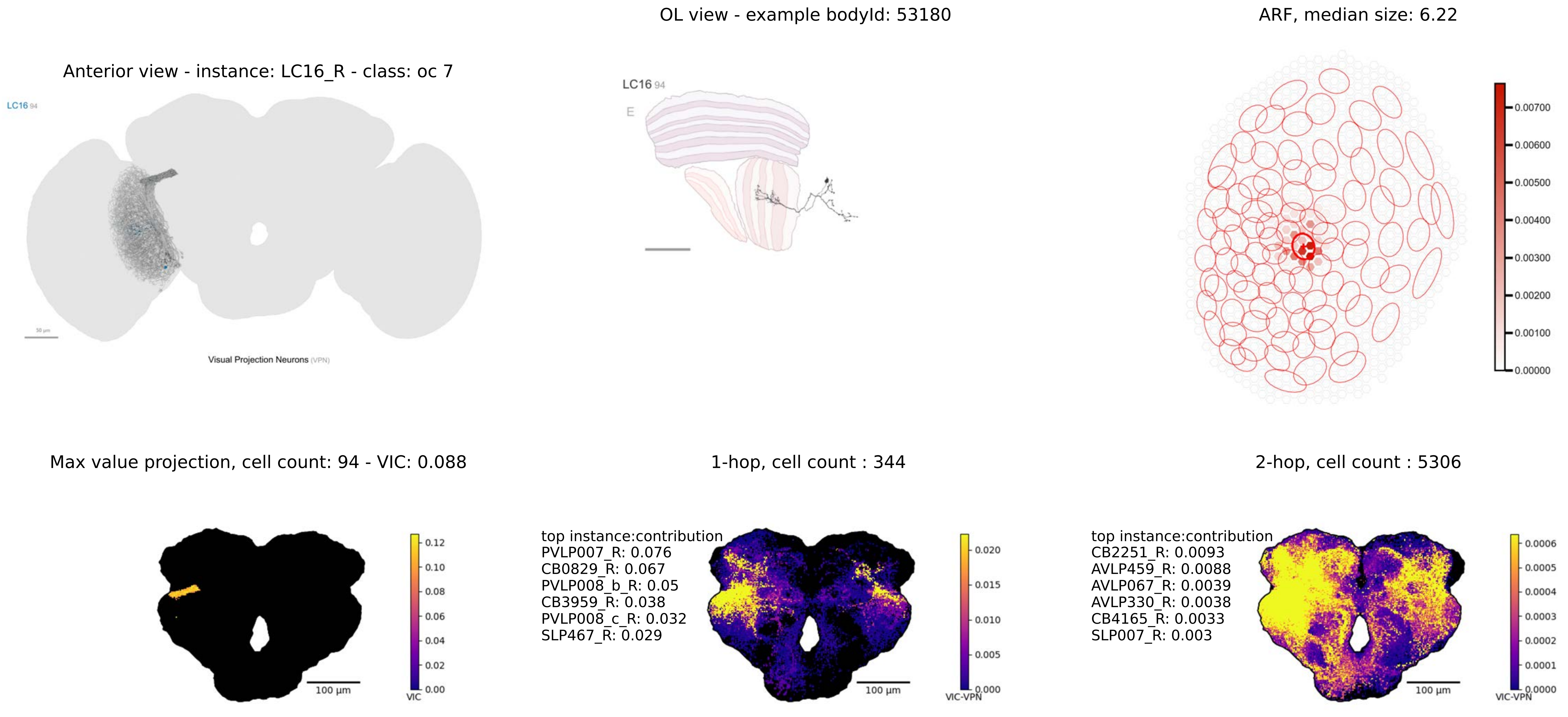

OL view - example bodyId: 53180
ARF, median size: 6.22
Anterior view - instance: LC16_R - class: oc 7
LC16 94
Visual Projection Neurons (VPN)
LC16 94
E
Max value projection, cell count: 94 - VIC: 0.088
1-hop, cell count : 344
2-hop, cell count : 5306
top instance:contribution
PVLP007_R: 0.076
CB0829_R: 0.067
PVLP008_b_R: 0.05
CB3959_R: 0.038
PVLP008_c_R: 0.032
SLP467_R: 0.029
top instance:contribution
CB2251_R: 0.0093
AVLP459_R: 0.0088
AVLP067_R: 0.0039
AVLP330_R: 0.0038
CB4165_R: 0.0033
SLP007_R: 0.003
100 µm
100 µm
100 µm
VIC
VIC-VPN
VIC-VPN

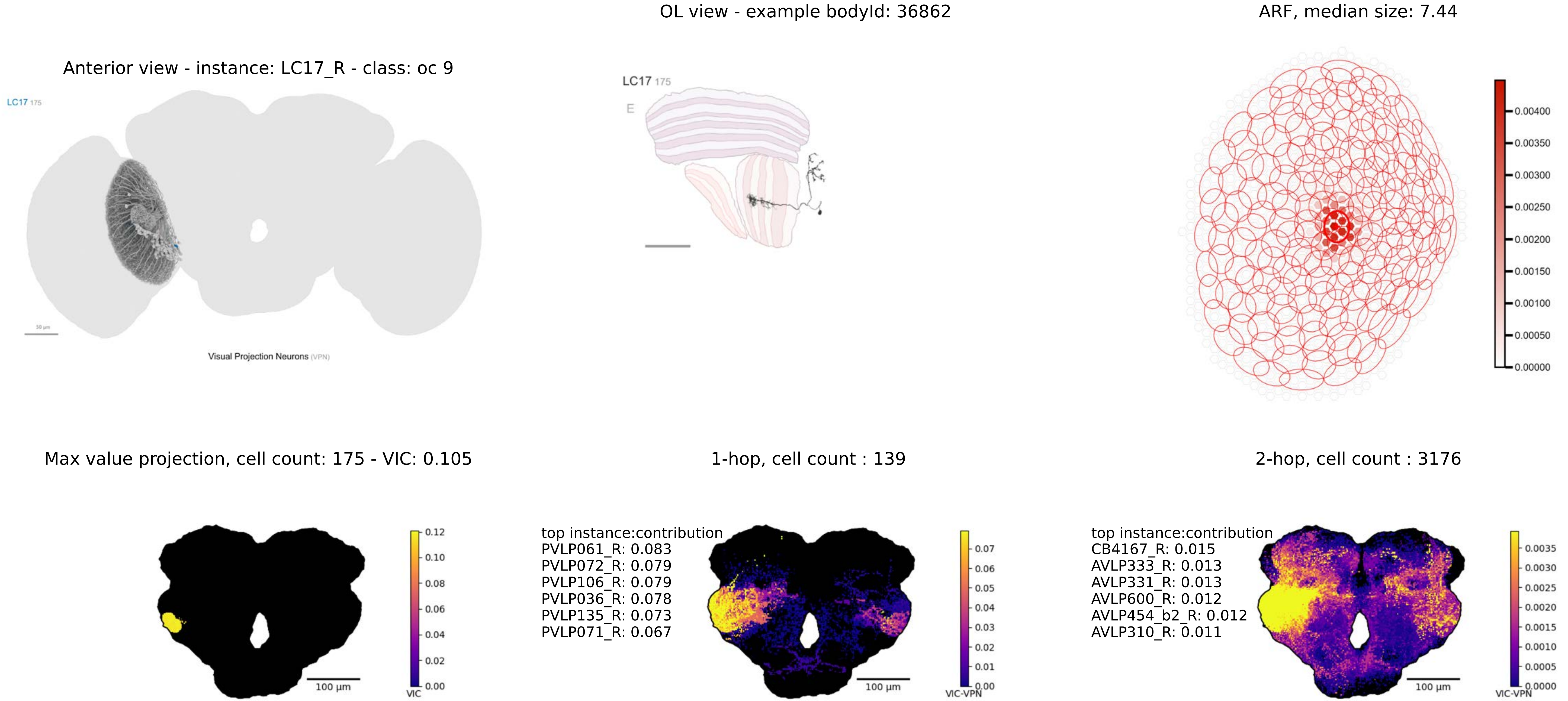
Anterior view - instance: LC17_R - class: oc 9
LC17 175
Visual Projection Neurons (VPN)
OL view - example bodyId: 36862
LC17 175
E
ARF, median size: 7.44
0.00400
0.00350
0.00300
0.00250
0.00200
0.00150
0.00100
0.00050
0.00000
Max value projection, cell count: 175 - VIC: 0.105
100 µm
VIC
0.12
0.10
0.08
0.06
0.04
0.02
0.00
1-hop, cell count : 139
top instance:contribution
PVLP061_R: 0.083
PVLP072_R: 0.079
PVLP106_R: 0.079
PVLP036_R: 0.078
PVLP135_R: 0.073
PVLP071_R: 0.067
100 µm
VIC-VPN
0.07
0.06
0.05
0.04
0.03
0.02
0.01
0.00
2-hop, cell count : 3176
top instance:contribution
CB4167_R: 0.015
AVLP333_R: 0.013
AVLP331_R: 0.013
AVLP600_R: 0.012
AVLP454_b2_R: 0.012
AVLP310_R: 0.011
100 µm
VIC-VPN
0.0035
0.0030
0.0025
0.0020
0.0015
0.0010
0.0005
0.0000

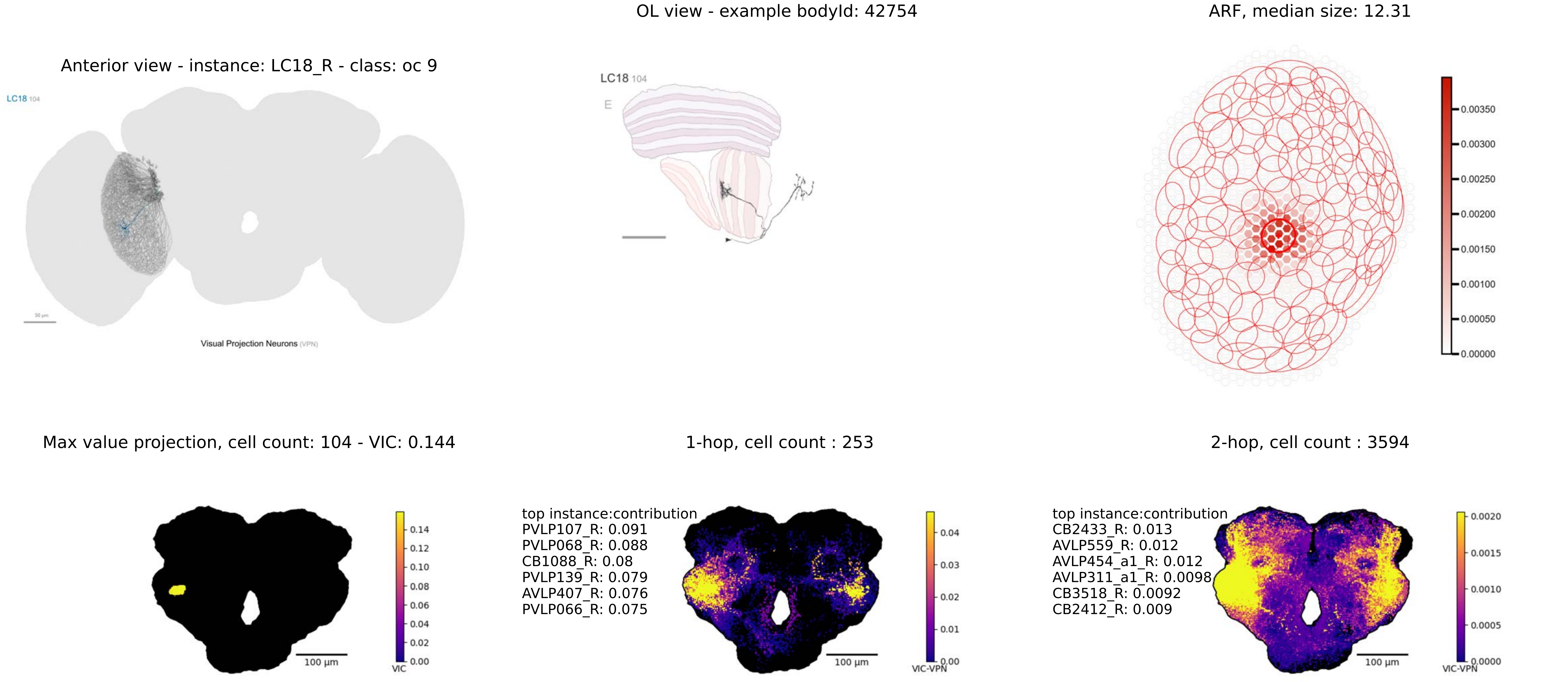
OL view - example bodyId: 42754
ARF, median size: 12.31
Anterior view - instance: LC18_R - class: oc 9
LC18 104
Visual Projection Neurons (VPN)
LC18 104
E
0.00350
0.00300
0.00250
0.00200
0.00150
0.00100
0.00050
0.00000
Max value projection, cell count: 104 - VIC: 0.144
1-hop, cell count : 253
2-hop, cell count : 3594
0.14
0.12
0.10
0.08
0.06
0.04
0.02
0.00
VIC
100 µm
top instance:contribution
PVLP107_R: 0.091
PVLP068_R: 0.088
CB1088_R: 0.08
PVLP139_R: 0.079
AVLP407_R: 0.076
PVLP066_R: 0.075
0.04
0.03
0.02
0.01
0.00
VIC-VPN
100 µm
top instance:contribution
CB2433_R: 0.013
AVLP559_R: 0.012
AVLP454_a1_R: 0.012
AVLP311_a1_R: 0.0098
CB3518_R: 0.0092
CB2412_R: 0.009
0.0020
0.0015
0.0010
0.0005
0.0000
VIC-VPN
100 µm

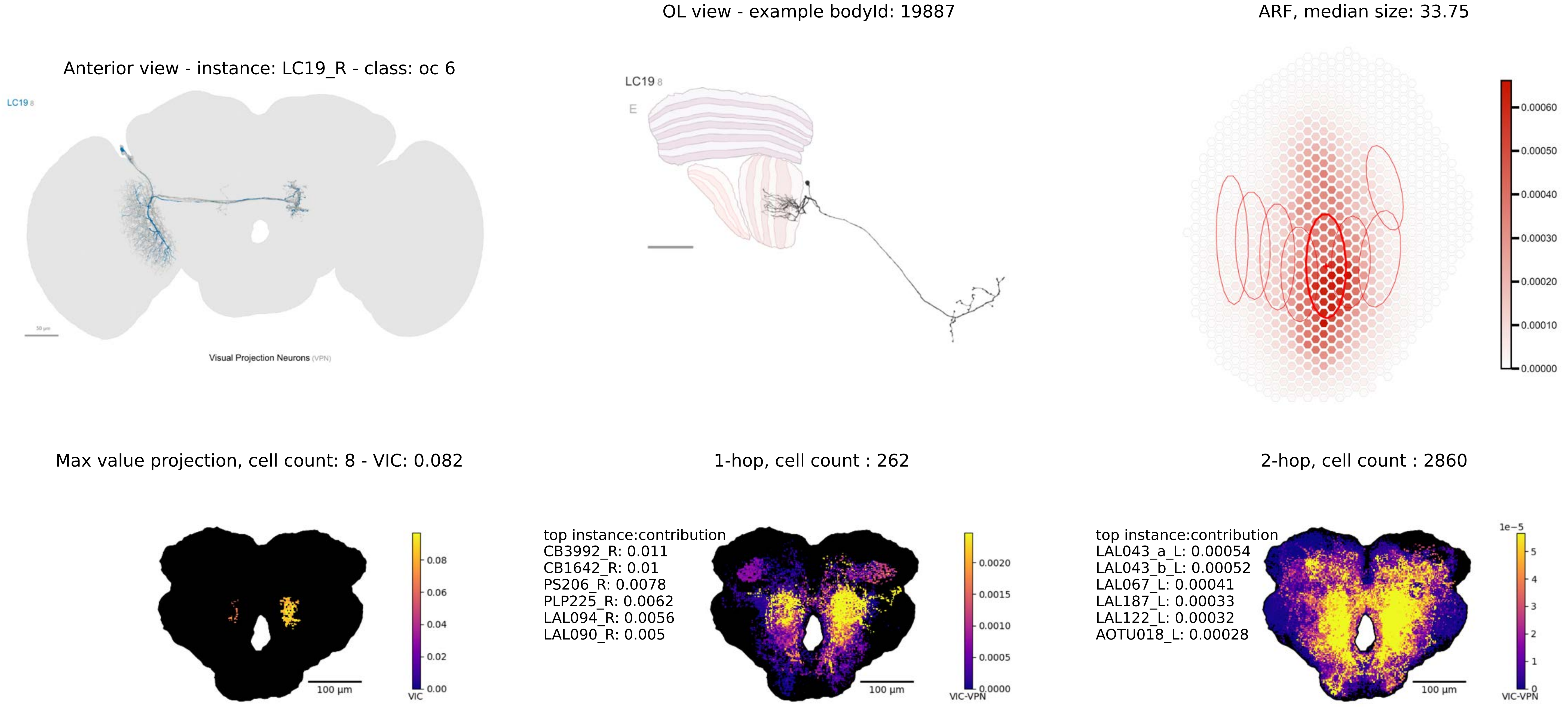
Anterior view - instance: LC19_R - class: oc 6
LC19 8
Visual Projection Neurons (VPN)
OL view - example bodyId: 19887
LC19 8
E
ARF, median size: 33.75
0.00060
0.00050
0.00040
0.00030
0.00020
0.00010
0.00000
Max value projection, cell count: 8 - VIC: 0.082
100 µm
0.08
0.06
0.04
0.02
0.00
VIC
1-hop, cell count : 262
top instance:contribution
CB3992_R: 0.011
CB1642_R: 0.01
PS206_R: 0.0078
PLP225_R: 0.0062
LAL094_R: 0.0056
LAL090_R: 0.005
100 µm
0.0020
0.0015
0.0010
0.0005
0.0000
VIC-VPN
2-hop, cell count : 2860
top instance:contribution
LAL043_a_L: 0.00054
LAL043_b_L: 0.00052
LAL067_L: 0.00041
LAL187_L: 0.00033
LAL122_L: 0.00032
AOTU018_L: 0.00028
100 µm
1e−5
5
4
3
2
1
0
VIC-VPN

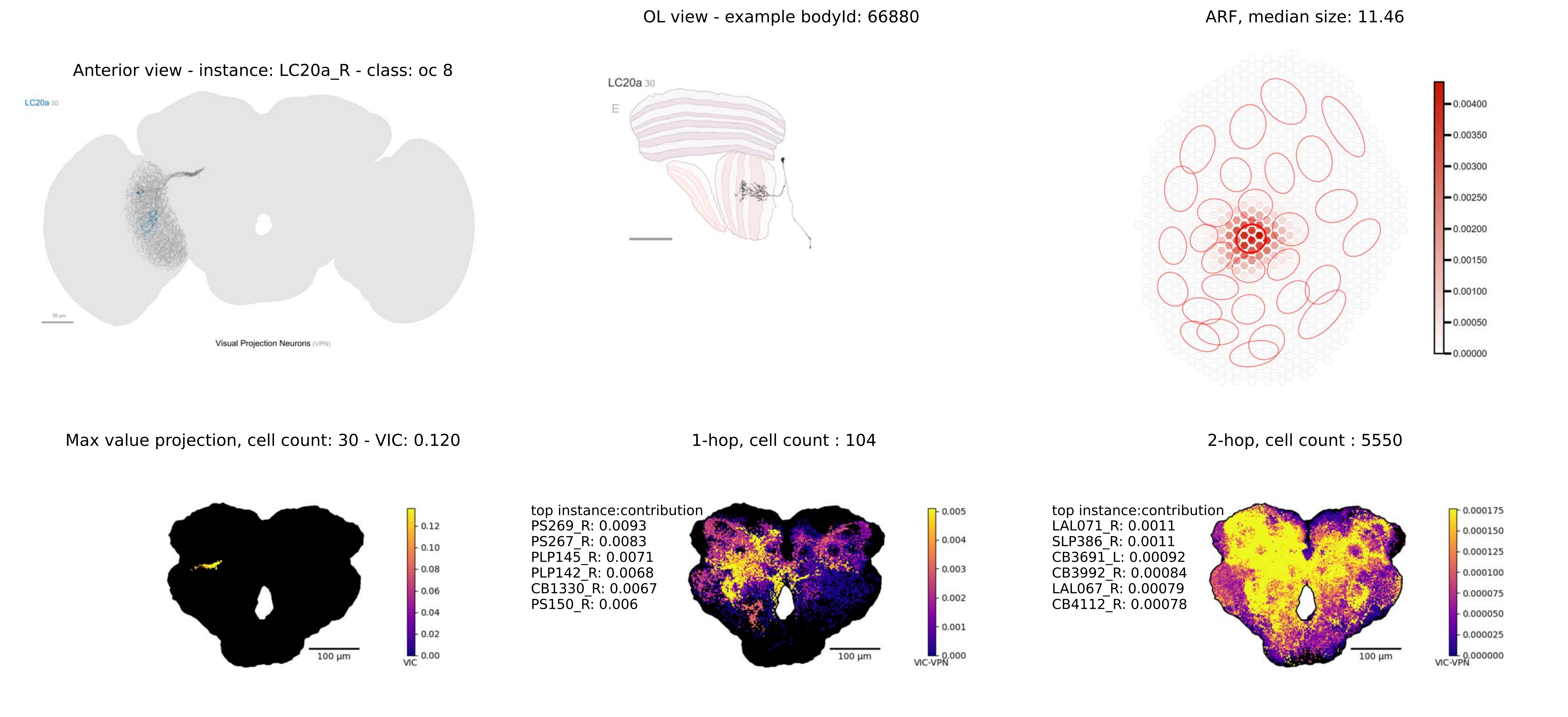
OL view - example bodyId: 66880
ARF, median size: 11.46
Anterior view - instance: LC20a_R - class: oc 8
LC20a 30
Visual Projection Neurons (VPN)
LC20a 30
E
0.00400
0.00350
0.00300
0.00250
0.00200
0.00150
0.00100
0.00050
0.00000
Max value projection, cell count: 30 - VIC: 0.120
1-hop, cell count : 104
2-hop, cell count : 5550
0.12
0.10
0.08
0.06
0.04
0.02
0.00
VIC
100 µm
top instance:contribution
PS269_R: 0.0093
PS267_R: 0.0083
PLP145_R: 0.0071
PLP142_R: 0.0068
CB1330_R: 0.0067
PS150_R: 0.006
0.005
0.004
0.003
0.002
0.001
0.000
VIC-VPN
100 µm
top instance:contribution
LAL071_R: 0.0011
SLP386_R: 0.0011
CB3691_L: 0.00092
CB3992_R: 0.00084
LAL067_R: 0.00079
CB4112_R: 0.00078
0.000175
0.000150
0.000125
0.000100
0.000075
0.000050
0.000025
0.000000
VIC-VPN
100 µm

OL view - example bodyId: 55338

ARF, median size: 13.19

Anterior view - instance: LC20b_R - class: oc 8

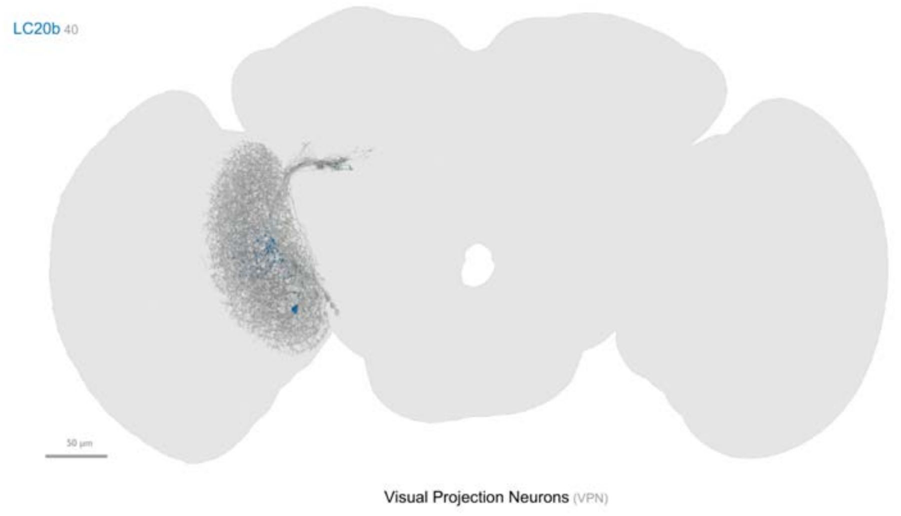


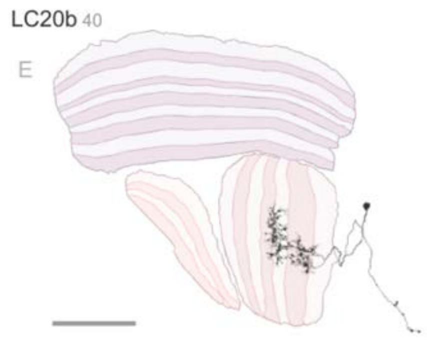


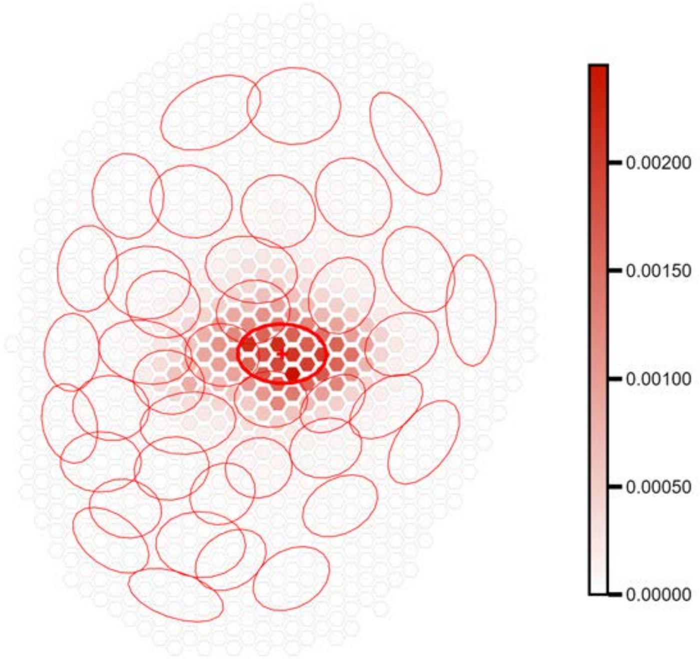


Max value projection, cell count: 40 - VIC: 0.109

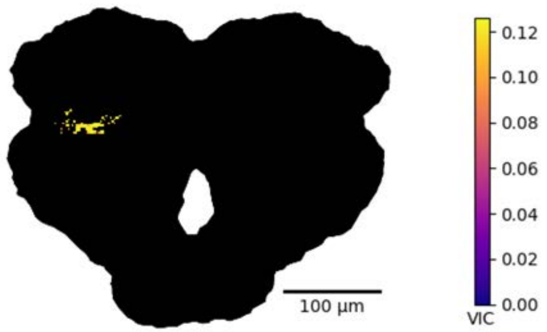


1-hop, cell count : 98

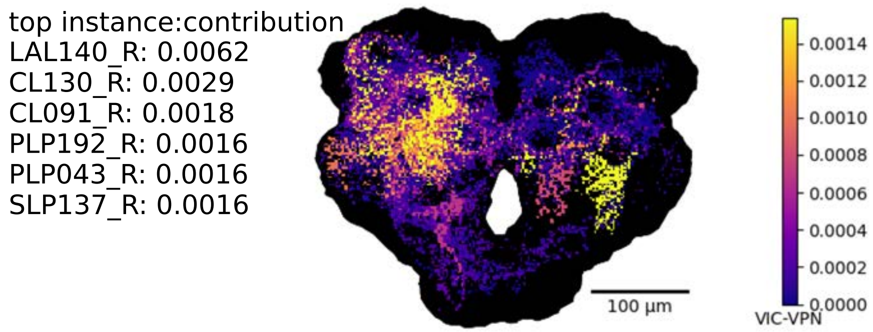


2-hop, cell count : 5368

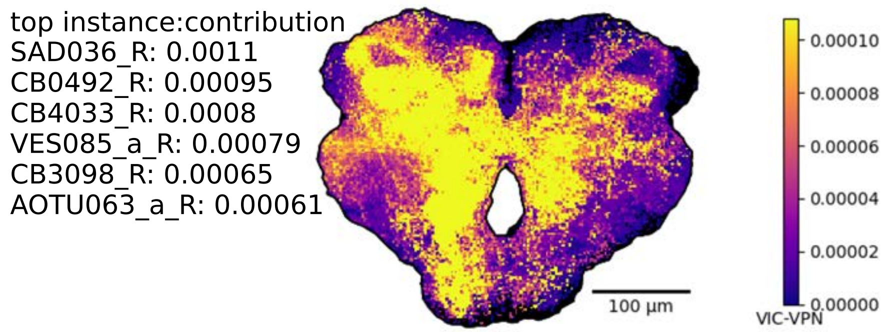

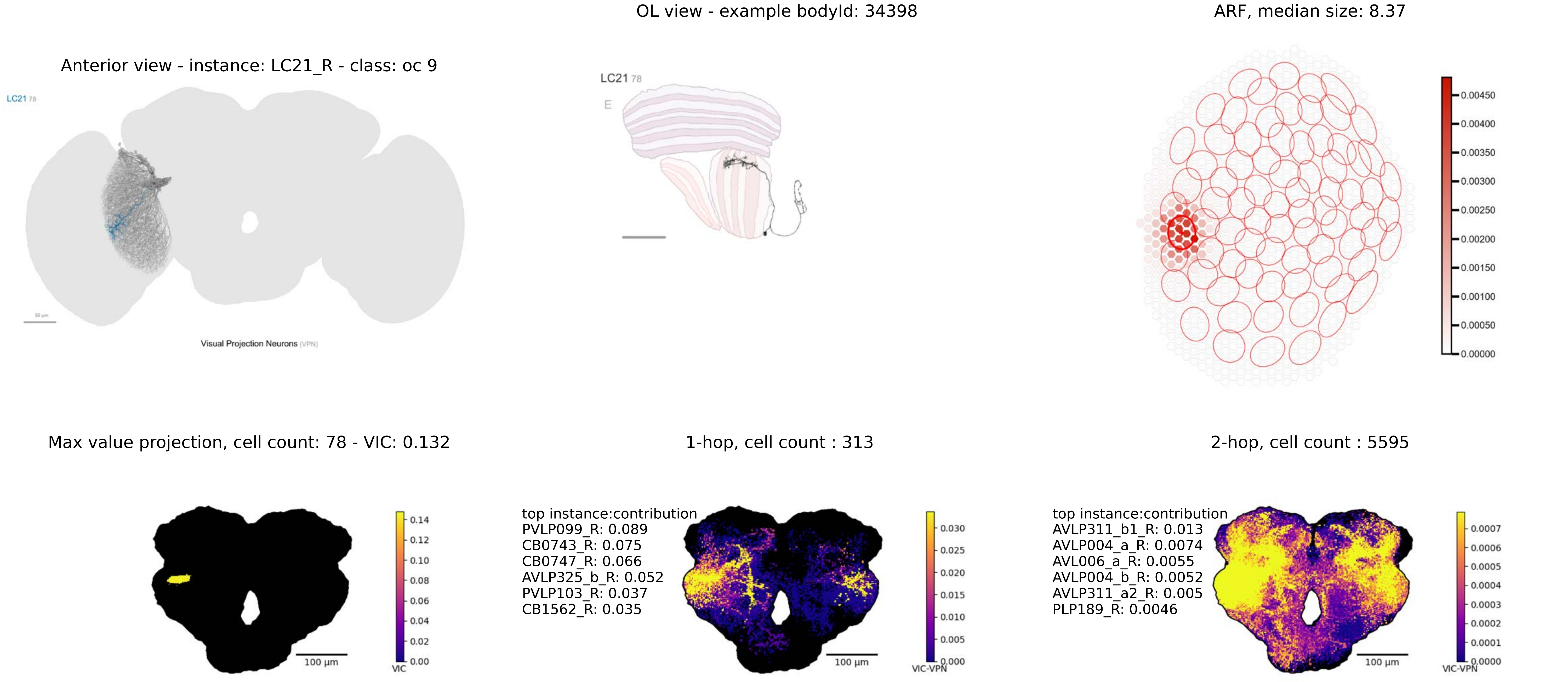
Anterior view - instance: LC21_R - class: oc 9
LC21 78
Visual Projection Neurons (VPN)
OL view - example bodyId: 34398
LC21 78
E
ARF, median size: 8.37
0.00450
0.00400
0.00350
0.00300
0.00250
0.00200
0.00150
0.00100
0.00050
0.00000
Max value projection, cell count: 78 - VIC: 0.132
100 µm
VIC
0.14
0.12
0.10
0.08
0.06
0.04
0.02
0.00
1-hop, cell count : 313
top instance:contribution
PVLP099_R: 0.089
CB0743_R: 0.075
CB0747_R: 0.066
AVLP325_b_R: 0.052
PVLP103_R: 0.037
CB1562_R: 0.035
100 µm
VIC-VPN
0.030
0.025
0.020
0.015
0.010
0.005
0.000
2-hop, cell count : 5595
top instance:contribution
AVLP311_b1_R: 0.013
AVLP004_a_R: 0.0074
AVL006_a_R: 0.0055
AVLP004_b_R: 0.0052
AVLP311_a2_R: 0.005
PLP189_R: 0.0046
100 µm
VIC-VPN
0.0007
0.0006
0.0005
0.0004
0.0003
0.0002
0.0001
0.0000

OL view - example bodyId: 36373

ARF, median size: 11.73

Anterior view - instance: LC22_R - class: oc 8

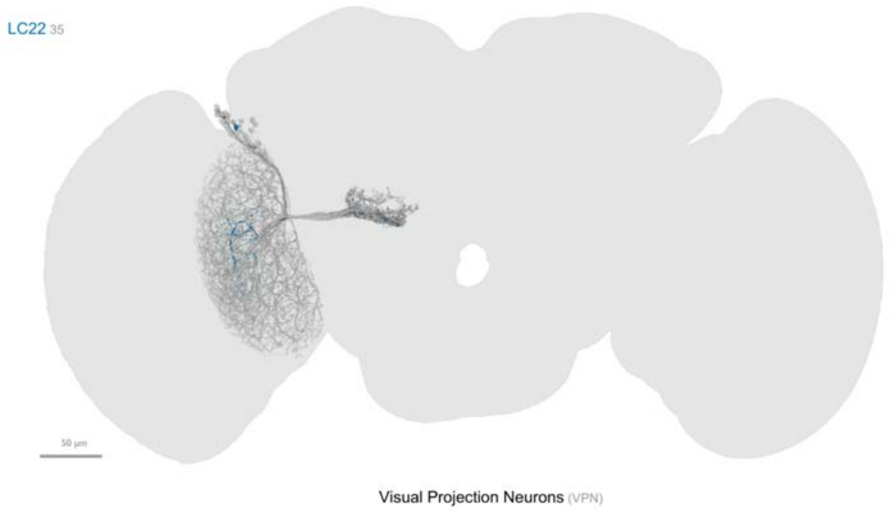


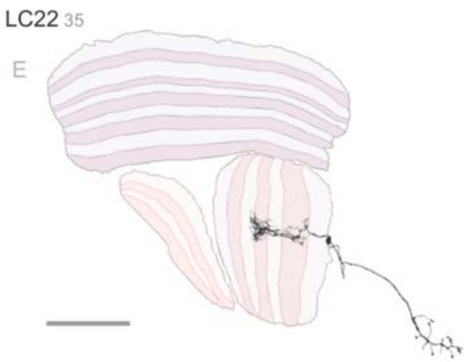


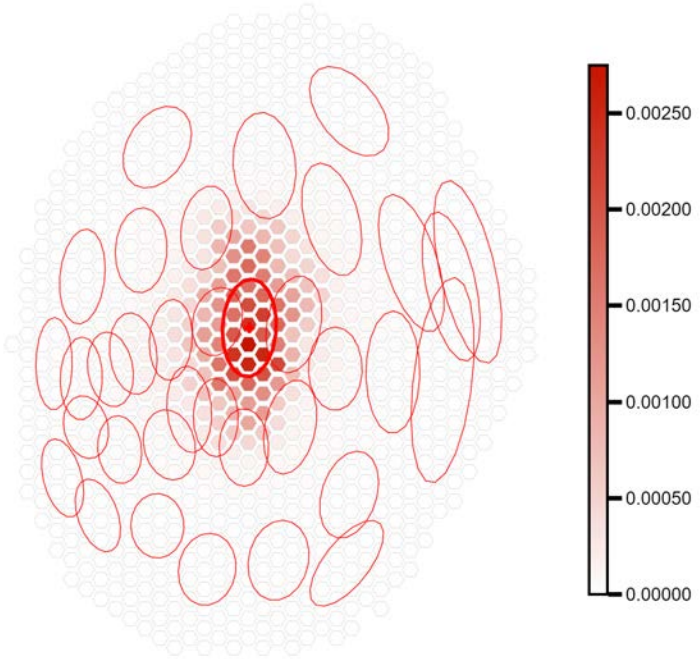


Max value projection, cell count: 35 - VIC: 0.115

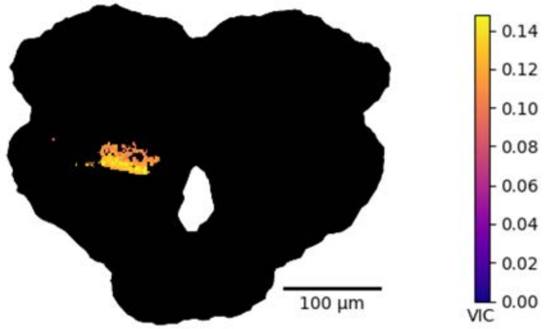


1-hop, cell count : 241

top instance:contribution
PLP009_R: 0.03
PLP172_R: 0.016
PS003_R: 0.014
PS021_R: 0.01
PLP093_R: 0.0097
PS007_R: 0.0096

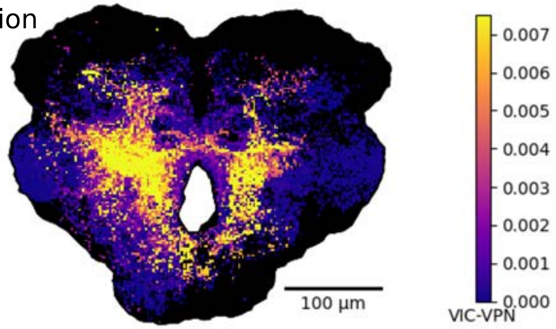


2-hop, cell count : 5749

top instance:contribution
DNa15_R: 0.0011
PS024_R: 0.001
DNg01_d_R: 0.00086
PS007_L: 0.00081
DNae002_R: 0.00074
DNge017_R: 0.00072

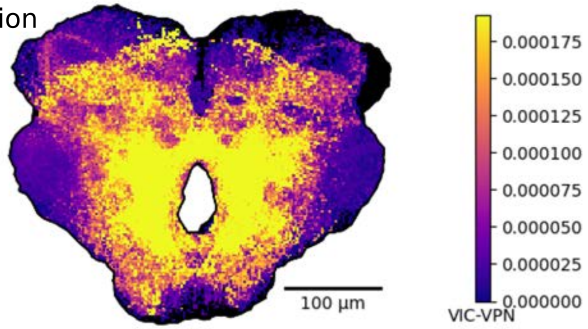

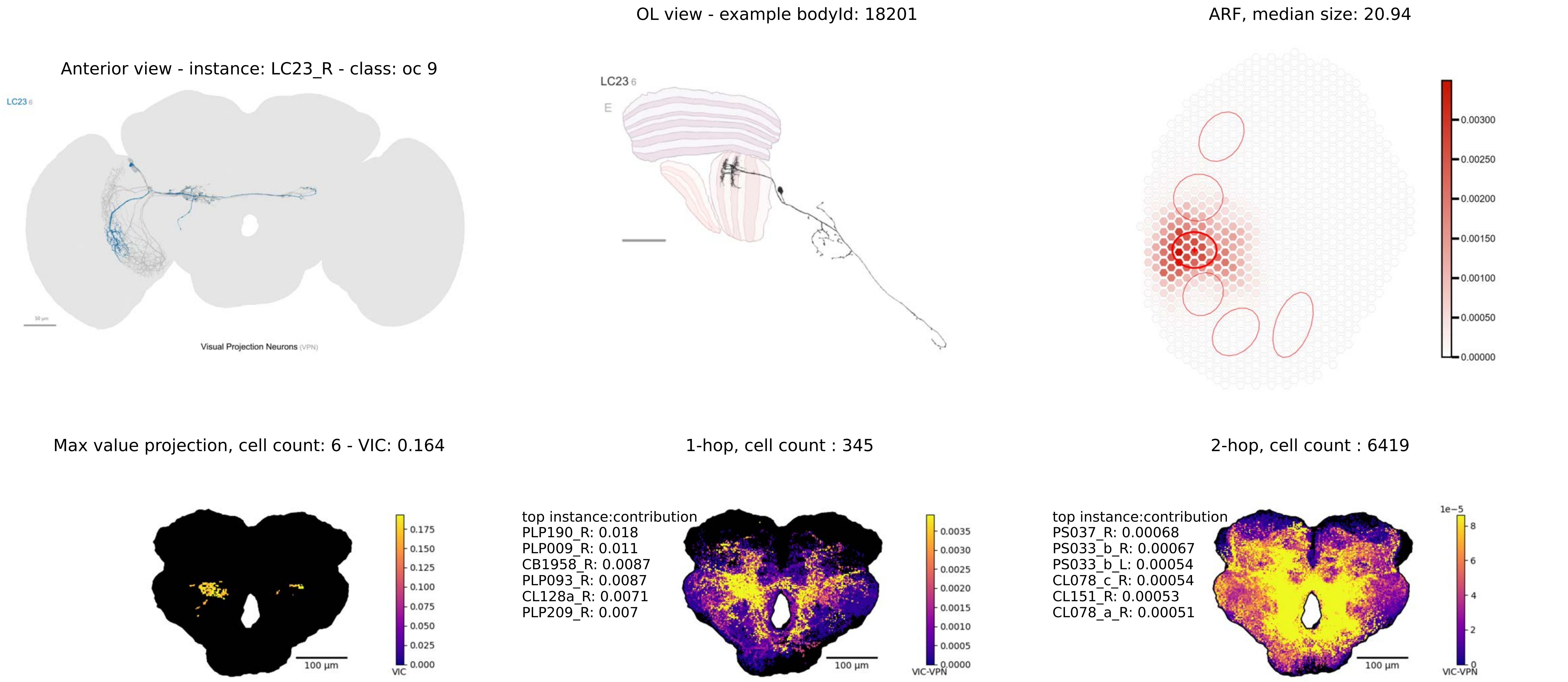

OL view - example bodyId: 18201
ARF, median size: 20.94
Anterior view - instance: LC23_R - class: oc 9
LC23 6
Visual Projection Neurons (VPN)
LC23 6
E
0.00300
0.00250
0.00200
0.00150
0.00100
0.00050
0.00000
Max value projection, cell count: 6 - VIC: 0.164
1-hop, cell count : 345
2-hop, cell count : 6419
0.175
0.150
0.125
0.100
0.075
0.050
0.025
0.000
VIC
100 µm
top instance:contribution
PLP190_R: 0.018
PLP009_R: 0.011
CB1958_R: 0.0087
PLP093_R: 0.0087
CL128a_R: 0.0071
PLP209_R: 0.007
100 µm
0.0035
0.0030
0.0025
0.0020
0.0015
0.0010
0.0005
0.0000
VIC-VPN
top instance:contribution
PS037_R: 0.00068
PS033_b_R: 0.00067
PS033_b_L: 0.00054
CL078_c_R: 0.00054
CL151_R: 0.00053
CL078_a_R: 0.00051
100 µm
1e−5
8
6
4
2
0
VIC-VPN

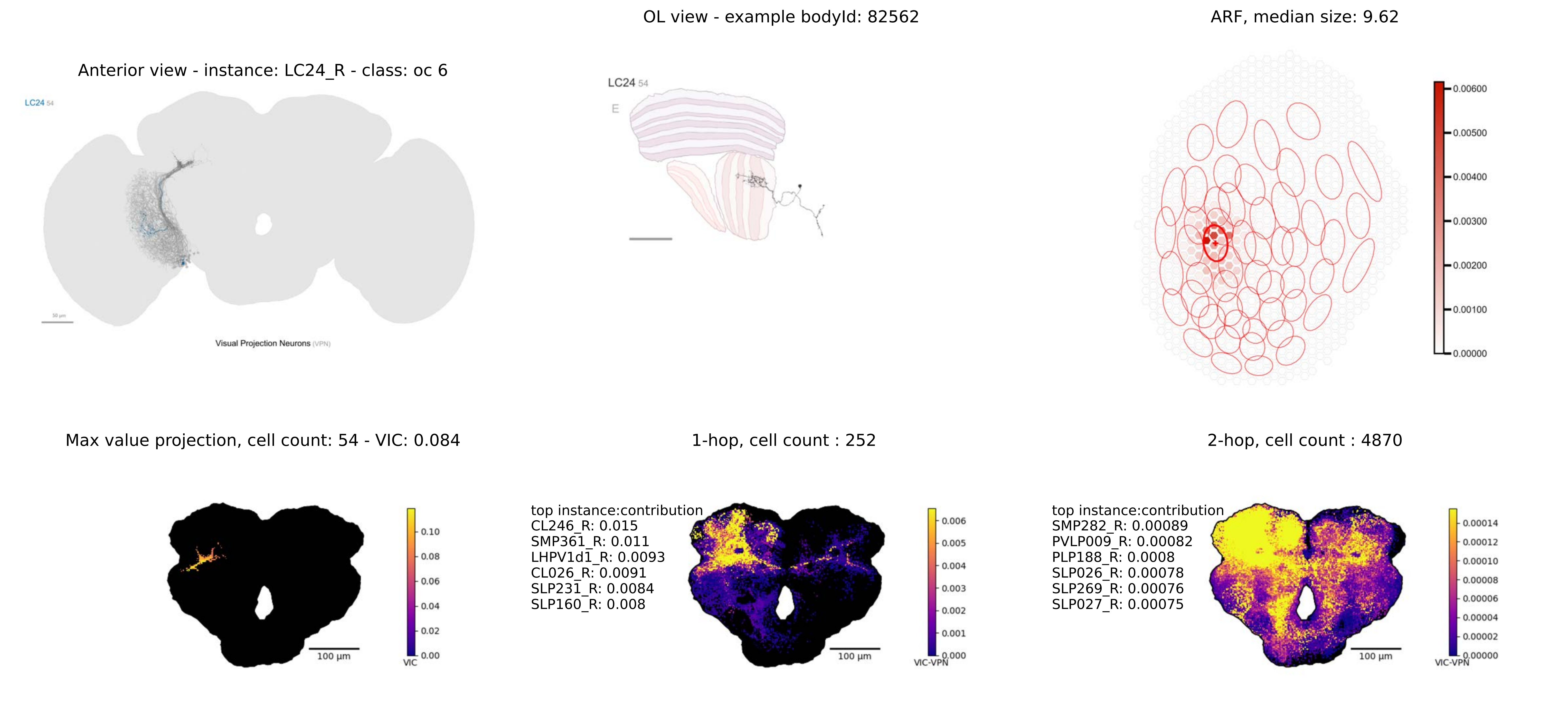

OL view - example bodyId: 82562
ARF, median size: 9.62
Anterior view - instance: LC24_R - class: oc 6
LC24 54
Visual Projection Neurons (VPN)
E
Max value projection, cell count: 54 - VIC: 0.084
1-hop, cell count : 252
2-hop, cell count : 4870
top instance:contribution
CL246_R: 0.015
SMP361_R: 0.011
LHPV1d1_R: 0.0093
CL026_R: 0.0091
SLP231_R: 0.0084
SLP160_R: 0.008
top instance:contribution
SMP282_R: 0.00089
PVLP009_R: 0.00082
PLP188_R: 0.0008
SLP026_R: 0.00078
SLP269_R: 0.00076
SLP027_R: 0.00075
100 µm
VIC
VIC-VPN

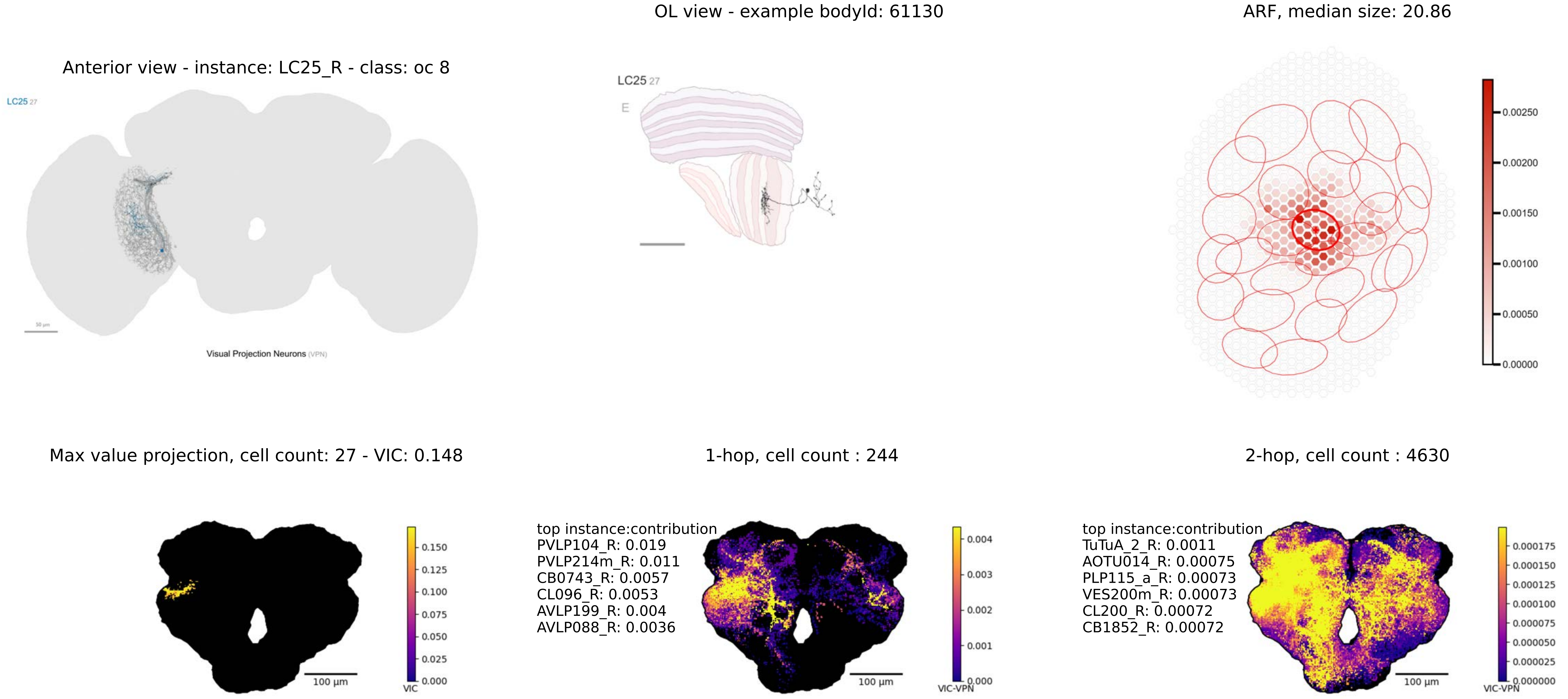

OL view - example bodyId: 61130
ARF, median size: 20.86
Anterior view - instance: LC25_R - class: oc 8
LC25 27
Visual Projection Neurons (VPN)
LC25 27
E
0.00250
0.00200
0.00150
0.00100
0.00050
0.00000
Max value projection, cell count: 27 - VIC: 0.148
1-hop, cell count : 244
2-hop, cell count : 4630
100 µm
VIC
0.150
0.125
0.100
0.075
0.050
0.025
0.000
top instance:contribution
PVLP104_R: 0.019
PVLP214m_R: 0.011
CB0743_R: 0.0057
CL096_R: 0.0053
AVLP199_R: 0.004
AVLP088_R: 0.0036
100 µm
VIC-VPN
0.004
0.003
0.002
0.001
0.000
top instance:contribution
TuTuA_2_R: 0.0011
AOTU014_R: 0.00075
PLP115_a_R: 0.00073
VES200m_R: 0.00073
CL200_R: 0.00072
CB1852_R: 0.00072
100 µm
VIC-VPN
0.000175
0.000150
0.000125
0.000100
0.000075
0.000050
0.000025
0.000000

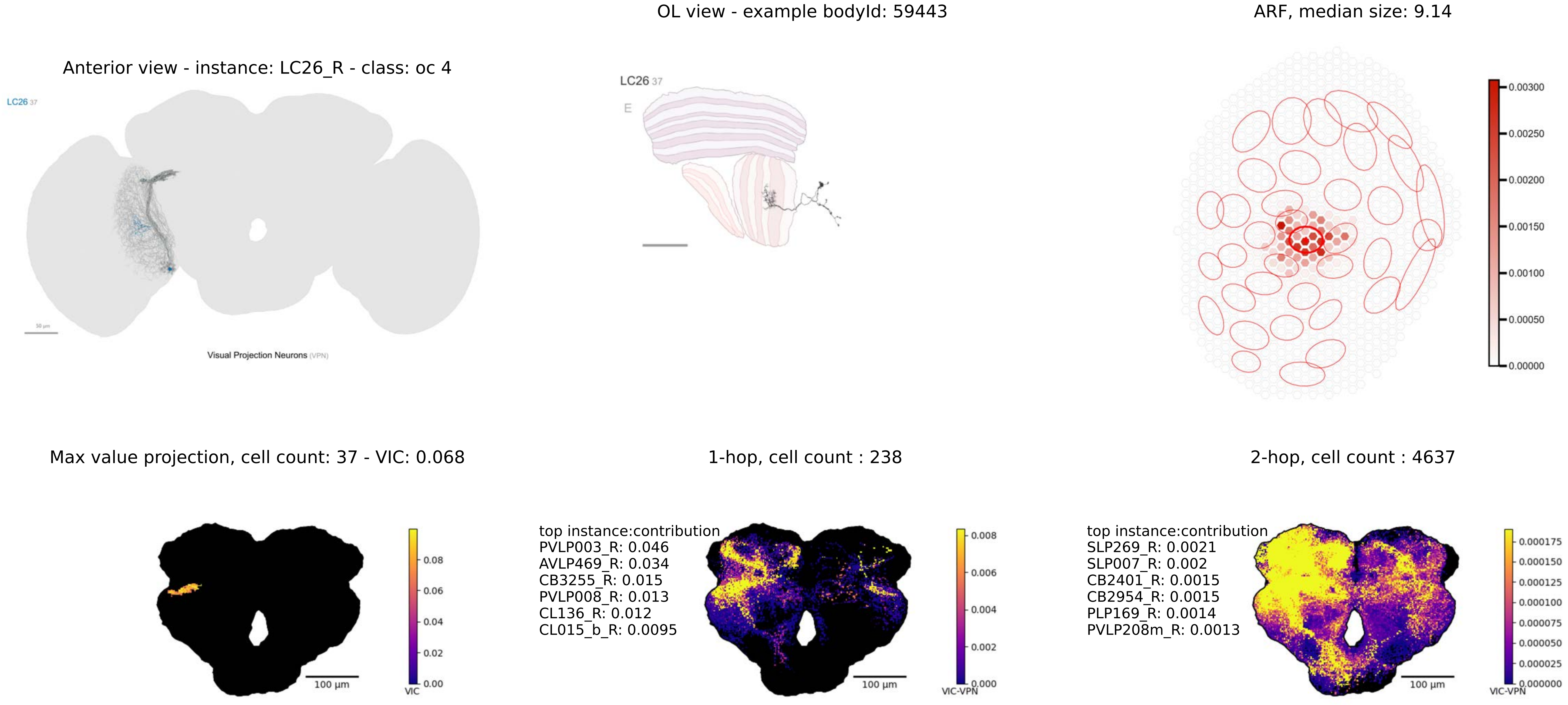
OL view - example bodyId: 59443
ARF, median size: 9.14
Anterior view - instance: LC26_R - class: oc 4
LC26 37
Visual Projection Neurons (VPN)
LC26 37
E
0.00300
0.00250
0.00200
0.00150
0.00100
0.00050
0.00000
Max value projection, cell count: 37 - VIC: 0.068
1-hop, cell count : 238
2-hop, cell count : 4637
0.08
0.06
0.04
0.02
0.00
VIC
100 µm
top instance:contribution
PVLP003_R: 0.046
AVLP469_R: 0.034
CB3255_R: 0.015
PVLP008_R: 0.013
CL136_R: 0.012
CL015_b_R: 0.0095
0.008
0.006
0.004
0.002
0.000
VIC-VPN
100 µm
top instance:contribution
SLP269_R: 0.0021
SLP007_R: 0.002
CB2401_R: 0.0015
CB2954_R: 0.0015
PLP169_R: 0.0014
PVLP208m_R: 0.0013
0.000175
0.000150
0.000125
0.000100
0.000075
0.000050
0.000025
0.000000
VIC-VPN
100 µm

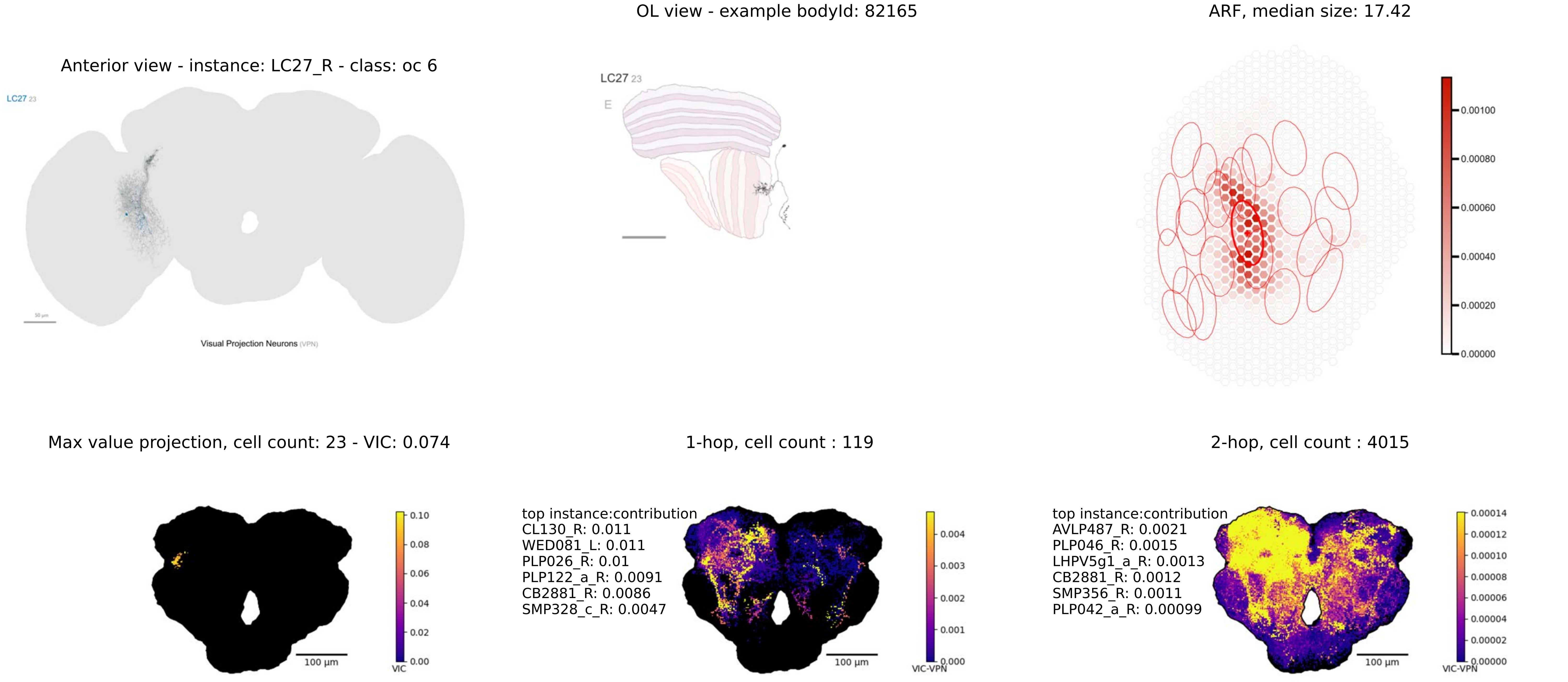

Anterior view - instance: LC27_R - class: oc 6
LC27 23
Visual Projection Neurons (VPN)
OL view - example bodyId: 82165
LC27 23
E
ARF, median size: 17.42
0.00100
0.00080
0.00060
0.00040
0.00020
0.00000
Max value projection, cell count: 23 - VIC: 0.074
0.10
0.08
0.06
0.04
0.02
0.00
VIC
100 µm
1-hop, cell count : 119
top instance:contribution
CL130_R: 0.011
WED081_L: 0.011
PLP026_R: 0.01
PLP122_a_R: 0.0091
CB2881_R: 0.0086
SMP328_c_R: 0.0047
0.004
0.003
0.002
0.001
0.000
VIC-VPN
100 µm
2-hop, cell count : 4015
top instance:contribution
AVLP487_R: 0.0021
PLP046_R: 0.0015
LHPV5g1_a_R: 0.0013
CB2881_R: 0.0012
SMP356_R: 0.0011
PLP042_a_R: 0.00099
0.00014
0.00012
0.00010
0.00008
0.00006
0.00004
0.00002
0.00000
VIC-VPN
100 µm

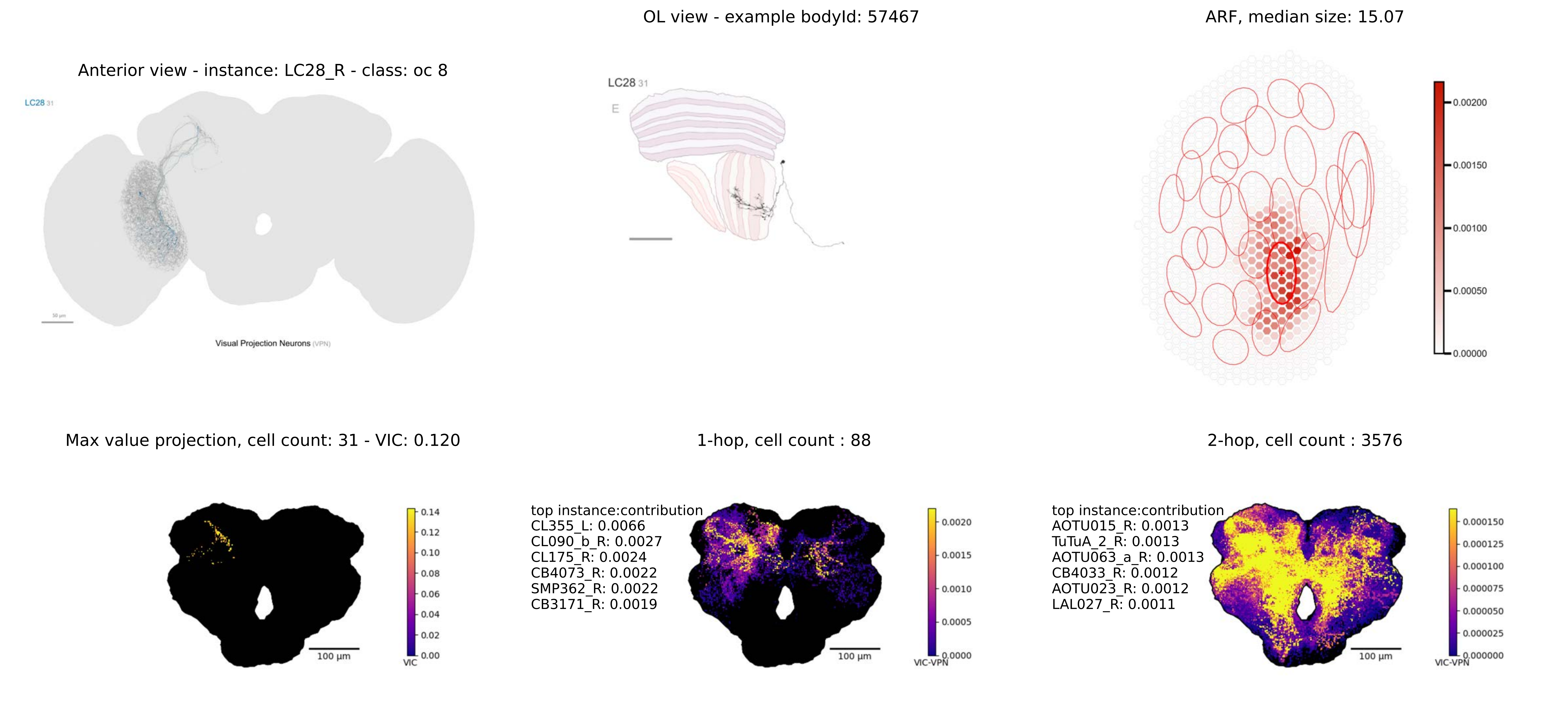
Anterior view - instance: LC28_R - class: oc 8
LC28 31
Visual Projection Neurons (VPN)
OL view - example bodyId: 57467
LC28 31
E
ARF, median size: 15.07
0.00200
0.00150
0.00100
0.00050
0.00000
Max value projection, cell count: 31 - VIC: 0.120
100 µm
0.14
0.12
0.10
0.08
0.06
0.04
0.02
0.00
VIC
1-hop, cell count : 88
top instance:contribution
CL355_L: 0.0066
CL090_b_R: 0.0027
CL175_R: 0.0024
CB4073_R: 0.0022
SMP362_R: 0.0022
CB3171_R: 0.0019
100 µm
0.0020
0.0015
0.0010
0.0005
0.0000
VIC-VPN
2-hop, cell count : 3576
top instance:contribution
AOTU015_R: 0.0013
TuTuA_2_R: 0.0013
AOTU063_a_R: 0.0013
CB4033_R: 0.0012
AOTU023_R: 0.0012
LAL027_R: 0.0011
100 µm
0.000150
0.000125
0.000100
0.000075
0.000050
0.000025
0.000000
VIC-VPN

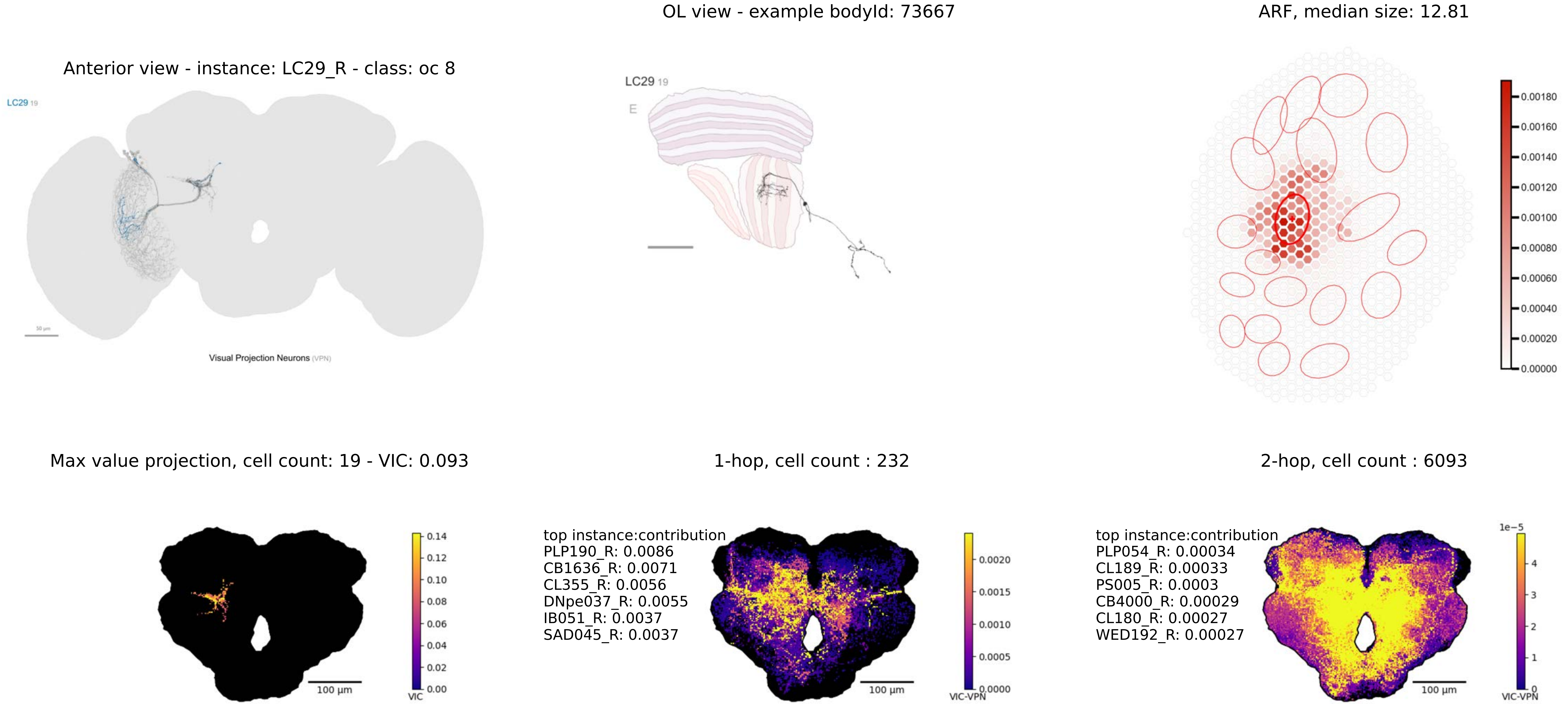
Anterior view - instance: LC29_R - class: oc 8
LC29 19
50 µm
Visual Projection Neurons (VPN)
OL view - example bodyId: 73667
LC29 19
E
ARF, median size: 12.81
0.00180
0.00160
0.00140
0.00120
0.00100
0.00080
0.00060
0.00040
0.00020
0.00000
Max value projection, cell count: 19 - VIC: 0.093
100 µm
0.14
0.12
0.10
0.08
0.06
0.04
0.02
0.00
VIC
1-hop, cell count : 232
top instance:contribution
PLP190_R: 0.0086
CB1636_R: 0.0071
CL355_R: 0.0056
DNpe037_R: 0.0055
IB051_R: 0.0037
SAD045_R: 0.0037
100 µm
0.0020
0.0015
0.0010
0.0005
0.0000
VIC-VPN
2-hop, cell count : 6093
top instance:contribution
PLP054_R: 0.00034
CL189_R: 0.00033
PS005_R: 0.0003
CB4000_R: 0.00029
CL180_R: 0.00027
WED192_R: 0.00027
100 µm
1e−5
4
3
2
1
0
VIC-VPN

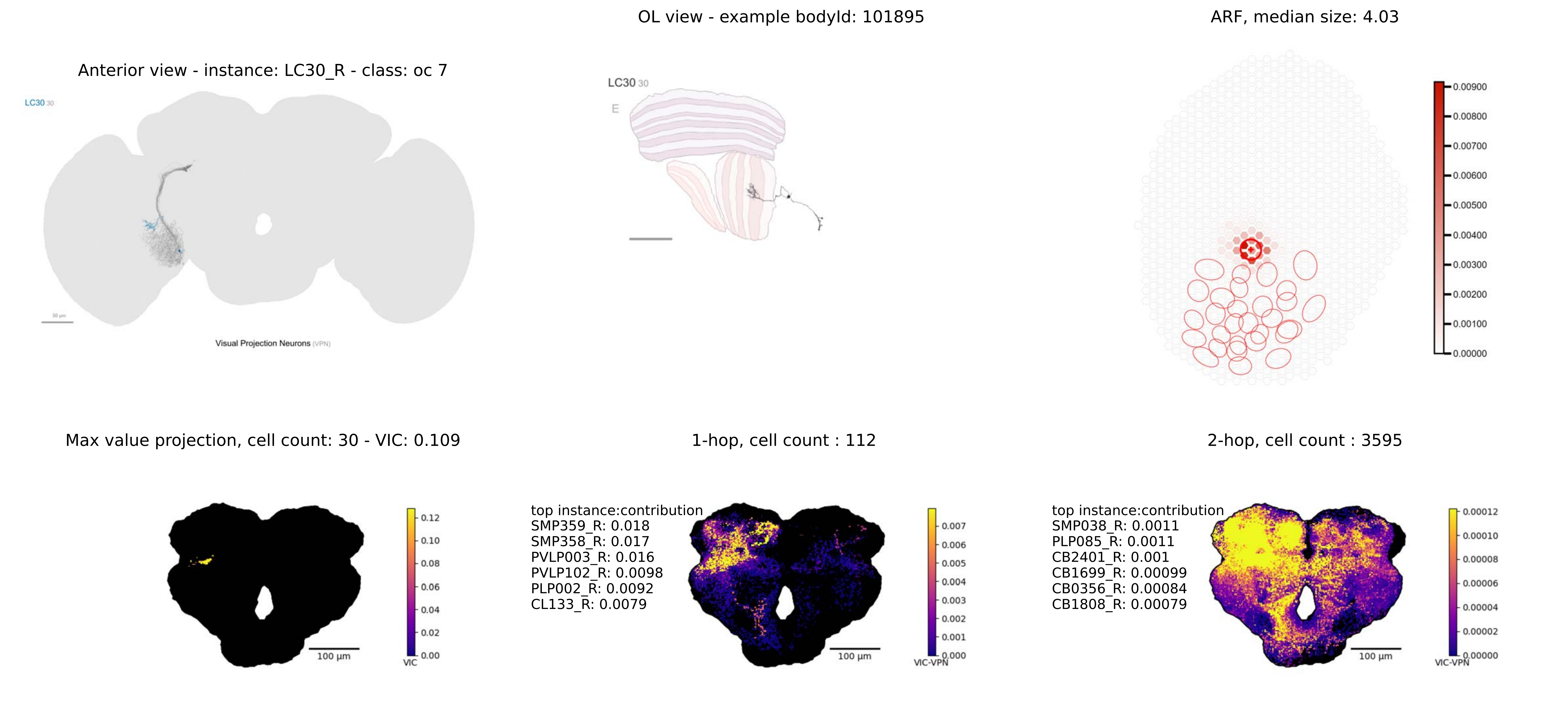

OL view - example bodyId: 101895
ARF, median size: 4.03
Anterior view - instance: LC30_R - class: oc 7
LC30 30
Visual Projection Neurons (VPN)
LC30 30
E
Max value projection, cell count: 30 - VIC: 0.109
1-hop, cell count : 112
2-hop, cell count : 3595
top instance:contribution
SMP359_R: 0.018
SMP358_R: 0.017
PVLP003_R: 0.016
PVLP102_R: 0.0098
PLP002_R: 0.0092
CL133_R: 0.0079
top instance:contribution
SMP038_R: 0.0011
PLP085_R: 0.0011
CB2401_R: 0.001
CB1699_R: 0.00099
CB0356_R: 0.00084
CB1808_R: 0.00079
100 µm
VIC
VIC-VPN

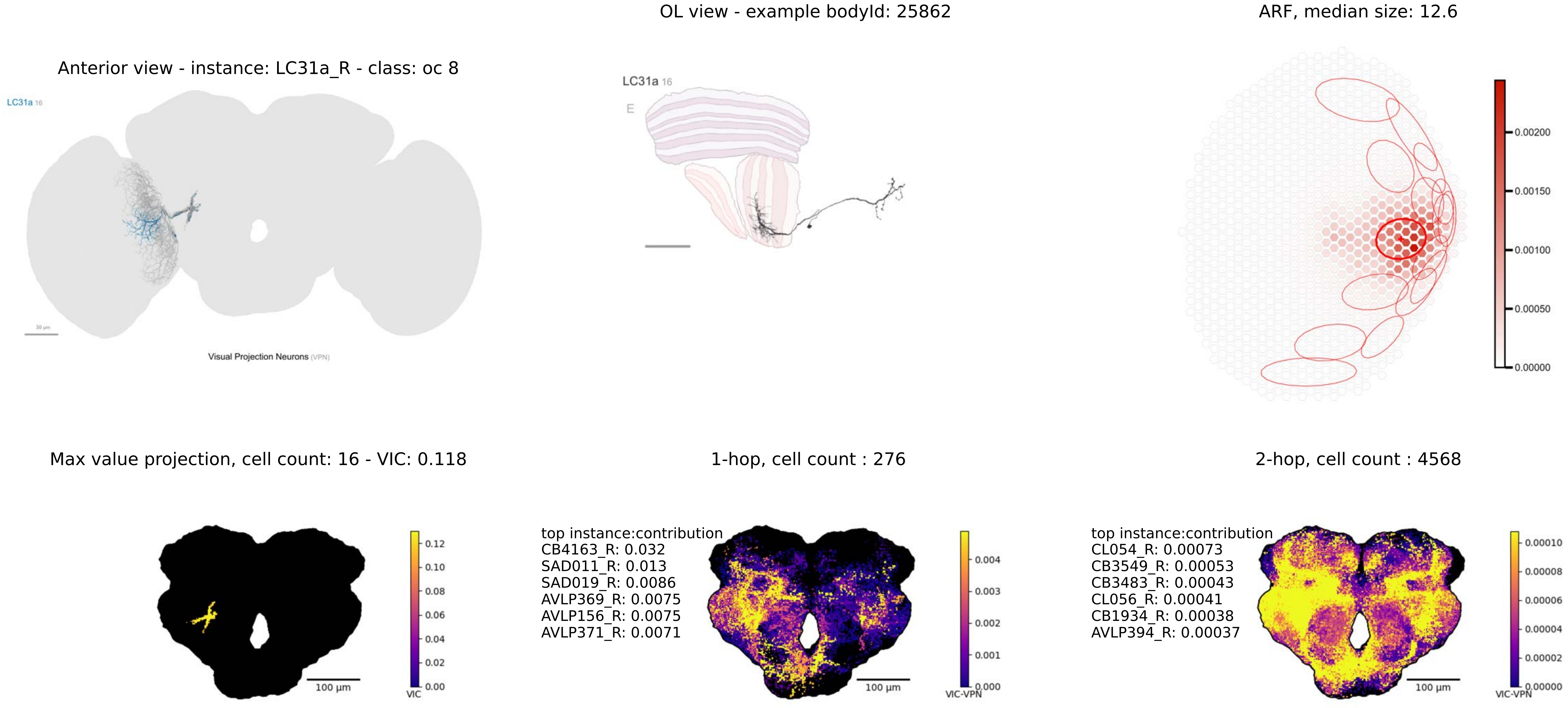
OL view - example bodyId: 25862
ARF, median size: 12.6
Anterior view - instance: LC31a_R - class: oc 8
LC31a 16
Visual Projection Neurons (VPN)
LC31a 16
E
0.00200
0.00150
0.00100
0.00050
0.00000
Max value projection, cell count: 16 - VIC: 0.118
1-hop, cell count : 276
2-hop, cell count : 4568
0.12
0.10
0.08
0.06
0.04
0.02
0.00
VIC
100 µm
top instance:contribution
CB4163_R: 0.032
SAD011_R: 0.013
SAD019_R: 0.0086
AVLP369_R: 0.0075
AVLP156_R: 0.0075
AVLP371_R: 0.0071
0.004
0.003
0.002
0.001
0.000
VIC-VPN
100 µm
top instance:contribution
CL054_R: 0.00073
CB3549_R: 0.00053
CB3483_R: 0.00043
CL056_R: 0.00041
CB1934_R: 0.00038
AVLP394_R: 0.00037
0.00010
0.00008
0.00006
0.00004
0.00002
0.00000
VIC-VPN
100 µm

OL view - example bodyId: 11812

ARF, median size: 41.27

Anterior view - instance: LC31b_R - class: oc 8

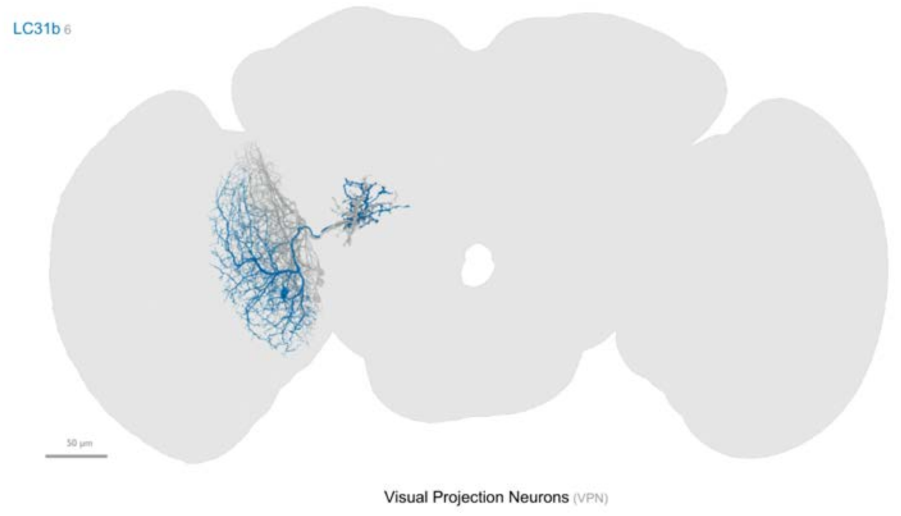


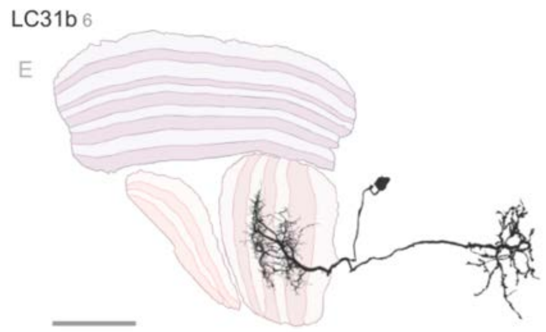


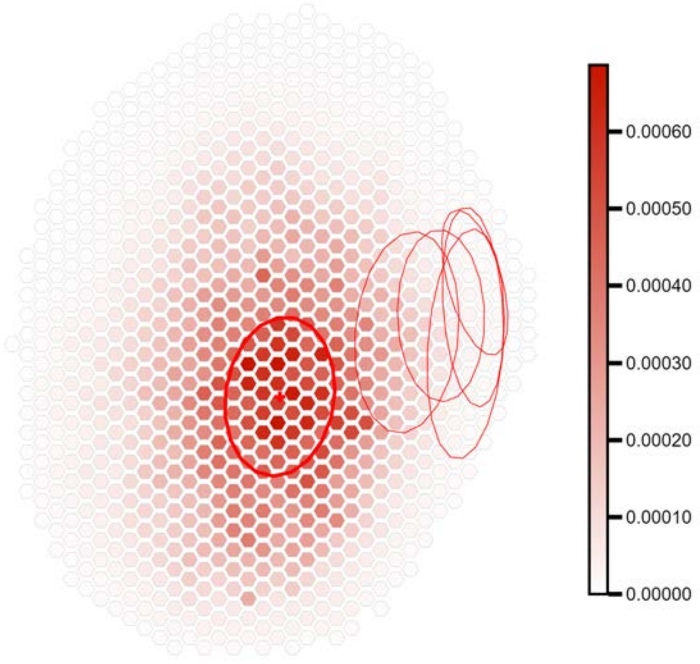


Max value projection, cell count: 6 - VIC: 0.114

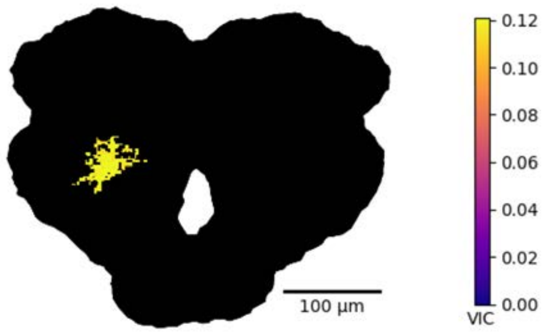


1-hop, cell count : 368

top instance:contribution
PVLP060_R: 0.019
CB3483_R: 0.018
AVLP734m_R: 0.016
CB3549_R: 0.013
CB3335_R: 0.012
DNp34_L: 0.01

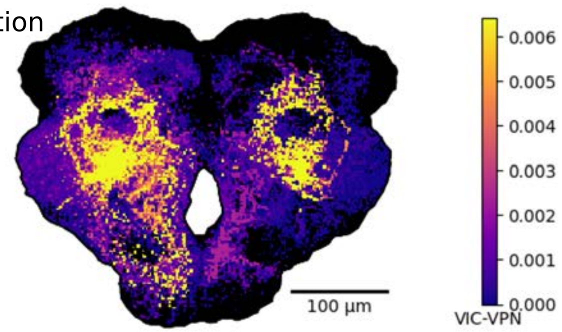


2-hop, cell count : 4974

top instance:contribution
P1_6b_R: 0.0018
CB1883_R: 0.0012
PVLP202m_R: 0.0011
ICL008m_R: 0.00086
AVLP733m_R: 0.00073
PVLP200m_a_R: 0.00065

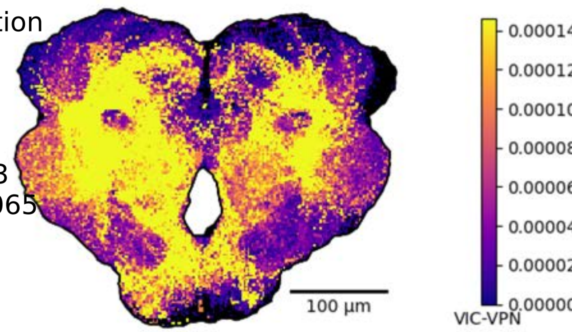

OL view - example bodyId: 14002

ARF, median size: 34.74

Anterior view - instance: LC33_R - class: oc 6

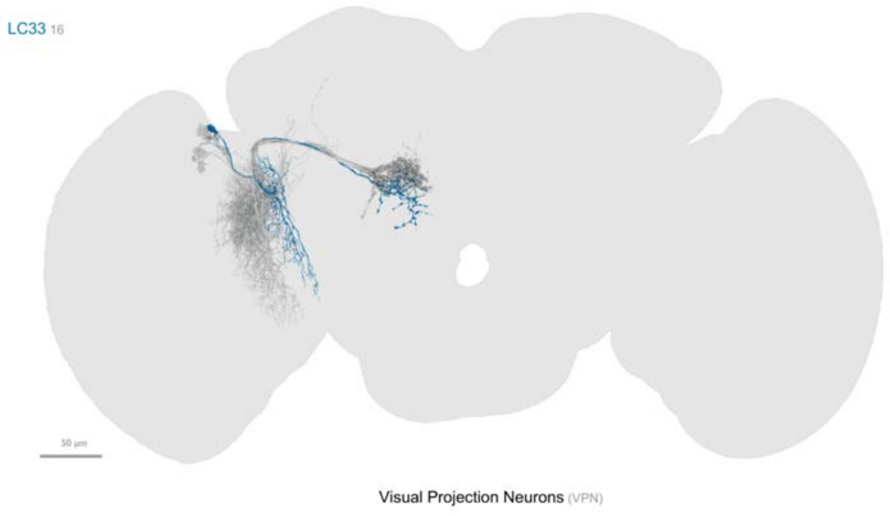


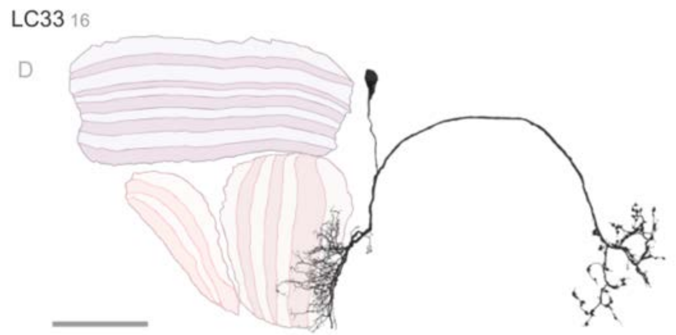


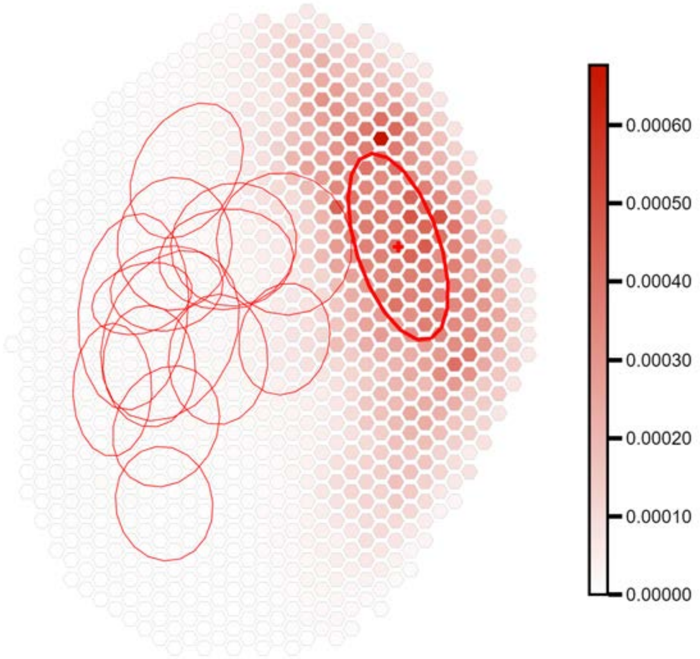


Max value projection, cell count: 16 - VIC: 0.061

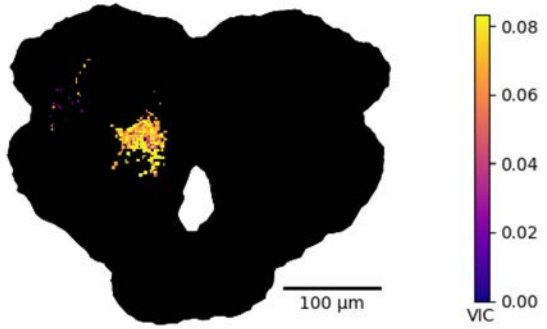


1-hop, cell count : 335

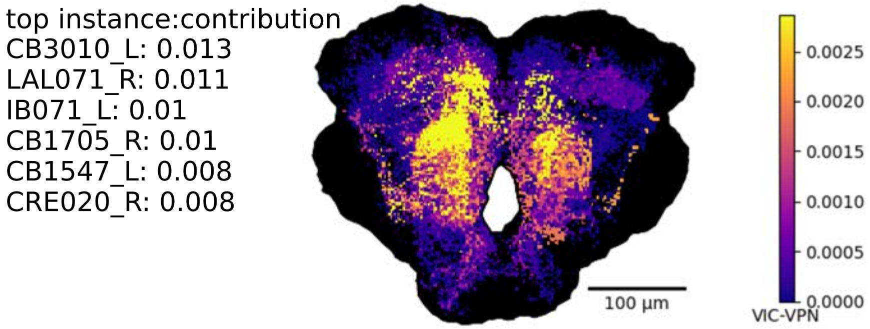


2-hop, cell count : 4412

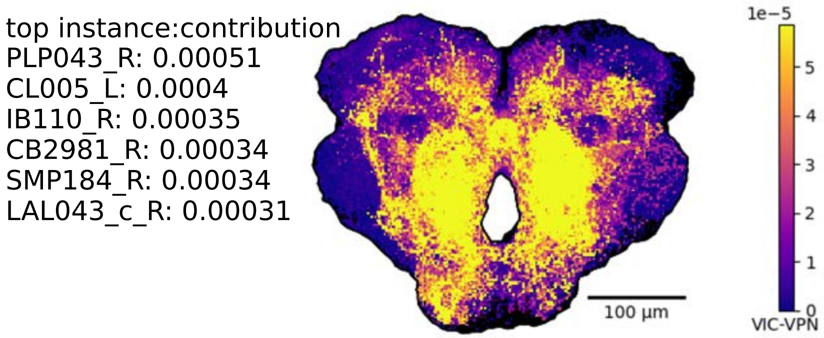

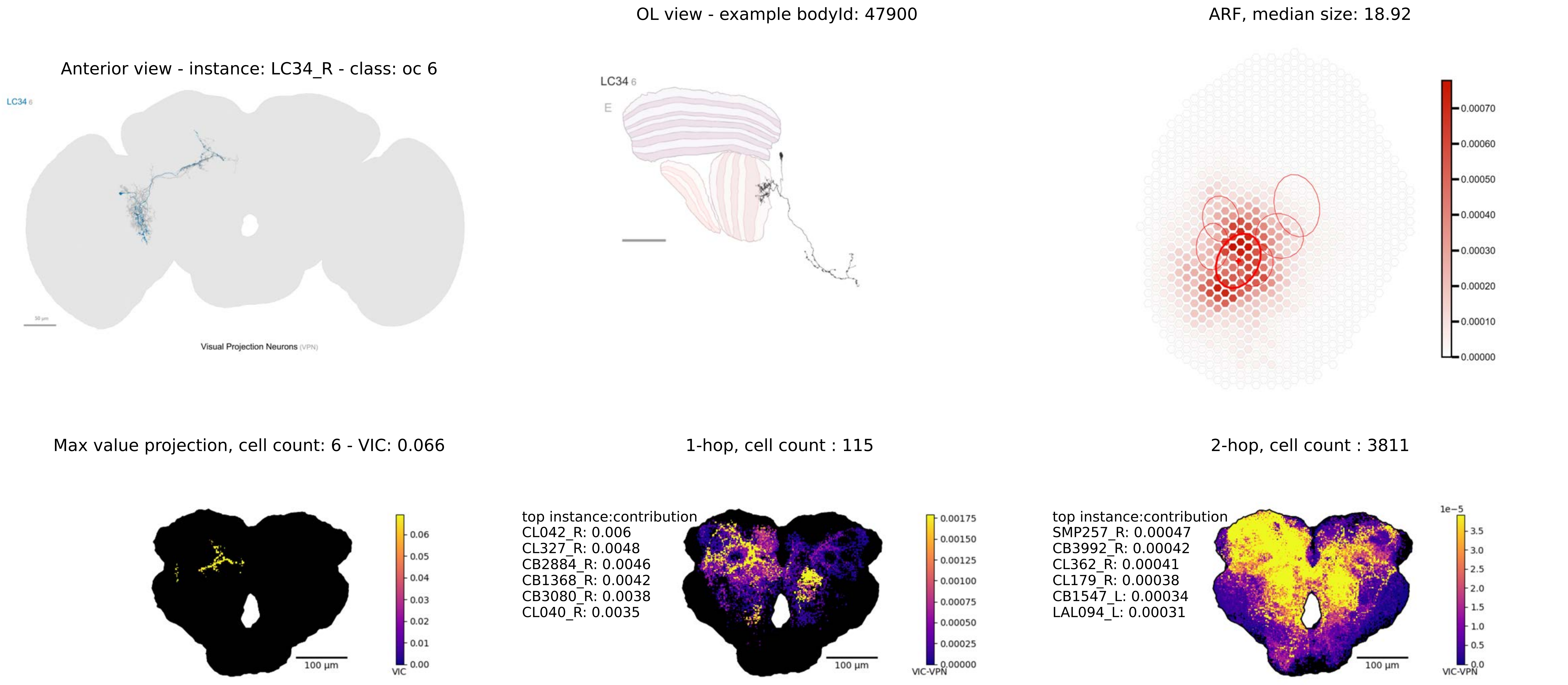

Anterior view - instance: LC34_R - class: oc 6
LC34 6
Visual Projection Neurons (VPN)
OL view - example bodyId: 47900
LC34 6
E
ARF, median size: 18.92
0.00070
0.00060
0.00050
0.00040
0.00030
0.00020
0.00010
0.00000
Max value projection, cell count: 6 - VIC: 0.066
100 µm
0.06
0.05
0.04
0.03
0.02
0.01
0.00
VIC
1-hop, cell count : 115
top instance:contribution
CL042_R: 0.006
CL327_R: 0.0048
CB2884_R: 0.0046
CB1368_R: 0.0042
CB3080_R: 0.0038
CL040_R: 0.0035
100 µm
0.00175
0.00150
0.00125
0.00100
0.00075
0.00050
0.00025
0.00000
VIC-VPN
2-hop, cell count : 3811
top instance:contribution
SMP257_R: 0.00047
CB3992_R: 0.00042
CL362_R: 0.00041
CL179_R: 0.00038
CB1547_L: 0.00034
LAL094_L: 0.00031
100 µm
1e−5
3.5
3.0
2.5
2.0
1.5
1.0
0.5
0.0
VIC-VPN

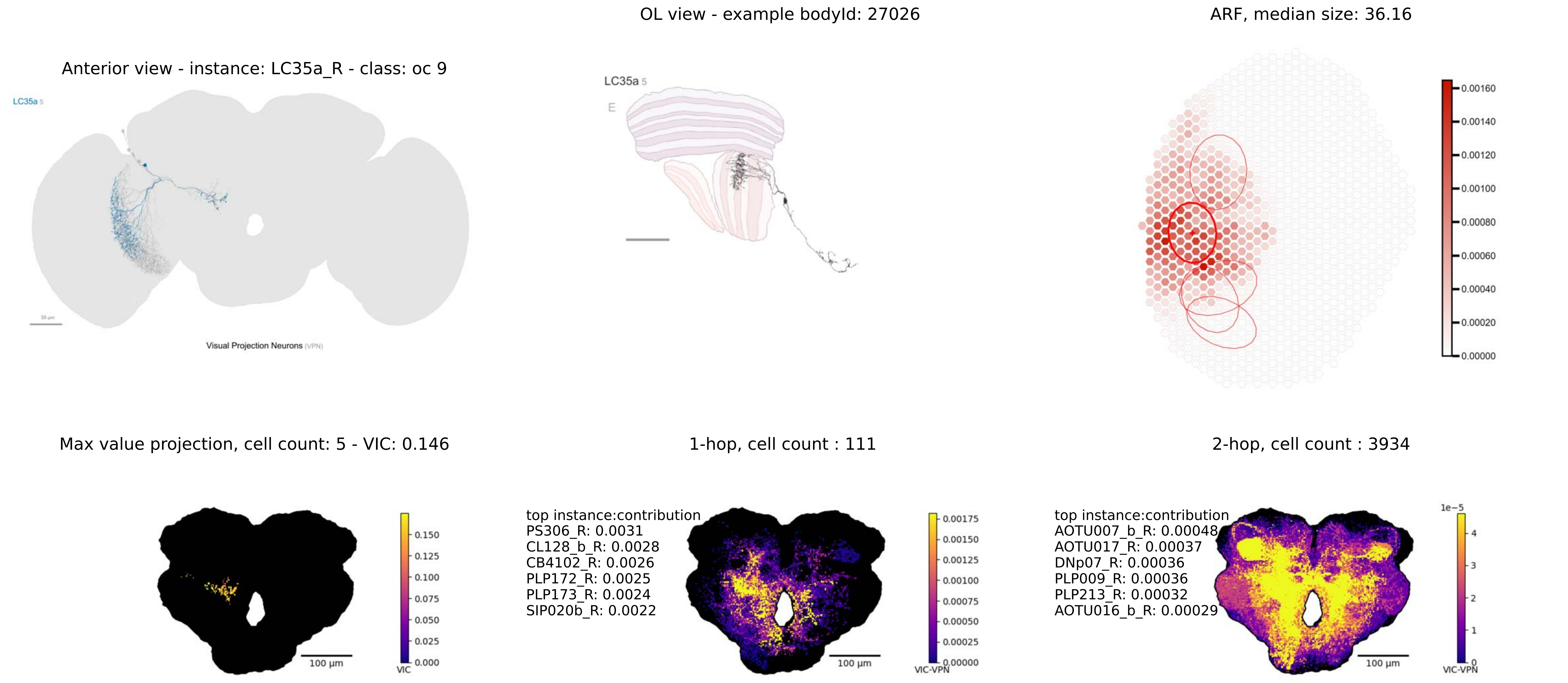
OL view - example bodyId: 27026
ARF, median size: 36.16
Anterior view - instance: LC35a_R - class: oc 9
LC35a 5
Visual Projection Neurons (VPN)
LC35a 5
E
0.00160
0.00140
0.00120
0.00100
0.00080
0.00060
0.00040
0.00020
0.00000
Max value projection, cell count: 5 - VIC: 0.146
1-hop, cell count : 111
2-hop, cell count : 3934
0.150
0.125
0.100
0.075
0.050
0.025
0.000
VIC
100 µm
top instance:contribution
PS306_R: 0.0031
CL128_b_R: 0.0028
CB4102_R: 0.0026
PLP172_R: 0.0025
PLP173_R: 0.0024
SIP020b_R: 0.0022
100 µm
0.00175
0.00150
0.00125
0.00100
0.00075
0.00050
0.00025
0.00000
VIC-VPN
top instance:contribution
AOTU007_b_R: 0.00048
AOTU017_R: 0.00037
DNp07_R: 0.00036
PLP009_R: 0.00036
PLP213_R: 0.00032
AOTU016_b_R: 0.00029
100 µm
1e−5
4
3
2
1
0
VIC-VPN

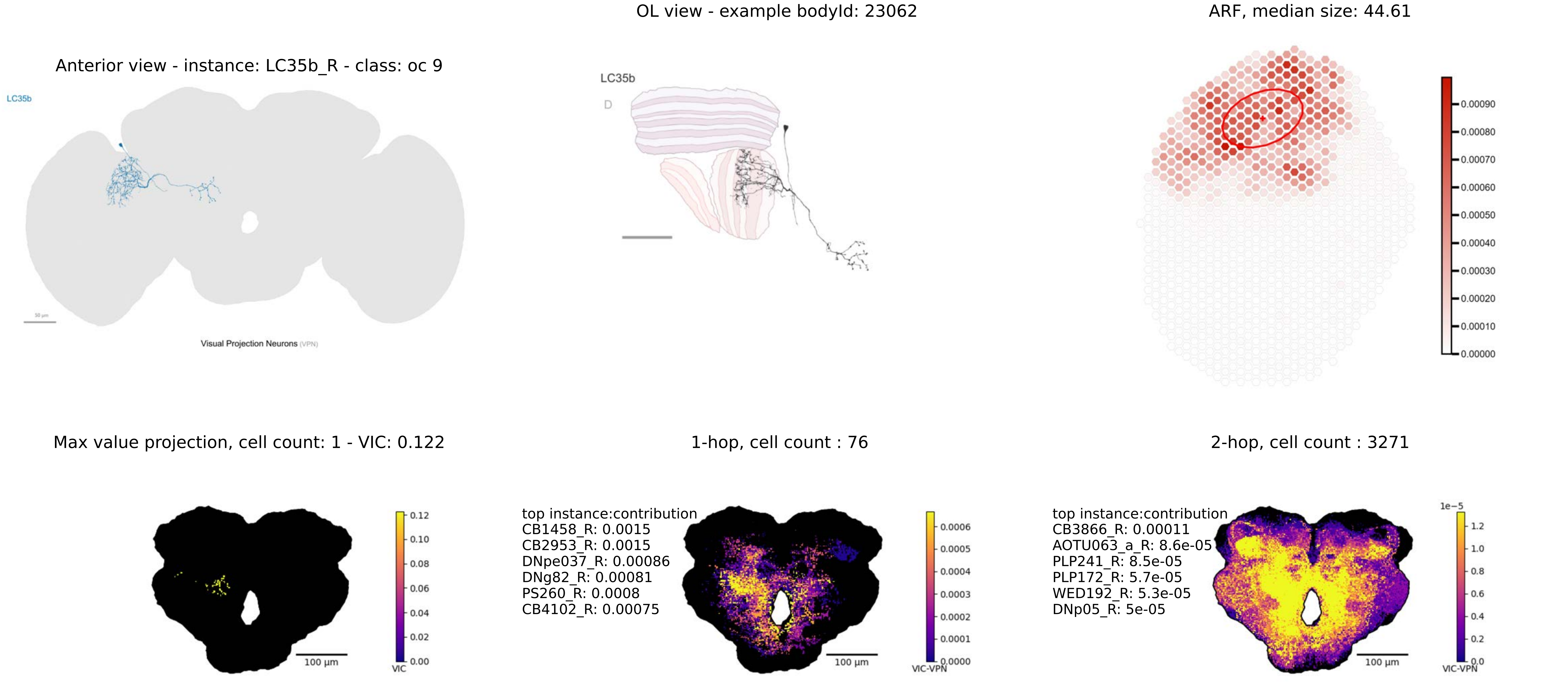
OL view - example bodyId: 23062
ARF, median size: 44.61
Anterior view - instance: LC35b_R - class: oc 9
LC35b
Visual Projection Neurons (VPN)
LC35b
D
0.00090
0.00080
0.00070
0.00060
0.00050
0.00040
0.00030
0.00020
0.00010
0.00000
Max value projection, cell count: 1 - VIC: 0.122
1-hop, cell count : 76
2-hop, cell count : 3271
0.12
0.10
0.08
0.06
0.04
0.02
0.00
VIC
100 µm
top instance:contribution
CB1458_R: 0.0015
CB2953_R: 0.0015
DNpe037_R: 0.00086
DNg82_R: 0.00081
PS260_R: 0.0008
CB4102_R: 0.00075
0.0006
0.0005
0.0004
0.0003
0.0002
0.0001
0.0000
VIC-VPN
100 µm
top instance:contribution
CB3866_R: 0.00011
AOTU063_a_R: 8.6e-05
PLP241_R: 8.5e-05
PLP172_R: 5.7e-05
WED192_R: 5.3e-05
DNp05_R: 5e-05
1e−5
1.2
1.0
0.8
0.6
0.4
0.2
0.0
VIC-VPN
100 µm

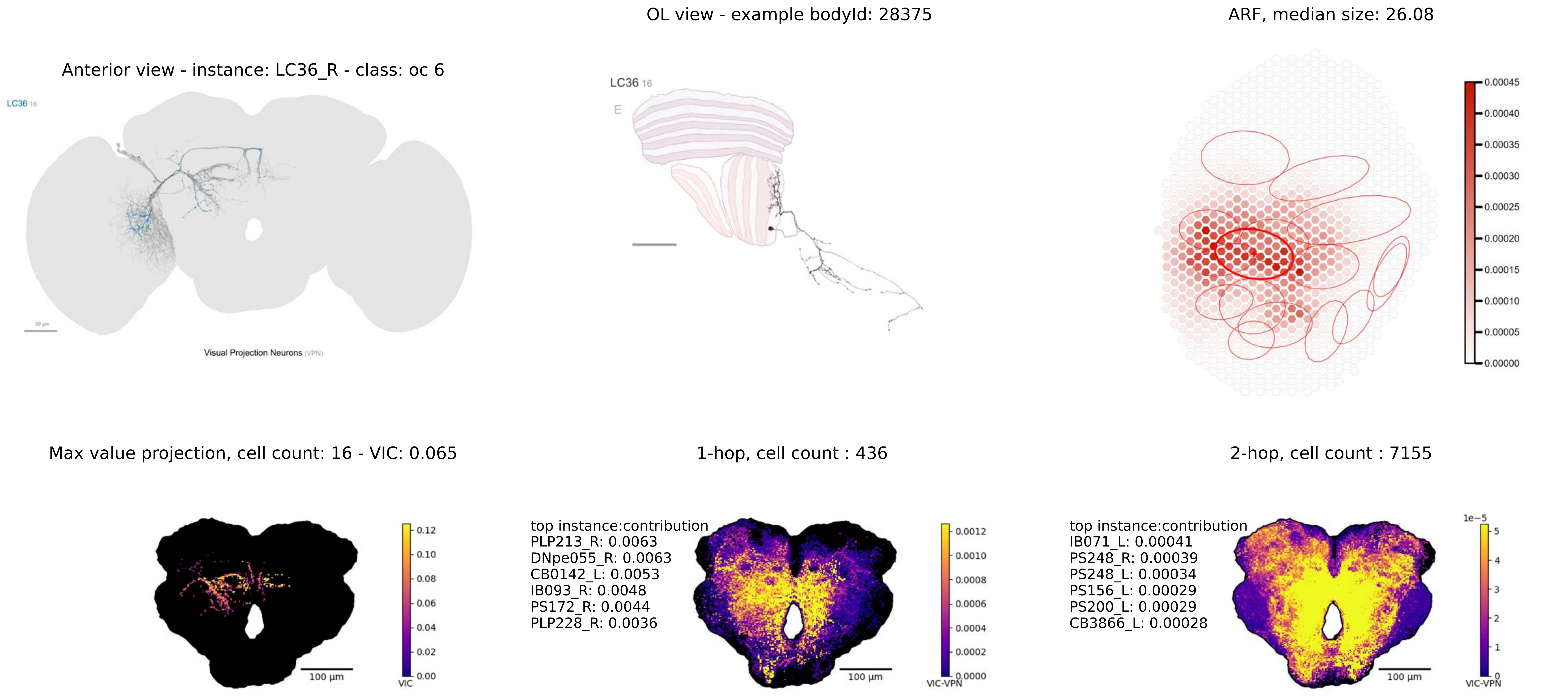

OL view - example bodyId: 28375
ARF, median size: 26.08
Anterior view - instance: LC36_R - class: oc 6
LC36 16
Visual Projection Neurons (VPN)
LC36 16
E
0.00045
0.00040
0.00035
0.00030
0.00025
0.00020
0.00015
0.00010
0.00005
0.00000
Max value projection, cell count: 16 - VIC: 0.065
1-hop, cell count : 436
2-hop, cell count : 7155
0.12
0.10
0.08
0.06
0.04
0.02
0.00
VIC
100 µm
top instance:contribution
PLP213_R: 0.0063
DNpe055_R: 0.0063
CB0142_L: 0.0053
IB093_R: 0.0048
PS172_R: 0.0044
PLP228_R: 0.0036
0.0012
0.0010
0.0008
0.0006
0.0004
0.0002
0.0000
VIC-VPN
100 µm
top instance:contribution
IB071_L: 0.00041
PS248_R: 0.00039
PS248_L: 0.00034
PS156_L: 0.00029
PS200_L: 0.00029
CB3866_L: 0.00028
1e−5
5
4
3
2
1
0
VIC-VPN
100 µm

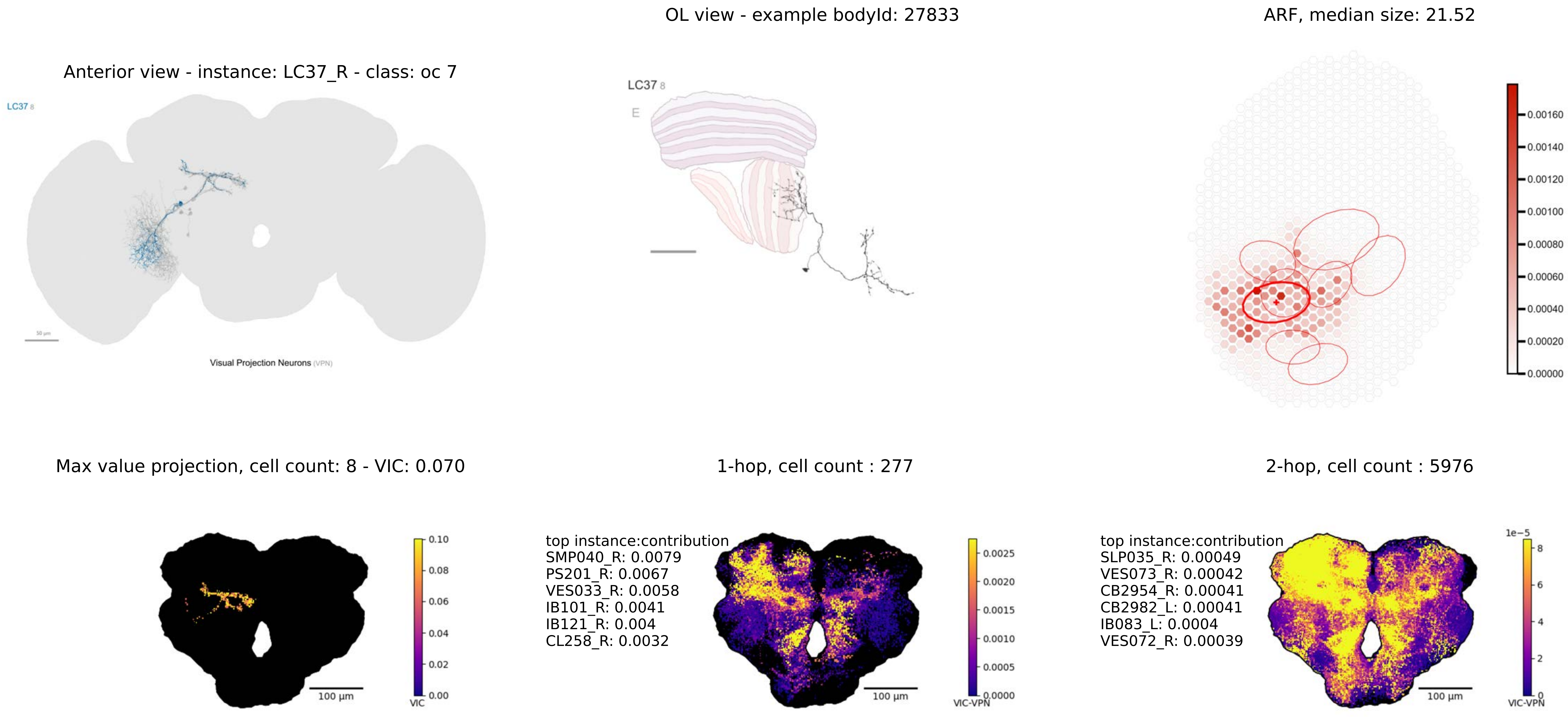

Anterior view - instance: LC37_R - class: oc 7
LC37 8
50 µm
Visual Projection Neurons (VPN)
OL view - example bodyId: 27833
LC37 8
E
ARF, median size: 21.52
0.00160
0.00140
0.00120
0.00100
0.00080
0.00060
0.00040
0.00020
0.00000
Max value projection, cell count: 8 - VIC: 0.070
100 µm
0.10
0.08
0.06
0.04
0.02
0.00
VIC
1-hop, cell count : 277
top instance:contribution
SMP040_R: 0.0079
PS201_R: 0.0067
VES033_R: 0.0058
IB101_R: 0.0041
IB121_R: 0.004
CL258_R: 0.0032
100 µm
0.0025
0.0020
0.0015
0.0010
0.0005
0.0000
VIC-VPN
2-hop, cell count : 5976
top instance:contribution
SLP035_R: 0.00049
VES073_R: 0.00042
CB2954_R: 0.00041
CB2982_L: 0.00041
IB083_L: 0.0004
VES072_R: 0.00039
100 µm
1e−5
8
6
4
2
0
VIC-VPN

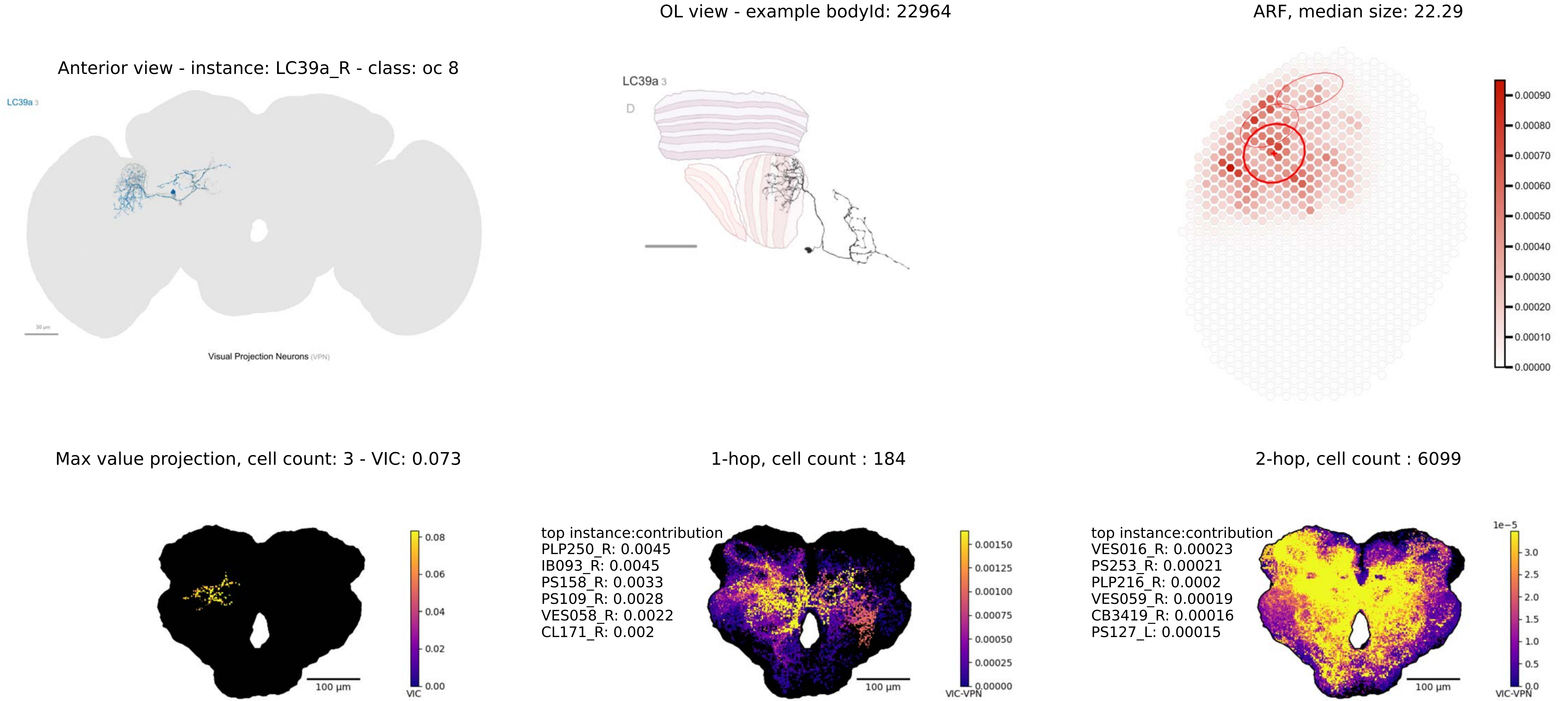

OL view - example bodyId: 22964
ARF, median size: 22.29
Anterior view - instance: LC39a_R - class: oc 8
LC39a 3
Visual Projection Neurons (VPN)
LC39a 3
D
0.00090
0.00080
0.00070
0.00060
0.00050
0.00040
0.00030
0.00020
0.00010
0.00000
Max value projection, cell count: 3 - VIC: 0.073
1-hop, cell count : 184
2-hop, cell count : 6099
0.08
0.06
0.04
0.02
0.00
VIC
100 µm
top instance:contribution
PLP250_R: 0.0045
IB093_R: 0.0045
PS158_R: 0.0033
PS109_R: 0.0028
VES058_R: 0.0022
CL171_R: 0.002
0.00150
0.00125
0.00100
0.00075
0.00050
0.00025
0.00000
VIC-VPN
100 µm
top instance:contribution
VES016_R: 0.00023
PS253_R: 0.00021
PLP216_R: 0.0002
VES059_R: 0.00019
CB3419_R: 0.00016
PS127_L: 0.00015
1e−5
3.0
2.5
2.0
1.5
1.0
0.5
0.0
VIC-VPN
100 µm

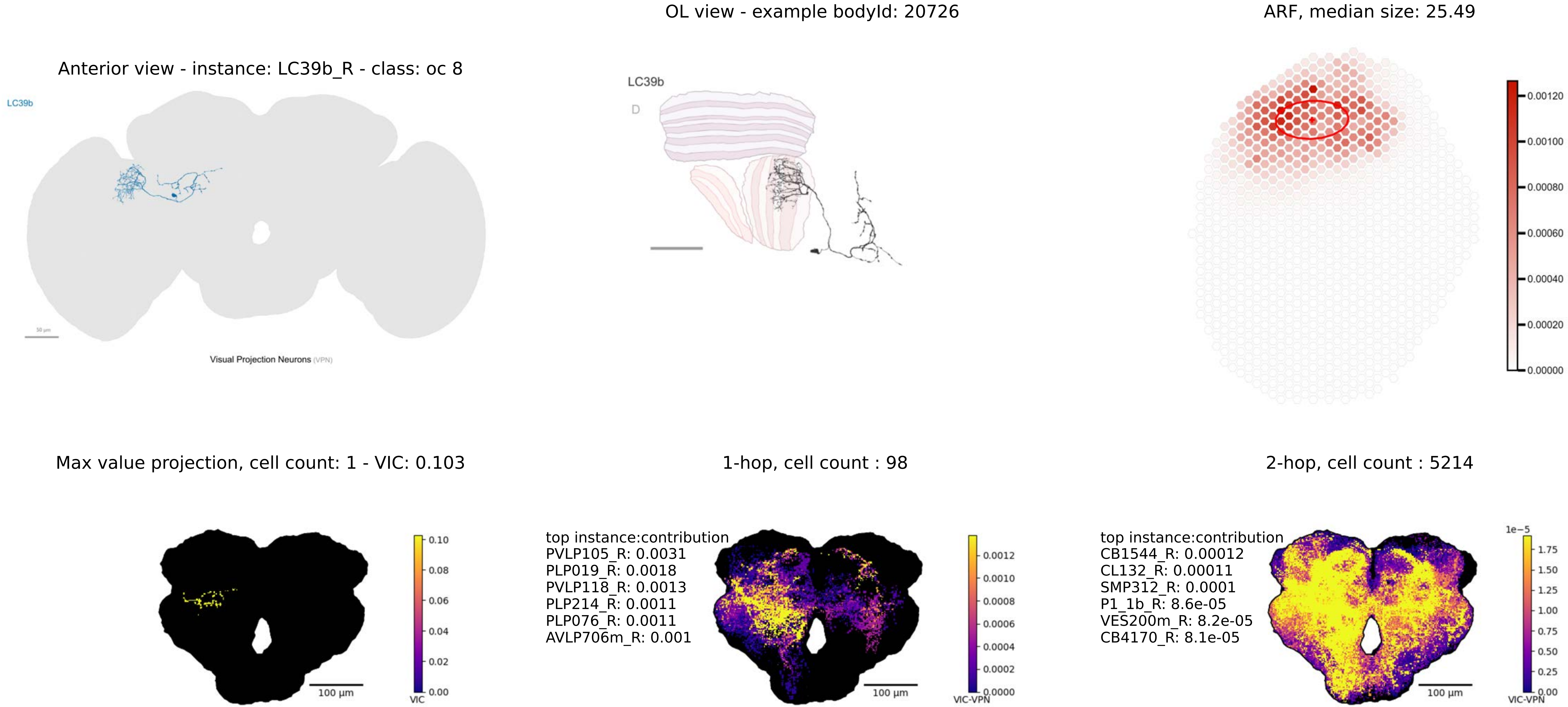

OL view - example bodyId: 20726
ARF, median size: 25.49
Anterior view - instance: LC39b_R - class: oc 8
LC39b
Visual Projection Neurons (VPN)
LC39b
D
0.00120
0.00100
0.00080
0.00060
0.00040
0.00020
0.00000
Max value projection, cell count: 1 - VIC: 0.103
1-hop, cell count : 98
2-hop, cell count : 5214
0.10
0.08
0.06
0.04
0.02
0.00
VIC
100 µm
top instance:contribution
PVLP105_R: 0.0031
PLP019_R: 0.0018
PVLP118_R: 0.0013
PLP214_R: 0.0011
PLP076_R: 0.0011
AVLP706m_R: 0.001
0.0012
0.0010
0.0008
0.0006
0.0004
0.0002
0.0000
VIC-VPN
100 µm
top instance:contribution
CB1544_R: 0.00012
CL132_R: 0.00011
SMP312_R: 0.0001
P1_1b_R: 8.6e-05
VES200m_R: 8.2e-05
CB4170_R: 8.1e-05
1e−5
1.75
1.50
1.25
1.00
0.75
0.50
0.25
0.00
VIC-VPN
100 µm

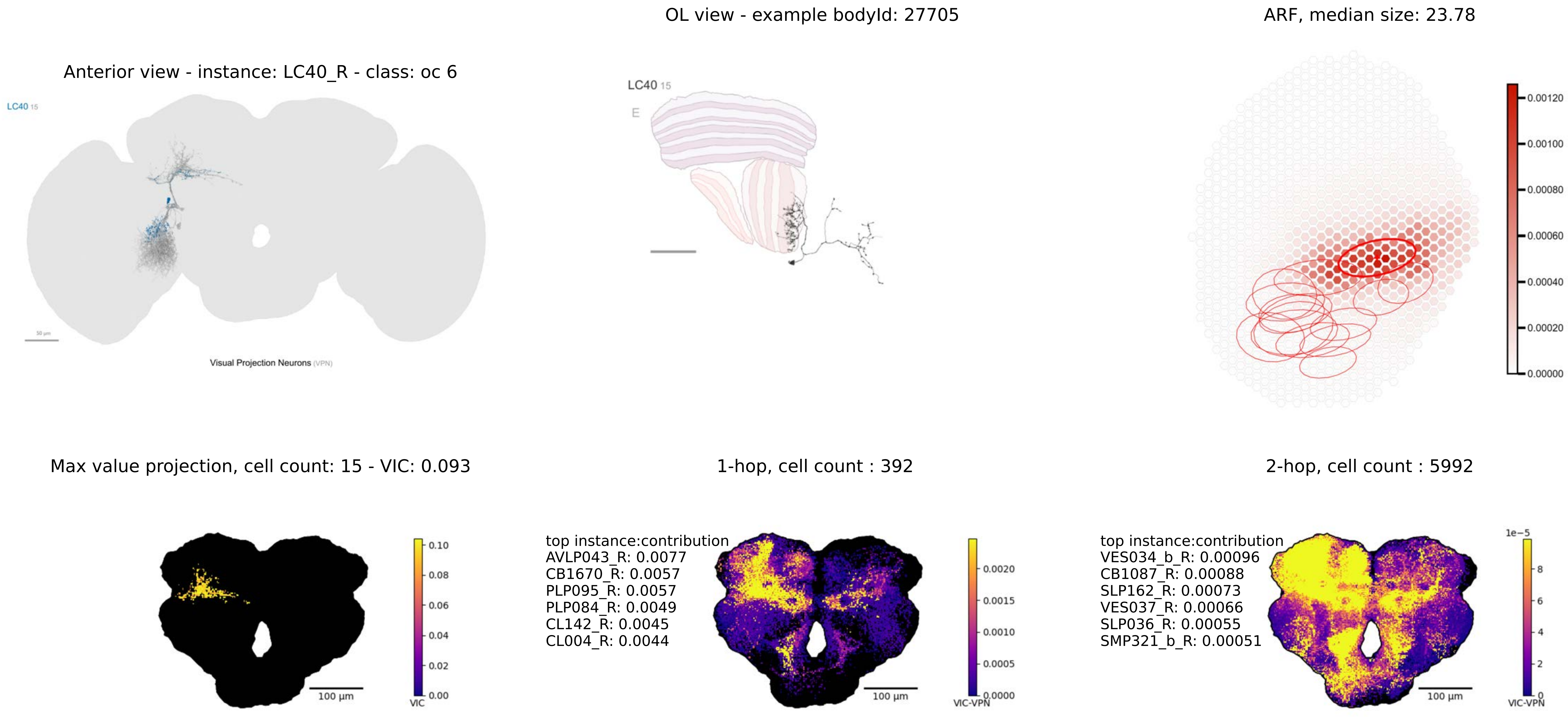

Anterior view - instance: LC40_R - class: oc 6
LC40 15
Visual Projection Neurons (VPN)
OL view - example bodyId: 27705
LC40 15
E
ARF, median size: 23.78
0.00120
0.00100
0.00080
0.00060
0.00040
0.00020
0.00000
Max value projection, cell count: 15 - VIC: 0.093
100 μm
0.10
0.08
0.06
0.04
0.02
0.00
VIC
1-hop, cell count : 392
top instance:contribution
AVLP043_R: 0.0077
CB1670_R: 0.0057
PLP095_R: 0.0057
PLP084_R: 0.0049
CL142_R: 0.0045
CL004_R: 0.0044
100 μm
0.0020
0.0015
0.0010
0.0005
0.0000
VIC-VPN
2-hop, cell count : 5992
top instance:contribution
VES034_b_R: 0.00096
CB1087_R: 0.00088
SLP162_R: 0.00073
VES037_R: 0.00066
SLP036_R: 0.00055
SMP321_b_R: 0.00051
100 μm
1e−5
8
6
4
2
0
VIC-VPN

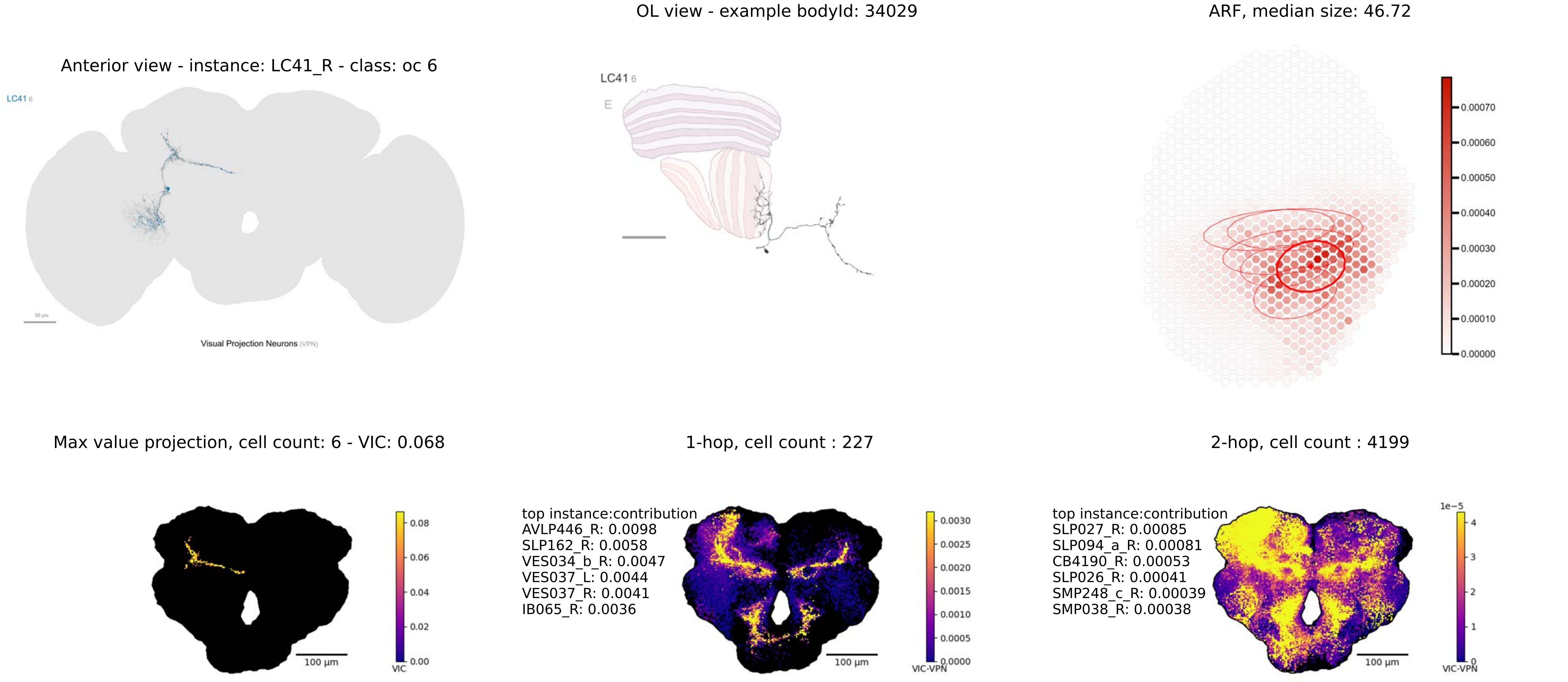
OL view - example bodyId: 34029
ARF, median size: 46.72
Anterior view - instance: LC41_R - class: oc 6
LC41 6
Visual Projection Neurons (VPN)
LC41 6
E
0.00070
0.00060
0.00050
0.00040
0.00030
0.00020
0.00010
0.00000
Max value projection, cell count: 6 - VIC: 0.068
1-hop, cell count : 227
2-hop, cell count : 4199
0.08
0.06
0.04
0.02
0.00
VIC
100 µm
top instance:contribution
AVLP446_R: 0.0098
SLP162_R: 0.0058
VES034_b_R: 0.0047
VES037_L: 0.0044
VES037_R: 0.0041
IB065_R: 0.0036
0.0030
0.0025
0.0020
0.0015
0.0010
0.0005
0.0000
VIC-VPN
100 µm
top instance:contribution
SLP027_R: 0.00085
SLP094_a_R: 0.00081
CB4190_R: 0.00053
SLP026_R: 0.00041
SMP248_c_R: 0.00039
SMP038_R: 0.00038
1e−5
4
3
2
1
0
VIC-VPN
100 µm

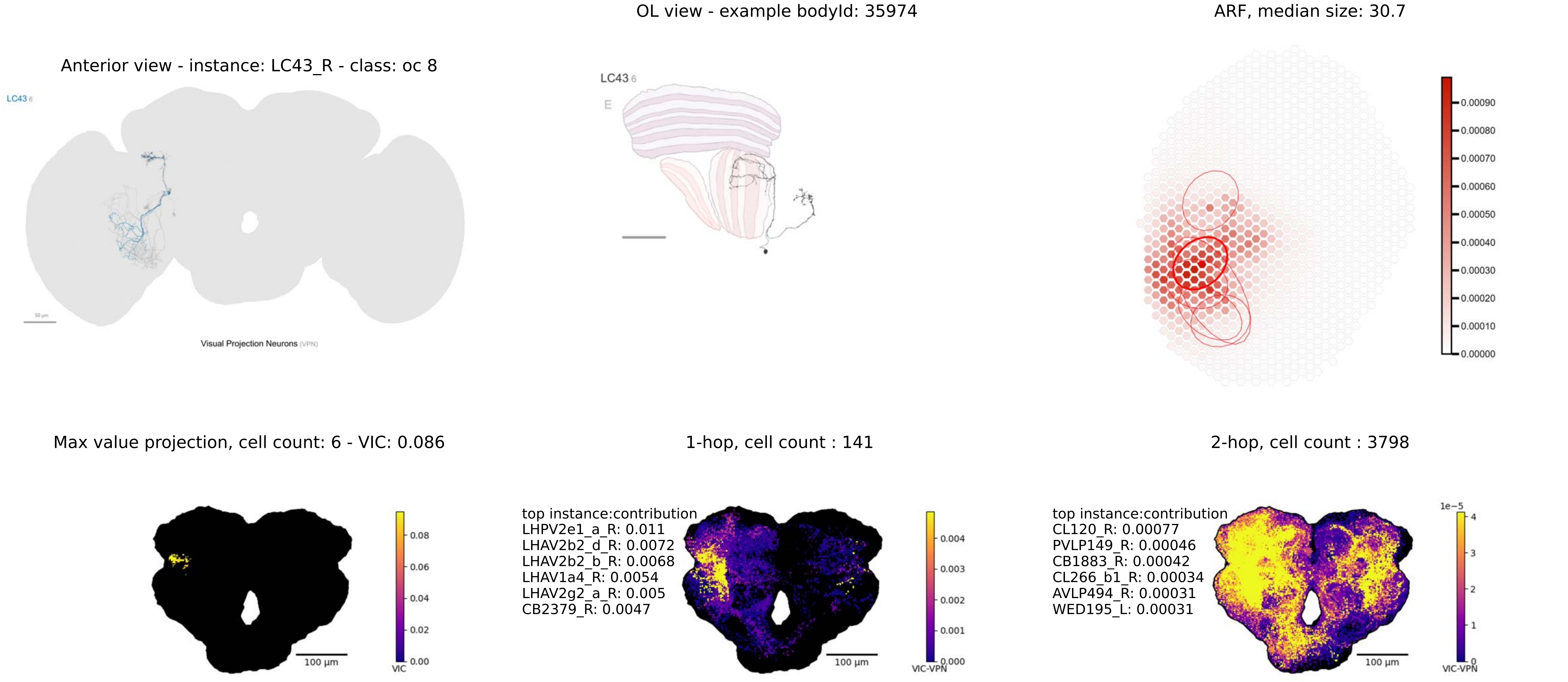
OL view - example bodyId: 35974
ARF, median size: 30.7
Anterior view - instance: LC43_R - class: oc 8
LC43 6
50 µm
Visual Projection Neurons (VPN)
LC43 6
E
0.00090
0.00080
0.00070
0.00060
0.00050
0.00040
0.00030
0.00020
0.00010
0.00000
Max value projection, cell count: 6 - VIC: 0.086
1-hop, cell count : 141
2-hop, cell count : 3798
0.08
0.06
0.04
0.02
0.00
VIC
100 µm
top instance:contribution
LHPV2e1_a_R: 0.011
LHAV2b2_d_R: 0.0072
LHAV2b2_b_R: 0.0068
LHAV1a4_R: 0.0054
LHAV2g2_a_R: 0.005
CB2379_R: 0.0047
0.004
0.003
0.002
0.001
0.000
VIC-VPN
100 µm
top instance:contribution
CL120_R: 0.00077
PVLP149_R: 0.00046
CB1883_R: 0.00042
CL266_b1_R: 0.00034
AVLP494_R: 0.00031
WED195_L: 0.00031
1e−5
4
3
2
1
0
VIC-VPN
100 µm

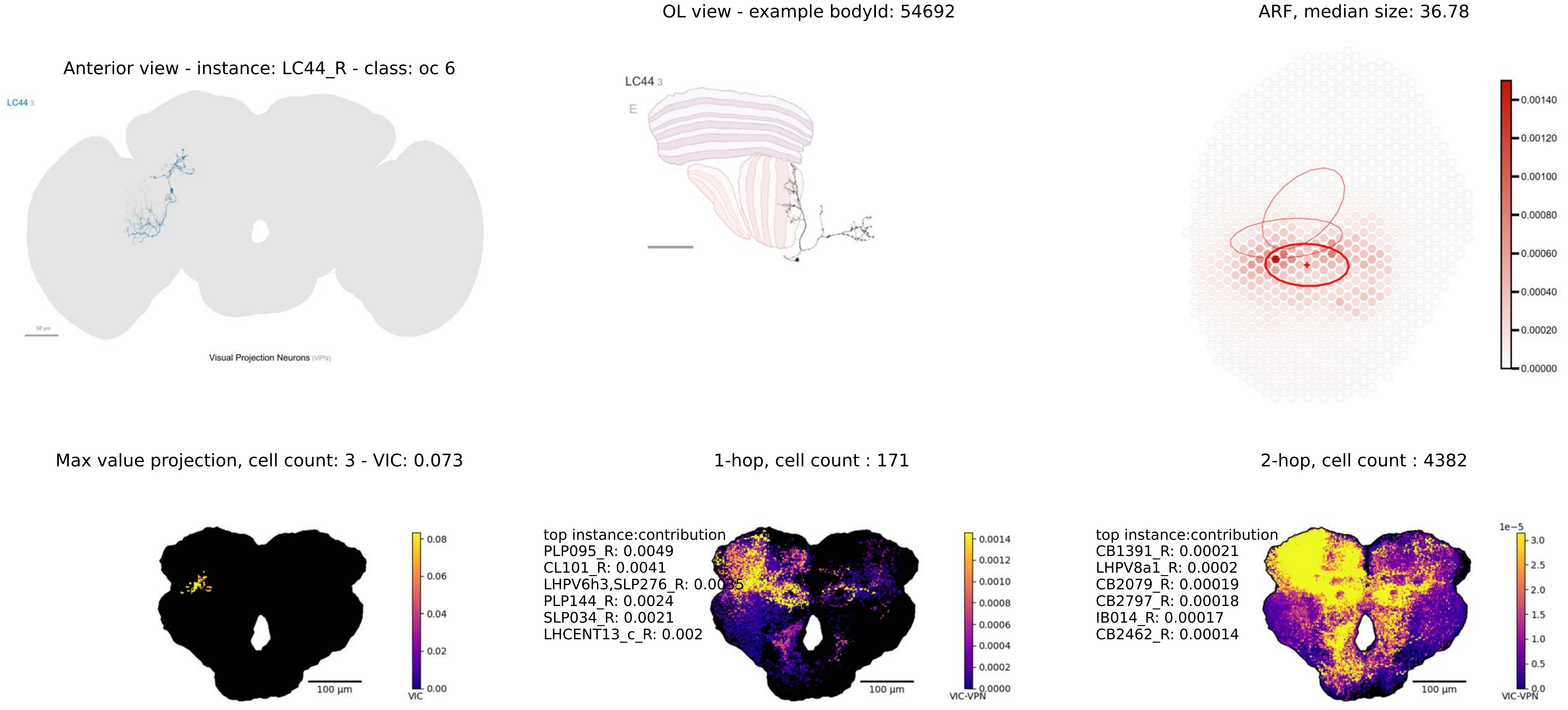

OL view - example bodyId: 54692
ARF, median size: 36.78
Anterior view - instance: LC44_R - class: oc 6
LC44 3
Visual Projection Neurons (VPN)
LC44 3
E
0.00140
0.00120
0.00100
0.00080
0.00060
0.00040
0.00020
0.00000
Max value projection, cell count: 3 - VIC: 0.073
1-hop, cell count : 171
2-hop, cell count : 4382
0.08
0.06
0.04
0.02
0.00
VIC
100 µm
top instance:contribution
PLP095_R: 0.0049
CL101_R: 0.0041
LHPV6h3,SLP276_R: 0.0035
PLP144_R: 0.0024
SLP034_R: 0.0021
LHCENT13_c_R: 0.002
0.0014
0.0012
0.0010
0.0008
0.0006
0.0004
0.0002
0.0000
VIC-VPN
100 µm
top instance:contribution
CB1391_R: 0.00021
LHPV8a1_R: 0.0002
CB2079_R: 0.00019
CB2797_R: 0.00018
IB014_R: 0.00017
CB2462_R: 0.00014
1e−5
3.0
2.5
2.0
1.5
1.0
0.5
0.0
VIC-VPN
100 µm

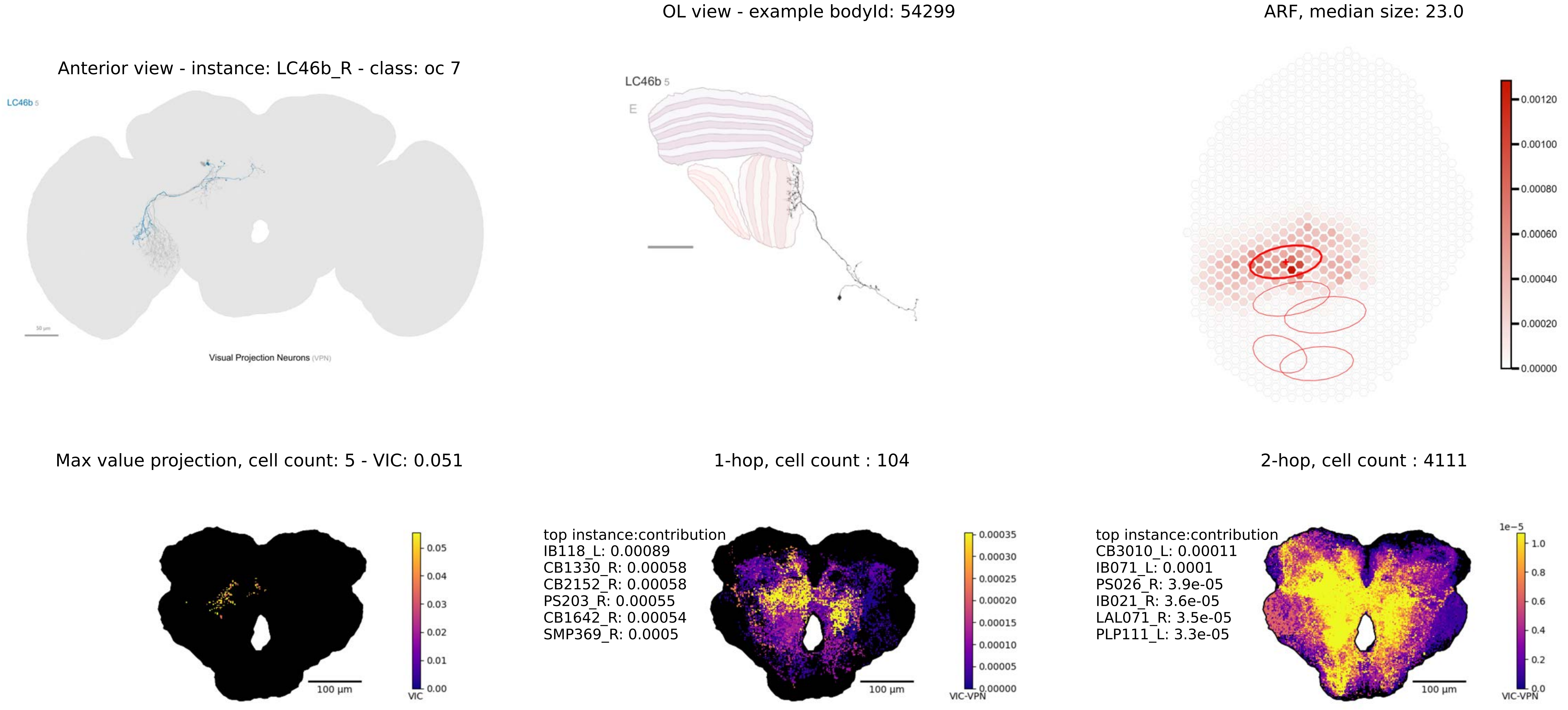

OL view - example bodyId: 54299
ARF, median size: 23.0
Anterior view - instance: LC46b_R - class: oc 7
LC46b 5
Visual Projection Neurons (VPN)
LC46b 5
E
0.00120
0.00100
0.00080
0.00060
0.00040
0.00020
0.00000
Max value projection, cell count: 5 - VIC: 0.051
1-hop, cell count : 104
2-hop, cell count : 4111
0.05
0.04
0.03
0.02
0.01
0.00
VIC
100 µm
top instance:contribution
IB118_L: 0.00089
CB1330_R: 0.00058
CB2152_R: 0.00058
PS203_R: 0.00055
CB1642_R: 0.00054
SMP369_R: 0.0005
0.00035
0.00030
0.00025
0.00020
0.00015
0.00010
0.00005
0.00000
VIC-VPN
100 µm
top instance:contribution
CB3010_L: 0.00011
IB071_L: 0.0001
PS026_R: 3.9e-05
IB021_R: 3.6e-05
LAL071_R: 3.5e-05
PLP111_L: 3.3e-05
1e−5
1.0
0.8
0.6
0.4
0.2
0.0
VIC-VPN
100 µm

Anterior view - instance: LLPC1_R - class: oc 9

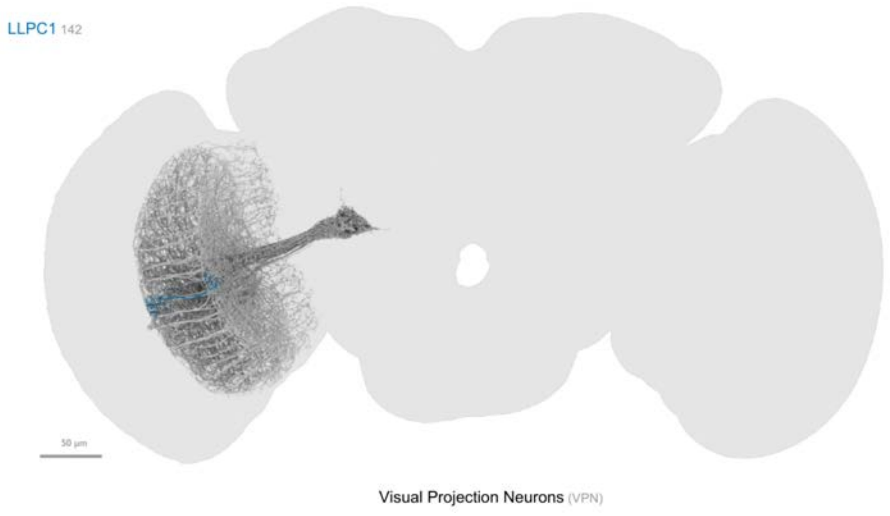


OL view - example bodyId: 26070

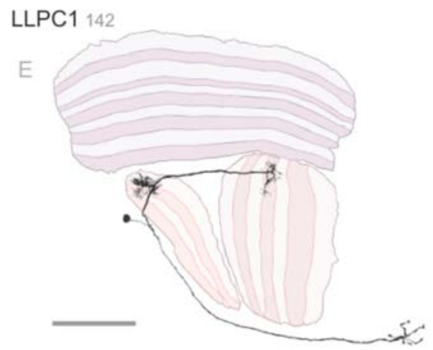


ARF, median size: 5.57

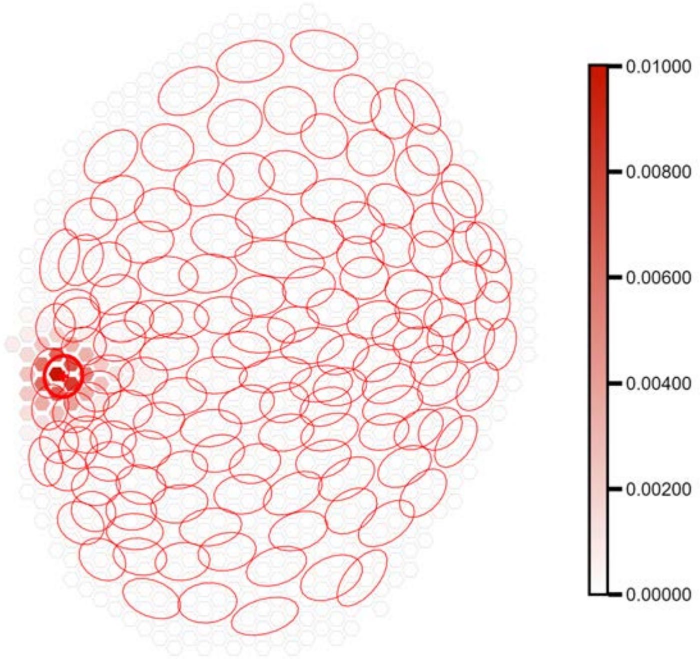


Max value projection, cell count: 142 - VIC: 0.119

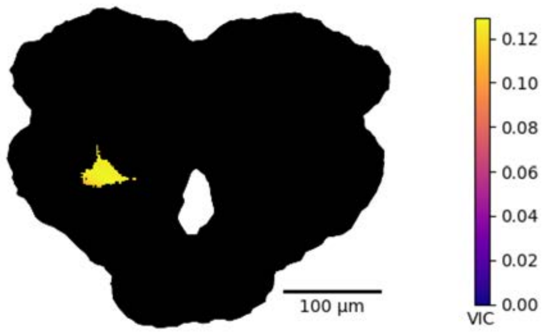


1-hop, cell count : 336

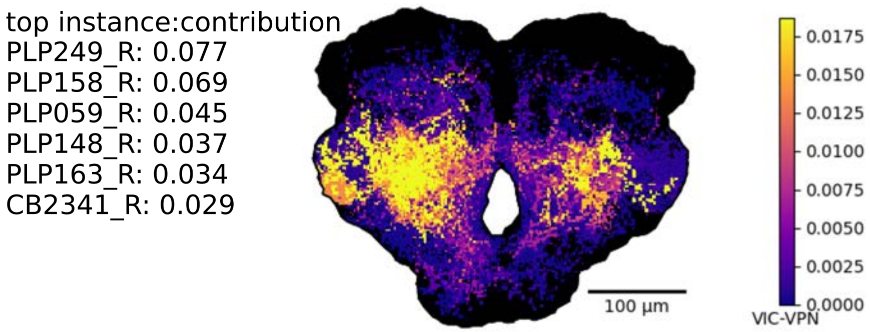


2-hop, cell count : 6800

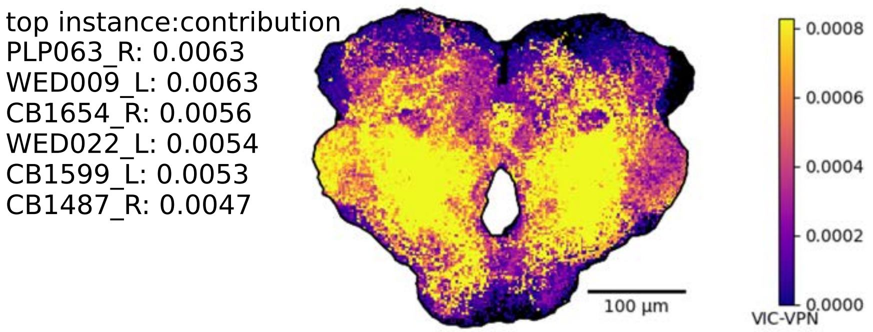

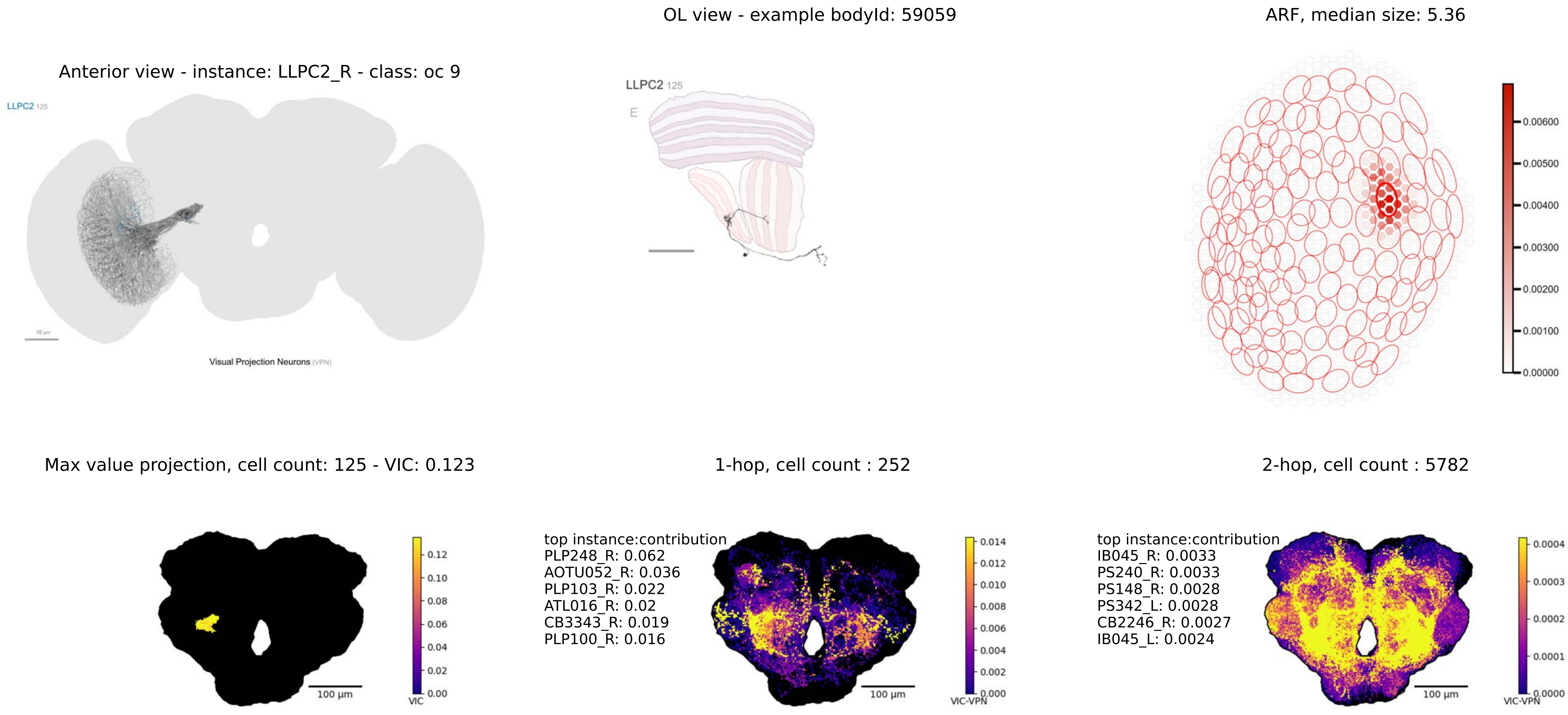
OL view - example bodyId: 59059
ARF, median size: 5.36
Anterior view - instance: LLPC2_R - class: oc 9
LLPC2 125
50 µm
Visual Projection Neurons (VPN)
LLPC2 125
E
0.00600
0.00500
0.00400
0.00300
0.00200
0.00100
0.00000
Max value projection, cell count: 125 - VIC: 0.123
1-hop, cell count : 252
2-hop, cell count : 5782
0.12
0.10
0.08
0.06
0.04
0.02
0.00
VIC
100 µm
top instance:contribution
PLP248_R: 0.062
AOTU052_R: 0.036
PLP103_R: 0.022
ATL016_R: 0.02
CB3343_R: 0.019
PLP100_R: 0.016
0.014
0.012
0.010
0.008
0.006
0.004
0.002
0.000
VIC-VPN
100 µm
top instance:contribution
IB045_R: 0.0033
PS240_R: 0.0033
PS148_R: 0.0028
PS342_L: 0.0028
CB2246_R: 0.0027
IB045_L: 0.0024
0.0004
0.0003
0.0002
0.0001
0.0000
VIC-VPN
100 µm

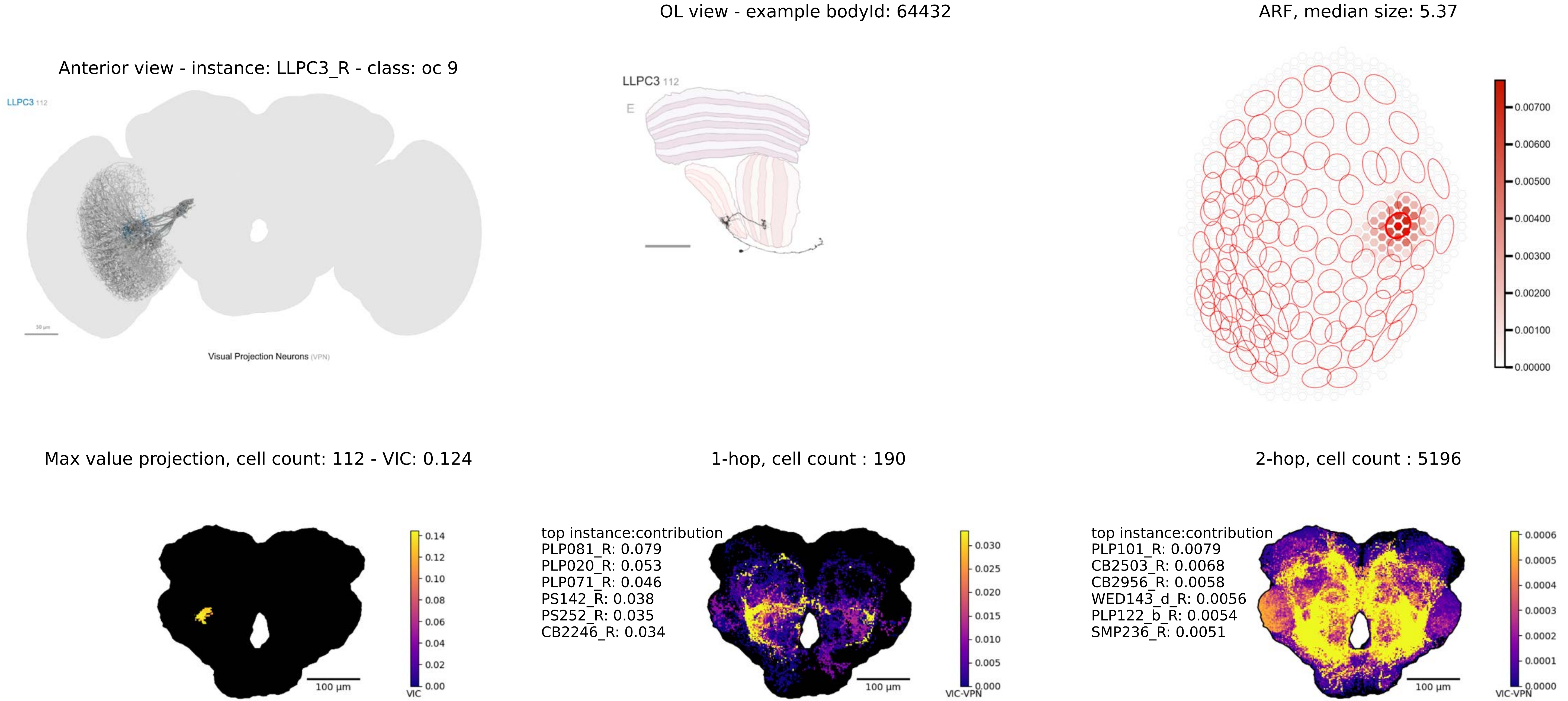

Anterior view - instance: LLPC3_R - class: oc 9
LLPC3 112
50 µm
Visual Projection Neurons (VPN)
OL view - example bodyId: 64432
LLPC3 112
E
ARF, median size: 5.37
0.00700
0.00600
0.00500
0.00400
0.00300
0.00200
0.00100
0.00000
Max value projection, cell count: 112 - VIC: 0.124
100 µm
0.14
0.12
0.10
0.08
0.06
0.04
0.02
0.00
VIC
1-hop, cell count : 190
top instance:contribution
PLP081_R: 0.079
PLP020_R: 0.053
PLP071_R: 0.046
PS142_R: 0.038
PS252_R: 0.035
CB2246_R: 0.034
100 µm
0.030
0.025
0.020
0.015
0.010
0.005
0.000
VIC-VPN
2-hop, cell count : 5196
top instance:contribution
PLP101_R: 0.0079
CB2503_R: 0.0068
CB2956_R: 0.0058
WED143_d_R: 0.0056
PLP122_b_R: 0.0054
SMP236_R: 0.0051
100 µm
0.0006
0.0005
0.0004
0.0003
0.0002
0.0001
0.0000
VIC-VPN

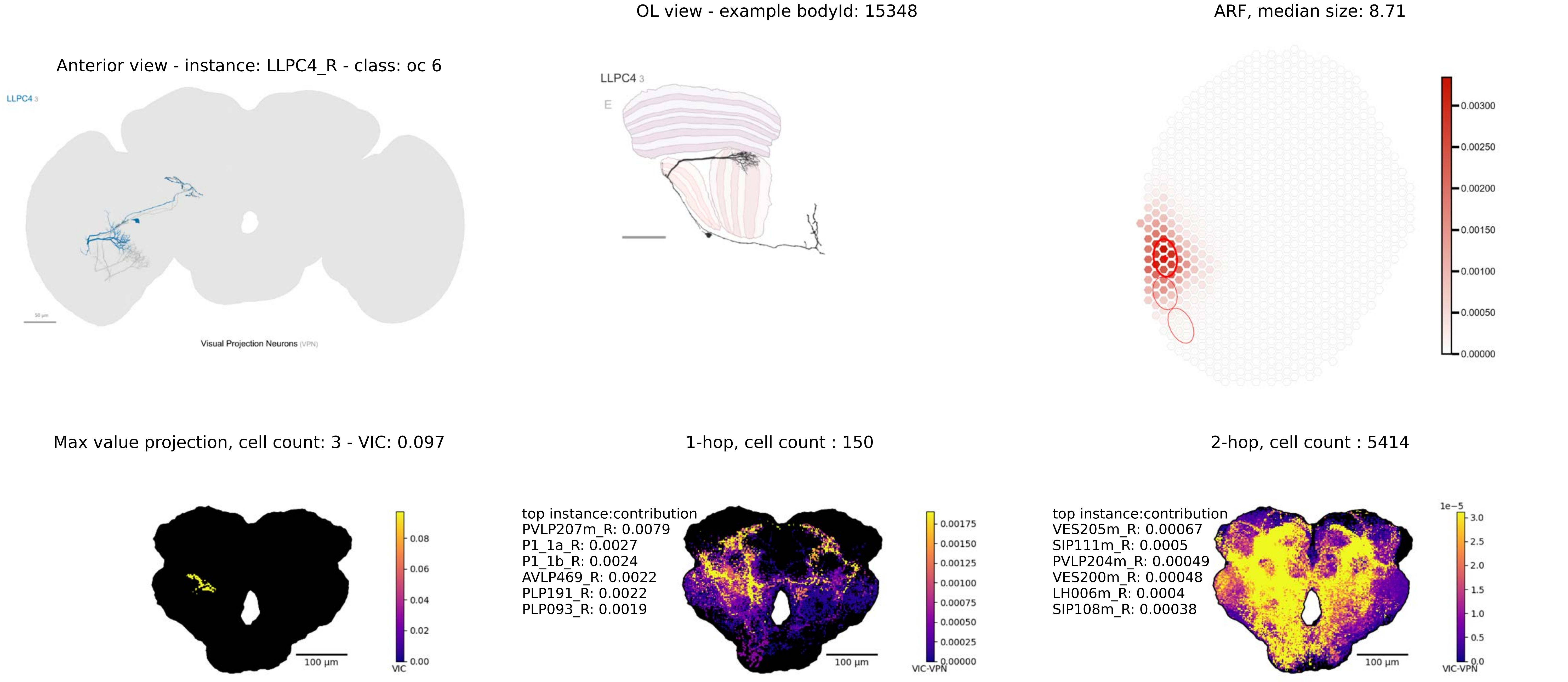

OL view - example bodyId: 15348
ARF, median size: 8.71
Anterior view - instance: LLPC4_R - class: oc 6
LLPC4 3
Visual Projection Neurons (VPN)
LLPC4 3
E
0.00300
0.00250
0.00200
0.00150
0.00100
0.00050
0.00000
Max value projection, cell count: 3 - VIC: 0.097
1-hop, cell count : 150
2-hop, cell count : 5414
top instance:contribution
PVLP207m_R: 0.0079
P1_1a_R: 0.0027
P1_1b_R: 0.0024
AVLP469_R: 0.0022
PLP191_R: 0.0022
PLP093_R: 0.0019
top instance:contribution
VES205m_R: 0.00067
SIP111m_R: 0.0005
PVLP204m_R: 0.00049
VES200m_R: 0.00048
LH006m_R: 0.0004
SIP108m_R: 0.00038
0.08
0.06
0.04
0.02
0.00
VIC
100 µm
0.00175
0.00150
0.00125
0.00100
0.00075
0.00050
0.00025
0.00000
VIC-VPN
1e−5
3.0
2.5
2.0
1.5
1.0
0.5
0.0

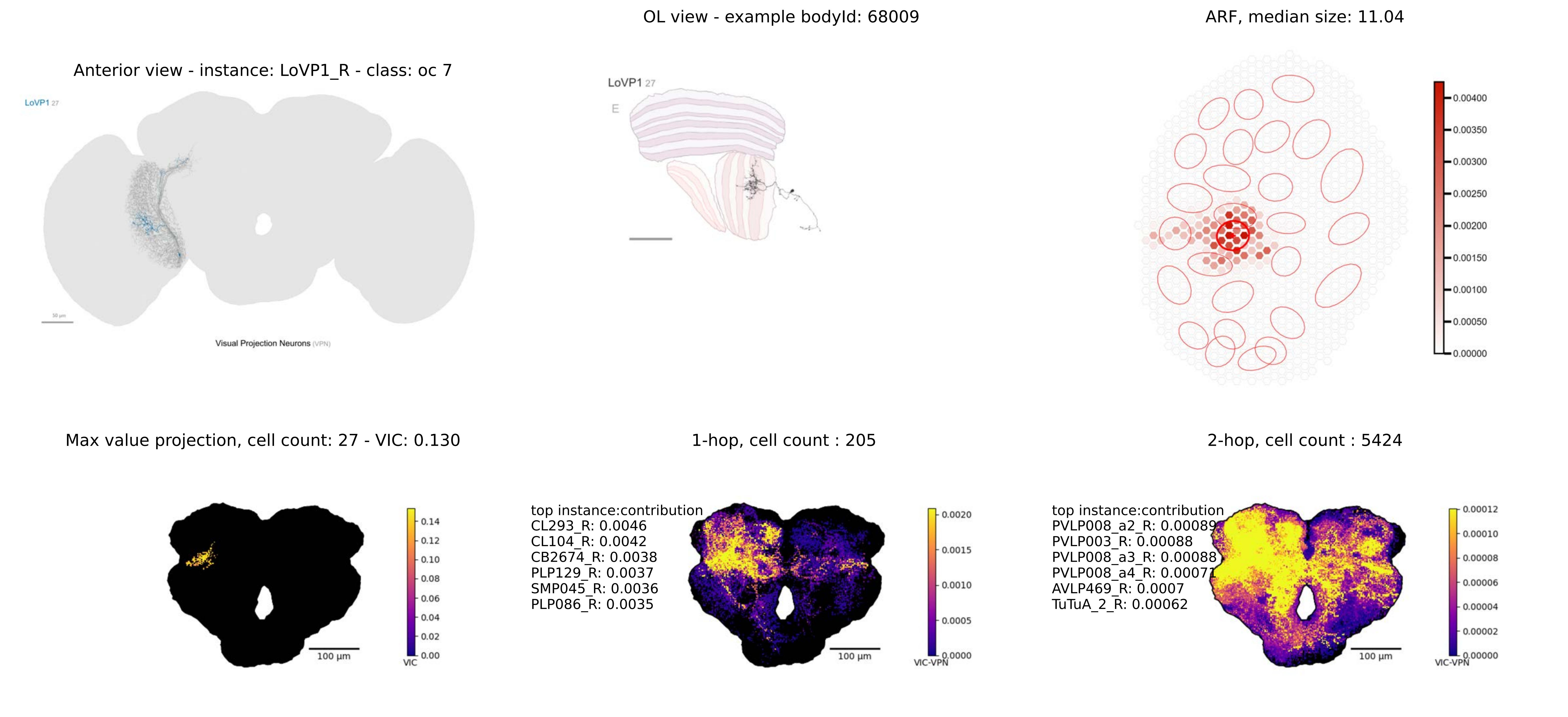

OL view - example bodyId: 68009
ARF, median size: 11.04
Anterior view - instance: LoVP1_R - class: oc 7
LoVP1 27
Visual Projection Neurons (VPN)
LoVP1 27
E
0.00400
0.00350
0.00300
0.00250
0.00200
0.00150
0.00100
0.00050
0.00000
Max value projection, cell count: 27 - VIC: 0.130
1-hop, cell count : 205
2-hop, cell count : 5424
100 µm
0.14
0.12
0.10
0.08
0.06
0.04
0.02
0.00
VIC
top instance:contribution
CL293_R: 0.0046
CL104_R: 0.0042
CB2674_R: 0.0038
PLP129_R: 0.0037
SMP045_R: 0.0036
PLP086_R: 0.0035
100 µm
0.0020
0.0015
0.0010
0.0005
0.0000
VIC-VPN
top instance:contribution
PVLP008_a2_R: 0.00089
PVLP003_R: 0.00088
PVLP008_a3_R: 0.00088
PVLP008_a4_R: 0.00071
AVLP469_R: 0.0007
TuTuA_2_R: 0.00062
100 µm
0.00012
0.00010
0.00008
0.00006
0.00004
0.00002
0.00000
VIC-VPN

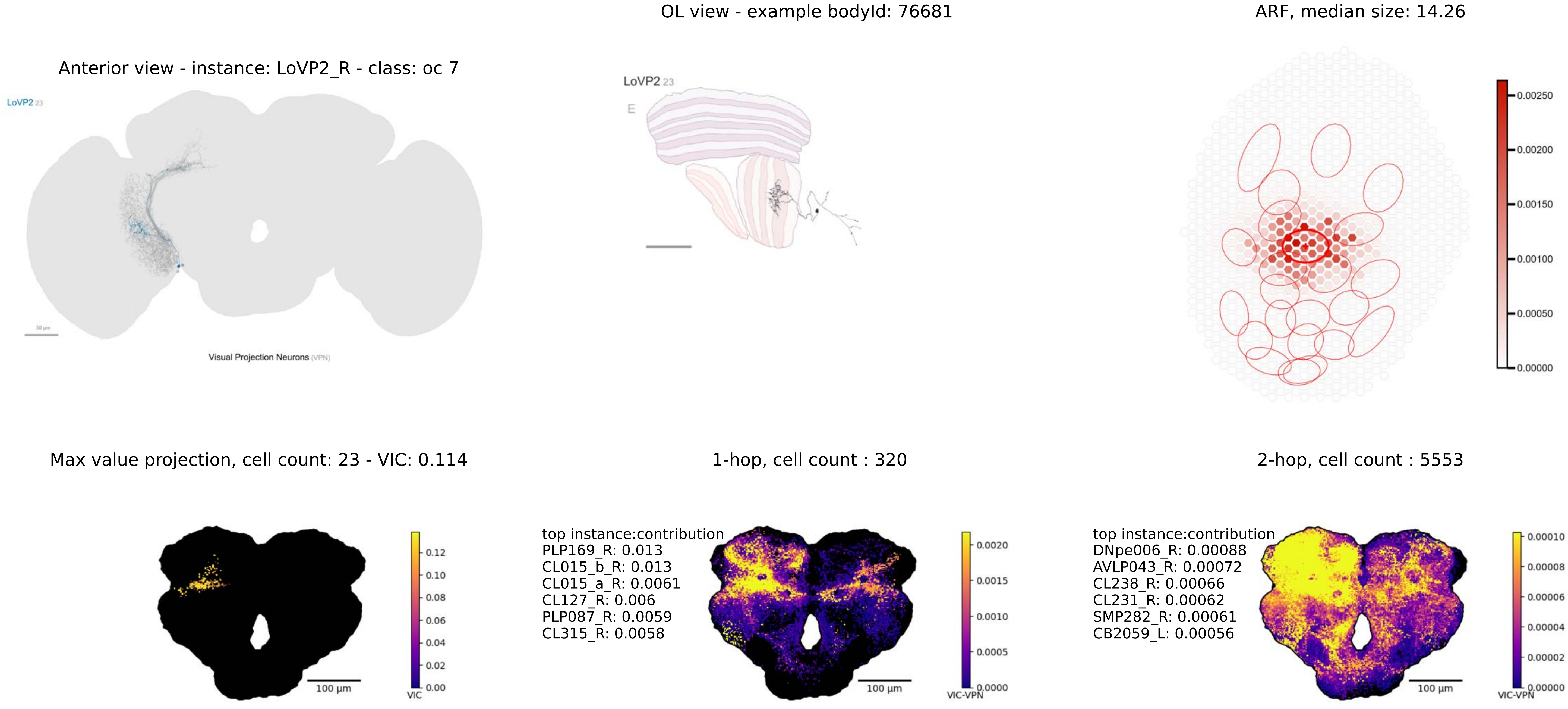
OL view - example bodyId: 76681
ARF, median size: 14.26
Anterior view - instance: LoVP2_R - class: oc 7
LoVP2 23
Visual Projection Neurons (VPN)
LoVP2 23
E
0.00250
0.00200
0.00150
0.00100
0.00050
0.00000
Max value projection, cell count: 23 - VIC: 0.114
1-hop, cell count : 320
2-hop, cell count : 5553
100 µm
0.12
0.10
0.08
0.06
0.04
0.02
0.00
VIC
top instance:contribution
PLP169_R: 0.013
CL015_b_R: 0.013
CL015_a_R: 0.0061
CL127_R: 0.006
PLP087_R: 0.0059
CL315_R: 0.0058
100 µm
0.0020
0.0015
0.0010
0.0005
0.0000
VIC-VPN
top instance:contribution
DNpe006_R: 0.00088
AVLP043_R: 0.00072
CL238_R: 0.00066
CL231_R: 0.00062
SMP282_R: 0.00061
CB2059_L: 0.00056
100 µm
0.00010
0.00008
0.00006
0.00004
0.00002
0.00000
VIC-VPN

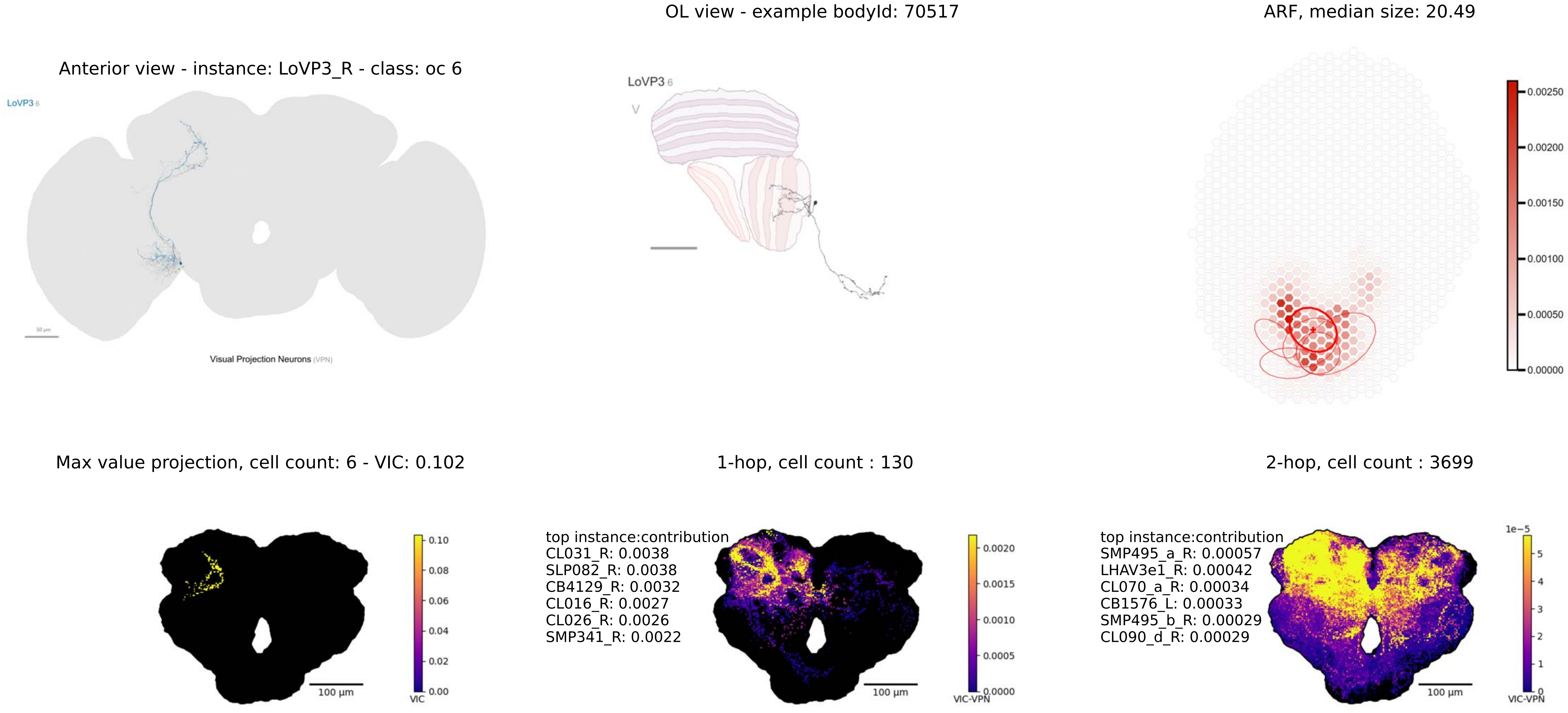

OL view - example bodyId: 70517
ARF, median size: 20.49
Anterior view - instance: LoVP3_R - class: oc 6
LoVP3 6
Visual Projection Neurons (VPN)
LoVP3 6
0.00250
0.00200
0.00150
0.00100
0.00050
0.00000
Max value projection, cell count: 6 - VIC: 0.102
1-hop, cell count : 130
2-hop, cell count : 3699
100 µm
0.10
0.08
0.06
0.04
0.02
0.00
VIC
top instance:contribution
CL031_R: 0.0038
SLP082_R: 0.0038
CB4129_R: 0.0032
CL016_R: 0.0027
CL026_R: 0.0026
SMP341_R: 0.0022
100 µm
0.0020
0.0015
0.0010
0.0005
0.0000
VIC-VPN
top instance:contribution
SMP495_a_R: 0.00057
LHAV3e1_R: 0.00042
CL070_a_R: 0.00034
CB1576_L: 0.00033
SMP495_b_R: 0.00029
CL090_d_R: 0.00029
100 µm
1e−5
5
4
3
2
1
0
VIC-VPN

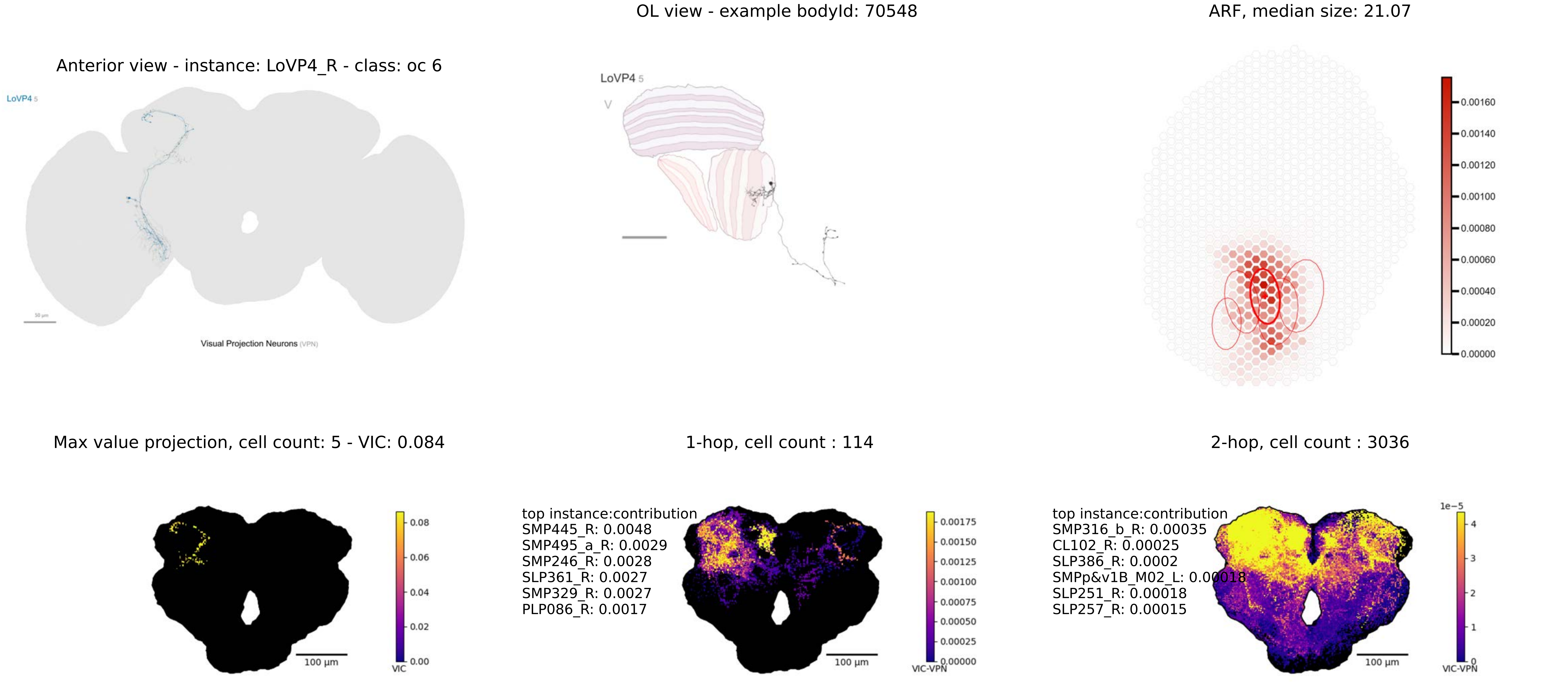
OL view - example bodyId: 70548
ARF, median size: 21.07
Anterior view - instance: LoVP4_R - class: oc 6
LoVP4 5
Visual Projection Neurons (VPN)
0.00160
0.00140
0.00120
0.00100
0.00080
0.00060
0.00040
0.00020
0.00000
Max value projection, cell count: 5 - VIC: 0.084
1-hop, cell count : 114
2-hop, cell count : 3036
0.08
0.06
0.04
0.02
0.00
VIC
100 µm
top instance:contribution
SMP445_R: 0.0048
SMP495_a_R: 0.0029
SMP246_R: 0.0028
SLP361_R: 0.0027
SMP329_R: 0.0027
PLP086_R: 0.0017
0.00175
0.00150
0.00125
0.00100
0.00075
0.00050
0.00025
0.00000
VIC-VPN
top instance:contribution
SMP316_b_R: 0.00035
CL102_R: 0.00025
SLP386_R: 0.0002
SMPp&v1B_M02_L: 0.00018
SLP251_R: 0.00018
SLP257_R: 0.00015
1e−5
4
3
2
1
0
VIC-VPN

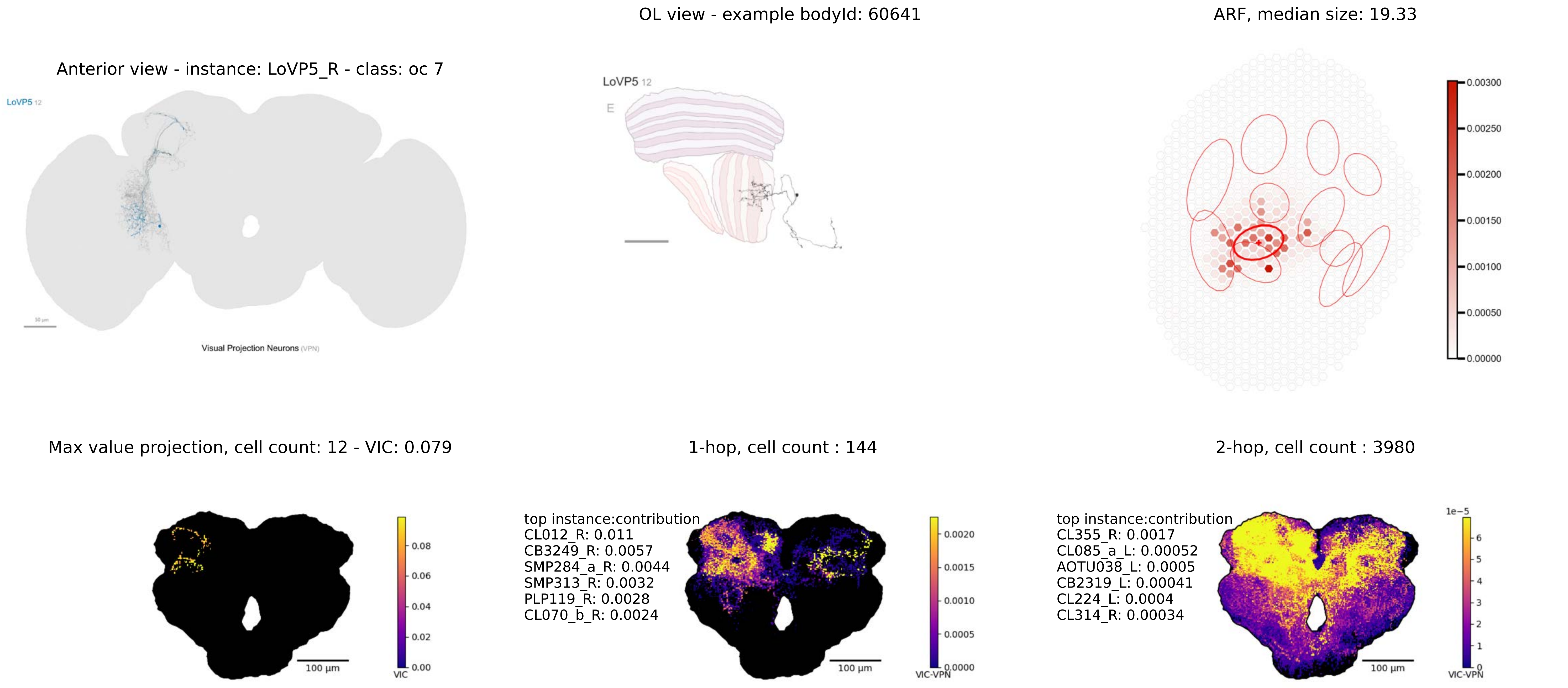

Anterior view - instance: LoVP5_R - class: oc 7
LoVP5 12
50 μm
Visual Projection Neurons (VPN)
OL view - example bodyId: 60641
LoVP5 12
E
ARF, median size: 19.33
0.00300
0.00250
0.00200
0.00150
0.00100
0.00050
0.00000
Max value projection, cell count: 12 - VIC: 0.079
100 μm
VIC
0.08
0.06
0.04
0.02
0.00
1-hop, cell count : 144
top instance:contribution
CL012_R: 0.011
CB3249_R: 0.0057
SMP284_a_R: 0.0044
SMP313_R: 0.0032
PLP119_R: 0.0028
CL070_b_R: 0.0024
100 μm
VIC-VPN
0.0020
0.0015
0.0010
0.0005
0.0000
2-hop, cell count : 3980
top instance:contribution
CL355_R: 0.0017
CL085_a_L: 0.00052
AOTU038_L: 0.0005
CB2319_L: 0.00041
CL224_L: 0.0004
CL314_R: 0.00034
100 μm
1e−5
VIC-VPN
6
5
4
3
2
1
0

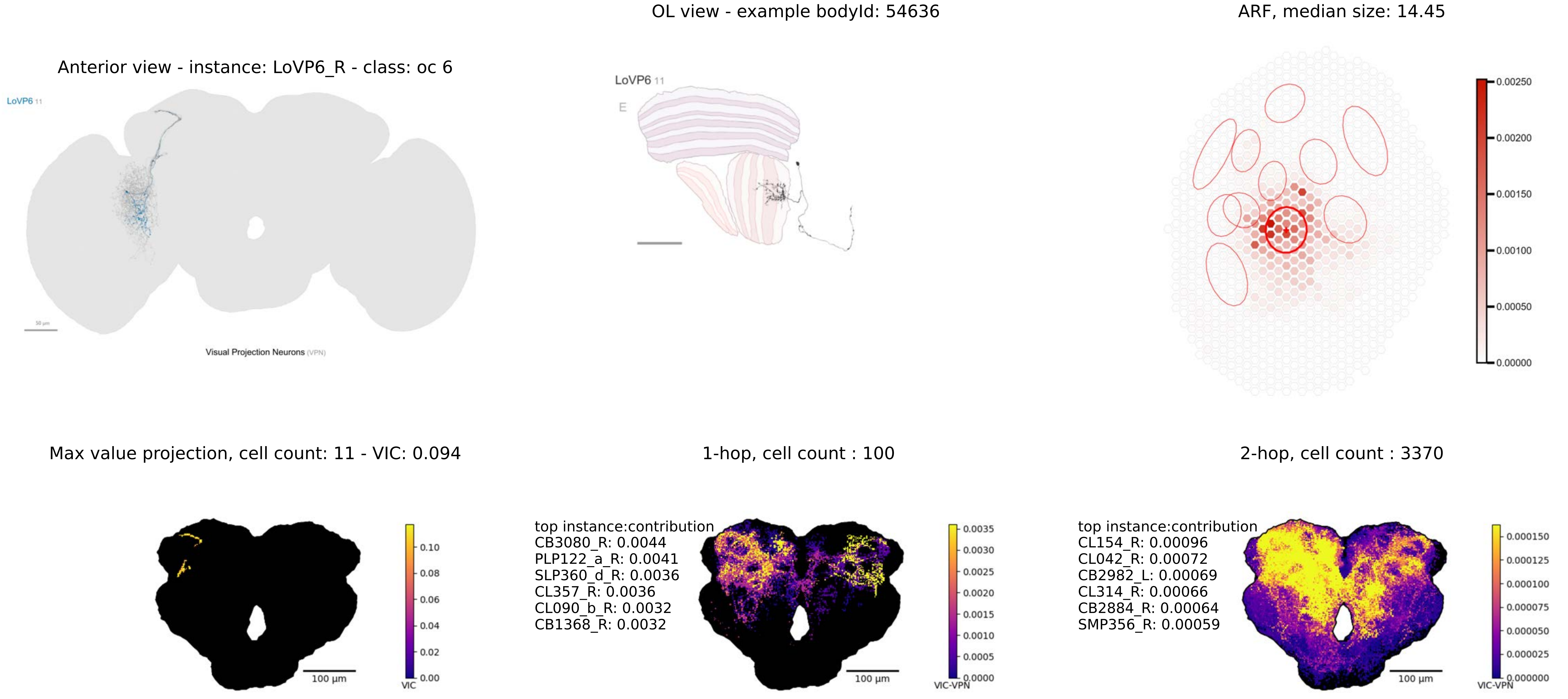
Anterior view - instance: LoVP6_R - class: oc 6
LoVP6 11
Visual Projection Neurons (VPN)
OL view - example bodyId: 54636
LoVP6 11
E
ARF, median size: 14.45
0.00250
0.00200
0.00150
0.00100
0.00050
0.00000
Max value projection, cell count: 11 - VIC: 0.094
0.10
0.08
0.06
0.04
0.02
0.00
VIC
100 µm
1-hop, cell count : 100
top instance:contribution
CB3080_R: 0.0044
PLP122_a_R: 0.0041
SLP360_d_R: 0.0036
CL357_R: 0.0036
CL090_b_R: 0.0032
CB1368_R: 0.0032
0.0035
0.0030
0.0025
0.0020
0.0015
0.0010
0.0005
0.0000
VIC-VPN
100 µm
2-hop, cell count : 3370
top instance:contribution
CL154_R: 0.00096
CL042_R: 0.00072
CB2982_L: 0.00069
CL314_R: 0.00066
CB2884_R: 0.00064
SMP356_R: 0.00059
0.000150
0.000125
0.000100
0.000075
0.000050
0.000025
0.000000
VIC-VPN
100 µm

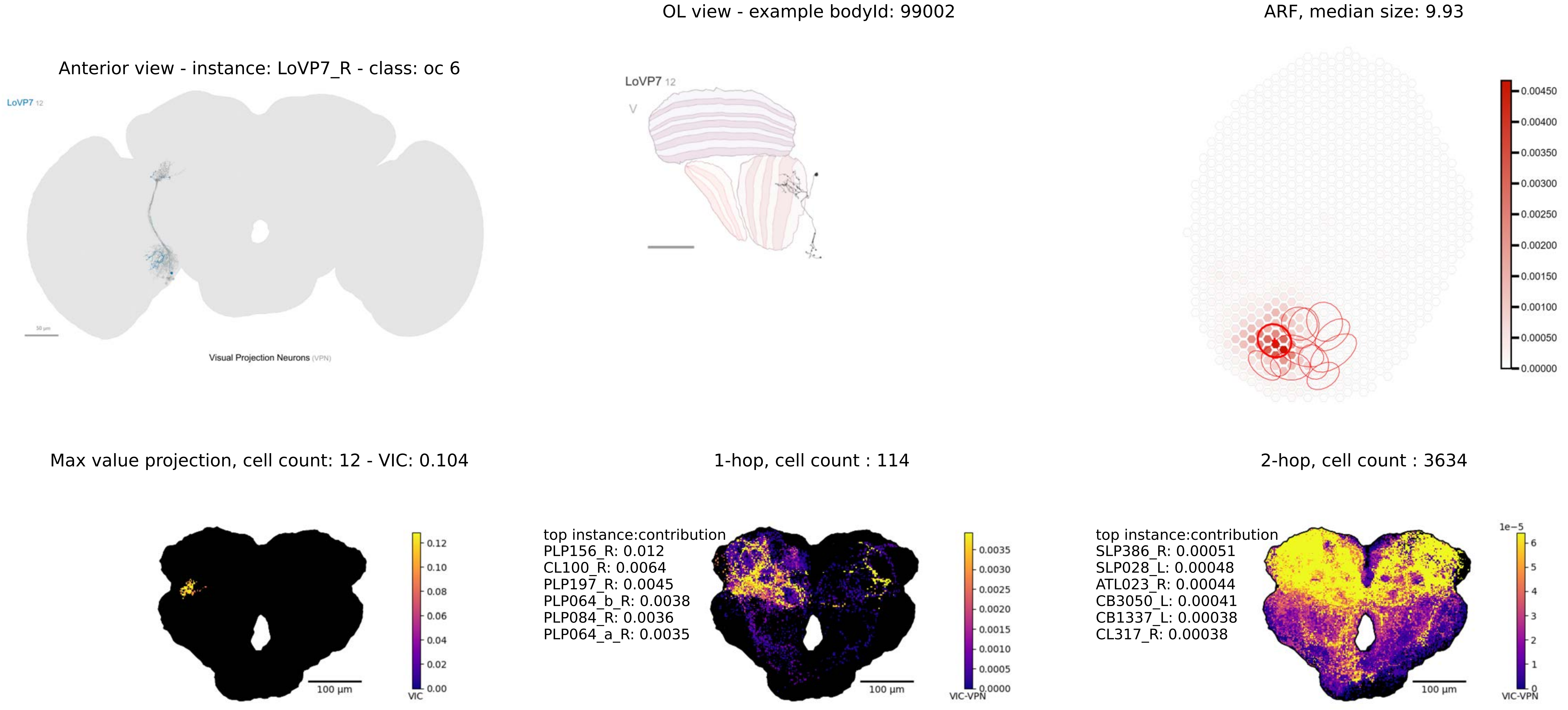
OL view - example bodyId: 99002
ARF, median size: 9.93
Anterior view - instance: LoVP7_R - class: oc 6
LoVP7 12
Visual Projection Neurons (VPN)
LoVP7 12
V
0.00450
0.00400
0.00350
0.00300
0.00250
0.00200
0.00150
0.00100
0.00050
0.00000
Max value projection, cell count: 12 - VIC: 0.104
1-hop, cell count : 114
2-hop, cell count : 3634
top instance:contribution
PLP156_R: 0.012
CL100_R: 0.0064
PLP197_R: 0.0045
PLP064_b_R: 0.0038
PLP084_R: 0.0036
PLP064_a_R: 0.0035
top instance:contribution
SLP386_R: 0.00051
SLP028_L: 0.00048
ATL023_R: 0.00044
CB3050_L: 0.00041
CB1337_L: 0.00038
CL317_R: 0.00038
100 μm
VIC
0.12
0.10
0.08
0.06
0.04
0.02
0.00
100 μm
VIC-VPN
0.0035
0.0030
0.0025
0.0020
0.0015
0.0010
0.0005
0.0000
100 μm
1e−5
6
5
4
3
2
1
0
VIC-VPN

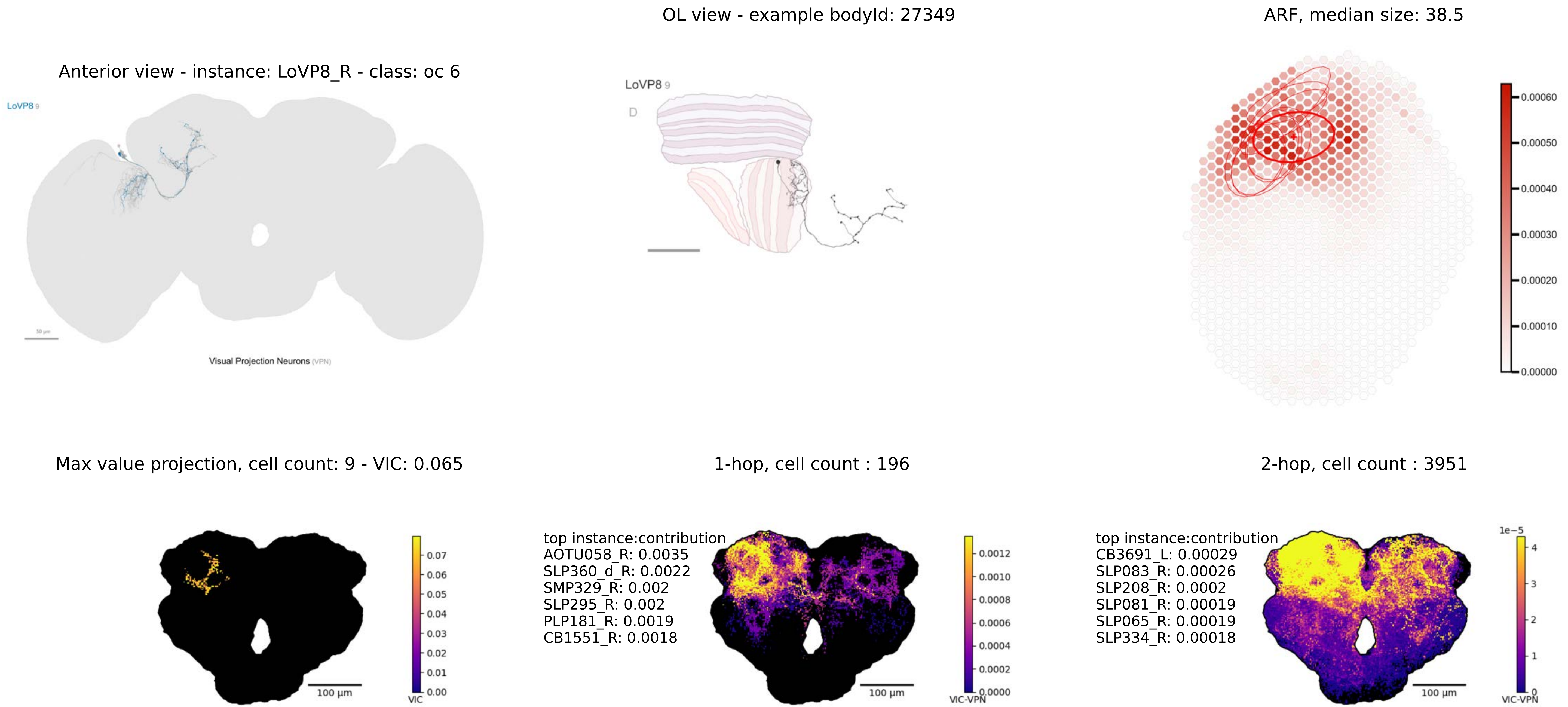

OL view - example bodyId: 27349
ARF, median size: 38.5
Anterior view - instance: LoVP8_R - class: oc 6
LoVP8 9
Visual Projection Neurons (VPN)
LoVP8 9
D
0.00060
0.00050
0.00040
0.00030
0.00020
0.00010
0.00000
Max value projection, cell count: 9 - VIC: 0.065
1-hop, cell count : 196
2-hop, cell count : 3951
100 µm
0.07
0.06
0.05
0.04
0.03
0.02
0.01
0.00
VIC
top instance:contribution
AOTU058_R: 0.0035
SLP360_d_R: 0.0022
SMP329_R: 0.002
SLP295_R: 0.002
PLP181_R: 0.0019
CB1551_R: 0.0018
100 µm
0.0012
0.0010
0.0008
0.0006
0.0004
0.0002
0.0000
VIC-VPN
top instance:contribution
CB3691_L: 0.00029
SLP083_R: 0.00026
SLP208_R: 0.0002
SLP081_R: 0.00019
SLP065_R: 0.00019
SLP334_R: 0.00018
100 µm
1e−5
4
3
2
1
0
VIC-VPN

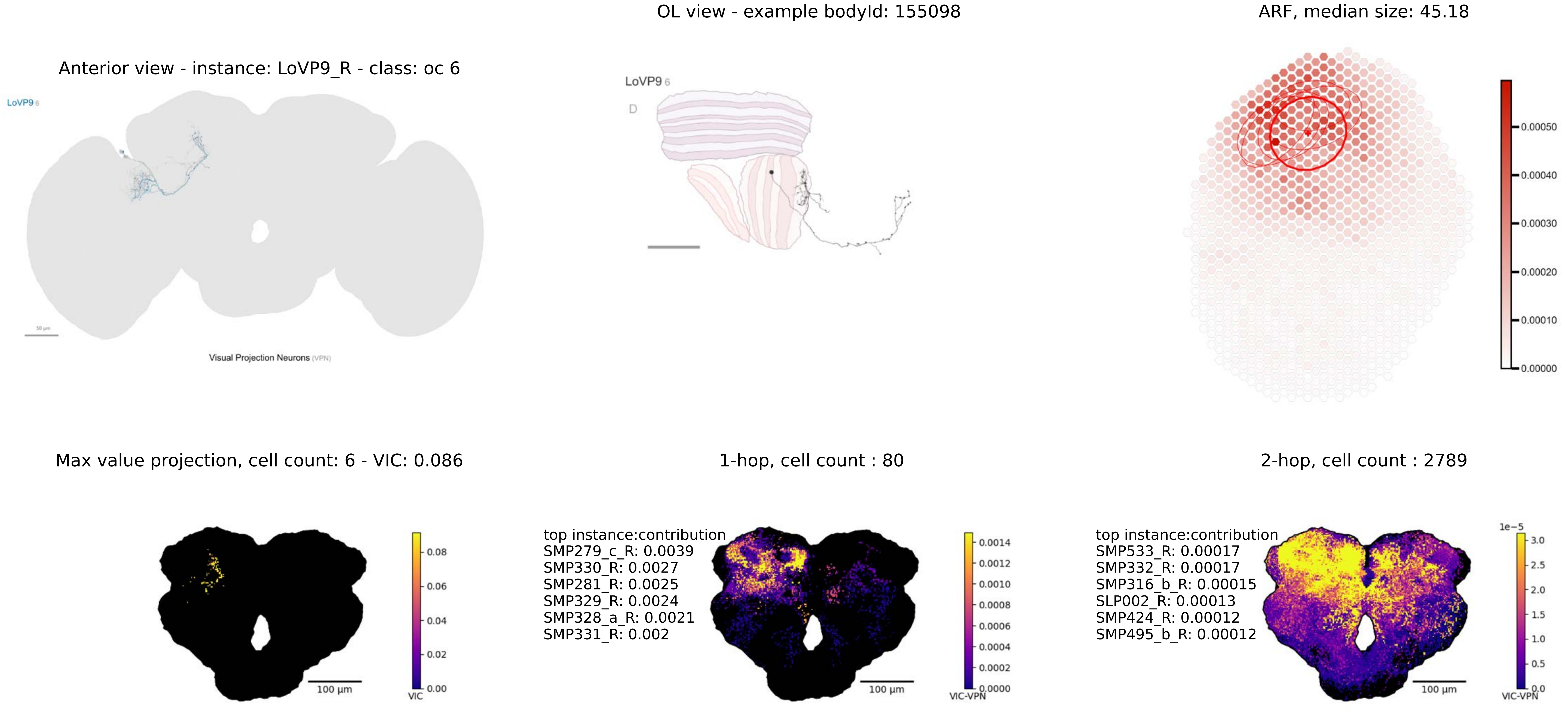

OL view - example bodyId: 155098
ARF, median size: 45.18
Anterior view - instance: LoVP9_R - class: oc 6
LoVP9 6
Visual Projection Neurons (VPN)
LoVP9 6
D
0.00050
0.00040
0.00030
0.00020
0.00010
0.00000
Max value projection, cell count: 6 - VIC: 0.086
1-hop, cell count : 80
2-hop, cell count : 2789
0.08
0.06
0.04
0.02
0.00
VIC
100 µm
top instance:contribution
SMP279_c_R: 0.0039
SMP330_R: 0.0027
SMP281_R: 0.0025
SMP329_R: 0.0024
SMP328_a_R: 0.0021
SMP331_R: 0.002
0.0014
0.0012
0.0010
0.0008
0.0006
0.0004
0.0002
0.0000
VIC-VPN
100 µm
top instance:contribution
SMP533_R: 0.00017
SMP332_R: 0.00017
SMP316_b_R: 0.00015
SLP002_R: 0.00013
SMP424_R: 0.00012
SMP495_b_R: 0.00012
1e−5
3.0
2.5
2.0
1.5
1.0
0.5
0.0
VIC-VPN
100 µm

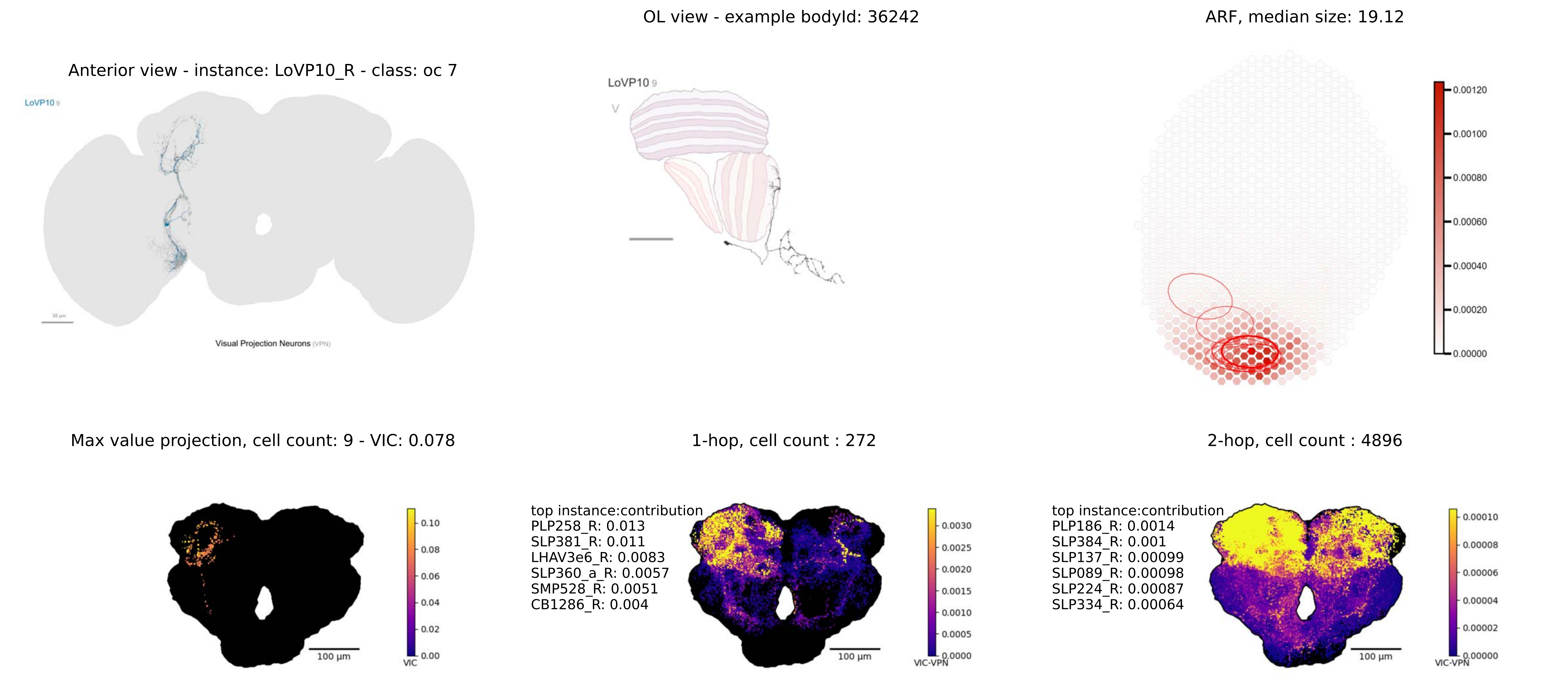

OL view - example bodyId: 36242
ARF, median size: 19.12
Anterior view - instance: LoVP10_R - class: oc 7
LoVP10 9
Visual Projection Neurons (VPN)
LoVP10 9
V
0.00120
0.00100
0.00080
0.00060
0.00040
0.00020
0.00000
Max value projection, cell count: 9 - VIC: 0.078
1-hop, cell count : 272
2-hop, cell count : 4896
100 µm
VIC
0.10
0.08
0.06
0.04
0.02
0.00
top instance:contribution
PLP258_R: 0.013
SLP381_R: 0.011
LHAV3e6_R: 0.0083
SLP360_a_R: 0.0057
SMP528_R: 0.0051
CB1286_R: 0.004
100 µm
VIC-VPN
0.0030
0.0025
0.0020
0.0015
0.0010
0.0005
0.0000
top instance:contribution
PLP186_R: 0.0014
SLP384_R: 0.001
SLP137_R: 0.00099
SLP089_R: 0.00098
SLP224_R: 0.00087
SLP334_R: 0.00064
100 µm
VIC-VPN
0.00010
0.00008
0.00006
0.00004
0.00002
0.00000

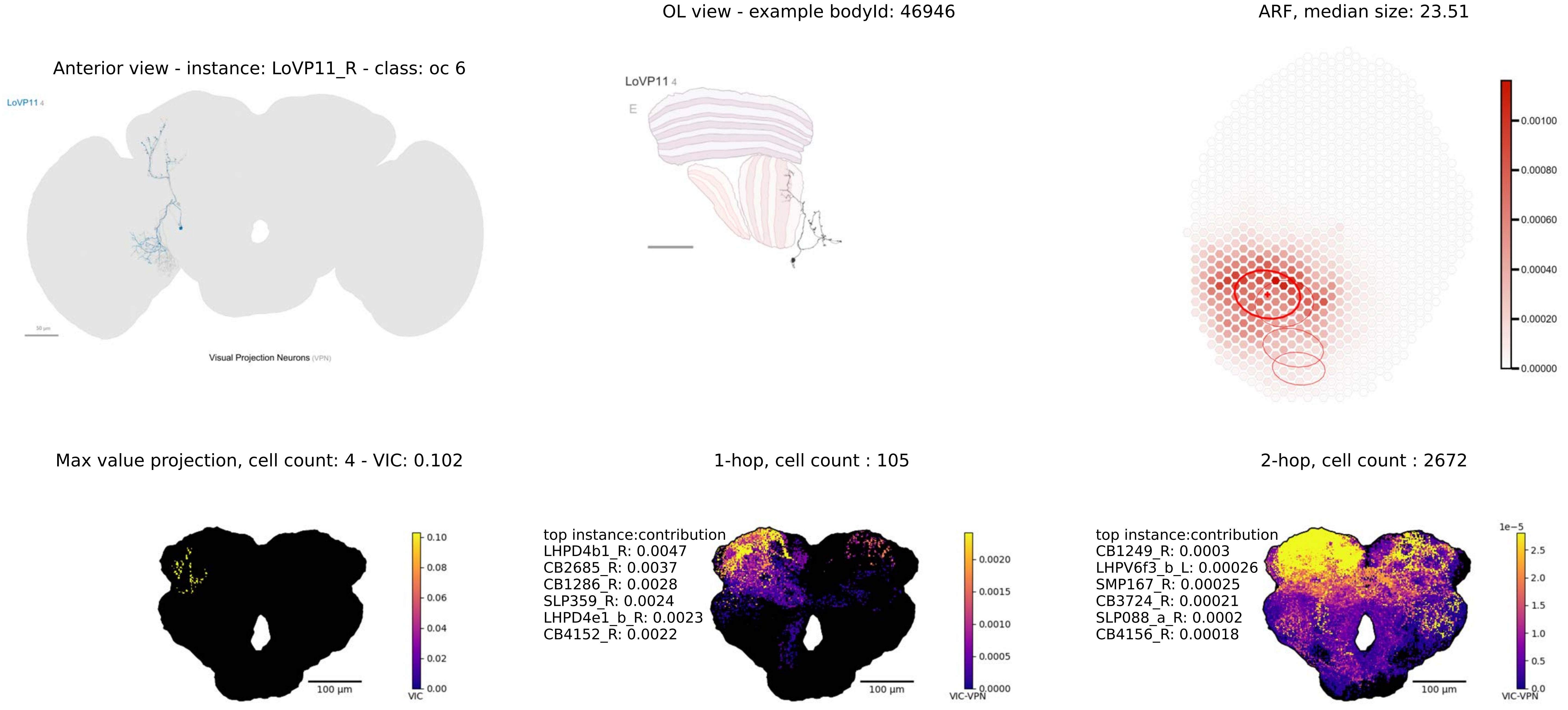
OL view - example bodyId: 46946
ARF, median size: 23.51
Anterior view - instance: LoVP11_R - class: oc 6
LoVP11 4
Visual Projection Neurons (VPN)
LoVP11 4
E
0.00100
0.00080
0.00060
0.00040
0.00020
0.00000
Max value projection, cell count: 4 - VIC: 0.102
1-hop, cell count : 105
2-hop, cell count : 2672
0.10
0.08
0.06
0.04
0.02
0.00
VIC
100 µm
top instance:contribution
LHPD4b1_R: 0.0047
CB2685_R: 0.0037
CB1286_R: 0.0028
SLP359_R: 0.0024
LHPD4e1_b_R: 0.0023
CB4152_R: 0.0022
0.0020
0.0015
0.0010
0.0005
0.0000
VIC-VPN
100 µm
top instance:contribution
CB1249_R: 0.0003
LHPV6f3_b_L: 0.00026
SMP167_R: 0.00025
CB3724_R: 0.00021
SLP088_a_R: 0.0002
CB4156_R: 0.00018
1e−5
2.5
2.0
1.5
1.0
0.5
0.0
VIC-VPN
100 µm

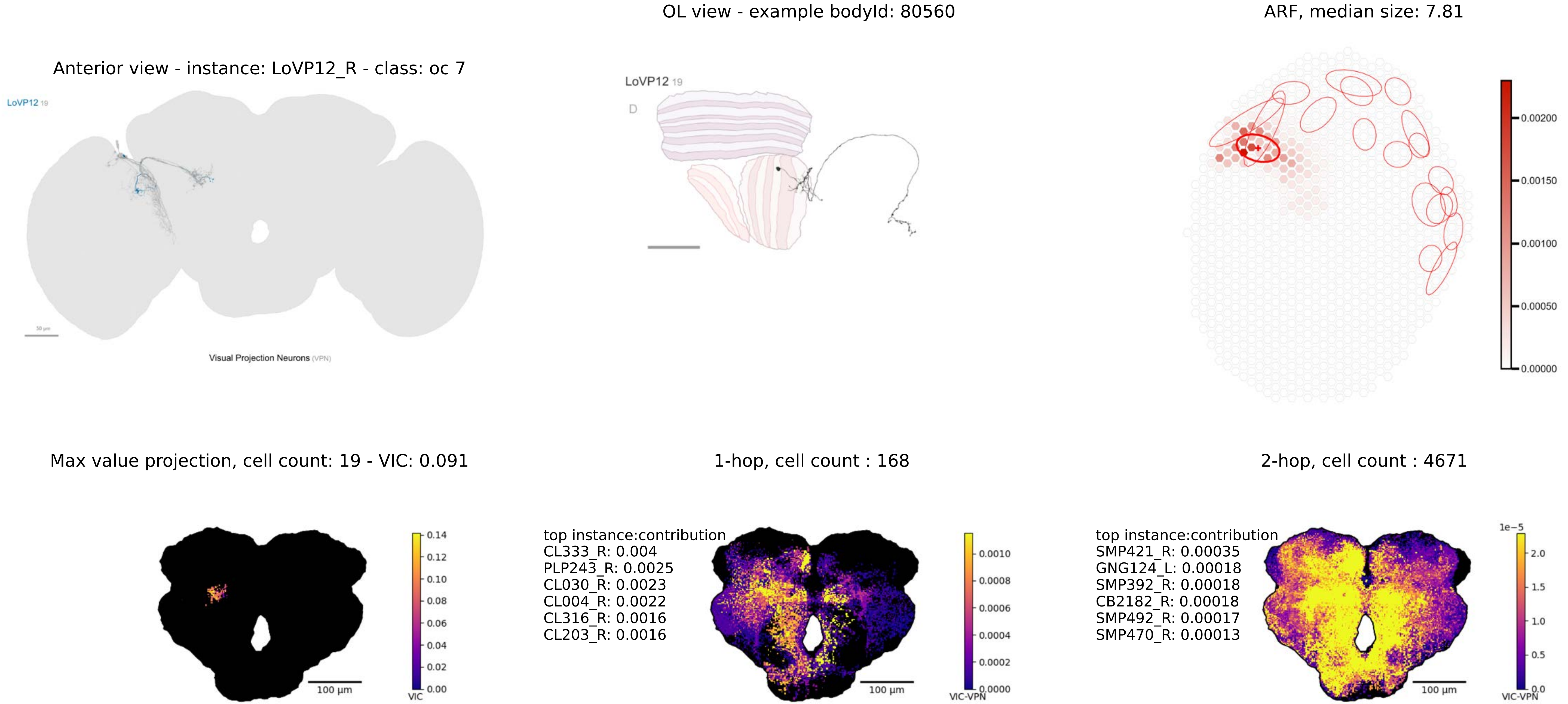

OL view - example bodyId: 80560
ARF, median size: 7.81
Anterior view - instance: LoVP12_R - class: oc 7
LoVP12 19
Visual Projection Neurons (VPN)
LoVP12 19
D
0.00200
0.00150
0.00100
0.00050
0.00000
Max value projection, cell count: 19 - VIC: 0.091
1-hop, cell count : 168
2-hop, cell count : 4671
100 µm
0.14
0.12
0.10
0.08
0.06
0.04
0.02
0.00
VIC
top instance:contribution
CL333_R: 0.004
PLP243_R: 0.0025
CL030_R: 0.0023
CL004_R: 0.0022
CL316_R: 0.0016
CL203_R: 0.0016
100 µm
0.0010
0.0008
0.0006
0.0004
0.0002
0.0000
VIC-VPN
top instance:contribution
SMP421_R: 0.00035
GNG124_L: 0.00018
SMP392_R: 0.00018
CB2182_R: 0.00018
SMP492_R: 0.00017
SMP470_R: 0.00013
100 µm
1e−5
2.0
1.5
1.0
0.5
0.0
VIC-VPN

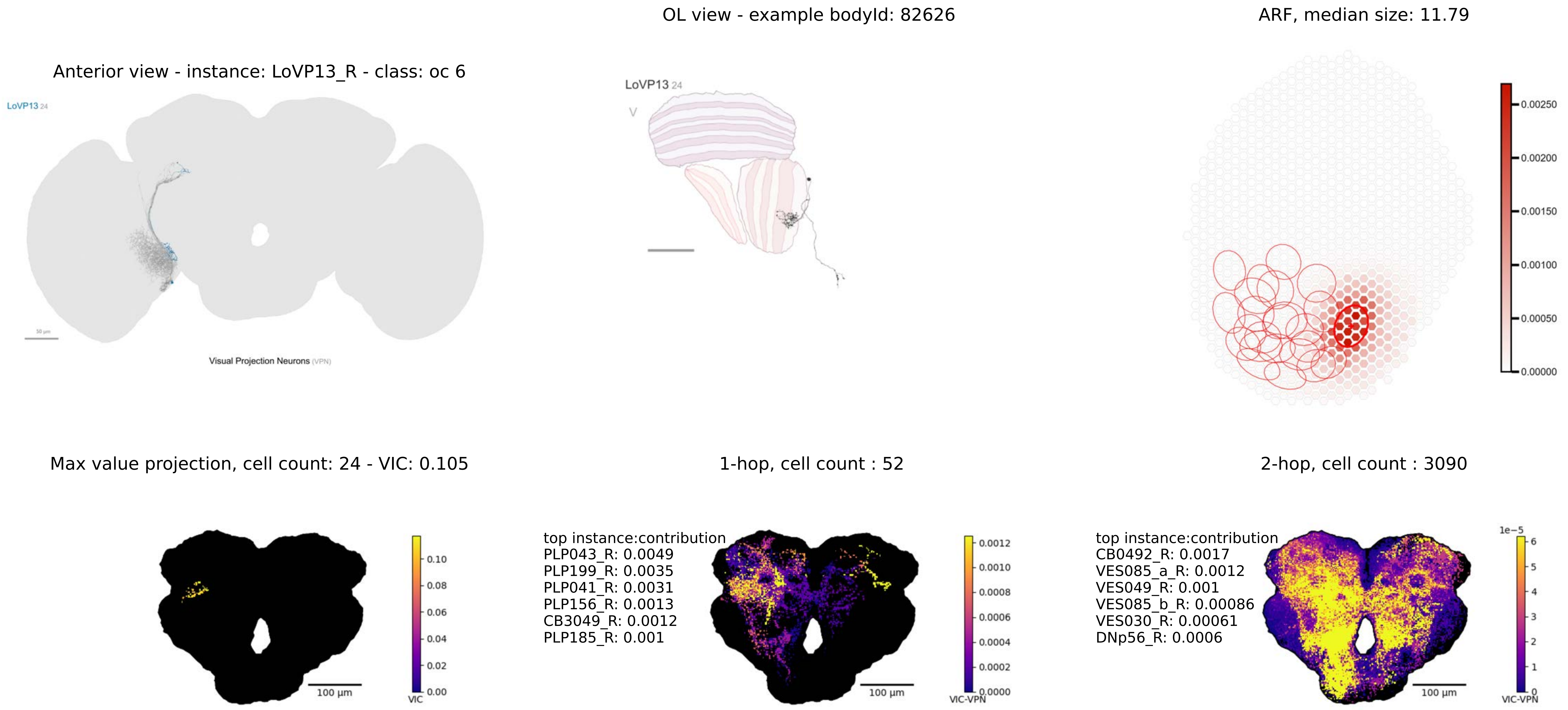
OL view - example bodyId: 82626
ARF, median size: 11.79
Anterior view - instance: LoVP13_R - class: oc 6
LoVP13 24
Visual Projection Neurons (VPN)
LoVP13 24
V
0.00250
0.00200
0.00150
0.00100
0.00050
0.00000
Max value projection, cell count: 24 - VIC: 0.105
1-hop, cell count : 52
2-hop, cell count : 3090
0.10
0.08
0.06
0.04
0.02
0.00
VIC
100 µm
top instance:contribution
PLP043_R: 0.0049
PLP199_R: 0.0035
PLP041_R: 0.0031
PLP156_R: 0.0013
CB3049_R: 0.0012
PLP185_R: 0.001
0.0012
0.0010
0.0008
0.0006
0.0004
0.0002
0.0000
VIC-VPN
100 µm
top instance:contribution
CB0492_R: 0.0017
VES085_a_R: 0.0012
VES049_R: 0.001
VES085_b_R: 0.00086
VES030_R: 0.00061
DNp56_R: 0.0006
1e−5
6
5
4
3
2
1
0
VIC-VPN
100 µm

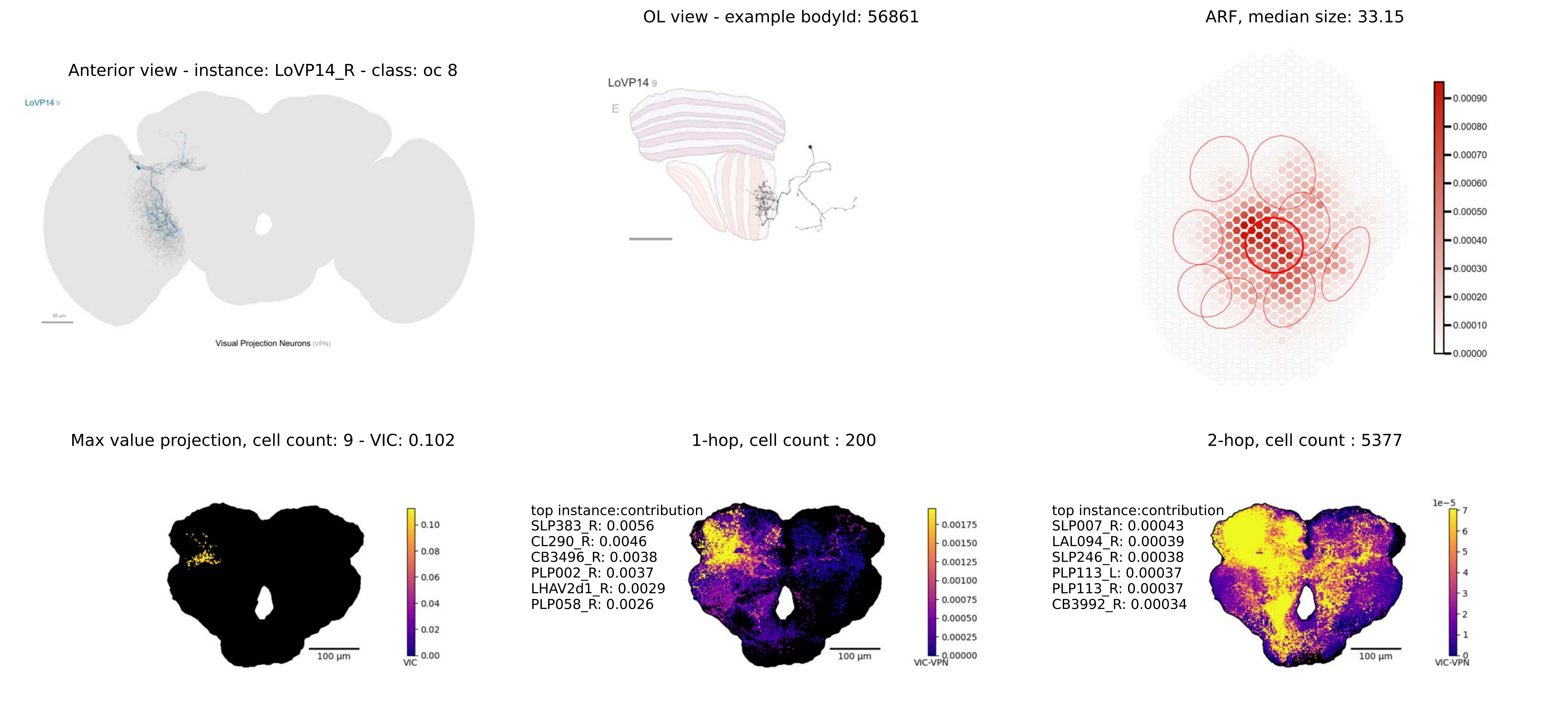

OL view - example bodyId: 56861
ARF, median size: 33.15
Anterior view - instance: LoVP14_R - class: oc 8
LoVP14 9
Visual Projection Neurons (VPN)
LoVP14 9
E
0.00090
0.00080
0.00070
0.00060
0.00050
0.00040
0.00030
0.00020
0.00010
0.00000
Max value projection, cell count: 9 - VIC: 0.102
1-hop, cell count : 200
2-hop, cell count : 5377
0.10
0.08
0.06
0.04
0.02
0.00
VIC
100 µm
top instance:contribution
SLP383_R: 0.0056
CL290_R: 0.0046
CB3496_R: 0.0038
PLP002_R: 0.0037
LHAV2d1_R: 0.0029
PLP058_R: 0.0026
0.00175
0.00150
0.00125
0.00100
0.00075
0.00050
0.00025
0.00000
VIC-VPN
100 µm
top instance:contribution
SLP007_R: 0.00043
LAL094_R: 0.00039
SLP246_R: 0.00038
PLP113_L: 0.00037
PLP113_R: 0.00037
CB3992_R: 0.00034
1e−5
7
6
5
4
3
2
1
0
VIC-VPN
100 µm

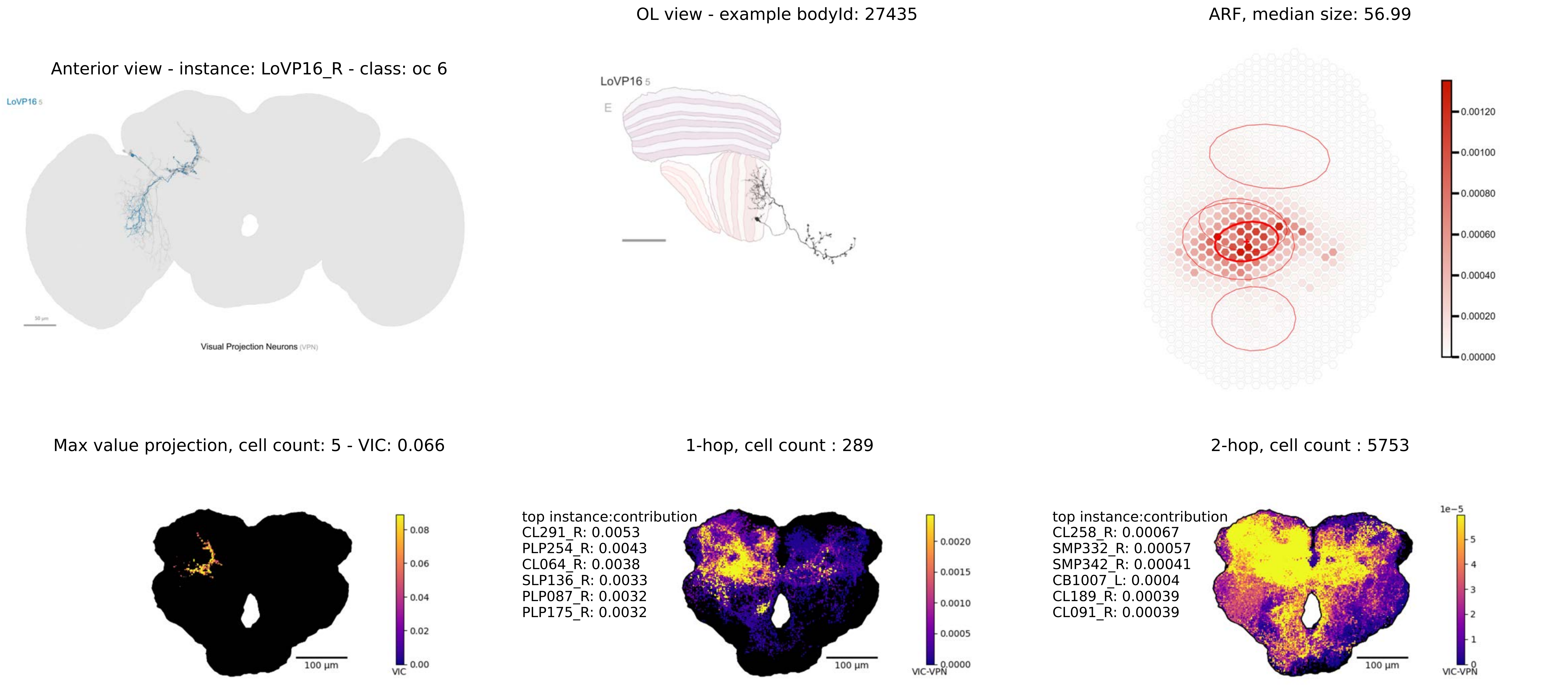
OL view - example bodyId: 27435
ARF, median size: 56.99
Anterior view - instance: LoVP16_R - class: oc 6
LoVP16 5
Visual Projection Neurons (VPN)
0.00120
0.00100
0.00080
0.00060
0.00040
0.00020
0.00000
Max value projection, cell count: 5 - VIC: 0.066
1-hop, cell count : 289
2-hop, cell count : 5753
top instance:contribution
CL291_R: 0.0053
PLP254_R: 0.0043
CL064_R: 0.0038
SLP136_R: 0.0033
PLP087_R: 0.0032
PLP175_R: 0.0032
top instance:contribution
CL258_R: 0.00067
SMP332_R: 0.00057
SMP342_R: 0.00041
CB1007_L: 0.0004
CL189_R: 0.00039
CL091_R: 0.00039
100 µm
VIC
VIC-VPN
1e−5

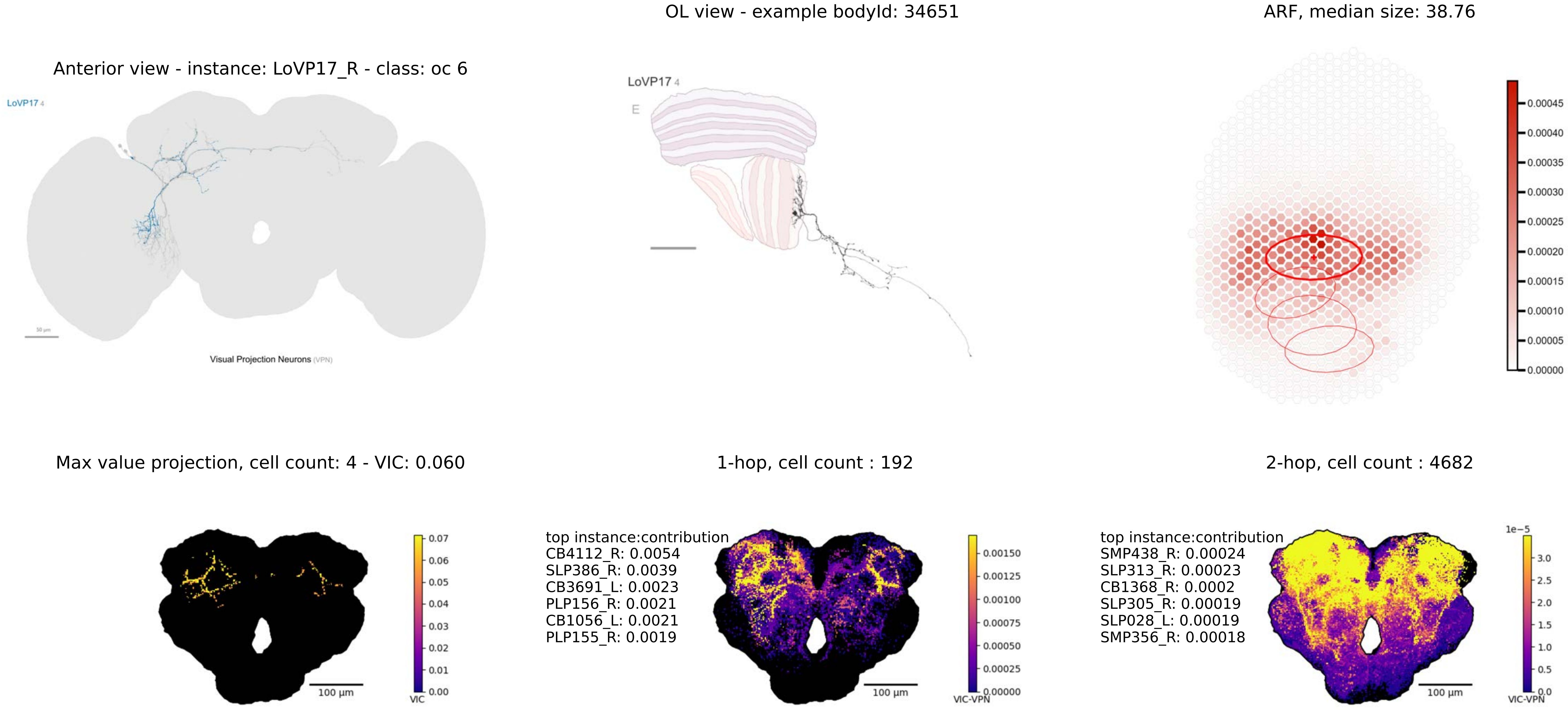

OL view - example bodyId: 34651
ARF, median size: 38.76
Anterior view - instance: LoVP17_R - class: oc 6
LoVP17 4
Visual Projection Neurons (VPN)
LoVP17 4
E
0.00045
0.00040
0.00035
0.00030
0.00025
0.00020
0.00015
0.00010
0.00005
0.00000
Max value projection, cell count: 4 - VIC: 0.060
1-hop, cell count : 192
2-hop, cell count : 4682
100 µm
VIC
0.07
0.06
0.05
0.04
0.03
0.02
0.01
0.00
top instance:contribution
CB4112_R: 0.0054
SLP386_R: 0.0039
CB3691_L: 0.0023
PLP156_R: 0.0021
CB1056_L: 0.0021
PLP155_R: 0.0019
100 µm
VIC-VPN
0.00150
0.00125
0.00100
0.00075
0.00050
0.00025
0.00000
top instance:contribution
SMP438_R: 0.00024
SLP313_R: 0.00023
CB1368_R: 0.0002
SLP305_R: 0.00019
SLP028_L: 0.00019
SMP356_R: 0.00018
100 µm
1e−5
3.0
2.5
2.0
1.5
1.0
0.5
0.0
VIC-VPN

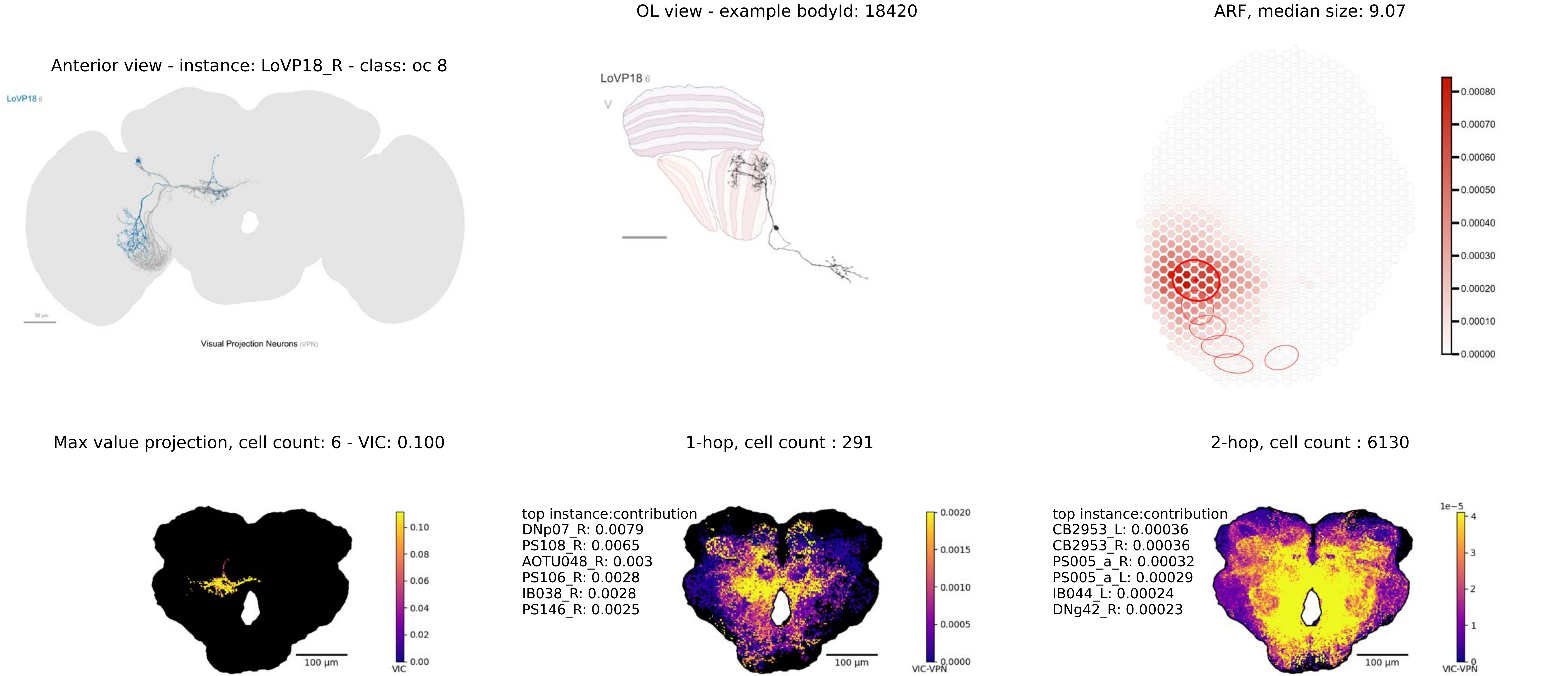

Anterior view - instance: LoVP18_R - class: oc 8
LoVP18 6
50 µm
Visual Projection Neurons (VPN)
OL view - example bodyId: 18420
LoVP18 6
V
ARF, median size: 9.07
0.00080
0.00070
0.00060
0.00050
0.00040
0.00030
0.00020
0.00010
0.00000
Max value projection, cell count: 6 - VIC: 0.100
0.10
0.08
0.06
0.04
0.02
0.00
VIC
100 µm
1-hop, cell count : 291
top instance:contribution
DNp07_R: 0.0079
PS108_R: 0.0065
AOTU048_R: 0.003
PS106_R: 0.0028
IB038_R: 0.0028
PS146_R: 0.0025
0.0020
0.0015
0.0010
0.0005
0.0000
VIC-VPN
100 µm
2-hop, cell count : 6130
top instance:contribution
CB2953_L: 0.00036
CB2953_R: 0.00036
PS005_a_R: 0.00032
PS005_a_L: 0.00029
IB044_L: 0.00024
DNg42_R: 0.00023
1e−5
4
3
2
1
0
VIC-VPN
100 µm

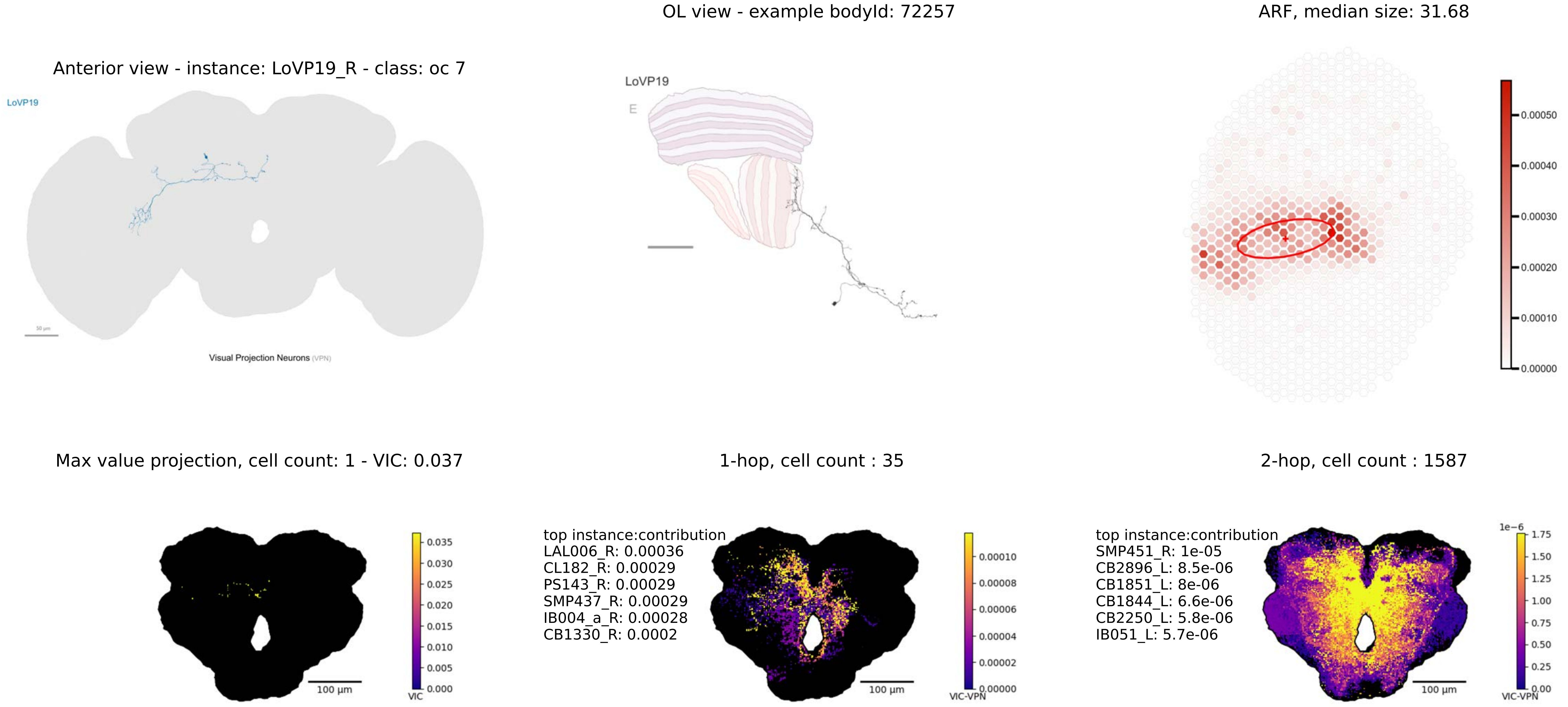

OL view - example bodyId: 72257
ARF, median size: 31.68
Anterior view - instance: LoVP19_R - class: oc 7
LoVP19
Visual Projection Neurons (VPN)
LoVP19
E
0.00050
0.00040
0.00030
0.00020
0.00010
0.00000
Max value projection, cell count: 1 - VIC: 0.037
1-hop, cell count : 35
2-hop, cell count : 1587
0.035
0.030
0.025
0.020
0.015
0.010
0.005
0.000
VIC
100 µm
top instance:contribution
LAL006_R: 0.00036
CL182_R: 0.00029
PS143_R: 0.00029
SMP437_R: 0.00029
IB004_a_R: 0.00028
CB1330_R: 0.0002
0.00010
0.00008
0.00006
0.00004
0.00002
0.00000
VIC-VPN
100 µm
top instance:contribution
SMP451_R: 1e-05
CB2896_L: 8.5e-06
CB1851_L: 8e-06
CB1844_L: 6.6e-06
CB2250_L: 5.8e-06
IB051_L: 5.7e-06
1e−6
1.75
1.50
1.25
1.00
0.75
0.50
0.25
0.00
VIC-VPN
100 µm

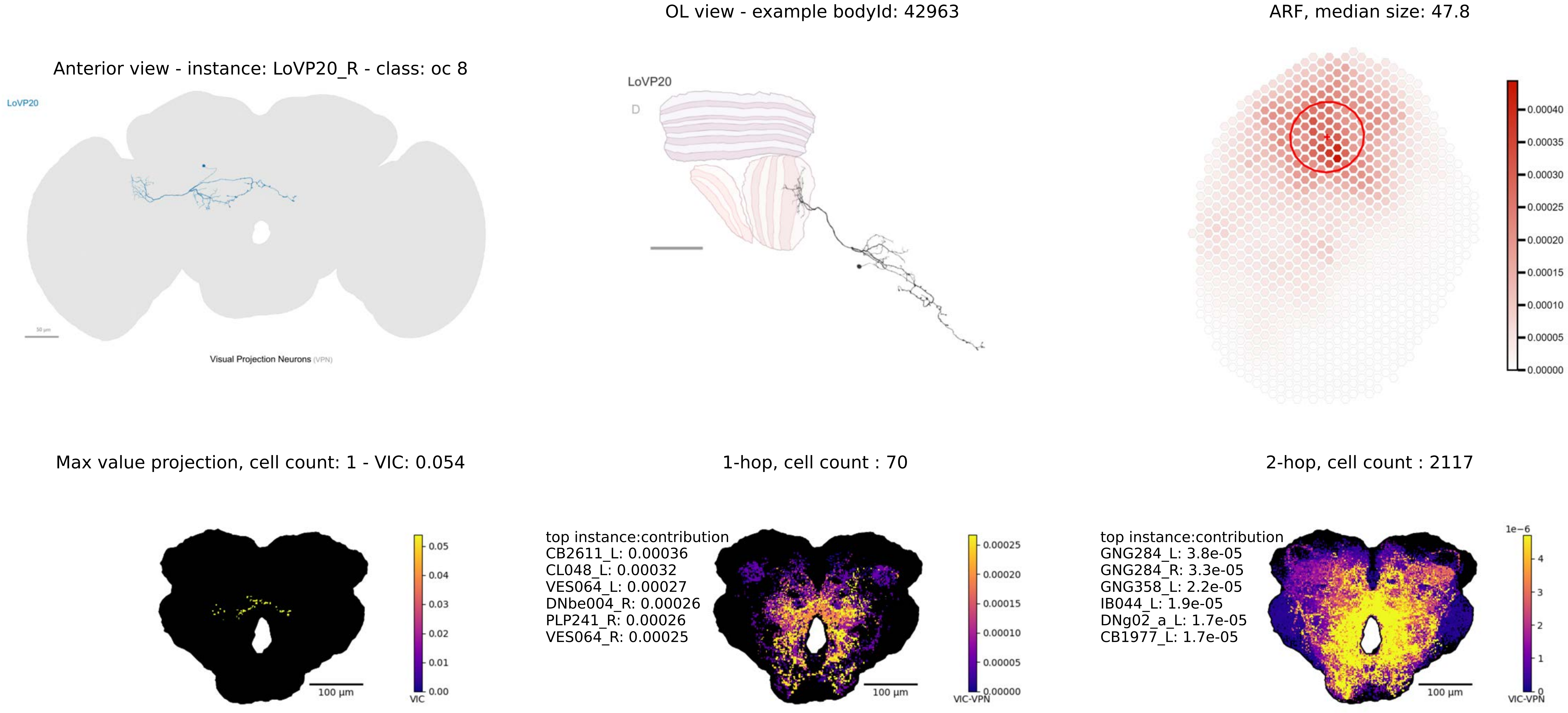
OL view - example bodyId: 42963
ARF, median size: 47.8
Anterior view - instance: LoVP20_R - class: oc 8
LoVP20
Visual Projection Neurons (VPN)
LoVP20
D
0.00040
0.00035
0.00030
0.00025
0.00020
0.00015
0.00010
0.00005
0.00000
Max value projection, cell count: 1 - VIC: 0.054
1-hop, cell count : 70
2-hop, cell count : 2117
0.05
0.04
0.03
0.02
0.01
0.00
VIC
100 µm
top instance:contribution
CB2611_L: 0.00036
CL048_L: 0.00032
VES064_L: 0.00027
DNbe004_R: 0.00026
PLP241_R: 0.00026
VES064_R: 0.00025
0.00025
0.00020
0.00015
0.00010
0.00005
0.00000
VIC-VPN
100 µm
top instance:contribution
GNG284_L: 3.8e-05
GNG284_R: 3.3e-05
GNG358_L: 2.2e-05
IB044_L: 1.9e-05
DNg02_a_L: 1.7e-05
CB1977_L: 1.7e-05
1e−6
4
3
2
1
0
VIC-VPN
100 µm

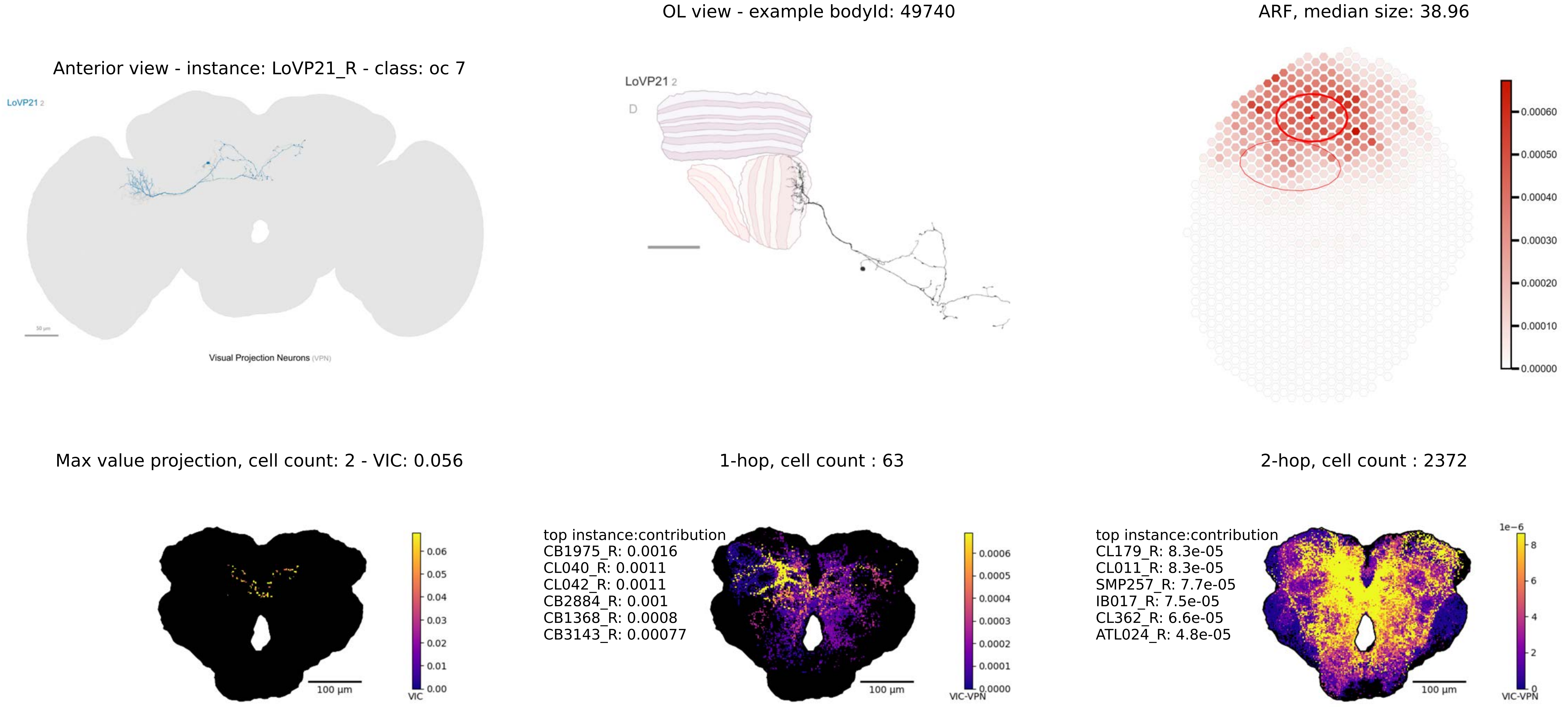
Anterior view - instance: LoVP21_R - class: oc 7
LoVP21 2
Visual Projection Neurons (VPN)
OL view - example bodyId: 49740
LoVP21 2
D
ARF, median size: 38.96
0.00060
0.00050
0.00040
0.00030
0.00020
0.00010
0.00000
Max value projection, cell count: 2 - VIC: 0.056
100 µm
VIC
1-hop, cell count : 63
top instance:contribution
CB1975_R: 0.0016
CL040_R: 0.0011
CL042_R: 0.0011
CB2884_R: 0.001
CB1368_R: 0.0008
CB3143_R: 0.00077
100 µm
VIC-VPN
2-hop, cell count : 2372
top instance:contribution
CL179_R: 8.3e-05
CL011_R: 8.3e-05
SMP257_R: 7.7e-05
IB017_R: 7.5e-05
CL362_R: 6.6e-05
ATL024_R: 4.8e-05
1e−6
100 µm
VIC-VPN

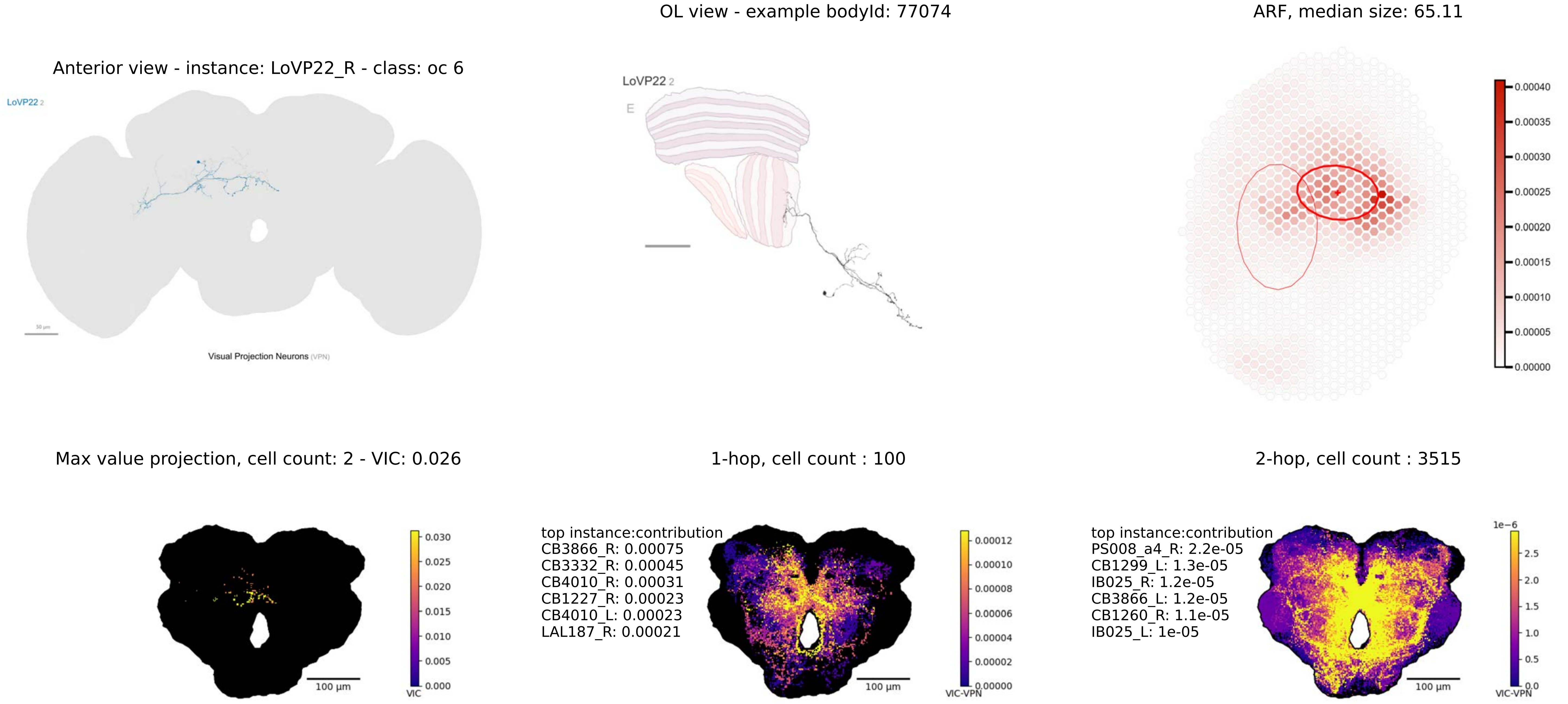

Anterior view - instance: LoVP22_R - class: oc 6
LoVP22 2
Visual Projection Neurons (VPN)
OL view - example bodyId: 77074
LoVP22 2
E
ARF, median size: 65.11
0.00040
0.00035
0.00030
0.00025
0.00020
0.00015
0.00010
0.00005
0.00000
Max value projection, cell count: 2 - VIC: 0.026
0.030
0.025
0.020
0.015
0.010
0.005
0.000
VIC
100 µm
1-hop, cell count : 100
top instance:contribution
CB3866_R: 0.00075
CB3332_R: 0.00045
CB4010_R: 0.00031
CB1227_R: 0.00023
CB4010_L: 0.00023
LAL187_R: 0.00021
0.00012
0.00010
0.00008
0.00006
0.00004
0.00002
0.00000
VIC-VPN
100 µm
2-hop, cell count : 3515
top instance:contribution
PS008_a4_R: 2.2e-05
CB1299_L: 1.3e-05
IB025_R: 1.2e-05
CB3866_L: 1.2e-05
CB1260_R: 1.1e-05
IB025_L: 1e-05
1e−6
2.5
2.0
1.5
1.0
0.5
0.0
VIC-VPN
100 µm

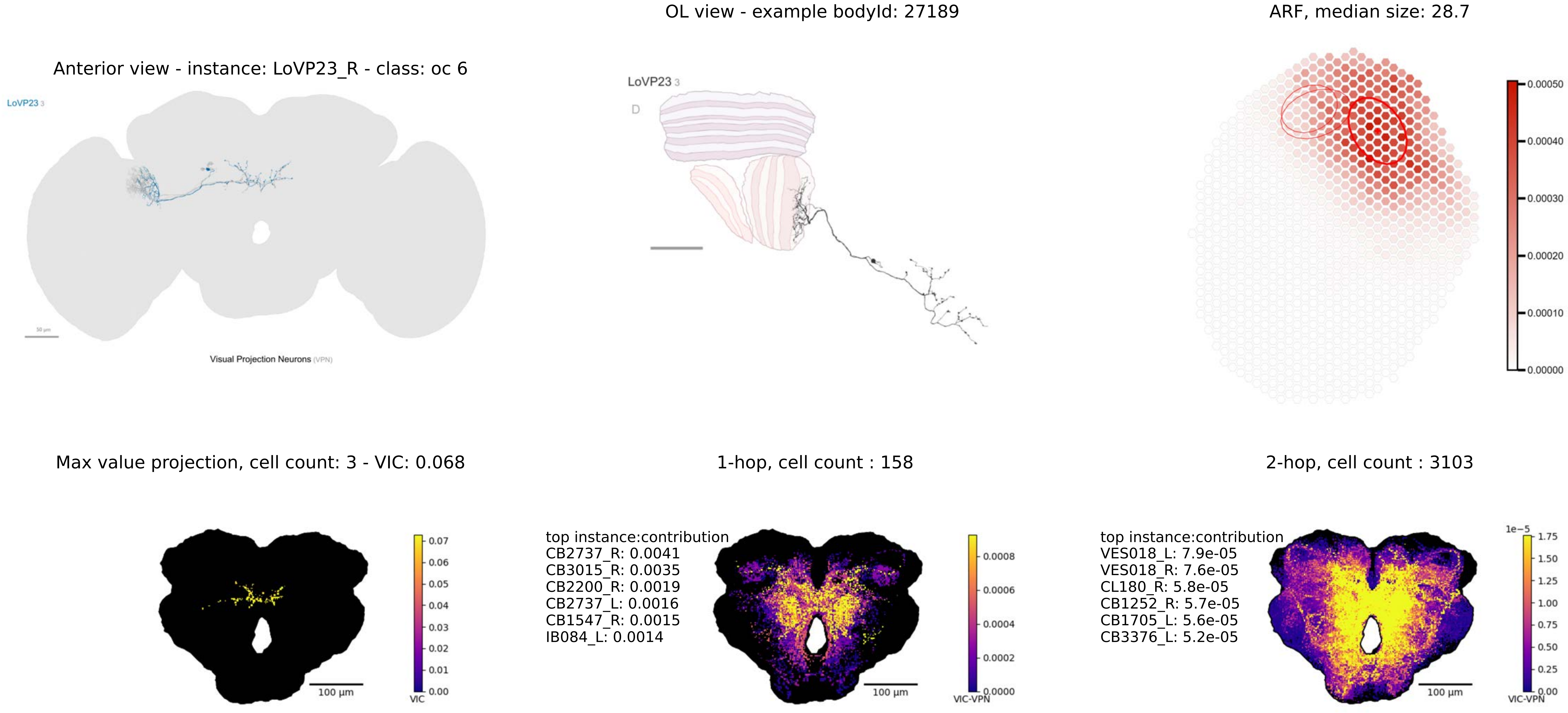

Anterior view - instance: LoVP23_R - class: oc 6
LoVP23 3
50 µm
Visual Projection Neurons (VPN)
OL view - example bodyId: 27189
LoVP23 3
D
ARF, median size: 28.7
0.00050
0.00040
0.00030
0.00020
0.00010
0.00000
Max value projection, cell count: 3 - VIC: 0.068
100 µm
VIC
0.07
0.06
0.05
0.04
0.03
0.02
0.01
0.00
1-hop, cell count : 158
top instance:contribution
CB2737_R: 0.0041
CB3015_R: 0.0035
CB2200_R: 0.0019
CB2737_L: 0.0016
CB1547_R: 0.0015
IB084_L: 0.0014
100 µm
VIC-VPN
0.0008
0.0006
0.0004
0.0002
0.0000
2-hop, cell count : 3103
top instance:contribution
VES018_L: 7.9e-05
VES018_R: 7.6e-05
CL180_R: 5.8e-05
CB1252_R: 5.7e-05
CB1705_L: 5.6e-05
CB3376_L: 5.2e-05
100 µm
1e−5
VIC-VPN
1.75
1.50
1.25
1.00
0.75
0.50
0.25
0.00

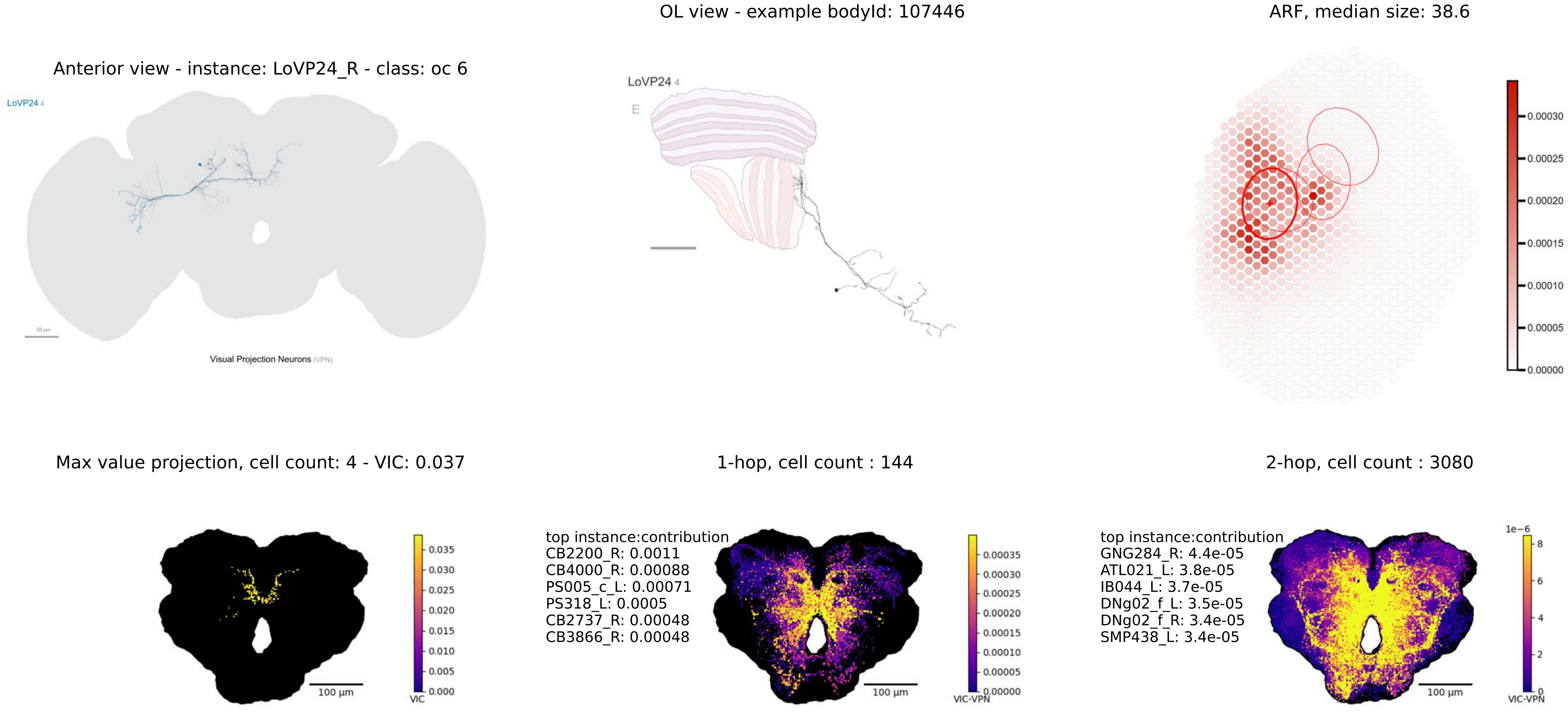

OL view - example bodyId: 107446
ARF, median size: 38.6
Anterior view - instance: LoVP24_R - class: oc 6
LoVP24 4
Visual Projection Neurons (VPN)
LoVP24 4
E
0.00030
0.00025
0.00020
0.00015
0.00010
0.00005
0.00000
Max value projection, cell count: 4 - VIC: 0.037
1-hop, cell count : 144
2-hop, cell count : 3080
0.035
0.030
0.025
0.020
0.015
0.010
0.005
0.000
VIC
100 µm
top instance:contribution
CB2200_R: 0.0011
CB4000_R: 0.00088
PS005_c_L: 0.00071
PS318_L: 0.0005
CB2737_R: 0.00048
CB3866_R: 0.00048
0.00035
0.00030
0.00025
0.00020
0.00015
0.00010
0.00005
0.00000
VIC-VPN
100 µm
top instance:contribution
GNG284_R: 4.4e-05
ATL021_L: 3.8e-05
IB044_L: 3.7e-05
DNg02_f_L: 3.5e-05
DNg02_f_R: 3.4e-05
SMP438_L: 3.4e-05
1e−6
8
6
4
2
0
VIC-VPN
100 µm

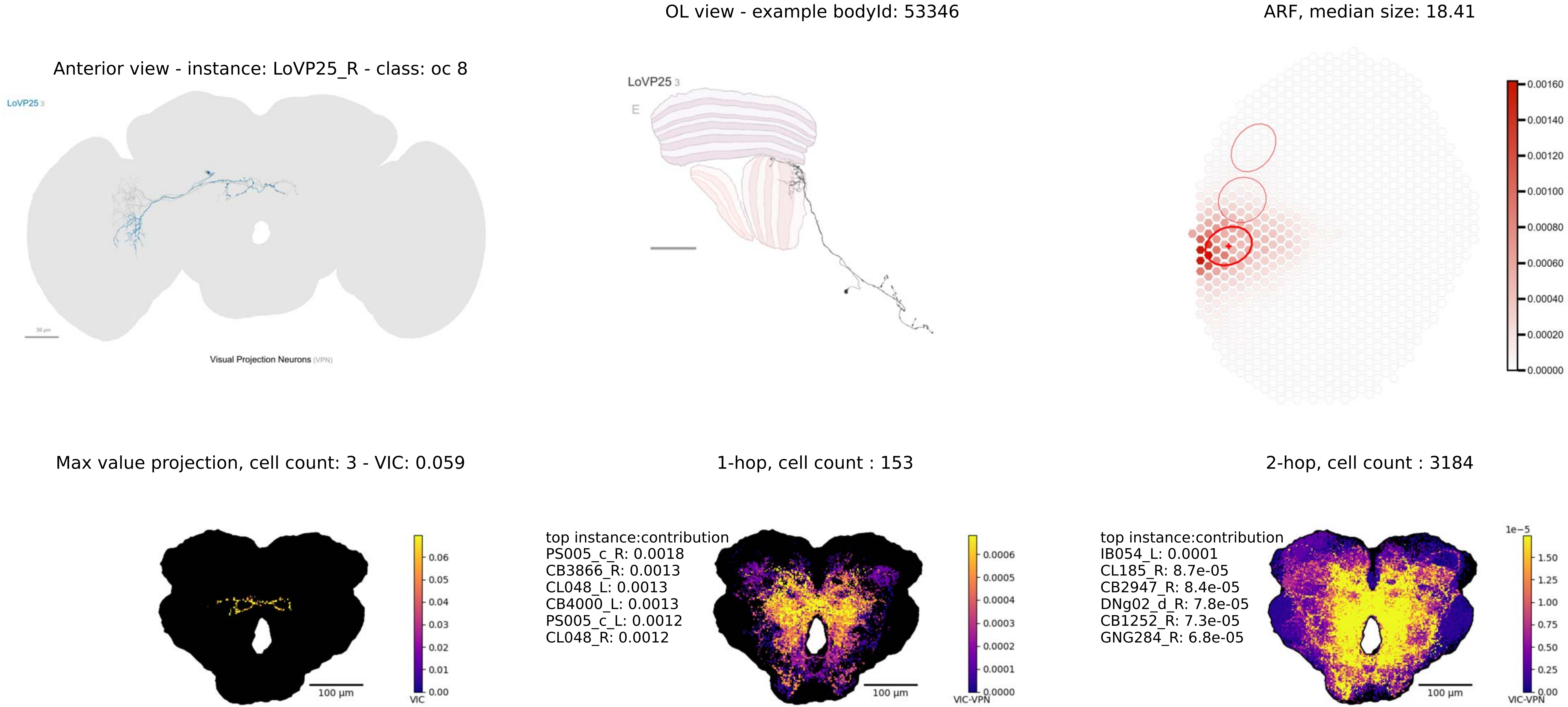
OL view - example bodyId: 53346
ARF, median size: 18.41
Anterior view - instance: LoVP25_R - class: oc 8
LoVP25 3
E
Visual Projection Neurons (VPN)
0.00160
0.00140
0.00120
0.00100
0.00080
0.00060
0.00040
0.00020
0.00000
Max value projection, cell count: 3 - VIC: 0.059
1-hop, cell count : 153
2-hop, cell count : 3184
top instance:contribution
PS005_c_R: 0.0018
CB3866_R: 0.0013
CL048_L: 0.0013
CB4000_L: 0.0013
PS005_c_L: 0.0012
CL048_R: 0.0012
top instance:contribution
IB054_L: 0.0001
CL185_R: 8.7e-05
CB2947_R: 8.4e-05
DNg02_d_R: 7.8e-05
CB1252_R: 7.3e-05
GNG284_R: 6.8e-05
100 µm
VIC
VIC-VPN
1e−5

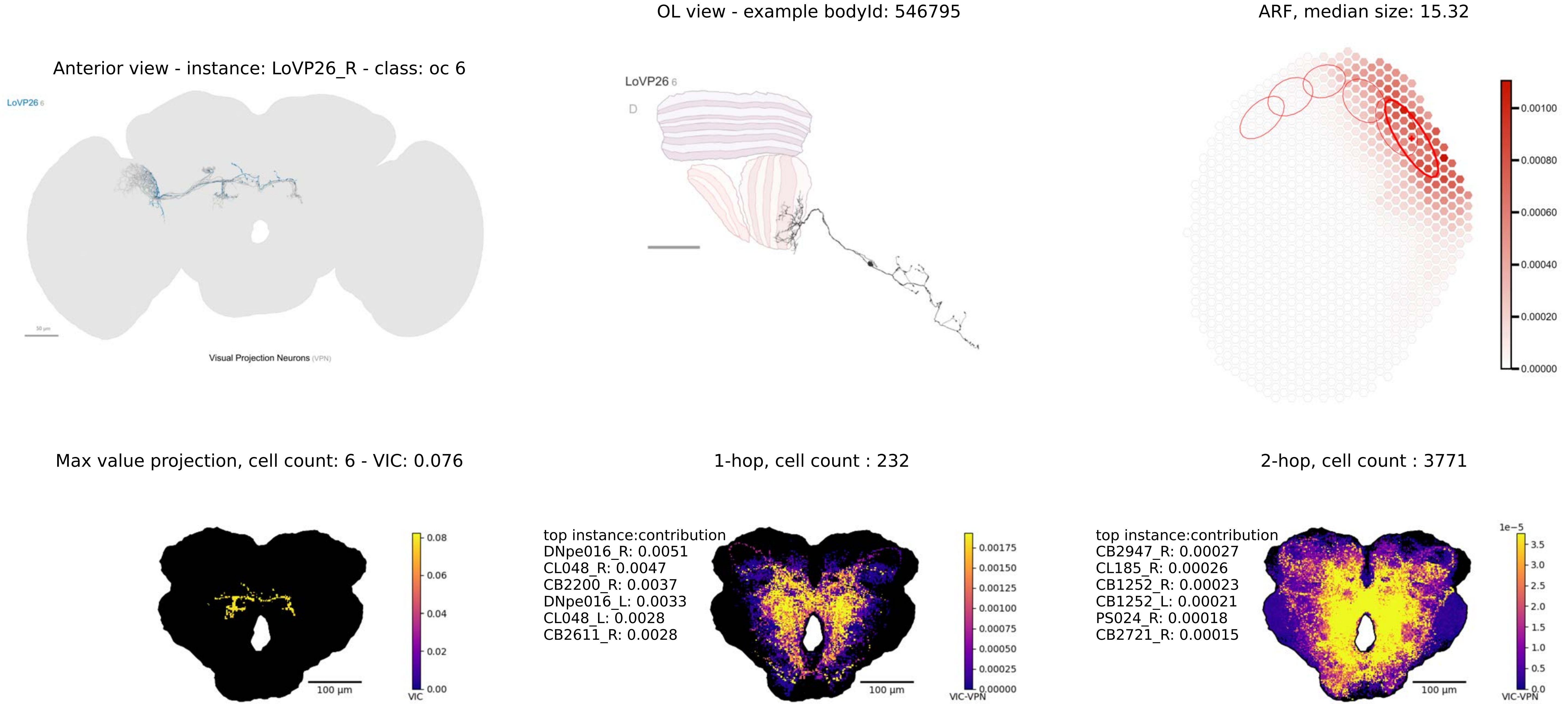
OL view - example bodyId: 546795
ARF, median size: 15.32
Anterior view - instance: LoVP26_R - class: oc 6
LoVP26 6
Visual Projection Neurons (VPN)
LoVP26 6
D
0.00100
0.00080
0.00060
0.00040
0.00020
0.00000
Max value projection, cell count: 6 - VIC: 0.076
1-hop, cell count : 232
2-hop, cell count : 3771
0.08
0.06
0.04
0.02
0.00
VIC
100 µm
top instance:contribution
DNpe016_R: 0.0051
CL048_R: 0.0047
CB2200_R: 0.0037
DNpe016_L: 0.0033
CL048_L: 0.0028
CB2611_R: 0.0028
0.00175
0.00150
0.00125
0.00100
0.00075
0.00050
0.00025
0.00000
VIC-VPN
100 µm
top instance:contribution
CB2947_R: 0.00027
CL185_R: 0.00026
CB1252_R: 0.00023
CB1252_L: 0.00021
PS024_R: 0.00018
CB2721_R: 0.00015
1e−5
3.5
3.0
2.5
2.0
1.5
1.0
0.5
0.0
VIC-VPN
100 µm

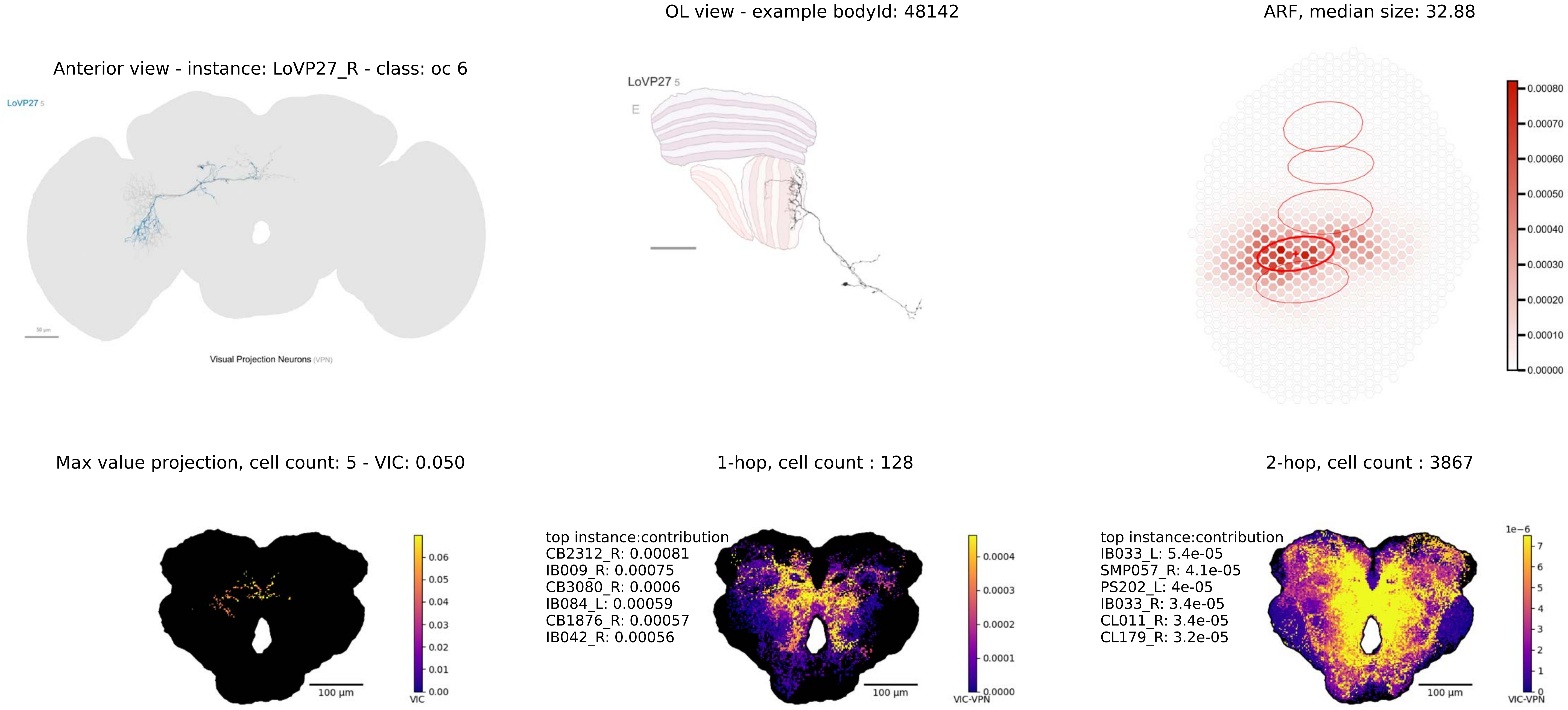
OL view - example bodyId: 48142
ARF, median size: 32.88
Anterior view - instance: LoVP27_R - class: oc 6
LoVP27 5
Visual Projection Neurons (VPN)
LoVP27 5
E
Max value projection, cell count: 5 - VIC: 0.050
VIC
100 µm
1-hop, cell count : 128
top instance:contribution
CB2312_R: 0.00081
IB009_R: 0.00075
CB3080_R: 0.0006
IB084_L: 0.00059
CB1876_R: 0.00057
IB042_R: 0.00056
VIC-VPN
100 µm
2-hop, cell count : 3867
top instance:contribution
IB033_L: 5.4e-05
SMP057_R: 4.1e-05
PS202_L: 4e-05
IB033_R: 3.4e-05
CL011_R: 3.4e-05
CL179_R: 3.2e-05
1e−6
VIC-VPN
100 µm

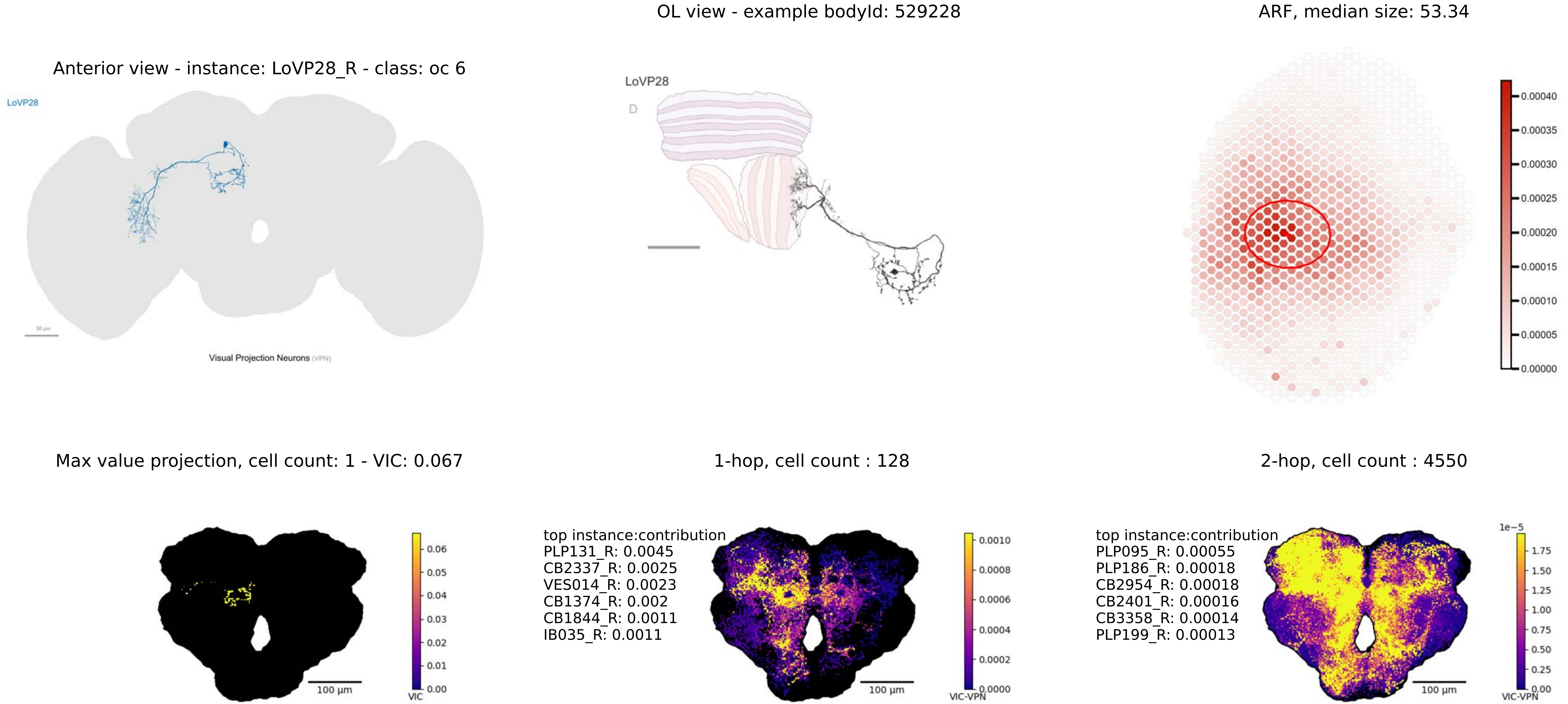

Anterior view - instance: LoVP28_R - class: oc 6
LoVP28
Visual Projection Neurons (VPN)
OL view - example bodyId: 529228
LoVP28
D
ARF, median size: 53.34
0.00040
0.00035
0.00030
0.00025
0.00020
0.00015
0.00010
0.00005
0.00000
Max value projection, cell count: 1 - VIC: 0.067
0.06
0.05
0.04
0.03
0.02
0.01
0.00
VIC
100 µm
1-hop, cell count : 128
top instance:contribution
PLP131_R: 0.0045
CB2337_R: 0.0025
VES014_R: 0.0023
CB1374_R: 0.002
CB1844_R: 0.0011
IB035_R: 0.0011
0.0010
0.0008
0.0006
0.0004
0.0002
0.0000
VIC-VPN
100 µm
2-hop, cell count : 4550
top instance:contribution
PLP095_R: 0.00055
PLP186_R: 0.00018
CB2954_R: 0.00018
CB2401_R: 0.00016
CB3358_R: 0.00014
PLP199_R: 0.00013
1e−5
1.75
1.50
1.25
1.00
0.75
0.50
0.25
0.00
VIC-VPN
100 µm

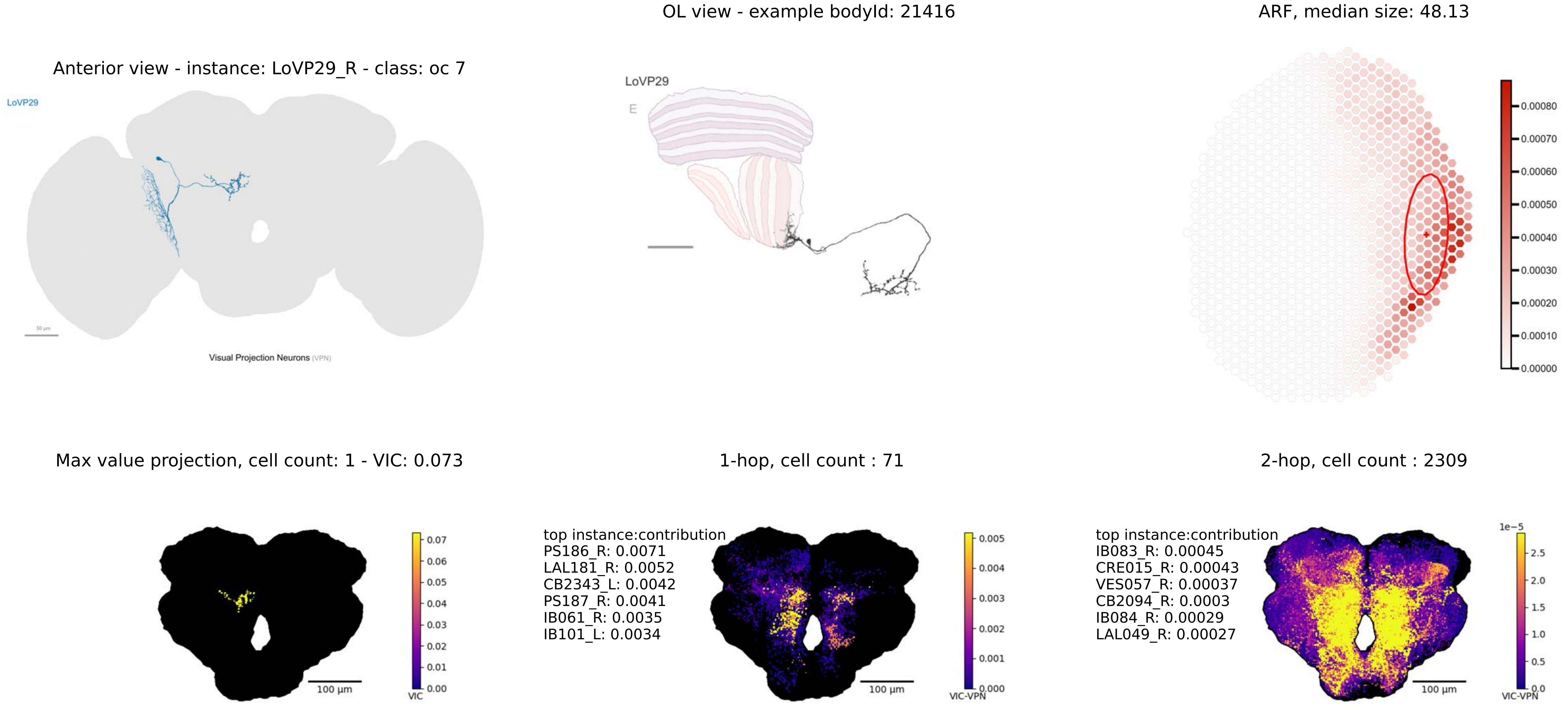
OL view - example bodyId: 21416
ARF, median size: 48.13
Anterior view - instance: LoVP29_R - class: oc 7
LoVP29
Visual Projection Neurons (VPN)
LoVP29
E
0.00080
0.00070
0.00060
0.00050
0.00040
0.00030
0.00020
0.00010
0.00000
Max value projection, cell count: 1 - VIC: 0.073
1-hop, cell count : 71
2-hop, cell count : 2309
100 µm
VIC
0.07
0.06
0.05
0.04
0.03
0.02
0.01
0.00
top instance:contribution
PS186_R: 0.0071
LAL181_R: 0.0052
CB2343_L: 0.0042
PS187_R: 0.0041
IB061_R: 0.0035
IB101_L: 0.0034
100 µm
VIC-VPN
0.005
0.004
0.003
0.002
0.001
0.000
top instance:contribution
IB083_R: 0.00045
CRE015_R: 0.00043
VES057_R: 0.00037
CB2094_R: 0.0003
IB084_R: 0.00029
LAL049_R: 0.00027
100 µm
1e−5
2.5
2.0
1.5
1.0
0.5
0.0
VIC-VPN

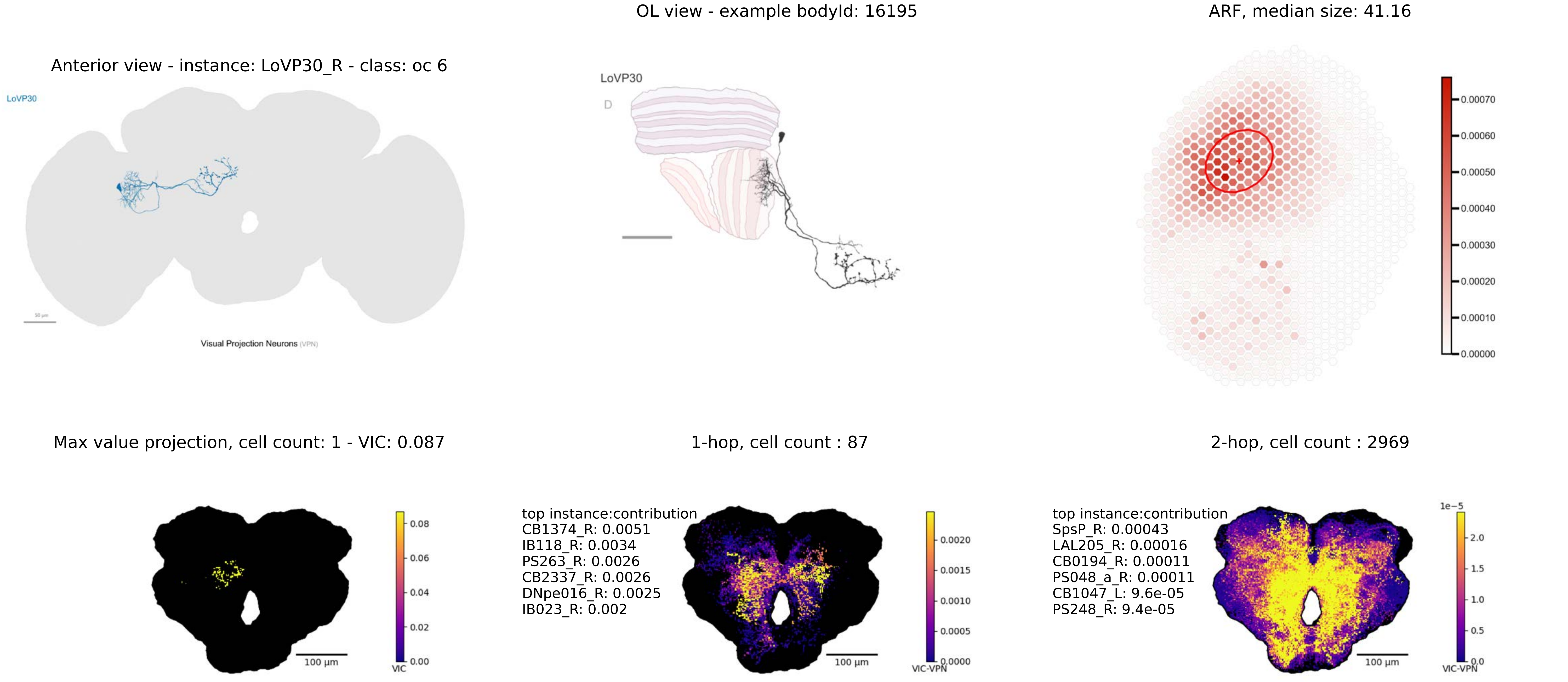

Anterior view - instance: LoVP30_R - class: oc 6
LoVP30
Visual Projection Neurons (VPN)
OL view - example bodyId: 16195
LoVP30
D
ARF, median size: 41.16
0.00070
0.00060
0.00050
0.00040
0.00030
0.00020
0.00010
0.00000
Max value projection, cell count: 1 - VIC: 0.087
0.08
0.06
0.04
0.02
0.00
VIC
100 µm
1-hop, cell count : 87
top instance:contribution
CB1374_R: 0.0051
IB118_R: 0.0034
PS263_R: 0.0026
CB2337_R: 0.0026
DNpe016_R: 0.0025
IB023_R: 0.002
0.0020
0.0015
0.0010
0.0005
0.0000
VIC-VPN
100 µm
2-hop, cell count : 2969
top instance:contribution
SpsP_R: 0.00043
LAL205_R: 0.00016
CB0194_R: 0.00011
PS048_a_R: 0.00011
CB1047_L: 9.6e-05
PS248_R: 9.4e-05
1e−5
2.0
1.5
1.0
0.5
0.0
VIC-VPN
100 µm

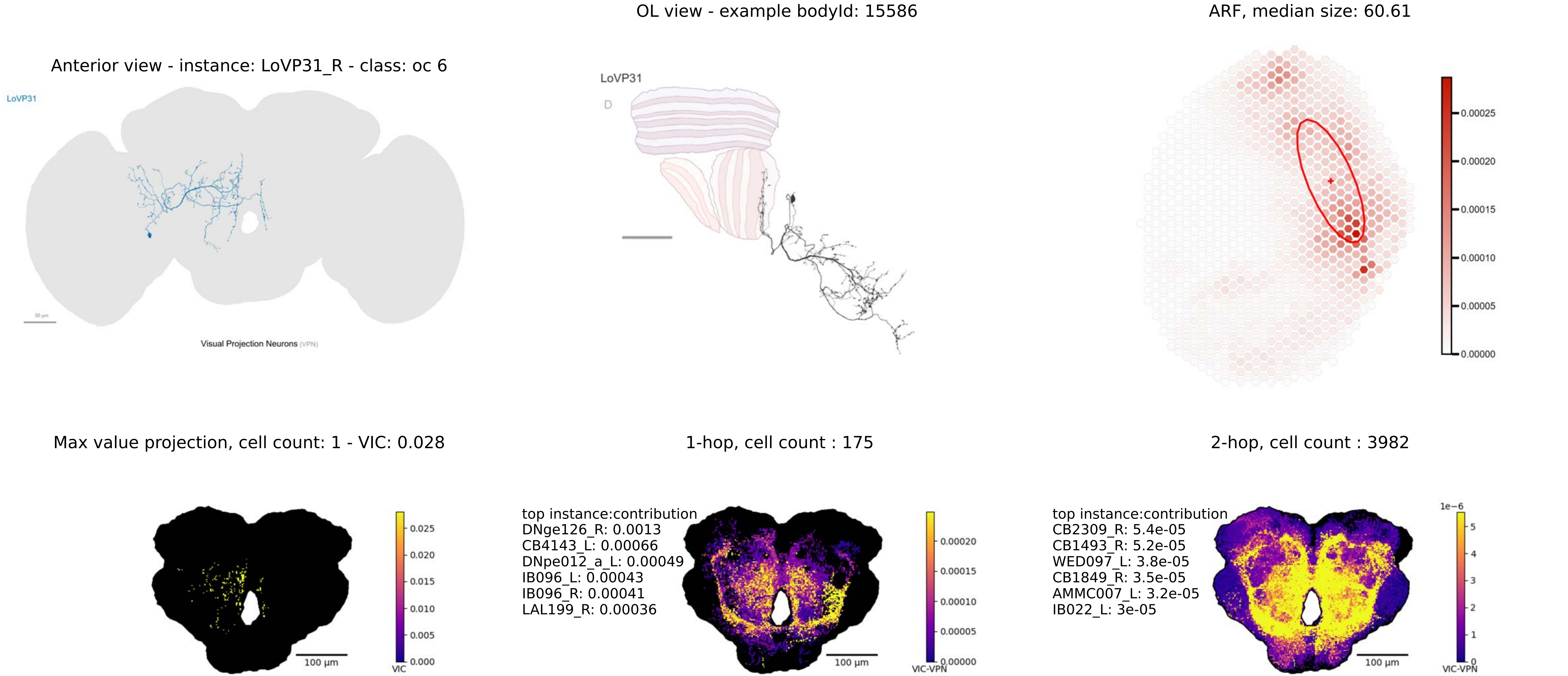
OL view - example bodyId: 15586
ARF, median size: 60.61
Anterior view - instance: LoVP31_R - class: oc 6
LoVP31
Visual Projection Neurons (VPN)
LoVP31
D
0.00025
0.00020
0.00015
0.00010
0.00005
0.00000
Max value projection, cell count: 1 - VIC: 0.028
1-hop, cell count : 175
2-hop, cell count : 3982
0.025
0.020
0.015
0.010
0.005
0.000
VIC
100 µm
top instance:contribution
DNge126_R: 0.0013
CB4143_L: 0.00066
DNpe012_a_L: 0.00049
IB096_L: 0.00043
IB096_R: 0.00041
LAL199_R: 0.00036
0.00020
0.00015
0.00010
0.00005
0.00000
VIC-VPN
100 µm
top instance:contribution
CB2309_R: 5.4e-05
CB1493_R: 5.2e-05
WED097_L: 3.8e-05
CB1849_R: 3.5e-05
AMMC007_L: 3.2e-05
IB022_L: 3e-05
1e−6
5
4
3
2
1
0
VIC-VPN
100 µm

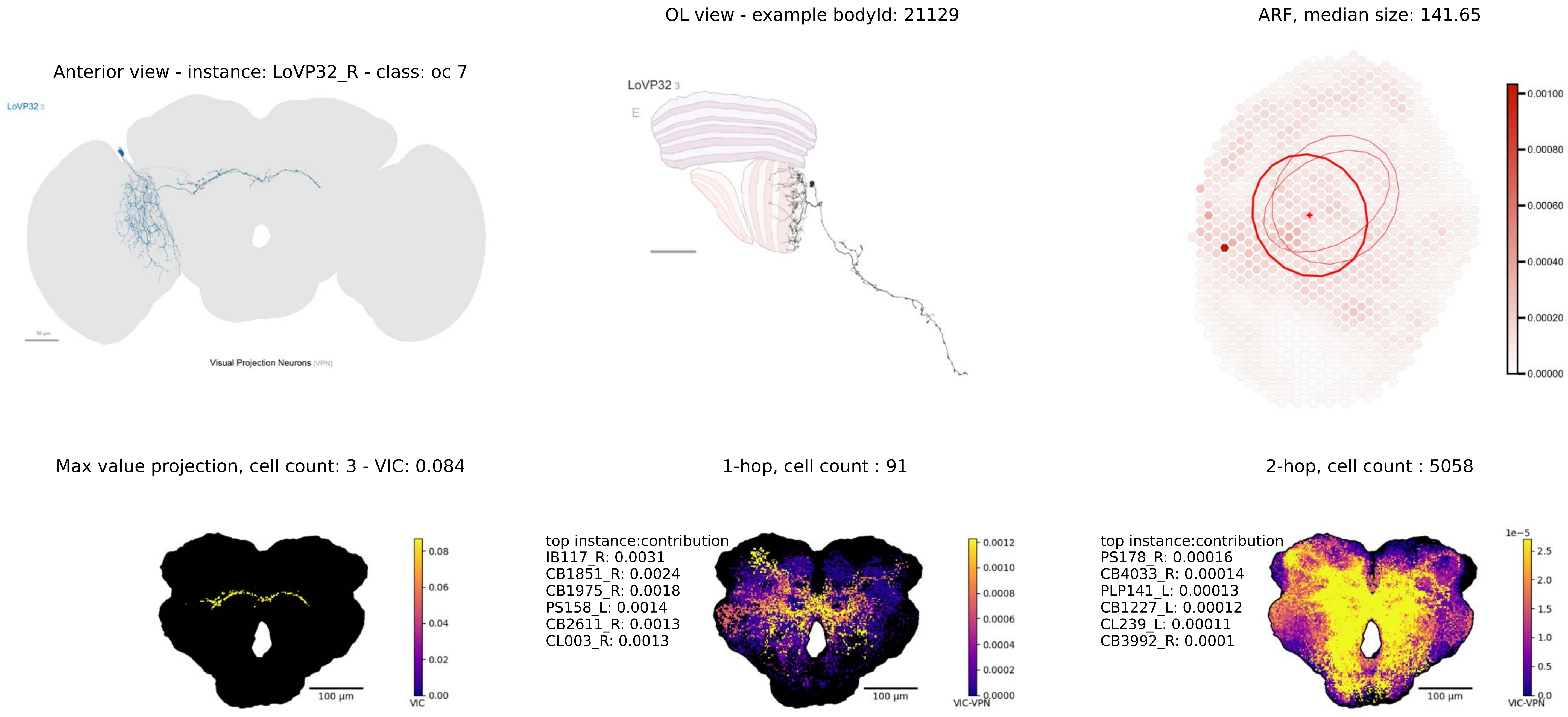

Anterior view - instance: LoVP32_R - class: oc 7
LoVP32 3
Visual Projection Neurons (VPN)
OL view - example bodyId: 21129
LoVP32 3
E
ARF, median size: 141.65
0.00100
0.00080
0.00060
0.00040
0.00020
0.00000
Max value projection, cell count: 3 - VIC: 0.084
100 µm
0.08
0.06
0.04
0.02
0.00
VIC
1-hop, cell count : 91
top instance:contribution
IB117_R: 0.0031
CB1851_R: 0.0024
CB1975_R: 0.0018
PS158_L: 0.0014
CB2611_R: 0.0013
CL003_R: 0.0013
100 µm
0.0012
0.0010
0.0008
0.0006
0.0004
0.0002
0.0000
VIC-VPN
2-hop, cell count : 5058
top instance:contribution
PS178_R: 0.00016
CB4033_R: 0.00014
PLP141_L: 0.00013
CB1227_L: 0.00012
CL239_L: 0.00011
CB3992_R: 0.0001
100 µm
1e−5
2.5
2.0
1.5
1.0
0.5
0.0
VIC-VPN

OL view - example bodyId: 36596

ARF, median size: 18.49

Anterior view - instance: LoVP33_R - class: oc 6

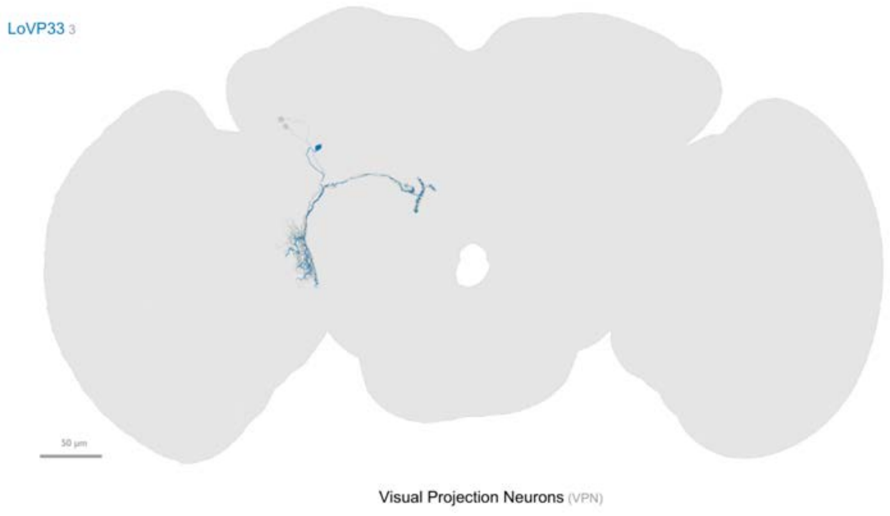


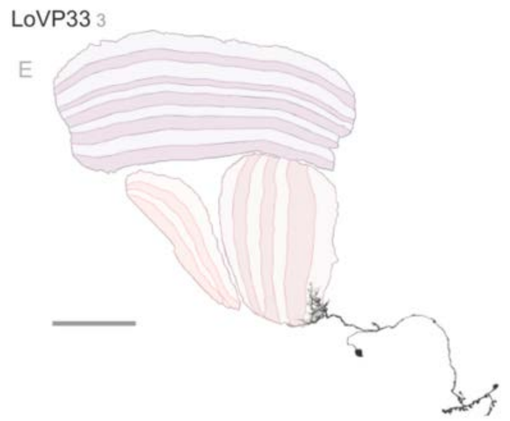


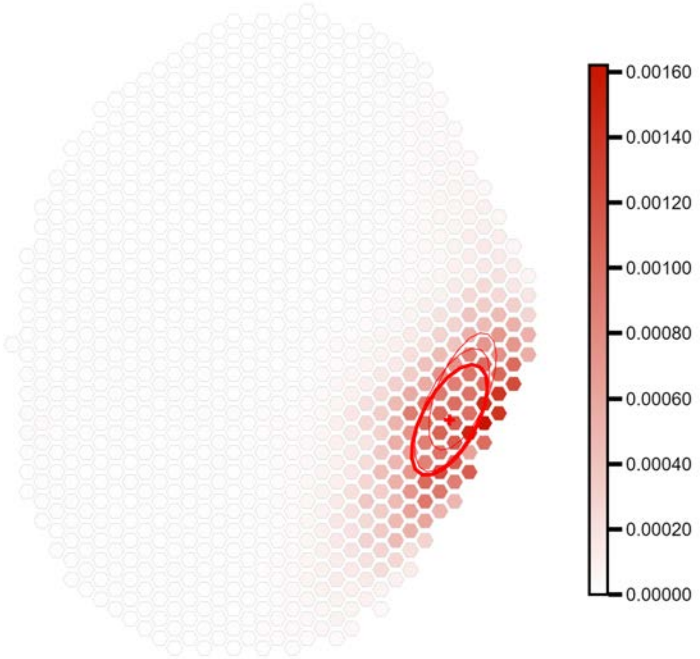


Max value projection, cell count: 3 - VIC: 0.084

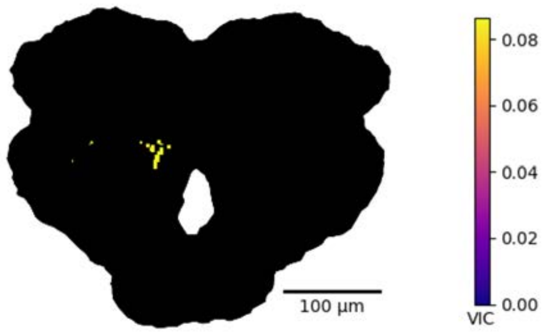


1-hop, cell count : 30

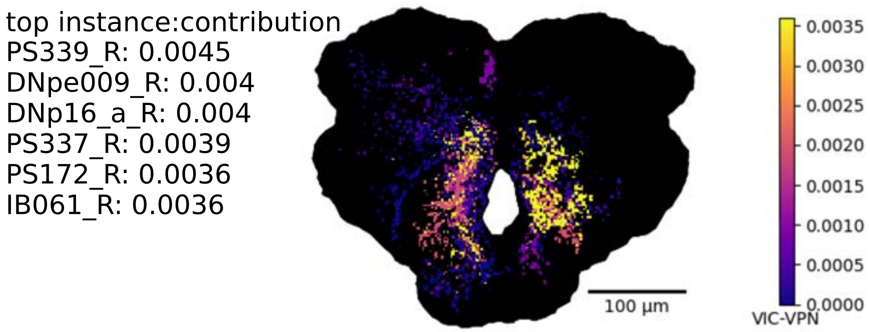


2-hop, cell count : 1138

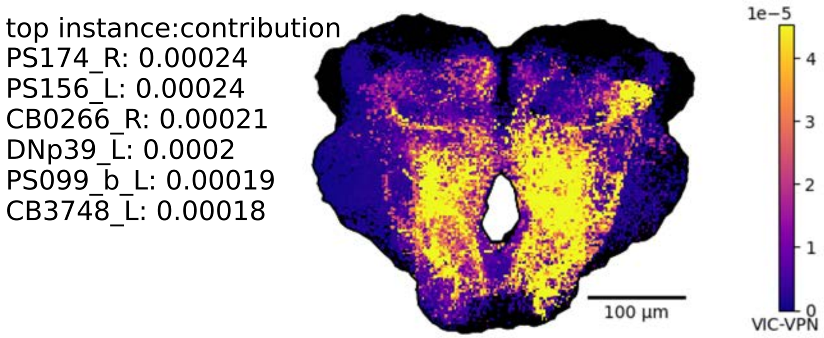

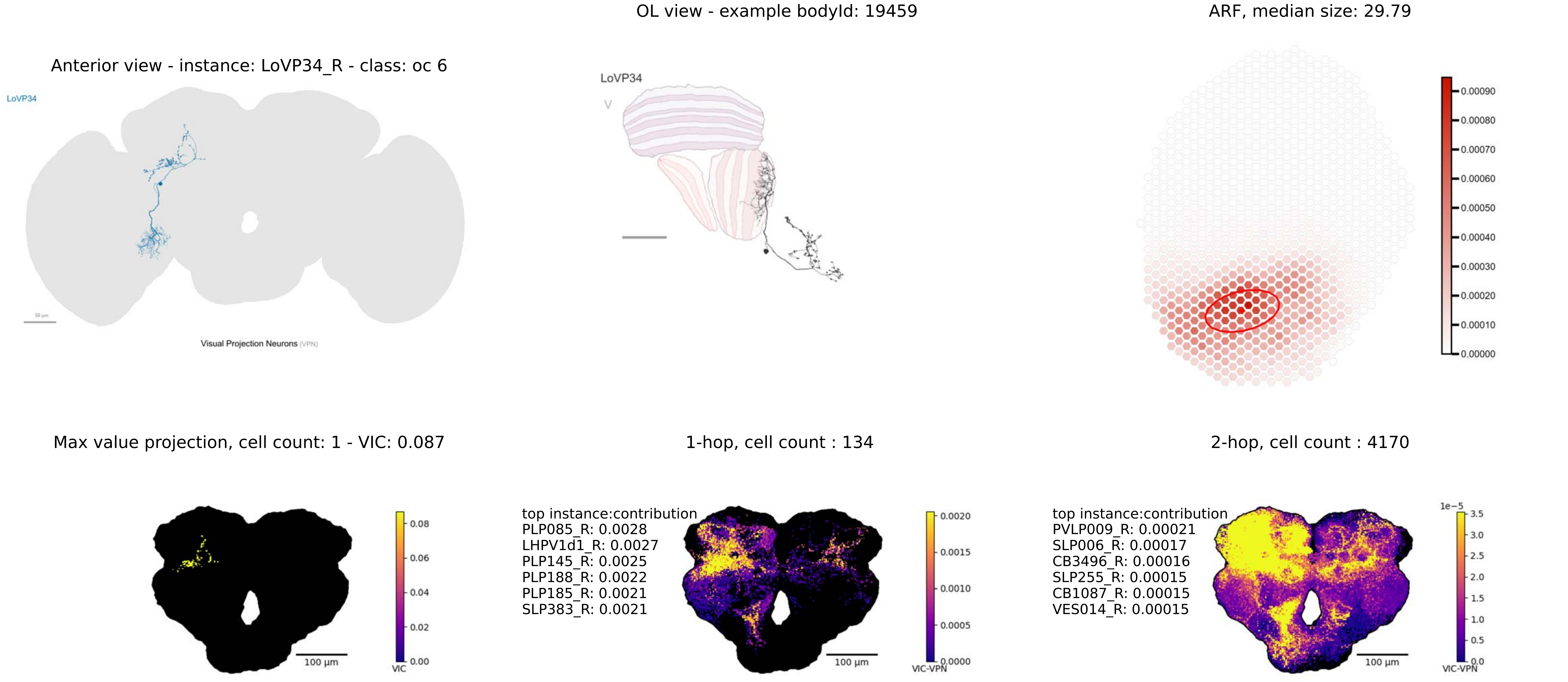
Anterior view - instance: LoVP34_R - class: oc 6
LoVP34
Visual Projection Neurons (VPN)
OL view - example bodyId: 19459
LoVP34
V
ARF, median size: 29.79
0.00090
0.00080
0.00070
0.00060
0.00050
0.00040
0.00030
0.00020
0.00010
0.00000
Max value projection, cell count: 1 - VIC: 0.087
0.08
0.06
0.04
0.02
0.00
VIC
100 µm
1-hop, cell count : 134
top instance:contribution
PLP085_R: 0.0028
LHPV1d1_R: 0.0027
PLP145_R: 0.0025
PLP188_R: 0.0022
PLP185_R: 0.0021
SLP383_R: 0.0021
100 µm
0.0020
0.0015
0.0010
0.0005
0.0000
VIC-VPN
2-hop, cell count : 4170
top instance:contribution
PVLP009_R: 0.00021
SLP006_R: 0.00017
CB3496_R: 0.00016
SLP255_R: 0.00015
CB1087_R: 0.00015
VES014_R: 0.00015
100 µm
1e−5
3.5
3.0
2.5
2.0
1.5
1.0
0.5
0.0
VIC-VPN

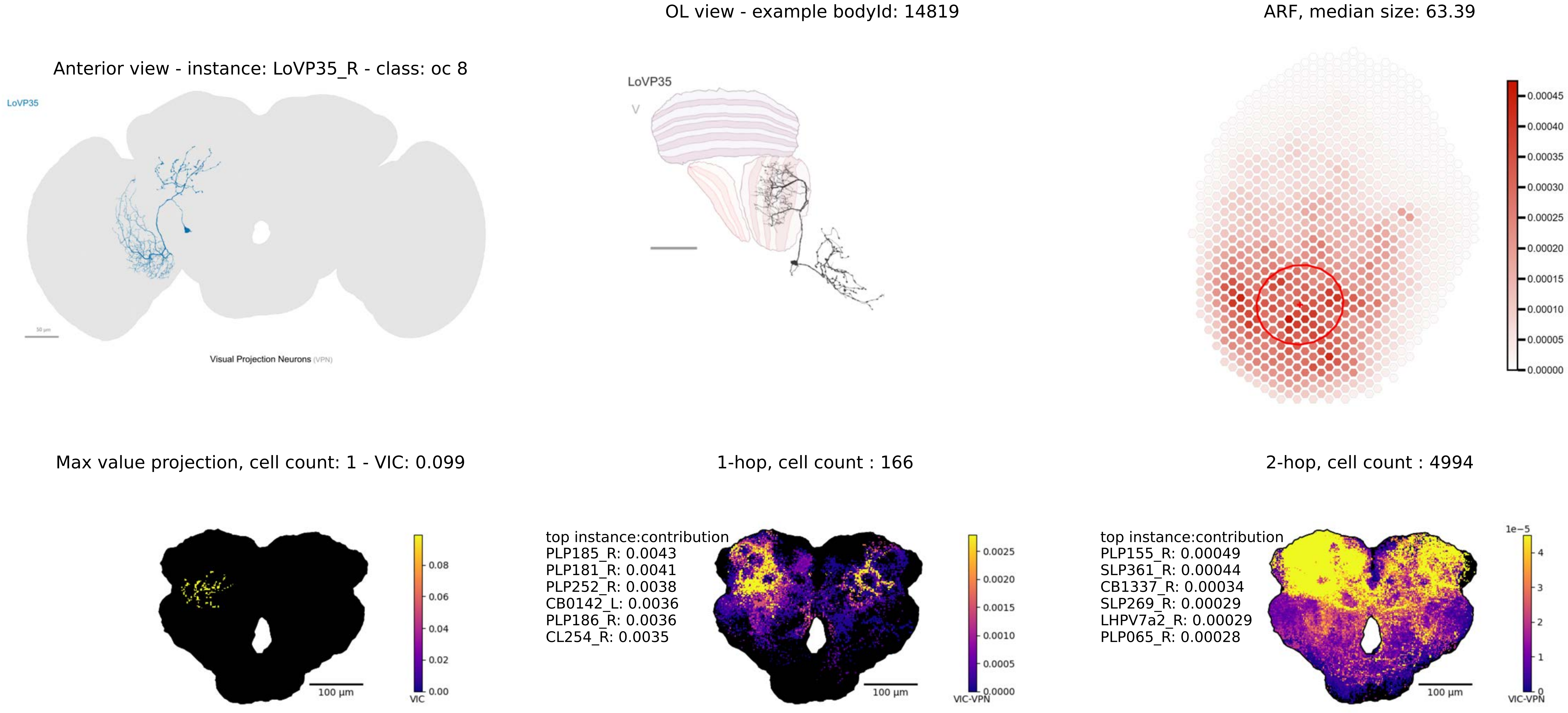
OL view - example bodyId: 14819
ARF, median size: 63.39
Anterior view - instance: LoVP35_R - class: oc 8
LoVP35
Visual Projection Neurons (VPN)
LoVP35
V
0.00045
0.00040
0.00035
0.00030
0.00025
0.00020
0.00015
0.00010
0.00005
0.00000
Max value projection, cell count: 1 - VIC: 0.099
1-hop, cell count : 166
2-hop, cell count : 4994
0.08
0.06
0.04
0.02
0.00
VIC
100 µm
top instance:contribution
PLP185_R: 0.0043
PLP181_R: 0.0041
PLP252_R: 0.0038
CB0142_L: 0.0036
PLP186_R: 0.0036
CL254_R: 0.0035
0.0025
0.0020
0.0015
0.0010
0.0005
0.0000
VIC-VPN
100 µm
top instance:contribution
PLP155_R: 0.00049
SLP361_R: 0.00044
CB1337_R: 0.00034
SLP269_R: 0.00029
LHPV7a2_R: 0.00029
PLP065_R: 0.00028
1e−5
4
3
2
1
0
VIC-VPN
100 µm

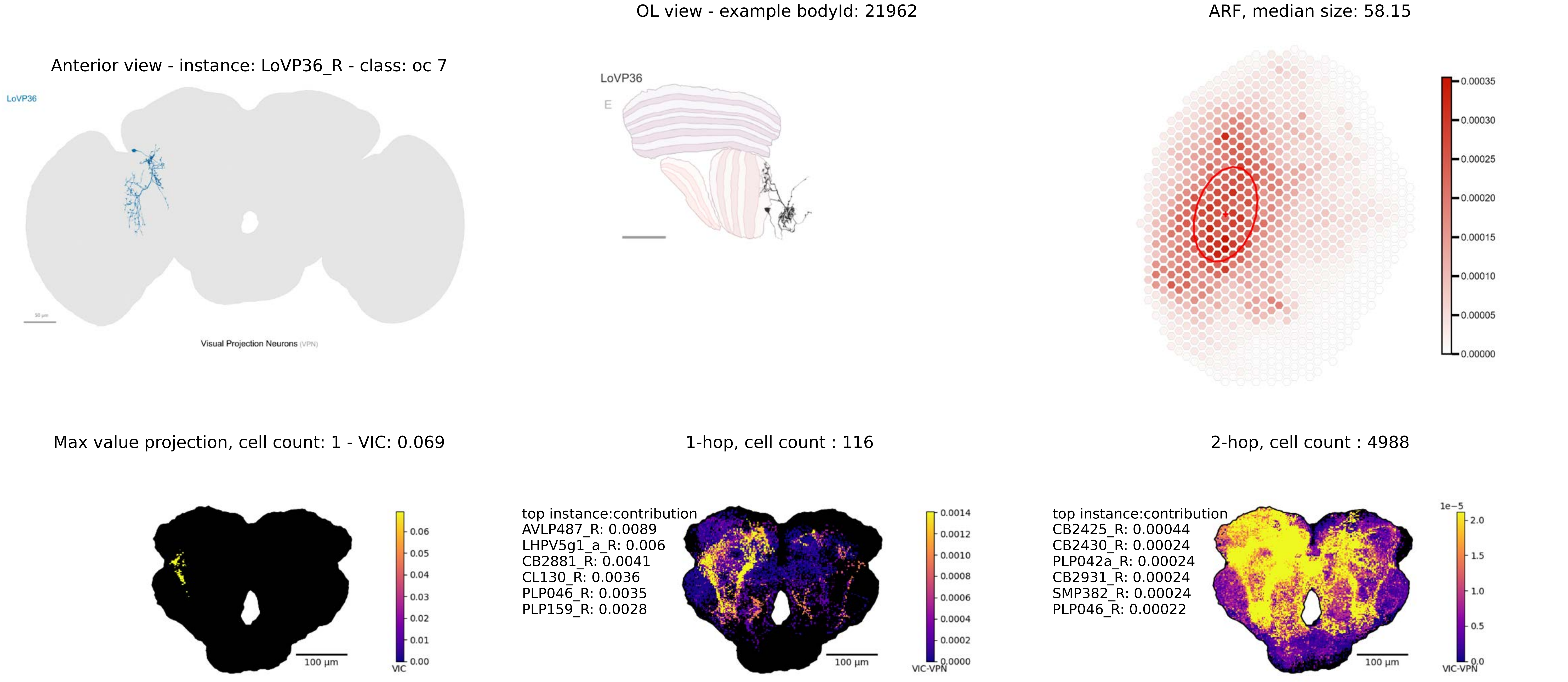
OL view - example bodyId: 21962
ARF, median size: 58.15
Anterior view - instance: LoVP36_R - class: oc 7
LoVP36
Visual Projection Neurons (VPN)
LoVP36
E
0.00035
0.00030
0.00025
0.00020
0.00015
0.00010
0.00005
0.00000
Max value projection, cell count: 1 - VIC: 0.069
1-hop, cell count : 116
2-hop, cell count : 4988
100 µm
VIC
0.06
0.05
0.04
0.03
0.02
0.01
0.00
top instance:contribution
AVLP487_R: 0.0089
LHPV5g1_a_R: 0.006
CB2881_R: 0.0041
CL130_R: 0.0036
PLP046_R: 0.0035
PLP159_R: 0.0028
100 µm
VIC-VPN
0.0014
0.0012
0.0010
0.0008
0.0006
0.0004
0.0002
0.0000
top instance:contribution
CB2425_R: 0.00044
CB2430_R: 0.00024
PLP042a_R: 0.00024
CB2931_R: 0.00024
SMP382_R: 0.00024
PLP046_R: 0.00022
100 µm
1e−5
2.0
1.5
1.0
0.5
0.0
VIC-VPN

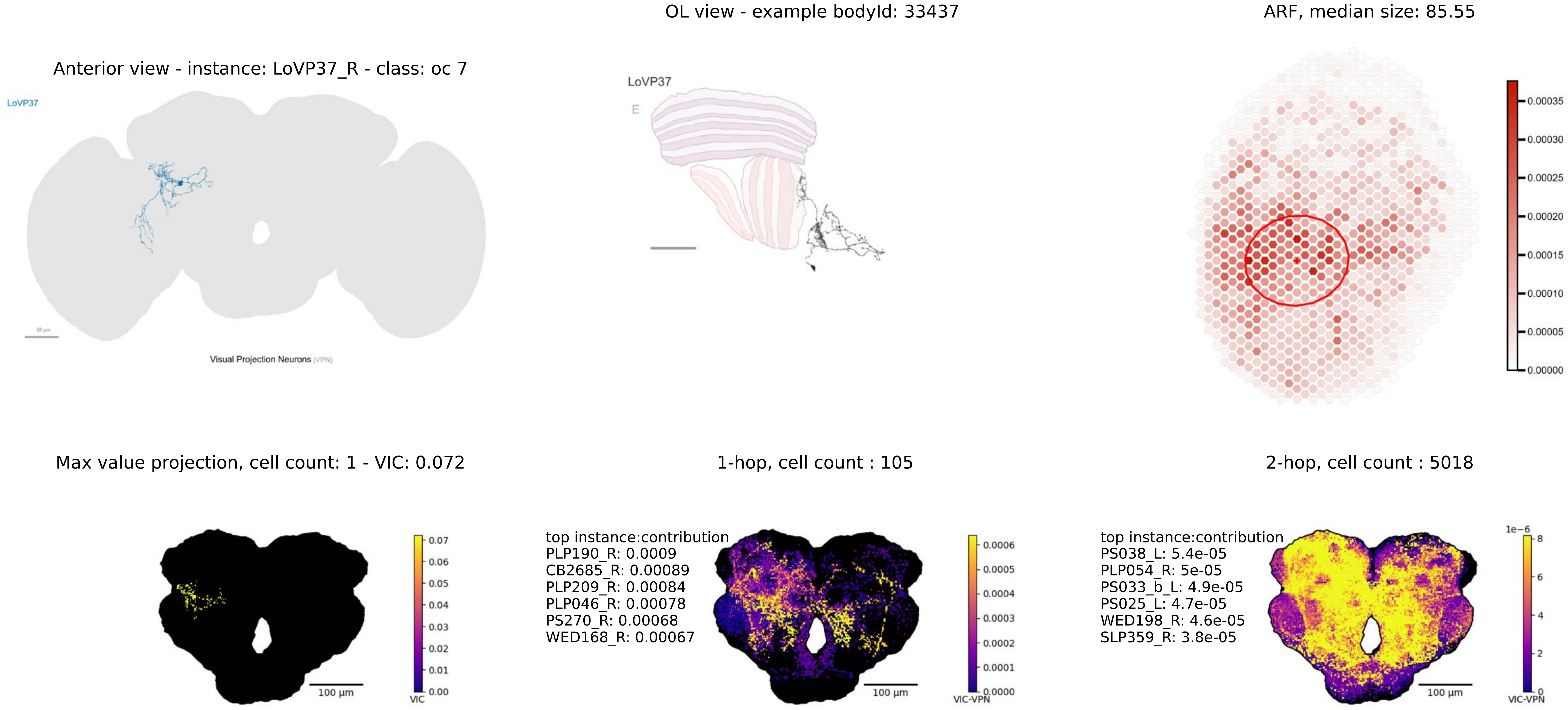
OL view - example bodyId: 33437
ARF, median size: 85.55
Anterior view - instance: LoVP37_R - class: oc 7
LoVP37
Visual Projection Neurons (VPN)
LoVP37
E
0.00035
0.00030
0.00025
0.00020
0.00015
0.00010
0.00005
0.00000
Max value projection, cell count: 1 - VIC: 0.072
1-hop, cell count : 105
2-hop, cell count : 5018
100 µm
VIC
0.07
0.06
0.05
0.04
0.03
0.02
0.01
0.00
top instance:contribution
PLP190_R: 0.0009
CB2685_R: 0.00089
PLP209_R: 0.00084
PLP046_R: 0.00078
PS270_R: 0.00068
WED168_R: 0.00067
100 µm
VIC-VPN
0.0006
0.0005
0.0004
0.0003
0.0002
0.0001
0.0000
top instance:contribution
PS038_L: 5.4e-05
PLP054_R: 5e-05
PS033_b_L: 4.9e-05
PS025_L: 4.7e-05
WED198_R: 4.6e-05
SLP359_R: 3.8e-05
100 µm
1e−6
8
6
4
2
0
VIC-VPN

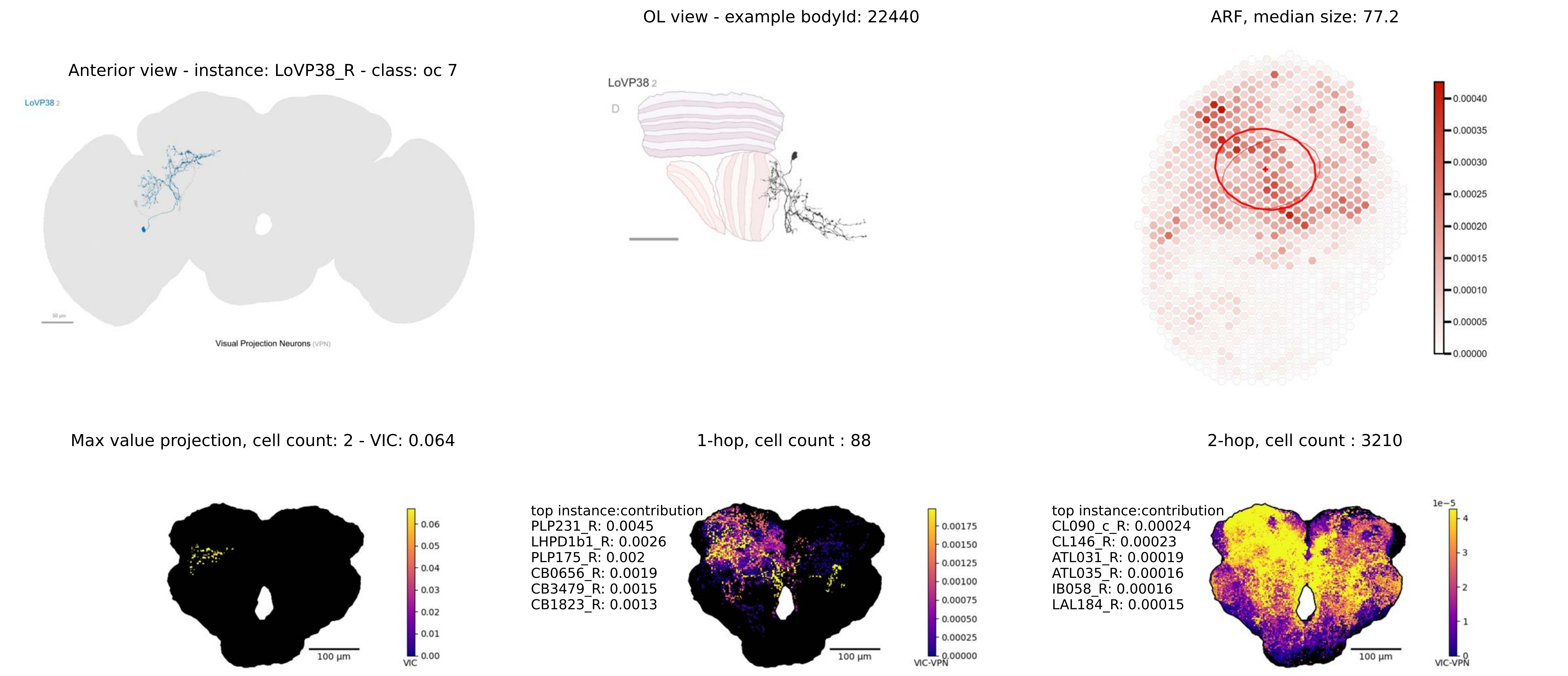

OL view - example bodyId: 22440
ARF, median size: 77.2
Anterior view - instance: LoVP38_R - class: oc 7
LoVP38 2
LoVP38 2
D
Visual Projection Neurons (VPN)
0.00040
0.00035
0.00030
0.00025
0.00020
0.00015
0.00010
0.00005
0.00000
Max value projection, cell count: 2 - VIC: 0.064
1-hop, cell count : 88
2-hop, cell count : 3210
top instance:contribution
PLP231_R: 0.0045
LHPD1b1_R: 0.0026
PLP175_R: 0.002
CB0656_R: 0.0019
CB3479_R: 0.0015
CB1823_R: 0.0013
top instance:contribution
CL090_c_R: 0.00024
CL146_R: 0.00023
ATL031_R: 0.00019
ATL035_R: 0.00016
IB058_R: 0.00016
LAL184_R: 0.00015
0.06
0.05
0.04
0.03
0.02
0.01
0.00
VIC
100 µm
0.00175
0.00150
0.00125
0.00100
0.00075
0.00050
0.00025
0.00000
VIC-VPN
100 µm
1e−5
4
3
2
1
0
VIC-VPN
100 µm

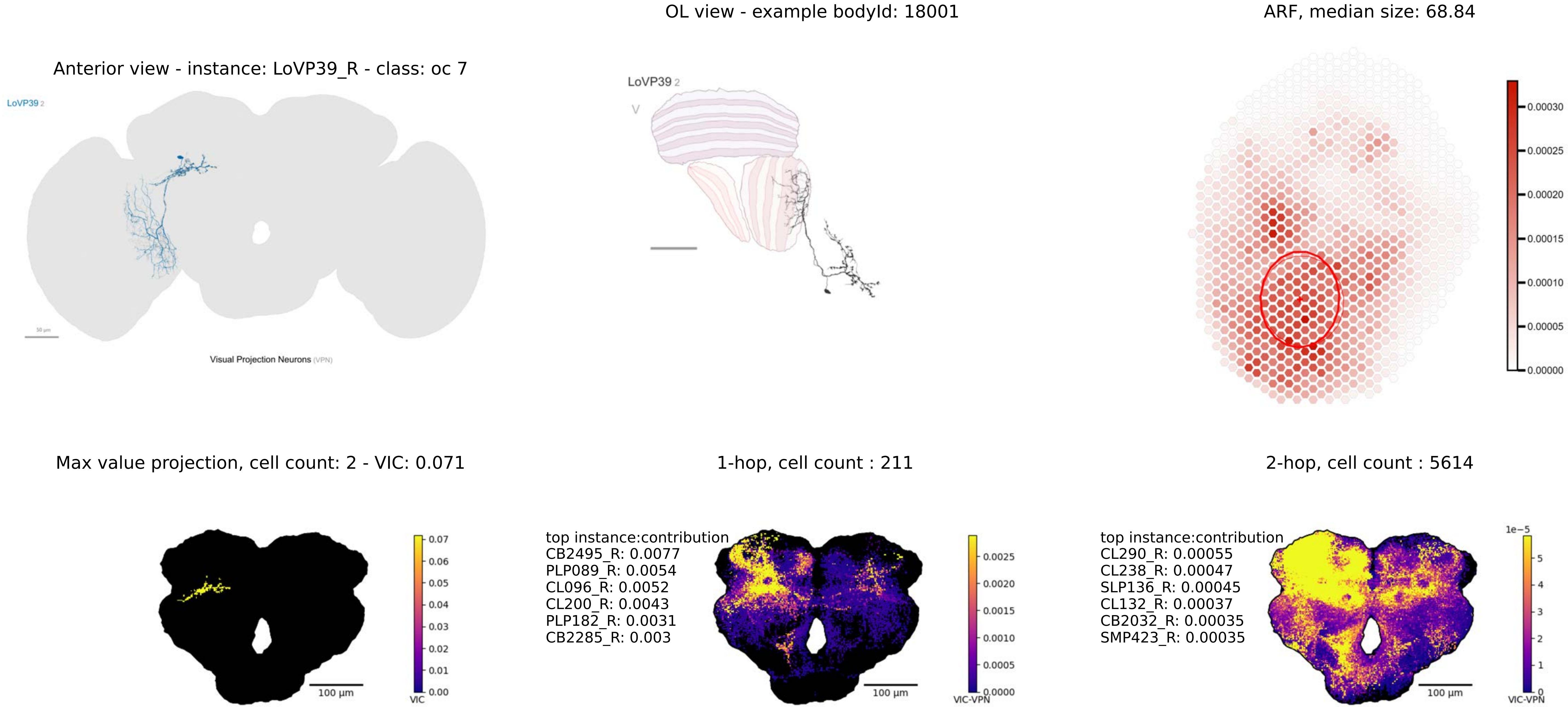
Anterior view - instance: LoVP39_R - class: oc 7
LoVP39 2
Visual Projection Neurons (VPN)
OL view - example bodyId: 18001
LoVP39 2
ARF, median size: 68.84
0.00030
0.00025
0.00020
0.00015
0.00010
0.00005
0.00000
Max value projection, cell count: 2 - VIC: 0.071
100 µm
VIC
0.07
0.06
0.05
0.04
0.03
0.02
0.01
0.00
1-hop, cell count : 211
top instance:contribution
CB2495_R: 0.0077
PLP089_R: 0.0054
CL096_R: 0.0052
CL200_R: 0.0043
PLP182_R: 0.0031
CB2285_R: 0.003
100 µm
VIC-VPN
0.0025
0.0020
0.0015
0.0010
0.0005
0.0000
2-hop, cell count : 5614
top instance:contribution
CL290_R: 0.00055
CL238_R: 0.00047
SLP136_R: 0.00045
CL132_R: 0.00037
CB2032_R: 0.00035
SMP423_R: 0.00035
100 µm
1e−5
VIC-VPN
5
4
3
2
1
0

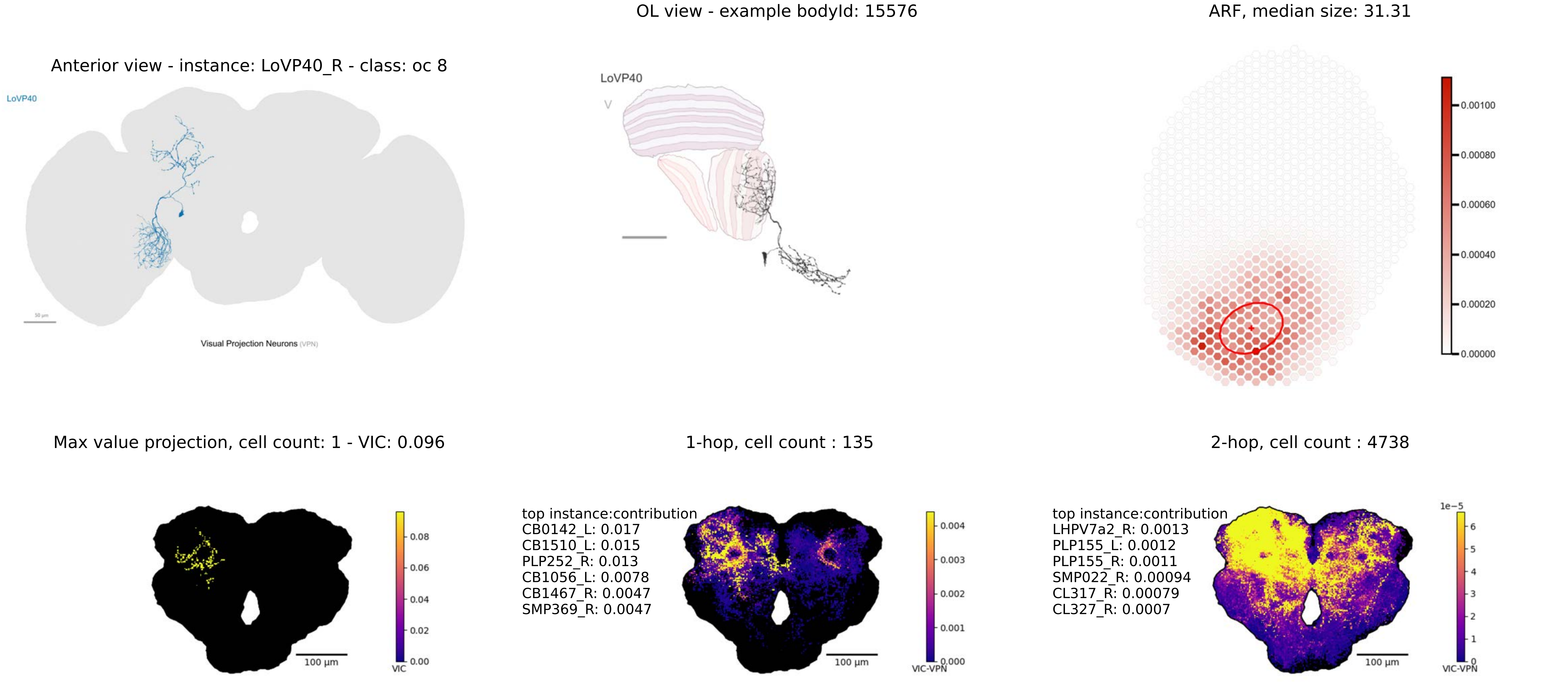

OL view - example bodyId: 15576
ARF, median size: 31.31
Anterior view - instance: LoVP40_R - class: oc 8
LoVP40
Visual Projection Neurons (VPN)
LoVP40
0.00100
0.00080
0.00060
0.00040
0.00020
0.00000
Max value projection, cell count: 1 - VIC: 0.096
1-hop, cell count : 135
2-hop, cell count : 4738
top instance:contribution
CB0142_L: 0.017
CB1510_L: 0.015
PLP252_R: 0.013
CB1056_L: 0.0078
CB1467_R: 0.0047
SMP369_R: 0.0047
top instance:contribution
LHPV7a2_R: 0.0013
PLP155_L: 0.0012
PLP155_R: 0.0011
SMP022_R: 0.00094
CL317_R: 0.00079
CL327_R: 0.0007
0.08
0.06
0.04
0.02
0.00
VIC
100 µm
0.004
0.003
0.002
0.001
0.000
VIC-VPN
100 µm
1e−5
6
5
4
3
2
1
0
VIC-VPN
100 µm

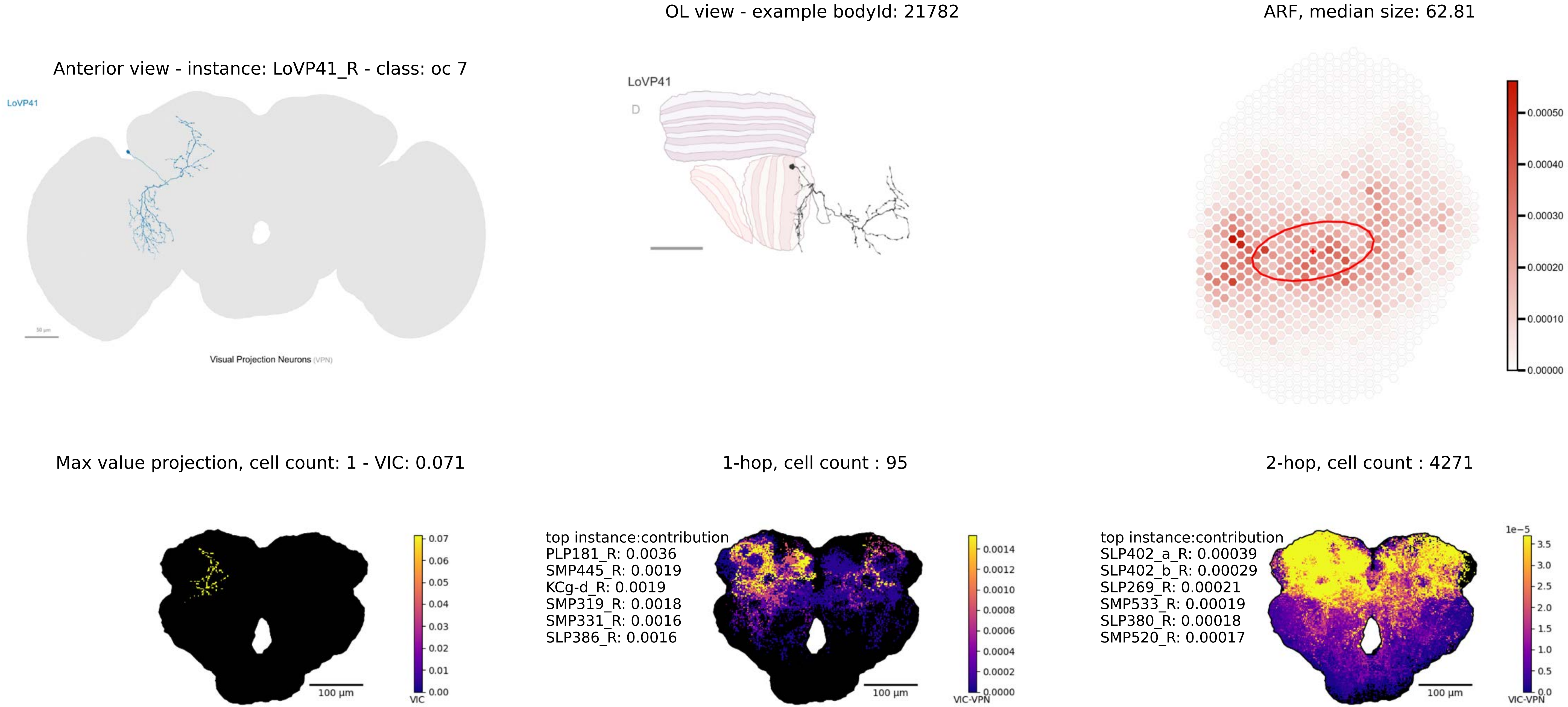
OL view - example bodyId: 21782
ARF, median size: 62.81
Anterior view - instance: LoVP41_R - class: oc 7
LoVP41
Visual Projection Neurons (VPN)
LoVP41
D
0.00050
0.00040
0.00030
0.00020
0.00010
0.00000
Max value projection, cell count: 1 - VIC: 0.071
1-hop, cell count : 95
2-hop, cell count : 4271
100 µm
VIC
0.07
0.06
0.05
0.04
0.03
0.02
0.01
0.00
top instance:contribution
PLP181_R: 0.0036
SMP445_R: 0.0019
KCg-d_R: 0.0019
SMP319_R: 0.0018
SMP331_R: 0.0016
SLP386_R: 0.0016
100 µm
VIC-VPN
0.0014
0.0012
0.0010
0.0008
0.0006
0.0004
0.0002
0.0000
top instance:contribution
SLP402_a_R: 0.00039
SLP402_b_R: 0.00029
SLP269_R: 0.00021
SMP533_R: 0.00019
SLP380_R: 0.00018
SMP520_R: 0.00017
100 µm
1e−5
VIC-VPN
3.5
3.0
2.5
2.0
1.5
1.0
0.5
0.0

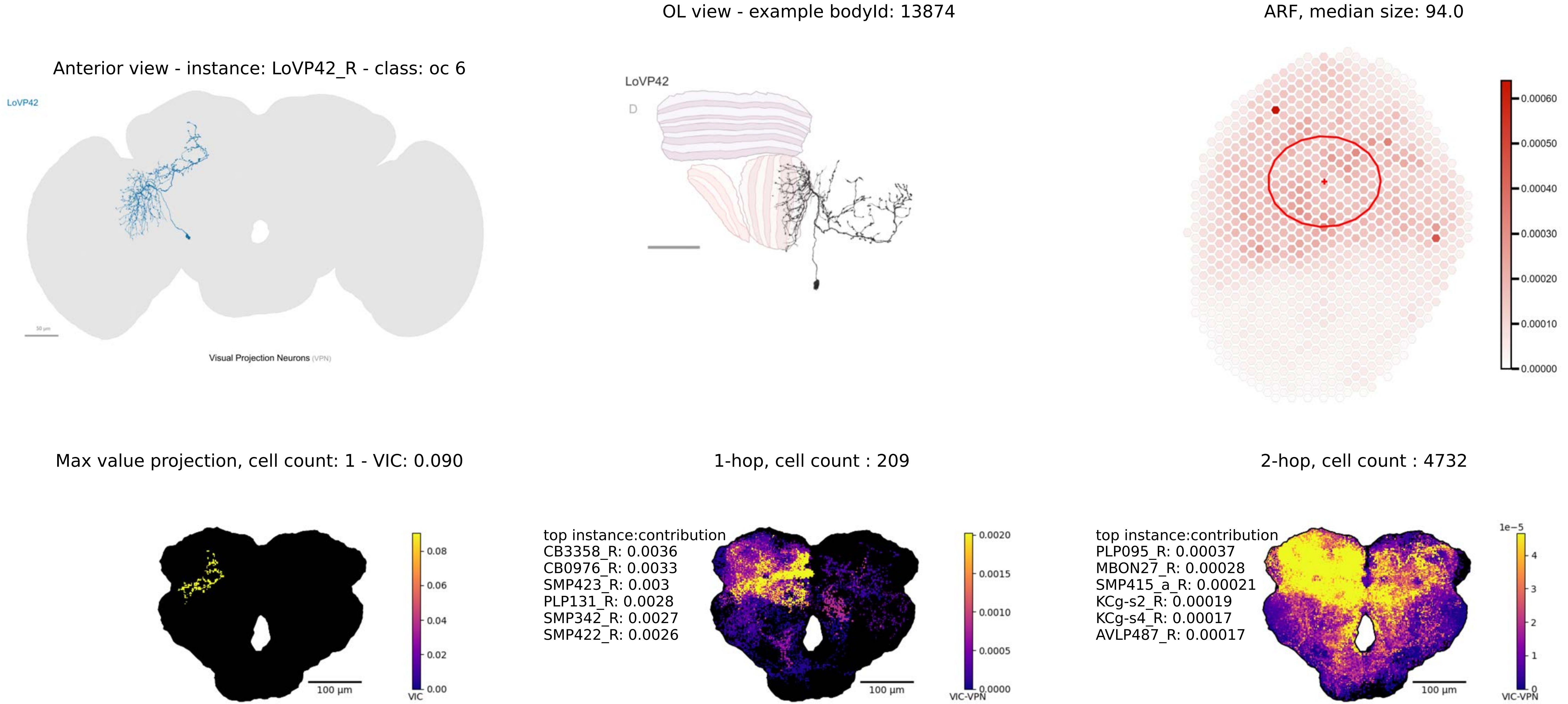

Anterior view - instance: LoVP42_R - class: oc 6
LoVP42
Visual Projection Neurons (VPN)
OL view - example bodyId: 13874
LoVP42
D
ARF, median size: 94.0
0.00060
0.00050
0.00040
0.00030
0.00020
0.00010
0.00000
Max value projection, cell count: 1 - VIC: 0.090
100 µm
VIC
0.08
0.06
0.04
0.02
0.00
1-hop, cell count : 209
top instance:contribution
CB3358_R: 0.0036
CB0976_R: 0.0033
SMP423_R: 0.003
PLP131_R: 0.0028
SMP342_R: 0.0027
SMP422_R: 0.0026
100 µm
VIC-VPN
0.0020
0.0015
0.0010
0.0005
0.0000
2-hop, cell count : 4732
top instance:contribution
PLP095_R: 0.00037
MBON27_R: 0.00028
SMP415_a_R: 0.00021
KCg-s2_R: 0.00019
KCg-s4_R: 0.00017
AVLP487_R: 0.00017
100 µm
1e−5
4
3
2
1
0
VIC-VPN

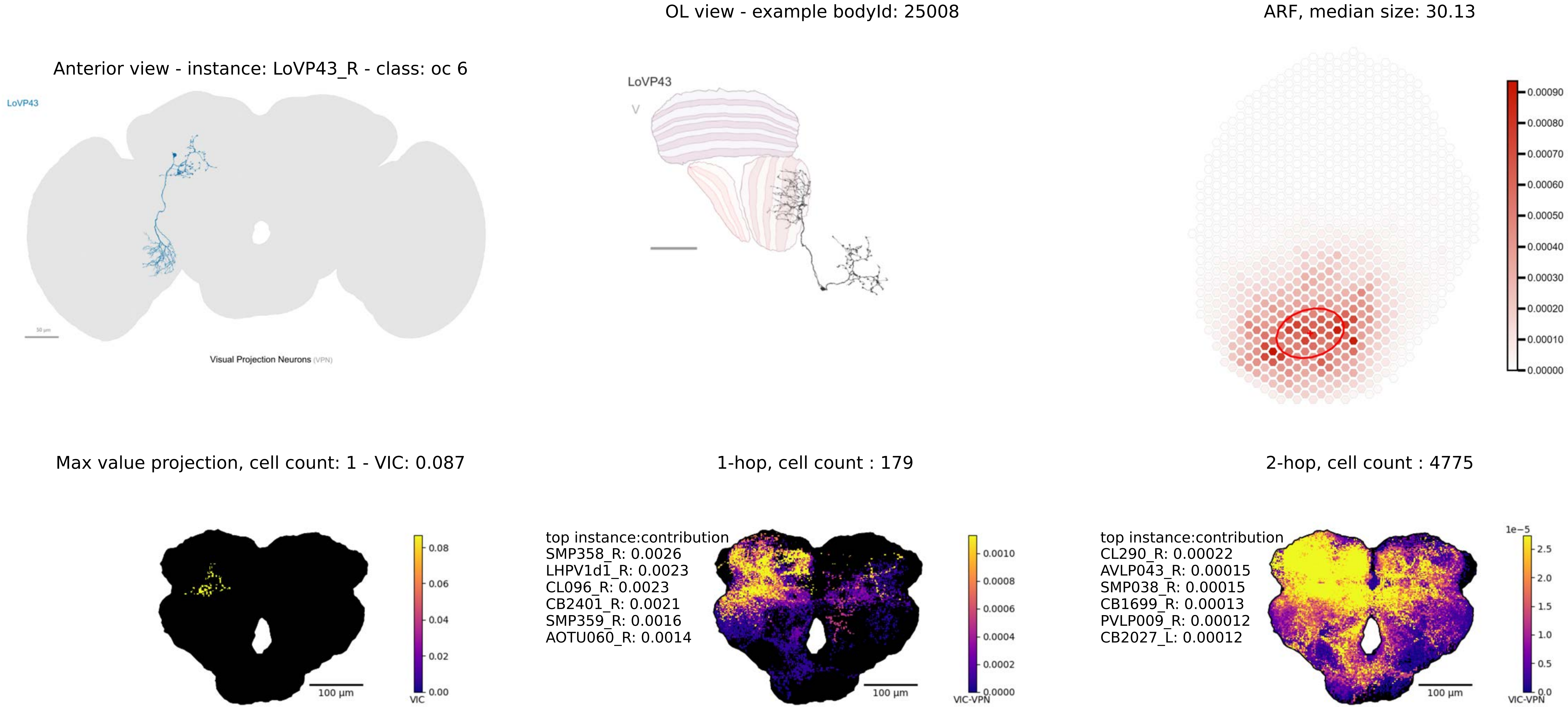
Anterior view - instance: LoVP43_R - class: oc 6
LoVP43
Visual Projection Neurons (VPN)
OL view - example bodyId: 25008
LoVP43
V
ARF, median size: 30.13
0.00090
0.00080
0.00070
0.00060
0.00050
0.00040
0.00030
0.00020
0.00010
0.00000
Max value projection, cell count: 1 - VIC: 0.087
100 µm
0.08
0.06
0.04
0.02
0.00
VIC
1-hop, cell count : 179
top instance:contribution
SMP358_R: 0.0026
LHPV1d1_R: 0.0023
CL096_R: 0.0023
CB2401_R: 0.0021
SMP359_R: 0.0016
AOTU060_R: 0.0014
100 µm
0.0010
0.0008
0.0006
0.0004
0.0002
0.0000
VIC-VPN
2-hop, cell count : 4775
top instance:contribution
CL290_R: 0.00022
AVLP043_R: 0.00015
SMP038_R: 0.00015
CB1699_R: 0.00013
PVLP009_R: 0.00012
CB2027_L: 0.00012
100 µm
1e−5
2.5
2.0
1.5
1.0
0.5
0.0
VIC-VPN

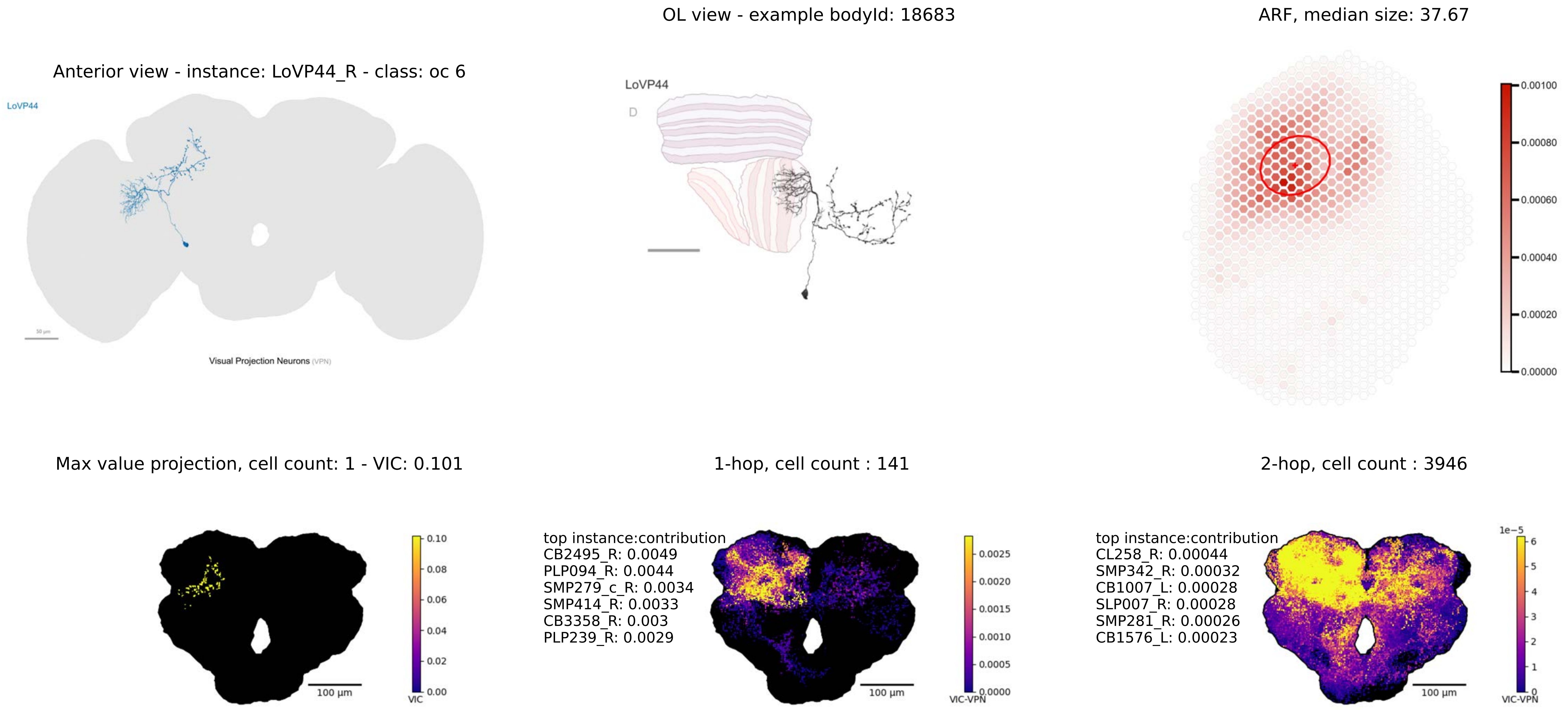
OL view - example bodyId: 18683
ARF, median size: 37.67
Anterior view - instance: LoVP44_R - class: oc 6
LoVP44
Visual Projection Neurons (VPN)
LoVP44
D
0.00100
0.00080
0.00060
0.00040
0.00020
0.00000
Max value projection, cell count: 1 - VIC: 0.101
1-hop, cell count : 141
2-hop, cell count : 3946
top instance:contribution
CB2495_R: 0.0049
PLP094_R: 0.0044
SMP279_c_R: 0.0034
SMP414_R: 0.0033
CB3358_R: 0.003
PLP239_R: 0.0029
top instance:contribution
CL258_R: 0.00044
SMP342_R: 0.00032
CB1007_L: 0.00028
SLP007_R: 0.00028
SMP281_R: 0.00026
CB1576_L: 0.00023
0.10
0.08
0.06
0.04
0.02
0.00
VIC
100 µm
0.0025
0.0020
0.0015
0.0010
0.0005
0.0000
VIC-VPN
100 µm
1e−5
6
5
4
3
2
1
0
VIC-VPN
100 µm

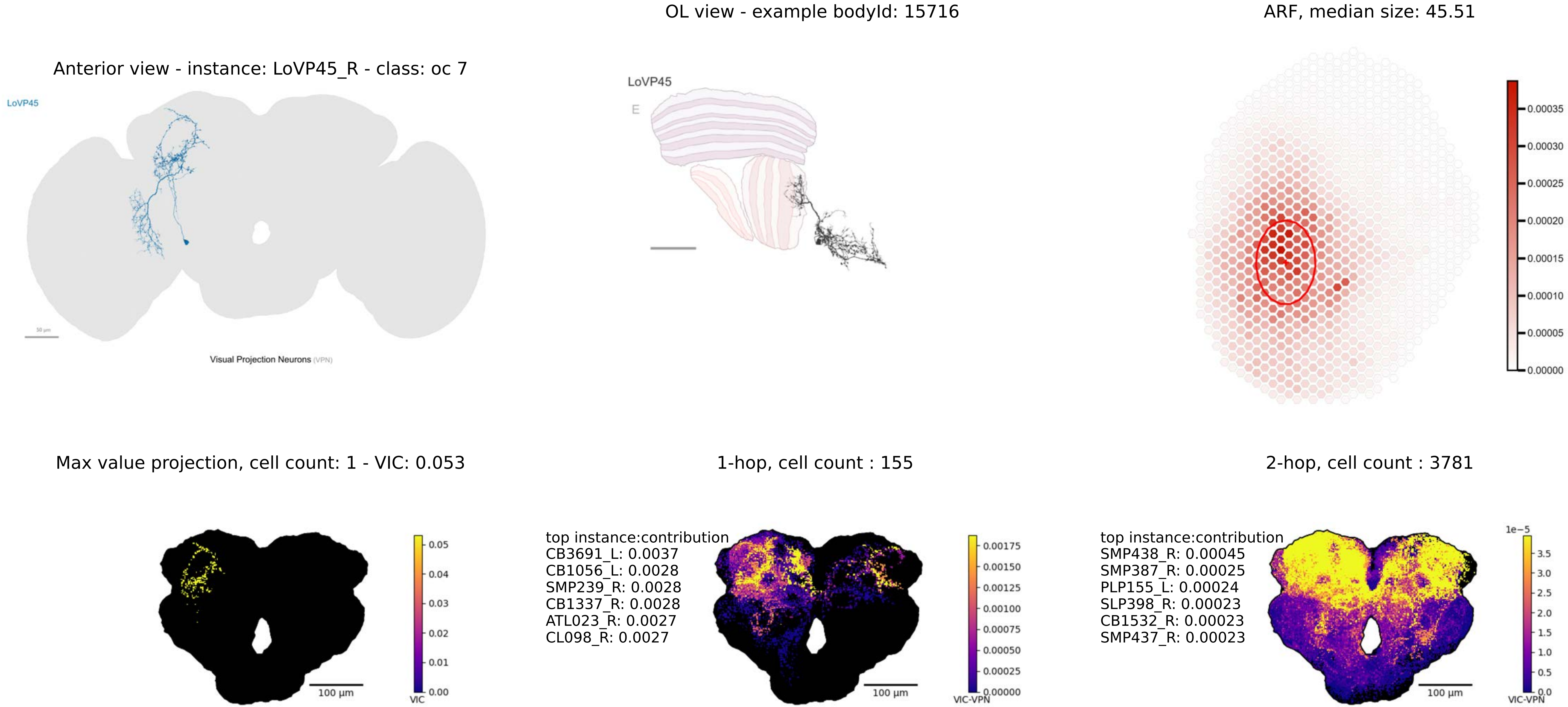
OL view - example bodyId: 15716
ARF, median size: 45.51
Anterior view - instance: LoVP45_R - class: oc 7
LoVP45
Visual Projection Neurons (VPN)
LoVP45
E
0.00035
0.00030
0.00025
0.00020
0.00015
0.00010
0.00005
0.00000
Max value projection, cell count: 1 - VIC: 0.053
1-hop, cell count : 155
2-hop, cell count : 3781
top instance:contribution
CB3691_L: 0.0037
CB1056_L: 0.0028
SMP239_R: 0.0028
CB1337_R: 0.0028
ATL023_R: 0.0027
CL098_R: 0.0027
top instance:contribution
SMP438_R: 0.00045
SMP387_R: 0.00025
PLP155_L: 0.00024
SLP398_R: 0.00023
CB1532_R: 0.00023
SMP437_R: 0.00023
100 µm
VIC
VIC-VPN
1e−5

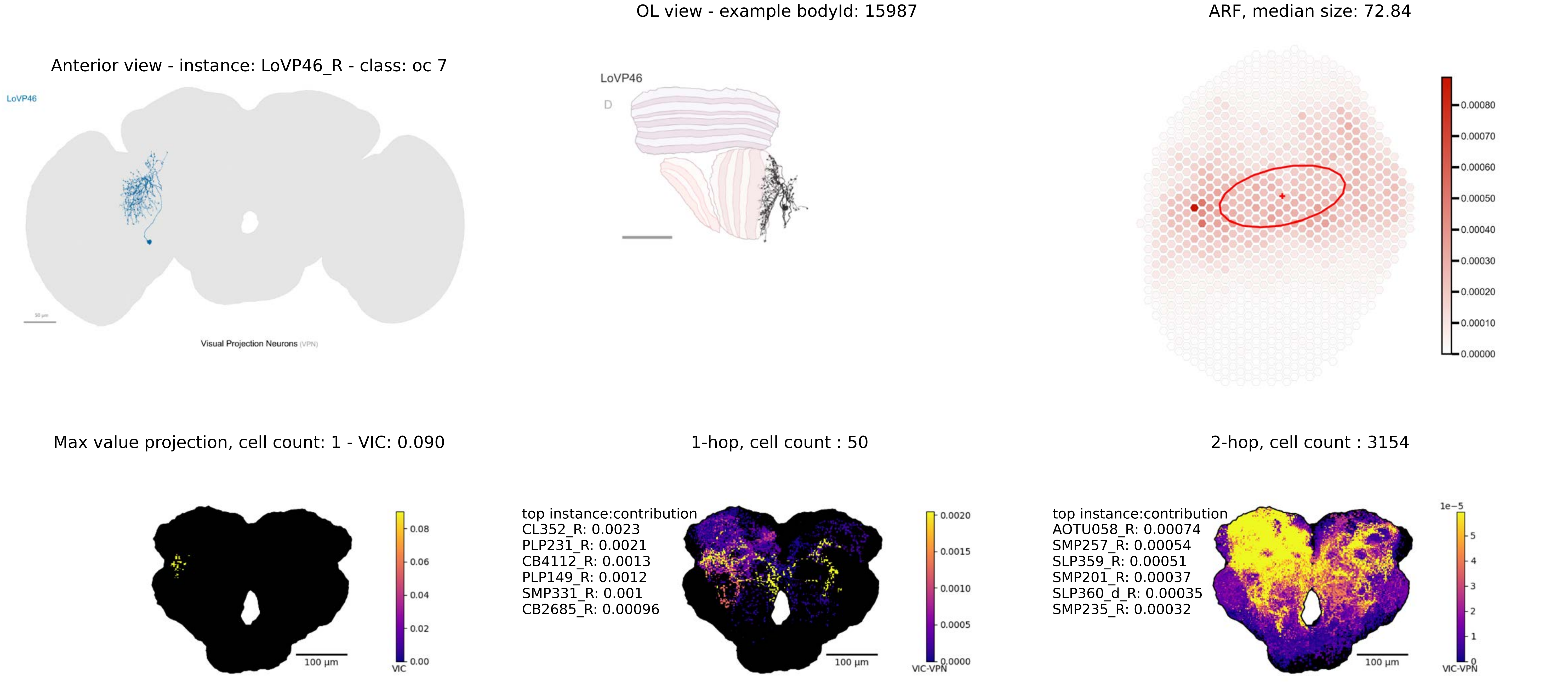
Anterior view - instance: LoVP46_R - class: oc 7
LoVP46
50 μm
Visual Projection Neurons (VPN)
OL view - example bodyId: 15987
LoVP46
D
ARF, median size: 72.84
0.00080
0.00070
0.00060
0.00050
0.00040
0.00030
0.00020
0.00010
0.00000
Max value projection, cell count: 1 - VIC: 0.090
100 μm
0.08
0.06
0.04
0.02
0.00
VIC
1-hop, cell count : 50
top instance:contribution
CL352_R: 0.0023
PLP231_R: 0.0021
CB4112_R: 0.0013
PLP149_R: 0.0012
SMP331_R: 0.001
CB2685_R: 0.00096
100 μm
0.0020
0.0015
0.0010
0.0005
0.0000
VIC-VPN
2-hop, cell count : 3154
top instance:contribution
AOTU058_R: 0.00074
SMP257_R: 0.00054
SLP359_R: 0.00051
SMP201_R: 0.00037
SLP360_d_R: 0.00035
SMP235_R: 0.00032
100 μm
1e−5
5
4
3
2
1
0
VIC-VPN

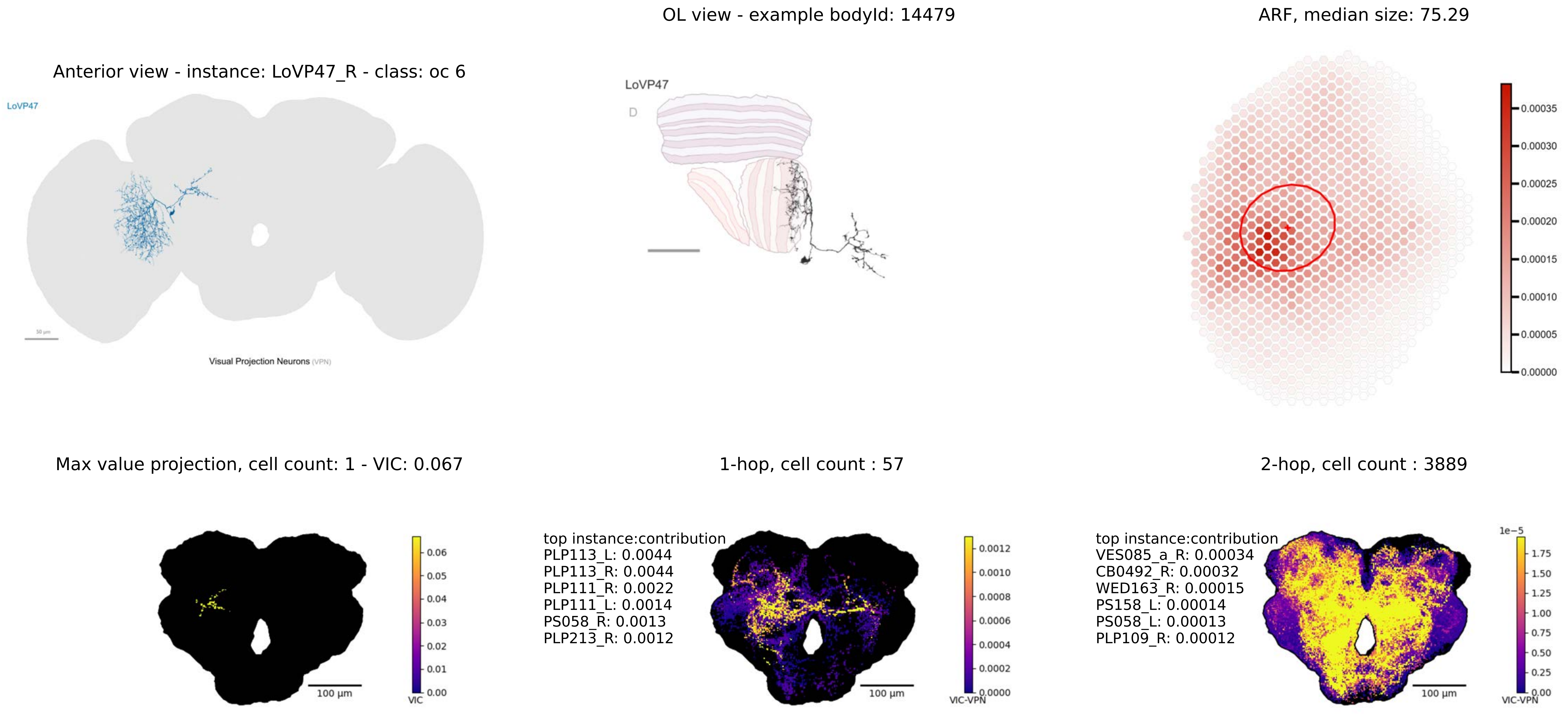
Anterior view - instance: LoVP47_R - class: oc 6
LoVP47
50 µm
Visual Projection Neurons (VPN)
OL view - example bodyId: 14479
LoVP47
D
ARF, median size: 75.29
0.00035
0.00030
0.00025
0.00020
0.00015
0.00010
0.00005
0.00000
Max value projection, cell count: 1 - VIC: 0.067
0.06
0.05
0.04
0.03
0.02
0.01
0.00
VIC
100 µm
1-hop, cell count : 57
top instance:contribution
PLP113_L: 0.0044
PLP113_R: 0.0044
PLP111_R: 0.0022
PLP111_L: 0.0014
PS058_R: 0.0013
PLP213_R: 0.0012
0.0012
0.0010
0.0008
0.0006
0.0004
0.0002
0.0000
VIC-VPN
100 µm
2-hop, cell count : 3889
top instance:contribution
VES085_a_R: 0.00034
CB0492_R: 0.00032
WED163_R: 0.00015
PS158_L: 0.00014
PS058_L: 0.00013
PLP109_R: 0.00012
1e−5
1.75
1.50
1.25
1.00
0.75
0.50
0.25
0.00
VIC-VPN
100 µm

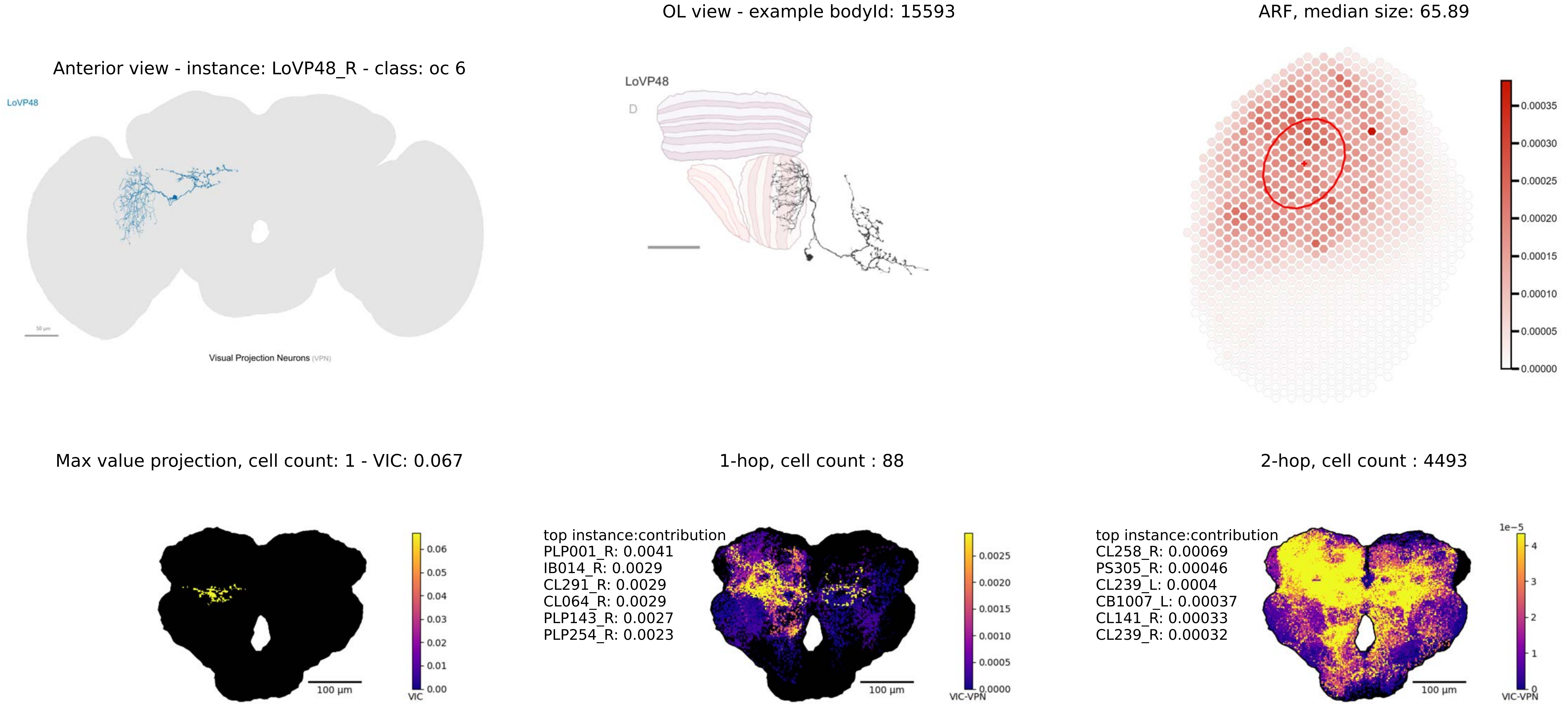
OL view - example bodyId: 15593
ARF, median size: 65.89
Anterior view - instance: LoVP48_R - class: oc 6
LoVP48
Visual Projection Neurons (VPN)
LoVP48
D
0.00035
0.00030
0.00025
0.00020
0.00015
0.00010
0.00005
0.00000
Max value projection, cell count: 1 - VIC: 0.067
1-hop, cell count : 88
2-hop, cell count : 4493
0.06
0.05
0.04
0.03
0.02
0.01
0.00
VIC
100 µm
top instance:contribution
PLP001_R: 0.0041
IB014_R: 0.0029
CL291_R: 0.0029
CL064_R: 0.0029
PLP143_R: 0.0027
PLP254_R: 0.0023
0.0025
0.0020
0.0015
0.0010
0.0005
0.0000
VIC-VPN
100 µm
top instance:contribution
CL258_R: 0.00069
PS305_R: 0.00046
CL239_L: 0.0004
CB1007_L: 0.00037
CL141_R: 0.00033
CL239_R: 0.00032
1e−5
4
3
2
1
0
VIC-VPN
100 µm

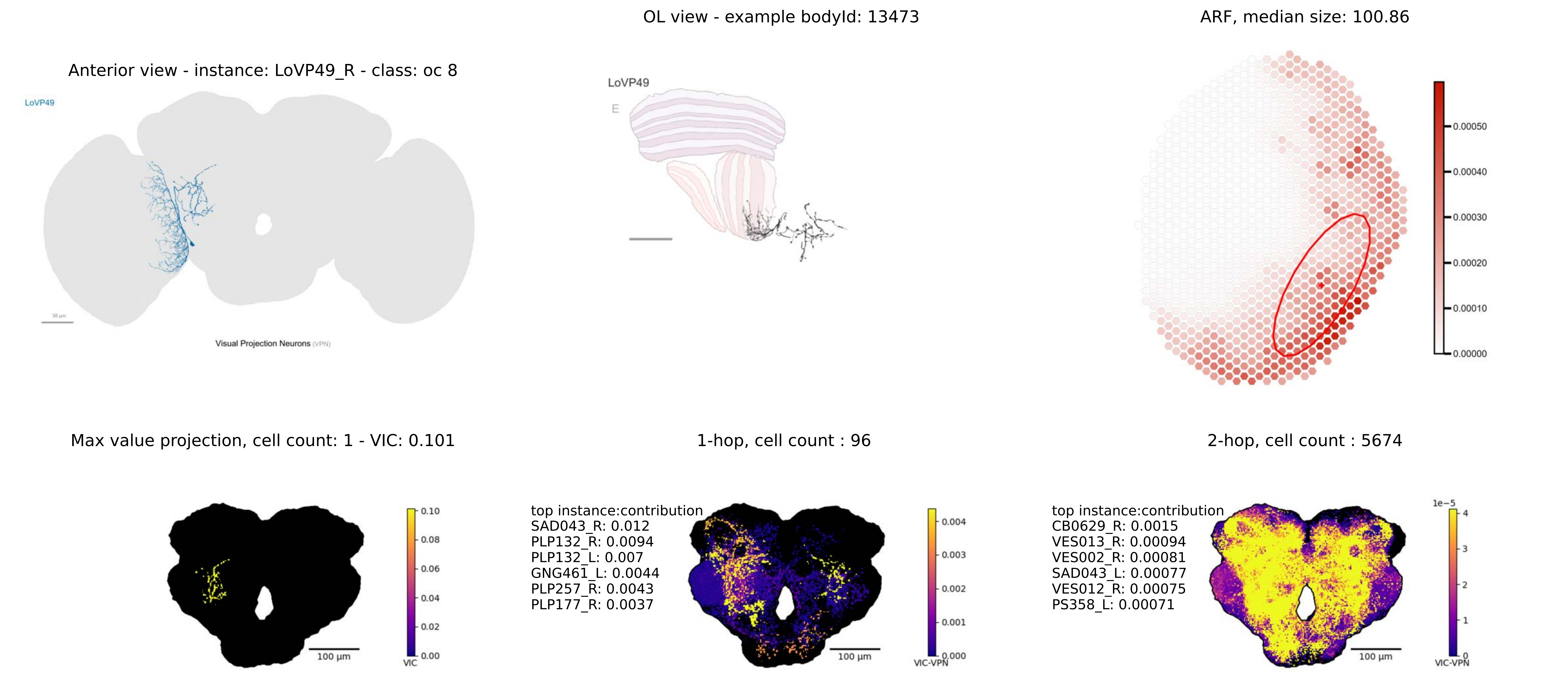

Anterior view - instance: LoVP49_R - class: oc 8
LoVP49
50 µm
Visual Projection Neurons (VPN)
OL view - example bodyId: 13473
LoVP49
E
ARF, median size: 100.86
0.00050
0.00040
0.00030
0.00020
0.00010
0.00000
Max value projection, cell count: 1 - VIC: 0.101
100 µm
0.10
0.08
0.06
0.04
0.02
0.00
VIC
1-hop, cell count : 96
top instance:contribution
SAD043_R: 0.012
PLP132_R: 0.0094
PLP132_L: 0.007
GNG461_L: 0.0044
PLP257_R: 0.0043
PLP177_R: 0.0037
100 µm
0.004
0.003
0.002
0.001
0.000
VIC-VPN
2-hop, cell count : 5674
top instance:contribution
CB0629_R: 0.0015
VES013_R: 0.00094
VES002_R: 0.00081
SAD043_L: 0.00077
VES012_R: 0.00075
PS358_L: 0.00071
100 µm
1e−5
4
3
2
1
0
VIC-VPN

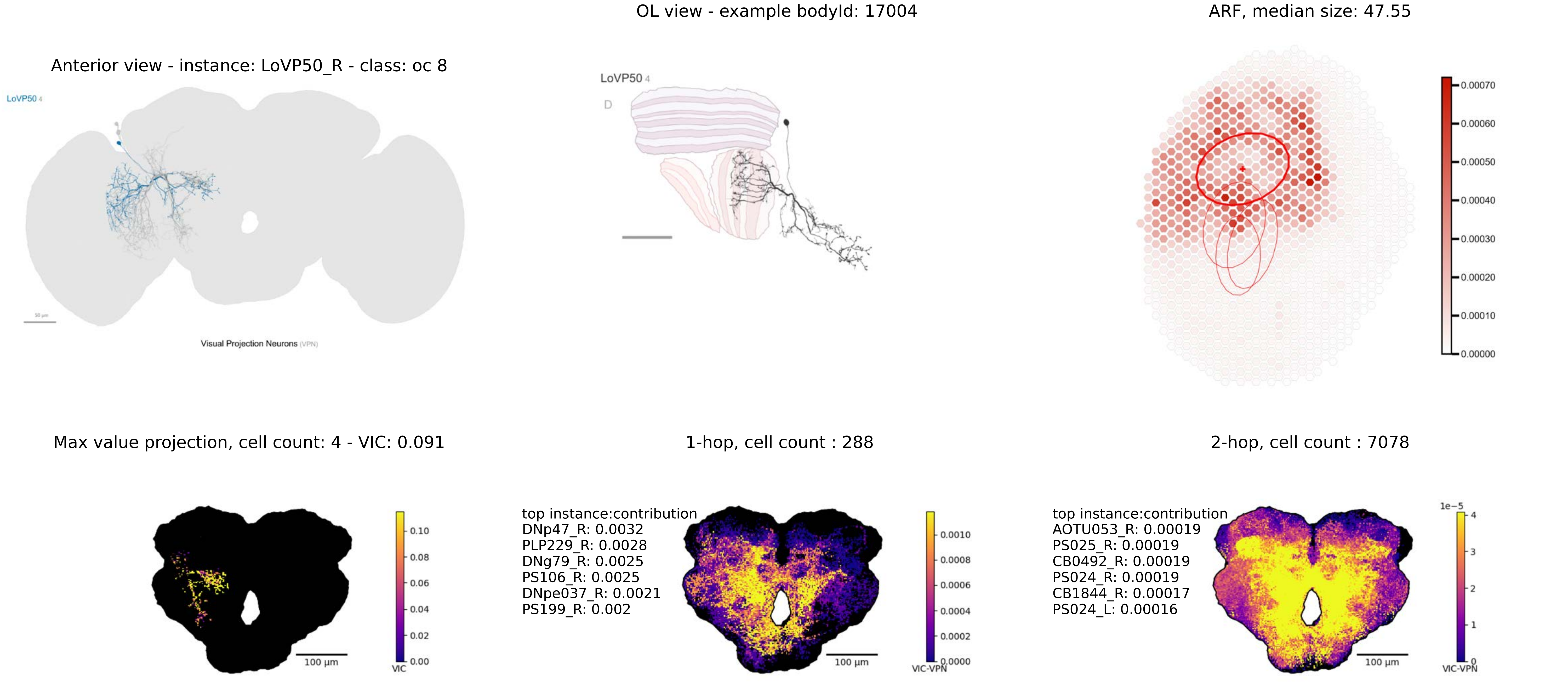
Anterior view - instance: LoVP50_R - class: oc 8
LoVP50 4
Visual Projection Neurons (VPN)
OL view - example bodyId: 17004
LoVP50 4
D
ARF, median size: 47.55
0.00070
0.00060
0.00050
0.00040
0.00030
0.00020
0.00010
0.00000
Max value projection, cell count: 4 - VIC: 0.091
100 µm
0.10
0.08
0.06
0.04
0.02
0.00
VIC
1-hop, cell count : 288
top instance:contribution
DNp47_R: 0.0032
PLP229_R: 0.0028
DNg79_R: 0.0025
PS106_R: 0.0025
DNpe037_R: 0.0021
PS199_R: 0.002
100 µm
0.0010
0.0008
0.0006
0.0004
0.0002
0.0000
VIC-VPN
2-hop, cell count : 7078
top instance:contribution
AOTU053_R: 0.00019
PS025_R: 0.00019
CB0492_R: 0.00019
PS024_R: 0.00019
CB1844_R: 0.00017
PS024_L: 0.00016
100 µm
1e−5
4
3
2
1
0
VIC-VPN

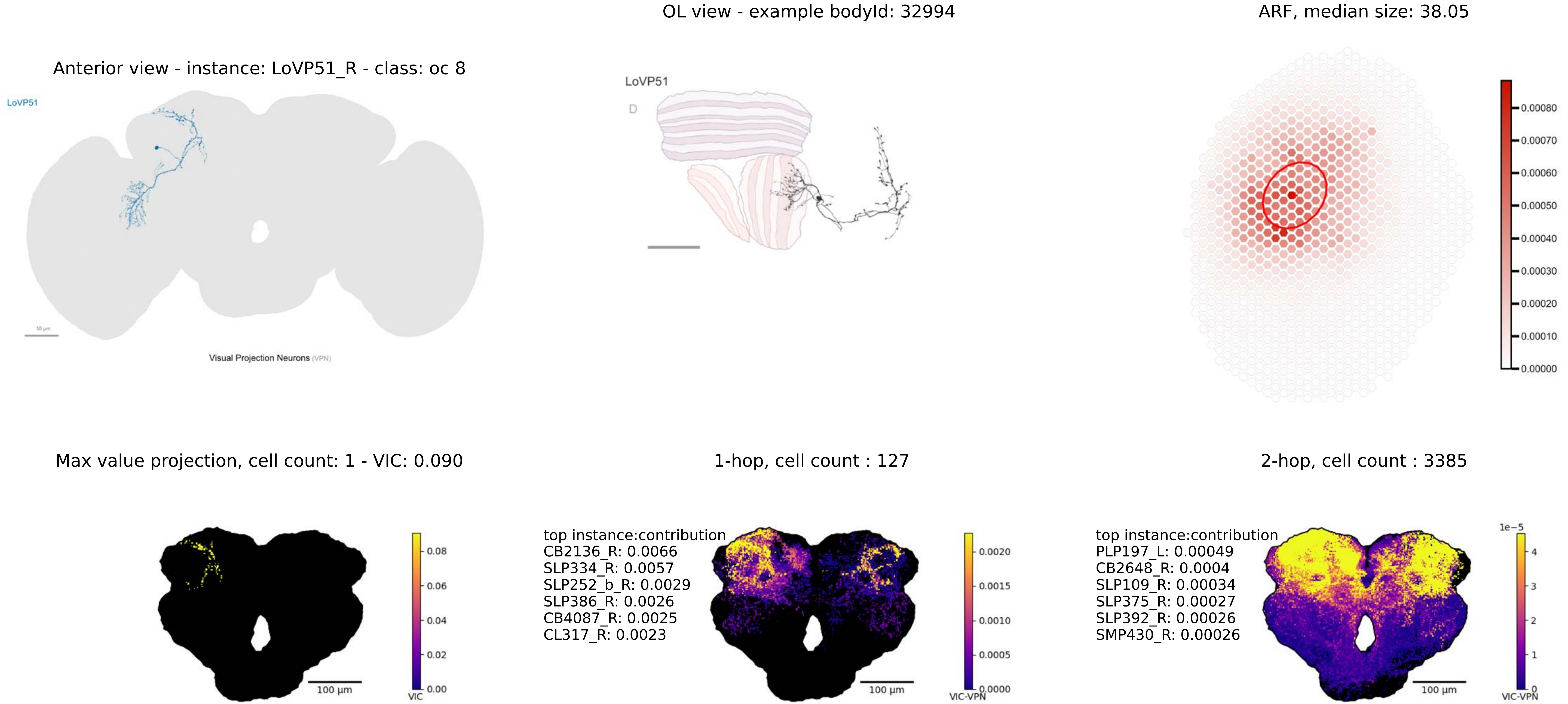
OL view - example bodyId: 32994
ARF, median size: 38.05
Anterior view - instance: LoVP51_R - class: oc 8
LoVP51
Visual Projection Neurons (VPN)
LoVP51
D
0.00080
0.00070
0.00060
0.00050
0.00040
0.00030
0.00020
0.00010
0.00000
Max value projection, cell count: 1 - VIC: 0.090
1-hop, cell count : 127
2-hop, cell count : 3385
top instance:contribution
CB2136_R: 0.0066
SLP334_R: 0.0057
SLP252_b_R: 0.0029
SLP386_R: 0.0026
CB4087_R: 0.0025
CL317_R: 0.0023
top instance:contribution
PLP197_L: 0.00049
CB2648_R: 0.0004
SLP109_R: 0.00034
SLP375_R: 0.00027
SLP392_R: 0.00026
SMP430_R: 0.00026
0.08
0.06
0.04
0.02
0.00
VIC
100 μm
0.0020
0.0015
0.0010
0.0005
0.0000
VIC-VPN
100 μm
1e−5
4
3
2
1
0
VIC-VPN
100 μm

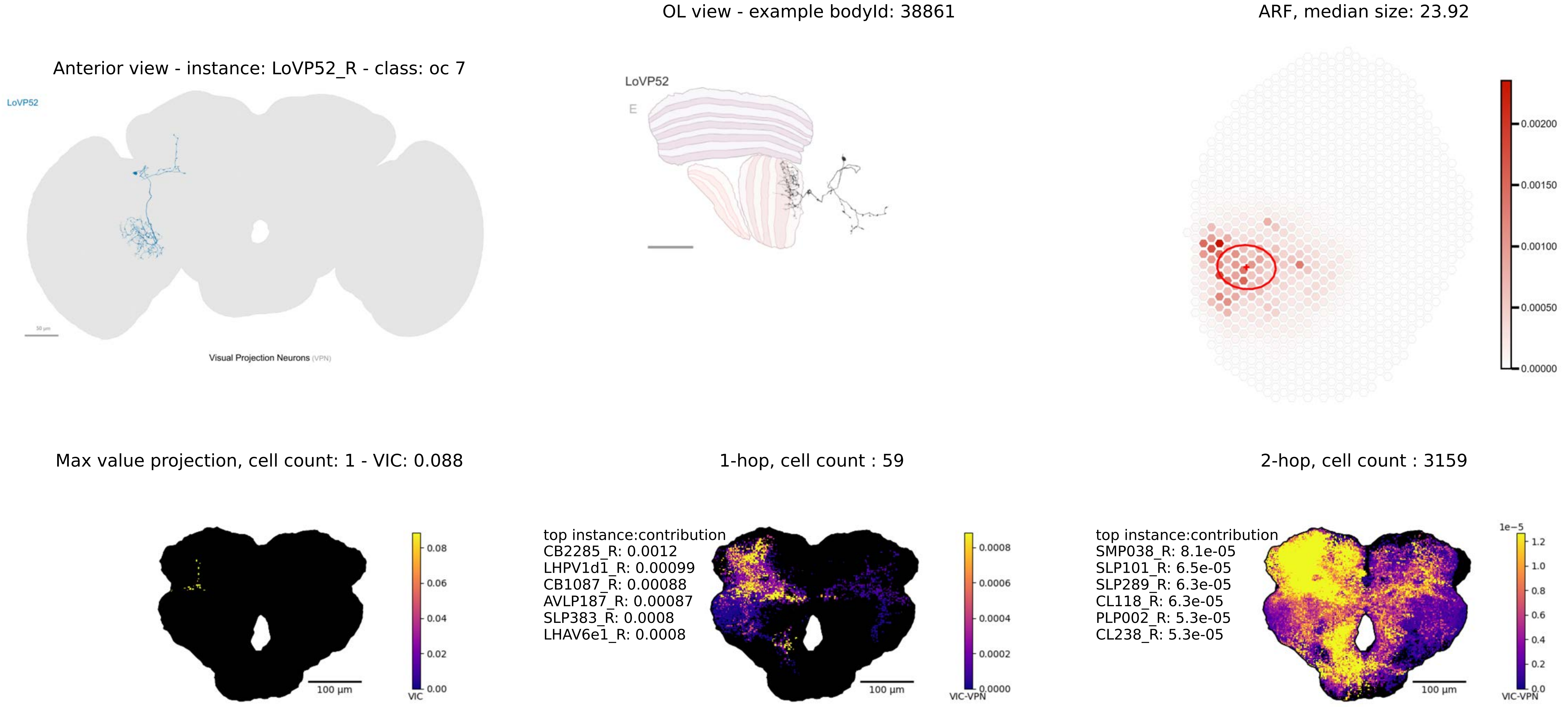
Anterior view - instance: LoVP52_R - class: oc 7
LoVP52
Visual Projection Neurons (VPN)
OL view - example bodyId: 38861
LoVP52
E
ARF, median size: 23.92
0.00200
0.00150
0.00100
0.00050
0.00000
Max value projection, cell count: 1 - VIC: 0.088
100 µm
VIC
0.08
0.06
0.04
0.02
0.00
1-hop, cell count : 59
top instance:contribution
CB2285_R: 0.0012
LHPV1d1_R: 0.00099
CB1087_R: 0.00088
AVLP187_R: 0.00087
SLP383_R: 0.0008
LHAV6e1_R: 0.0008
100 µm
VIC-VPN
0.0008
0.0006
0.0004
0.0002
0.0000
2-hop, cell count : 3159
top instance:contribution
SMP038_R: 8.1e-05
SLP101_R: 6.5e-05
SLP289_R: 6.3e-05
CL118_R: 6.3e-05
PLP002_R: 5.3e-05
CL238_R: 5.3e-05
100 µm
1e−5
1.2
1.0
0.8
0.6
0.4
0.2
0.0
VIC-VPN

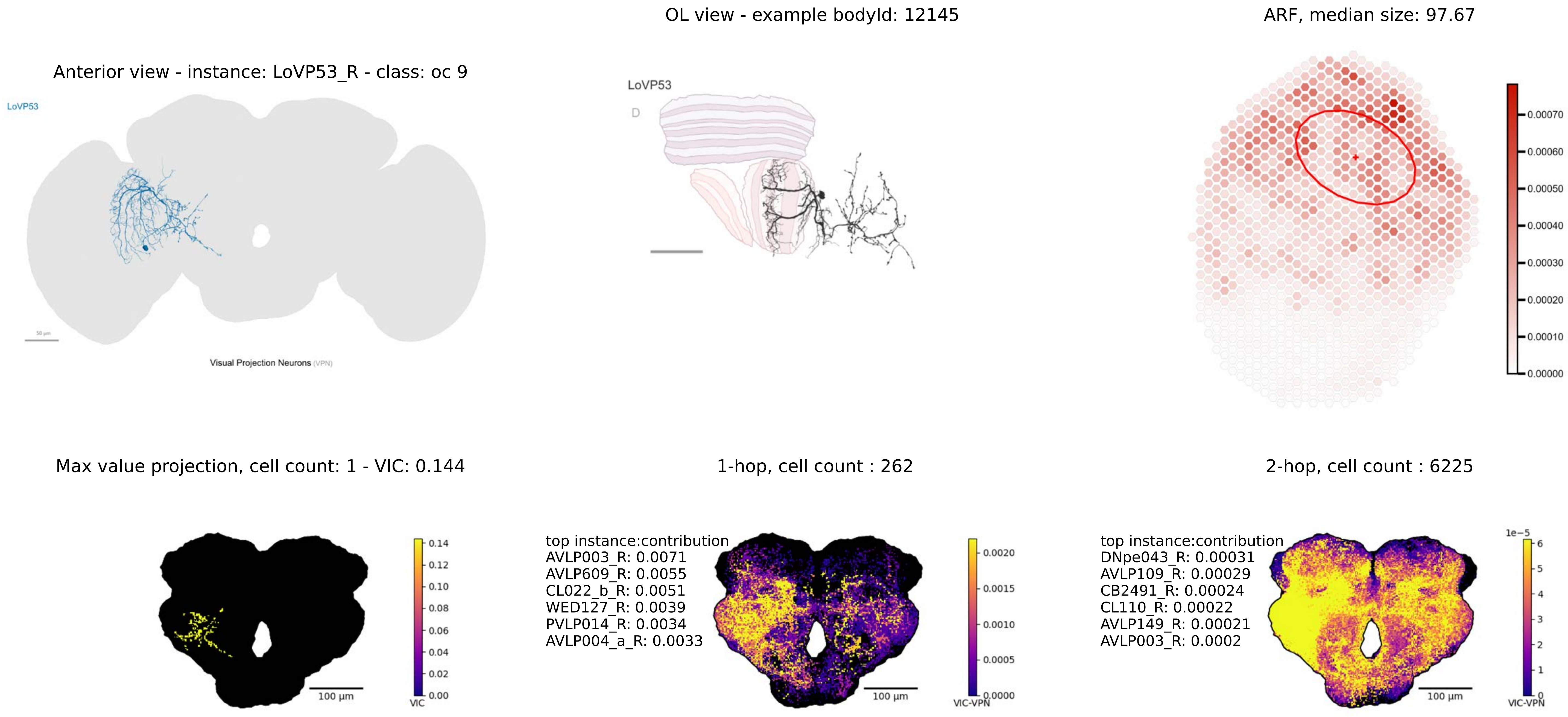
OL view - example bodyId: 12145
ARF, median size: 97.67
Anterior view - instance: LoVP53_R - class: oc 9
LoVP53
Visual Projection Neurons (VPN)
LoVP53
D
0.00070
0.00060
0.00050
0.00040
0.00030
0.00020
0.00010
0.00000
Max value projection, cell count: 1 - VIC: 0.144
1-hop, cell count : 262
2-hop, cell count : 6225
0.14
0.12
0.10
0.08
0.06
0.04
0.02
0.00
VIC
100 µm
top instance:contribution
AVLP003_R: 0.0071
AVLP609_R: 0.0055
CL022_b_R: 0.0051
WED127_R: 0.0039
PVLP014_R: 0.0034
AVLP004_a_R: 0.0033
0.0020
0.0015
0.0010
0.0005
0.0000
VIC-VPN
100 µm
top instance:contribution
DNpe043_R: 0.00031
AVLP109_R: 0.00029
CB2491_R: 0.00024
CL110_R: 0.00022
AVLP149_R: 0.00021
AVLP003_R: 0.0002
1e−5
6
5
4
3
2
1
0
VIC-VPN
100 µm

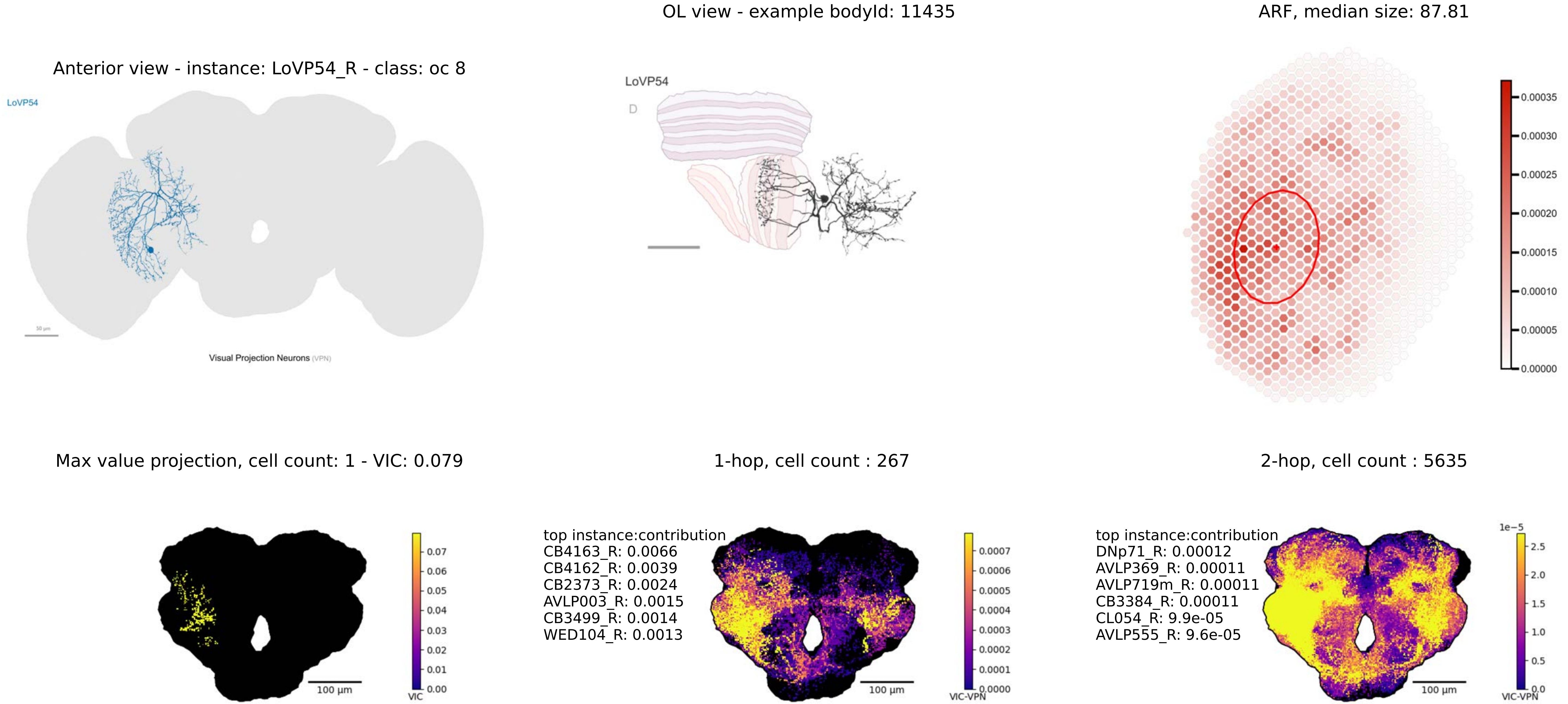
OL view - example bodyId: 11435
ARF, median size: 87.81
Anterior view - instance: LoVP54_R - class: oc 8
LoVP54
LoVP54
D
Visual Projection Neurons (VPN)
0.00035
0.00030
0.00025
0.00020
0.00015
0.00010
0.00005
0.00000
Max value projection, cell count: 1 - VIC: 0.079
1-hop, cell count : 267
2-hop, cell count : 5635
top instance:contribution
CB4163_R: 0.0066
CB4162_R: 0.0039
CB2373_R: 0.0024
AVLP003_R: 0.0015
CB3499_R: 0.0014
WED104_R: 0.0013
top instance:contribution
DNp71_R: 0.00012
AVLP369_R: 0.00011
AVLP719m_R: 0.00011
CB3384_R: 0.00011
CL054_R: 9.9e-05
AVLP555_R: 9.6e-05
100 µm
100 µm
100 µm
VIC
VIC-VPN
VIC-VPN
1e−5

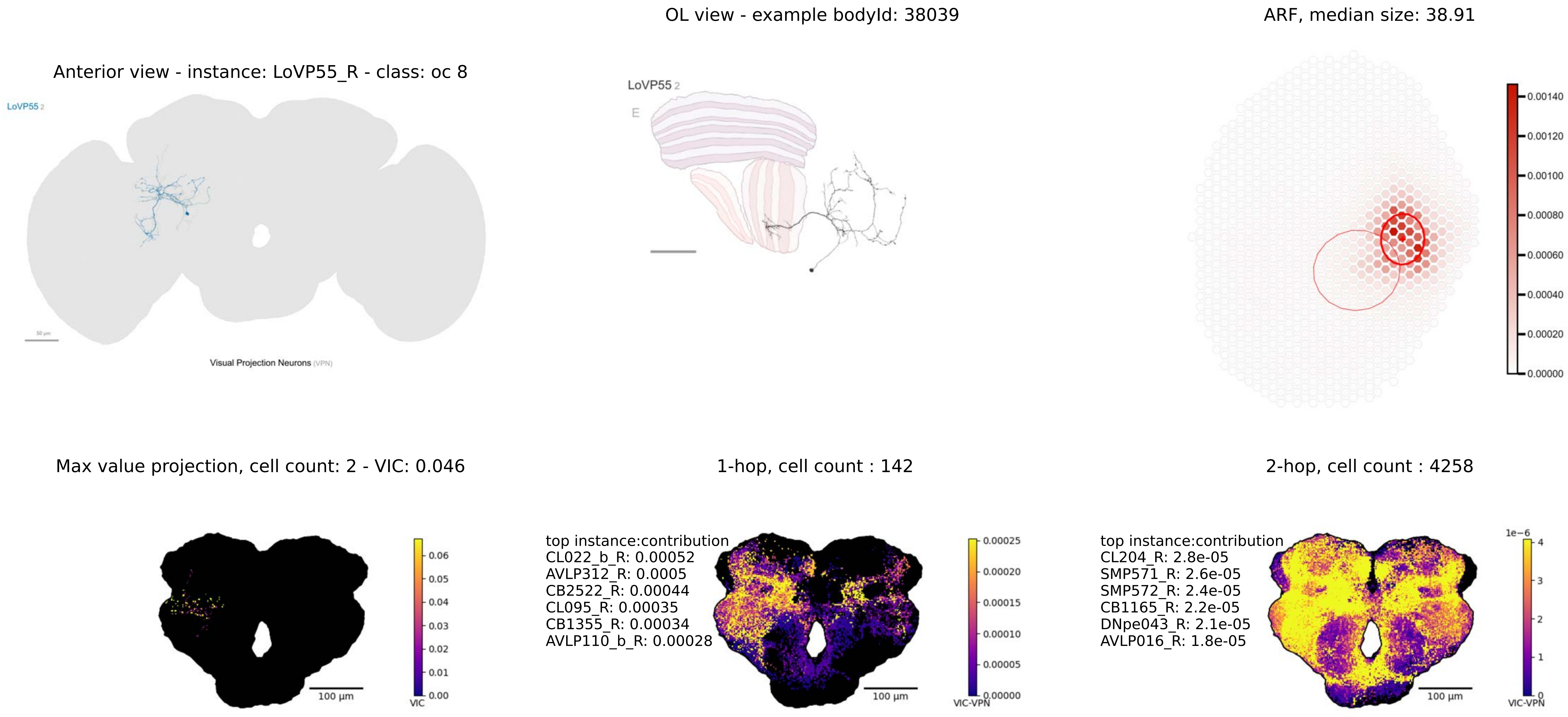
OL view - example bodyId: 38039
ARF, median size: 38.91
Anterior view - instance: LoVP55_R - class: oc 8
LoVP55 2
Visual Projection Neurons (VPN)
LoVP55 2
E
0.00140
0.00120
0.00100
0.00080
0.00060
0.00040
0.00020
0.00000
Max value projection, cell count: 2 - VIC: 0.046
1-hop, cell count : 142
2-hop, cell count : 4258
top instance:contribution
CL022_b_R: 0.00052
AVLP312_R: 0.0005
CB2522_R: 0.00044
CL095_R: 0.00035
CB1355_R: 0.00034
AVLP110_b_R: 0.00028
top instance:contribution
CL204_R: 2.8e-05
SMP571_R: 2.6e-05
SMP572_R: 2.4e-05
CB1165_R: 2.2e-05
DNpe043_R: 2.1e-05
AVLP016_R: 1.8e-05
100 µm
100 µm
100 µm
VIC
VIC-VPN
VIC-VPN
1e-6

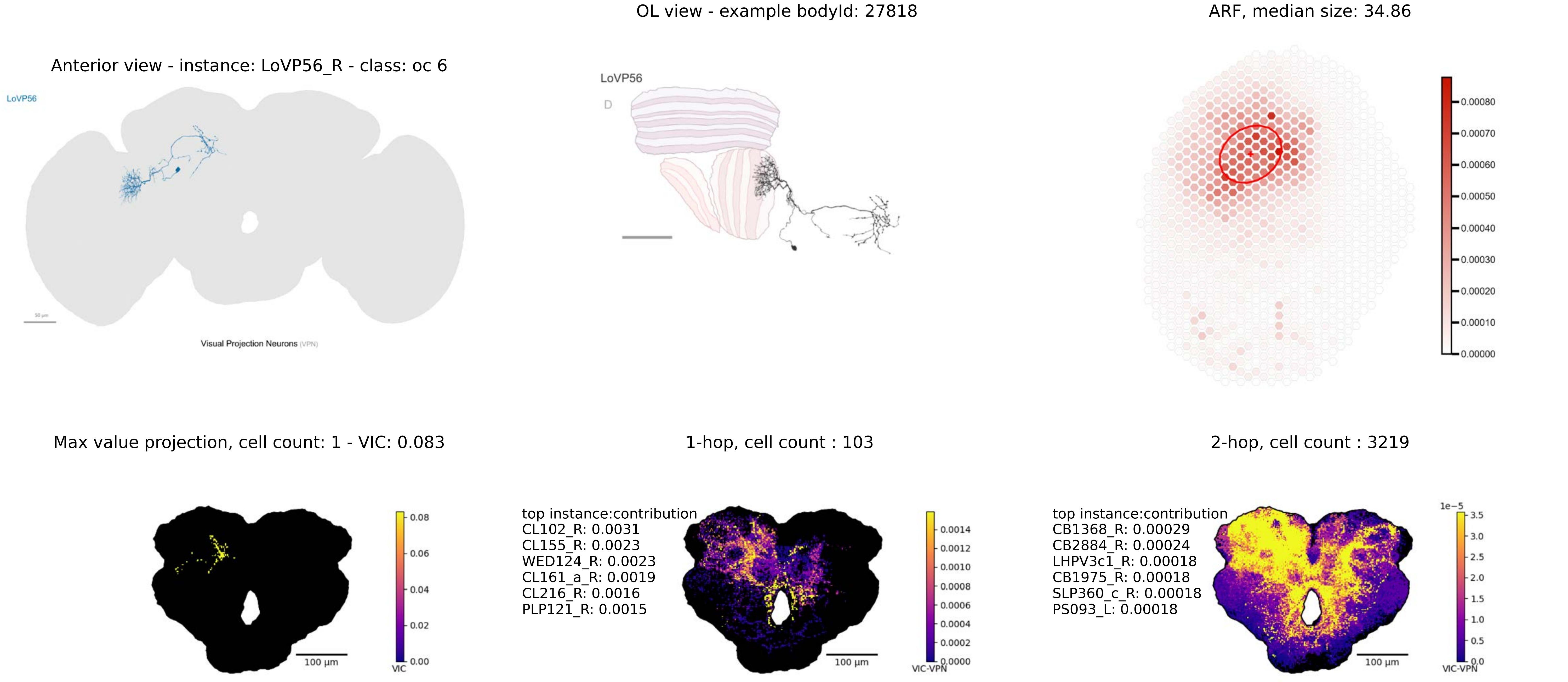
OL view - example bodyId: 27818
ARF, median size: 34.86
Anterior view - instance: LoVP56_R - class: oc 6
LoVP56
LoVP56
D
Visual Projection Neurons (VPN)
0.00080
0.00070
0.00060
0.00050
0.00040
0.00030
0.00020
0.00010
0.00000
Max value projection, cell count: 1 - VIC: 0.083
1-hop, cell count : 103
2-hop, cell count : 3219
0.08
0.06
0.04
0.02
0.00
VIC
100 µm
top instance:contribution
CL102_R: 0.0031
CL155_R: 0.0023
WED124_R: 0.0023
CL161_a_R: 0.0019
CL216_R: 0.0016
PLP121_R: 0.0015
0.0014
0.0012
0.0010
0.0008
0.0006
0.0004
0.0002
0.0000
VIC-VPN
100 µm
top instance:contribution
CB1368_R: 0.00029
CB2884_R: 0.00024
LHPV3c1_R: 0.00018
CB1975_R: 0.00018
SLP360_c_R: 0.00018
PS093_L: 0.00018
1e−5
3.5
3.0
2.5
2.0
1.5
1.0
0.5
0.0
VIC-VPN
100 µm

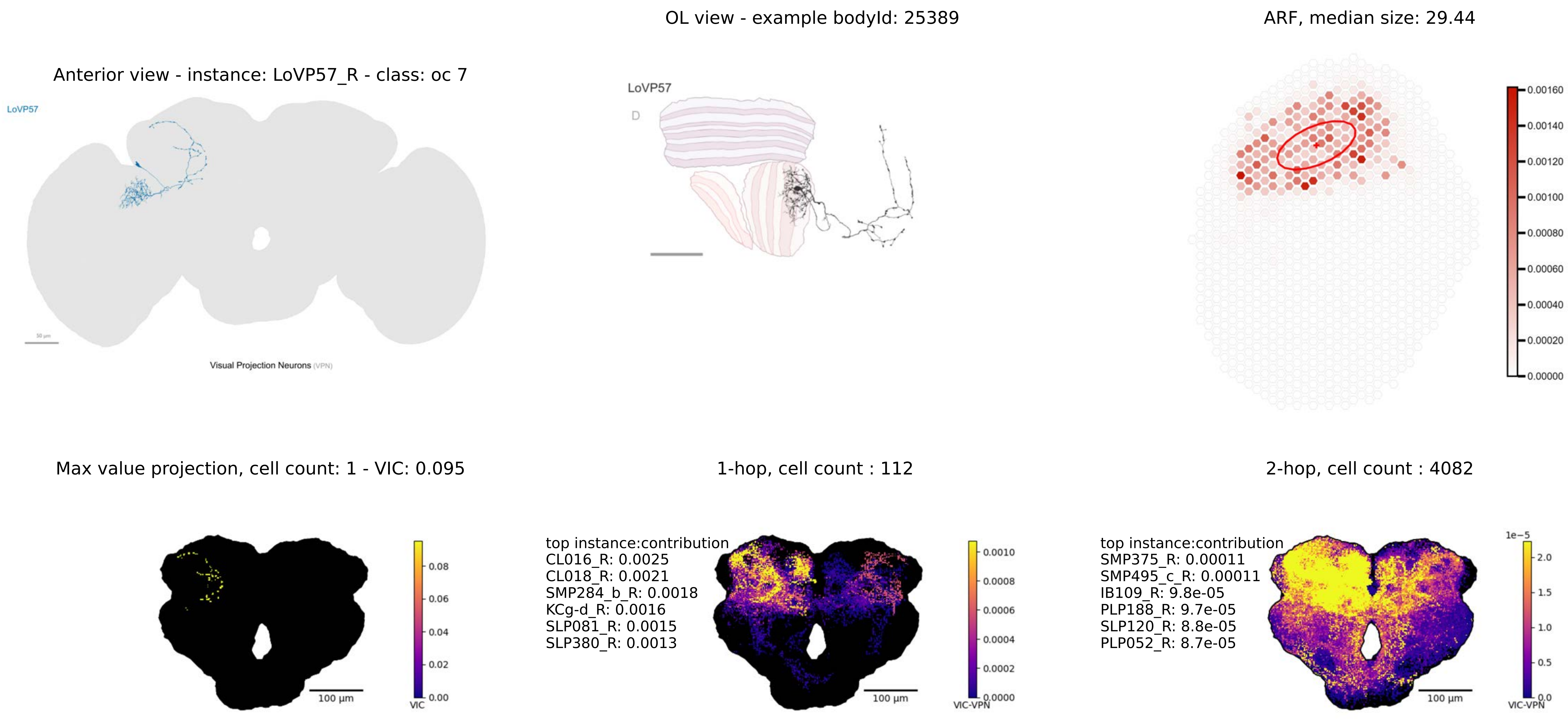
OL view - example bodyId: 25389
ARF, median size: 29.44
Anterior view - instance: LoVP57_R - class: oc 7
LoVP57
Visual Projection Neurons (VPN)
LoVP57
D
0.00160
0.00140
0.00120
0.00100
0.00080
0.00060
0.00040
0.00020
0.00000
Max value projection, cell count: 1 - VIC: 0.095
1-hop, cell count : 112
2-hop, cell count : 4082
0.08
0.06
0.04
0.02
0.00
VIC
100 µm
top instance:contribution
CL016_R: 0.0025
CL018_R: 0.0021
SMP284_b_R: 0.0018
KCg-d_R: 0.0016
SLP081_R: 0.0015
SLP380_R: 0.0013
0.0010
0.0008
0.0006
0.0004
0.0002
0.0000
VIC-VPN
100 µm
top instance:contribution
SMP375_R: 0.00011
SMP495_c_R: 0.00011
IB109_R: 9.8e-05
PLP188_R: 9.7e-05
SLP120_R: 8.8e-05
PLP052_R: 8.7e-05
1e−5
2.0
1.5
1.0
0.5
0.0
VIC-VPN
100 µm

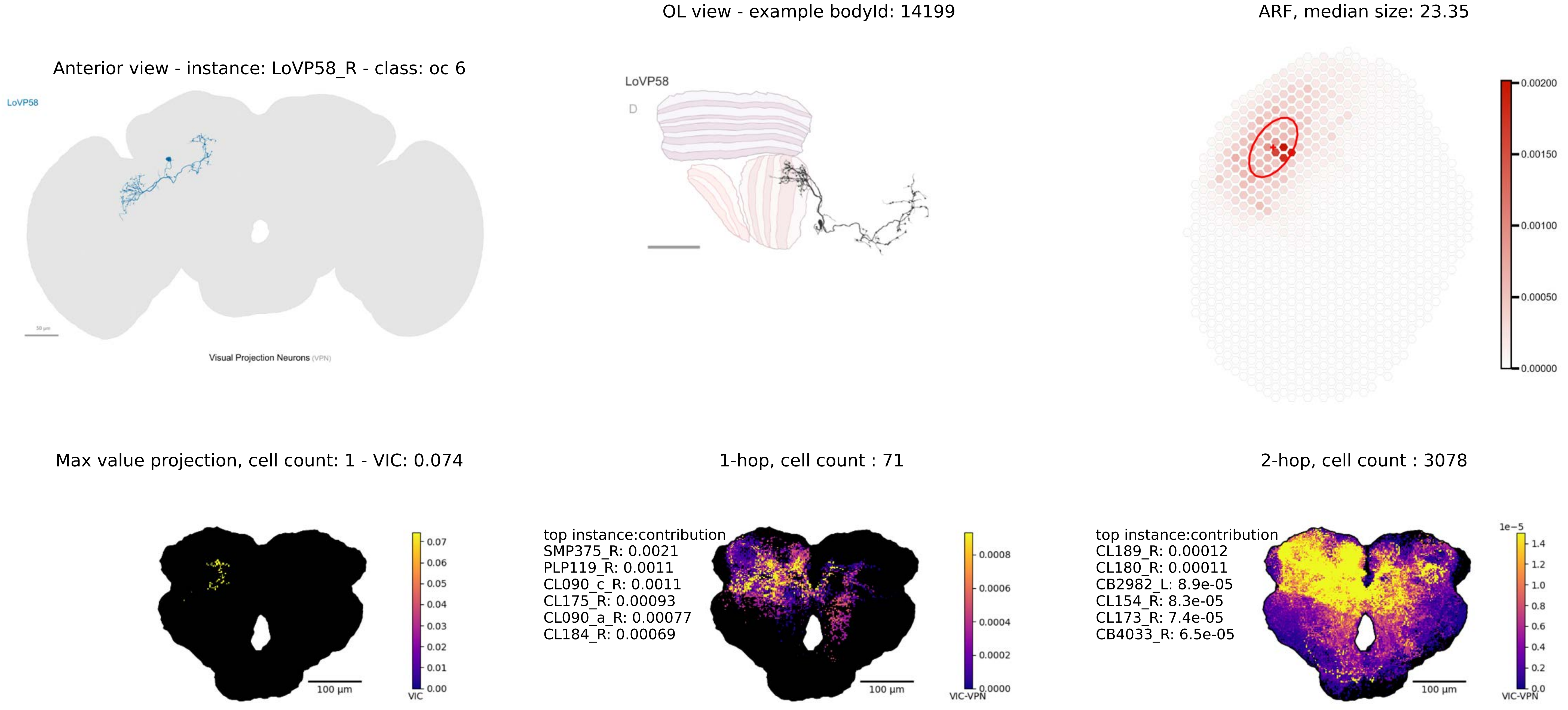
OL view - example bodyId: 14199
ARF, median size: 23.35
Anterior view - instance: LoVP58_R - class: oc 6
LoVP58
Visual Projection Neurons (VPN)
LoVP58
D
0.00200
0.00150
0.00100
0.00050
0.00000
Max value projection, cell count: 1 - VIC: 0.074
1-hop, cell count : 71
2-hop, cell count : 3078
top instance:contribution
SMP375_R: 0.0021
PLP119_R: 0.0011
CL090_c_R: 0.0011
CL175_R: 0.00093
CL090_a_R: 0.00077
CL184_R: 0.00069
top instance:contribution
CL189_R: 0.00012
CL180_R: 0.00011
CB2982_L: 8.9e-05
CL154_R: 8.3e-05
CL173_R: 7.4e-05
CB4033_R: 6.5e-05
100 µm
VIC
VIC-VPN
1e−5

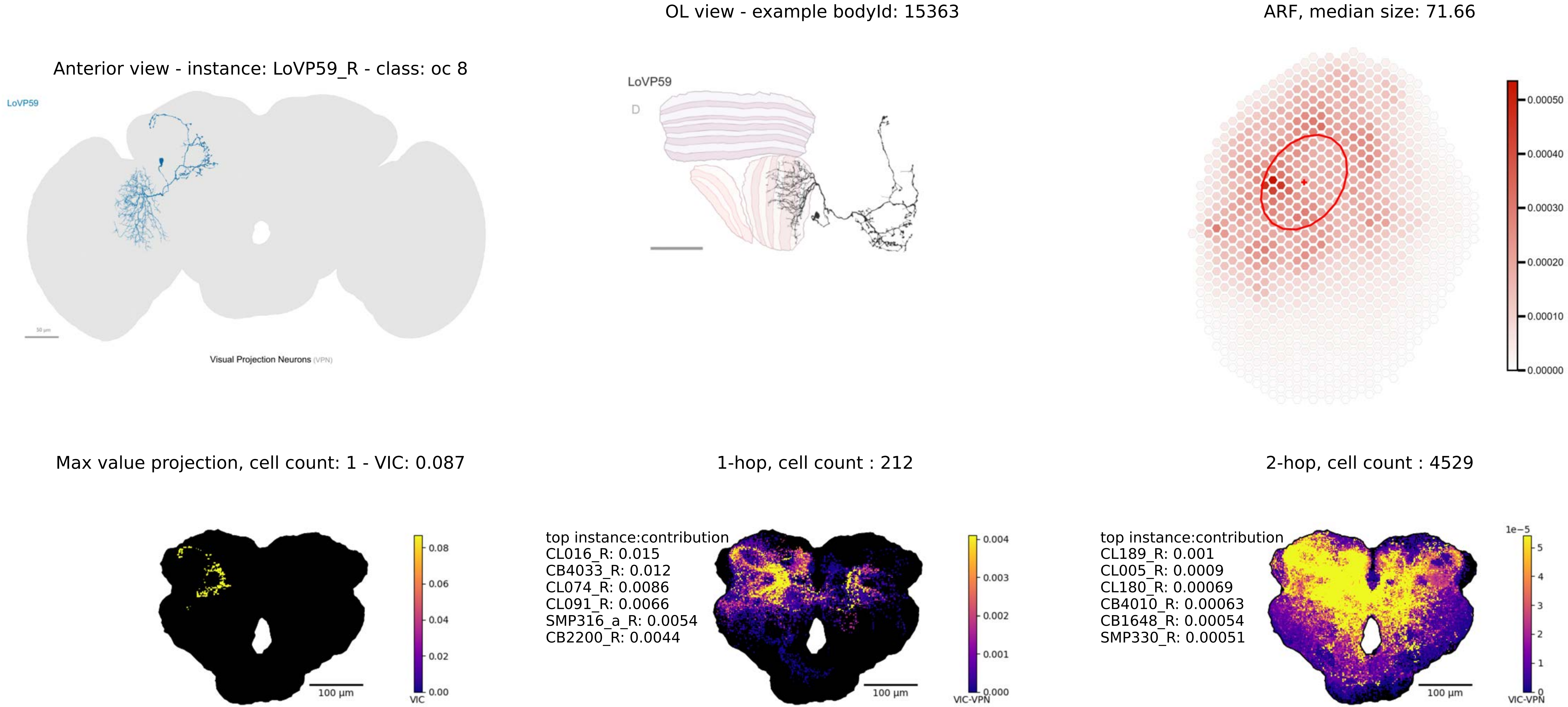
Anterior view - instance: LoVP59_R - class: oc 8
LoVP59
Visual Projection Neurons (VPN)
OL view - example bodyId: 15363
LoVP59
D
ARF, median size: 71.66
0.00050
0.00040
0.00030
0.00020
0.00010
0.00000
Max value projection, cell count: 1 - VIC: 0.087
100 µm
VIC
0.08
0.06
0.04
0.02
0.00
1-hop, cell count : 212
top instance:contribution
CL016_R: 0.015
CB4033_R: 0.012
CL074_R: 0.0086
CL091_R: 0.0066
SMP316_a_R: 0.0054
CB2200_R: 0.0044
100 µm
VIC-VPN
0.004
0.003
0.002
0.001
0.000
2-hop, cell count : 4529
top instance:contribution
CL189_R: 0.001
CL005_R: 0.0009
CL180_R: 0.00069
CB4010_R: 0.00063
CB1648_R: 0.00054
SMP330_R: 0.00051
100 µm
1e−5
5
4
3
2
1
0
VIC-VPN

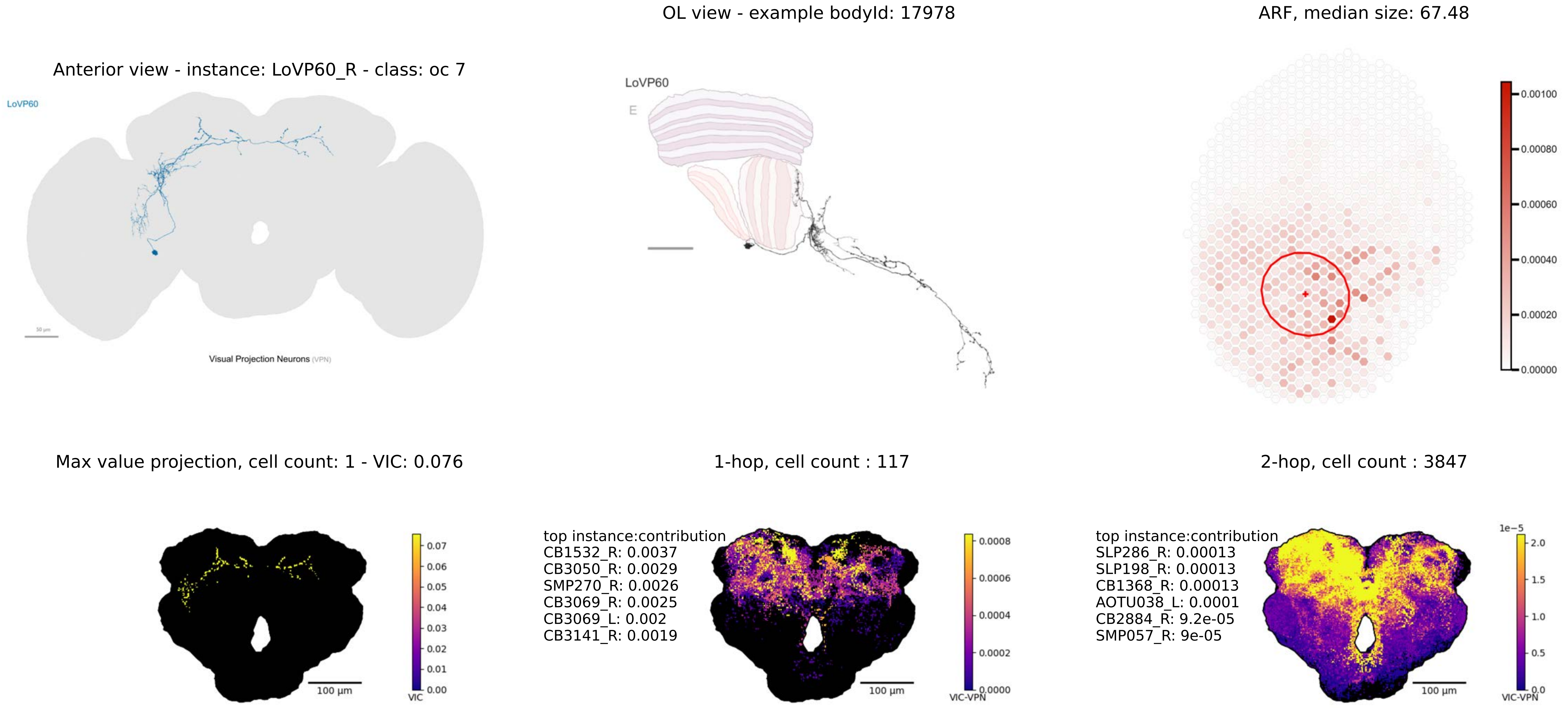
OL view - example bodyId: 17978
ARF, median size: 67.48
Anterior view - instance: LoVP60_R - class: oc 7
LoVP60
50 µm
Visual Projection Neurons (VPN)
LoVP60
E
0.00100
0.00080
0.00060
0.00040
0.00020
0.00000
Max value projection, cell count: 1 - VIC: 0.076
1-hop, cell count : 117
2-hop, cell count : 3847
0.07
0.06
0.05
0.04
0.03
0.02
0.01
0.00
VIC
100 µm
top instance:contribution
CB1532_R: 0.0037
CB3050_R: 0.0029
SMP270_R: 0.0026
CB3069_R: 0.0025
CB3069_L: 0.002
CB3141_R: 0.0019
0.0008
0.0006
0.0004
0.0002
0.0000
VIC-VPN
100 µm
top instance:contribution
SLP286_R: 0.00013
SLP198_R: 0.00013
CB1368_R: 0.00013
AOTU038_L: 0.0001
CB2884_R: 9.2e-05
SMP057_R: 9e-05
1e−5
2.0
1.5
1.0
0.5
0.0
VIC-VPN
100 µm

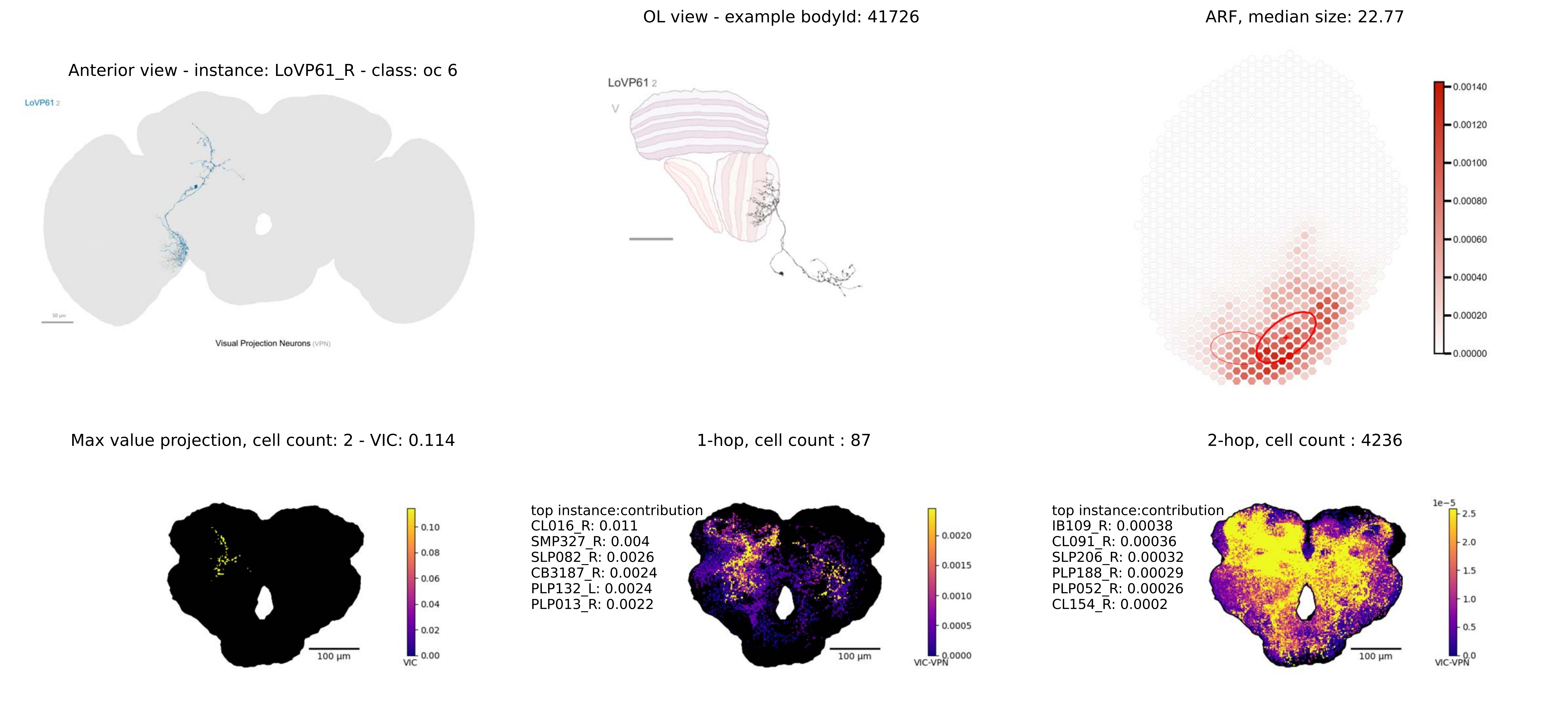

OL view - example bodyId: 41726
ARF, median size: 22.77
Anterior view - instance: LoVP61_R - class: oc 6
LoVP61 2
Visual Projection Neurons (VPN)
LoVP61 2
V
0.00140
0.00120
0.00100
0.00080
0.00060
0.00040
0.00020
0.00000
Max value projection, cell count: 2 - VIC: 0.114
1-hop, cell count : 87
2-hop, cell count : 4236
0.10
0.08
0.06
0.04
0.02
0.00
VIC
100 µm
top instance:contribution
CL016_R: 0.011
SMP327_R: 0.004
SLP082_R: 0.0026
CB3187_R: 0.0024
PLP132_L: 0.0024
PLP013_R: 0.0022
0.0020
0.0015
0.0010
0.0005
0.0000
VIC-VPN
100 µm
top instance:contribution
IB109_R: 0.00038
CL091_R: 0.00036
SLP206_R: 0.00032
PLP188_R: 0.00029
PLP052_R: 0.00026
CL154_R: 0.0002
1e−5
2.5
2.0
1.5
1.0
0.5
0.0
VIC-VPN
100 µm

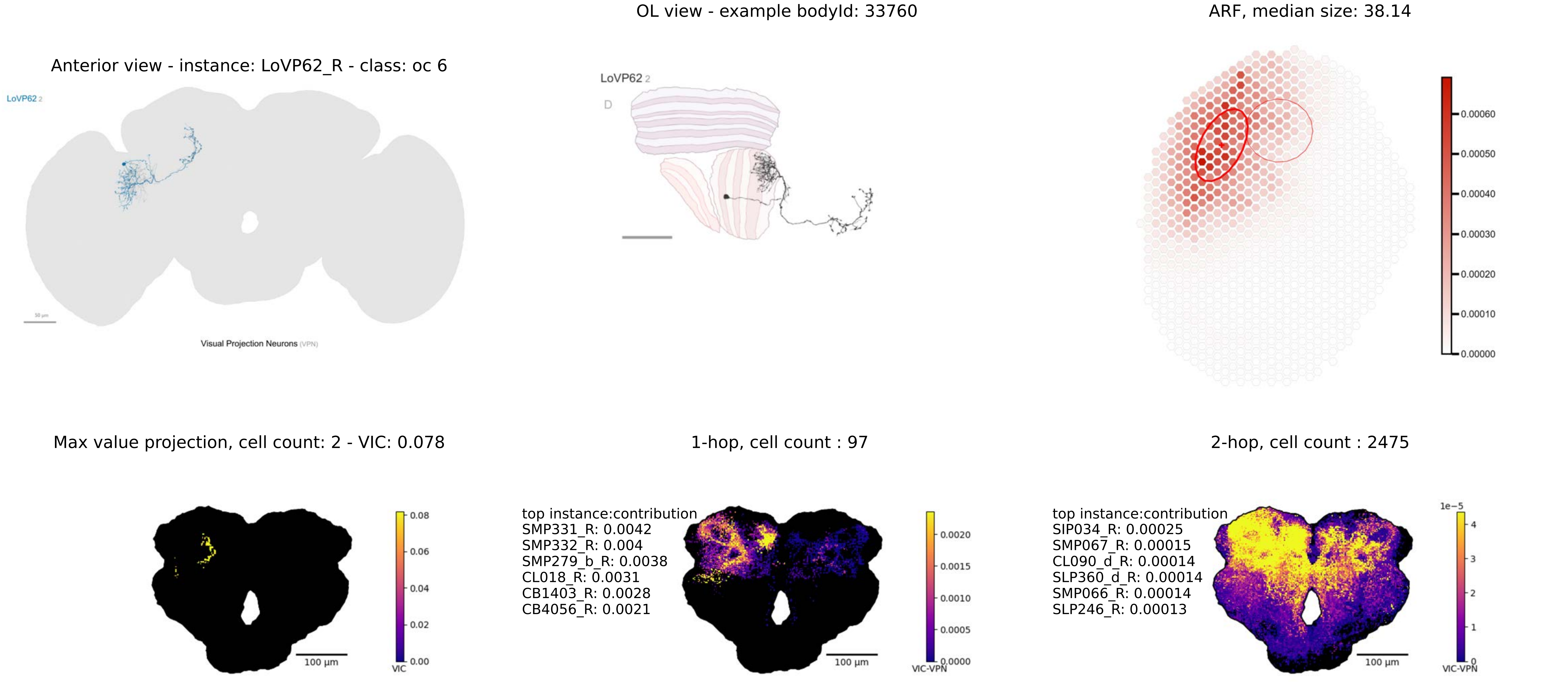
Anterior view - instance: LoVP62_R - class: oc 6
LoVP62 2
Visual Projection Neurons (VPN)
OL view - example bodyId: 33760
LoVP62 2
D
ARF, median size: 38.14
0.00060
0.00050
0.00040
0.00030
0.00020
0.00010
0.00000
Max value projection, cell count: 2 - VIC: 0.078
0.08
0.06
0.04
0.02
0.00
VIC
100 μm
1-hop, cell count : 97
top instance:contribution
SMP331_R: 0.0042
SMP332_R: 0.004
SMP279_b_R: 0.0038
CL018_R: 0.0031
CB1403_R: 0.0028
CB4056_R: 0.0021
0.0020
0.0015
0.0010
0.0005
0.0000
VIC-VPN
100 μm
2-hop, cell count : 2475
top instance:contribution
SIP034_R: 0.00025
SMP067_R: 0.00015
CL090_d_R: 0.00014
SLP360_d_R: 0.00014
SMP066_R: 0.00014
SLP246_R: 0.00013
1e−5
4
3
2
1
0
VIC-VPN
100 μm

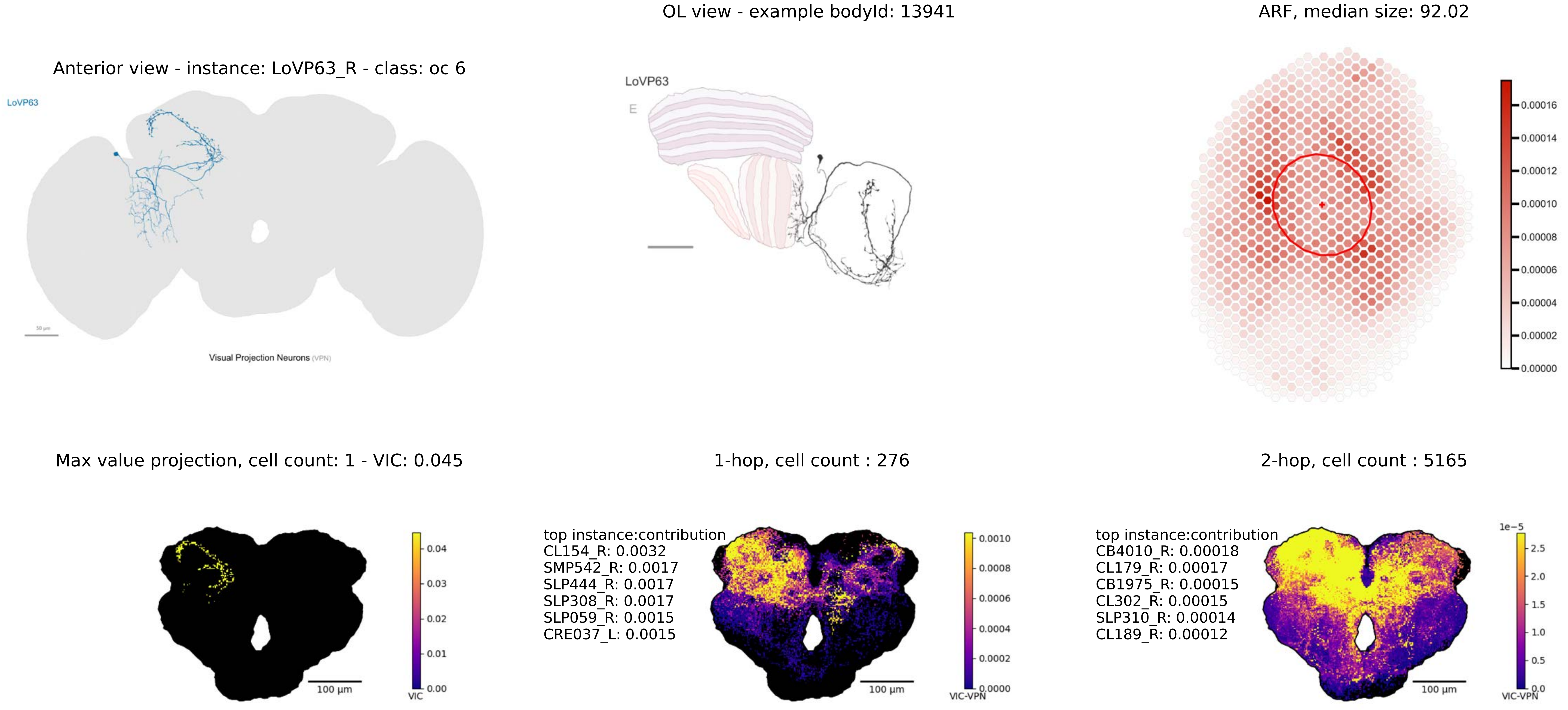

OL view - example bodyId: 13941
ARF, median size: 92.02
Anterior view - instance: LoVP63_R - class: oc 6
LoVP63
Visual Projection Neurons (VPN)
LoVP63
E
0.00016
0.00014
0.00012
0.00010
0.00008
0.00006
0.00004
0.00002
0.00000
Max value projection, cell count: 1 - VIC: 0.045
1-hop, cell count : 276
2-hop, cell count : 5165
0.04
0.03
0.02
0.01
0.00
VIC
100 µm
top instance:contribution
CL154_R: 0.0032
SMP542_R: 0.0017
SLP444_R: 0.0017
SLP308_R: 0.0017
SLP059_R: 0.0015
CRE037_L: 0.0015
0.0010
0.0008
0.0006
0.0004
0.0002
0.0000
VIC-VPN
100 µm
top instance:contribution
CB4010_R: 0.00018
CL179_R: 0.00017
CB1975_R: 0.00015
CL302_R: 0.00015
SLP310_R: 0.00014
CL189_R: 0.00012
1e−5
2.5
2.0
1.5
1.0
0.5
0.0
VIC-VPN
100 µm

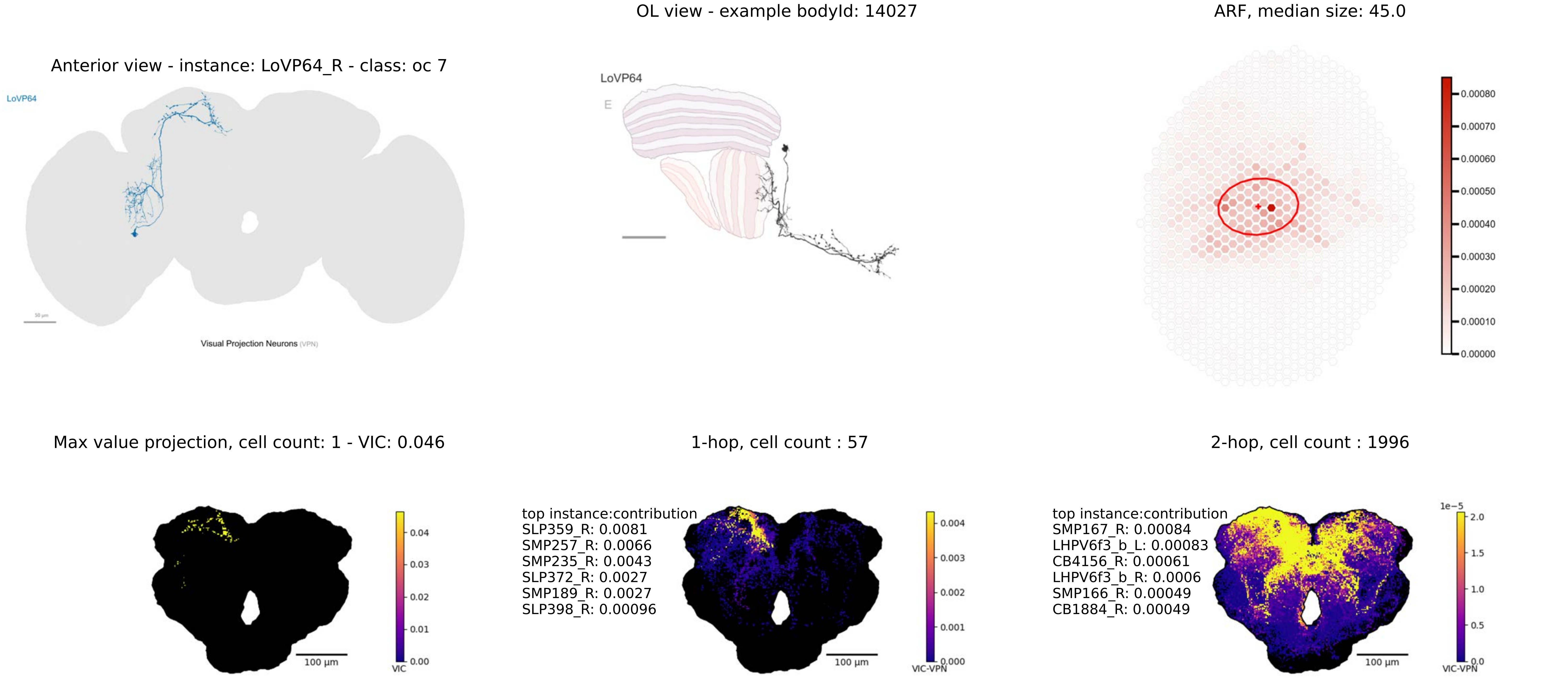
OL view - example bodyId: 14027
ARF, median size: 45.0
Anterior view - instance: LoVP64_R - class: oc 7
LoVP64
Visual Projection Neurons (VPN)
LoVP64
E
0.00080
0.00070
0.00060
0.00050
0.00040
0.00030
0.00020
0.00010
0.00000
Max value projection, cell count: 1 - VIC: 0.046
1-hop, cell count : 57
2-hop, cell count : 1996
0.04
0.03
0.02
0.01
0.00
VIC
100 µm
top instance:contribution
SLP359_R: 0.0081
SMP257_R: 0.0066
SMP235_R: 0.0043
SLP372_R: 0.0027
SMP189_R: 0.0027
SLP398_R: 0.00096
0.004
0.003
0.002
0.001
0.000
VIC-VPN
100 µm
top instance:contribution
SMP167_R: 0.00084
LHPV6f3_b_L: 0.00083
CB4156_R: 0.00061
LHPV6f3_b_R: 0.0006
SMP166_R: 0.00049
CB1884_R: 0.00049
1e−5
2.0
1.5
1.0
0.5
0.0
VIC-VPN
100 µm

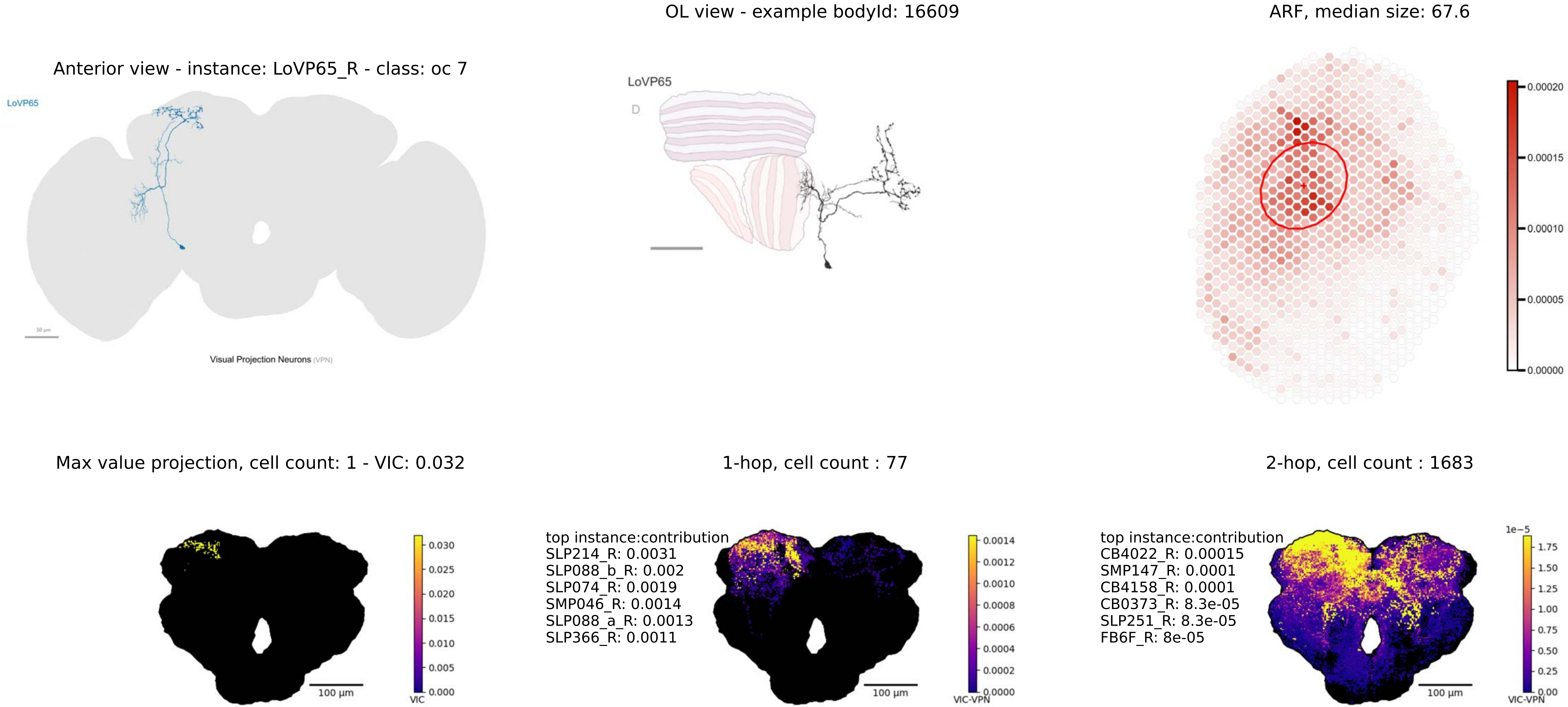

OL view - example bodyId: 16609
ARF, median size: 67.6
Anterior view - instance: LoVP65_R - class: oc 7
LoVP65
Visual Projection Neurons (VPN)
LoVP65
D
0.00020
0.00015
0.00010
0.00005
0.00000
Max value projection, cell count: 1 - VIC: 0.032
1-hop, cell count : 77
2-hop, cell count : 1683
0.030
0.025
0.020
0.015
0.010
0.005
0.000
VIC
100 µm
top instance:contribution
SLP214_R: 0.0031
SLP088_b_R: 0.002
SLP074_R: 0.0019
SMP046_R: 0.0014
SLP088_a_R: 0.0013
SLP366_R: 0.0011
0.0014
0.0012
0.0010
0.0008
0.0006
0.0004
0.0002
0.0000
VIC-VPN
100 µm
top instance:contribution
CB4022_R: 0.00015
SMP147_R: 0.0001
CB4158_R: 0.0001
CB0373_R: 8.3e-05
SLP251_R: 8.3e-05
FB6F_R: 8e-05
1e−5
1.75
1.50
1.25
1.00
0.75
0.50
0.25
0.00
VIC-VPN
100 µm

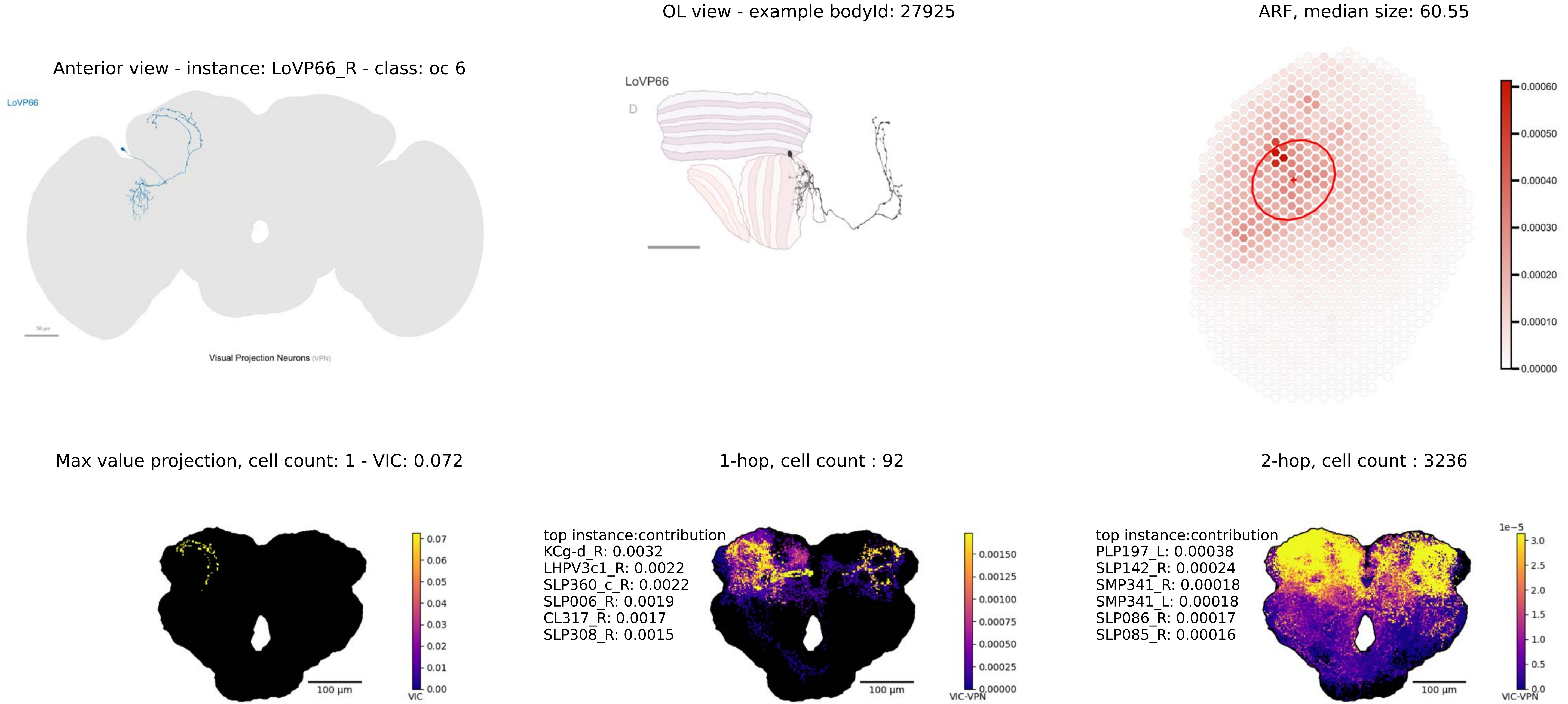
OL view - example bodyId: 27925
ARF, median size: 60.55
Anterior view - instance: LoVP66_R - class: oc 6
LoVP66
Visual Projection Neurons (VPN)
LoVP66
D
0.00060
0.00050
0.00040
0.00030
0.00020
0.00010
0.00000
Max value projection, cell count: 1 - VIC: 0.072
1-hop, cell count : 92
2-hop, cell count : 3236
top instance:contribution
KCg-d_R: 0.0032
LHPV3c1_R: 0.0022
SLP360_c_R: 0.0022
SLP006_R: 0.0019
CL317_R: 0.0017
SLP308_R: 0.0015
top instance:contribution
PLP197_L: 0.00038
SLP142_R: 0.00024
SMP341_R: 0.00018
SMP341_L: 0.00018
SLP086_R: 0.00017
SLP085_R: 0.00016
100 µm
VIC
VIC-VPN
1e−5

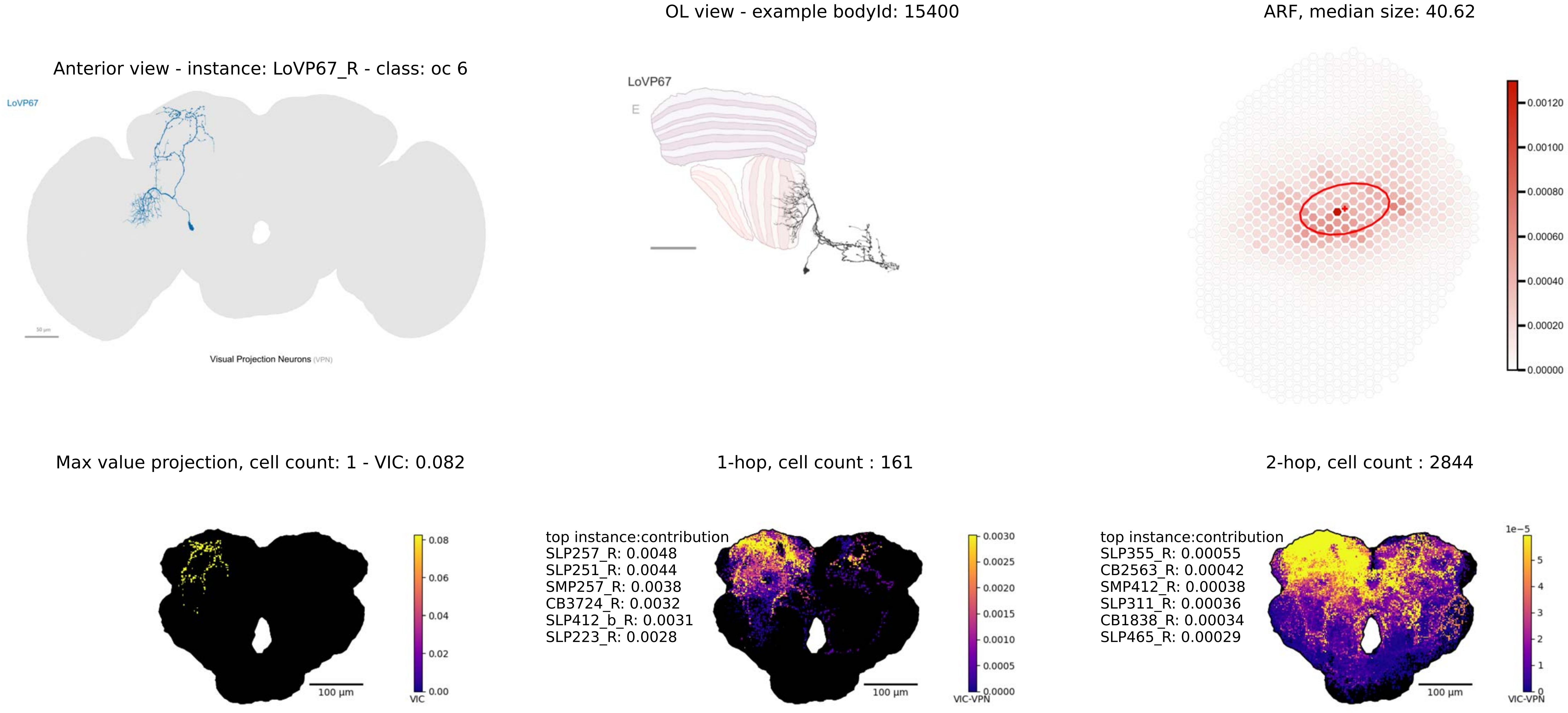

OL view - example bodyId: 15400
ARF, median size: 40.62
Anterior view - instance: LoVP67_R - class: oc 6
LoVP67
Visual Projection Neurons (VPN)
LoVP67
E
0.00120
0.00100
0.00080
0.00060
0.00040
0.00020
0.00000
Max value projection, cell count: 1 - VIC: 0.082
1-hop, cell count : 161
2-hop, cell count : 2844
0.08
0.06
0.04
0.02
0.00
VIC
100 μm
top instance:contribution
SLP257_R: 0.0048
SLP251_R: 0.0044
SMP257_R: 0.0038
CB3724_R: 0.0032
SLP412_b_R: 0.0031
SLP223_R: 0.0028
0.0030
0.0025
0.0020
0.0015
0.0010
0.0005
0.0000
VIC-VPN
100 μm
top instance:contribution
SLP355_R: 0.00055
CB2563_R: 0.00042
SMP412_R: 0.00038
SLP311_R: 0.00036
CB1838_R: 0.00034
SLP465_R: 0.00029
1e−5
5
4
3
2
1
0
VIC-VPN
100 μm

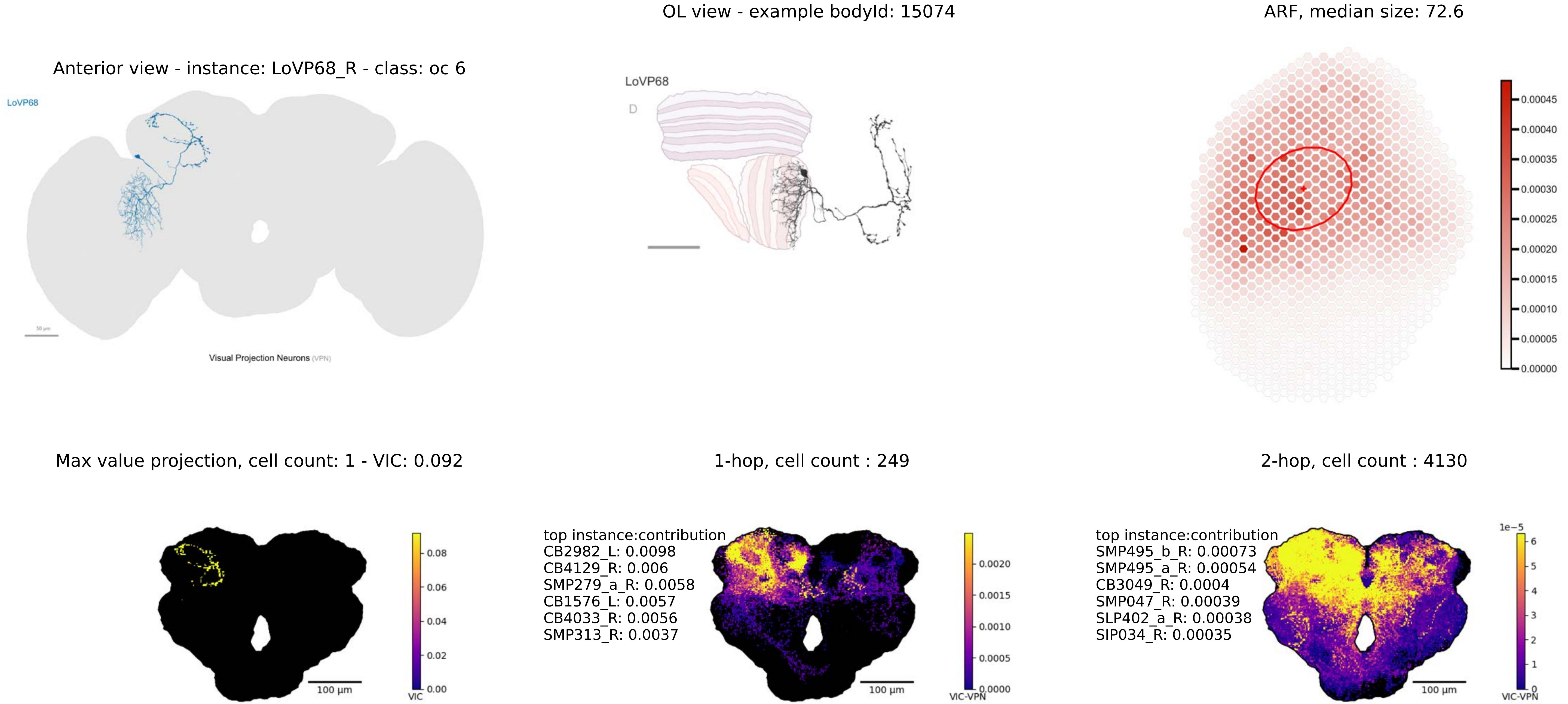

Anterior view - instance: LoVP68_R - class: oc 6
LoVP68
Visual Projection Neurons (VPN)
OL view - example bodyId: 15074
LoVP68
D
ARF, median size: 72.6
0.00045
0.00040
0.00035
0.00030
0.00025
0.00020
0.00015
0.00010
0.00005
0.00000
Max value projection, cell count: 1 - VIC: 0.092
100 µm
VIC
0.08
0.06
0.04
0.02
0.00
1-hop, cell count : 249
top instance:contribution
CB2982_L: 0.0098
CB4129_R: 0.006
SMP279_a_R: 0.0058
CB1576_L: 0.0057
CB4033_R: 0.0056
SMP313_R: 0.0037
100 µm
VIC-VPN
0.0020
0.0015
0.0010
0.0005
0.0000
2-hop, cell count : 4130
top instance:contribution
SMP495_b_R: 0.00073
SMP495_a_R: 0.00054
CB3049_R: 0.0004
SMP047_R: 0.00039
SLP402_a_R: 0.00038
SIP034_R: 0.00035
100 µm
1e−5
6
5
4
3
2
1
0
VIC-VPN

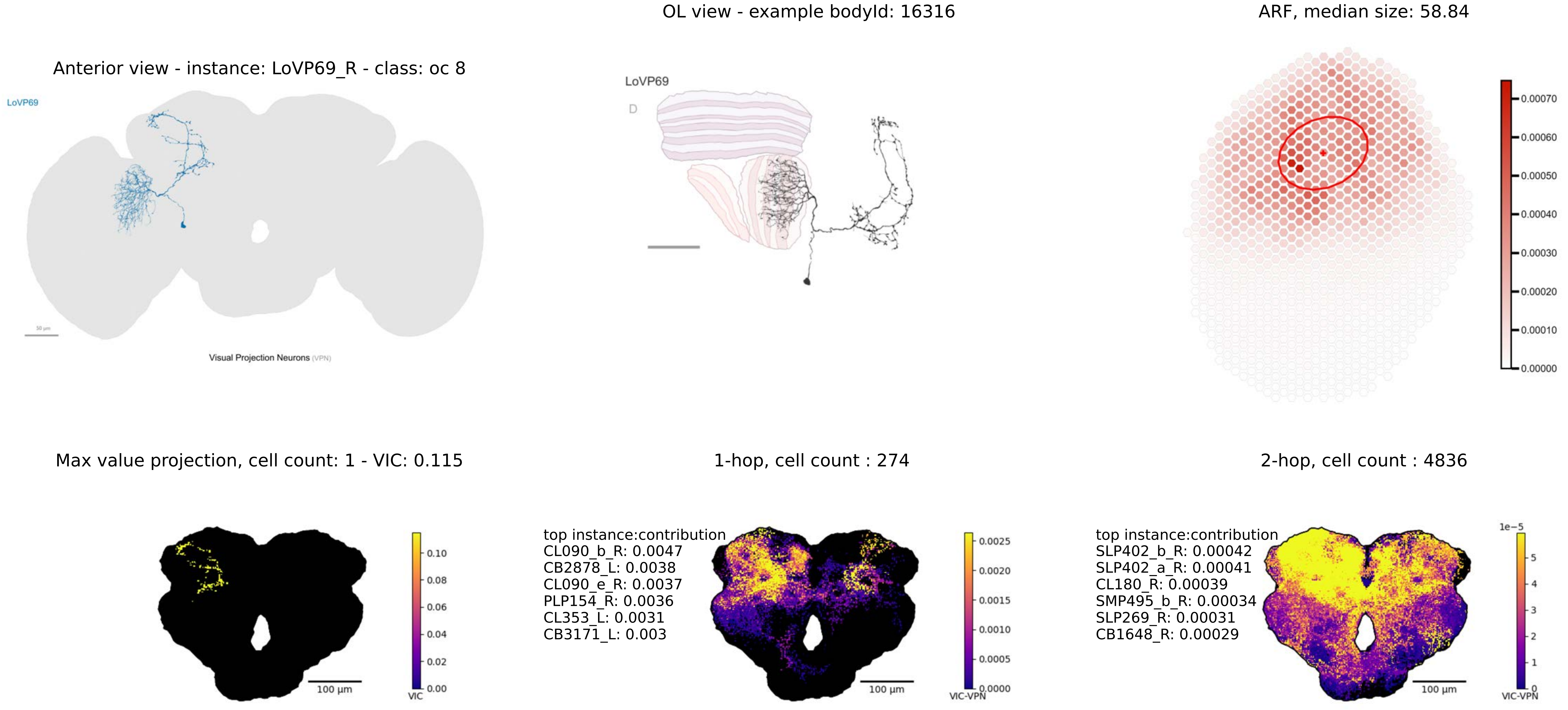

OL view - example bodyId: 16316
ARF, median size: 58.84
Anterior view - instance: LoVP69_R - class: oc 8
LoVP69
Visual Projection Neurons (VPN)
LoVP69
D
0.00070
0.00060
0.00050
0.00040
0.00030
0.00020
0.00010
0.00000
Max value projection, cell count: 1 - VIC: 0.115
1-hop, cell count : 274
2-hop, cell count : 4836
0.10
0.08
0.06
0.04
0.02
0.00
VIC
100 µm
top instance:contribution
CL090_b_R: 0.0047
CB2878_L: 0.0038
CL090_e_R: 0.0037
PLP154_R: 0.0036
CL353_L: 0.0031
CB3171_L: 0.003
0.0025
0.0020
0.0015
0.0010
0.0005
0.0000
VIC-VPN
100 µm
top instance:contribution
SLP402_b_R: 0.00042
SLP402_a_R: 0.00041
CL180_R: 0.00039
SMP495_b_R: 0.00034
SLP269_R: 0.00031
CB1648_R: 0.00029
1e−5
5
4
3
2
1
0
VIC-VPN
100 µm

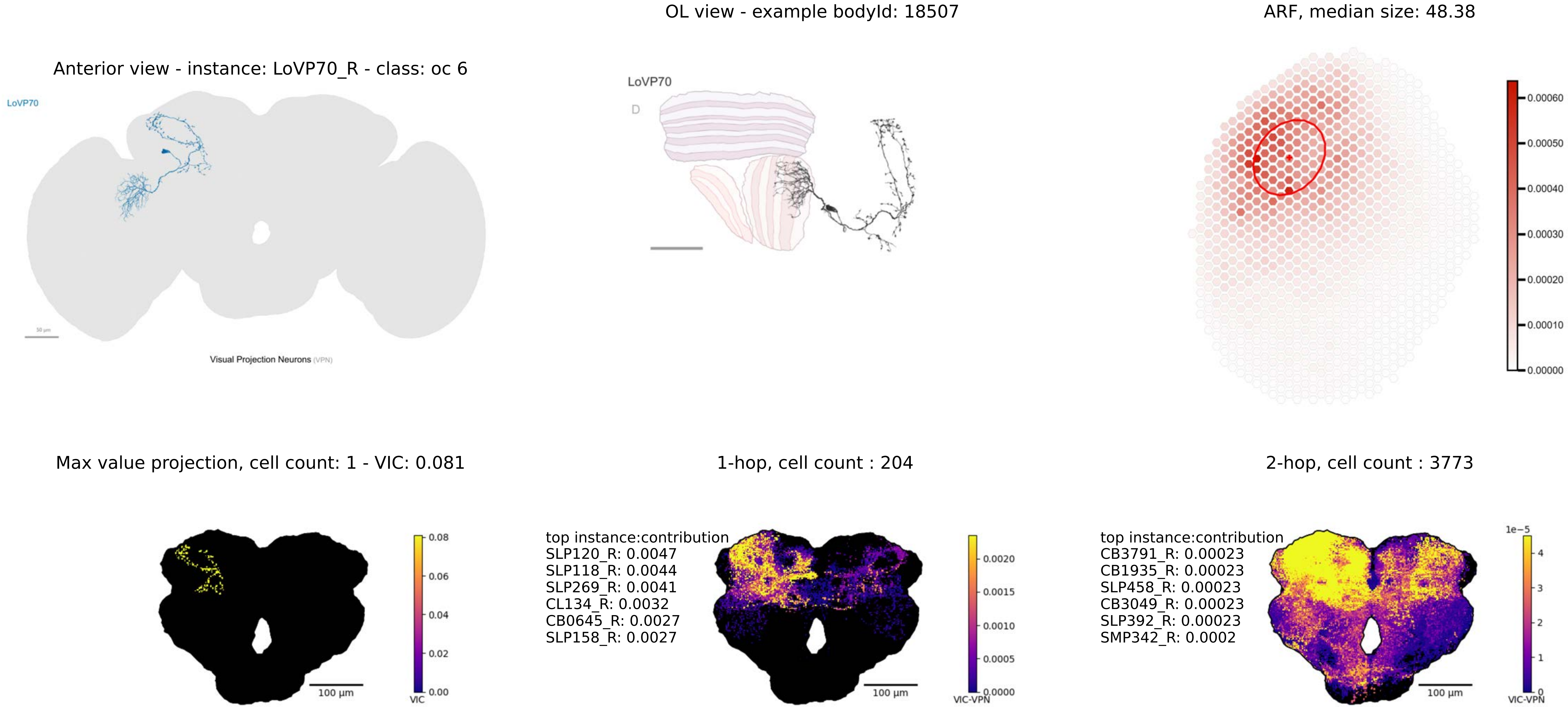
OL view - example bodyId: 18507
ARF, median size: 48.38
Anterior view - instance: LoVP70_R - class: oc 6
LoVP70
Visual Projection Neurons (VPN)
LoVP70
D
0.00060
0.00050
0.00040
0.00030
0.00020
0.00010
0.00000
Max value projection, cell count: 1 - VIC: 0.081
1-hop, cell count : 204
2-hop, cell count : 3773
100 µm
VIC
0.08
0.06
0.04
0.02
0.00
top instance:contribution
SLP120_R: 0.0047
SLP118_R: 0.0044
SLP269_R: 0.0041
CL134_R: 0.0032
CB0645_R: 0.0027
SLP158_R: 0.0027
VIC-VPN
0.0020
0.0015
0.0010
0.0005
0.0000
top instance:contribution
CB3791_R: 0.00023
CB1935_R: 0.00023
SLP458_R: 0.00023
CB3049_R: 0.00023
SLP392_R: 0.00023
SMP342_R: 0.0002
1e−5
4
3
2
1
0

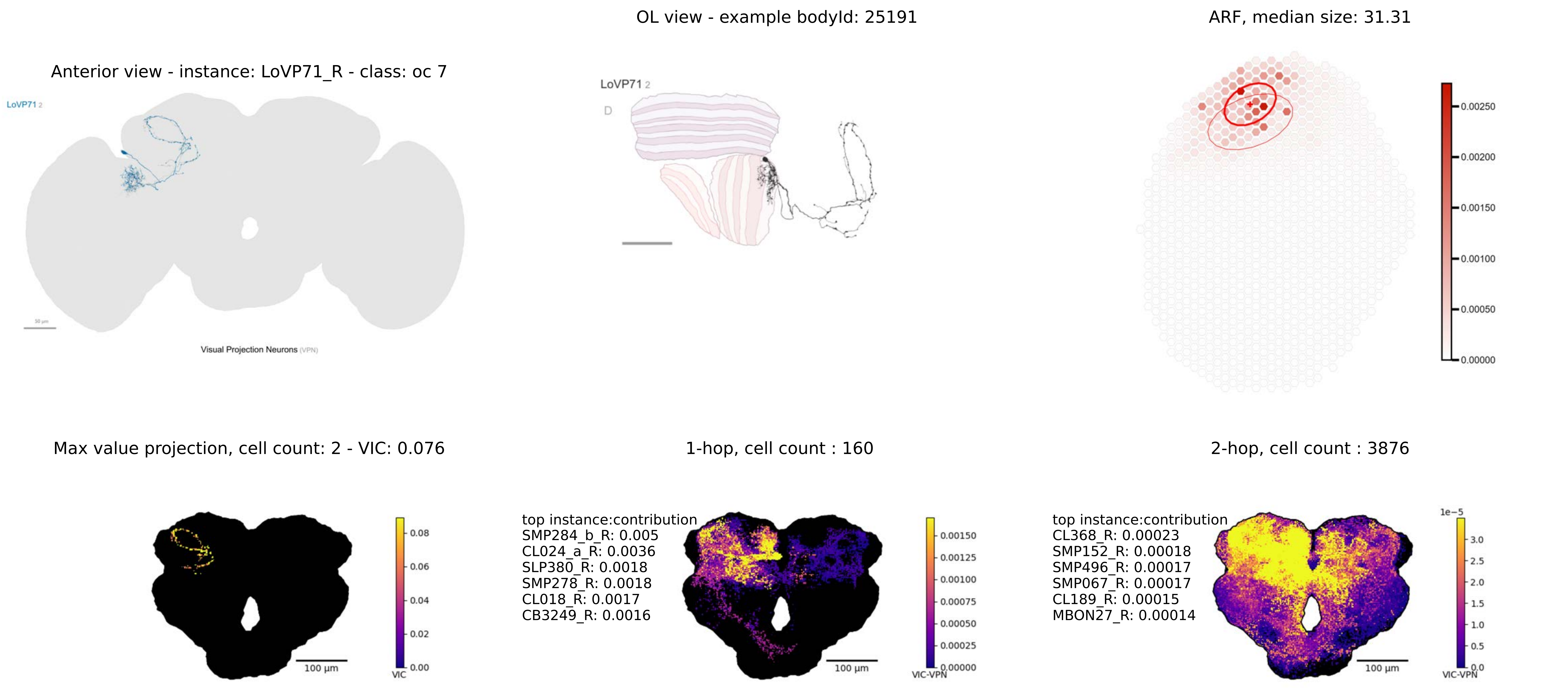

OL view - example bodyId: 25191
ARF, median size: 31.31
Anterior view - instance: LoVP71_R - class: oc 7
LoVP71 2
D
Visual Projection Neurons (VPN)
Max value projection, cell count: 2 - VIC: 0.076
1-hop, cell count : 160
2-hop, cell count : 3876
top instance:contribution
SMP284_b_R: 0.005
CL024_a_R: 0.0036
SLP380_R: 0.0018
SMP278_R: 0.0018
CL018_R: 0.0017
CB3249_R: 0.0016
top instance:contribution
CL368_R: 0.00023
SMP152_R: 0.00018
SMP496_R: 0.00017
SMP067_R: 0.00017
CL189_R: 0.00015
MBON27_R: 0.00014
100 µm
VIC
VIC-VPN
1e−5

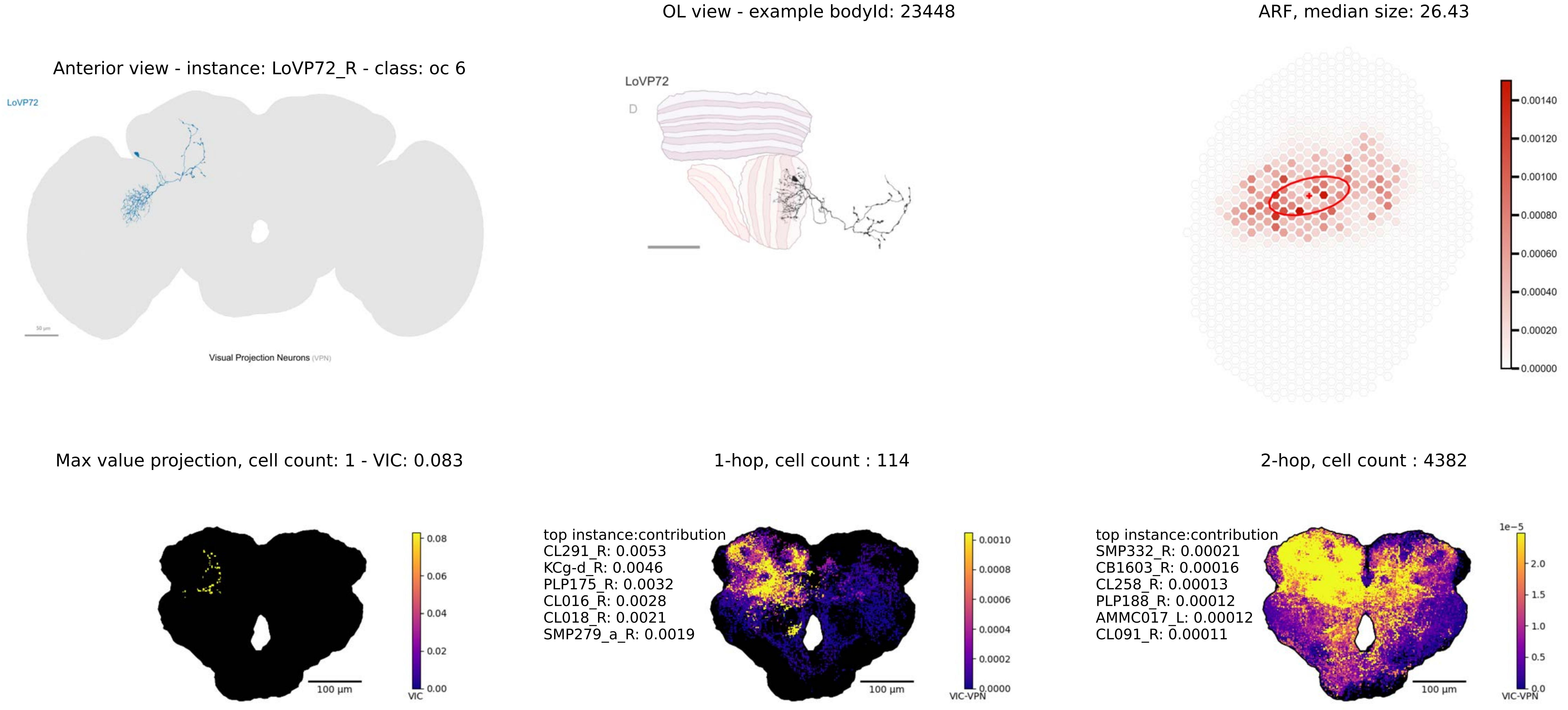

Anterior view - instance: LoVP72_R - class: oc 6
LoVP72
50 µm
Visual Projection Neurons (VPN)
OL view - example bodyId: 23448
LoVP72
D
ARF, median size: 26.43
0.00140
0.00120
0.00100
0.00080
0.00060
0.00040
0.00020
0.00000
Max value projection, cell count: 1 - VIC: 0.083
0.08
0.06
0.04
0.02
0.00
VIC
100 µm
1-hop, cell count : 114
top instance:contribution
CL291_R: 0.0053
KCg-d_R: 0.0046
PLP175_R: 0.0032
CL016_R: 0.0028
CL018_R: 0.0021
SMP279_a_R: 0.0019
0.0010
0.0008
0.0006
0.0004
0.0002
0.0000
VIC-VPN
100 µm
2-hop, cell count : 4382
top instance:contribution
SMP332_R: 0.00021
CB1603_R: 0.00016
CL258_R: 0.00013
PLP188_R: 0.00012
AMMC017_L: 0.00012
CL091_R: 0.00011
1e−5
2.0
1.5
1.0
0.5
0.0
VIC-VPN
100 µm

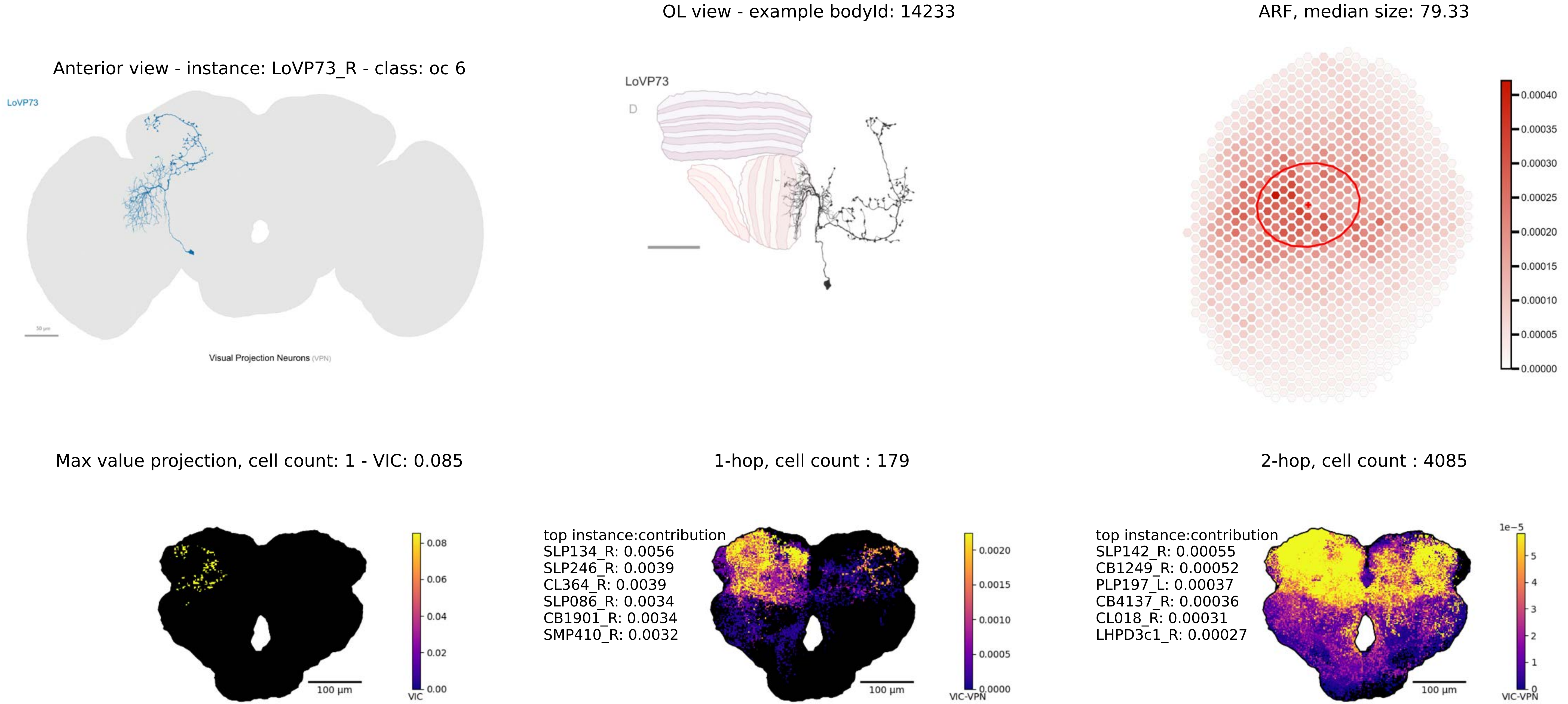
OL view - example bodyId: 14233
ARF, median size: 79.33
Anterior view - instance: LoVP73_R - class: oc 6
LoVP73
Visual Projection Neurons (VPN)
LoVP73
D
0.00040
0.00035
0.00030
0.00025
0.00020
0.00015
0.00010
0.00005
0.00000
Max value projection, cell count: 1 - VIC: 0.085
1-hop, cell count : 179
2-hop, cell count : 4085
0.08
0.06
0.04
0.02
0.00
VIC
100 µm
top instance:contribution
SLP134_R: 0.0056
SLP246_R: 0.0039
CL364_R: 0.0039
SLP086_R: 0.0034
CB1901_R: 0.0034
SMP410_R: 0.0032
0.0020
0.0015
0.0010
0.0005
0.0000
VIC-VPN
100 µm
top instance:contribution
SLP142_R: 0.00055
CB1249_R: 0.00052
PLP197_L: 0.00037
CB4137_R: 0.00036
CL018_R: 0.00031
LHPD3c1_R: 0.00027
1e−5
5
4
3
2
1
0
VIC-VPN
100 µm

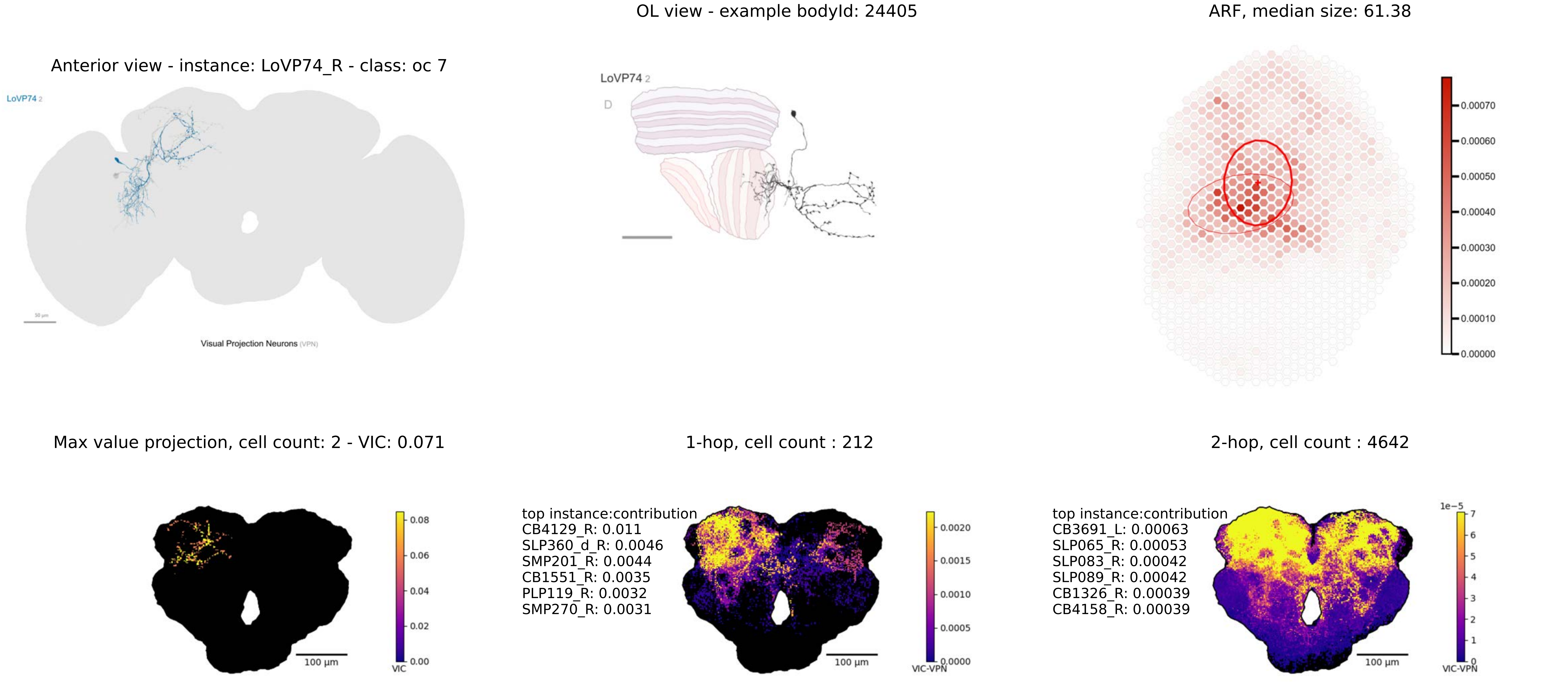

Anterior view - instance: LoVP74_R - class: oc 7
LoVP74 2
Visual Projection Neurons (VPN)
OL view - example bodyId: 24405
LoVP74 2
D
ARF, median size: 61.38
0.00070
0.00060
0.00050
0.00040
0.00030
0.00020
0.00010
0.00000
Max value projection, cell count: 2 - VIC: 0.071
100 µm
VIC
0.08
0.06
0.04
0.02
0.00
1-hop, cell count : 212
top instance:contribution
CB4129_R: 0.011
SLP360_d_R: 0.0046
SMP201_R: 0.0044
CB1551_R: 0.0035
PLP119_R: 0.0032
SMP270_R: 0.0031
100 µm
VIC-VPN
0.0020
0.0015
0.0010
0.0005
0.0000
2-hop, cell count : 4642
top instance:contribution
CB3691_L: 0.00063
SLP065_R: 0.00053
SLP083_R: 0.00042
SLP089_R: 0.00042
CB1326_R: 0.00039
CB4158_R: 0.00039
100 µm
1e-5
7
6
5
4
3
2
1
0
VIC-VPN

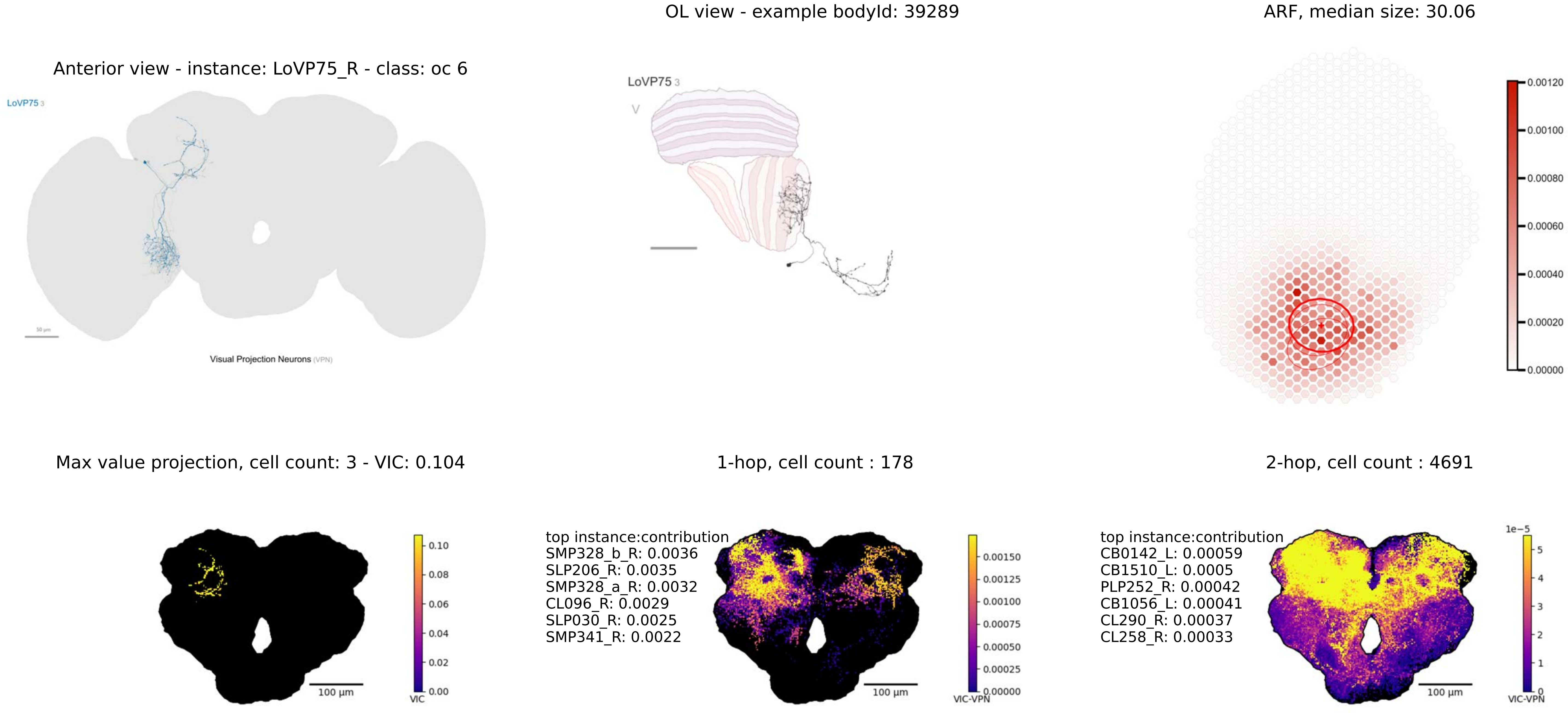
OL view - example bodyId: 39289
ARF, median size: 30.06
Anterior view - instance: LoVP75_R - class: oc 6
LoVP75 3
Visual Projection Neurons (VPN)
LoVP75 3
V
0.00120
0.00100
0.00080
0.00060
0.00040
0.00020
0.00000
Max value projection, cell count: 3 - VIC: 0.104
1-hop, cell count : 178
2-hop, cell count : 4691
0.10
0.08
0.06
0.04
0.02
0.00
VIC
100 µm
top instance:contribution
SMP328_b_R: 0.0036
SLP206_R: 0.0035
SMP328_a_R: 0.0032
CL096_R: 0.0029
SLP030_R: 0.0025
SMP341_R: 0.0022
0.00150
0.00125
0.00100
0.00075
0.00050
0.00025
0.00000
VIC-VPN
100 µm
top instance:contribution
CB0142_L: 0.00059
CB1510_L: 0.0005
PLP252_R: 0.00042
CB1056_L: 0.00041
CL290_R: 0.00037
CL258_R: 0.00033
1e−5
5
4
3
2
1
0
VIC-VPN
100 µm

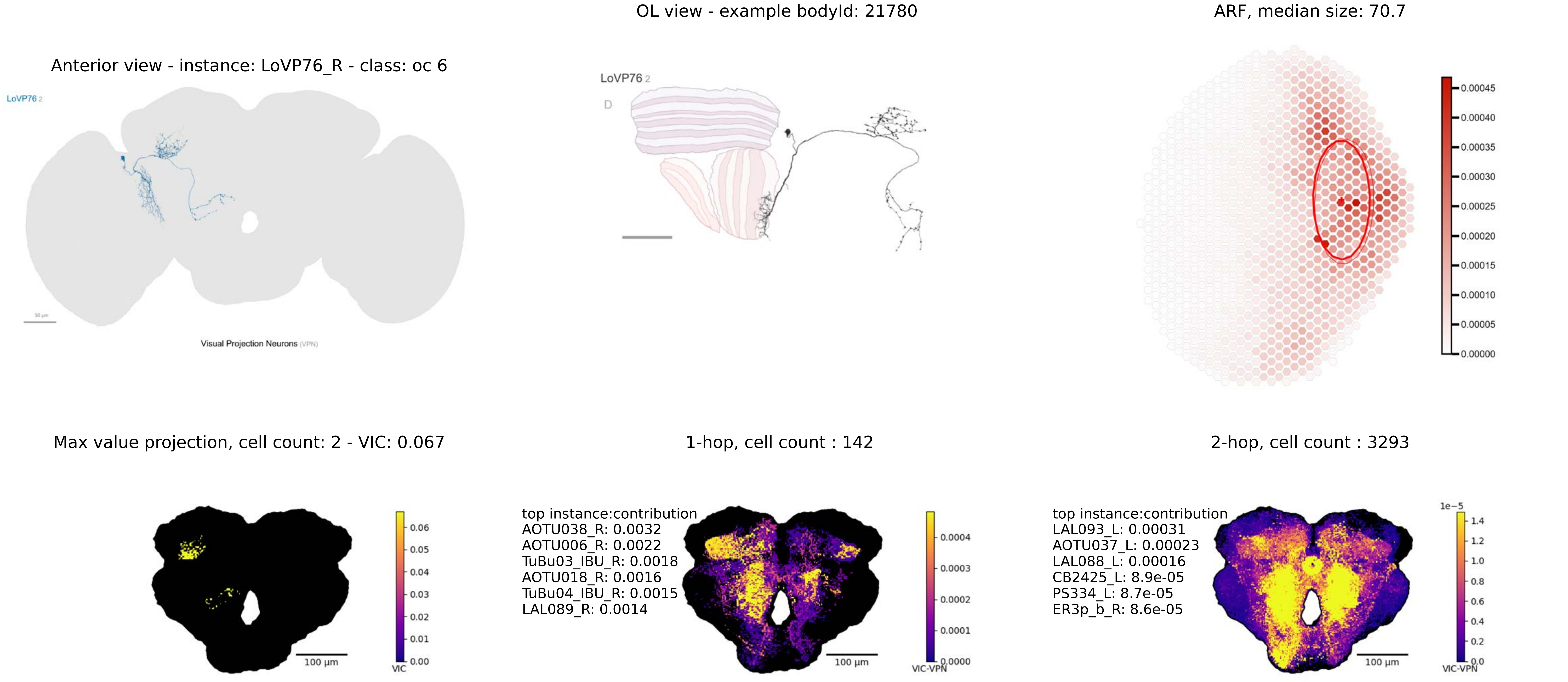

OL view - example bodyId: 21780
ARF, median size: 70.7
Anterior view - instance: LoVP76_R - class: oc 6
LoVP76 2
Visual Projection Neurons (VPN)
LoVP76 2
D
0.00045
0.00040
0.00035
0.00030
0.00025
0.00020
0.00015
0.00010
0.00005
0.00000
Max value projection, cell count: 2 - VIC: 0.067
1-hop, cell count : 142
2-hop, cell count : 3293
top instance:contribution
AOTU038_R: 0.0032
AOTU006_R: 0.0022
TuBu03_IBU_R: 0.0018
AOTU018_R: 0.0016
TuBu04_IBU_R: 0.0015
LAL089_R: 0.0014
top instance:contribution
LAL093_L: 0.00031
AOTU037_L: 0.00023
LAL088_L: 0.00016
CB2425_L: 8.9e-05
PS334_L: 8.7e-05
ER3p_b_R: 8.6e-05
100 µm
100 µm
100 µm
VIC
VIC-VPN
VIC-VPN
1e−5

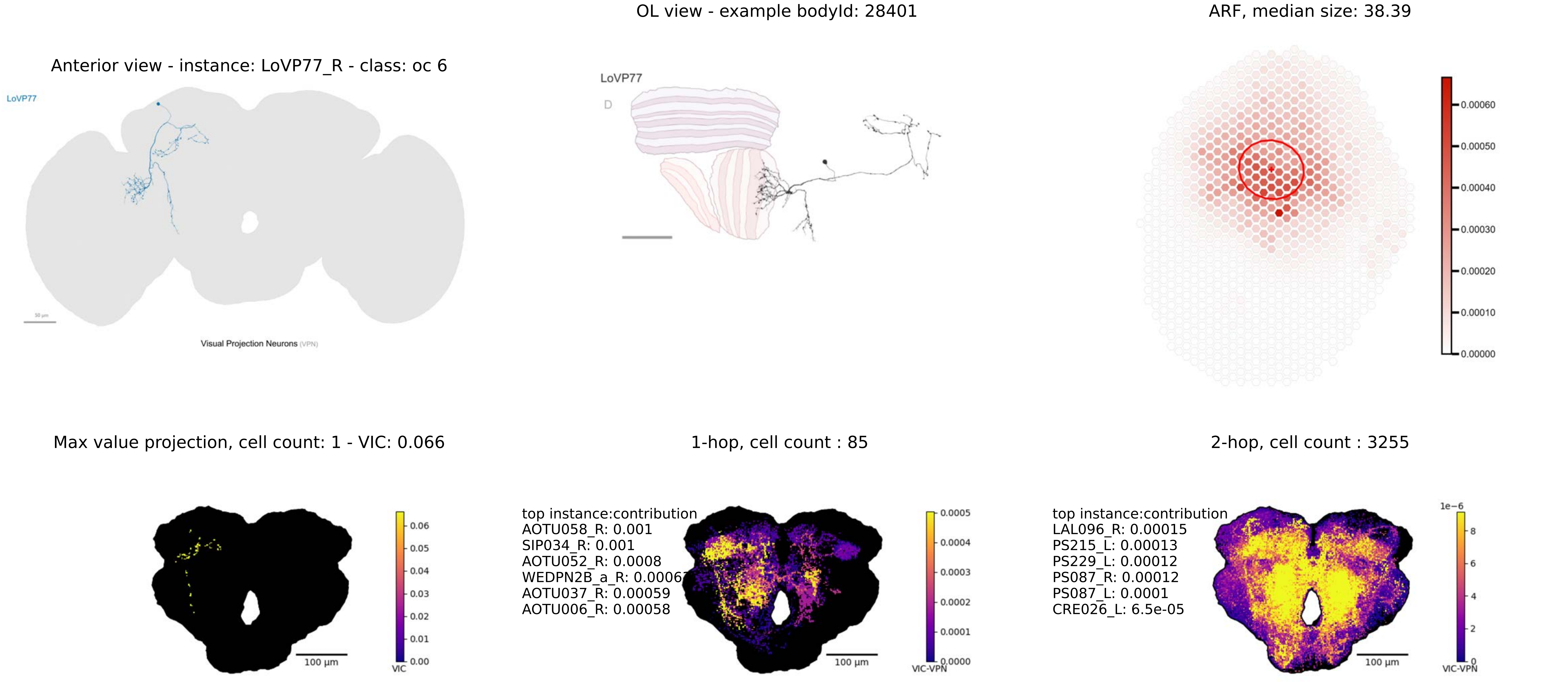

Anterior view - instance: LoVP77_R - class: oc 6
LoVP77
Visual Projection Neurons (VPN)
OL view - example bodyId: 28401
LoVP77
D
ARF, median size: 38.39
0.00060
0.00050
0.00040
0.00030
0.00020
0.00010
0.00000
Max value projection, cell count: 1 - VIC: 0.066
0.06
0.05
0.04
0.03
0.02
0.01
0.00
VIC
100 µm
1-hop, cell count : 85
top instance:contribution
AOTU058_R: 0.001
SIP034_R: 0.001
AOTU052_R: 0.0008
WEDPN2B_a_R: 0.0006
AOTU037_R: 0.00059
AOTU006_R: 0.00058
0.0005
0.0004
0.0003
0.0002
0.0001
0.0000
VIC-VPN
100 µm
2-hop, cell count : 3255
top instance:contribution
LAL096_R: 0.00015
PS215_L: 0.00013
PS229_L: 0.00012
PS087_R: 0.00012
PS087_L: 0.0001
CRE026_L: 6.5e-05
1e−6
8
6
4
2
0
VIC-VPN
100 µm

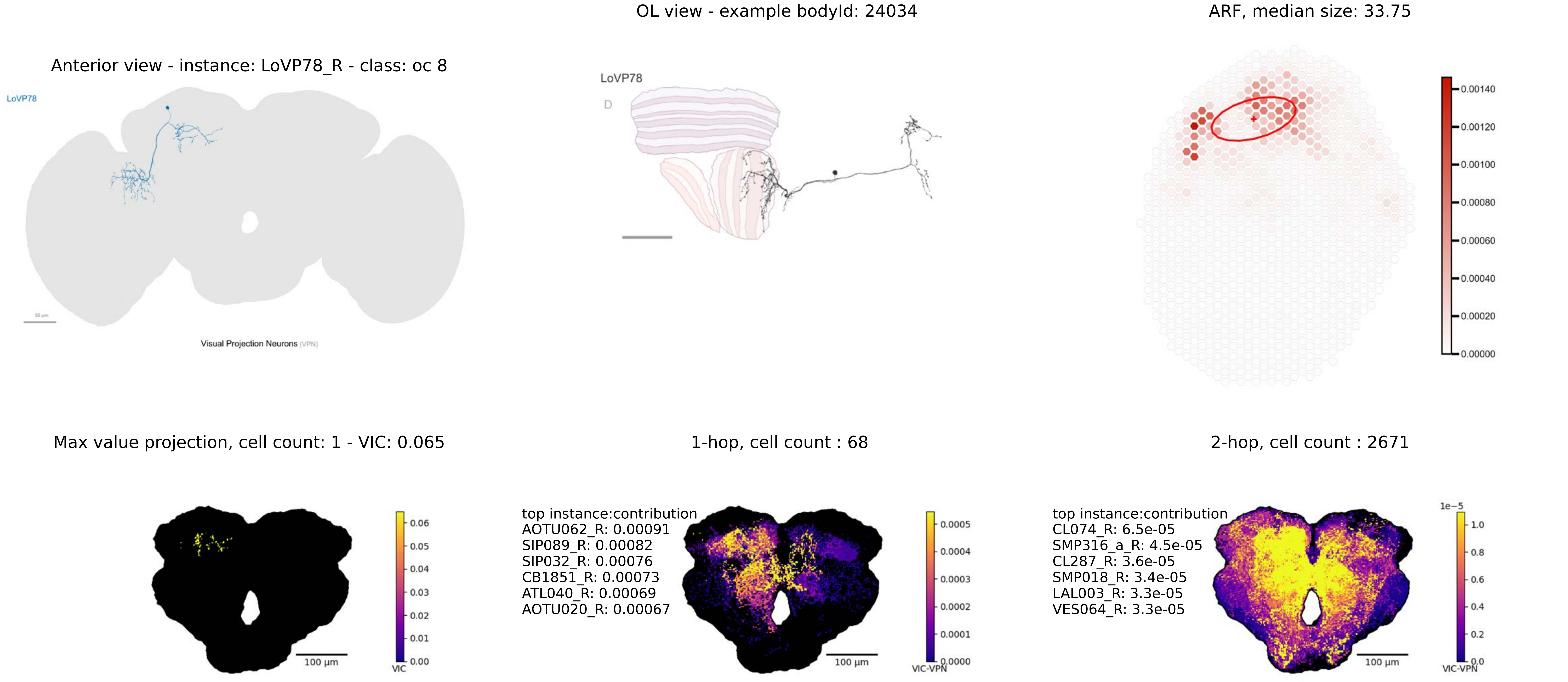

OL view - example bodyId: 24034
ARF, median size: 33.75
Anterior view - instance: LoVP78_R - class: oc 8
LoVP78
LoVP78
D
Visual Projection Neurons (VPN)
0.00140
0.00120
0.00100
0.00080
0.00060
0.00040
0.00020
0.00000
Max value projection, cell count: 1 - VIC: 0.065
1-hop, cell count : 68
2-hop, cell count : 2671
0.06
0.05
0.04
0.03
0.02
0.01
0.00
VIC
100 µm
top instance:contribution
AOTU062_R: 0.00091
SIP089_R: 0.00082
SIP032_R: 0.00076
CB1851_R: 0.00073
ATL040_R: 0.00069
AOTU020_R: 0.00067
0.0005
0.0004
0.0003
0.0002
0.0001
0.0000
VIC-VPN
100 µm
top instance:contribution
CL074_R: 6.5e-05
SMP316_a_R: 4.5e-05
CL287_R: 3.6e-05
SMP018_R: 3.4e-05
LAL003_R: 3.3e-05
VES064_R: 3.3e-05
1e−5
1.0
0.8
0.6
0.4
0.2
0.0
VIC-VPN
100 µm

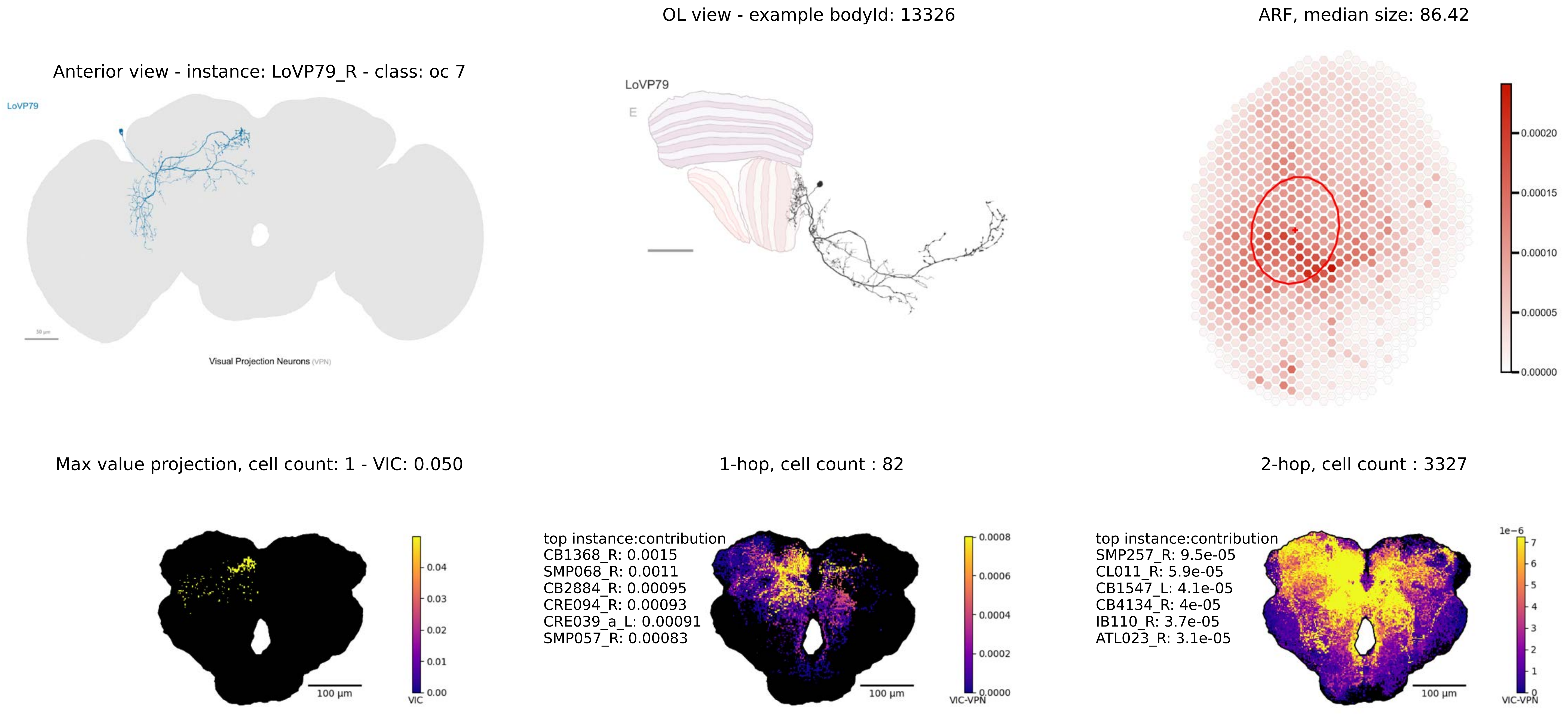

OL view - example bodyId: 13326
ARF, median size: 86.42
Anterior view - instance: LoVP79_R - class: oc 7
LoVP79
Visual Projection Neurons (VPN)
LoVP79
E
0.00020
0.00015
0.00010
0.00005
0.00000
Max value projection, cell count: 1 - VIC: 0.050
1-hop, cell count : 82
2-hop, cell count : 3327
0.04
0.03
0.02
0.01
0.00
VIC
100 µm
top instance:contribution
CB1368_R: 0.0015
SMP068_R: 0.0011
CB2884_R: 0.00095
CRE094_R: 0.00093
CRE039_a_L: 0.00091
SMP057_R: 0.00083
0.0008
0.0006
0.0004
0.0002
0.0000
VIC-VPN
100 µm
top instance:contribution
SMP257_R: 9.5e-05
CL011_R: 5.9e-05
CB1547_L: 4.1e-05
CB4134_R: 4e-05
IB110_R: 3.7e-05
ATL023_R: 3.1e-05
1e−6
7
6
5
4
3
2
1
0
VIC-VPN
100 µm

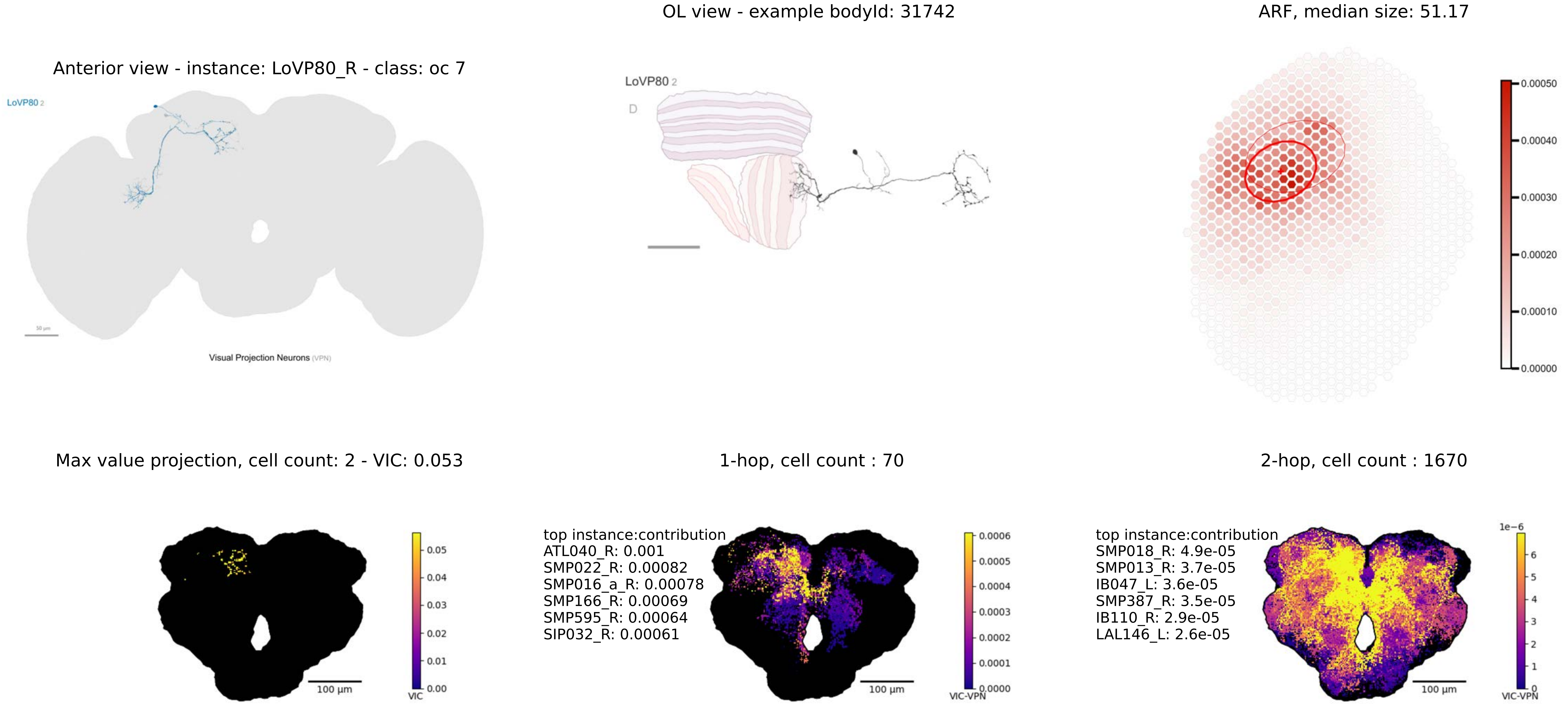

OL view - example bodyId: 31742
ARF, median size: 51.17
Anterior view - instance: LoVP80_R - class: oc 7
LoVP80 2
LoVP80 2
D
0.00050
0.00040
0.00030
0.00020
0.00010
0.00000
Visual Projection Neurons (VPN)
Max value projection, cell count: 2 - VIC: 0.053
1-hop, cell count : 70
2-hop, cell count : 1670
0.05
0.04
0.03
0.02
0.01
0.00
VIC
100 µm
top instance:contribution
ATL040_R: 0.001
SMP022_R: 0.00082
SMP016_a_R: 0.00078
SMP166_R: 0.00069
SMP595_R: 0.00064
SIP032_R: 0.00061
0.0006
0.0005
0.0004
0.0003
0.0002
0.0001
0.0000
VIC-VPN
100 µm
top instance:contribution
SMP018_R: 4.9e-05
SMP013_R: 3.7e-05
IB047_L: 3.6e-05
SMP387_R: 3.5e-05
IB110_R: 2.9e-05
LAL146_L: 2.6e-05
1e−6
6
5
4
3
2
1
0
VIC-VPN
100 µm

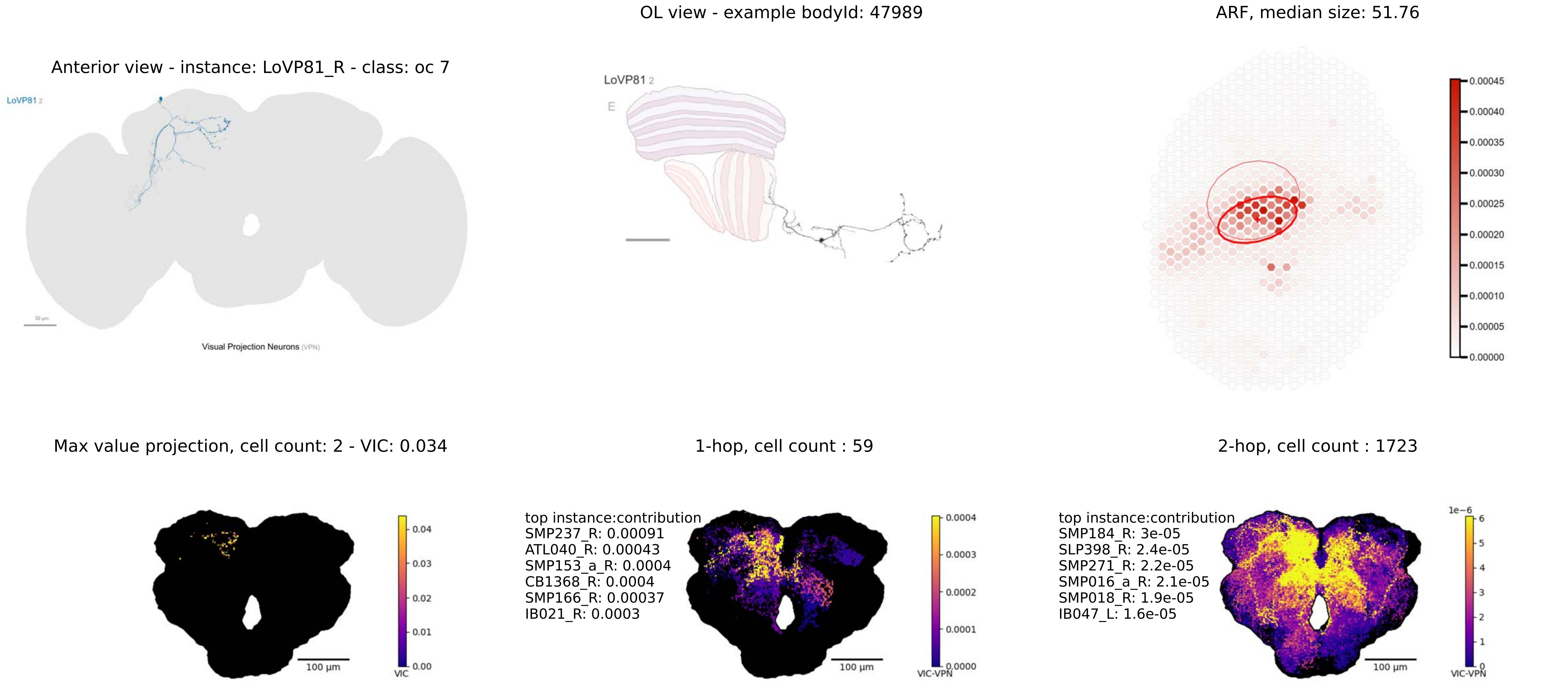
OL view - example bodyId: 47989
ARF, median size: 51.76
Anterior view - instance: LoVP81_R - class: oc 7
LoVP81 2
Visual Projection Neurons (VPN)
LoVP81 2
E
0.00045
0.00040
0.00035
0.00030
0.00025
0.00020
0.00015
0.00010
0.00005
0.00000
Max value projection, cell count: 2 - VIC: 0.034
1-hop, cell count : 59
2-hop, cell count : 1723
0.04
0.03
0.02
0.01
0.00
VIC
100 µm
top instance:contribution
SMP237_R: 0.00091
ATL040_R: 0.00043
SMP153_a_R: 0.0004
CB1368_R: 0.0004
SMP166_R: 0.00037
IB021_R: 0.0003
0.0004
0.0003
0.0002
0.0001
0.0000
VIC-VPN
100 µm
top instance:contribution
SMP184_R: 3e-05
SLP398_R: 2.4e-05
SMP271_R: 2.2e-05
SMP016_a_R: 2.1e-05
SMP018_R: 1.9e-05
IB047_L: 1.6e-05
1e−6
6
5
4
3
2
1
0
VIC-VPN
100 µm

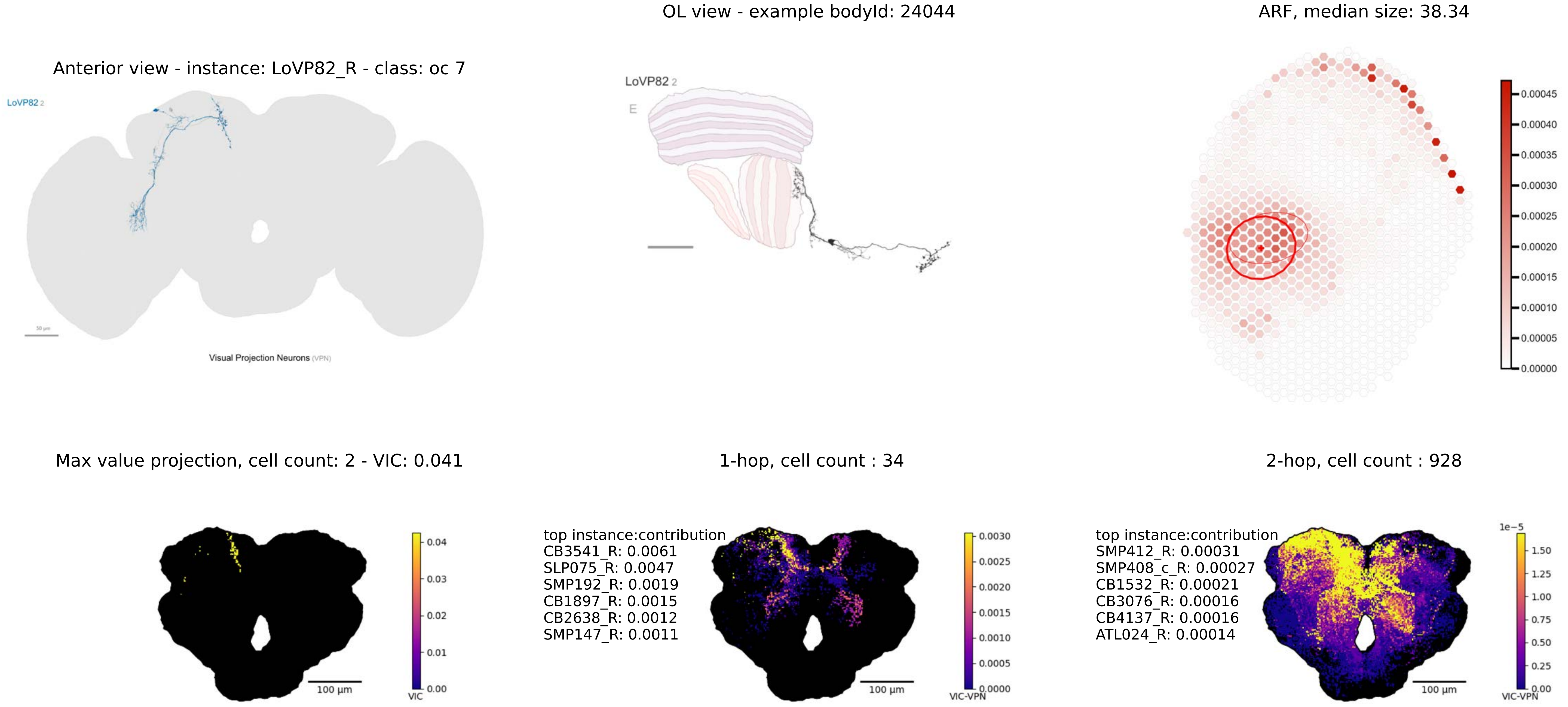
OL view - example bodyId: 24044
ARF, median size: 38.34
Anterior view - instance: LoVP82_R - class: oc 7
LoVP82 2
Visual Projection Neurons (VPN)
LoVP82 2
E
Max value projection, cell count: 2 - VIC: 0.041
1-hop, cell count : 34
2-hop, cell count : 928
top instance:contribution
CB3541_R: 0.0061
SLP075_R: 0.0047
SMP192_R: 0.0019
CB1897_R: 0.0015
CB2638_R: 0.0012
SMP147_R: 0.0011
top instance:contribution
SMP412_R: 0.00031
SMP408_c_R: 0.00027
CB1532_R: 0.00021
CB3076_R: 0.00016
CB4137_R: 0.00016
ATL024_R: 0.00014
100 µm
VIC
VIC-VPN
1e−5

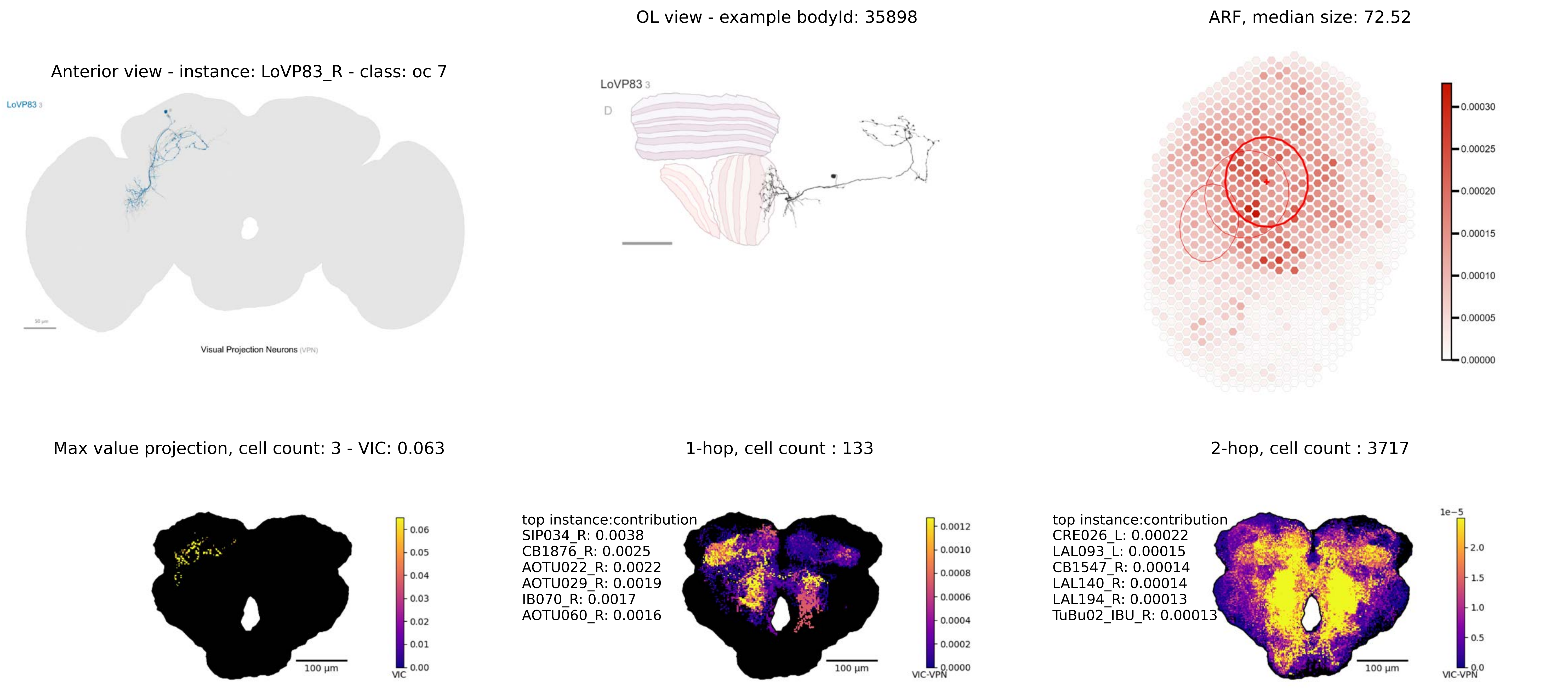
OL view - example bodyId: 35898
ARF, median size: 72.52
Anterior view - instance: LoVP83_R - class: oc 7
LoVP83 3
Visual Projection Neurons (VPN)
D
0.00030
0.00025
0.00020
0.00015
0.00010
0.00005
0.00000
Max value projection, cell count: 3 - VIC: 0.063
1-hop, cell count : 133
2-hop, cell count : 3717
VIC
VIC-VPN
100 µm
top instance:contribution
SIP034_R: 0.0038
CB1876_R: 0.0025
AOTU022_R: 0.0022
AOTU029_R: 0.0019
IB070_R: 0.0017
AOTU060_R: 0.0016
top instance:contribution
CRE026_L: 0.00022
LAL093_L: 0.00015
CB1547_R: 0.00014
LAL140_R: 0.00014
LAL194_R: 0.00013
TuBu02_IBU_R: 0.00013
1e−5

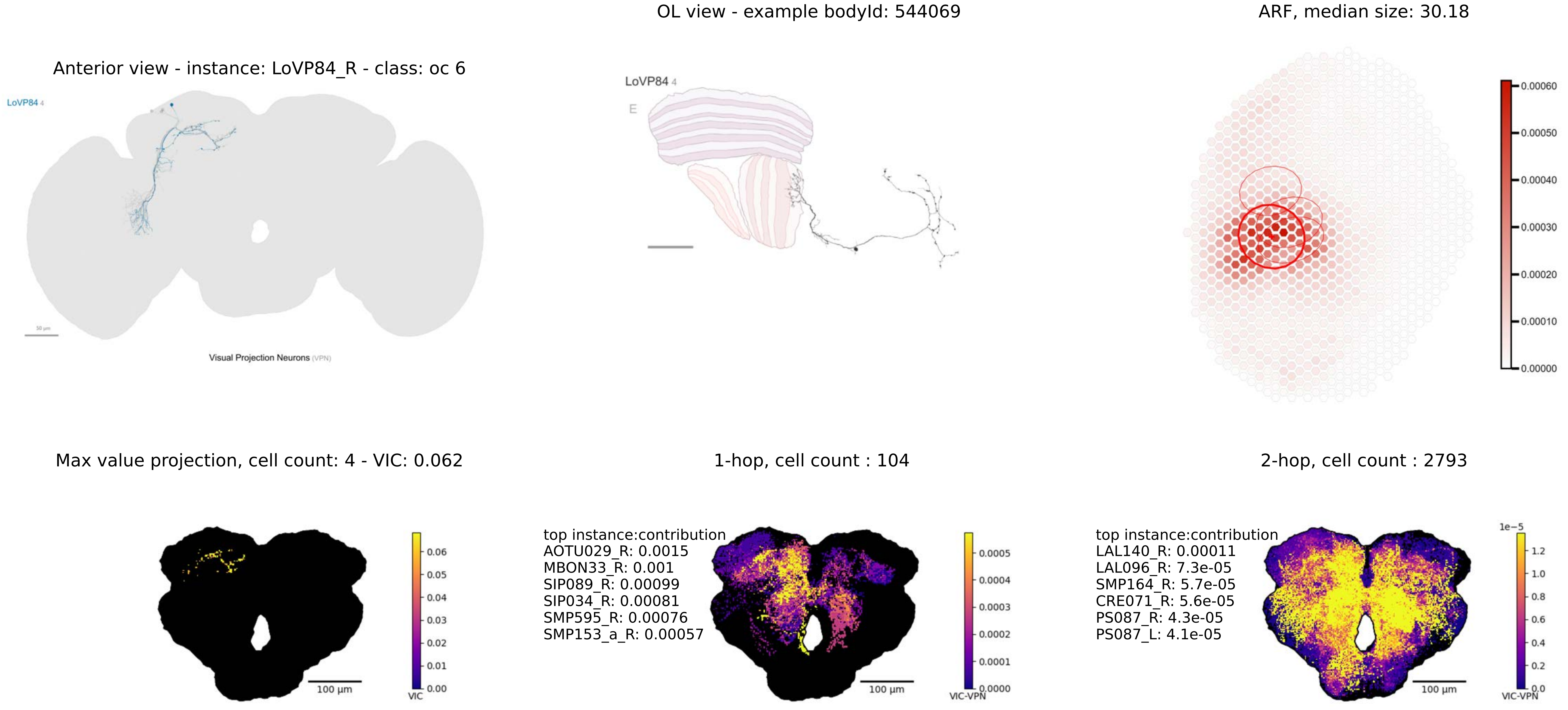

OL view - example bodyId: 544069
ARF, median size: 30.18
Anterior view - instance: LoVP84_R - class: oc 6
LoVP84 4
Visual Projection Neurons (VPN)
LoVP84 4
E
0.00060
0.00050
0.00040
0.00030
0.00020
0.00010
0.00000
Max value projection, cell count: 4 - VIC: 0.062
1-hop, cell count : 104
2-hop, cell count : 2793
100 µm
VIC
0.06
0.05
0.04
0.03
0.02
0.01
0.00
top instance:contribution
AOTU029_R: 0.0015
MBON33_R: 0.001
SIP089_R: 0.00099
SIP034_R: 0.00081
SMP595_R: 0.00076
SMP153_a_R: 0.00057
100 µm
VIC-VPN
0.0005
0.0004
0.0003
0.0002
0.0001
0.0000
top instance:contribution
LAL140_R: 0.00011
LAL096_R: 7.3e-05
SMP164_R: 5.7e-05
CRE071_R: 5.6e-05
PS087_R: 4.3e-05
PS087_L: 4.1e-05
100 µm
1e−5
1.2
1.0
0.8
0.6
0.4
0.2
0.0
VIC-VPN

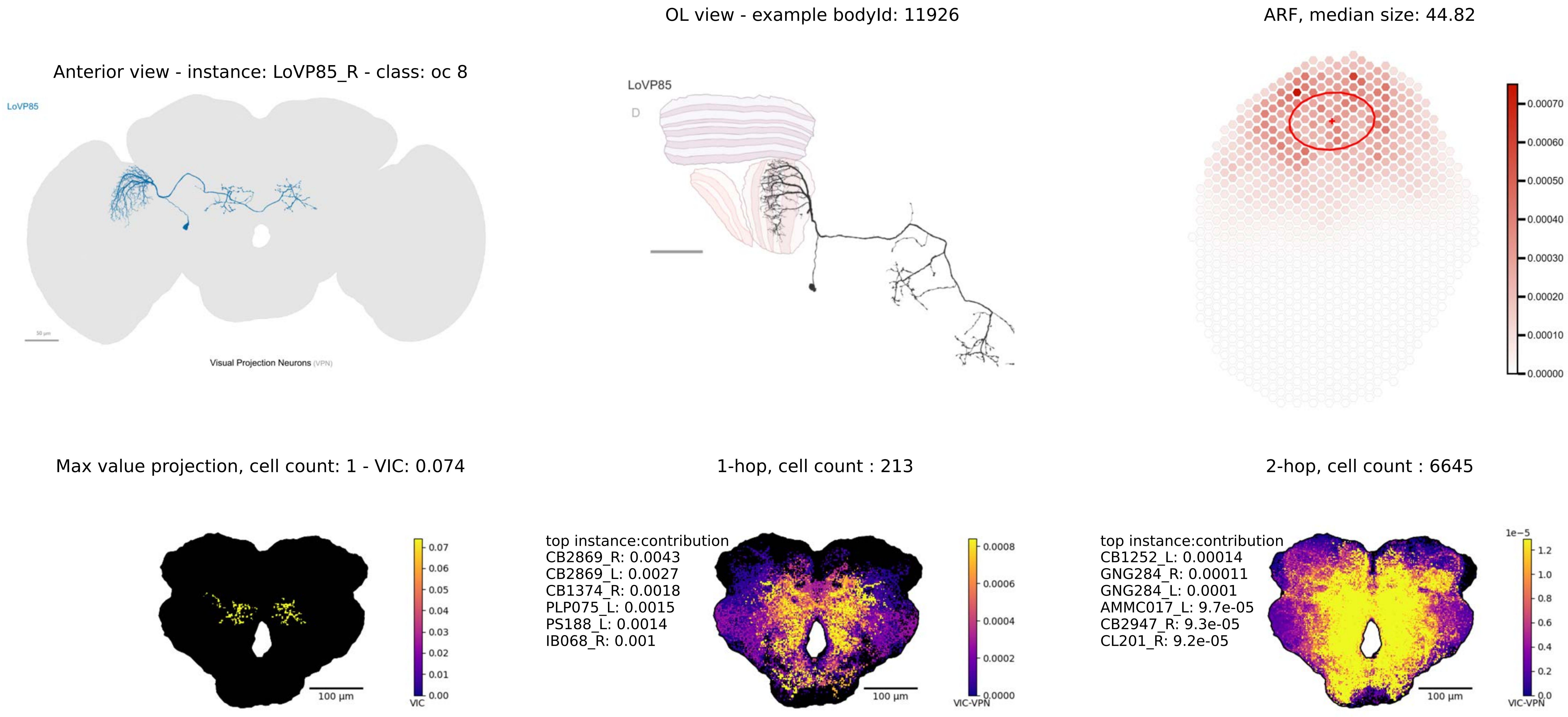

OL view - example bodyId: 11926
ARF, median size: 44.82
Anterior view - instance: LoVP85_R - class: oc 8
LoVP85
Visual Projection Neurons (VPN)
LoVP85
D
0.00070
0.00060
0.00050
0.00040
0.00030
0.00020
0.00010
0.00000
Max value projection, cell count: 1 - VIC: 0.074
1-hop, cell count : 213
2-hop, cell count : 6645
0.07
0.06
0.05
0.04
0.03
0.02
0.01
0.00
VIC
100 µm
top instance:contribution
CB2869_R: 0.0043
CB2869_L: 0.0027
CB1374_R: 0.0018
PLP075_L: 0.0015
PS188_L: 0.0014
IB068_R: 0.001
0.0008
0.0006
0.0004
0.0002
0.0000
VIC-VPN
100 µm
top instance:contribution
CB1252_L: 0.00014
GNG284_R: 0.00011
GNG284_L: 0.0001
AMMC017_L: 9.7e-05
CB2947_R: 9.3e-05
CL201_R: 9.2e-05
1e−5
1.2
1.0
0.8
0.6
0.4
0.2
0.0
VIC-VPN
100 µm

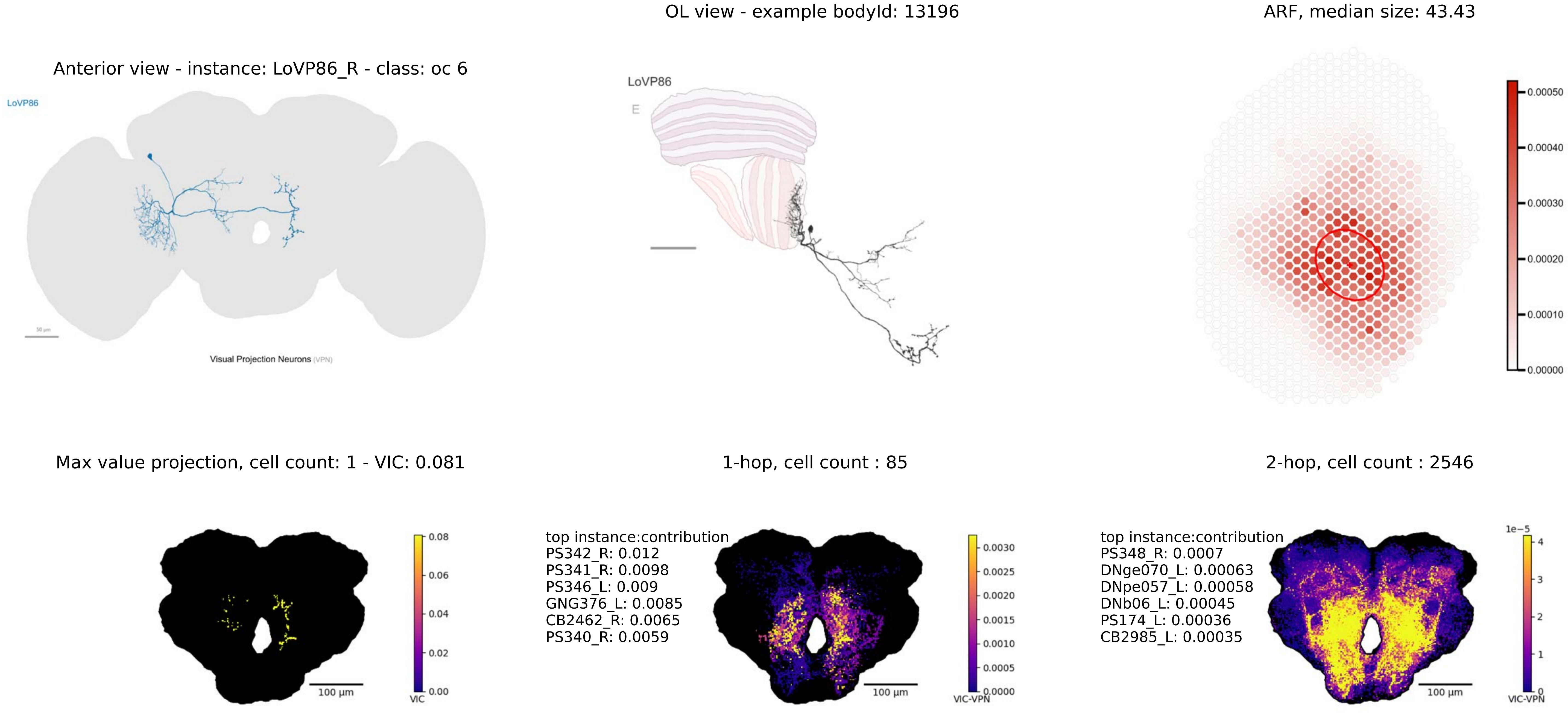

OL view - example bodyId: 13196
ARF, median size: 43.43
Anterior view - instance: LoVP86_R - class: oc 6
LoVP86
Visual Projection Neurons (VPN)
LoVP86
E
0.00050
0.00040
0.00030
0.00020
0.00010
0.00000
Max value projection, cell count: 1 - VIC: 0.081
1-hop, cell count : 85
2-hop, cell count : 2546
100 µm
VIC
0.08
0.06
0.04
0.02
0.00
top instance:contribution
PS342_R: 0.012
PS341_R: 0.0098
PS346_L: 0.009
GNG376_L: 0.0085
CB2462_R: 0.0065
PS340_R: 0.0059
VIC-VPN
0.0030
0.0025
0.0020
0.0015
0.0010
0.0005
0.0000
top instance:contribution
PS348_R: 0.0007
DNge070_L: 0.00063
DNpe057_L: 0.00058
DNb06_L: 0.00045
PS174_L: 0.00036
CB2985_L: 0.00035
1e−5
4
3
2
1
0

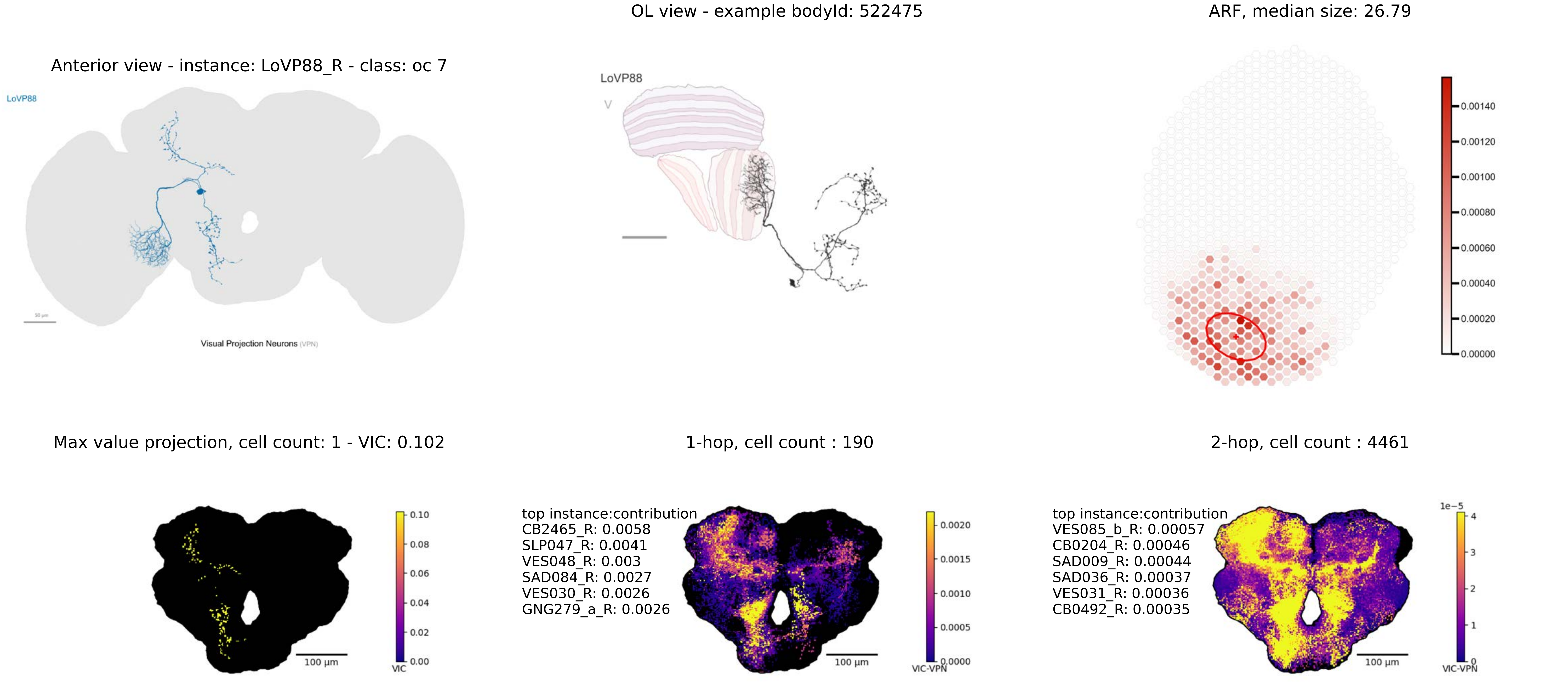
OL view - example bodyId: 522475
ARF, median size: 26.79
Anterior view - instance: LoVP88_R - class: oc 7
LoVP88
Visual Projection Neurons (VPN)
LoVP88
V
0.00140
0.00120
0.00100
0.00080
0.00060
0.00040
0.00020
0.00000
Max value projection, cell count: 1 - VIC: 0.102
1-hop, cell count : 190
2-hop, cell count : 4461
0.10
0.08
0.06
0.04
0.02
0.00
VIC
100 µm
top instance:contribution
CB2465_R: 0.0058
SLP047_R: 0.0041
VES048_R: 0.003
SAD084_R: 0.0027
VES030_R: 0.0026
GNG279_a_R: 0.0026
0.0020
0.0015
0.0010
0.0005
0.0000
VIC-VPN
100 µm
top instance:contribution
VES085_b_R: 0.00057
CB0204_R: 0.00046
SAD009_R: 0.00044
SAD036_R: 0.00037
VES031_R: 0.00036
CB0492_R: 0.00035
1e−5
4
3
2
1
0
VIC-VPN
100 µm

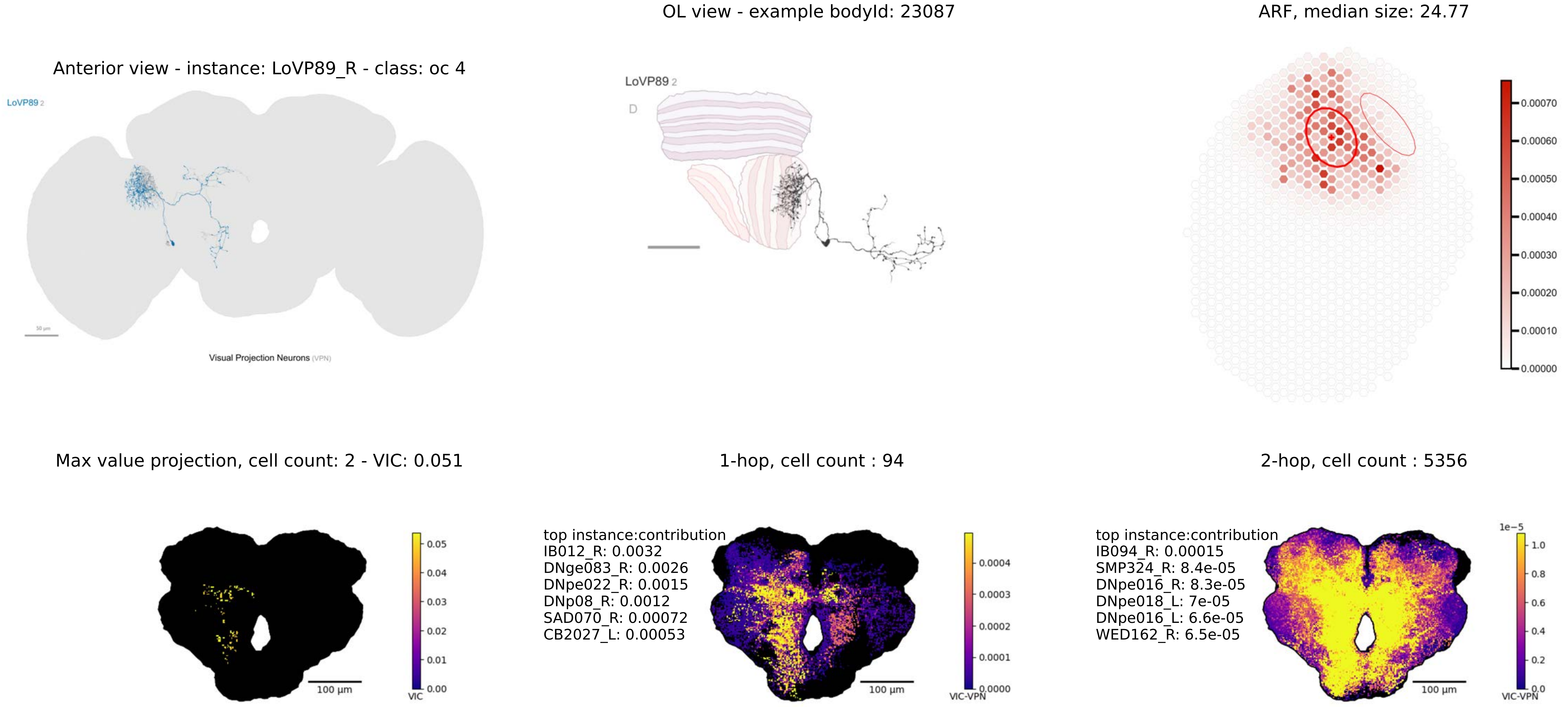

OL view - example bodyId: 23087
ARF, median size: 24.77
Anterior view - instance: LoVP89_R - class: oc 4
LoVP89 2
50 μm
Visual Projection Neurons (VPN)
LoVP89 2
D
0.00070
0.00060
0.00050
0.00040
0.00030
0.00020
0.00010
0.00000
Max value projection, cell count: 2 - VIC: 0.051
1-hop, cell count : 94
2-hop, cell count : 5356
0.05
0.04
0.03
0.02
0.01
0.00
VIC
100 μm
top instance:contribution
IB012_R: 0.0032
DNge083_R: 0.0026
DNpe022_R: 0.0015
DNp08_R: 0.0012
SAD070_R: 0.00072
CB2027_L: 0.00053
0.0004
0.0003
0.0002
0.0001
0.0000
VIC-VPN
100 μm
top instance:contribution
IB094_R: 0.00015
SMP324_R: 8.4e-05
DNpe016_R: 8.3e-05
DNpe018_L: 7e-05
DNpe016_L: 6.6e-05
WED162_R: 6.5e-05
1e−5
1.0
0.8
0.6
0.4
0.2
0.0
VIC-VPN
100 μm

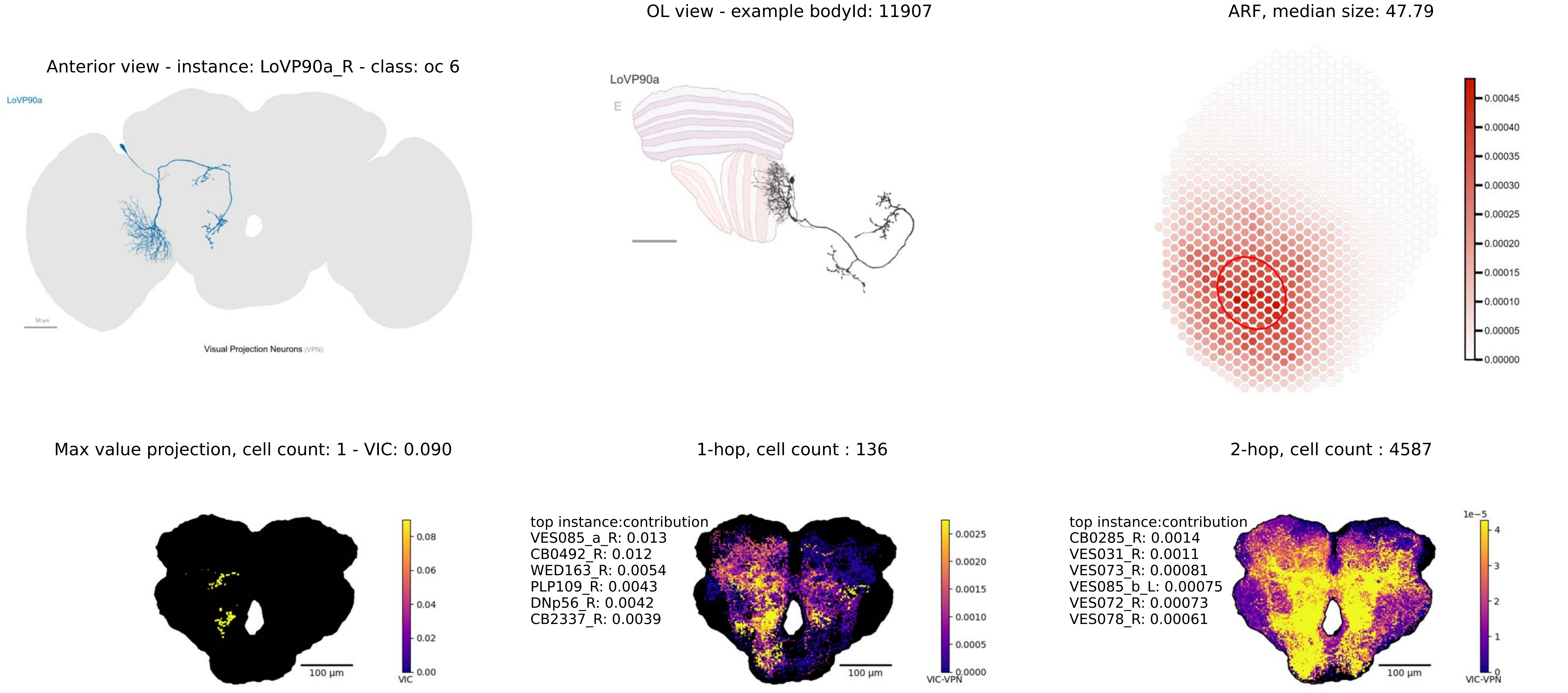

OL view - example bodyId: 11907
ARF, median size: 47.79
Anterior view - instance: LoVP90a_R - class: oc 6
LoVP90a
LoVP90a
E
Visual Projection Neurons (VPN)
0.00045
0.00040
0.00035
0.00030
0.00025
0.00020
0.00015
0.00010
0.00005
0.00000
Max value projection, cell count: 1 - VIC: 0.090
1-hop, cell count : 136
2-hop, cell count : 4587
top instance:contribution
VES085_a_R: 0.013
CB0492_R: 0.012
WED163_R: 0.0054
PLP109_R: 0.0043
DNp56_R: 0.0042
CB2337_R: 0.0039
top instance:contribution
CB0285_R: 0.0014
VES031_R: 0.0011
VES073_R: 0.00081
VES085_b_L: 0.00075
VES072_R: 0.00073
VES078_R: 0.00061
0.08
0.06
0.04
0.02
0.00
VIC
0.0025
0.0020
0.0015
0.0010
0.0005
0.0000
VIC-VPN
1e−5
4
3
2
1
0
VIC-VPN
100 µm
100 µm
100 µm

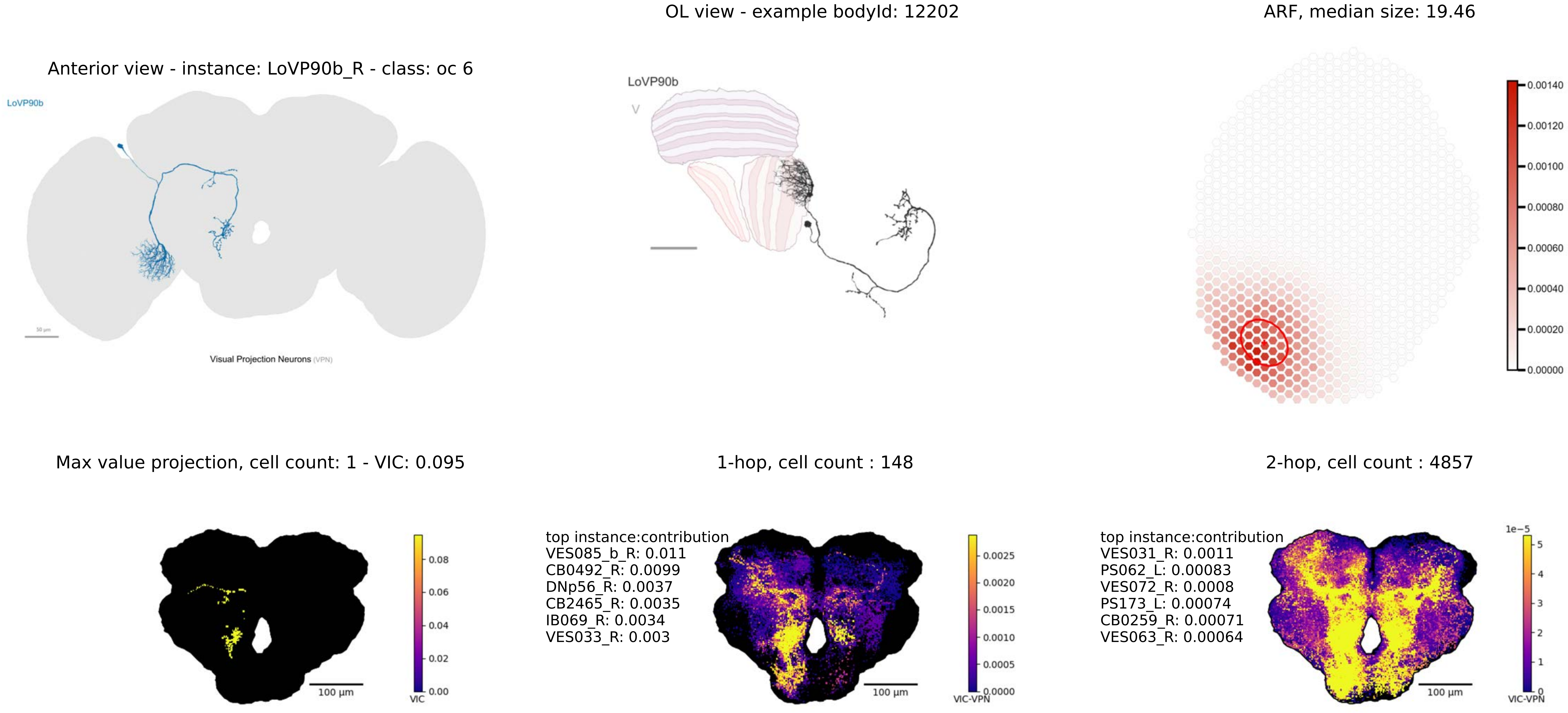

Anterior view - instance: LoVP90b_R - class: oc 6
LoVP90b
50 μm
Visual Projection Neurons (VPN)
OL view - example bodyId: 12202
LoVP90b
V
ARF, median size: 19.46
0.00140
0.00120
0.00100
0.00080
0.00060
0.00040
0.00020
0.00000
Max value projection, cell count: 1 - VIC: 0.095
100 μm
VIC
0.08
0.06
0.04
0.02
0.00
1-hop, cell count : 148
top instance:contribution
VES085_b_R: 0.011
CB0492_R: 0.0099
DNp56_R: 0.0037
CB2465_R: 0.0035
IB069_R: 0.0034
VES033_R: 0.003
100 μm
VIC-VPN
0.0025
0.0020
0.0015
0.0010
0.0005
0.0000
2-hop, cell count : 4857
top instance:contribution
VES031_R: 0.0011
PS062_L: 0.00083
VES072_R: 0.0008
PS173_L: 0.00074
CB0259_R: 0.00071
VES063_R: 0.00064
100 μm
1e−5
5
4
3
2
1
0
VIC-VPN

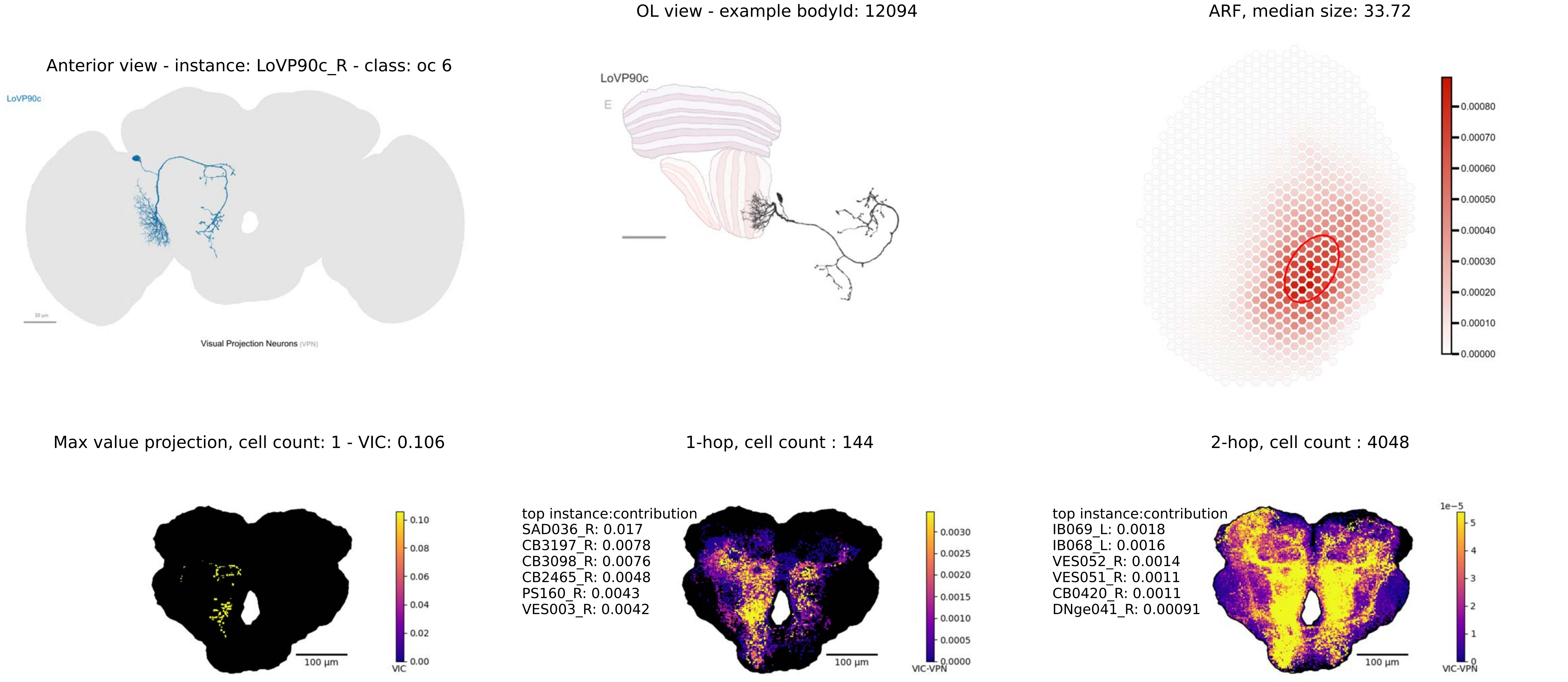
OL view - example bodyId: 12094
ARF, median size: 33.72
Anterior view - instance: LoVP90c_R - class: oc 6
LoVP90c
Visual Projection Neurons (VPN)
LoVP90c
E
0.00080
0.00070
0.00060
0.00050
0.00040
0.00030
0.00020
0.00010
0.00000
Max value projection, cell count: 1 - VIC: 0.106
1-hop, cell count : 144
2-hop, cell count : 4048
0.10
0.08
0.06
0.04
0.02
0.00
VIC
100 µm
top instance:contribution
SAD036_R: 0.017
CB3197_R: 0.0078
CB3098_R: 0.0076
CB2465_R: 0.0048
PS160_R: 0.0043
VES003_R: 0.0042
0.0030
0.0025
0.0020
0.0015
0.0010
0.0005
0.0000
VIC-VPN
100 µm
top instance:contribution
IB069_L: 0.0018
IB068_L: 0.0016
VES052_R: 0.0014
VES051_R: 0.0011
CB0420_R: 0.0011
DNge041_R: 0.00091
1e−5
5
4
3
2
1
0
VIC-VPN
100 µm

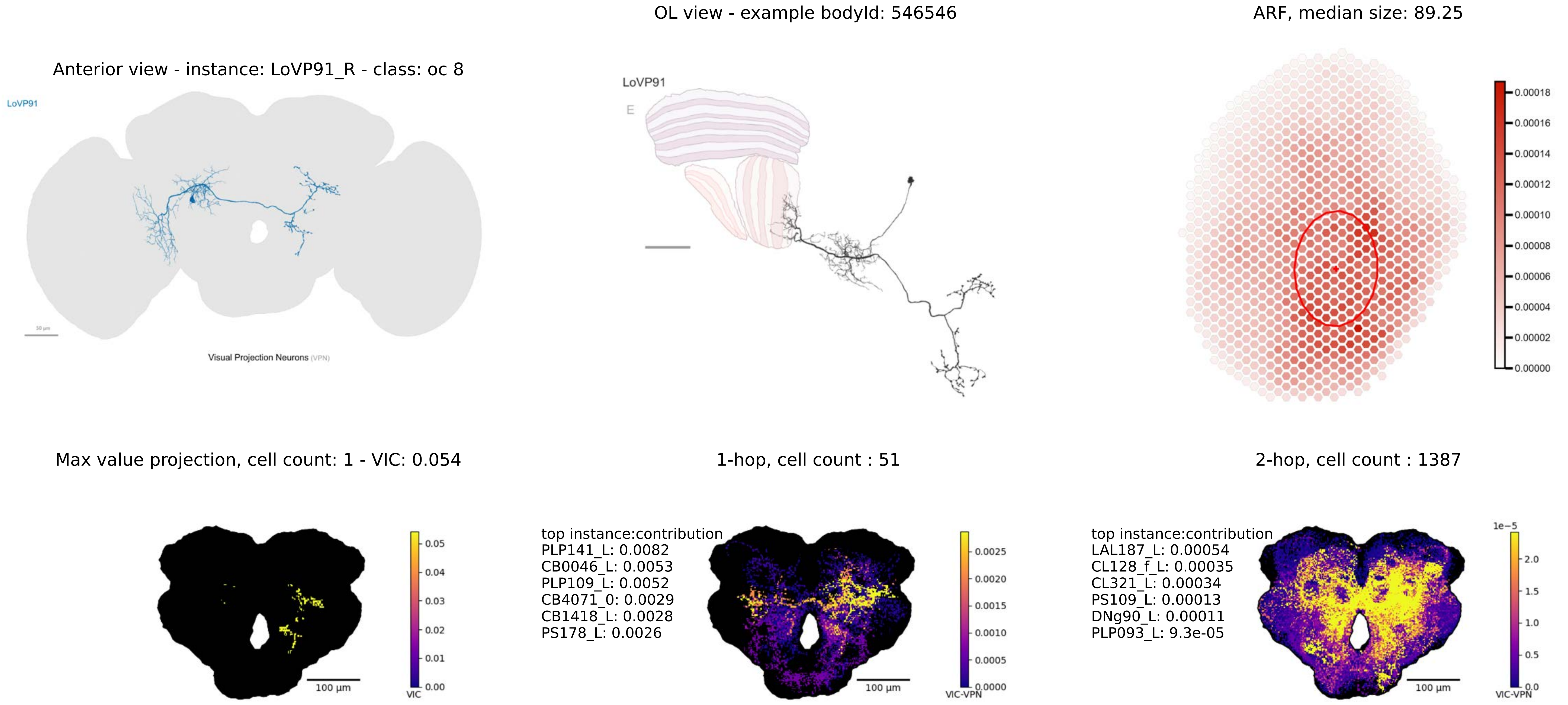
OL view - example bodyId: 546546
ARF, median size: 89.25
Anterior view - instance: LoVP91_R - class: oc 8
LoVP91
Visual Projection Neurons (VPN)
LoVP91
E
0.00018
0.00016
0.00014
0.00012
0.00010
0.00008
0.00006
0.00004
0.00002
0.00000
Max value projection, cell count: 1 - VIC: 0.054
1-hop, cell count : 51
2-hop, cell count : 1387
0.05
0.04
0.03
0.02
0.01
0.00
VIC
100 µm
top instance:contribution
PLP141_L: 0.0082
CB0046_L: 0.0053
PLP109_L: 0.0052
CB4071_0: 0.0029
CB1418_L: 0.0028
PS178_L: 0.0026
0.0025
0.0020
0.0015
0.0010
0.0005
0.0000
VIC-VPN
100 µm
top instance:contribution
LAL187_L: 0.00054
CL128_f_L: 0.00035
CL321_L: 0.00034
PS109_L: 0.00013
DNg90_L: 0.00011
PLP093_L: 9.3e-05
1e−5
2.0
1.5
1.0
0.5
0.0
VIC-VPN
100 µm

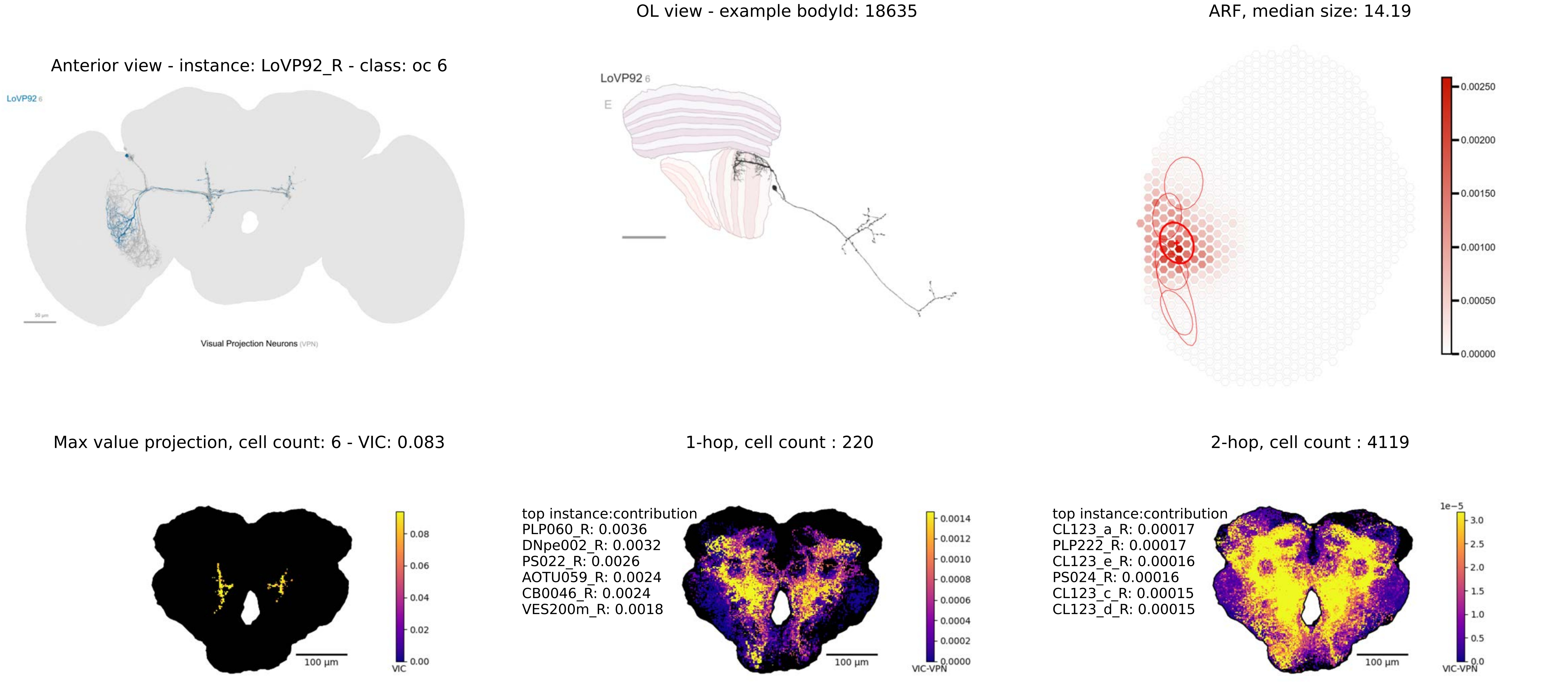

Anterior view - instance: LoVP92_R - class: oc 6
LoVP92 6
Visual Projection Neurons (VPN)
OL view - example bodyId: 18635
LoVP92 6
E
ARF, median size: 14.19
0.00250
0.00200
0.00150
0.00100
0.00050
0.00000
Max value projection, cell count: 6 - VIC: 0.083
100 µm
0.08
0.06
0.04
0.02
0.00
VIC
1-hop, cell count : 220
top instance:contribution
PLP060_R: 0.0036
DNpe002_R: 0.0032
PS022_R: 0.0026
AOTU059_R: 0.0024
CB0046_R: 0.0024
VES200m_R: 0.0018
100 µm
0.0014
0.0012
0.0010
0.0008
0.0006
0.0004
0.0002
0.0000
VIC-VPN
2-hop, cell count : 4119
top instance:contribution
CL123_a_R: 0.00017
PLP222_R: 0.00017
CL123_e_R: 0.00016
PS024_R: 0.00016
CL123_c_R: 0.00015
CL123_d_R: 0.00015
100 µm
1e−5
3.0
2.5
2.0
1.5
1.0
0.5
0.0
VIC-VPN

Anterior view - instance: LoVP93_R - class: oc 8

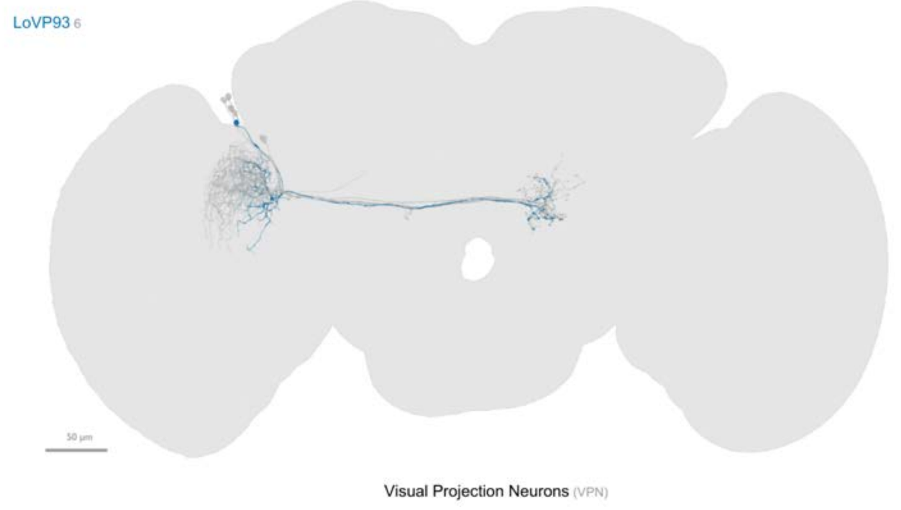


OL view - example bodyId: 41512

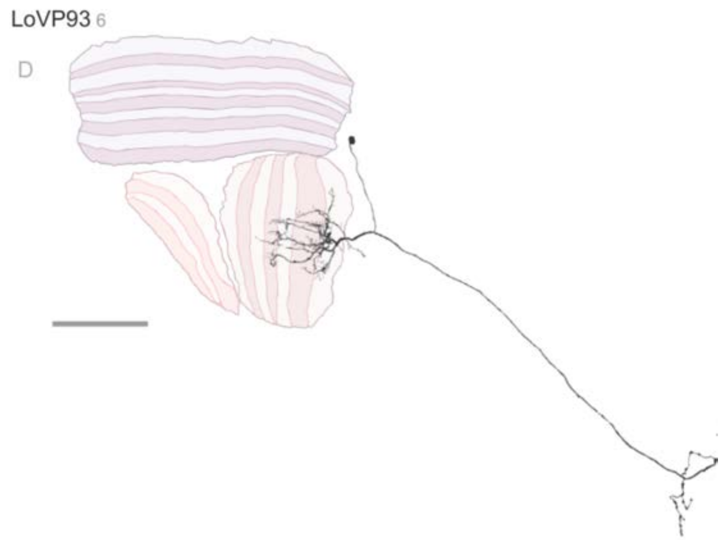


ARF, median size: 17.68

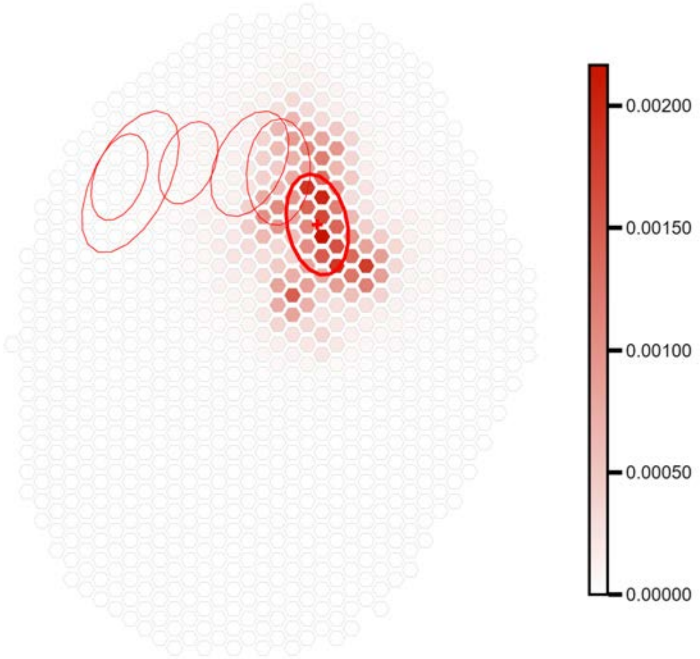


Max value projection, cell count: 6 - VIC: 0.095

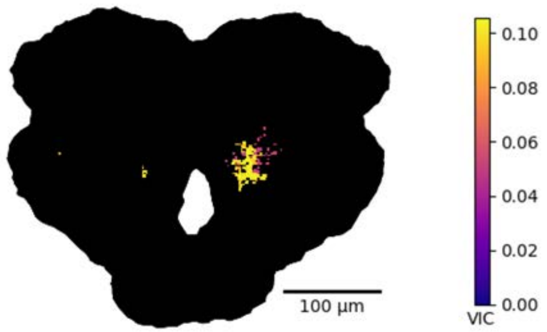


1-hop, cell count : 185

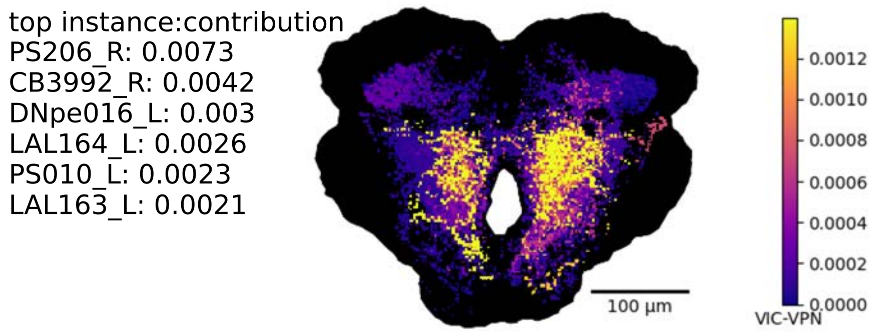


2-hop, cell count : 3014

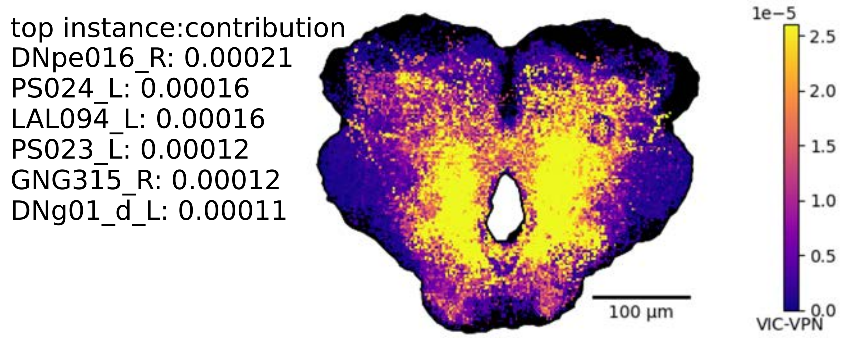

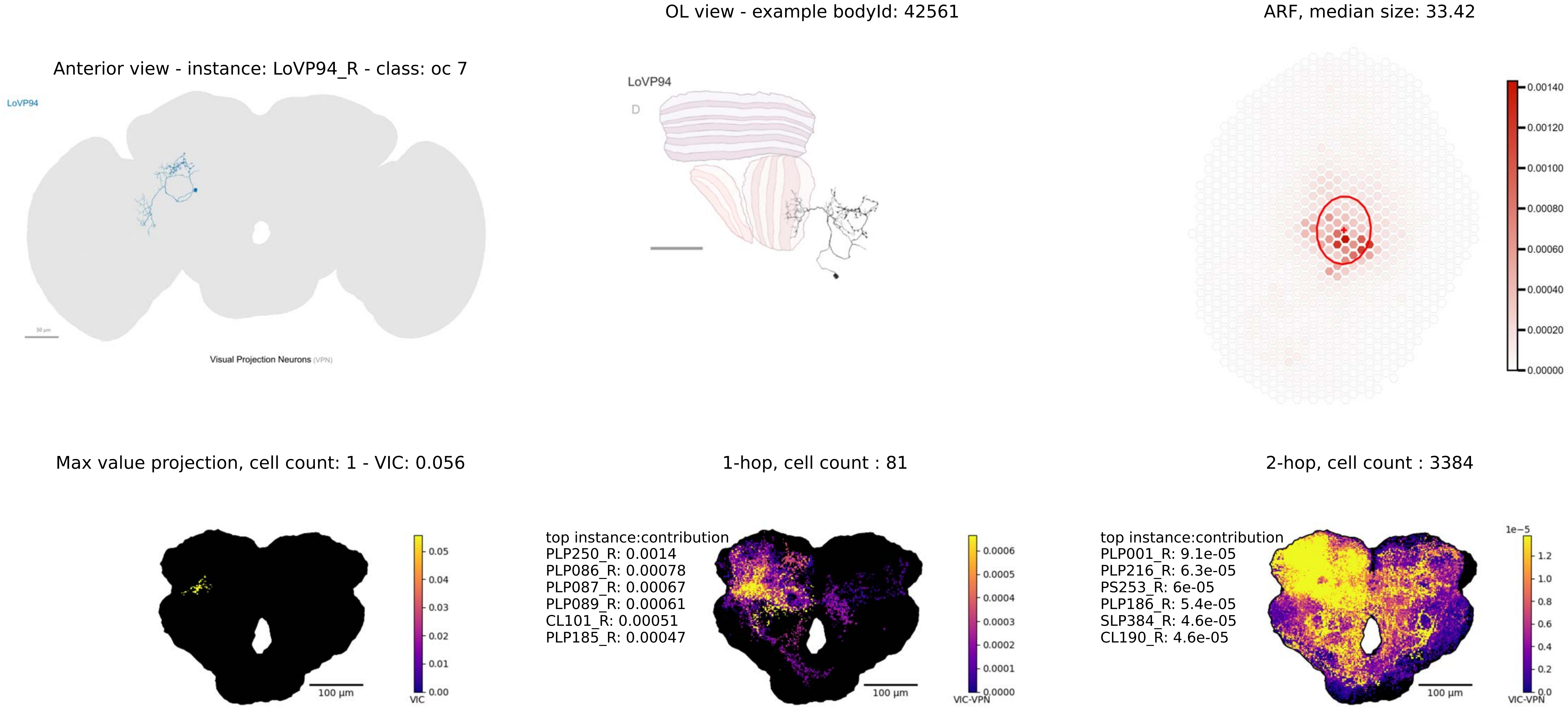

OL view - example bodyId: 42561
ARF, median size: 33.42
Anterior view - instance: LoVP94_R - class: oc 7
LoVP94
Visual Projection Neurons (VPN)
LoVP94
D
0.00140
0.00120
0.00100
0.00080
0.00060
0.00040
0.00020
0.00000
Max value projection, cell count: 1 - VIC: 0.056
1-hop, cell count : 81
2-hop, cell count : 3384
0.05
0.04
0.03
0.02
0.01
0.00
VIC
100 µm
top instance:contribution
PLP250_R: 0.0014
PLP086_R: 0.00078
PLP087_R: 0.00067
PLP089_R: 0.00061
CL101_R: 0.00051
PLP185_R: 0.00047
0.0006
0.0005
0.0004
0.0003
0.0002
0.0001
0.0000
VIC-VPN
100 µm
top instance:contribution
PLP001_R: 9.1e-05
PLP216_R: 6.3e-05
PS253_R: 6e-05
PLP186_R: 5.4e-05
SLP384_R: 4.6e-05
CL190_R: 4.6e-05
1e−5
1.2
1.0
0.8
0.6
0.4
0.2
0.0
VIC-VPN
100 µm

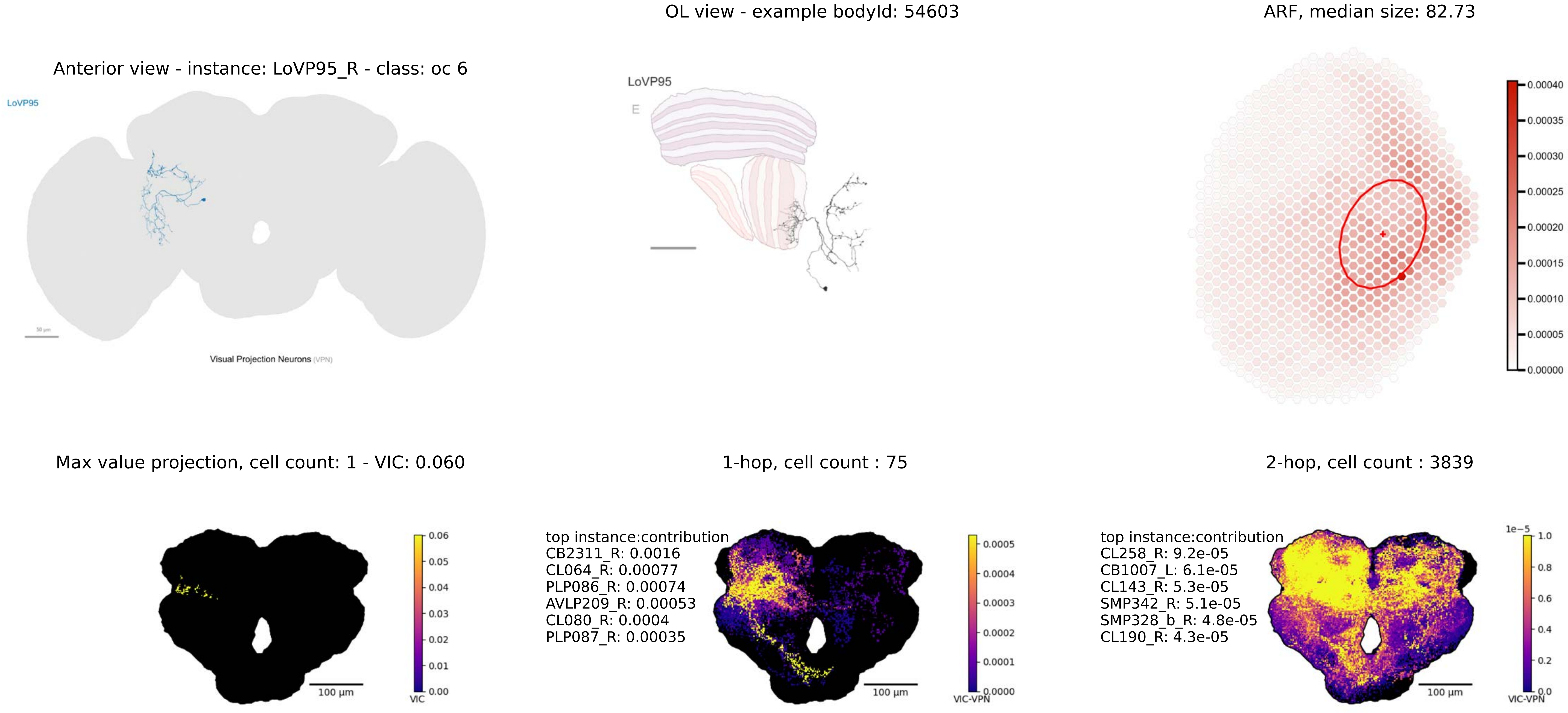
Anterior view - instance: LoVP95_R - class: oc 6
LoVP95
Visual Projection Neurons (VPN)
OL view - example bodyId: 54603
LoVP95
E
ARF, median size: 82.73
0.00040
0.00035
0.00030
0.00025
0.00020
0.00015
0.00010
0.00005
0.00000
Max value projection, cell count: 1 - VIC: 0.060
0.06
0.05
0.04
0.03
0.02
0.01
0.00
VIC
100 µm
1-hop, cell count : 75
top instance:contribution
CB2311_R: 0.0016
CL064_R: 0.00077
PLP086_R: 0.00074
AVLP209_R: 0.00053
CL080_R: 0.0004
PLP087_R: 0.00035
0.0005
0.0004
0.0003
0.0002
0.0001
0.0000
VIC-VPN
100 µm
2-hop, cell count : 3839
top instance:contribution
CL258_R: 9.2e-05
CB1007_L: 6.1e-05
CL143_R: 5.3e-05
SMP342_R: 5.1e-05
SMP328_b_R: 4.8e-05
CL190_R: 4.3e-05
1e−5
1.0
0.8
0.6
0.4
0.2
0.0
VIC-VPN
100 µm

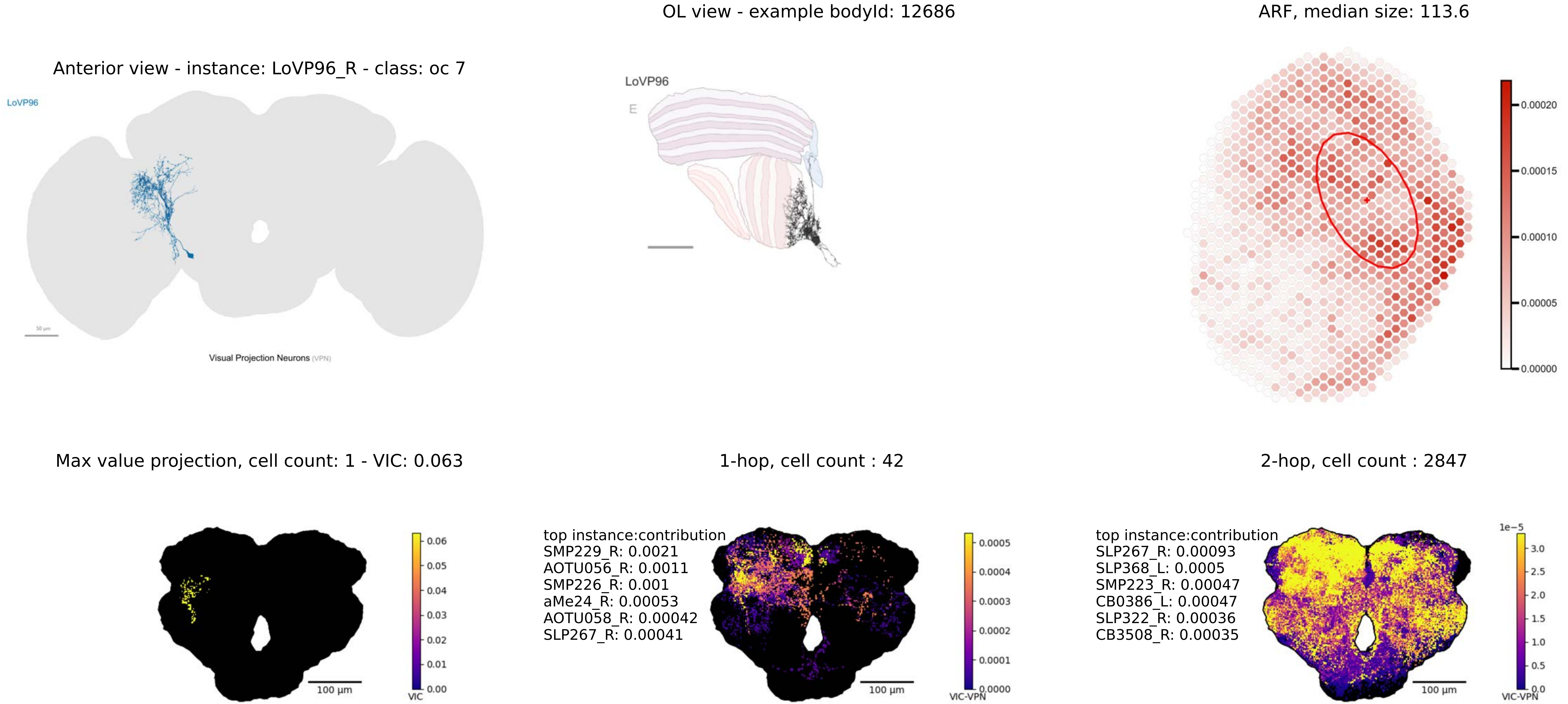

OL view - example bodyId: 12686
ARF, median size: 113.6
Anterior view - instance: LoVP96_R - class: oc 7
LoVP96
LoVP96
E
Visual Projection Neurons (VPN)
0.00020
0.00015
0.00010
0.00005
0.00000
Max value projection, cell count: 1 - VIC: 0.063
1-hop, cell count : 42
2-hop, cell count : 2847
0.06
0.05
0.04
0.03
0.02
0.01
0.00
VIC
100 µm
top instance:contribution
SMP229_R: 0.0021
AOTU056_R: 0.0011
SMP226_R: 0.001
aMe24_R: 0.00053
AOTU058_R: 0.00042
SLP267_R: 0.00041
0.0005
0.0004
0.0003
0.0002
0.0001
0.0000
VIC-VPN
100 µm
top instance:contribution
SLP267_R: 0.00093
SLP368_L: 0.0005
SMP223_R: 0.00047
CB0386_L: 0.00047
SLP322_R: 0.00036
CB3508_R: 0.00035
1e−5
3.0
2.5
2.0
1.5
1.0
0.5
0.0
VIC-VPN
100 µm

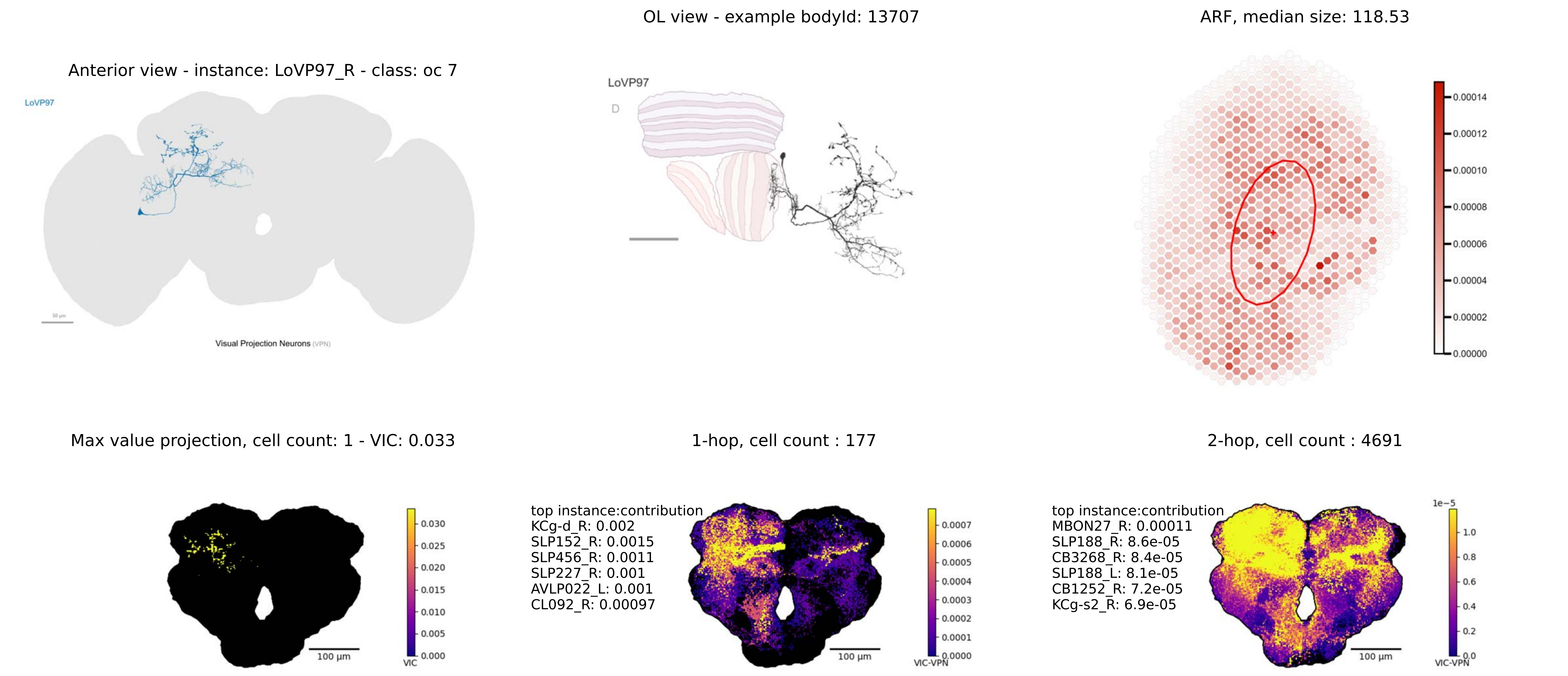
Anterior view - instance: LoVP97_R - class: oc 7
LoVP97
Visual Projection Neurons (VPN)
OL view - example bodyId: 13707
LoVP97
D
ARF, median size: 118.53
0.00014
0.00012
0.00010
0.00008
0.00006
0.00004
0.00002
0.00000
Max value projection, cell count: 1 - VIC: 0.033
100 µm
VIC
0.030
0.025
0.020
0.015
0.010
0.005
0.000
1-hop, cell count : 177
top instance:contribution
KCg-d_R: 0.002
SLP152_R: 0.0015
SLP456_R: 0.0011
SLP227_R: 0.001
AVLP022_L: 0.001
CL092_R: 0.00097
100 µm
VIC-VPN
0.0007
0.0006
0.0005
0.0004
0.0003
0.0002
0.0001
0.0000
2-hop, cell count : 4691
top instance:contribution
MBON27_R: 0.00011
SLP188_R: 8.6e-05
CB3268_R: 8.4e-05
SLP188_L: 8.1e-05
CB1252_R: 7.2e-05
KCg-s2_R: 6.9e-05
100 µm
1e−5
1.0
0.8
0.6
0.4
0.2
0.0
VIC-VPN

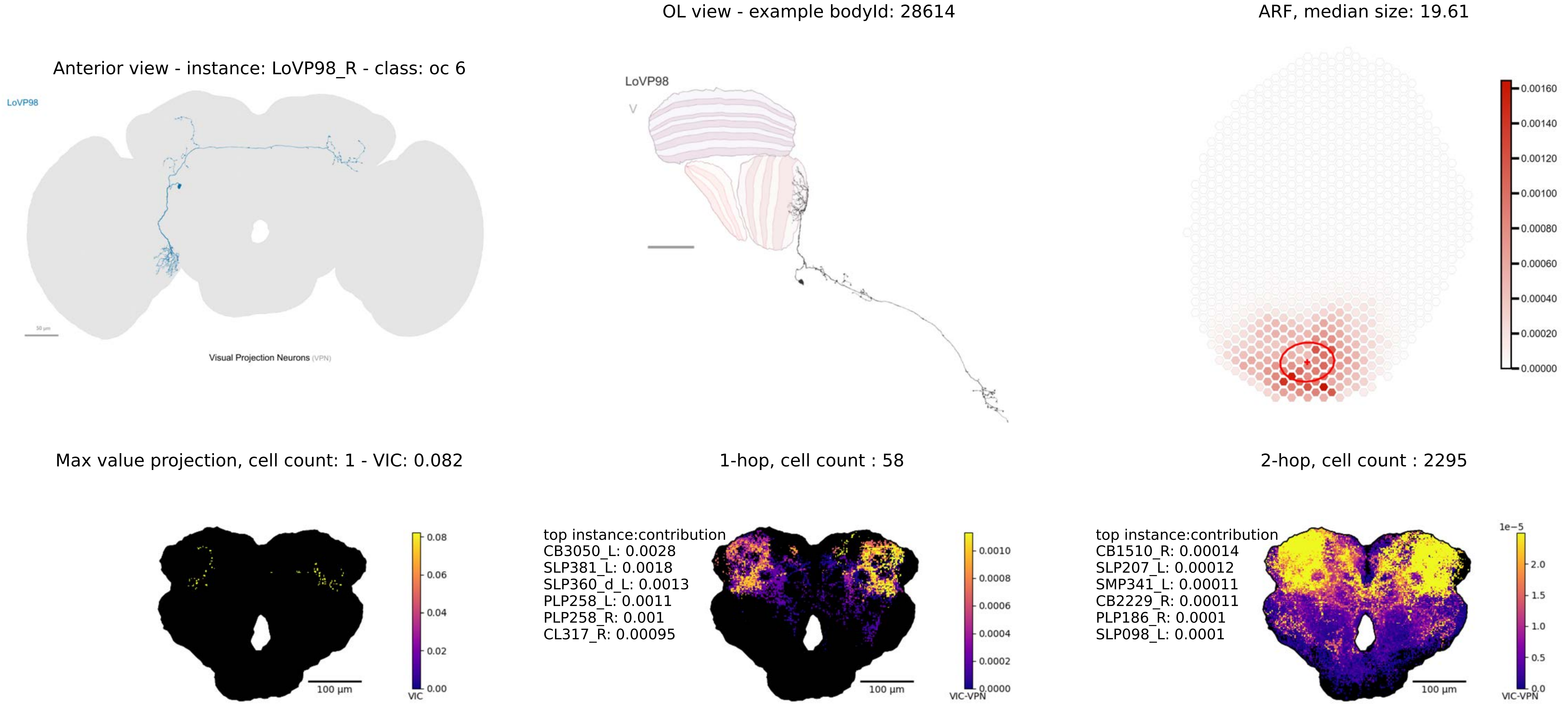

OL view - example bodyId: 28614
ARF, median size: 19.61
Anterior view - instance: LoVP98_R - class: oc 6
LoVP98
Visual Projection Neurons (VPN)
LoVP98
V
0.00160
0.00140
0.00120
0.00100
0.00080
0.00060
0.00040
0.00020
0.00000
Max value projection, cell count: 1 - VIC: 0.082
1-hop, cell count : 58
2-hop, cell count : 2295
0.08
0.06
0.04
0.02
0.00
VIC
100 µm
top instance:contribution
CB3050_L: 0.0028
SLP381_L: 0.0018
SLP360_d_L: 0.0013
PLP258_L: 0.0011
PLP258_R: 0.001
CL317_R: 0.00095
0.0010
0.0008
0.0006
0.0004
0.0002
0.0000
VIC-VPN
100 µm
top instance:contribution
CB1510_R: 0.00014
SLP207_L: 0.00012
SMP341_L: 0.00011
CB2229_R: 0.00011
PLP186_R: 0.0001
SLP098_L: 0.0001
1e−5
2.0
1.5
1.0
0.5
0.0
VIC-VPN
100 µm

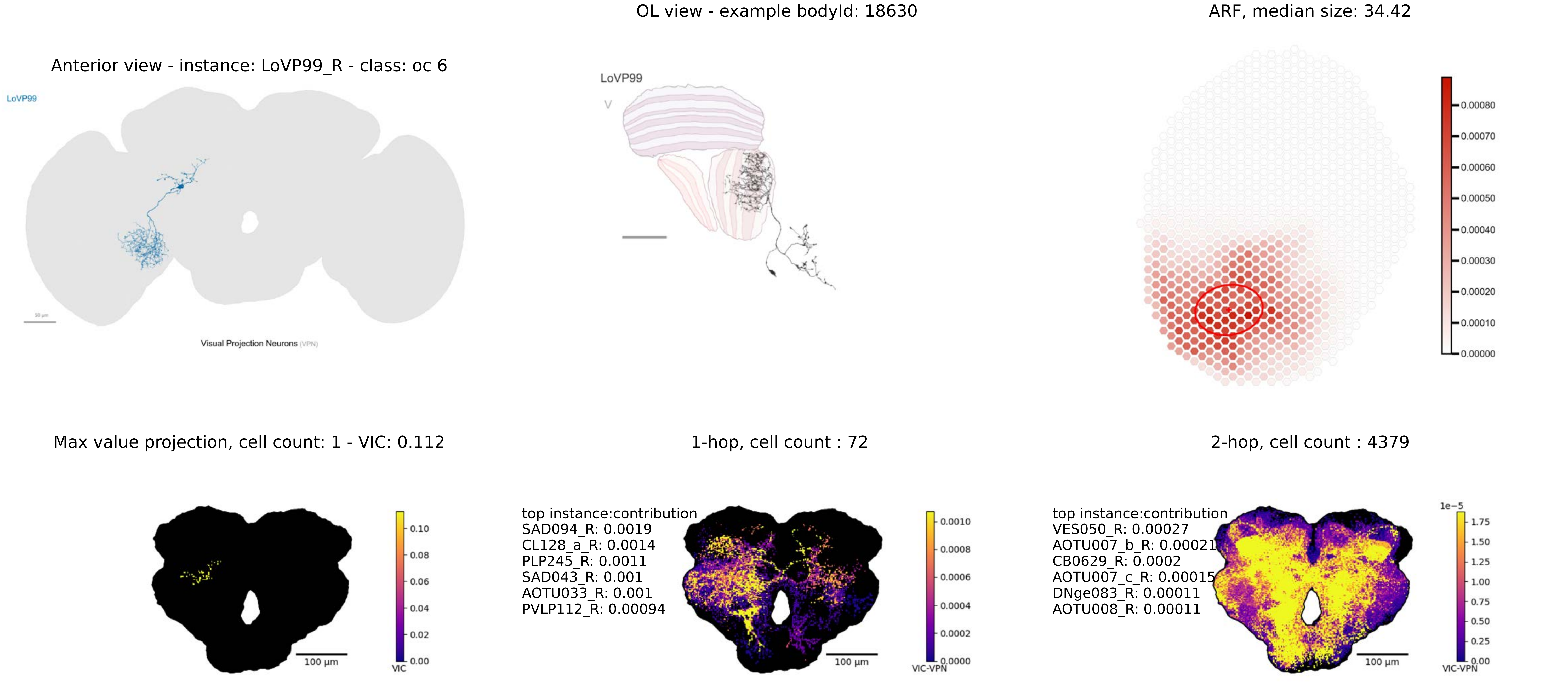
Anterior view - instance: LoVP99_R - class: oc 6
LoVP99
Visual Projection Neurons (VPN)
OL view - example bodyId: 18630
LoVP99
ARF, median size: 34.42
0.00080
0.00070
0.00060
0.00050
0.00040
0.00030
0.00020
0.00010
0.00000
Max value projection, cell count: 1 - VIC: 0.112
100 µm
VIC
0.10
0.08
0.06
0.04
0.02
0.00
1-hop, cell count : 72
top instance:contribution
SAD094_R: 0.0019
CL128_a_R: 0.0014
PLP245_R: 0.0011
SAD043_R: 0.001
AOTU033_R: 0.001
PVLP112_R: 0.00094
100 µm
VIC-VPN
0.0010
0.0008
0.0006
0.0004
0.0002
0.0000
2-hop, cell count : 4379
top instance:contribution
VES050_R: 0.00027
AOTU007_b_R: 0.00021
CB0629_R: 0.0002
AOTU007_c_R: 0.00015
DNge083_R: 0.00011
AOTU008_R: 0.00011
100 µm
1e−5
VIC-VPN
1.75
1.50
1.25
1.00
0.75
0.50
0.25
0.00

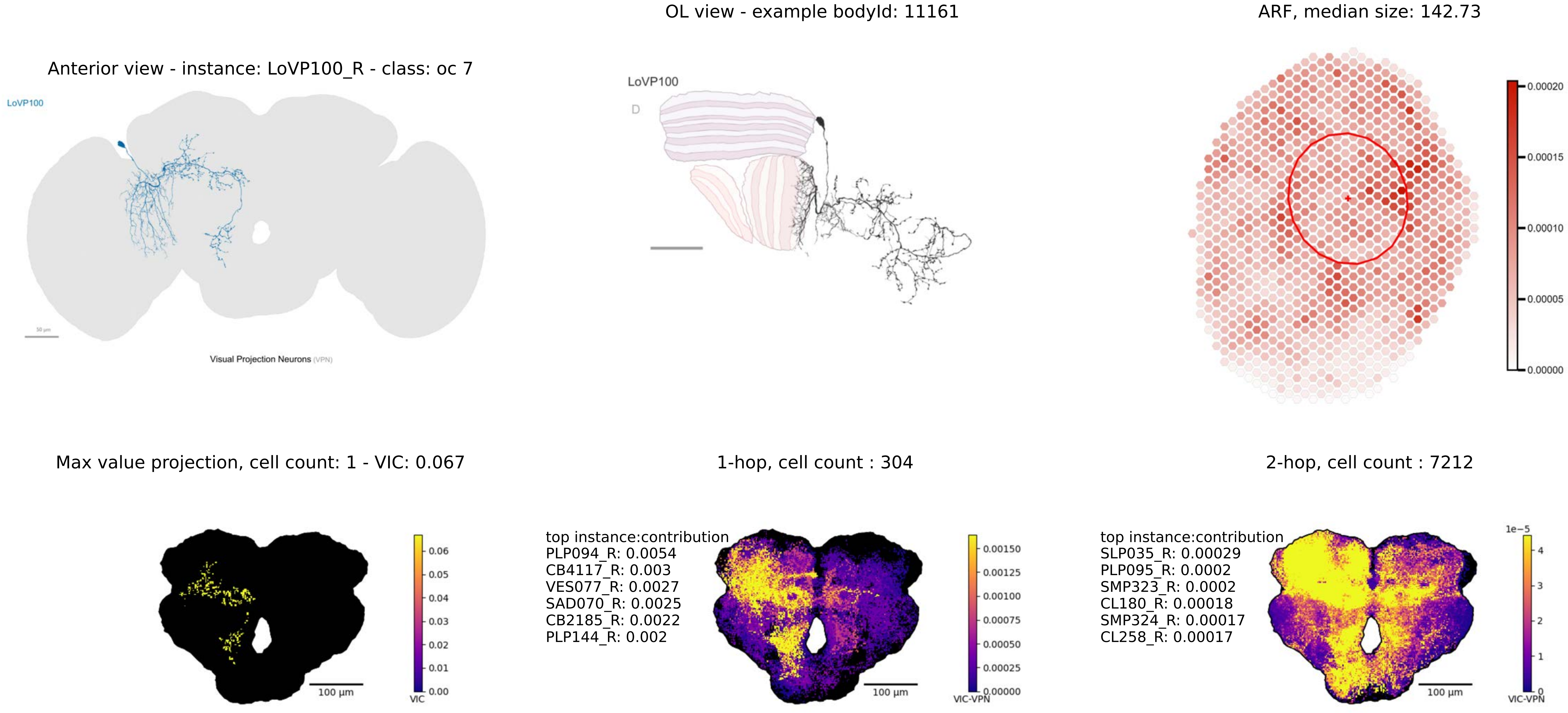

OL view - example bodyId: 11161
ARF, median size: 142.73
Anterior view - instance: LoVP100_R - class: oc 7
LoVP100
Visual Projection Neurons (VPN)
LoVP100
D
0.00020
0.00015
0.00010
0.00005
0.00000
Max value projection, cell count: 1 - VIC: 0.067
1-hop, cell count : 304
2-hop, cell count : 7212
0.06
0.05
0.04
0.03
0.02
0.01
0.00
VIC
100 µm
top instance:contribution
PLP094_R: 0.0054
CB4117_R: 0.003
VES077_R: 0.0027
SAD070_R: 0.0025
CB2185_R: 0.0022
PLP144_R: 0.002
0.00150
0.00125
0.00100
0.00075
0.00050
0.00025
0.00000
VIC-VPN
100 µm
top instance:contribution
SLP035_R: 0.00029
PLP095_R: 0.0002
SMP323_R: 0.0002
CL180_R: 0.00018
SMP324_R: 0.00017
CL258_R: 0.00017
1e−5
4
3
2
1
0
VIC-VPN
100 µm

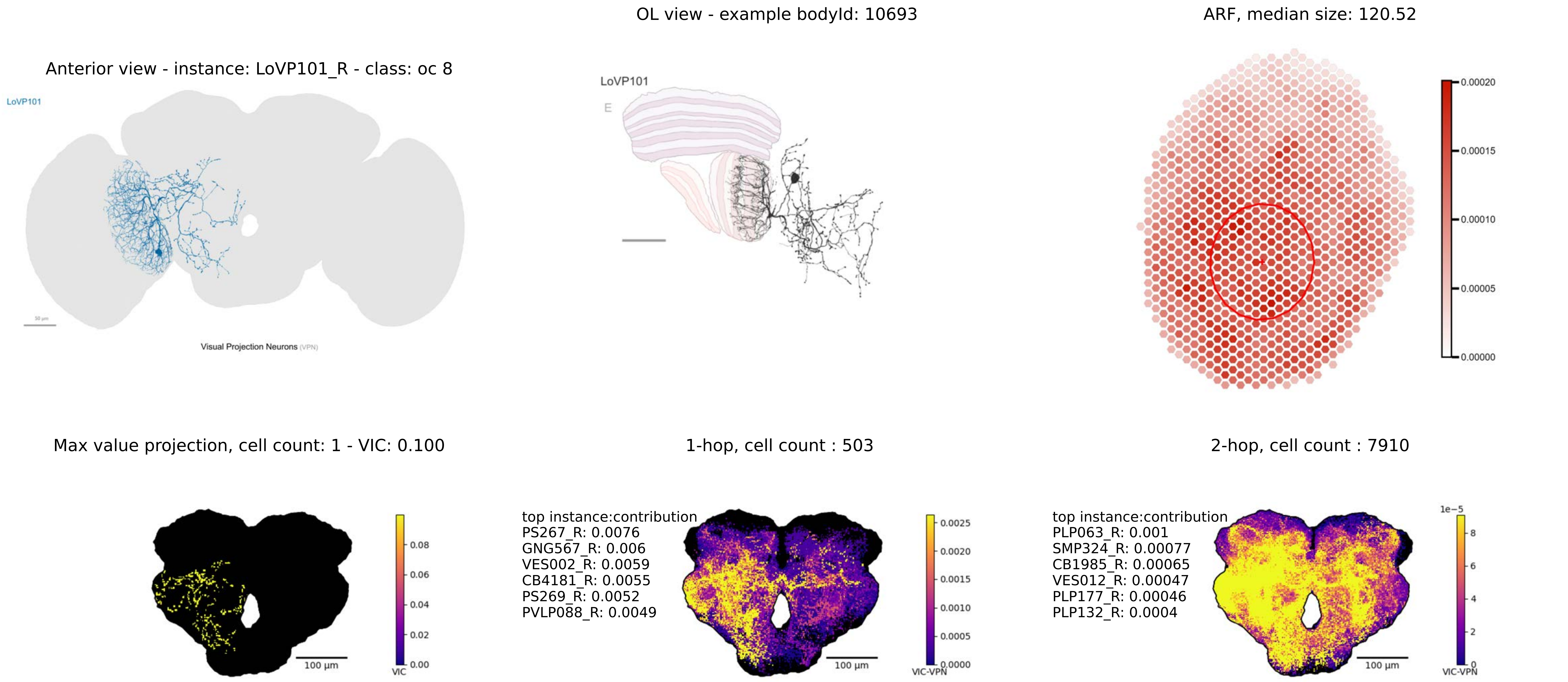

Anterior view - instance: LoVP101_R - class: oc 8
LoVP101
Visual Projection Neurons (VPN)
OL view - example bodyId: 10693
LoVP101
E
ARF, median size: 120.52
0.00020
0.00015
0.00010
0.00005
0.00000
Max value projection, cell count: 1 - VIC: 0.100
0.08
0.06
0.04
0.02
0.00
VIC
100 µm
1-hop, cell count : 503
top instance:contribution
PS267_R: 0.0076
GNG567_R: 0.006
VES002_R: 0.0059
CB4181_R: 0.0055
PS269_R: 0.0052
PVLP088_R: 0.0049
0.0025
0.0020
0.0015
0.0010
0.0005
0.0000
VIC-VPN
100 µm
2-hop, cell count : 7910
top instance:contribution
PLP063_R: 0.001
SMP324_R: 0.00077
CB1985_R: 0.00065
VES012_R: 0.00047
PLP177_R: 0.00046
PLP132_R: 0.0004
1e−5
8
6
4
2
0
VIC-VPN
100 µm

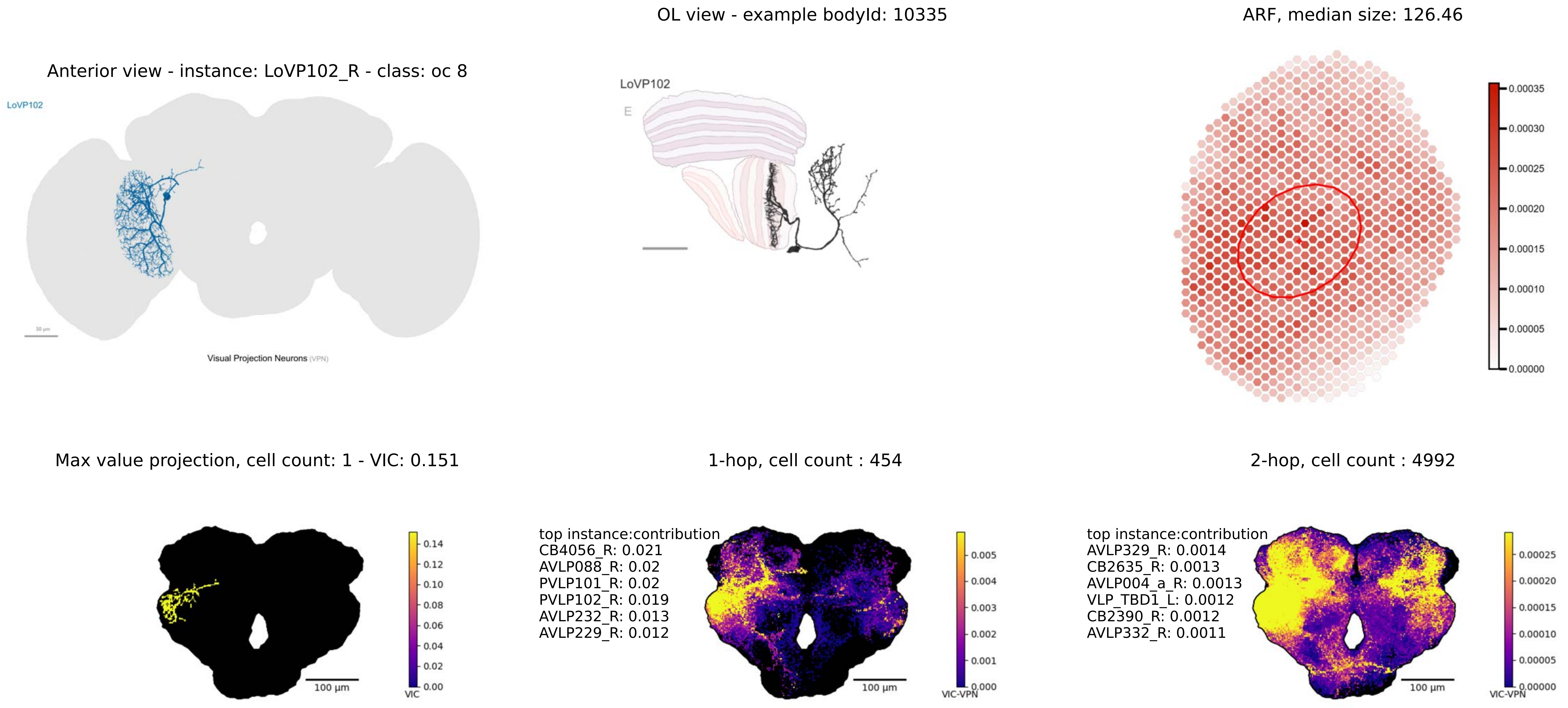

OL view - example bodyId: 10335
ARF, median size: 126.46
Anterior view - instance: LoVP102_R - class: oc 8
LoVP102
Visual Projection Neurons (VPN)
LoVP102
E
Max value projection, cell count: 1 - VIC: 0.151
1-hop, cell count : 454
2-hop, cell count : 4992
100 μm
VIC
top instance:contribution
CB4056_R: 0.021
AVLP088_R: 0.02
PVLP101_R: 0.02
PVLP102_R: 0.019
AVLP232_R: 0.013
AVLP229_R: 0.012
VIC-VPN
top instance:contribution
AVLP329_R: 0.0014
CB2635_R: 0.0013
AVLP004_a_R: 0.0013
VLP_TBD1_L: 0.0012
CB2390_R: 0.0012
AVLP332_R: 0.0011

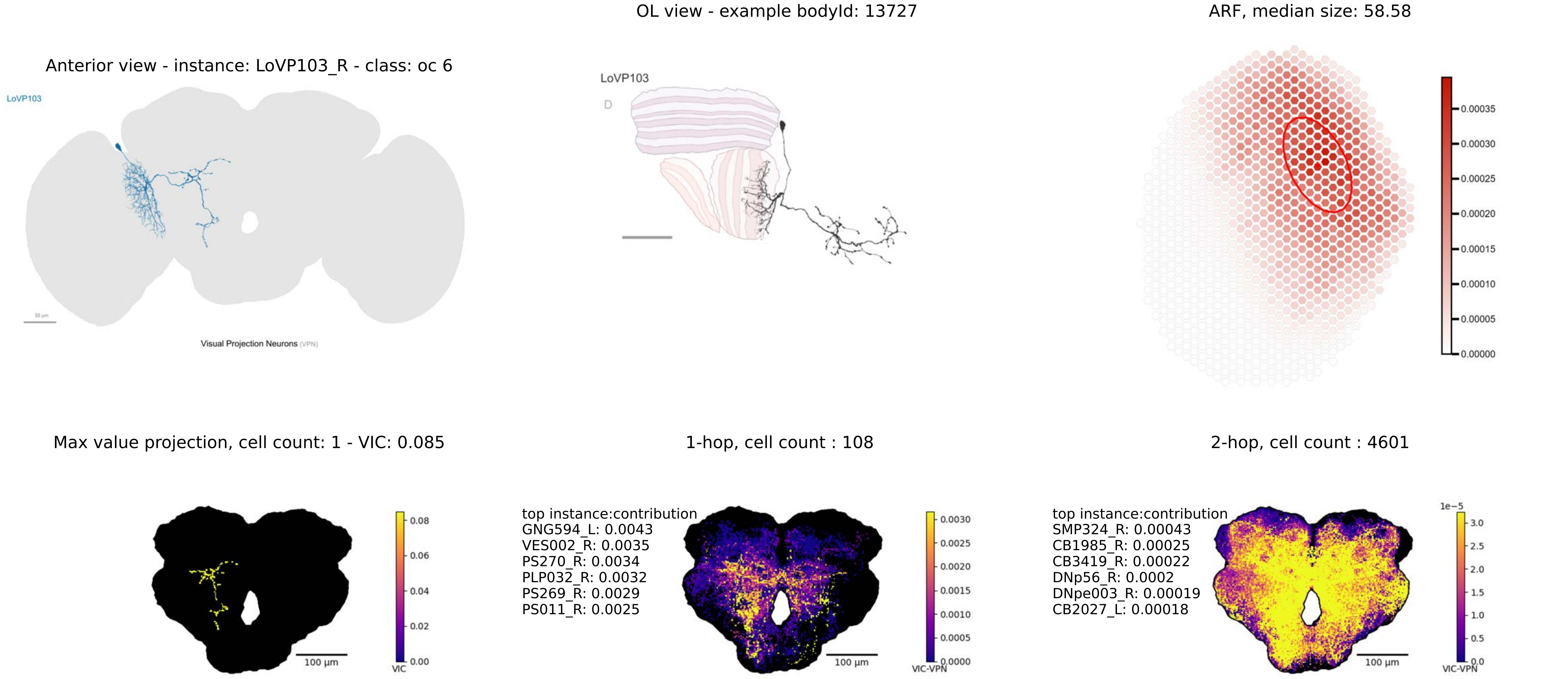

Anterior view - instance: LoVP103_R - class: oc 6
LoVP103
Visual Projection Neurons (VPN)
OL view - example bodyId: 13727
LoVP103
D
ARF, median size: 58.58
0.00035
0.00030
0.00025
0.00020
0.00015
0.00010
0.00005
0.00000
Max value projection, cell count: 1 - VIC: 0.085
0.08
0.06
0.04
0.02
0.00
VIC
100 μm
1-hop, cell count : 108
top instance:contribution
GNG594_L: 0.0043
VES002_R: 0.0035
PS270_R: 0.0034
PLP032_R: 0.0032
PS269_R: 0.0029
PS011_R: 0.0025
0.0030
0.0025
0.0020
0.0015
0.0010
0.0005
0.0000
VIC-VPN
100 μm
2-hop, cell count : 4601
top instance:contribution
SMP324_R: 0.00043
CB1985_R: 0.00025
CB3419_R: 0.00022
DNp56_R: 0.0002
DNpe003_R: 0.00019
CB2027_L: 0.00018
1e−5
3.0
2.5
2.0
1.5
1.0
0.5
0.0
VIC-VPN
100 μm

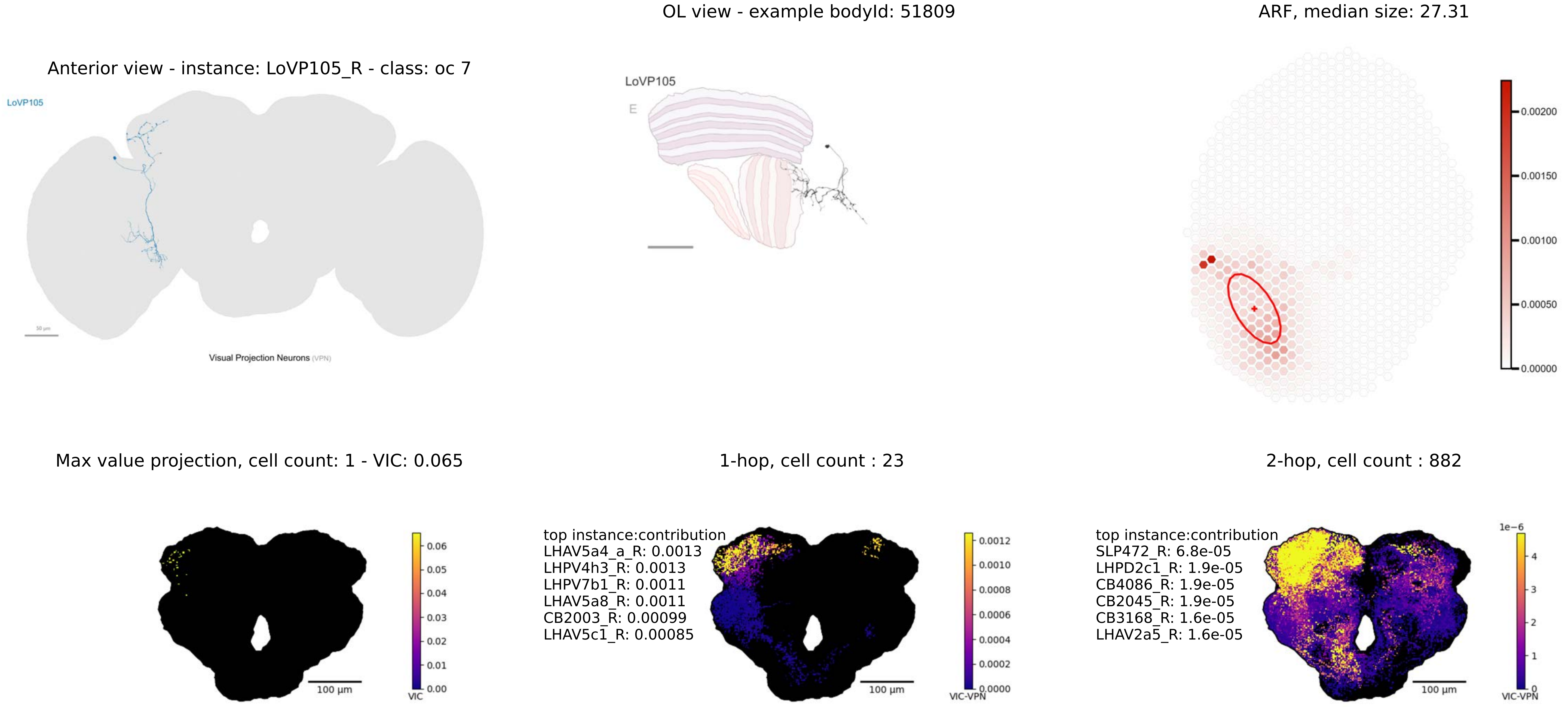
OL view - example bodyId: 51809
ARF, median size: 27.31
Anterior view - instance: LoVP105_R - class: oc 7
LoVP105
Visual Projection Neurons (VPN)
LoVP105
E
0.00200
0.00150
0.00100
0.00050
0.00000
Max value projection, cell count: 1 - VIC: 0.065
1-hop, cell count : 23
2-hop, cell count : 882
0.06
0.05
0.04
0.03
0.02
0.01
0.00
VIC
100 μm
top instance:contribution
LHAV5a4_a_R: 0.0013
LHPV4h3_R: 0.0013
LHPV7b1_R: 0.0011
LHAV5a8_R: 0.0011
CB2003_R: 0.00099
LHAV5c1_R: 0.00085
0.0012
0.0010
0.0008
0.0006
0.0004
0.0002
0.0000
VIC-VPN
100 μm
top instance:contribution
SLP472_R: 6.8e-05
LHPD2c1_R: 1.9e-05
CB4086_R: 1.9e-05
CB2045_R: 1.9e-05
CB3168_R: 1.6e-05
LHAV2a5_R: 1.6e-05
1e−6
4
3
2
1
0
VIC-VPN
100 μm

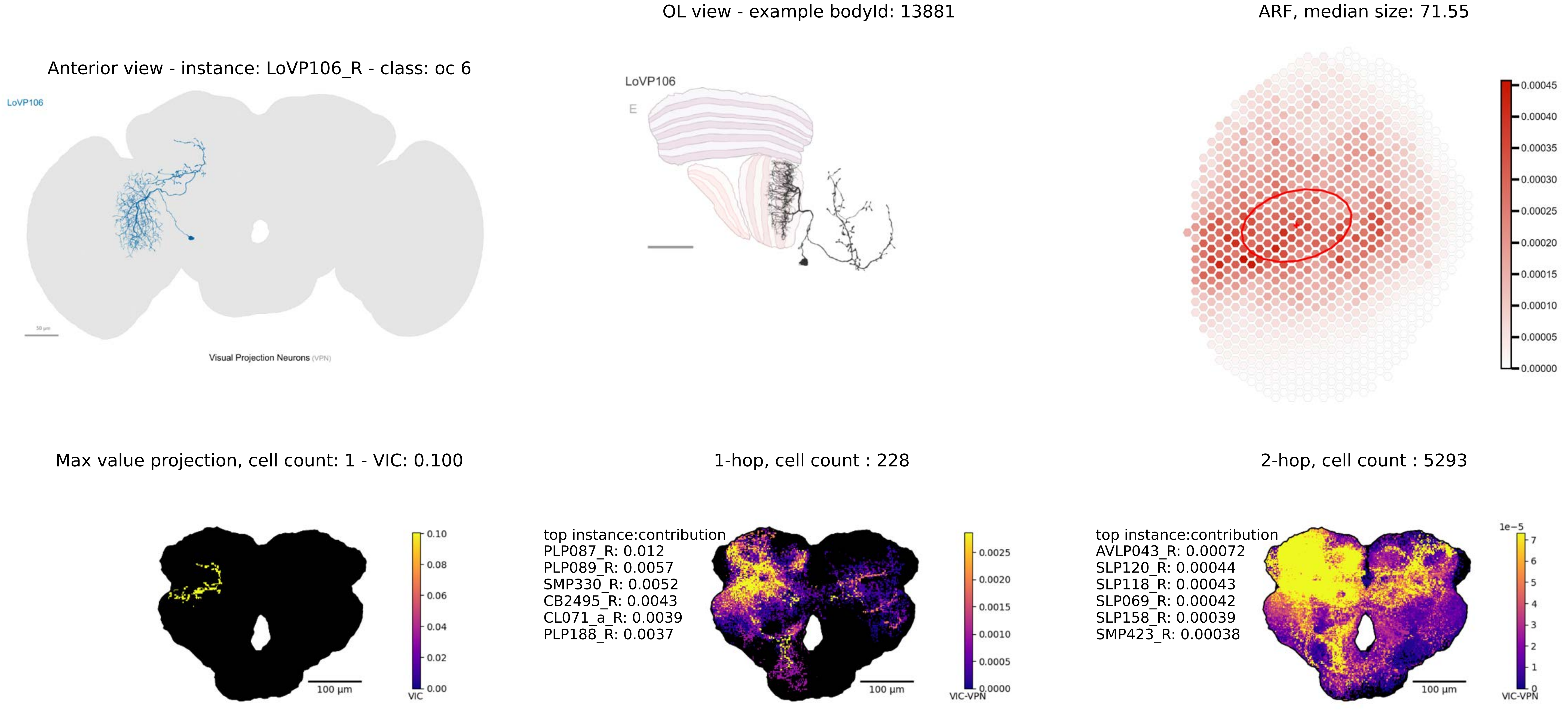

Anterior view - instance: LoVP106_R - class: oc 6
LoVP106
50 µm
Visual Projection Neurons (VPN)
OL view - example bodyId: 13881
LoVP106
E
ARF, median size: 71.55
0.00045
0.00040
0.00035
0.00030
0.00025
0.00020
0.00015
0.00010
0.00005
0.00000
Max value projection, cell count: 1 - VIC: 0.100
0.10
0.08
0.06
0.04
0.02
0.00
VIC
100 µm
1-hop, cell count : 228
top instance:contribution
PLP087_R: 0.012
PLP089_R: 0.0057
SMP330_R: 0.0052
CB2495_R: 0.0043
CL071_a_R: 0.0039
PLP188_R: 0.0037
0.0025
0.0020
0.0015
0.0010
0.0005
0.0000
VIC-VPN
100 µm
2-hop, cell count : 5293
top instance:contribution
AVLP043_R: 0.00072
SLP120_R: 0.00044
SLP118_R: 0.00043
SLP069_R: 0.00042
SLP158_R: 0.00039
SMP423_R: 0.00038
1e−5
7
6
5
4
3
2
1
0
VIC-VPN
100 µm

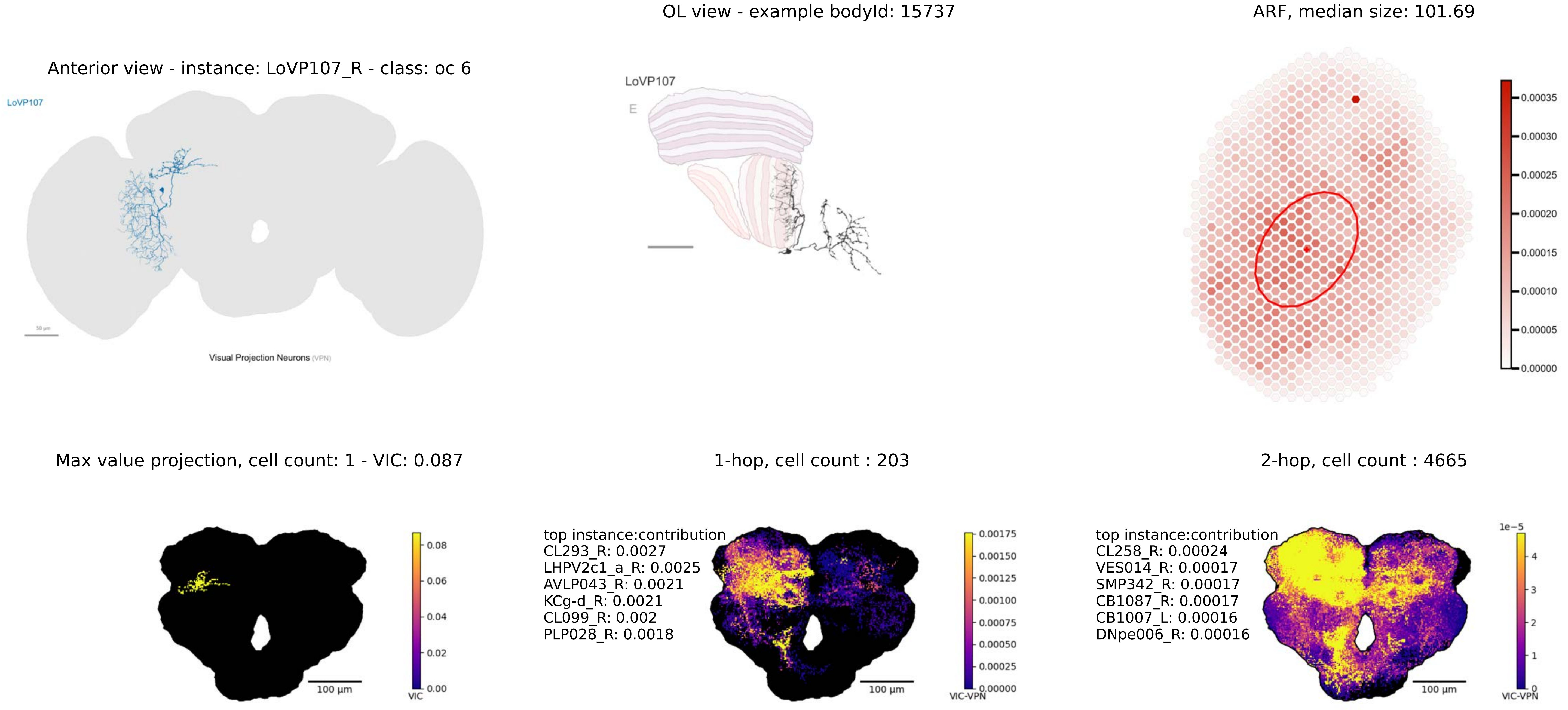
OL view - example bodyId: 15737
ARF, median size: 101.69
Anterior view - instance: LoVP107_R - class: oc 6
LoVP107
Visual Projection Neurons (VPN)
LoVP107
E
0.00035
0.00030
0.00025
0.00020
0.00015
0.00010
0.00005
0.00000
Max value projection, cell count: 1 - VIC: 0.087
1-hop, cell count : 203
2-hop, cell count : 4665
0.08
0.06
0.04
0.02
0.00
VIC
100 µm
top instance:contribution
CL293_R: 0.0027
LHPV2c1_a_R: 0.0025
AVLP043_R: 0.0021
KCg-d_R: 0.0021
CL099_R: 0.002
PLP028_R: 0.0018
0.00175
0.00150
0.00125
0.00100
0.00075
0.00050
0.00025
0.00000
VIC-VPN
100 µm
top instance:contribution
CL258_R: 0.00024
VES014_R: 0.00017
SMP342_R: 0.00017
CB1087_R: 0.00017
CB1007_L: 0.00016
DNpe006_R: 0.00016
1e−5
4
3
2
1
0
VIC-VPN
100 µm

OL view - example bodyId: 57463

ARF, median size: 89.04

Anterior view - instance: LoVP108_R - class: oc 8

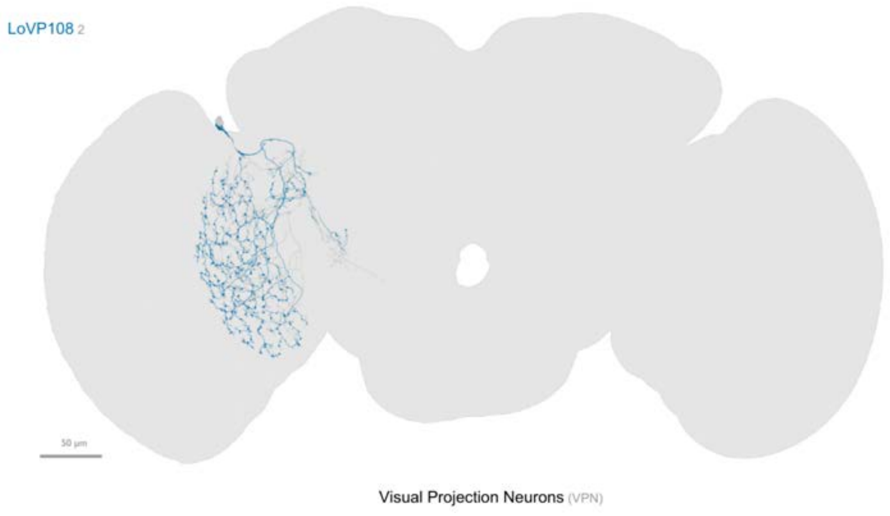


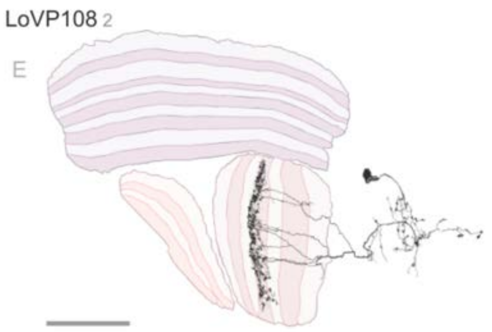


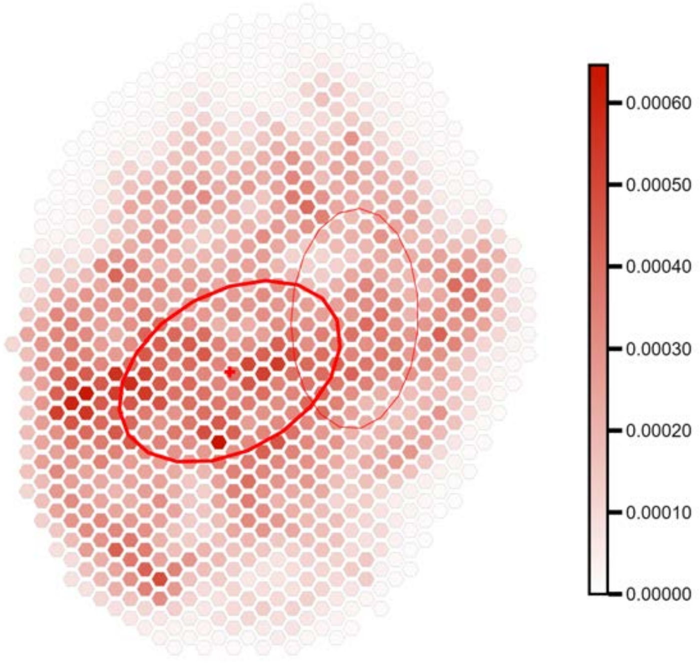


Max value projection, cell count: 2 - VIC: 0.159

1-hop, cell count : 114

2-hop, cell count : 3797

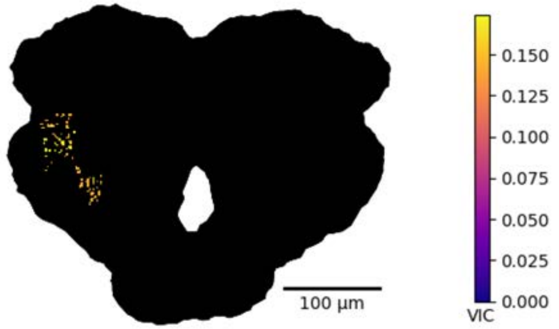


top instance:contribution
CB4165_R: 0.0013
PVLP105_R: 0.0011
PLP191_R: 0.00099
AVLP613_R: 0.00092
PVLP008_a3_R: 0.00081
PLP165_R: 0.0007

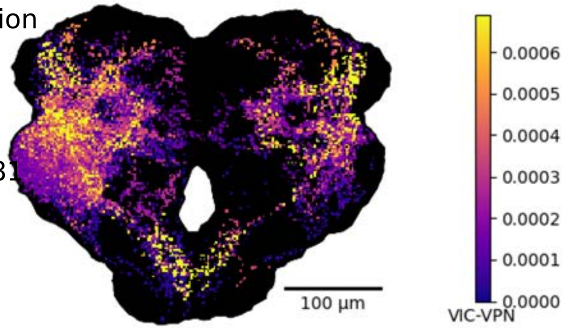


top instance:contribution
PVLP098_R: 0.00024
CB1109_R: 0.00022
AVLP282_R: 0.00021
CB1428_R: 0.00017
PLP192_R: 0.00014
CB2175_R: 0.00013

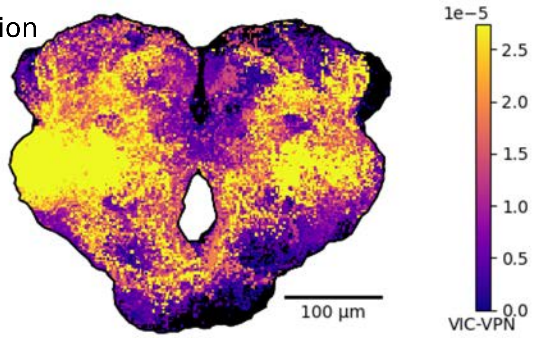

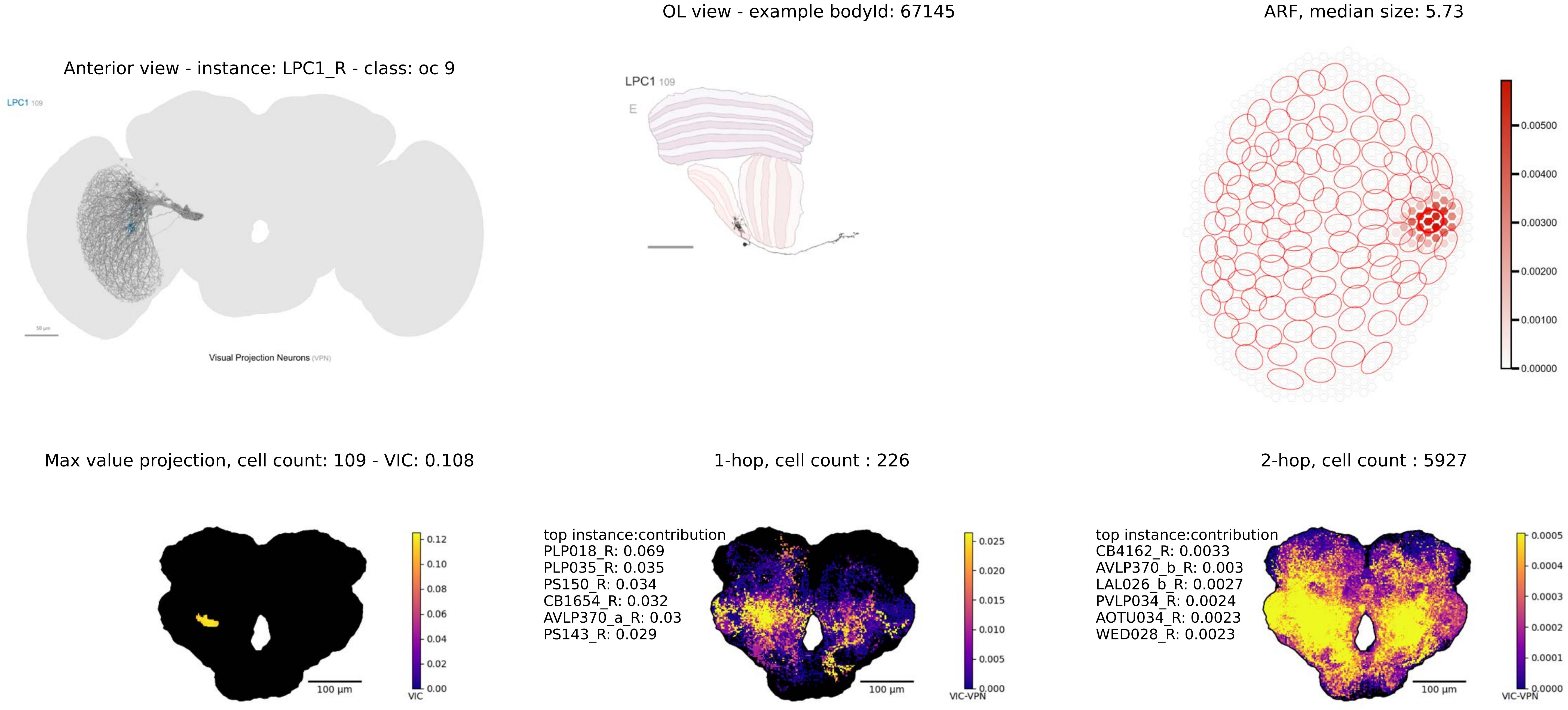

Anterior view - instance: LPC1_R - class: oc 9
LPC1 109
Visual Projection Neurons (VPN)
OL view - example bodyId: 67145
LPC1 109
E
ARF, median size: 5.73
0.00500
0.00400
0.00300
0.00200
0.00100
0.00000
Max value projection, cell count: 109 - VIC: 0.108
100 µm
VIC
0.12
0.10
0.08
0.06
0.04
0.02
0.00
1-hop, cell count : 226
top instance:contribution
PLP018_R: 0.069
PLP035_R: 0.035
PS150_R: 0.034
CB1654_R: 0.032
AVLP370_a_R: 0.03
PS143_R: 0.029
100 µm
VIC-VPN
0.025
0.020
0.015
0.010
0.005
0.000
2-hop, cell count : 5927
top instance:contribution
CB4162_R: 0.0033
AVLP370_b_R: 0.003
LAL026_b_R: 0.0027
PVLP034_R: 0.0024
AOTU034_R: 0.0023
WED028_R: 0.0023
100 µm
VIC-VPN
0.0005
0.0004
0.0003
0.0002
0.0001
0.0000

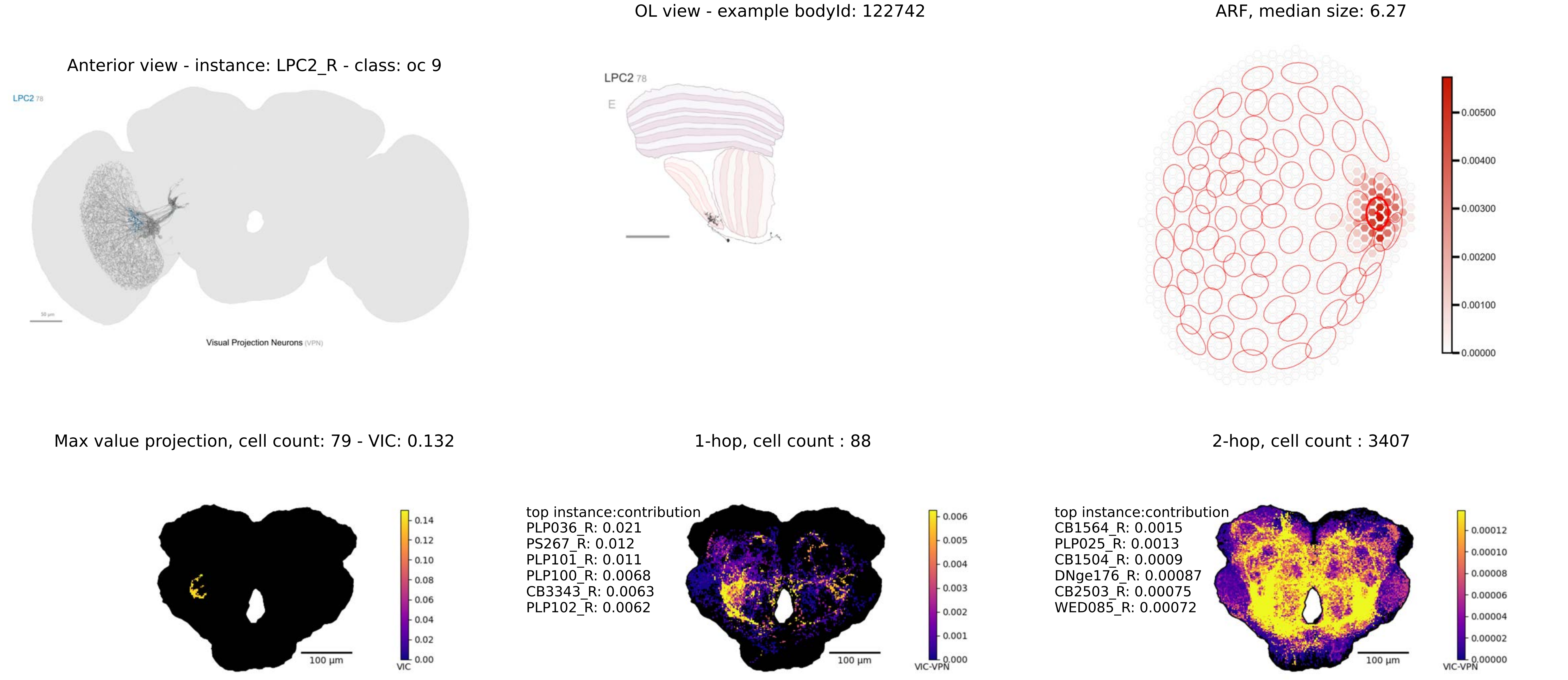

OL view - example bodyId: 122742
ARF, median size: 6.27
Anterior view - instance: LPC2_R - class: oc 9
LPC2 78
Visual Projection Neurons (VPN)
LPC2 78
E
0.00500
0.00400
0.00300
0.00200
0.00100
0.00000
Max value projection, cell count: 79 - VIC: 0.132
1-hop, cell count : 88
2-hop, cell count : 3407
0.14
0.12
0.10
0.08
0.06
0.04
0.02
0.00
VIC
100 µm
top instance:contribution
PLP036_R: 0.021
PS267_R: 0.012
PLP101_R: 0.011
PLP100_R: 0.0068
CB3343_R: 0.0063
PLP102_R: 0.0062
0.006
0.005
0.004
0.003
0.002
0.001
0.000
VIC-VPN
100 µm
top instance:contribution
CB1564_R: 0.0015
PLP025_R: 0.0013
CB1504_R: 0.0009
DNge176_R: 0.00087
CB2503_R: 0.00075
WED085_R: 0.00072
0.00012
0.00010
0.00008
0.00006
0.00004
0.00002
0.00000
VIC-VPN
100 µm

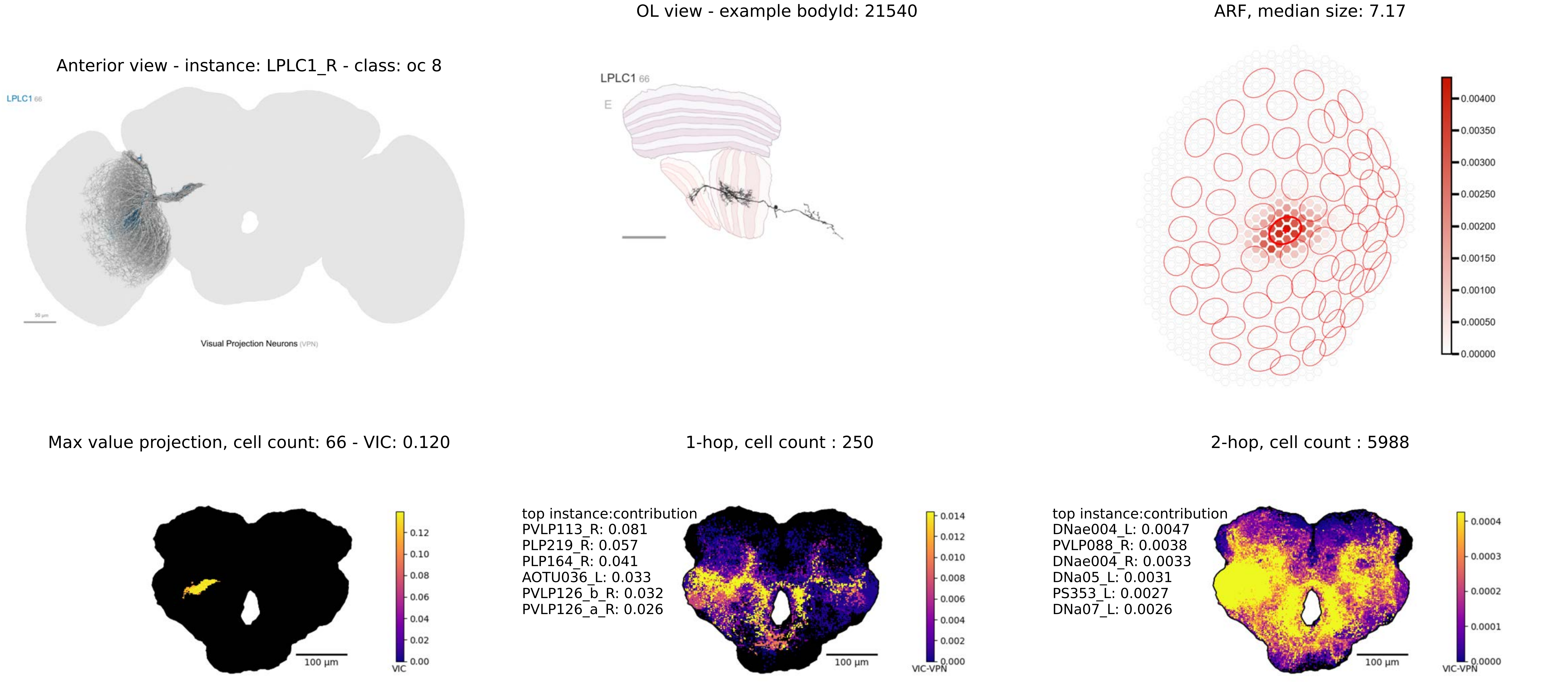

OL view - example bodyId: 21540
ARF, median size: 7.17
Anterior view - instance: LPLC1_R - class: oc 8
LPLC1 66
Visual Projection Neurons (VPN)
LPLC1 66
E
0.00400
0.00350
0.00300
0.00250
0.00200
0.00150
0.00100
0.00050
0.00000
Max value projection, cell count: 66 - VIC: 0.120
1-hop, cell count : 250
2-hop, cell count : 5988
100 µm
VIC
0.12
0.10
0.08
0.06
0.04
0.02
0.00
top instance:contribution
PVLP113_R: 0.081
PLP219_R: 0.057
PLP164_R: 0.041
AOTU036_L: 0.033
PVLP126_b_R: 0.032
PVLP126_a_R: 0.026
100 µm
VIC-VPN
0.014
0.012
0.010
0.008
0.006
0.004
0.002
0.000
top instance:contribution
DNae004_L: 0.0047
PVLP088_R: 0.0038
DNae004_R: 0.0033
DNa05_L: 0.0031
PS353_L: 0.0027
DNa07_L: 0.0026
100 µm
VIC-VPN
0.0004
0.0003
0.0002
0.0001
0.0000

OL view - example bodyId: 37431

ARF, median size: 10.24

Anterior view - instance: LPLC2_R - class: oc 8

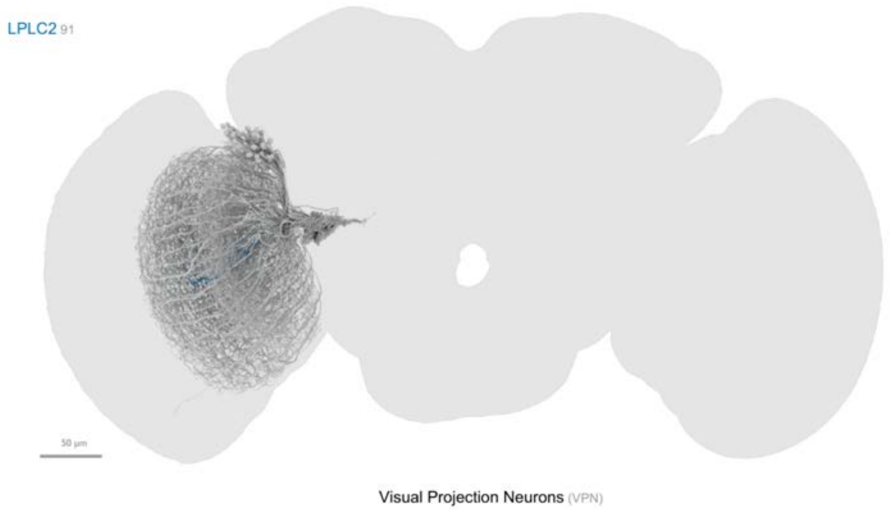


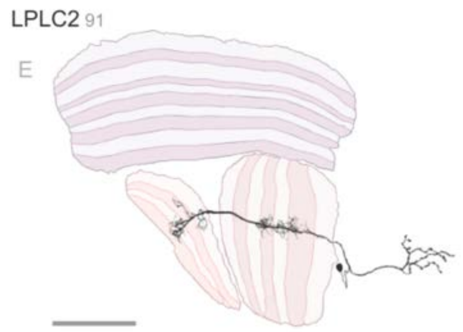


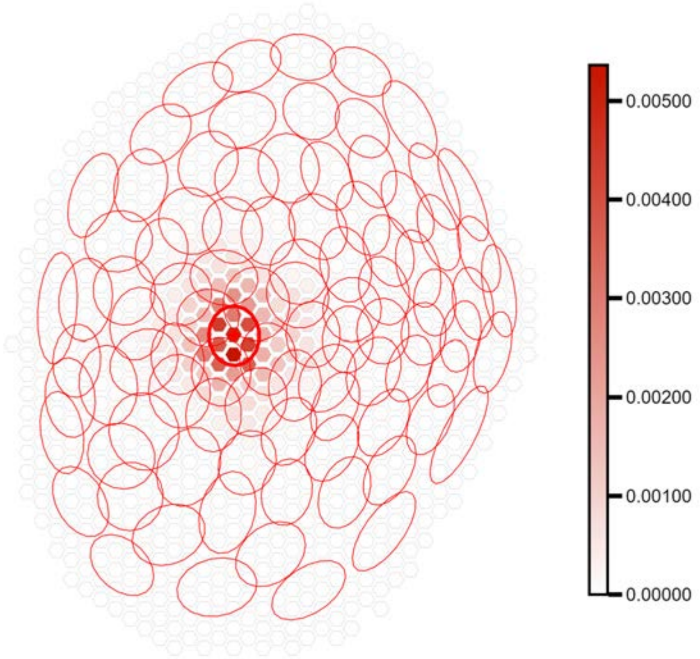


Max value projection, cell count: 91 - VIC: 0.125

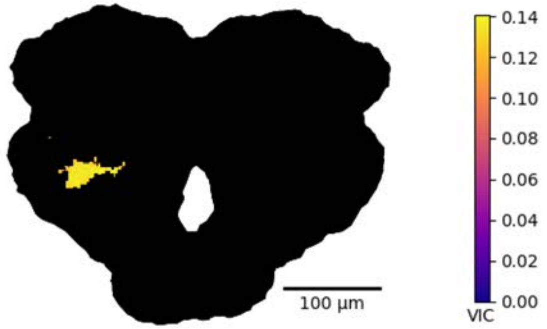


1-hop, cell count : 283

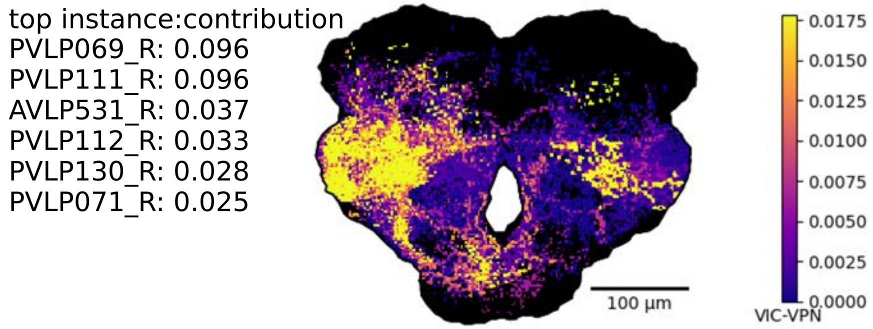


2-hop, cell count : 5478

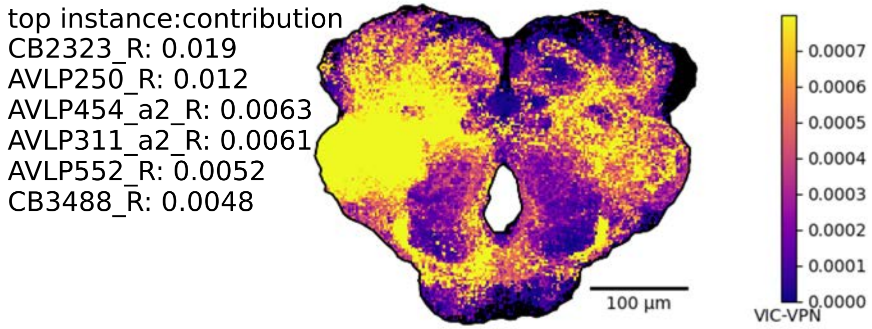

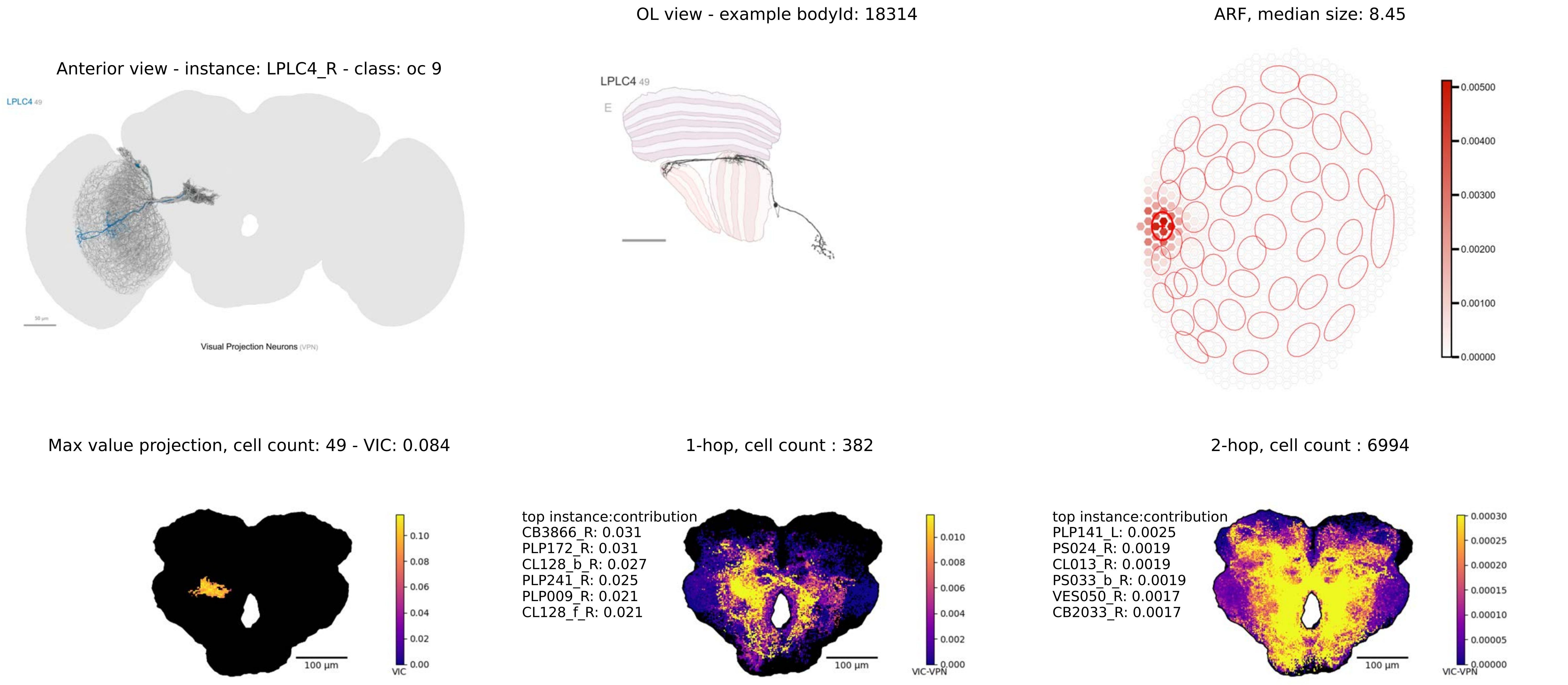

OL view - example bodyId: 18314
ARF, median size: 8.45
Anterior view - instance: LPLC4_R - class: oc 9
LPLC4 49
Visual Projection Neurons (VPN)
LPLC4 49
E
0.00500
0.00400
0.00300
0.00200
0.00100
0.00000
Max value projection, cell count: 49 - VIC: 0.084
1-hop, cell count : 382
2-hop, cell count : 6994
100 µm
VIC
0.10
0.08
0.06
0.04
0.02
0.00
top instance:contribution
CB3866_R: 0.031
PLP172_R: 0.031
CL128_b_R: 0.027
PLP241_R: 0.025
PLP009_R: 0.021
CL128_f_R: 0.021
100 µm
VIC-VPN
0.010
0.008
0.006
0.004
0.002
0.000
top instance:contribution
PLP141_L: 0.0025
PS024_R: 0.0019
CL013_R: 0.0019
PS033_b_R: 0.0019
VES050_R: 0.0017
CB2033_R: 0.0017
100 µm
VIC-VPN
0.00030
0.00025
0.00020
0.00015
0.00010
0.00005
0.00000

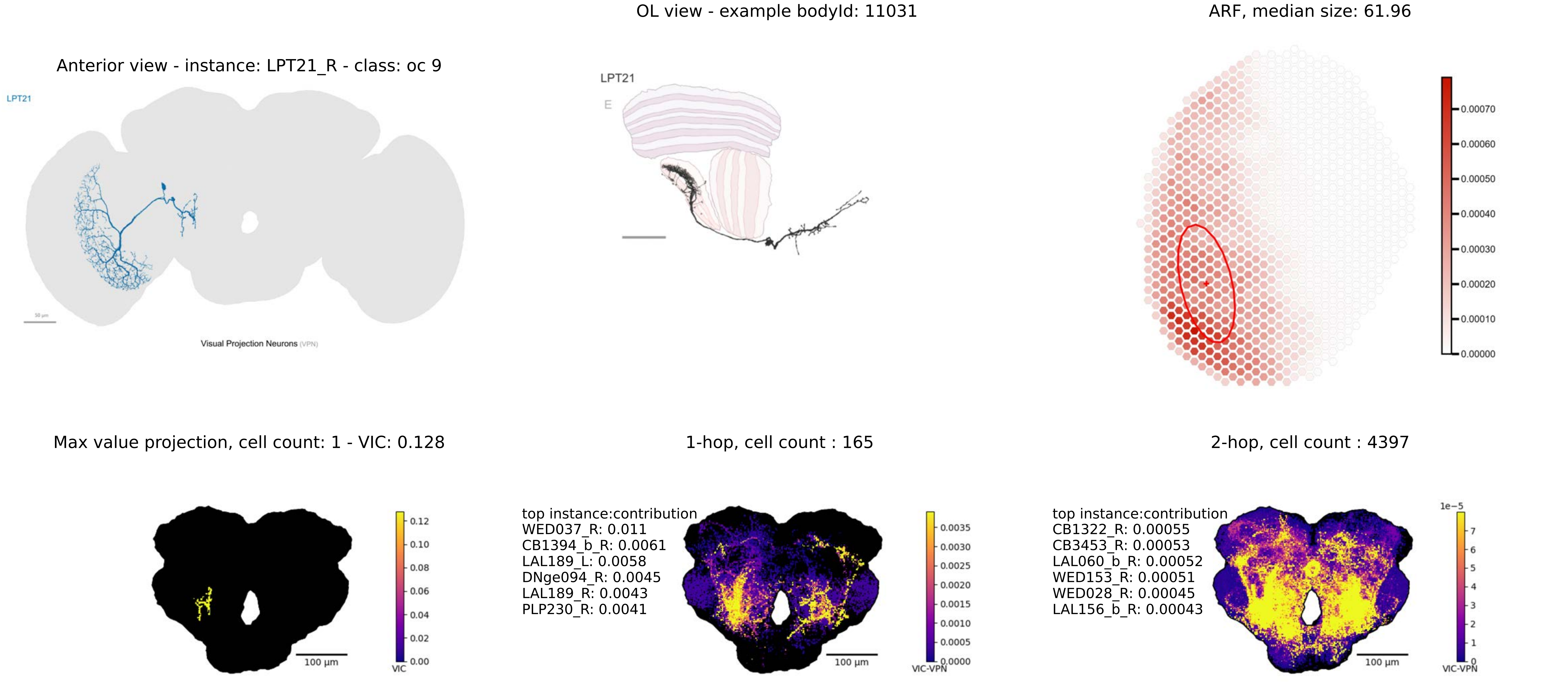

OL view - example bodyId: 11031
ARF, median size: 61.96
Anterior view - instance: LPT21_R - class: oc 9
LPT21
Visual Projection Neurons (VPN)
LPT21
E
0.00070
0.00060
0.00050
0.00040
0.00030
0.00020
0.00010
0.00000
Max value projection, cell count: 1 - VIC: 0.128
1-hop, cell count : 165
2-hop, cell count : 4397
0.12
0.10
0.08
0.06
0.04
0.02
0.00
VIC
100 µm
top instance:contribution
WED037_R: 0.011
CB1394_b_R: 0.0061
LAL189_L: 0.0058
DNge094_R: 0.0045
LAL189_R: 0.0043
PLP230_R: 0.0041
0.0035
0.0030
0.0025
0.0020
0.0015
0.0010
0.0005
0.0000
VIC-VPN
100 µm
top instance:contribution
CB1322_R: 0.00055
CB3453_R: 0.00053
LAL060_b_R: 0.00052
WED153_R: 0.00051
WED028_R: 0.00045
LAL156_b_R: 0.00043
1e−5
7
6
5
4
3
2
1
0
VIC-VPN
100 µm

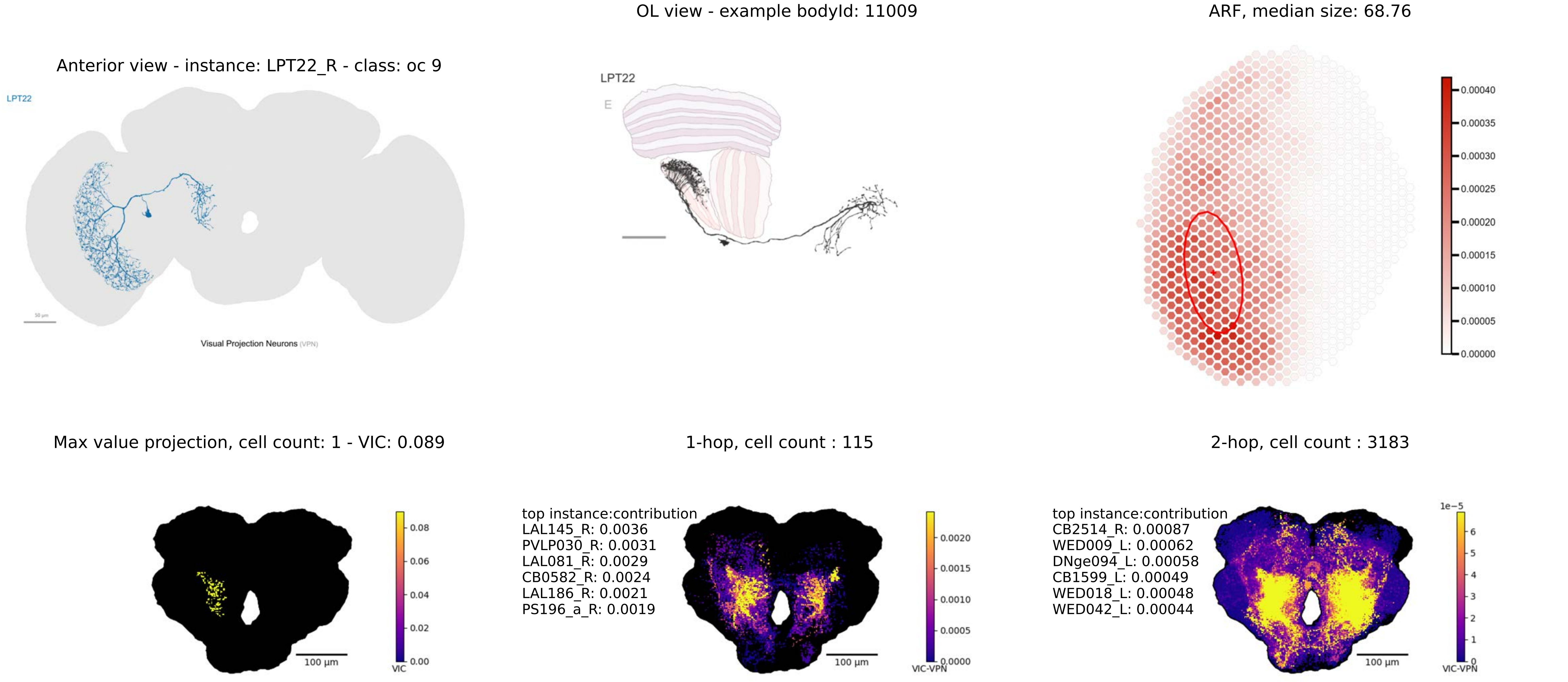
OL view - example bodyId: 11009
ARF, median size: 68.76
Anterior view - instance: LPT22_R - class: oc 9
LPT22
Visual Projection Neurons (VPN)
LPT22
E
0.00040
0.00035
0.00030
0.00025
0.00020
0.00015
0.00010
0.00005
0.00000
Max value projection, cell count: 1 - VIC: 0.089
1-hop, cell count : 115
2-hop, cell count : 3183
0.08
0.06
0.04
0.02
0.00
VIC
100 µm
top instance:contribution
LAL145_R: 0.0036
PVLP030_R: 0.0031
LAL081_R: 0.0029
CB0582_R: 0.0024
LAL186_R: 0.0021
PS196_a_R: 0.0019
0.0020
0.0015
0.0010
0.0005
0.0000
VIC-VPN
100 µm
top instance:contribution
CB2514_R: 0.00087
WED009_L: 0.00062
DNge094_L: 0.00058
CB1599_L: 0.00049
WED018_L: 0.00048
WED042_L: 0.00044
1e−5
6
5
4
3
2
1
0
VIC-VPN
100 µm

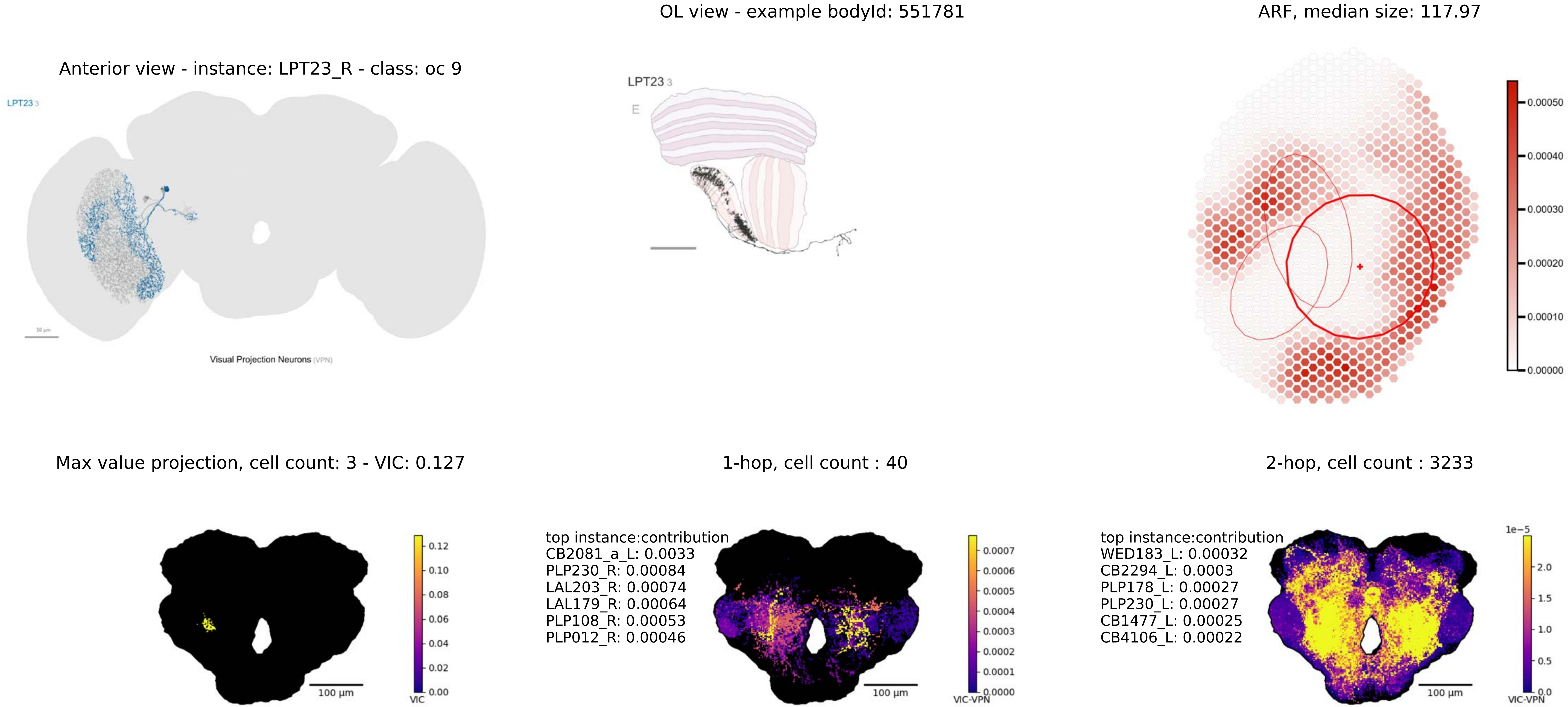
Anterior view - instance: LPT23_R - class: oc 9
LPT23 3
Visual Projection Neurons (VPN)
OL view - example bodyId: 551781
LPT23 3
E
ARF, median size: 117.97
0.00050
0.00040
0.00030
0.00020
0.00010
0.00000
Max value projection, cell count: 3 - VIC: 0.127
100 µm
VIC
0.12
0.10
0.08
0.06
0.04
0.02
0.00
1-hop, cell count : 40
top instance:contribution
CB2081_a_L: 0.0033
PLP230_R: 0.00084
LAL203_R: 0.00074
LAL179_R: 0.00064
PLP108_R: 0.00053
PLP012_R: 0.00046
100 µm
VIC-VPN
0.0007
0.0006
0.0005
0.0004
0.0003
0.0002
0.0001
0.0000
2-hop, cell count : 3233
top instance:contribution
WED183_L: 0.00032
CB2294_L: 0.0003
PLP178_L: 0.00027
PLP230_L: 0.00027
CB1477_L: 0.00025
CB4106_L: 0.00022
100 µm
1e−5
VIC-VPN
2.0
1.5
1.0
0.5
0.0

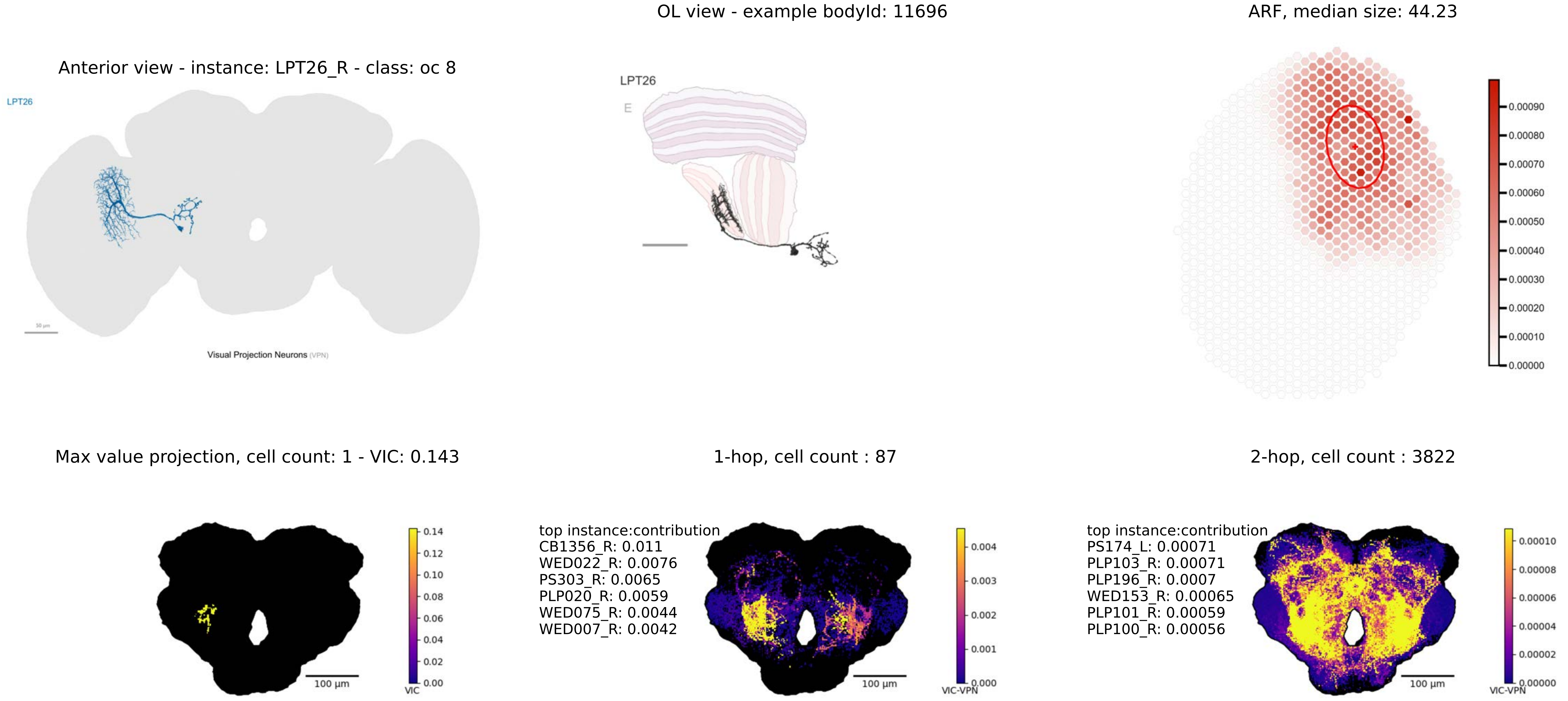
OL view - example bodyId: 11696
ARF, median size: 44.23
Anterior view - instance: LPT26_R - class: oc 8
LPT26
50 μm
Visual Projection Neurons (VPN)
LPT26
E
0.00090
0.00080
0.00070
0.00060
0.00050
0.00040
0.00030
0.00020
0.00010
0.00000
Max value projection, cell count: 1 - VIC: 0.143
1-hop, cell count : 87
2-hop, cell count : 3822
100 μm
VIC
0.14
0.12
0.10
0.08
0.06
0.04
0.02
0.00
top instance:contribution
CB1356_R: 0.011
WED022_R: 0.0076
PS303_R: 0.0065
PLP020_R: 0.0059
WED075_R: 0.0044
WED007_R: 0.0042
100 μm
VIC-VPN
0.004
0.003
0.002
0.001
0.000
top instance:contribution
PS174_L: 0.00071
PLP103_R: 0.00071
PLP196_R: 0.0007
WED153_R: 0.00065
PLP101_R: 0.00059
PLP100_R: 0.00056
100 μm
VIC-VPN
0.00010
0.00008
0.00006
0.00004
0.00002
0.00000

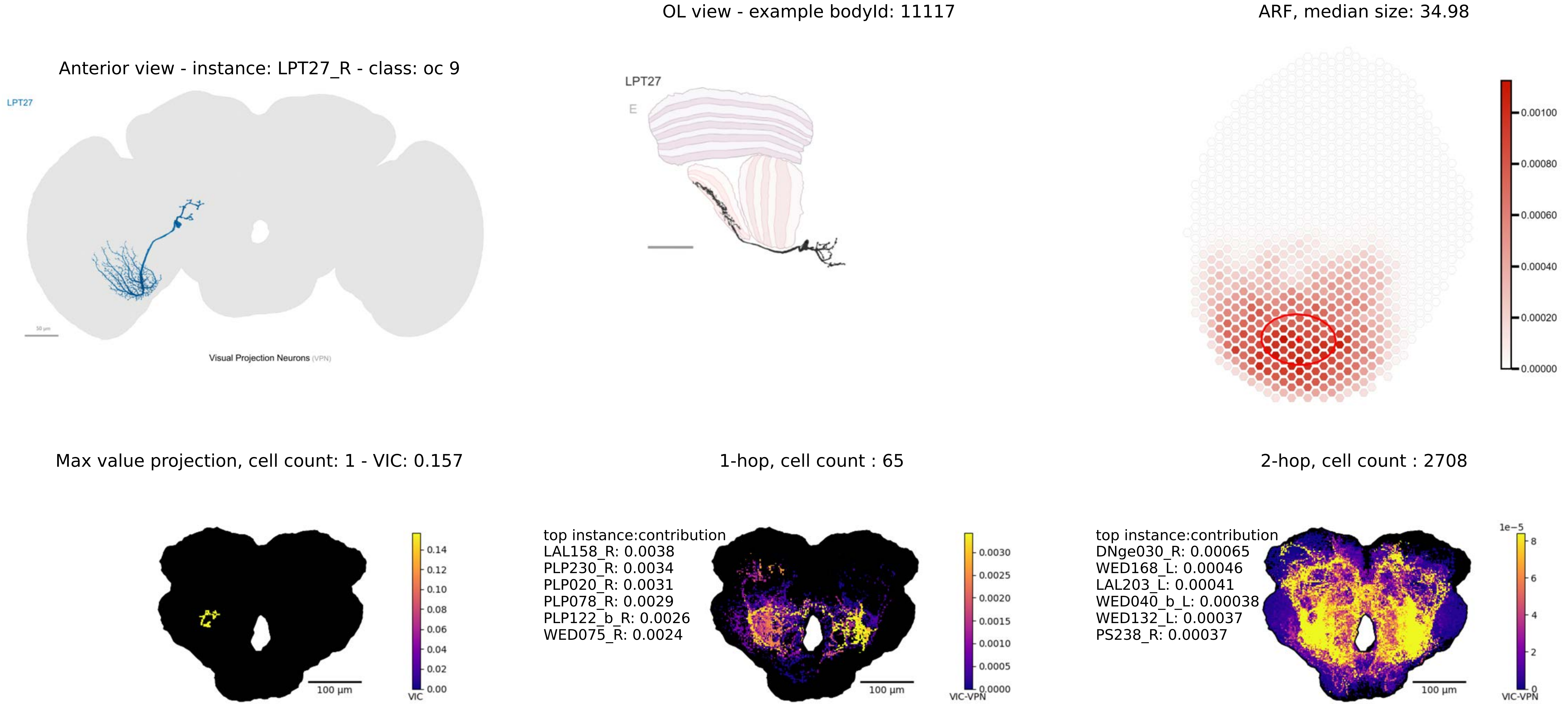

OL view - example bodyId: 11117
ARF, median size: 34.98
Anterior view - instance: LPT27_R - class: oc 9
LPT27
Visual Projection Neurons (VPN)
LPT27
E
0.00100
0.00080
0.00060
0.00040
0.00020
0.00000
Max value projection, cell count: 1 - VIC: 0.157
1-hop, cell count : 65
2-hop, cell count : 2708
0.14
0.12
0.10
0.08
0.06
0.04
0.02
0.00
VIC
100 µm
top instance:contribution
LAL158_R: 0.0038
PLP230_R: 0.0034
PLP020_R: 0.0031
PLP078_R: 0.0029
PLP122_b_R: 0.0026
WED075_R: 0.0024
0.0030
0.0025
0.0020
0.0015
0.0010
0.0005
0.0000
VIC-VPN
100 µm
top instance:contribution
DNge030_R: 0.00065
WED168_L: 0.00046
LAL203_L: 0.00041
WED040_b_L: 0.00038
WED132_L: 0.00037
PS238_R: 0.00037
1e−5
8
6
4
2
0
VIC-VPN
100 µm

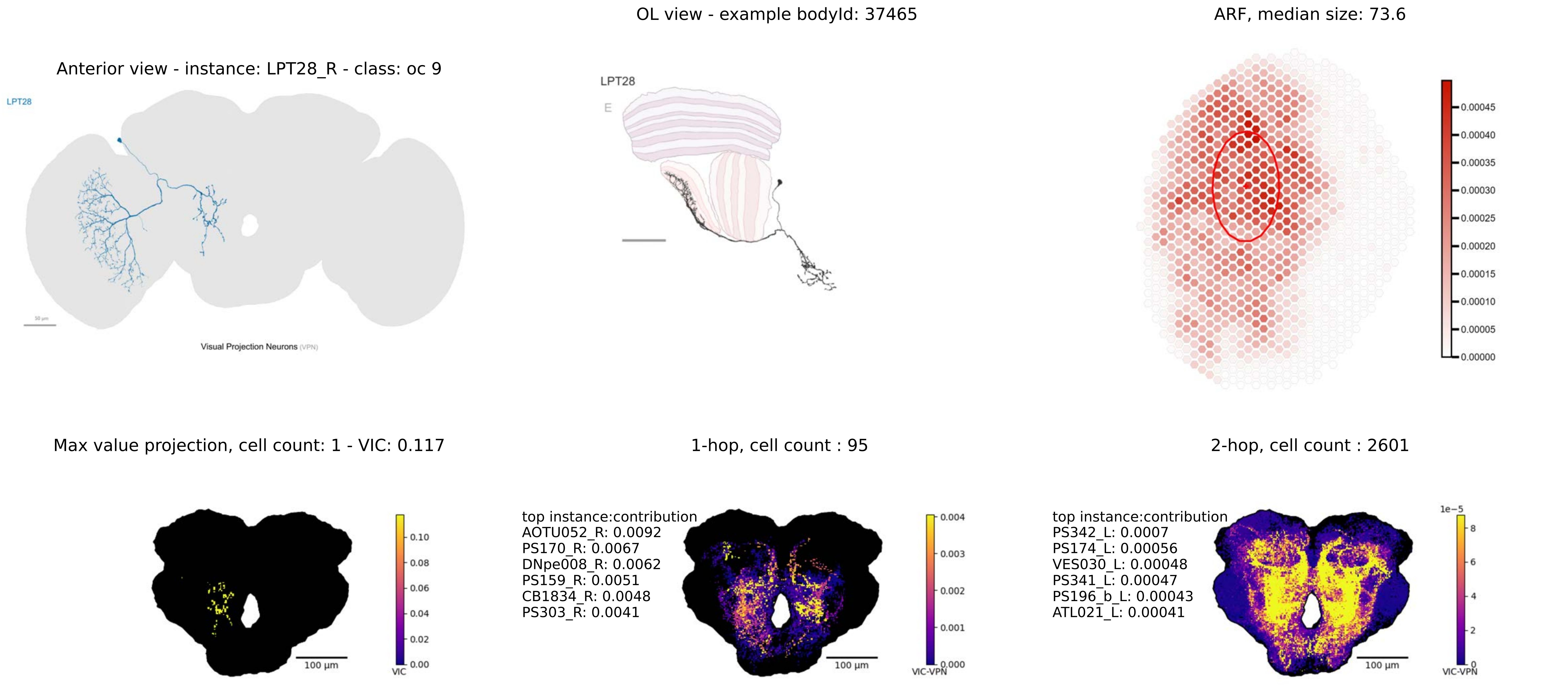

Anterior view - instance: LPT28_R - class: oc 9
LPT28
Visual Projection Neurons (VPN)
OL view - example bodyId: 37465
LPT28
E
ARF, median size: 73.6
0.00045
0.00040
0.00035
0.00030
0.00025
0.00020
0.00015
0.00010
0.00005
0.00000
Max value projection, cell count: 1 - VIC: 0.117
0.10
0.08
0.06
0.04
0.02
0.00
VIC
100 μm
1-hop, cell count : 95
top instance:contribution
AOTU052_R: 0.0092
PS170_R: 0.0067
DNpe008_R: 0.0062
PS159_R: 0.0051
CB1834_R: 0.0048
PS303_R: 0.0041
0.004
0.003
0.002
0.001
0.000
VIC-VPN
100 μm
2-hop, cell count : 2601
top instance:contribution
PS342_L: 0.0007
PS174_L: 0.00056
VES030_L: 0.00048
PS341_L: 0.00047
PS196_b_L: 0.00043
ATL021_L: 0.00041
1e−5
8
6
4
2
0
VIC-VPN
100 μm

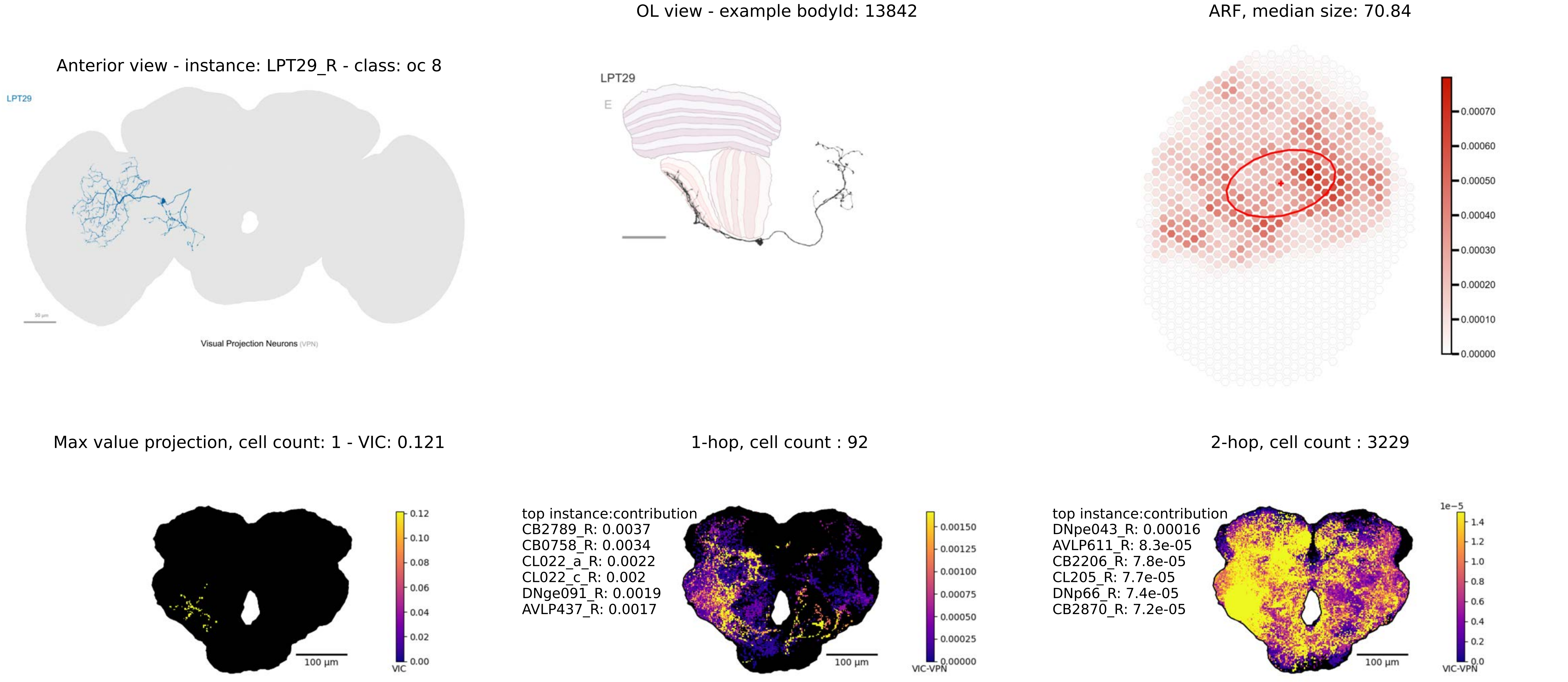
OL view - example bodyId: 13842
ARF, median size: 70.84
Anterior view - instance: LPT29_R - class: oc 8
LPT29
LPT29
E
Visual Projection Neurons (VPN)
0.00070
0.00060
0.00050
0.00040
0.00030
0.00020
0.00010
0.00000
Max value projection, cell count: 1 - VIC: 0.121
1-hop, cell count : 92
2-hop, cell count : 3229
top instance:contribution
CB2789_R: 0.0037
CB0758_R: 0.0034
CL022_a_R: 0.0022
CL022_c_R: 0.002
DNge091_R: 0.0019
AVLP437_R: 0.0017
top instance:contribution
DNpe043_R: 0.00016
AVLP611_R: 8.3e-05
CB2206_R: 7.8e-05
CL205_R: 7.7e-05
DNp66_R: 7.4e-05
CB2870_R: 7.2e-05
100 µm
100 µm
100 µm
VIC
VIC-VPN
VIC-VPN
1e−5

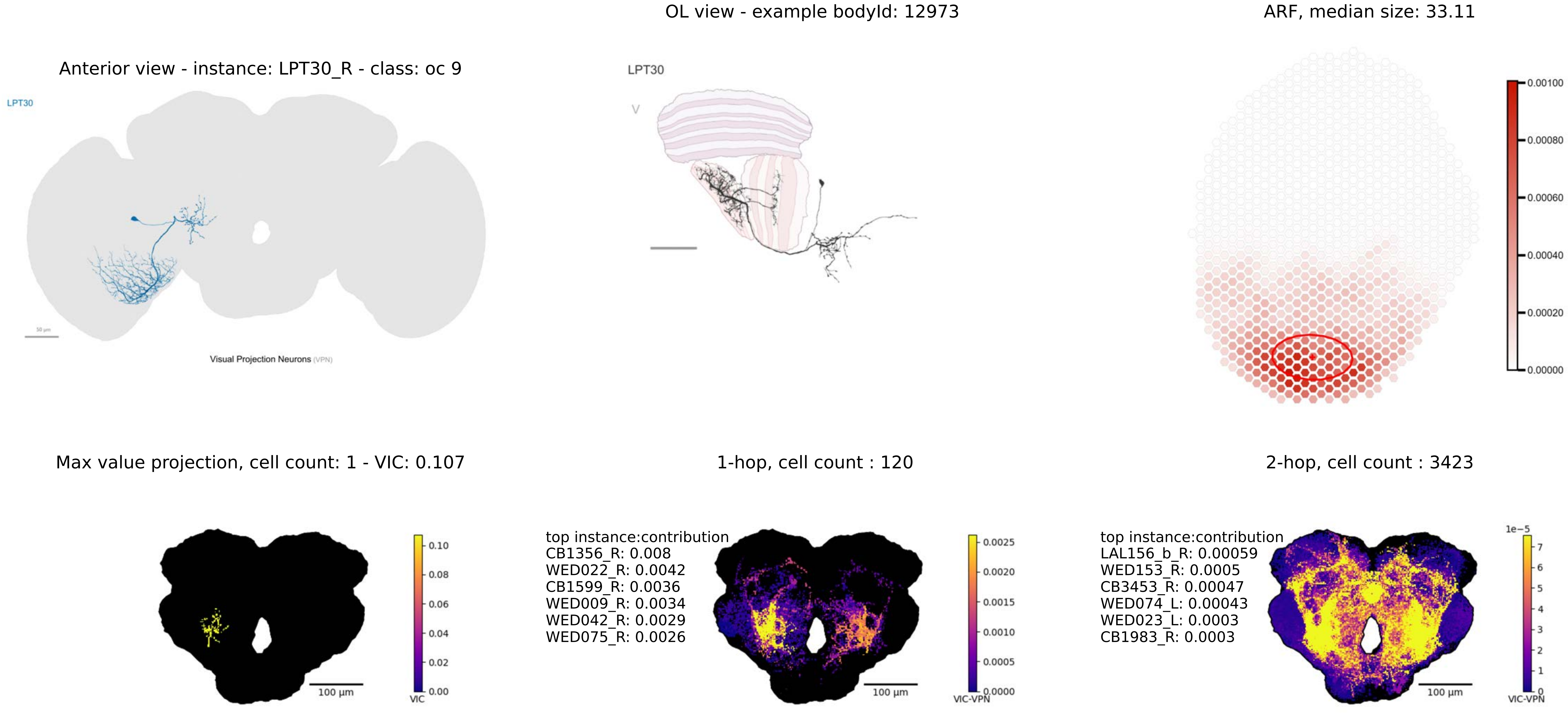
OL view - example bodyId: 12973
ARF, median size: 33.11
Anterior view - instance: LPT30_R - class: oc 9
LPT30
Visual Projection Neurons (VPN)
LPT30
V
0.00100
0.00080
0.00060
0.00040
0.00020
0.00000
Max value projection, cell count: 1 - VIC: 0.107
1-hop, cell count : 120
2-hop, cell count : 3423
0.10
0.08
0.06
0.04
0.02
0.00
VIC
100 µm
top instance:contribution
CB1356_R: 0.008
WED022_R: 0.0042
CB1599_R: 0.0036
WED009_R: 0.0034
WED042_R: 0.0029
WED075_R: 0.0026
0.0025
0.0020
0.0015
0.0010
0.0005
0.0000
VIC-VPN
100 µm
top instance:contribution
LAL156_b_R: 0.00059
WED153_R: 0.0005
CB3453_R: 0.00047
WED074_L: 0.00043
WED023_L: 0.0003
CB1983_R: 0.0003
1e−5
7
6
5
4
3
2
1
0
VIC-VPN
100 µm

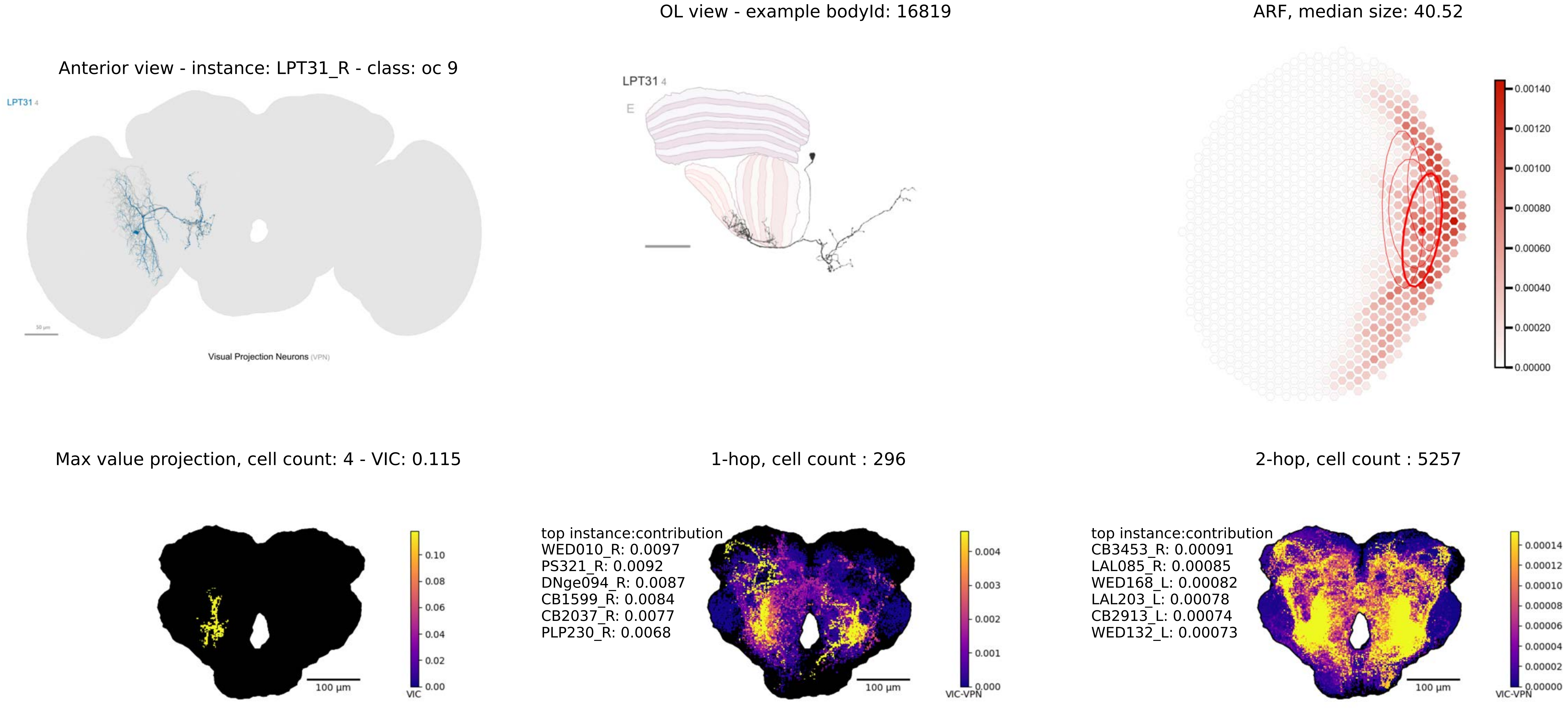

OL view - example bodyId: 16819
ARF, median size: 40.52
Anterior view - instance: LPT31_R - class: oc 9
LPT31 4
LPT31 4
E
0.00140
0.00120
0.00100
0.00080
0.00060
0.00040
0.00020
0.00000
50 µm
Visual Projection Neurons (VPN)
Max value projection, cell count: 4 - VIC: 0.115
1-hop, cell count : 296
2-hop, cell count : 5257
top instance:contribution
WED010_R: 0.0097
PS321_R: 0.0092
DNge094_R: 0.0087
CB1599_R: 0.0084
CB2037_R: 0.0077
PLP230_R: 0.0068
top instance:contribution
CB3453_R: 0.00091
LAL085_R: 0.00085
WED168_L: 0.00082
LAL203_L: 0.00078
CB2913_L: 0.00074
WED132_L: 0.00073
0.10
0.08
0.06
0.04
0.02
0.00
VIC
100 µm
0.004
0.003
0.002
0.001
0.000
VIC-VPN
100 µm
0.00014
0.00012
0.00010
0.00008
0.00006
0.00004
0.00002
0.00000
VIC-VPN
100 µm

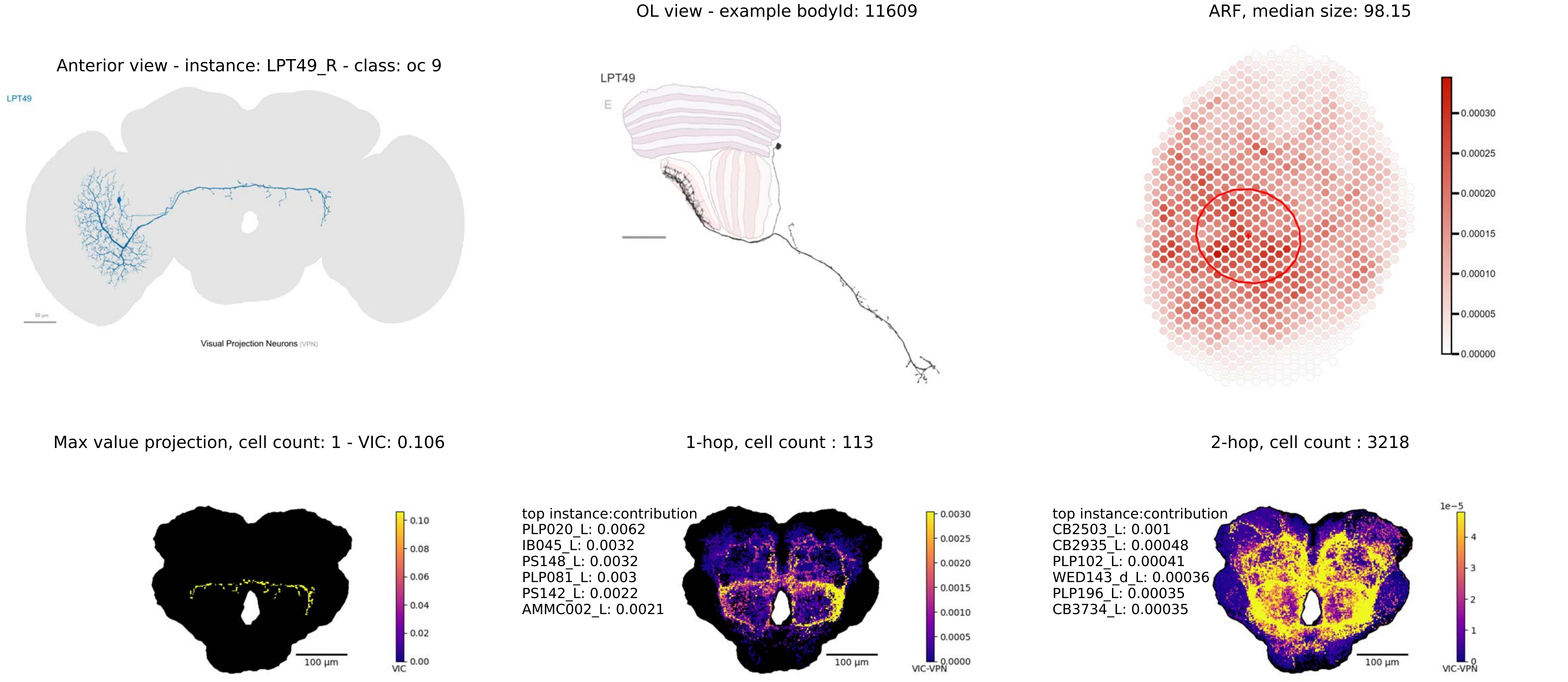

OL view - example bodyId: 11609
ARF, median size: 98.15
Anterior view - instance: LPT49_R - class: oc 9
LPT49
Visual Projection Neurons (VPN)
LPT49
E
0.00030
0.00025
0.00020
0.00015
0.00010
0.00005
0.00000
Max value projection, cell count: 1 - VIC: 0.106
1-hop, cell count : 113
2-hop, cell count : 3218
100 µm
VIC
0.10
0.08
0.06
0.04
0.02
0.00
top instance:contribution
PLP020_L: 0.0062
IB045_L: 0.0032
PS148_L: 0.0032
PLP081_L: 0.003
PS142_L: 0.0022
AMMC002_L: 0.0021
100 µm
VIC-VPN
0.0030
0.0025
0.0020
0.0015
0.0010
0.0005
0.0000
top instance:contribution
CB2503_L: 0.001
CB2935_L: 0.00048
PLP102_L: 0.00041
WED143_d_L: 0.00036
PLP196_L: 0.00035
CB3734_L: 0.00035
100 µm
1e−5
4
3
2
1
0
VIC-VPN

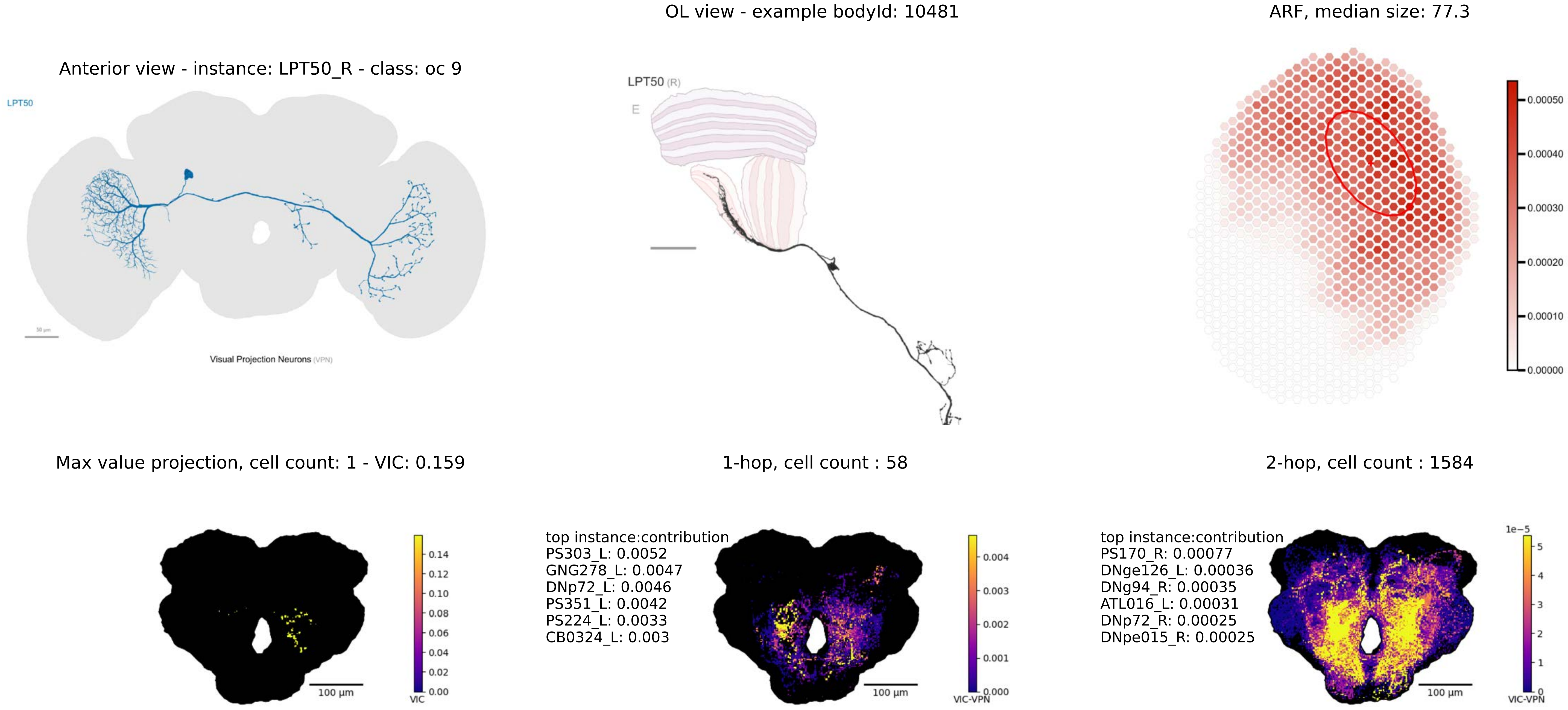
OL view - example bodyId: 10481
ARF, median size: 77.3
Anterior view - instance: LPT50_R - class: oc 9
LPT50
Visual Projection Neurons (VPN)
LPT50 (R)
E
0.00050
0.00040
0.00030
0.00020
0.00010
0.00000
Max value projection, cell count: 1 - VIC: 0.159
1-hop, cell count : 58
2-hop, cell count : 1584
100 µm
VIC
0.14
0.12
0.10
0.08
0.06
0.04
0.02
0.00
top instance:contribution
PS303_L: 0.0052
GNG278_L: 0.0047
DNp72_L: 0.0046
PS351_L: 0.0042
PS224_L: 0.0033
CB0324_L: 0.003
0.004
0.003
0.002
0.001
0.000
VIC-VPN
top instance:contribution
PS170_R: 0.00077
DNge126_L: 0.00036
DNg94_R: 0.00035
ATL016_L: 0.00031
DNp72_R: 0.00025
DNpe015_R: 0.00025
1e−5
5
4
3
2
1
0
VIC-VPN

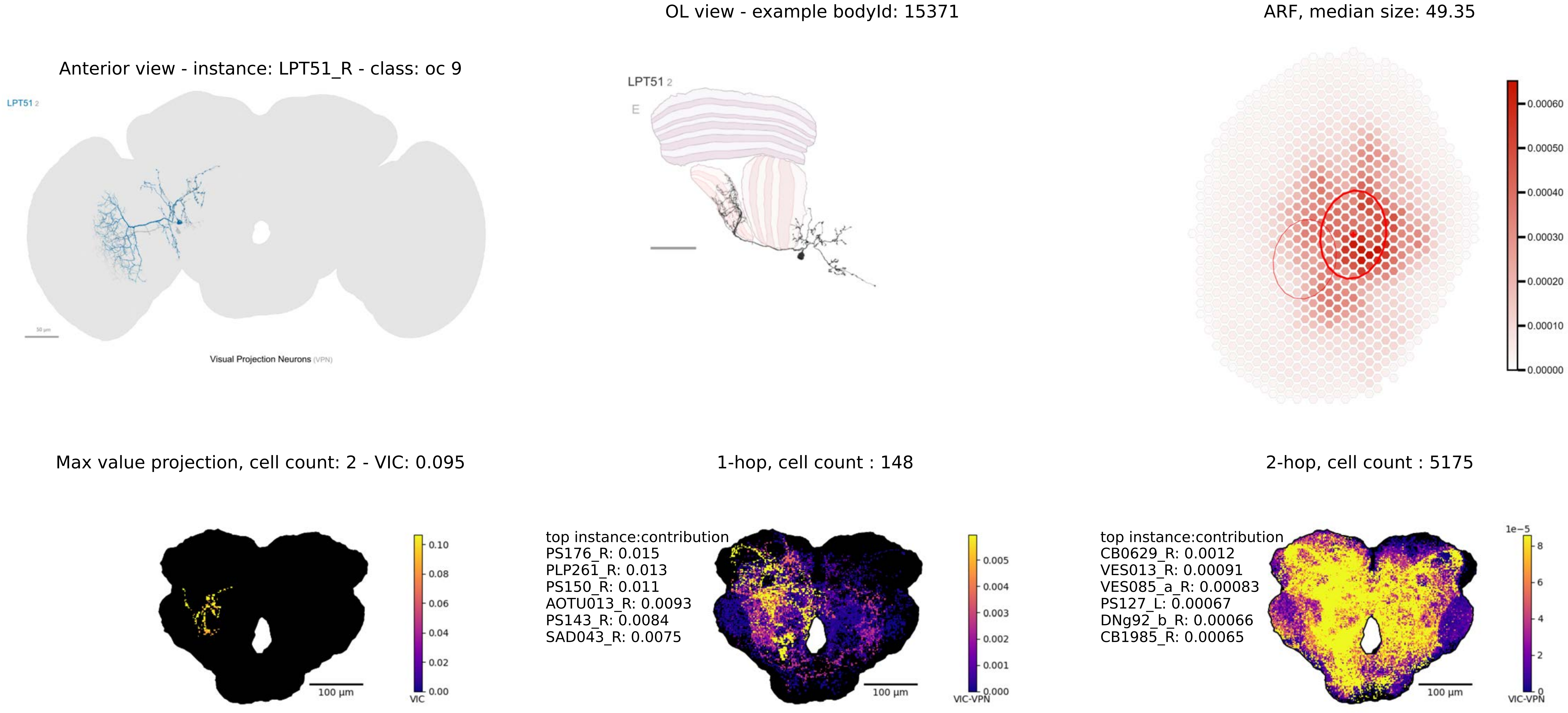
OL view - example bodyId: 15371
ARF, median size: 49.35
Anterior view - instance: LPT51_R - class: oc 9
LPT51 2
Visual Projection Neurons (VPN)
LPT51 2
E
0.00060
0.00050
0.00040
0.00030
0.00020
0.00010
0.00000
Max value projection, cell count: 2 - VIC: 0.095
1-hop, cell count : 148
2-hop, cell count : 5175
top instance:contribution
PS176_R: 0.015
PLP261_R: 0.013
PS150_R: 0.011
AOTU013_R: 0.0093
PS143_R: 0.0084
SAD043_R: 0.0075
top instance:contribution
CB0629_R: 0.0012
VES013_R: 0.00091
VES085_a_R: 0.00083
PS127_L: 0.00067
DNg92_b_R: 0.00066
CB1985_R: 0.00065
0.10
0.08
0.06
0.04
0.02
0.00
VIC
100 µm
0.005
0.004
0.003
0.002
0.001
0.000
VIC-VPN
100 µm
1e−5
8
6
4
2
0
VIC-VPN
100 µm

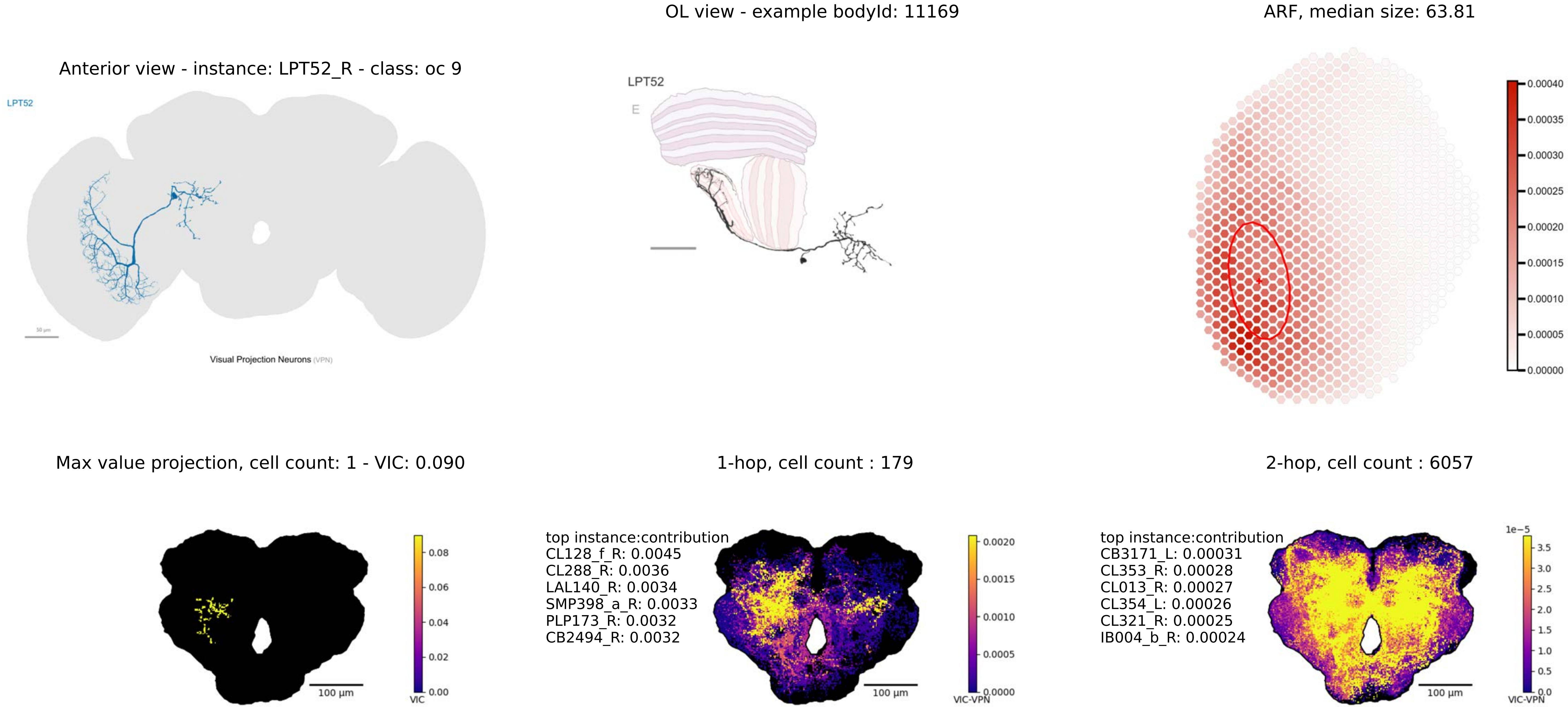
OL view - example bodyId: 11169
ARF, median size: 63.81
Anterior view - instance: LPT52_R - class: oc 9
LPT52
Visual Projection Neurons (VPN)
LPT52
E
0.00040
0.00035
0.00030
0.00025
0.00020
0.00015
0.00010
0.00005
0.00000
Max value projection, cell count: 1 - VIC: 0.090
1-hop, cell count : 179
2-hop, cell count : 6057
0.08
0.06
0.04
0.02
0.00
VIC
100 µm
top instance:contribution
CL128_f_R: 0.0045
CL288_R: 0.0036
LAL140_R: 0.0034
SMP398_a_R: 0.0033
PLP173_R: 0.0032
CB2494_R: 0.0032
0.0020
0.0015
0.0010
0.0005
0.0000
VIC-VPN
100 µm
top instance:contribution
CB3171_L: 0.00031
CL353_R: 0.00028
CL013_R: 0.00027
CL354_L: 0.00026
CL321_R: 0.00025
IB004_b_R: 0.00024
1e−5
3.5
3.0
2.5
2.0
1.5
1.0
0.5
0.0
VIC-VPN
100 µm

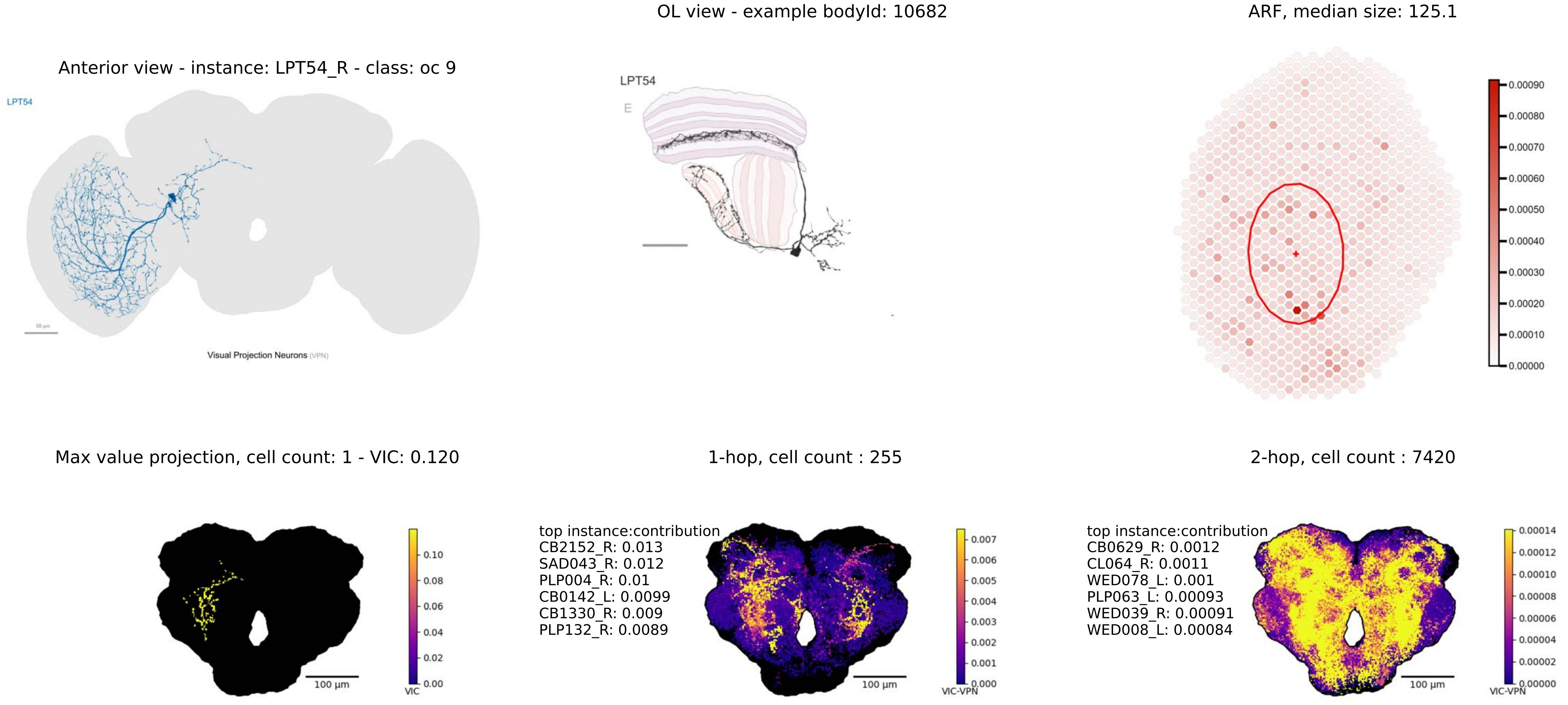
Anterior view - instance: LPT54_R - class: oc 9
LPT54
Visual Projection Neurons (VPN)
OL view - example bodyId: 10682
LPT54
E
ARF, median size: 125.1
0.00090
0.00080
0.00070
0.00060
0.00050
0.00040
0.00030
0.00020
0.00010
0.00000
Max value projection, cell count: 1 - VIC: 0.120
100 µm
VIC
0.10
0.08
0.06
0.04
0.02
0.00
1-hop, cell count : 255
top instance:contribution
CB2152_R: 0.013
SAD043_R: 0.012
PLP004_R: 0.01
CB0142_L: 0.0099
CB1330_R: 0.009
PLP132_R: 0.0089
100 µm
VIC-VPN
0.007
0.006
0.005
0.004
0.003
0.002
0.001
0.000
2-hop, cell count : 7420
top instance:contribution
CB0629_R: 0.0012
CL064_R: 0.0011
WED078_L: 0.001
PLP063_L: 0.00093
WED039_R: 0.00091
WED008_L: 0.00084
100 µm
VIC-VPN
0.00014
0.00012
0.00010
0.00008
0.00006
0.00004
0.00002
0.00000

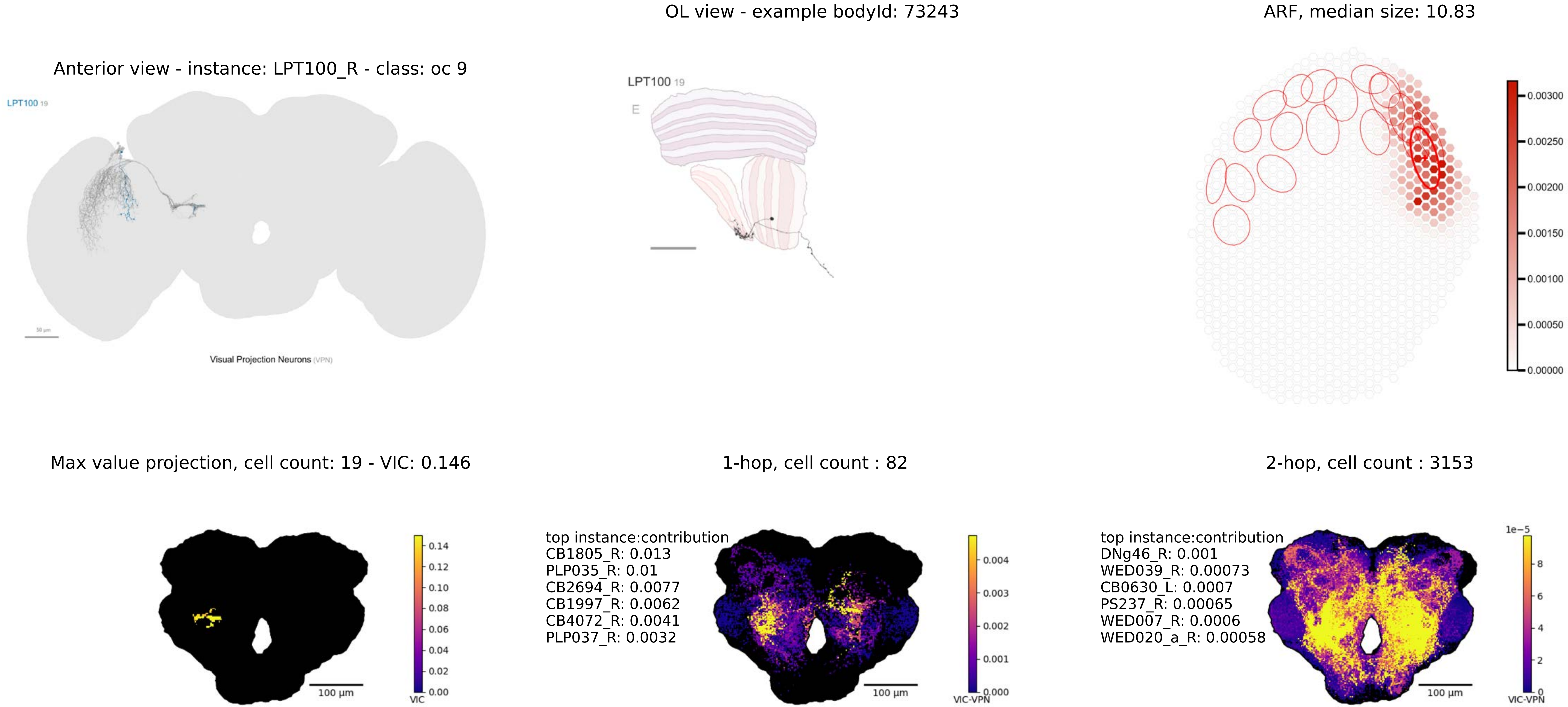
OL view - example bodyId: 73243
ARF, median size: 10.83
Anterior view - instance: LPT100_R - class: oc 9
LPT100 19
LPT100 19
E
0.00300
0.00250
0.00200
0.00150
0.00100
0.00050
0.00000
Visual Projection Neurons (VPN)
Max value projection, cell count: 19 - VIC: 0.146
1-hop, cell count : 82
2-hop, cell count : 3153
top instance:contribution
CB1805_R: 0.013
PLP035_R: 0.01
CB2694_R: 0.0077
CB1997_R: 0.0062
CB4072_R: 0.0041
PLP037_R: 0.0032
top instance:contribution
DNg46_R: 0.001
WED039_R: 0.00073
CB0630_L: 0.0007
PS237_R: 0.00065
WED007_R: 0.0006
WED020_a_R: 0.00058
0.14
0.12
0.10
0.08
0.06
0.04
0.02
0.00
VIC
0.004
0.003
0.002
0.001
0.000
VIC-VPN
1e−5
8
6
4
2
0
VIC-VPN
100 µm
100 µm
100 µm

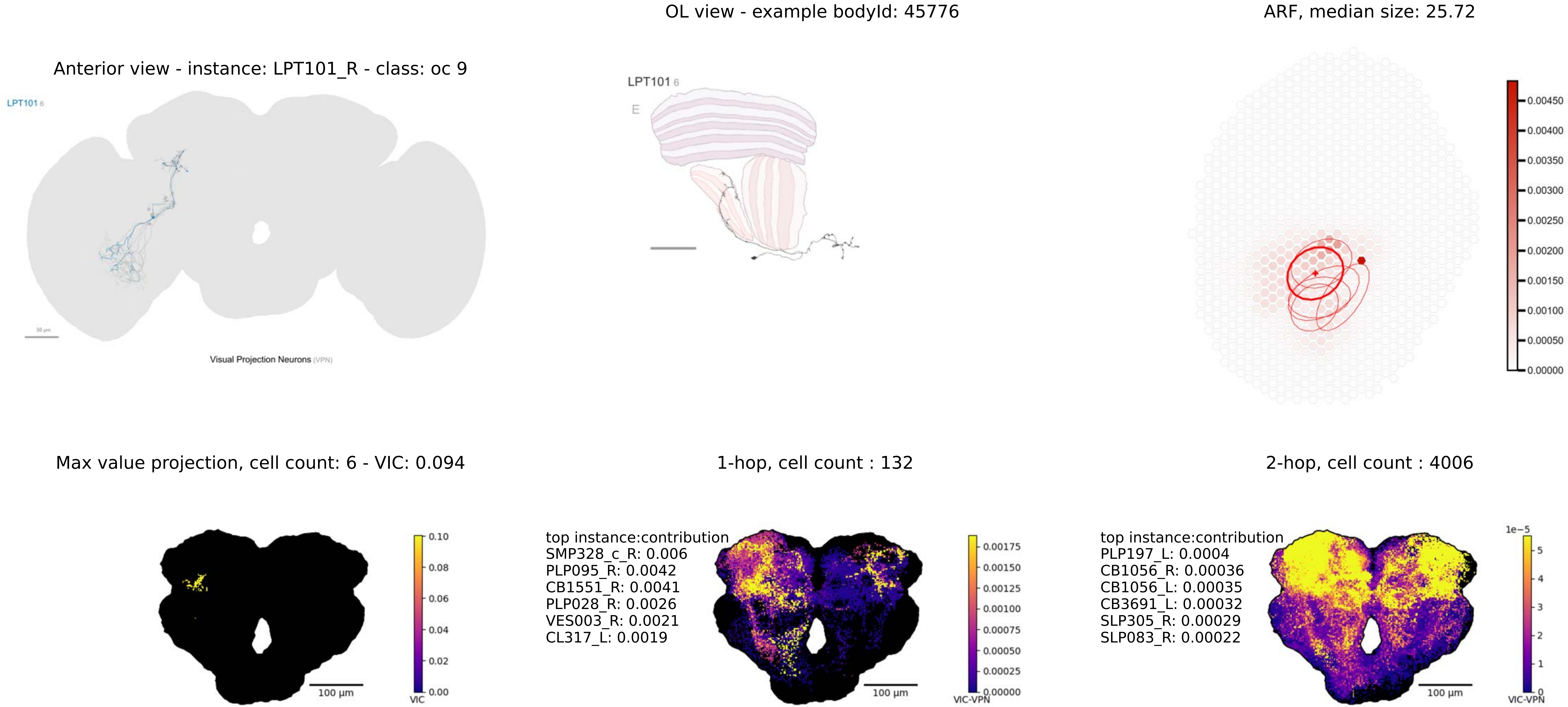

OL view - example bodyId: 45776
ARF, median size: 25.72
Anterior view - instance: LPT101_R - class: oc 9
LPT101 6
Visual Projection Neurons (VPN)
LPT101 6
E
0.00450
0.00400
0.00350
0.00300
0.00250
0.00200
0.00150
0.00100
0.00050
0.00000
Max value projection, cell count: 6 - VIC: 0.094
1-hop, cell count : 132
2-hop, cell count : 4006
0.10
0.08
0.06
0.04
0.02
0.00
VIC
100 µm
top instance:contribution
SMP328_c_R: 0.006
PLP095_R: 0.0042
CB1551_R: 0.0041
PLP028_R: 0.0026
VES003_R: 0.0021
CL317_L: 0.0019
0.00175
0.00150
0.00125
0.00100
0.00075
0.00050
0.00025
0.00000
VIC-VPN
100 µm
top instance:contribution
PLP197_L: 0.0004
CB1056_R: 0.00036
CB1056_L: 0.00035
CB3691_L: 0.00032
SLP305_R: 0.00029
SLP083_R: 0.00022
1e−5
5
4
3
2
1
0
VIC-VPN
100 µm

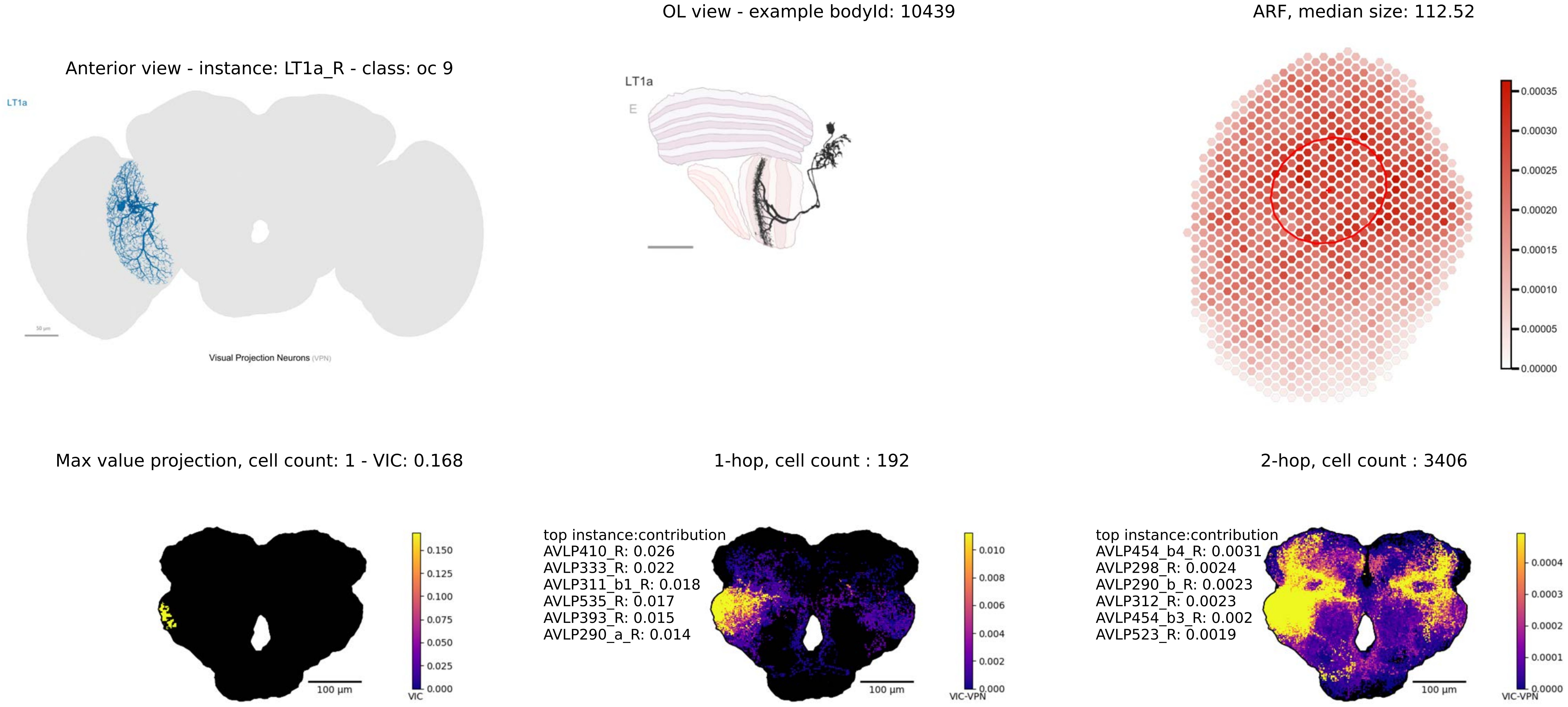

Anterior view - instance: LT1a_R - class: oc 9
LT1a
Visual Projection Neurons (VPN)
OL view - example bodyId: 10439
LT1a
E
ARF, median size: 112.52
0.00035
0.00030
0.00025
0.00020
0.00015
0.00010
0.00005
0.00000
Max value projection, cell count: 1 - VIC: 0.168
100 µm
VIC
0.150
0.125
0.100
0.075
0.050
0.025
0.000
1-hop, cell count : 192
top instance:contribution
AVLP410_R: 0.026
AVLP333_R: 0.022
AVLP311_b1_R: 0.018
AVLP535_R: 0.017
AVLP393_R: 0.015
AVLP290_a_R: 0.014
100 µm
VIC-VPN
0.010
0.008
0.006
0.004
0.002
0.000
2-hop, cell count : 3406
top instance:contribution
AVLP454_b4_R: 0.0031
AVLP298_R: 0.0024
AVLP290_b_R: 0.0023
AVLP312_R: 0.0023
AVLP454_b3_R: 0.002
AVLP523_R: 0.0019
100 µm
VIC-VPN
0.0004
0.0003
0.0002
0.0001
0.0000

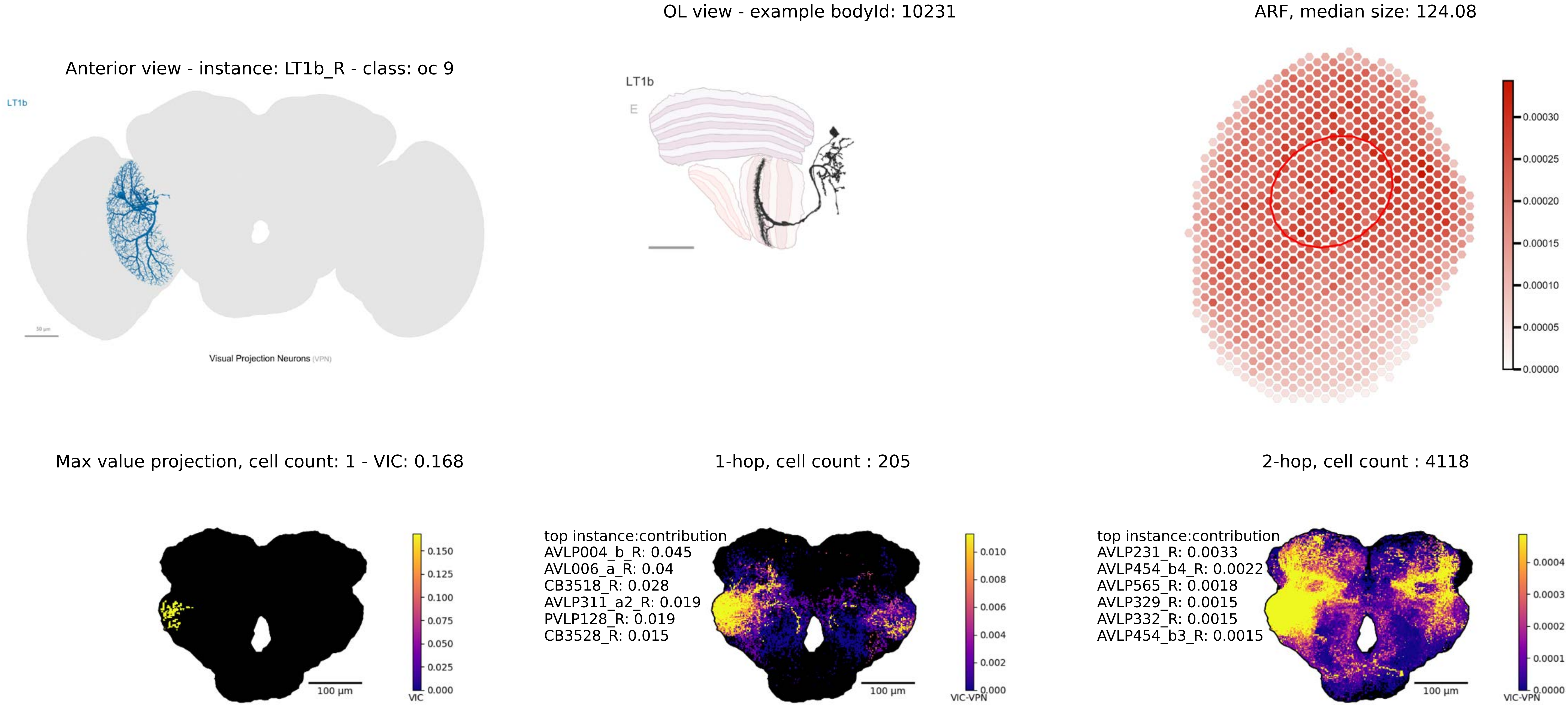
Anterior view - instance: LT1b_R - class: oc 9
LT1b
50 µm
Visual Projection Neurons (VPN)
OL view - example bodyId: 10231
LT1b
E
ARF, median size: 124.08
0.00030
0.00025
0.00020
0.00015
0.00010
0.00005
0.00000
Max value projection, cell count: 1 - VIC: 0.168
100 µm
0.150
0.125
0.100
0.075
0.050
0.025
0.000
VIC
1-hop, cell count : 205
top instance:contribution
AVLP004_b_R: 0.045
AVL006_a_R: 0.04
CB3518_R: 0.028
AVLP311_a2_R: 0.019
PVLP128_R: 0.019
CB3528_R: 0.015
100 µm
0.010
0.008
0.006
0.004
0.002
0.000
VIC-VPN
2-hop, cell count : 4118
top instance:contribution
AVLP231_R: 0.0033
AVLP454_b4_R: 0.0022
AVLP565_R: 0.0018
AVLP329_R: 0.0015
AVLP332_R: 0.0015
AVLP454_b3_R: 0.0015
100 µm
0.0004
0.0003
0.0002
0.0001
0.0000
VIC-VPN

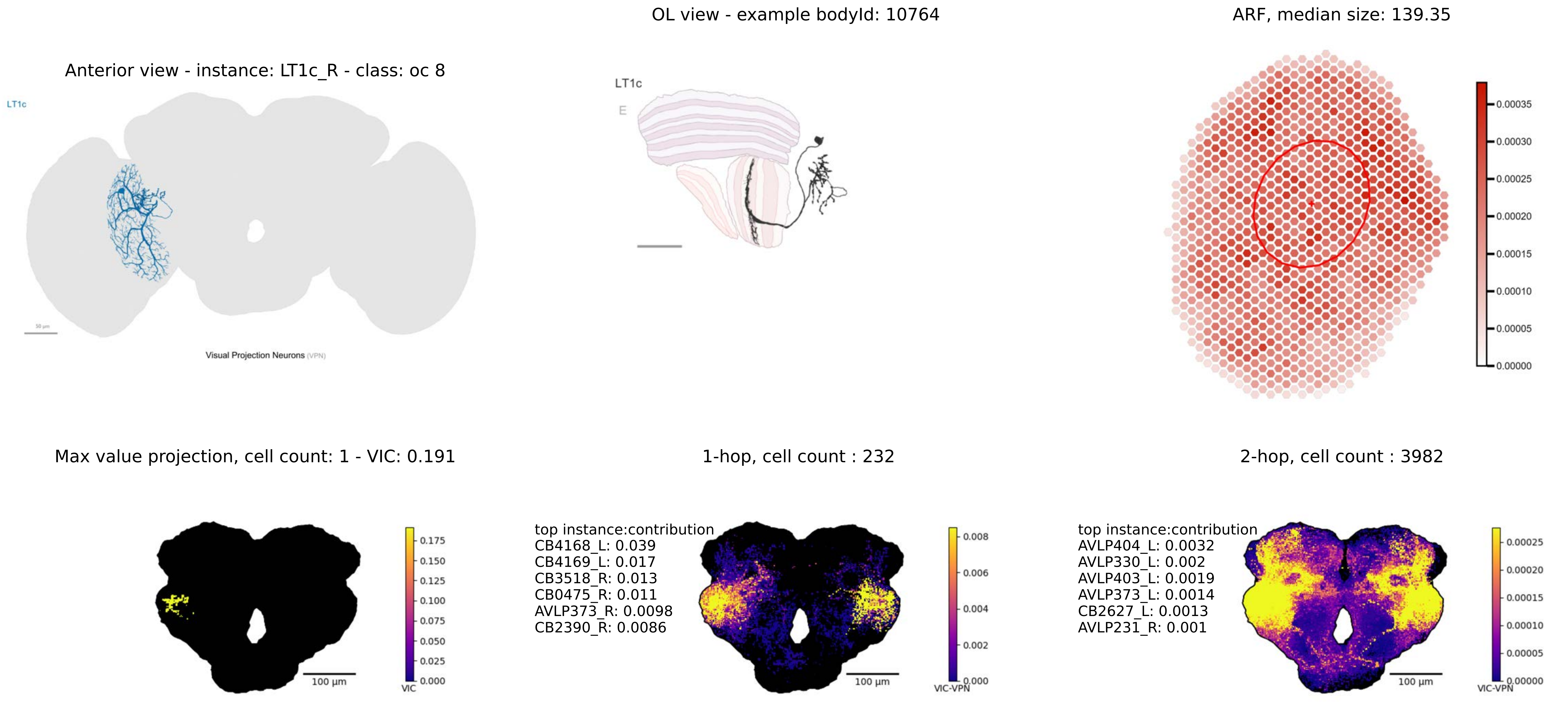
Anterior view - instance: LT1c_R - class: oc 8
LT1c
Visual Projection Neurons (VPN)
OL view - example bodyId: 10764
LT1c
E
ARF, median size: 139.35
0.00035
0.00030
0.00025
0.00020
0.00015
0.00010
0.00005
0.00000
Max value projection, cell count: 1 - VIC: 0.191
100 µm
VIC
0.175
0.150
0.125
0.100
0.075
0.050
0.025
0.000
1-hop, cell count : 232
top instance:contribution
CB4168_L: 0.039
CB4169_L: 0.017
CB3518_R: 0.013
CB0475_R: 0.011
AVLP373_R: 0.0098
CB2390_R: 0.0086
100 µm
VIC-VPN
0.008
0.006
0.004
0.002
0.000
2-hop, cell count : 3982
top instance:contribution
AVLP404_L: 0.0032
AVLP330_L: 0.002
AVLP403_L: 0.0019
AVLP373_L: 0.0014
CB2627_L: 0.0013
AVLP231_R: 0.001
100 µm
VIC-VPN
0.00025
0.00020
0.00015
0.00010
0.00005
0.00000

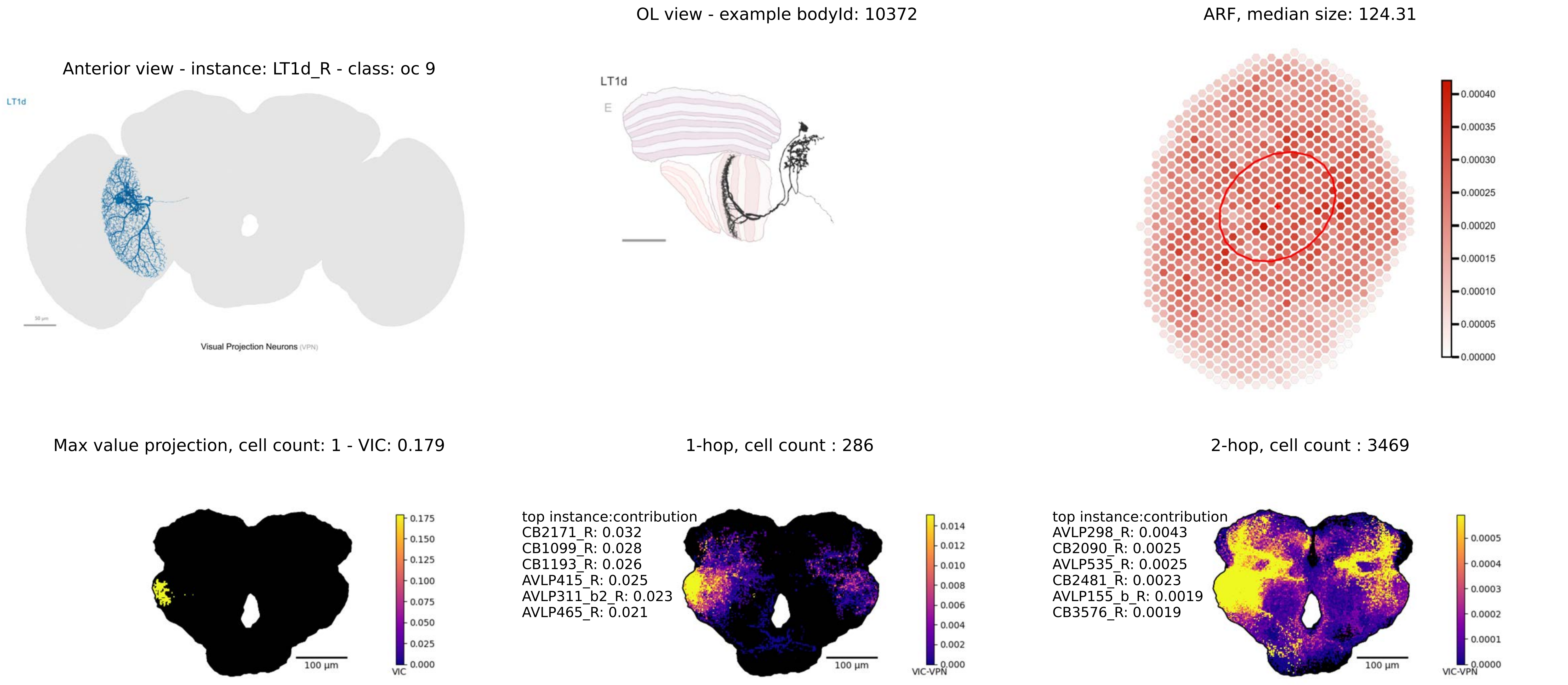
OL view - example bodyId: 10372
ARF, median size: 124.31
Anterior view - instance: LT1d_R - class: oc 9
LT1d
Visual Projection Neurons (VPN)
LT1d
E
0.00040
0.00035
0.00030
0.00025
0.00020
0.00015
0.00010
0.00005
0.00000
Max value projection, cell count: 1 - VIC: 0.179
1-hop, cell count : 286
2-hop, cell count : 3469
100 µm
VIC
0.175
0.150
0.125
0.100
0.075
0.050
0.025
0.000
top instance:contribution
CB2171_R: 0.032
CB1099_R: 0.028
CB1193_R: 0.026
AVLP415_R: 0.025
AVLP311_b2_R: 0.023
AVLP465_R: 0.021
100 µm
VIC-VPN
0.014
0.012
0.010
0.008
0.006
0.004
0.002
0.000
top instance:contribution
AVLP298_R: 0.0043
CB2090_R: 0.0025
AVLP535_R: 0.0025
CB2481_R: 0.0023
AVLP155_b_R: 0.0019
CB3576_R: 0.0019
100 µm
VIC-VPN
0.0005
0.0004
0.0003
0.0002
0.0001
0.0000

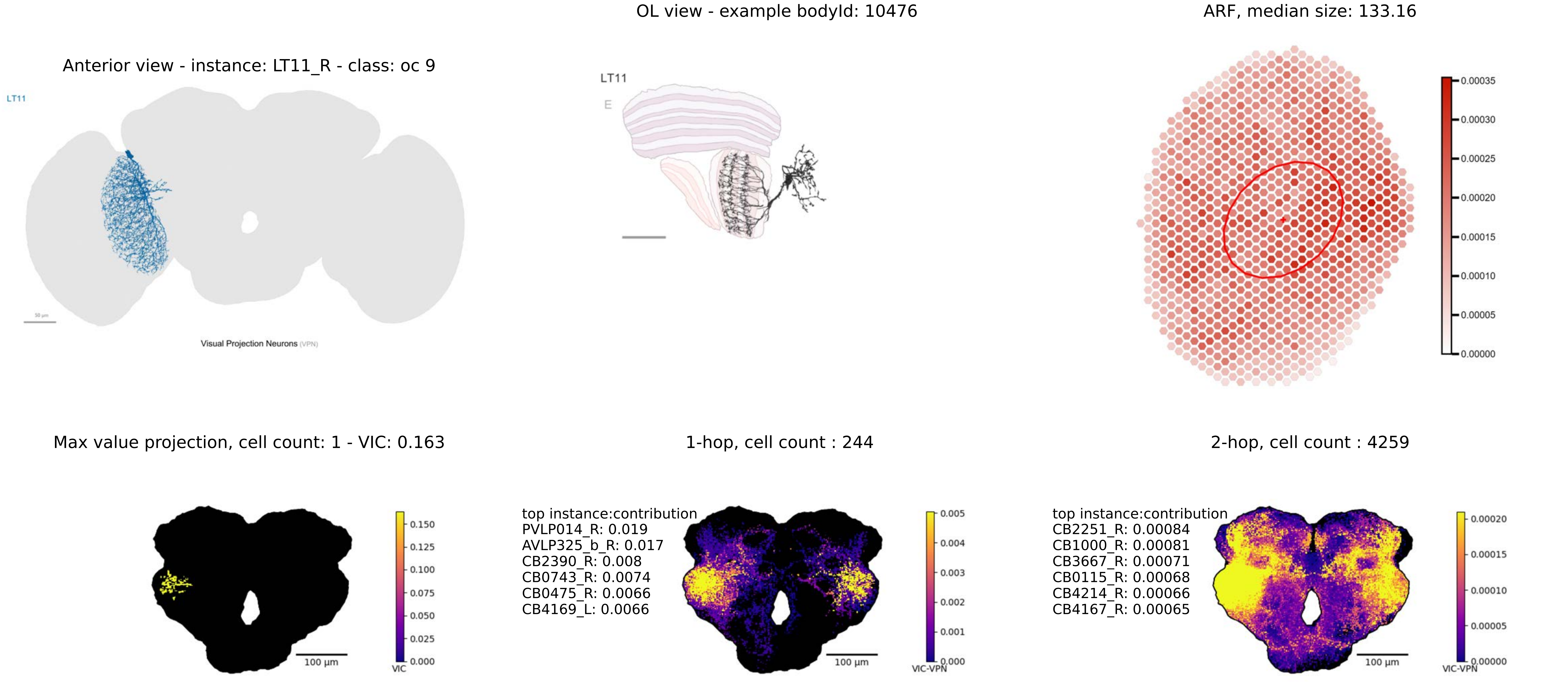
Anterior view - instance: LT11_R - class: oc 9
LT11
Visual Projection Neurons (VPN)
OL view - example bodyId: 10476
LT11
E
ARF, median size: 133.16
0.00035
0.00030
0.00025
0.00020
0.00015
0.00010
0.00005
0.00000
Max value projection, cell count: 1 - VIC: 0.163
100 µm
VIC
1-hop, cell count : 244
top instance:contribution
PVLP014_R: 0.019
AVLP325_b_R: 0.017
CB2390_R: 0.008
CB0743_R: 0.0074
CB0475_R: 0.0066
CB4169_L: 0.0066
100 µm
VIC-VPN
2-hop, cell count : 4259
top instance:contribution
CB2251_R: 0.00084
CB1000_R: 0.00081
CB3667_R: 0.00071
CB0115_R: 0.00068
CB4214_R: 0.00066
CB4167_R: 0.00065
100 µm
VIC-VPN

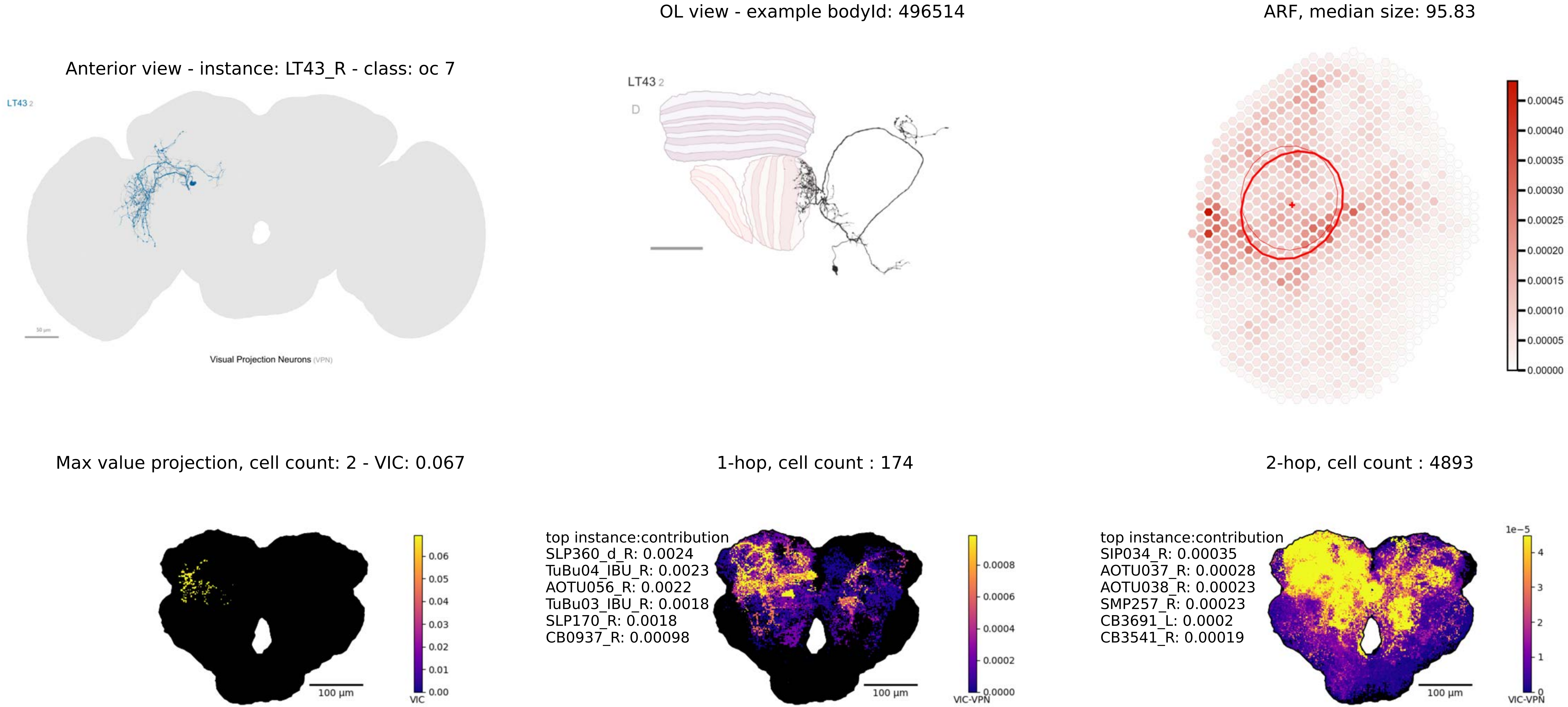

OL view - example bodyId: 496514
ARF, median size: 95.83
Anterior view - instance: LT43_R - class: oc 7
LT43 2
Visual Projection Neurons (VPN)
LT43 2
D
0.00045
0.00040
0.00035
0.00030
0.00025
0.00020
0.00015
0.00010
0.00005
0.00000
Max value projection, cell count: 2 - VIC: 0.067
1-hop, cell count : 174
2-hop, cell count : 4893
0.06
0.05
0.04
0.03
0.02
0.01
0.00
VIC
100 µm
top instance:contribution
SLP360_d_R: 0.0024
TuBu04_IBU_R: 0.0023
AOTU056_R: 0.0022
TuBu03_IBU_R: 0.0018
SLP170_R: 0.0018
CB0937_R: 0.00098
0.0008
0.0006
0.0004
0.0002
0.0000
VIC-VPN
100 µm
top instance:contribution
SIP034_R: 0.00035
AOTU037_R: 0.00028
AOTU038_R: 0.00023
SMP257_R: 0.00023
CB3691_L: 0.0002
CB3541_R: 0.00019
1e−5
4
3
2
1
0
VIC-VPN
100 µm

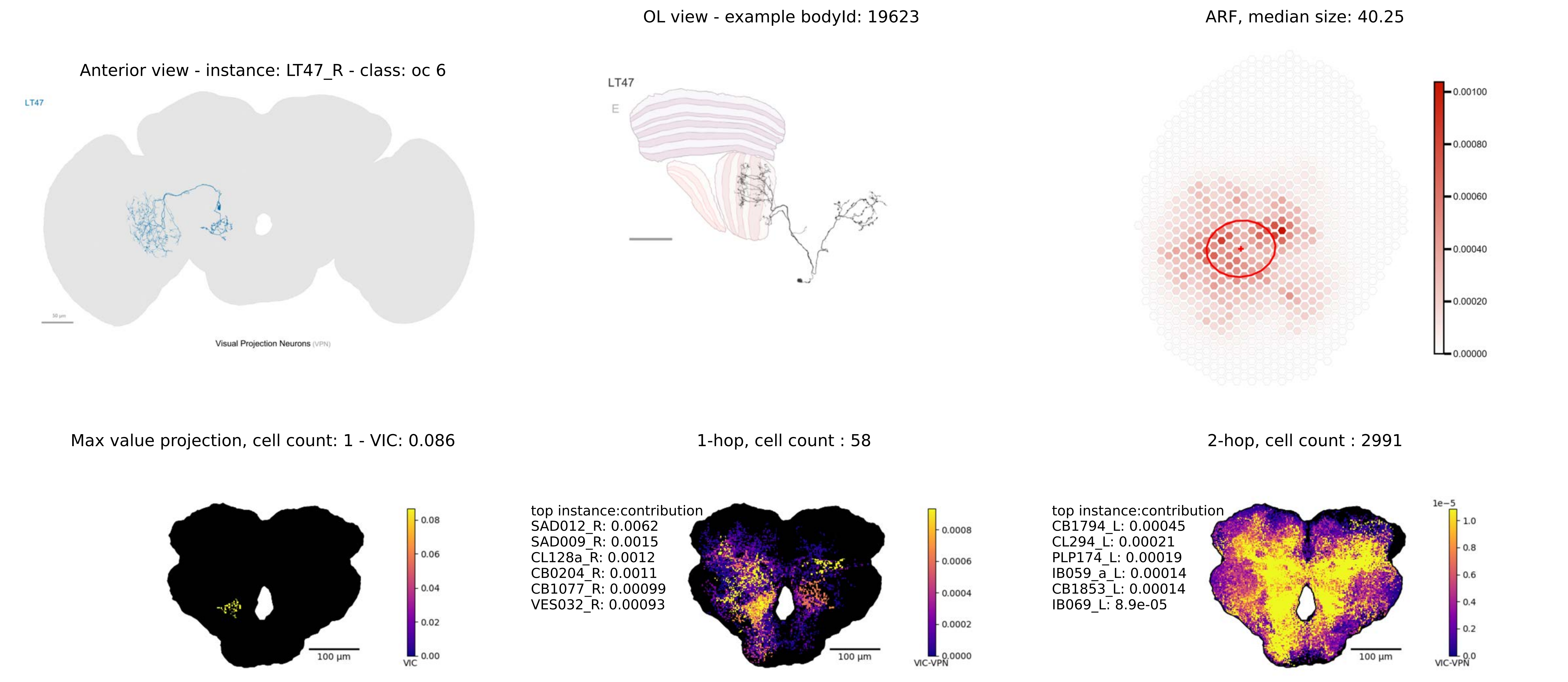
OL view - example bodyId: 19623
ARF, median size: 40.25
Anterior view - instance: LT47_R - class: oc 6
LT47
Visual Projection Neurons (VPN)
LT47
E
0.00100
0.00080
0.00060
0.00040
0.00020
0.00000
Max value projection, cell count: 1 - VIC: 0.086
1-hop, cell count : 58
2-hop, cell count : 2991
0.08
0.06
0.04
0.02
0.00
VIC
100 µm
top instance:contribution
SAD012_R: 0.0062
SAD009_R: 0.0015
CL128a_R: 0.0012
CB0204_R: 0.0011
CB1077_R: 0.00099
VES032_R: 0.00093
0.0008
0.0006
0.0004
0.0002
0.0000
VIC-VPN
100 µm
top instance:contribution
CB1794_L: 0.00045
CL294_L: 0.00021
PLP174_L: 0.00019
IB059_a_L: 0.00014
CB1853_L: 0.00014
IB069_L: 8.9e-05
1e−5
1.0
0.8
0.6
0.4
0.2
0.0
VIC-VPN
100 µm

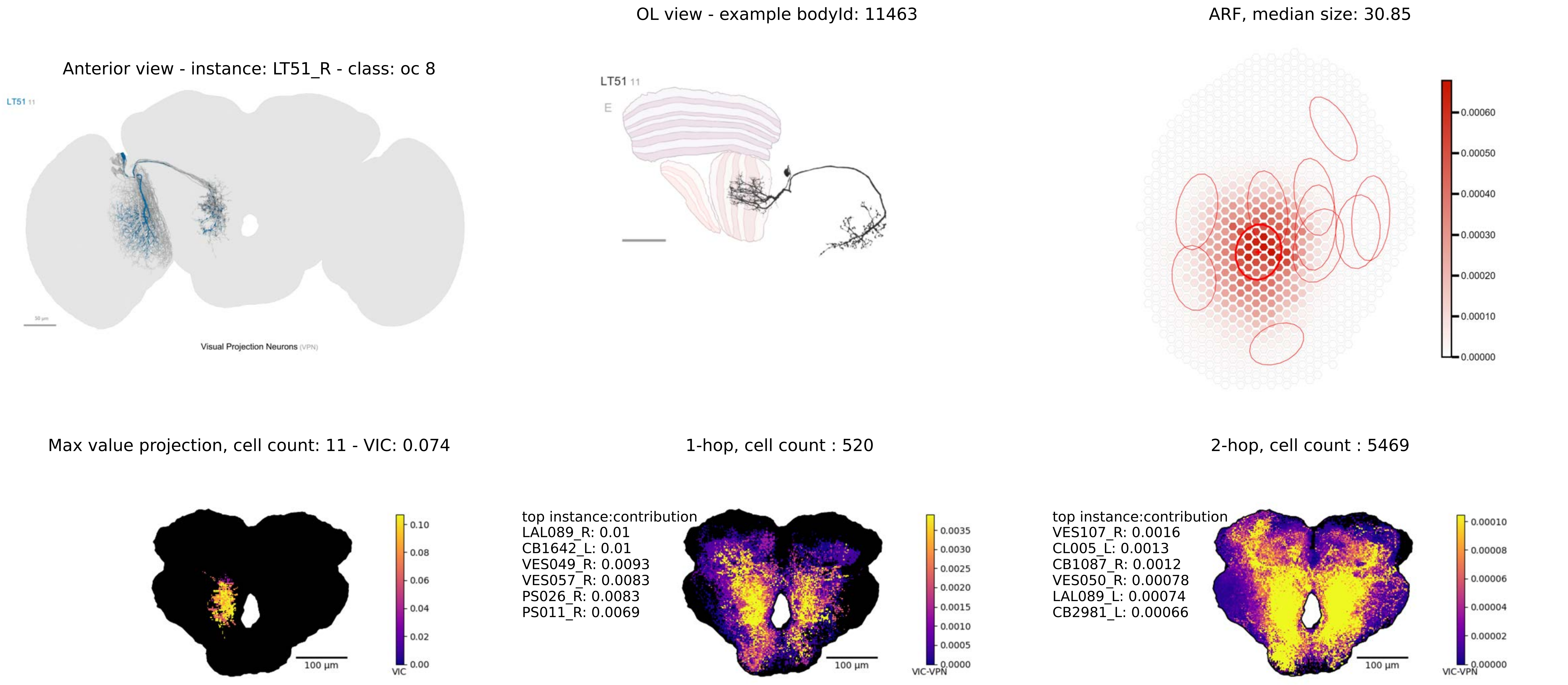

OL view - example bodyId: 11463
ARF, median size: 30.85
Anterior view - instance: LT51_R - class: oc 8
LT51 11
Visual Projection Neurons (VPN)
LT51 11
E
0.00060
0.00050
0.00040
0.00030
0.00020
0.00010
0.00000
Max value projection, cell count: 11 - VIC: 0.074
1-hop, cell count : 520
2-hop, cell count : 5469
0.10
0.08
0.06
0.04
0.02
0.00
VIC
100 µm
top instance:contribution
LAL089_R: 0.01
CB1642_L: 0.01
VES049_R: 0.0093
VES057_R: 0.0083
PS026_R: 0.0083
PS011_R: 0.0069
0.0035
0.0030
0.0025
0.0020
0.0015
0.0010
0.0005
0.0000
VIC-VPN
100 µm
top instance:contribution
VES107_R: 0.0016
CL005_L: 0.0013
CB1087_R: 0.0012
VES050_R: 0.00078
LAL089_L: 0.00074
CB2981_L: 0.00066
0.00010
0.00008
0.00006
0.00004
0.00002
0.00000
VIC-VPN
100 µm

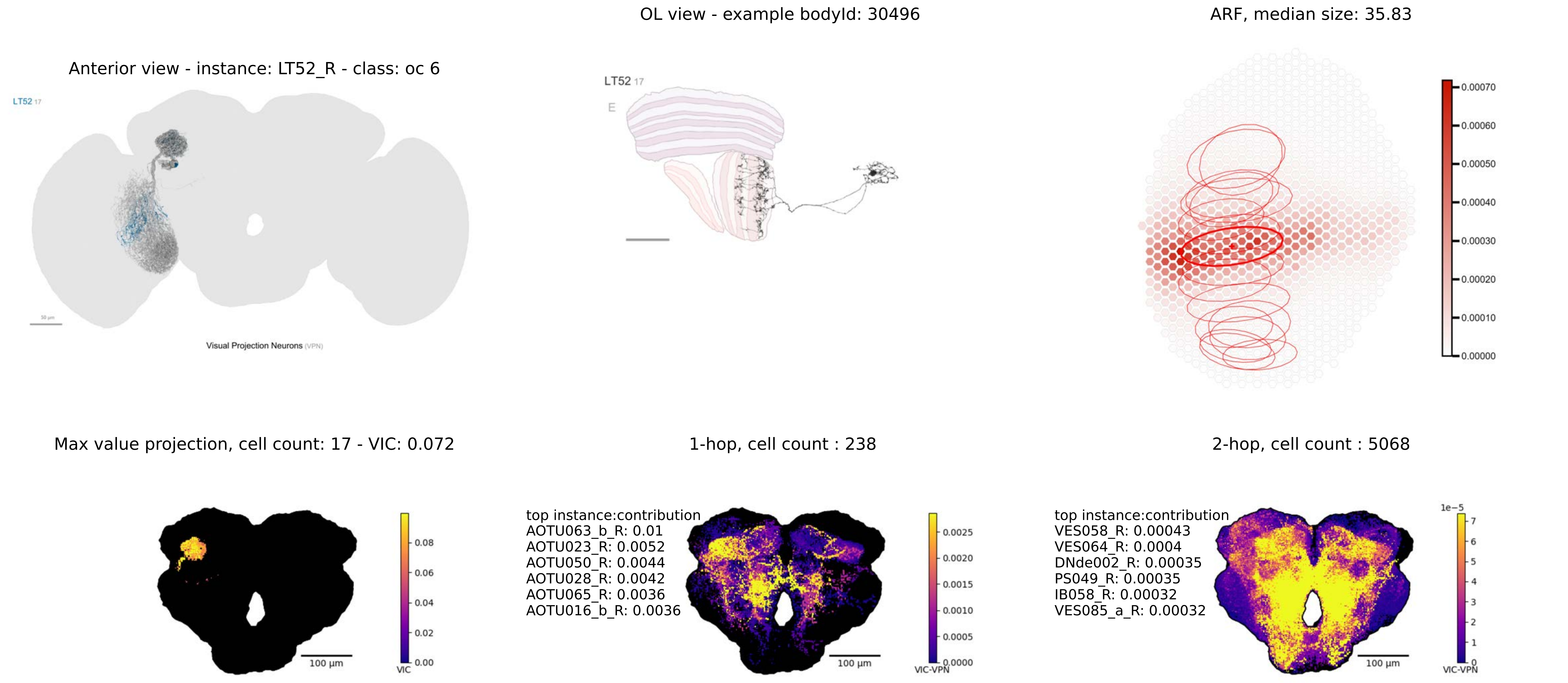

OL view - example bodyId: 30496
ARF, median size: 35.83
Anterior view - instance: LT52_R - class: oc 6
LT52 17
LT52 17
E
Visual Projection Neurons (VPN)
0.00070
0.00060
0.00050
0.00040
0.00030
0.00020
0.00010
0.00000
Max value projection, cell count: 17 - VIC: 0.072
1-hop, cell count : 238
2-hop, cell count : 5068
0.08
0.06
0.04
0.02
0.00
VIC
100 µm
top instance:contribution
AOTU063_b_R: 0.01
AOTU023_R: 0.0052
AOTU050_R: 0.0044
AOTU028_R: 0.0042
AOTU065_R: 0.0036
AOTU016_b_R: 0.0036
100 µm
0.0025
0.0020
0.0015
0.0010
0.0005
0.0000
VIC-VPN
top instance:contribution
VES058_R: 0.00043
VES064_R: 0.0004
DNde002_R: 0.00035
PS049_R: 0.00035
IB058_R: 0.00032
VES085_a_R: 0.00032
100 µm
1e−5
7
6
5
4
3
2
1
0
VIC-VPN

Anterior view - instance: LT54_R - class: oc 8

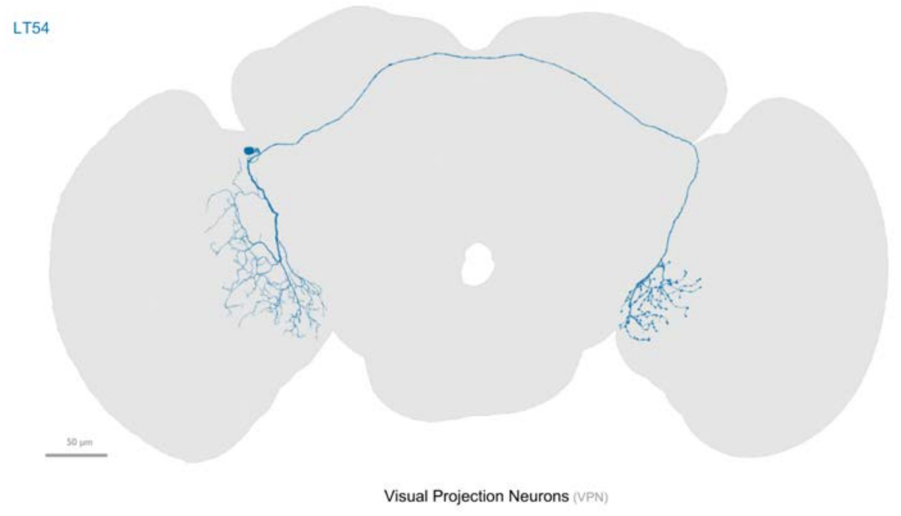


OL view - example bodyId: 18774

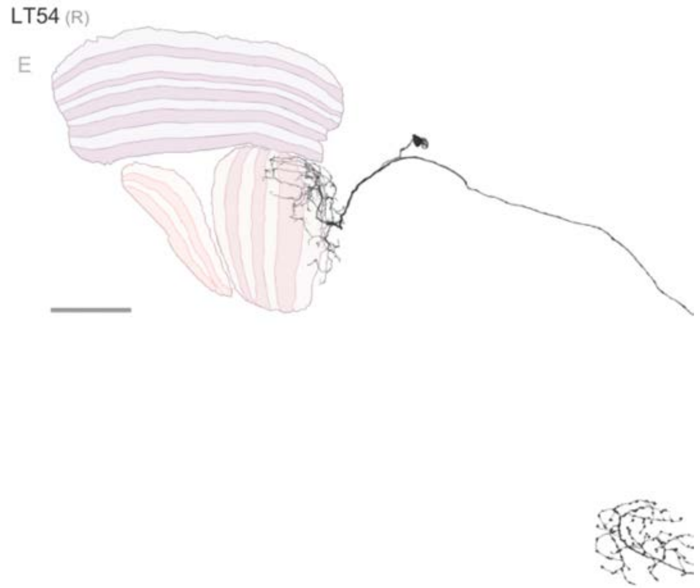


ARF, median size: 56.97

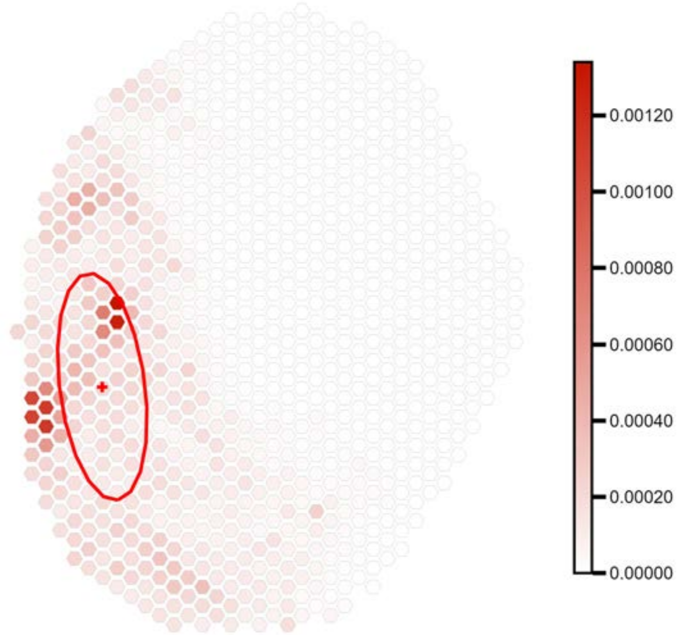


fewer than 100 output connections in central brain

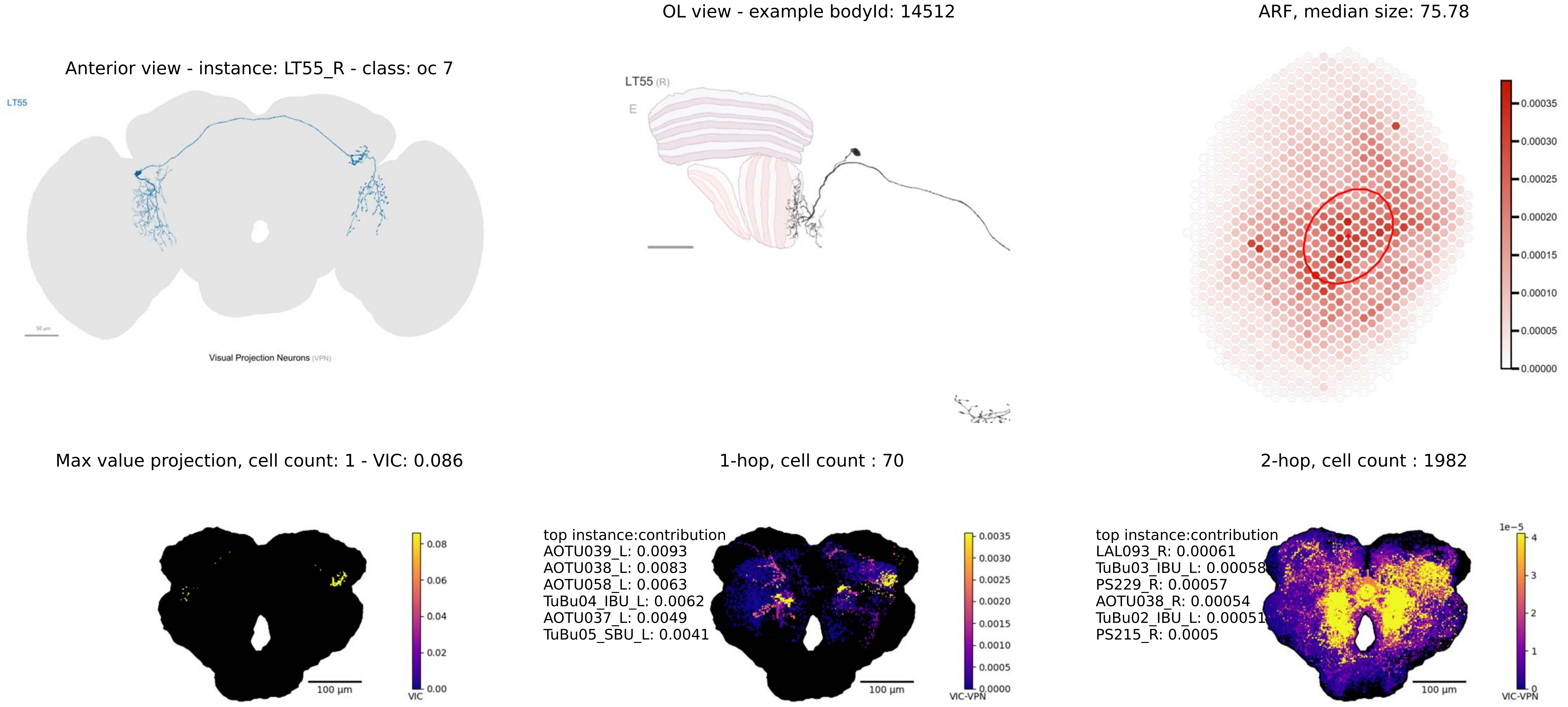

OL view - example bodyId: 14512
ARF, median size: 75.78
Anterior view - instance: LT55_R - class: oc 7
LT55
Visual Projection Neurons (VPN)
LT55 (R)
E
0.00035
0.00030
0.00025
0.00020
0.00015
0.00010
0.00005
0.00000
Max value projection, cell count: 1 - VIC: 0.086
1-hop, cell count : 70
2-hop, cell count : 1982
0.08
0.06
0.04
0.02
0.00
VIC
100 µm
top instance:contribution
AOTU039_L: 0.0093
AOTU038_L: 0.0083
AOTU058_L: 0.0063
TuBu04_IBU_L: 0.0062
AOTU037_L: 0.0049
TuBu05_SBU_L: 0.0041
0.0035
0.0030
0.0025
0.0020
0.0015
0.0010
0.0005
0.0000
VIC-VPN
100 µm
top instance:contribution
LAL093_R: 0.00061
TuBu03_IBU_L: 0.00058
PS229_R: 0.00057
AOTU038_R: 0.00054
TuBu02_IBU_L: 0.00051
PS215_R: 0.0005
1e−5
4
3
2
1
0
VIC-VPN
100 µm

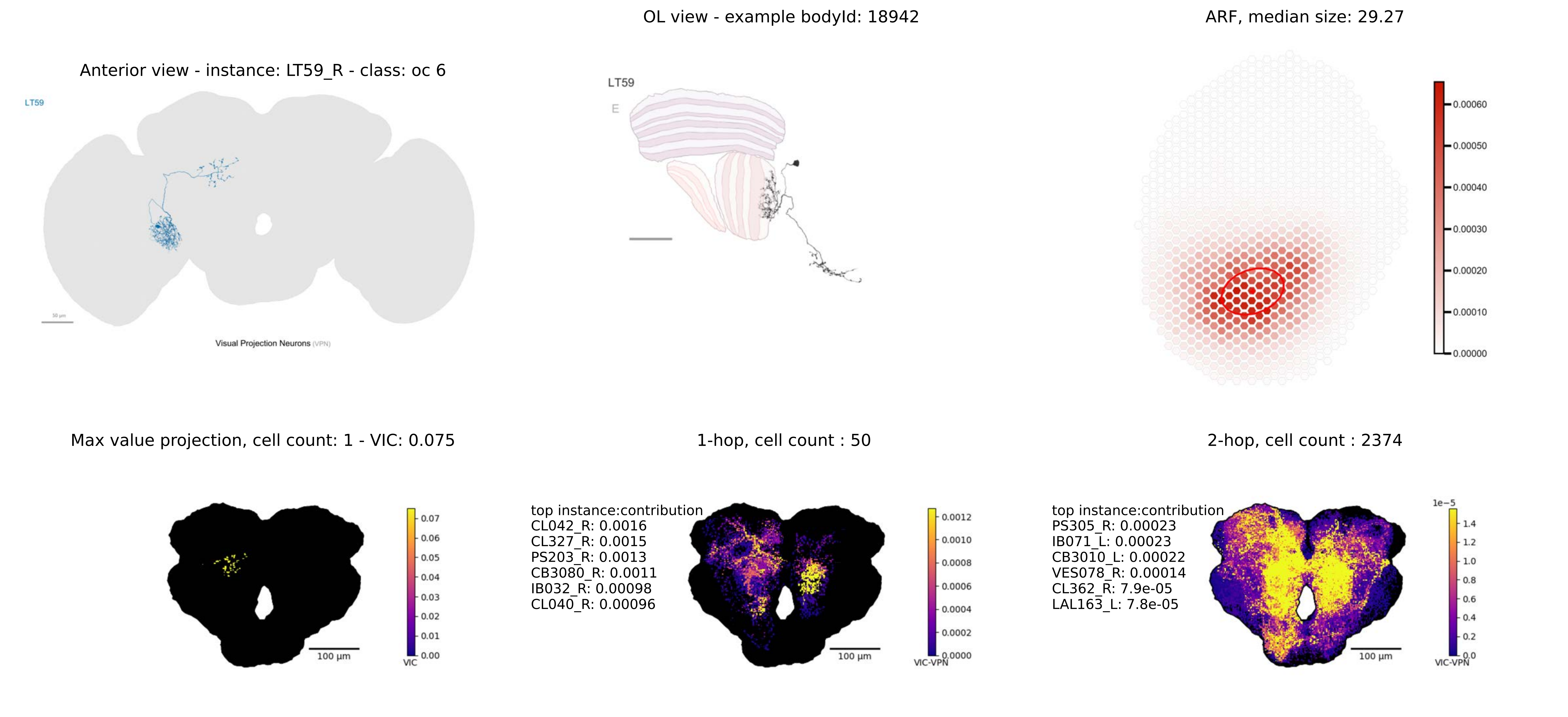

OL view - example bodyId: 18942
ARF, median size: 29.27
Anterior view - instance: LT59_R - class: oc 6
LT59
50 µm
Visual Projection Neurons (VPN)
LT59
E
0.00060
0.00050
0.00040
0.00030
0.00020
0.00010
0.00000
Max value projection, cell count: 1 - VIC: 0.075
1-hop, cell count : 50
2-hop, cell count : 2374
0.07
0.06
0.05
0.04
0.03
0.02
0.01
0.00
VIC
100 µm
top instance:contribution
CL042_R: 0.0016
CL327_R: 0.0015
PS203_R: 0.0013
CB3080_R: 0.0011
IB032_R: 0.00098
CL040_R: 0.00096
0.0012
0.0010
0.0008
0.0006
0.0004
0.0002
0.0000
VIC-VPN
100 µm
top instance:contribution
PS305_R: 0.00023
IB071_L: 0.00023
CB3010_L: 0.00022
VES078_R: 0.00014
CL362_R: 7.9e-05
LAL163_L: 7.8e-05
1e−5
1.4
1.2
1.0
0.8
0.6
0.4
0.2
0.0
VIC-VPN
100 µm

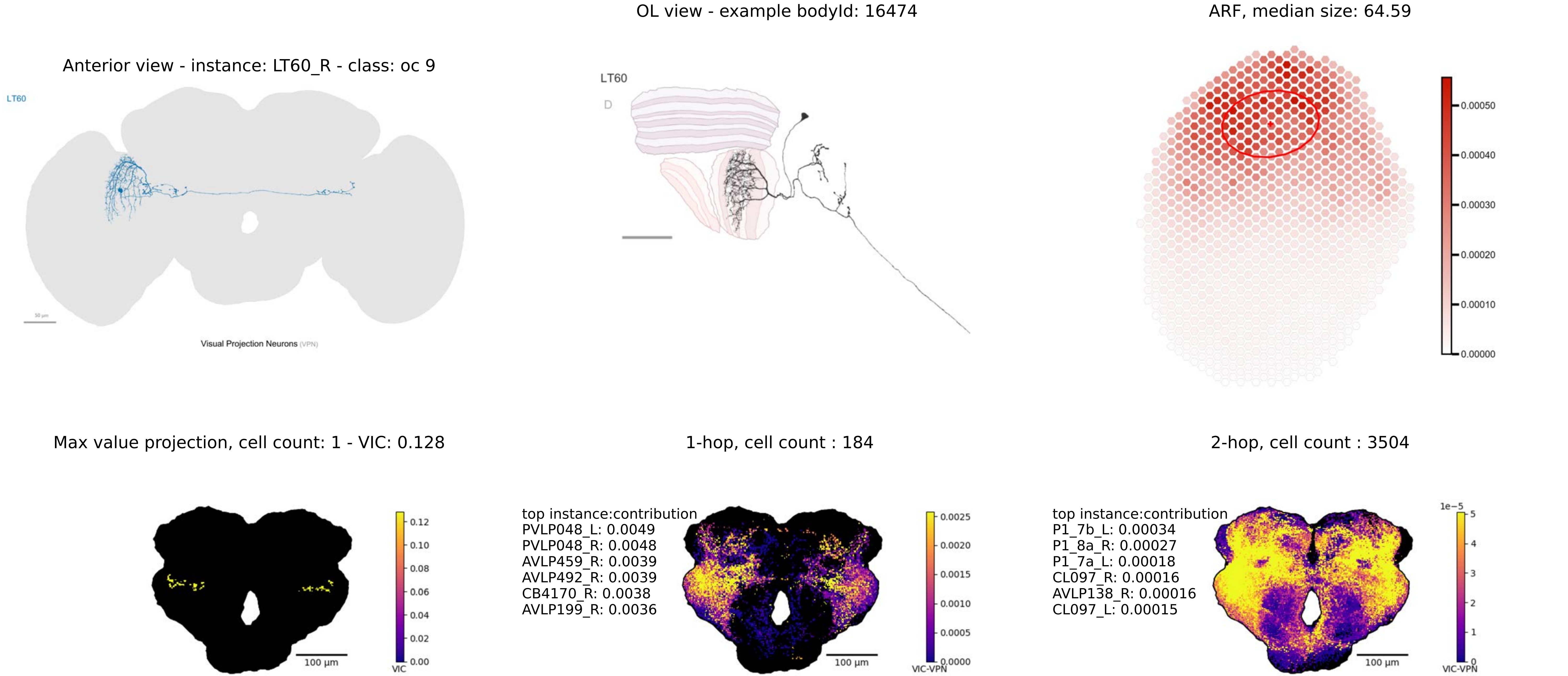

OL view - example bodyId: 16474
ARF, median size: 64.59
Anterior view - instance: LT60_R - class: oc 9
LT60
Visual Projection Neurons (VPN)
LT60
D
0.00050
0.00040
0.00030
0.00020
0.00010
0.00000
Max value projection, cell count: 1 - VIC: 0.128
1-hop, cell count : 184
2-hop, cell count : 3504
100 µm
VIC
0.12
0.10
0.08
0.06
0.04
0.02
0.00
top instance:contribution
PVLP048_L: 0.0049
PVLP048_R: 0.0048
AVLP459_R: 0.0039
AVLP492_R: 0.0039
CB4170_R: 0.0038
AVLP199_R: 0.0036
100 µm
VIC-VPN
0.0025
0.0020
0.0015
0.0010
0.0005
0.0000
top instance:contribution
P1_7b_L: 0.00034
P1_8a_R: 0.00027
P1_7a_L: 0.00018
CL097_R: 0.00016
AVLP138_R: 0.00016
CL097_L: 0.00015
100 µm
1e−5
5
4
3
2
1
0
VIC-VPN

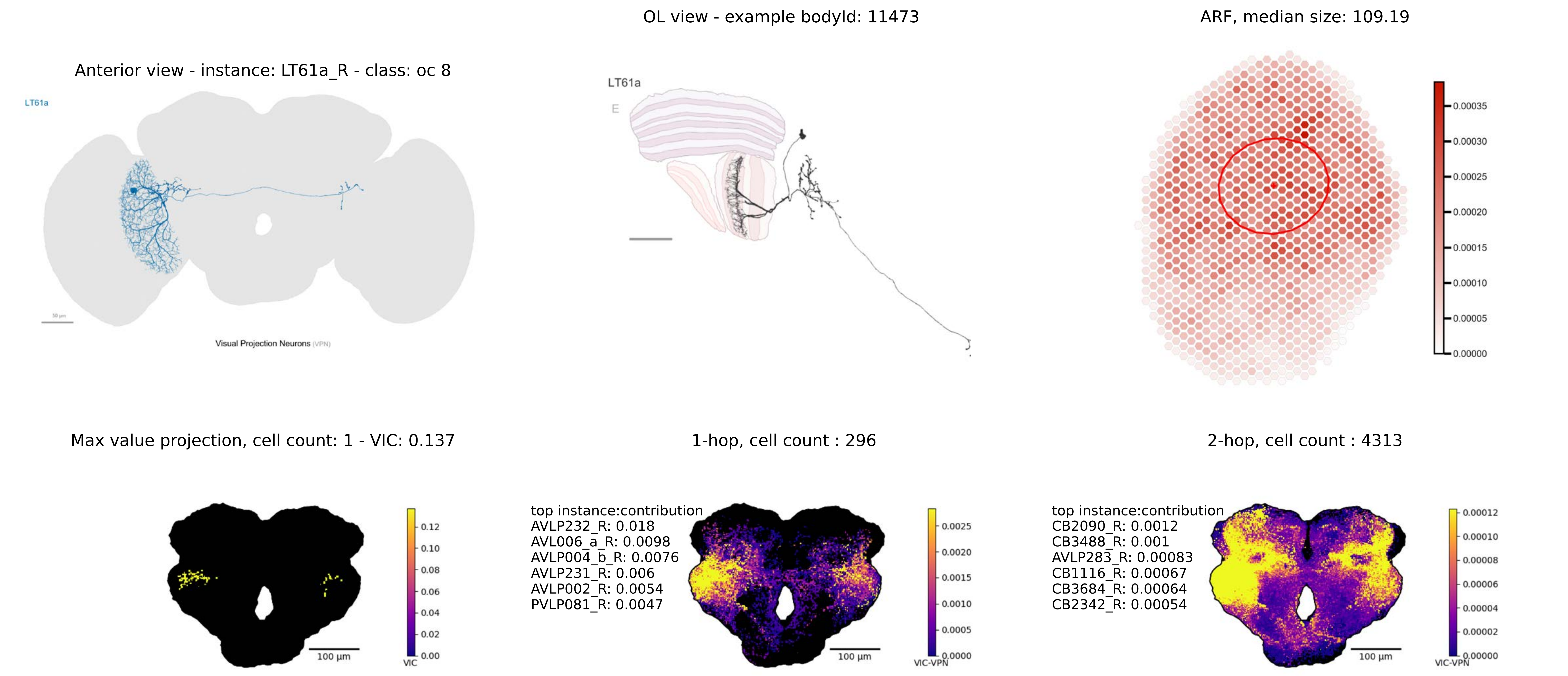

OL view - example bodyId: 11473
ARF, median size: 109.19
Anterior view - instance: LT61a_R - class: oc 8
LT61a
Visual Projection Neurons (VPN)
LT61a
E
0.00035
0.00030
0.00025
0.00020
0.00015
0.00010
0.00005
0.00000
Max value projection, cell count: 1 - VIC: 0.137
1-hop, cell count : 296
2-hop, cell count : 4313
100 µm
0.12
0.10
0.08
0.06
0.04
0.02
0.00
VIC
top instance:contribution
AVLP232_R: 0.018
AVL006_a_R: 0.0098
AVLP004_b_R: 0.0076
AVLP231_R: 0.006
AVLP002_R: 0.0054
PVLP081_R: 0.0047
100 µm
0.0025
0.0020
0.0015
0.0010
0.0005
0.0000
VIC-VPN
top instance:contribution
CB2090_R: 0.0012
CB3488_R: 0.001
AVLP283_R: 0.00083
CB1116_R: 0.00067
CB3684_R: 0.00064
CB2342_R: 0.00054
100 µm
0.00012
0.00010
0.00008
0.00006
0.00004
0.00002
0.00000
VIC-VPN

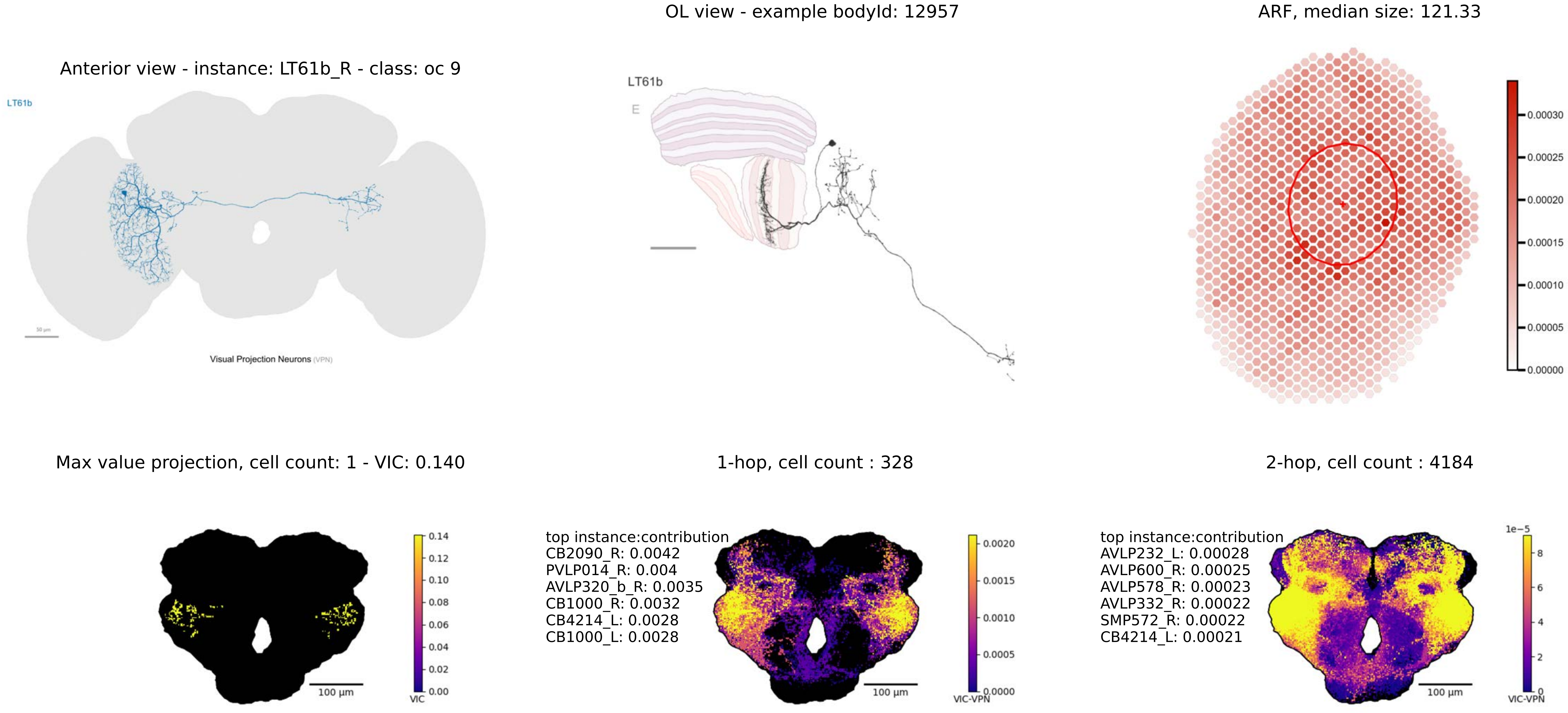

OL view - example bodyId: 12957
ARF, median size: 121.33
Anterior view - instance: LT61b_R - class: oc 9
LT61b
Visual Projection Neurons (VPN)
LT61b
E
0.00030
0.00025
0.00020
0.00015
0.00010
0.00005
0.00000
Max value projection, cell count: 1 - VIC: 0.140
1-hop, cell count : 328
2-hop, cell count : 4184
0.14
0.12
0.10
0.08
0.06
0.04
0.02
0.00
VIC
100 µm
top instance:contribution
CB2090_R: 0.0042
PVLP014_R: 0.004
AVLP320_b_R: 0.0035
CB1000_R: 0.0032
CB4214_L: 0.0028
CB1000_L: 0.0028
0.0020
0.0015
0.0010
0.0005
0.0000
VIC-VPN
100 µm
top instance:contribution
AVLP232_L: 0.00028
AVLP600_R: 0.00025
AVLP578_R: 0.00023
AVLP332_R: 0.00022
SMP572_R: 0.00022
CB4214_L: 0.00021
1e−5
8
6
4
2
0
VIC-VPN
100 µm

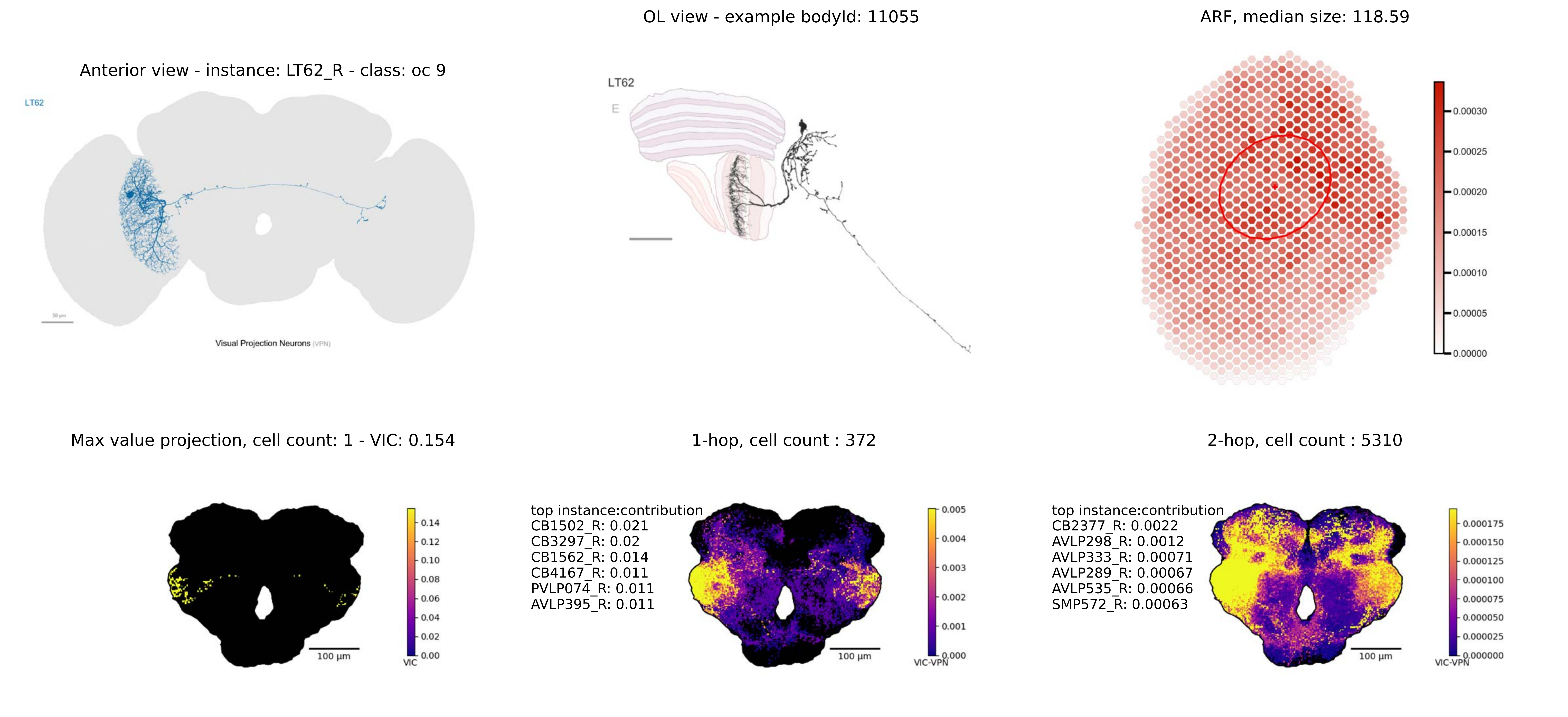

Anterior view - instance: LT62_R - class: oc 9
LT62
Visual Projection Neurons (VPN)
OL view - example bodyId: 11055
LT62
E
ARF, median size: 118.59
0.00030
0.00025
0.00020
0.00015
0.00010
0.00005
0.00000
Max value projection, cell count: 1 - VIC: 0.154
100 µm
VIC
0.14
0.12
0.10
0.08
0.06
0.04
0.02
0.00
1-hop, cell count : 372
top instance:contribution
CB1502_R: 0.021
CB3297_R: 0.02
CB1562_R: 0.014
CB4167_R: 0.011
PVLP074_R: 0.011
AVLP395_R: 0.011
100 µm
VIC-VPN
0.005
0.004
0.003
0.002
0.001
0.000
2-hop, cell count : 5310
top instance:contribution
CB2377_R: 0.0022
AVLP298_R: 0.0012
AVLP333_R: 0.00071
AVLP289_R: 0.00067
AVLP535_R: 0.00066
SMP572_R: 0.00063
100 µm
VIC-VPN
0.000175
0.000150
0.000125
0.000100
0.000075
0.000050
0.000025
0.000000

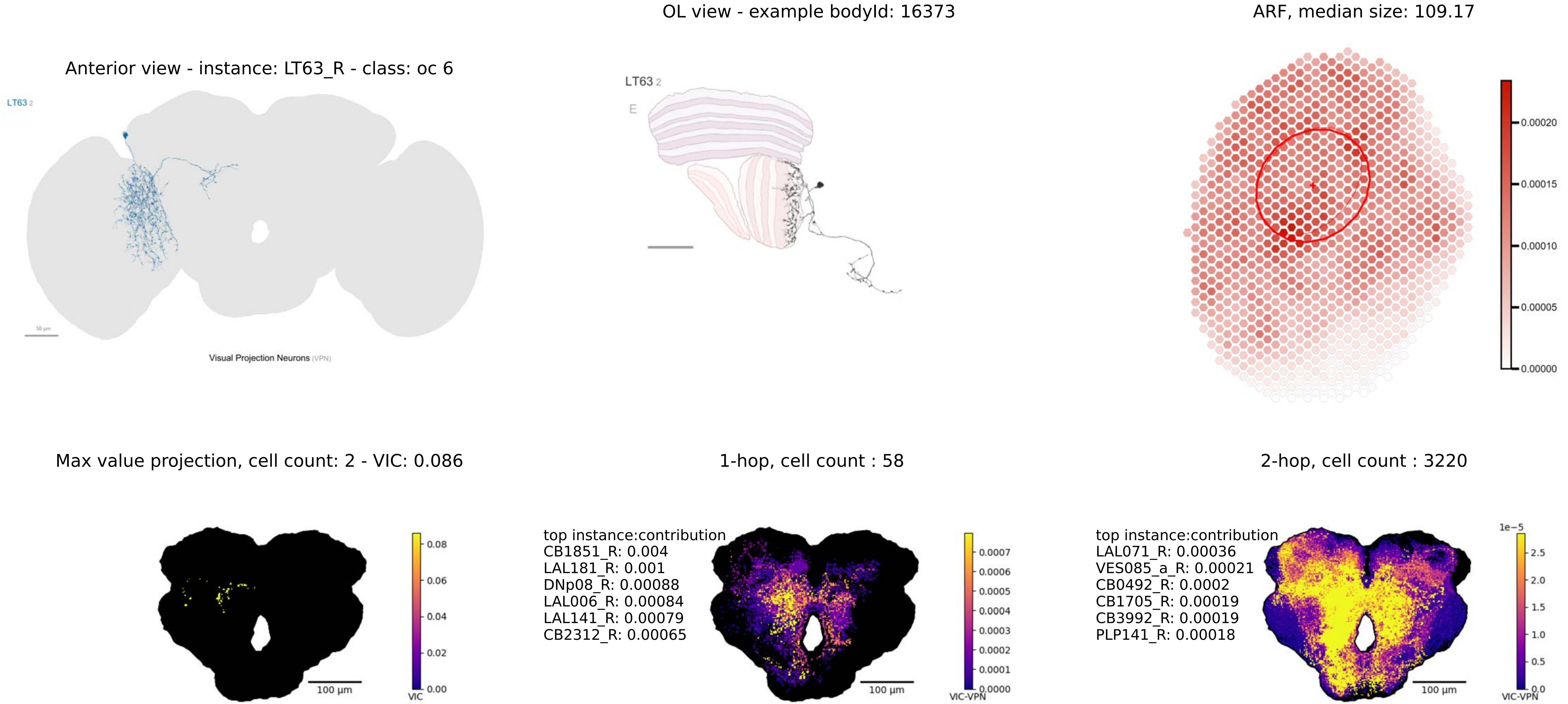
OL view - example bodyId: 16373
ARF, median size: 109.17
Anterior view - instance: LT63_R - class: oc 6
LT63 2
Visual Projection Neurons (VPN)
LT63 2
E
0.00020
0.00015
0.00010
0.00005
0.00000
Max value projection, cell count: 2 - VIC: 0.086
1-hop, cell count : 58
2-hop, cell count : 3220
0.08
0.06
0.04
0.02
0.00
100 µm
VIC
top instance:contribution
CB1851_R: 0.004
LAL181_R: 0.001
DNp08_R: 0.00088
LAL006_R: 0.00084
LAL141_R: 0.00079
CB2312_R: 0.00065
100 µm
0.0007
0.0006
0.0005
0.0004
0.0003
0.0002
0.0001
0.0000
VIC-VPN
top instance:contribution
LAL071_R: 0.00036
VES085_a_R: 0.00021
CB0492_R: 0.0002
CB1705_R: 0.00019
CB3992_R: 0.00019
PLP141_R: 0.00018
100 µm
1e−5
2.5
2.0
1.5
1.0
0.5
0.0
VIC-VPN

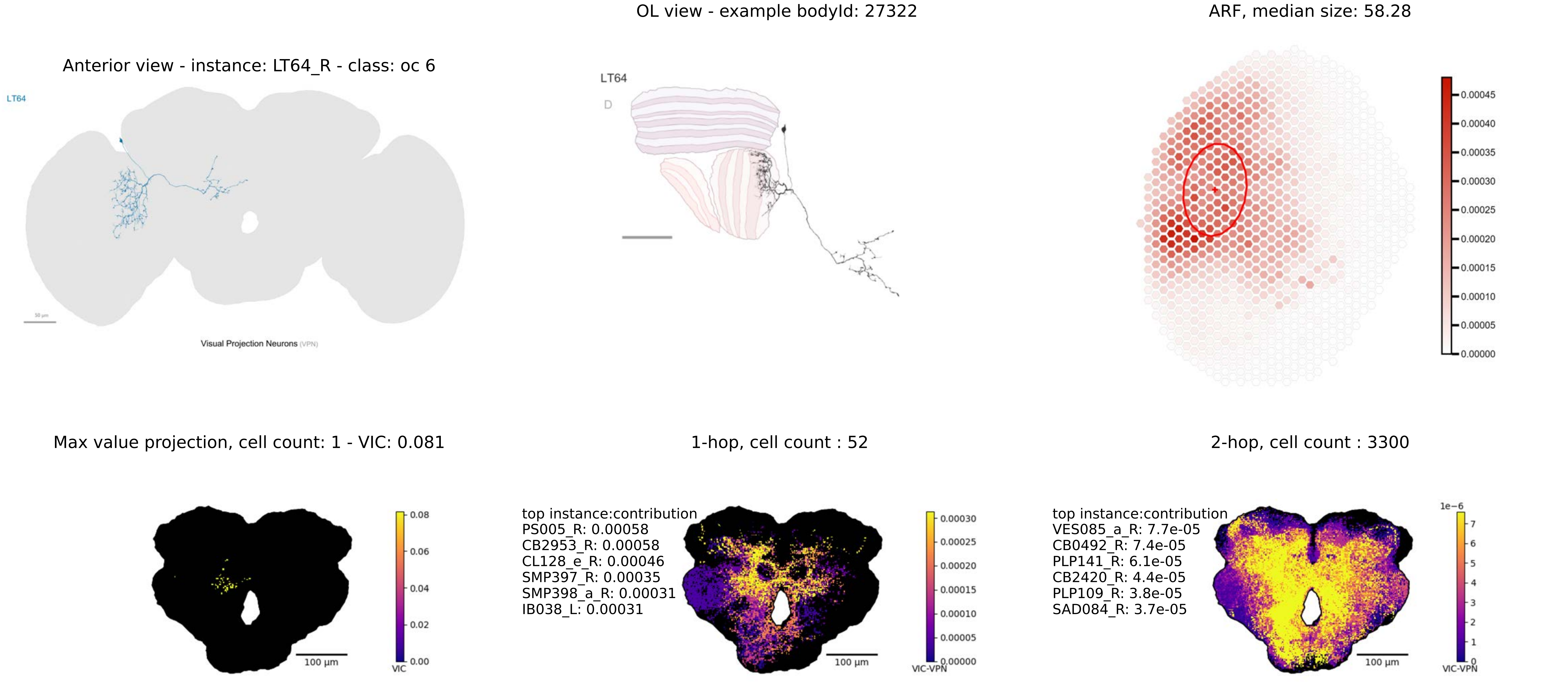
Anterior view - instance: LT64_R - class: oc 6
LT64
Visual Projection Neurons (VPN)
OL view - example bodyId: 27322
LT64
D
ARF, median size: 58.28
0.00045
0.00040
0.00035
0.00030
0.00025
0.00020
0.00015
0.00010
0.00005
0.00000
Max value projection, cell count: 1 - VIC: 0.081
100 µm
0.08
0.06
0.04
0.02
0.00
VIC
1-hop, cell count : 52
top instance:contribution
PS005_R: 0.00058
CB2953_R: 0.00058
CL128_e_R: 0.00046
SMP397_R: 0.00035
SMP398_a_R: 0.00031
IB038_L: 0.00031
100 µm
0.00030
0.00025
0.00020
0.00015
0.00010
0.00005
0.00000
VIC-VPN
2-hop, cell count : 3300
top instance:contribution
VES085_a_R: 7.7e-05
CB0492_R: 7.4e-05
PLP141_R: 6.1e-05
CB2420_R: 4.4e-05
PLP109_R: 3.8e-05
SAD084_R: 3.7e-05
100 µm
1e−6
7
6
5
4
3
2
1
0
VIC-VPN

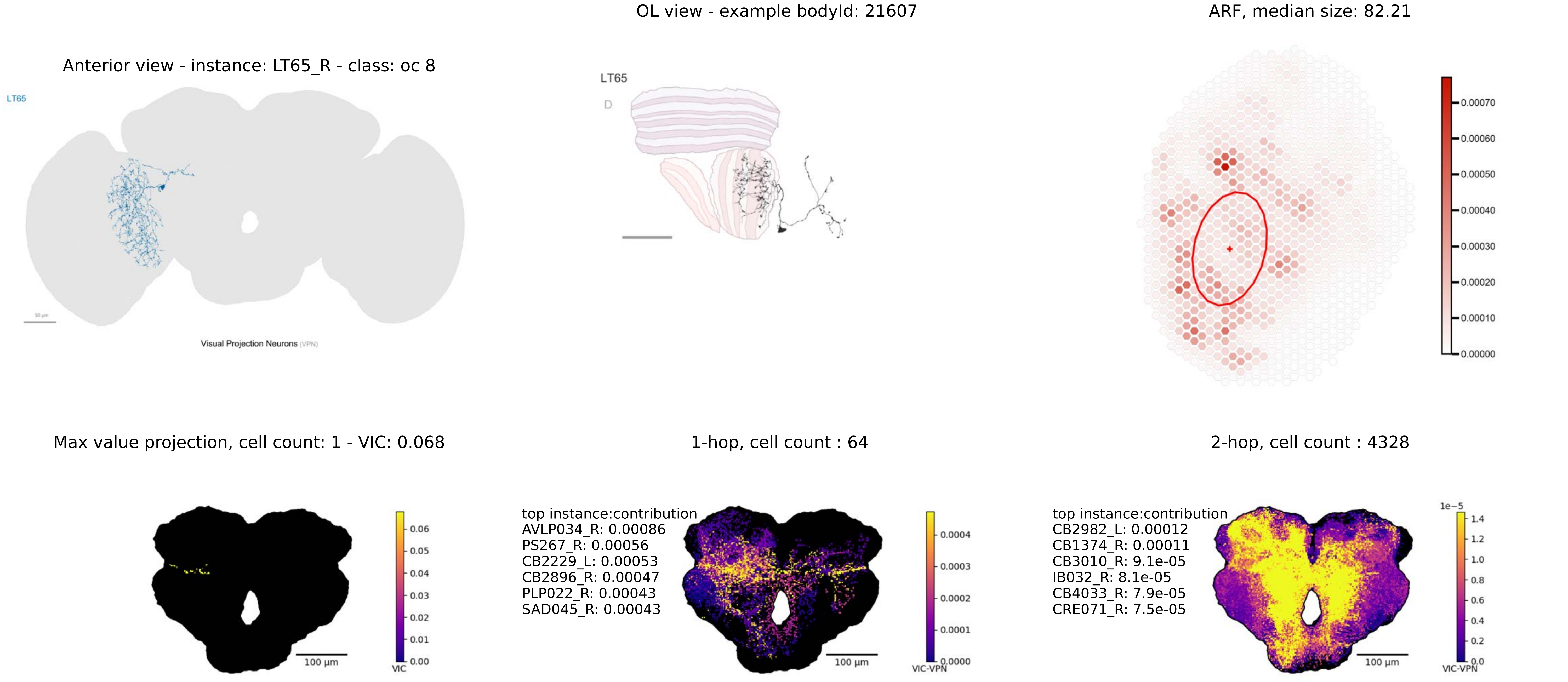

OL view - example bodyId: 21607
ARF, median size: 82.21
Anterior view - instance: LT65_R - class: oc 8
LT65
Visual Projection Neurons (VPN)
LT65
D
0.00070
0.00060
0.00050
0.00040
0.00030
0.00020
0.00010
0.00000
Max value projection, cell count: 1 - VIC: 0.068
1-hop, cell count : 64
2-hop, cell count : 4328
100 µm
VIC
0.06
0.05
0.04
0.03
0.02
0.01
0.00
top instance:contribution
AVLP034_R: 0.00086
PS267_R: 0.00056
CB2229_L: 0.00053
CB2896_R: 0.00047
PLP022_R: 0.00043
SAD045_R: 0.00043
100 µm
VIC-VPN
0.0004
0.0003
0.0002
0.0001
0.0000
top instance:contribution
CB2982_L: 0.00012
CB1374_R: 0.00011
CB3010_R: 9.1e-05
IB032_R: 8.1e-05
CB4033_R: 7.9e-05
CRE071_R: 7.5e-05
100 µm
1e−5
1.4
1.2
1.0
0.8
0.6
0.4
0.2
0.0
VIC-VPN

Anterior view - instance: LT66_R - class: oc 9

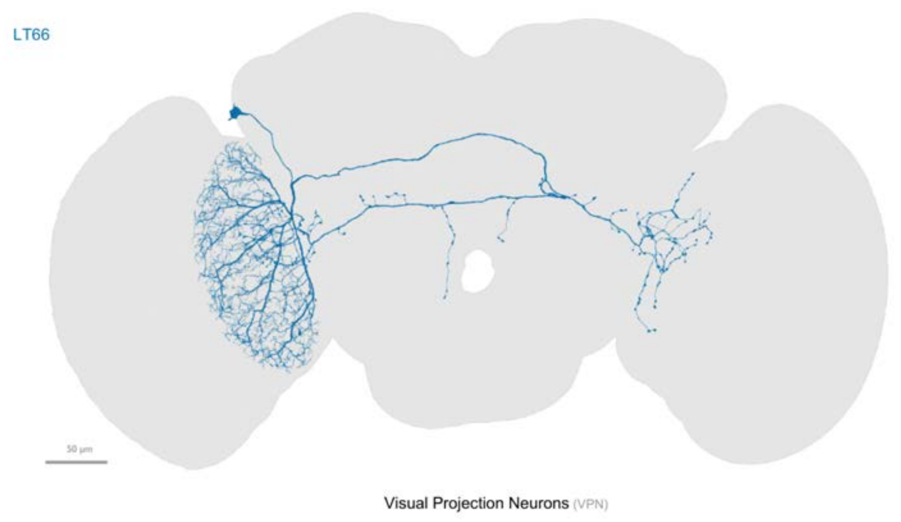


OL view - example bodyId: 10771

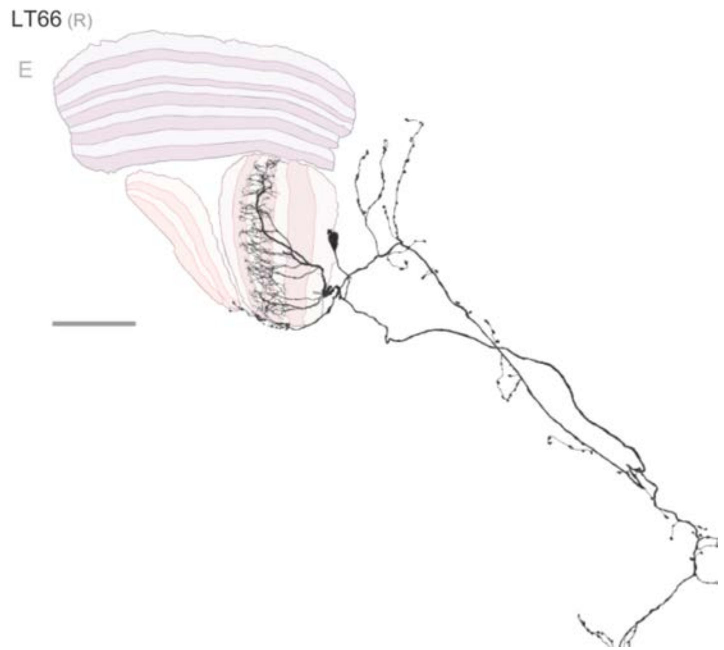


ARF, median size: 150.97

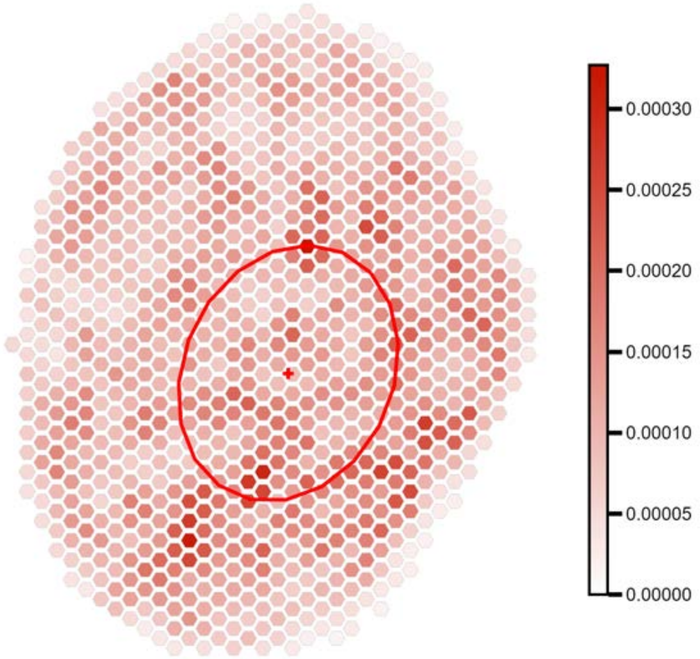


Max value projection, cell count: 1 - VIC: 0.102

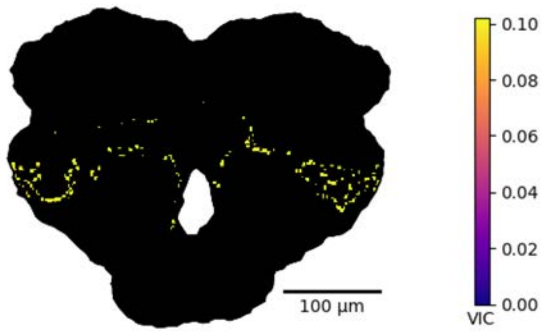


1-hop, cell count : 128

top instance:contribution
aMe_TBD1_L: 0.0036
PVLP024_R: 0.0022
PVLP024_L: 0.0021
PLP191_R: 0.0016
GNG282_L: 0.0016
GNG282_R: 0.0015

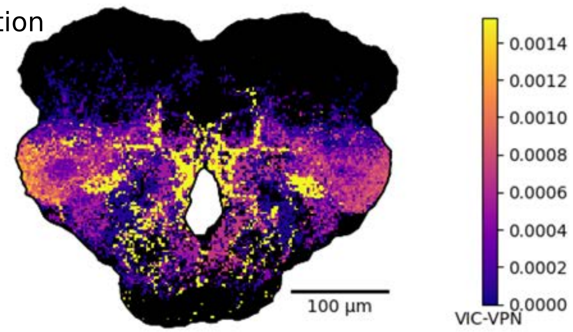


2-hop, cell count : 4761

top instance:contribution
CB2646_R: 0.00046
CB1787_R: 0.00029
CB3332_R: 0.00027
DNg95_R: 0.0002
AOTU064_R: 0.0002
PS149_L: 0.00019

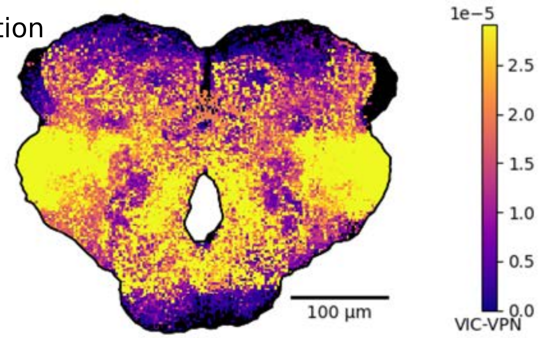

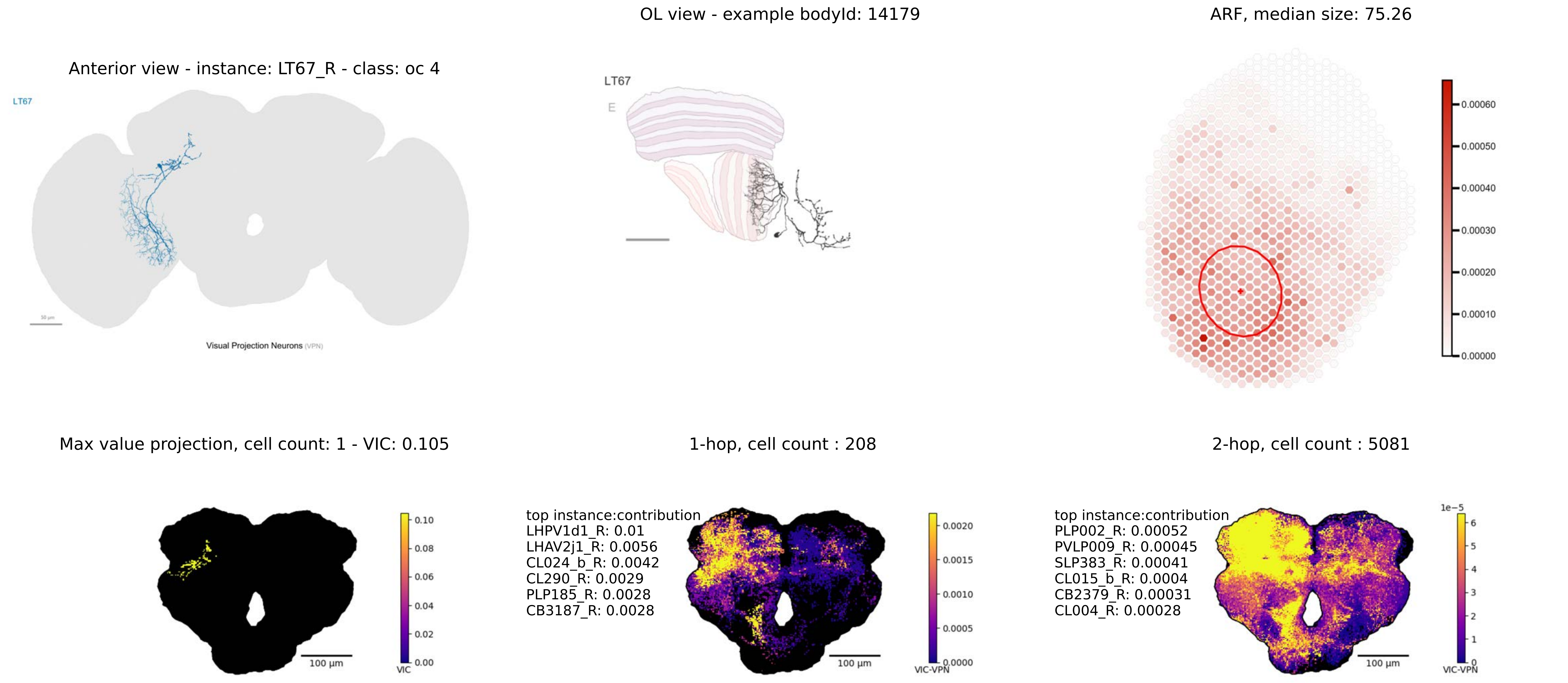
Anterior view - instance: LT67_R - class: oc 4
LT67
Visual Projection Neurons (VPN)
OL view - example bodyId: 14179
LT67
E
ARF, median size: 75.26
0.00060
0.00050
0.00040
0.00030
0.00020
0.00010
0.00000
Max value projection, cell count: 1 - VIC: 0.105
100 µm
0.10
0.08
0.06
0.04
0.02
0.00
VIC
1-hop, cell count : 208
top instance:contribution
LHPV1d1_R: 0.01
LHAV2j1_R: 0.0056
CL024_b_R: 0.0042
CL290_R: 0.0029
PLP185_R: 0.0028
CB3187_R: 0.0028
100 µm
0.0020
0.0015
0.0010
0.0005
0.0000
VIC-VPN
2-hop, cell count : 5081
top instance:contribution
PLP002_R: 0.00052
PVLP009_R: 0.00045
SLP383_R: 0.00041
CL015_b_R: 0.0004
CB2379_R: 0.00031
CL004_R: 0.00028
100 µm
1e−5
6
5
4
3
2
1
0
VIC-VPN

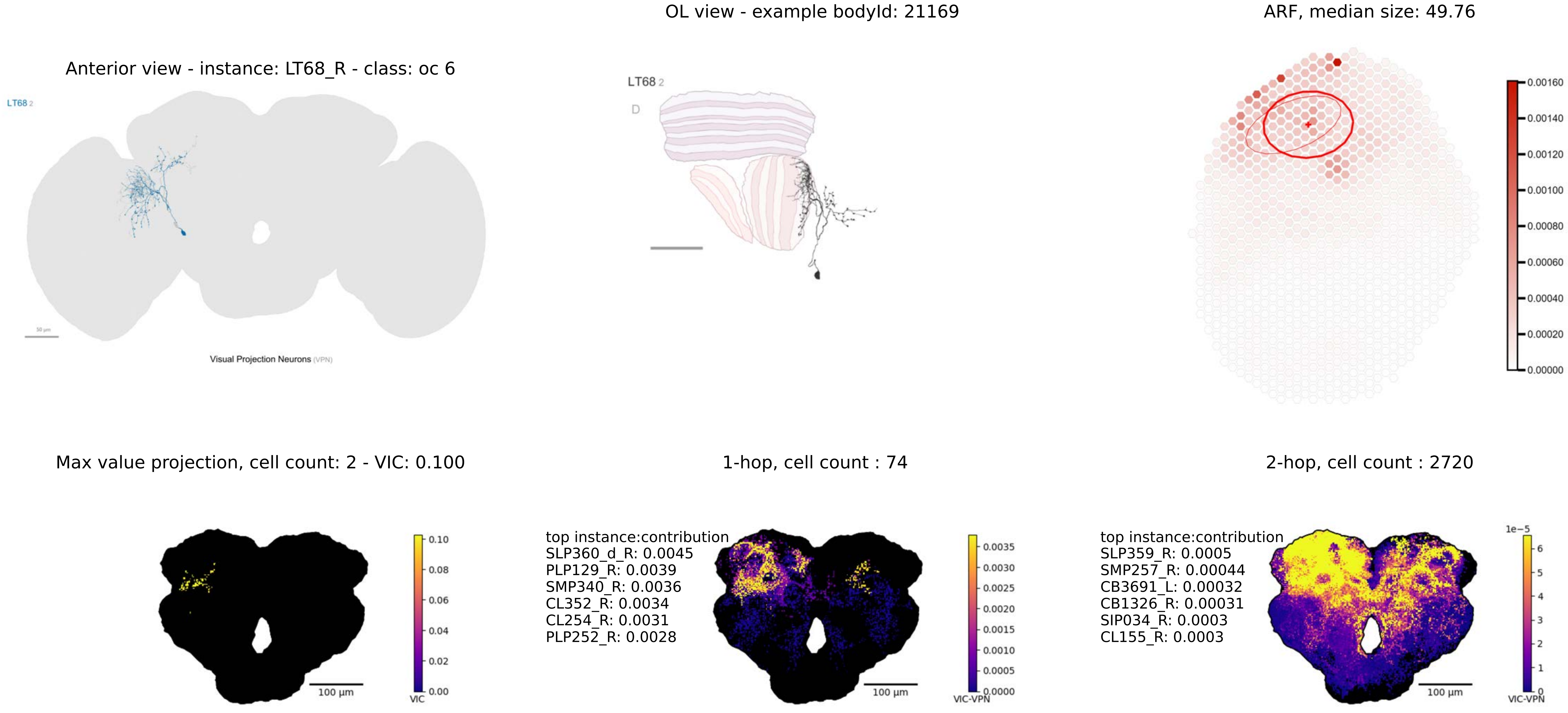
OL view - example bodyId: 21169
ARF, median size: 49.76
Anterior view - instance: LT68_R - class: oc 6
LT68 2
Visual Projection Neurons (VPN)
LT68 2
D
0.00160
0.00140
0.00120
0.00100
0.00080
0.00060
0.00040
0.00020
0.00000
Max value projection, cell count: 2 - VIC: 0.100
1-hop, cell count : 74
2-hop, cell count : 2720
0.10
0.08
0.06
0.04
0.02
0.00
VIC
100 µm
top instance:contribution
SLP360_d_R: 0.0045
PLP129_R: 0.0039
SMP340_R: 0.0036
CL352_R: 0.0034
CL254_R: 0.0031
PLP252_R: 0.0028
0.0035
0.0030
0.0025
0.0020
0.0015
0.0010
0.0005
0.0000
VIC-VPN
100 µm
top instance:contribution
SLP359_R: 0.0005
SMP257_R: 0.00044
CB3691_L: 0.00032
CB1326_R: 0.00031
SIP034_R: 0.0003
CL155_R: 0.0003
1e−5
6
5
4
3
2
1
0
VIC-VPN
100 µm

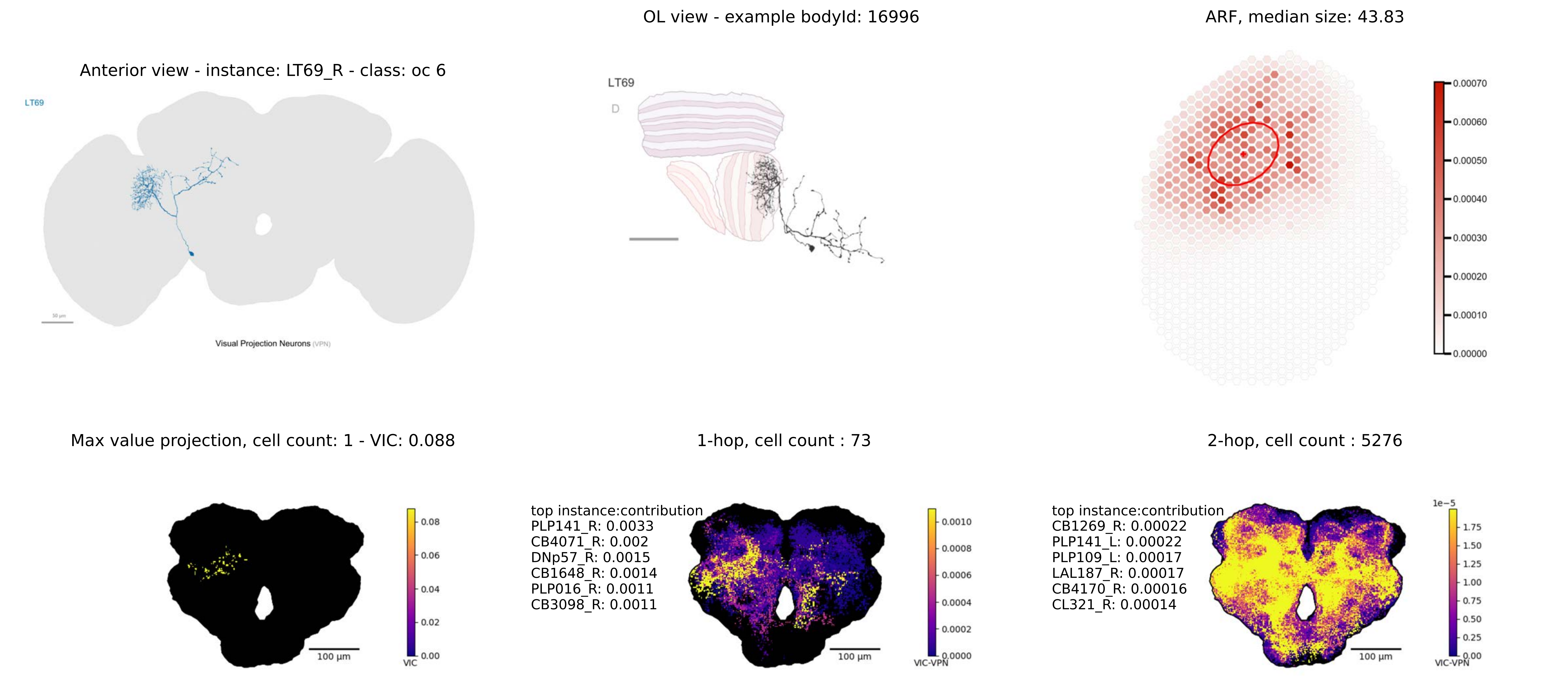

Anterior view - instance: LT69_R - class: oc 6
LT69
Visual Projection Neurons (VPN)
OL view - example bodyId: 16996
LT69
D
ARF, median size: 43.83
0.00070
0.00060
0.00050
0.00040
0.00030
0.00020
0.00010
0.00000
Max value projection, cell count: 1 - VIC: 0.088
0.08
0.06
0.04
0.02
0.00
VIC
100 µm
1-hop, cell count : 73
top instance:contribution
PLP141_R: 0.0033
CB4071_R: 0.002
DNp57_R: 0.0015
CB1648_R: 0.0014
PLP016_R: 0.0011
CB3098_R: 0.0011
0.0010
0.0008
0.0006
0.0004
0.0002
0.0000
VIC-VPN
100 µm
2-hop, cell count : 5276
top instance:contribution
CB1269_R: 0.00022
PLP141_L: 0.00022
PLP109_L: 0.00017
LAL187_R: 0.00017
CB4170_R: 0.00016
CL321_R: 0.00014
1e−5
1.75
1.50
1.25
1.00
0.75
0.50
0.25
0.00
VIC-VPN
100 µm

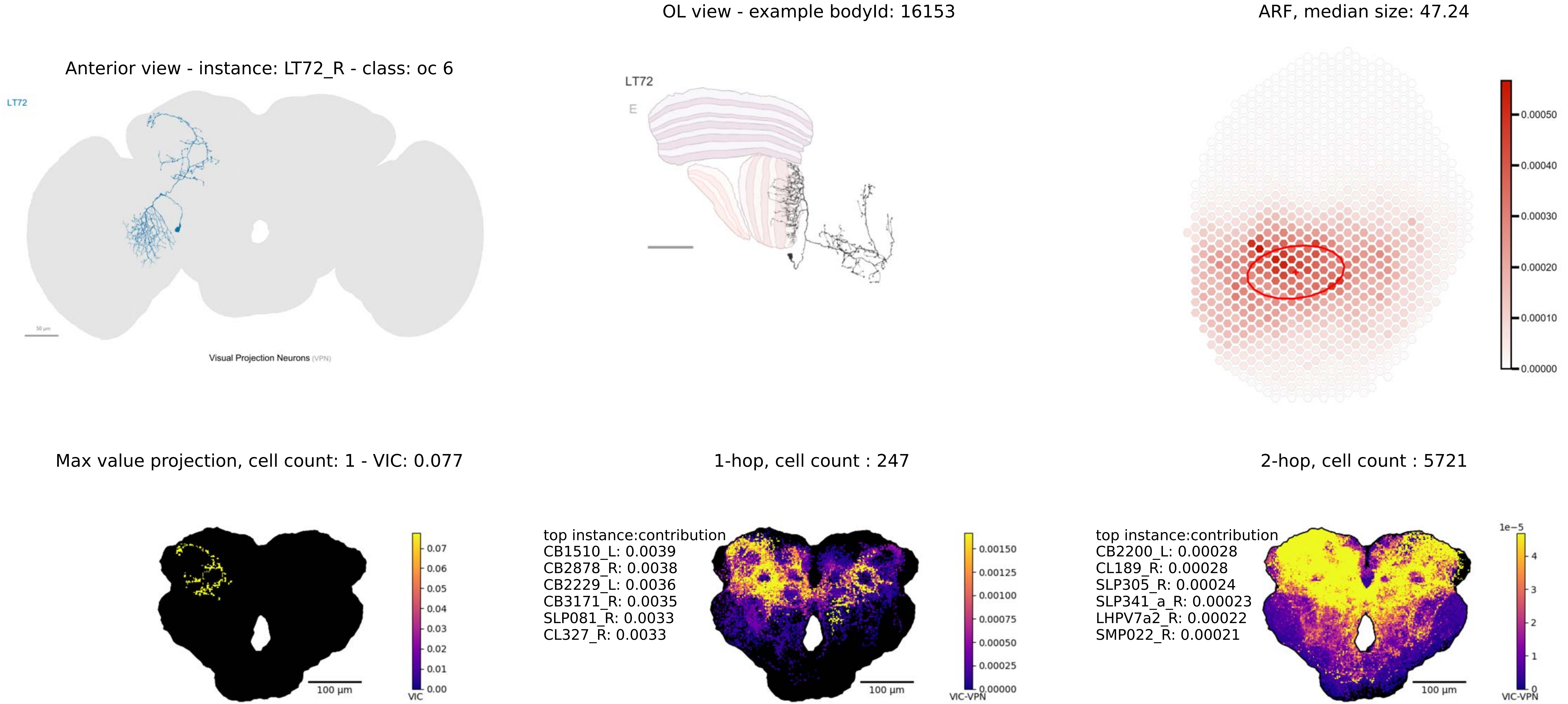

OL view - example bodyId: 16153
ARF, median size: 47.24
Anterior view - instance: LT72_R - class: oc 6
LT72
Visual Projection Neurons (VPN)
LT72
E
0.00050
0.00040
0.00030
0.00020
0.00010
0.00000
Max value projection, cell count: 1 - VIC: 0.077
1-hop, cell count : 247
2-hop, cell count : 5721
0.07
0.06
0.05
0.04
0.03
0.02
0.01
0.00
VIC
100 µm
top instance:contribution
CB1510_L: 0.0039
CB2878_R: 0.0038
CB2229_L: 0.0036
CB3171_R: 0.0035
SLP081_R: 0.0033
CL327_R: 0.0033
0.00150
0.00125
0.00100
0.00075
0.00050
0.00025
0.00000
VIC-VPN
100 µm
top instance:contribution
CB2200_L: 0.00028
CL189_R: 0.00028
SLP305_R: 0.00024
SLP341_a_R: 0.00023
LHPV7a2_R: 0.00022
SMP022_R: 0.00021
1e−5
4
3
2
1
0
VIC-VPN
100 µm

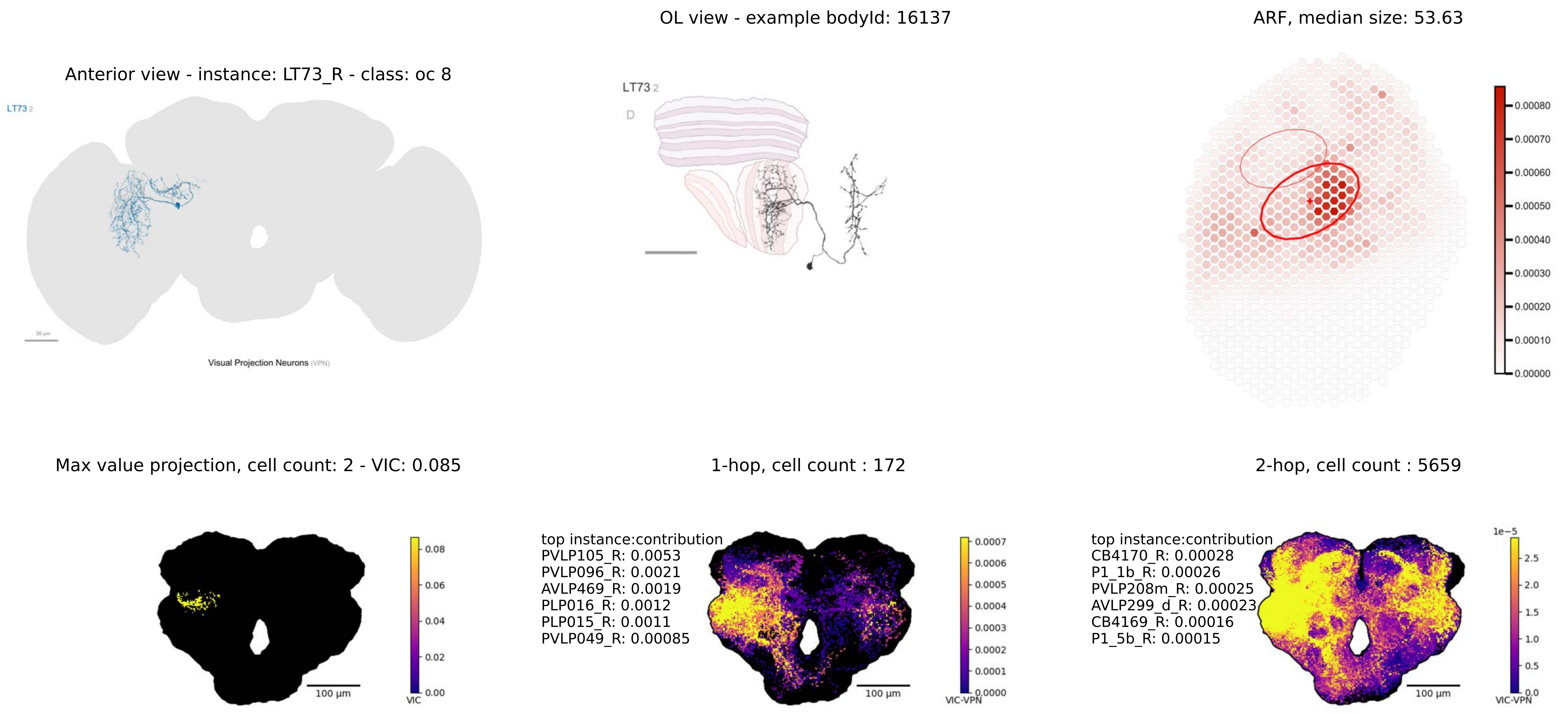
Anterior view - instance: LT73_R - class: oc 8
LT73 2
Visual Projection Neurons (VPN)
OL view - example bodyId: 16137
LT73 2
D
ARF, median size: 53.63
0.00080
0.00070
0.00060
0.00050
0.00040
0.00030
0.00020
0.00010
0.00000
Max value projection, cell count: 2 - VIC: 0.085
100 µm
0.08
0.06
0.04
0.02
0.00
VIC
1-hop, cell count : 172
top instance:contribution
PVLP105_R: 0.0053
PVLP096_R: 0.0021
AVLP469_R: 0.0019
PLP016_R: 0.0012
PLP015_R: 0.0011
PVLP049_R: 0.00085
100 µm
0.0007
0.0006
0.0005
0.0004
0.0003
0.0002
0.0001
0.0000
VIC-VPN
2-hop, cell count : 5659
top instance:contribution
CB4170_R: 0.00028
P1_1b_R: 0.00026
PVLP208m_R: 0.00025
AVLP299_d_R: 0.00023
CB4169_R: 0.00016
P1_5b_R: 0.00015
100 µm
1e−5
2.5
2.0
1.5
1.0
0.5
0.0
VIC-VPN

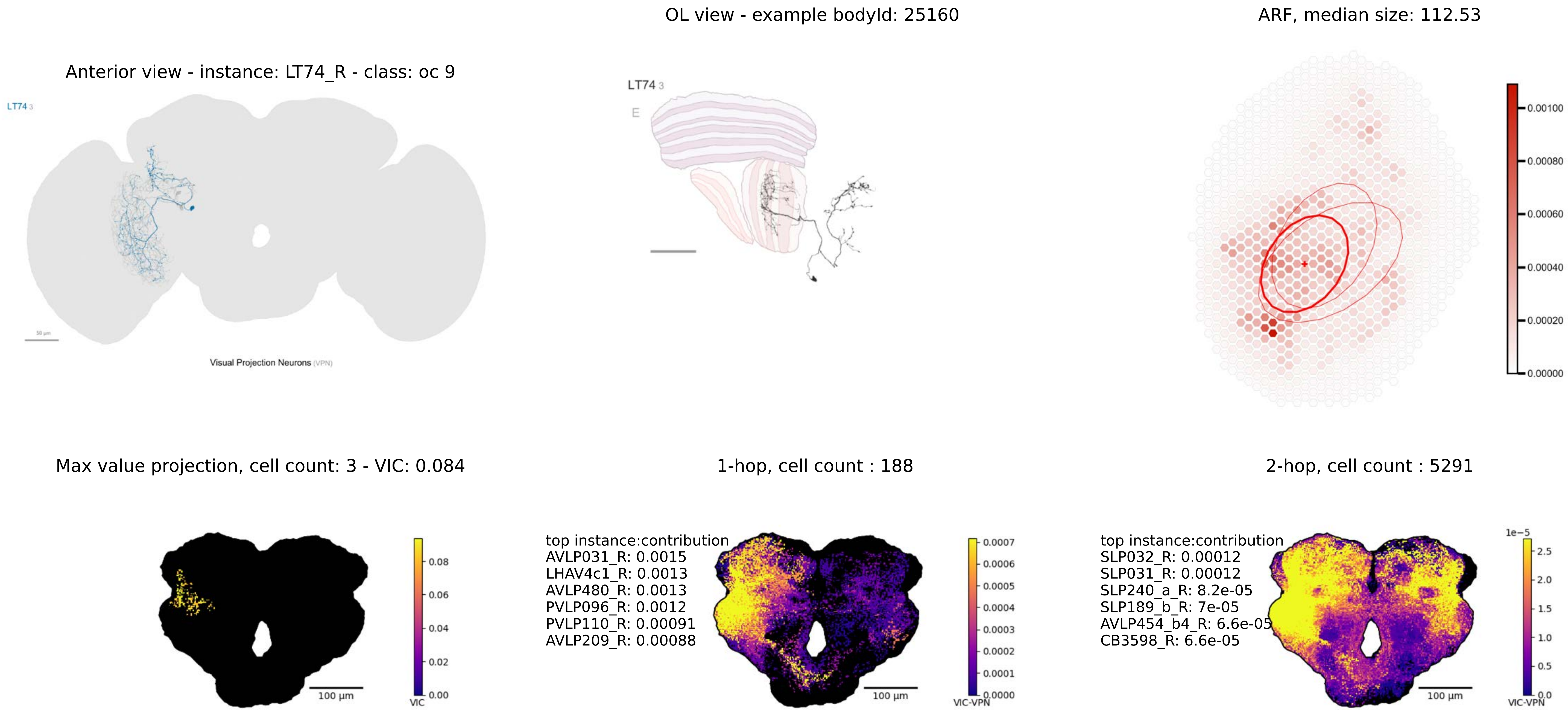
OL view - example bodyId: 25160
ARF, median size: 112.53
Anterior view - instance: LT74_R - class: oc 9
LT74 3
Visual Projection Neurons (VPN)
LT74 3
E
0.00100
0.00080
0.00060
0.00040
0.00020
0.00000
Max value projection, cell count: 3 - VIC: 0.084
1-hop, cell count : 188
2-hop, cell count : 5291
0.08
0.06
0.04
0.02
0.00
VIC
100 µm
top instance:contribution
AVLP031_R: 0.0015
LHAV4c1_R: 0.0013
AVLP480_R: 0.0013
PVLP096_R: 0.0012
PVLP110_R: 0.00091
AVLP209_R: 0.00088
100 µm
0.0007
0.0006
0.0005
0.0004
0.0003
0.0002
0.0001
0.0000
VIC-VPN
top instance:contribution
SLP032_R: 0.00012
SLP031_R: 0.00012
SLP240_a_R: 8.2e-05
SLP189_b_R: 7e-05
AVLP454_b4_R: 6.6e-05
CB3598_R: 6.6e-05
100 µm
1e−5
2.5
2.0
1.5
1.0
0.5
0.0
VIC-VPN

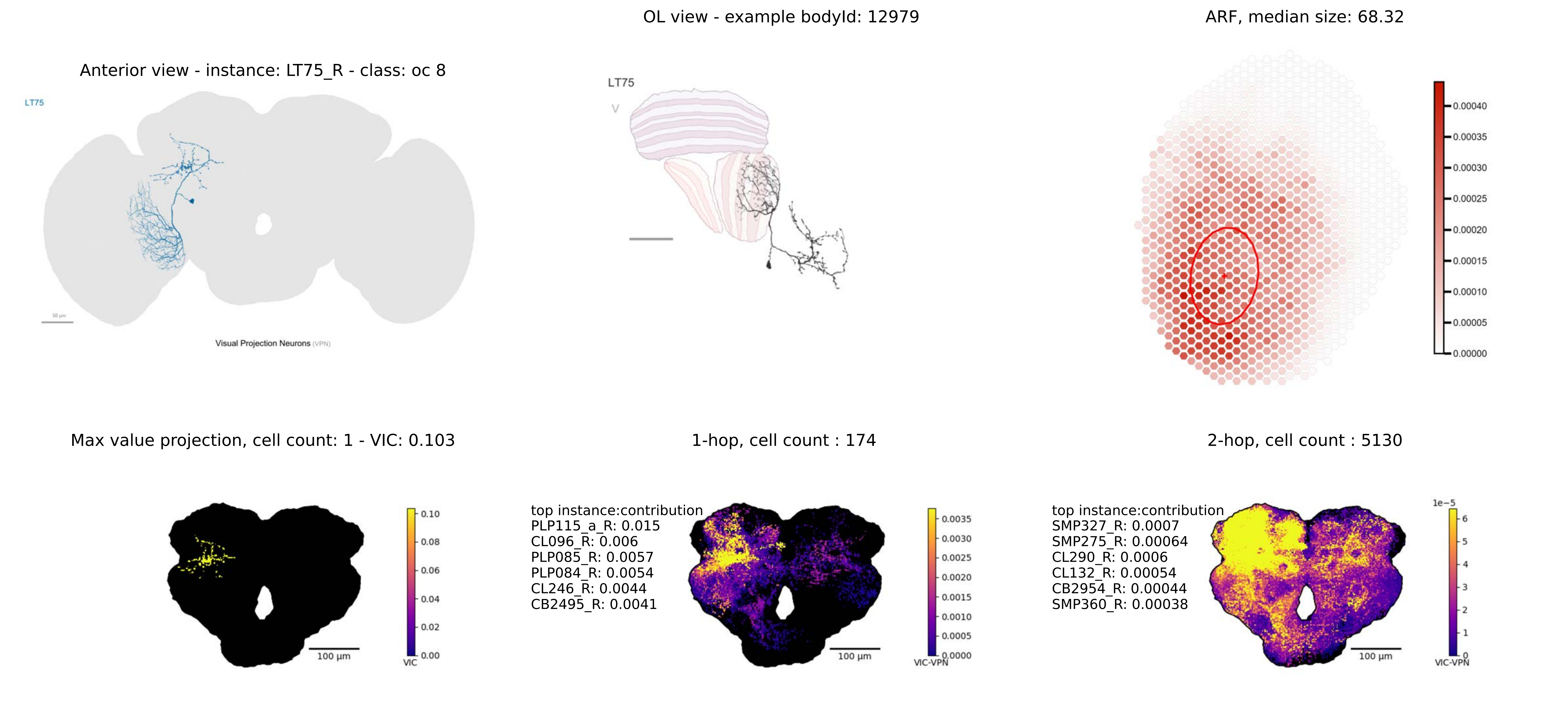
OL view - example bodyId: 12979
ARF, median size: 68.32
Anterior view - instance: LT75_R - class: oc 8
LT75
Visual Projection Neurons (VPN)
LT75
Max value projection, cell count: 1 - VIC: 0.103
1-hop, cell count : 174
2-hop, cell count : 5130
top instance:contribution
PLP115_a_R: 0.015
CL096_R: 0.006
PLP085_R: 0.0057
PLP084_R: 0.0054
CL246_R: 0.0044
CB2495_R: 0.0041
top instance:contribution
SMP327_R: 0.0007
SMP275_R: 0.00064
CL290_R: 0.0006
CL132_R: 0.00054
CB2954_R: 0.00044
SMP360_R: 0.00038
100 µm
100 µm
100 µm
VIC
VIC-VPN
VIC-VPN
1e−5

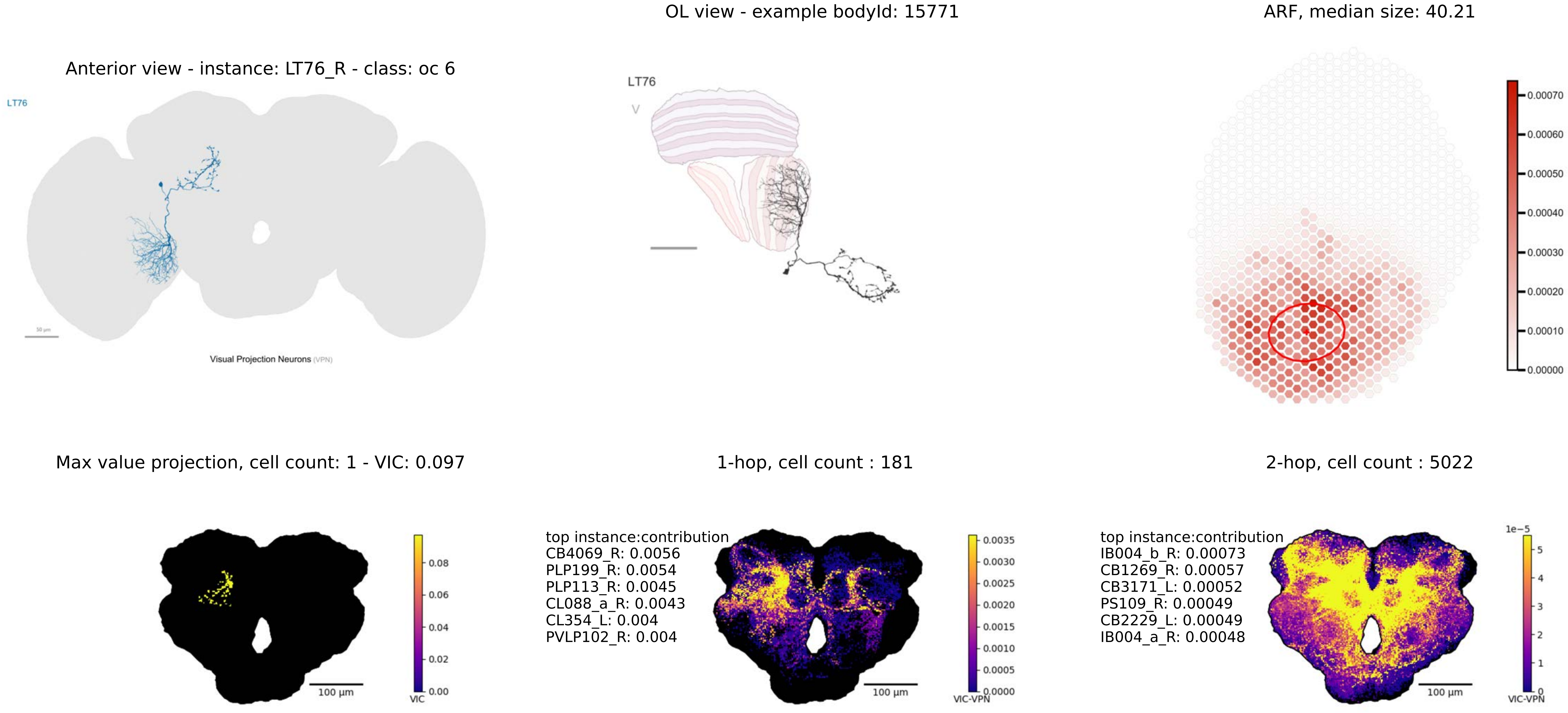
OL view - example bodyId: 15771
ARF, median size: 40.21
Anterior view - instance: LT76_R - class: oc 6
LT76
Visual Projection Neurons (VPN)
LT76
Max value projection, cell count: 1 - VIC: 0.097
1-hop, cell count : 181
2-hop, cell count : 5022
top instance:contribution
CB4069_R: 0.0056
PLP199_R: 0.0054
PLP113_R: 0.0045
CL088_a_R: 0.0043
CL354_L: 0.004
PVLP102_R: 0.004
top instance:contribution
IB004_b_R: 0.00073
CB1269_R: 0.00057
CB3171_L: 0.00052
PS109_R: 0.00049
CB2229_L: 0.00049
IB004_a_R: 0.00048
100 µm
VIC
VIC-VPN
1e−5

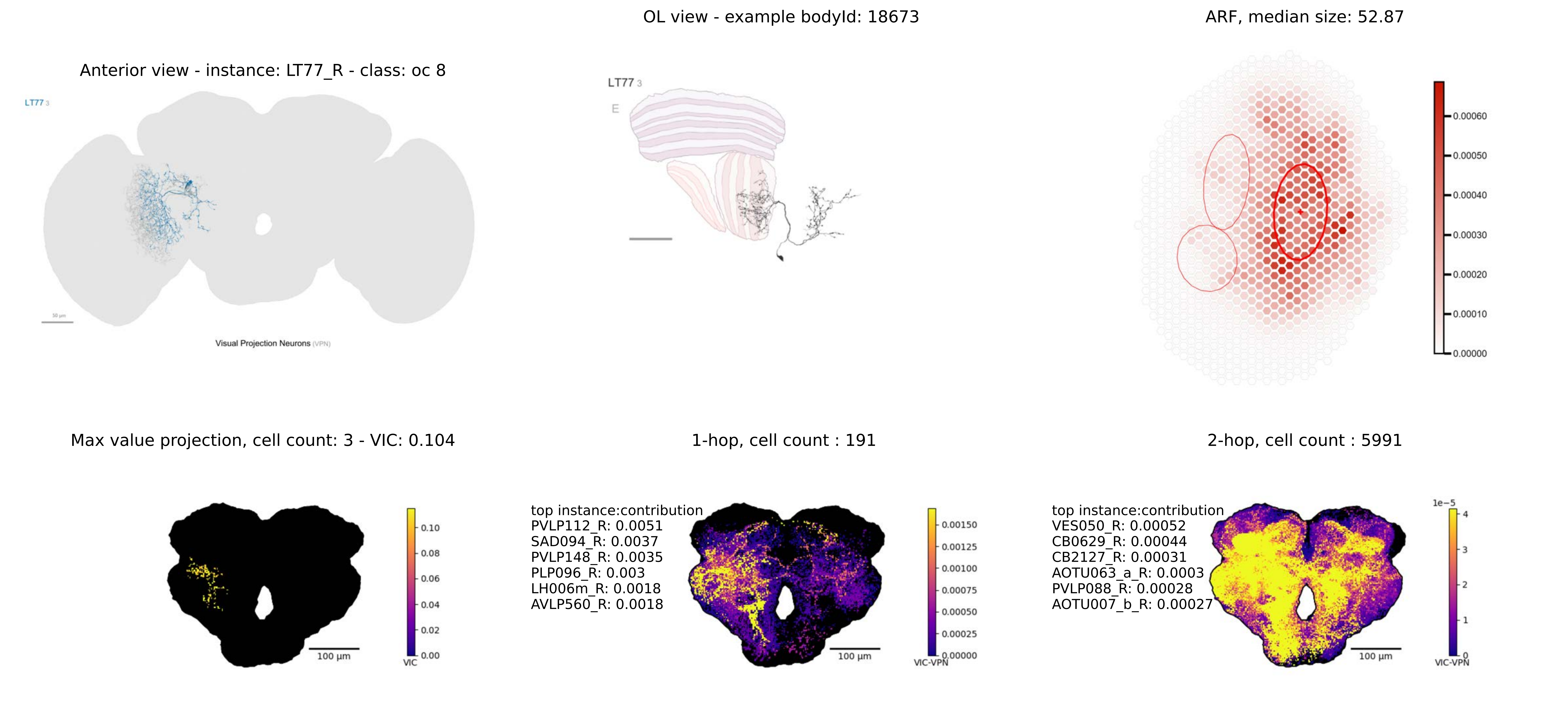
OL view - example bodyId: 18673
ARF, median size: 52.87
Anterior view - instance: LT77_R - class: oc 8
LT77 3
Visual Projection Neurons (VPN)
LT77 3
E
0.00060
0.00050
0.00040
0.00030
0.00020
0.00010
0.00000
Max value projection, cell count: 3 - VIC: 0.104
1-hop, cell count : 191
2-hop, cell count : 5991
0.10
0.08
0.06
0.04
0.02
0.00
VIC
100 µm
top instance:contribution
PVLP112_R: 0.0051
SAD094_R: 0.0037
PVLP148_R: 0.0035
PLP096_R: 0.003
LH006m_R: 0.0018
AVLP560_R: 0.0018
0.00150
0.00125
0.00100
0.00075
0.00050
0.00025
0.00000
VIC-VPN
100 µm
top instance:contribution
VES050_R: 0.00052
CB0629_R: 0.00044
CB2127_R: 0.00031
AOTU063_a_R: 0.0003
PVLP088_R: 0.00028
AOTU007_b_R: 0.00027
1e−5
4
3
2
1
0
VIC-VPN
100 µm

Anterior view - instance: LT78_R - class: oc 8

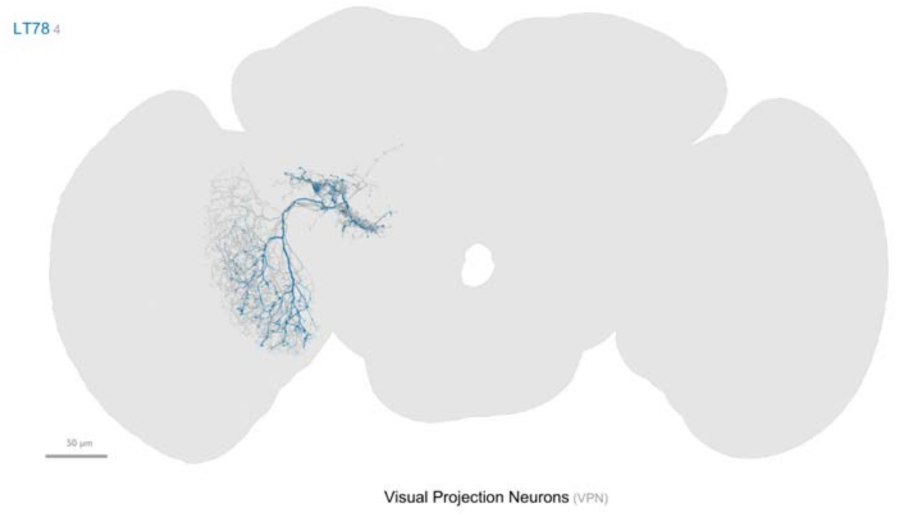


OL view - example bodyId: 16876

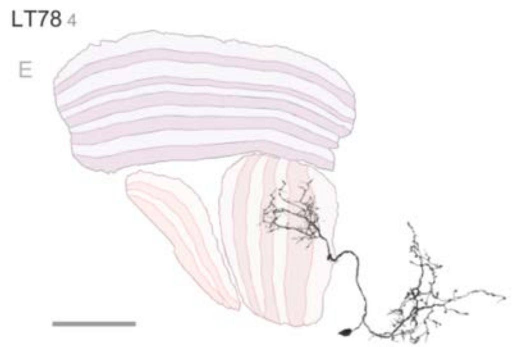


ARF, median size: 32.29

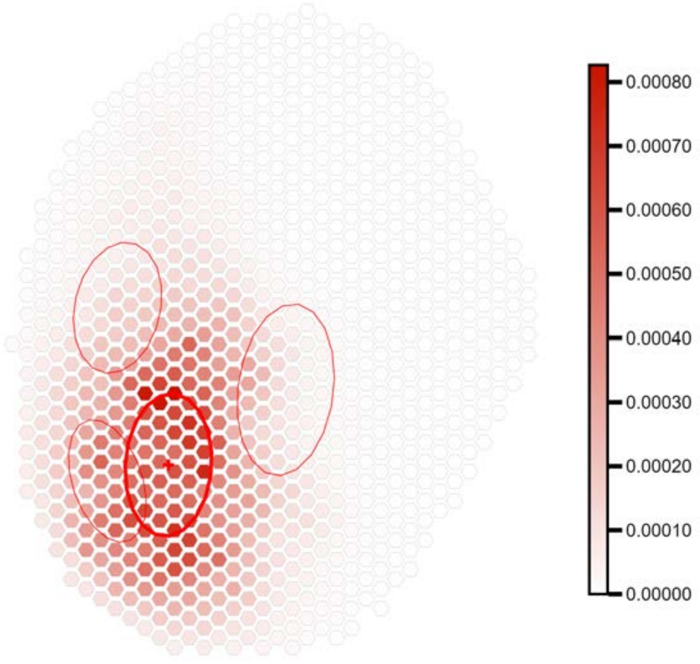


Max value projection, cell count: 4 - VIC: 0.098

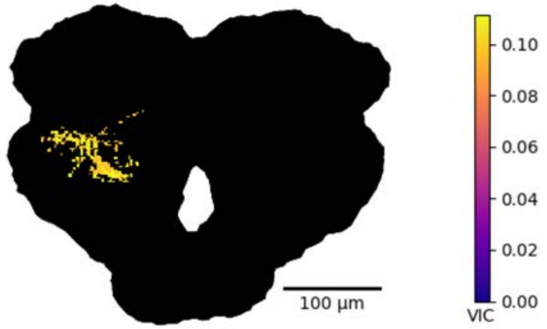


1-hop, cell count : 333

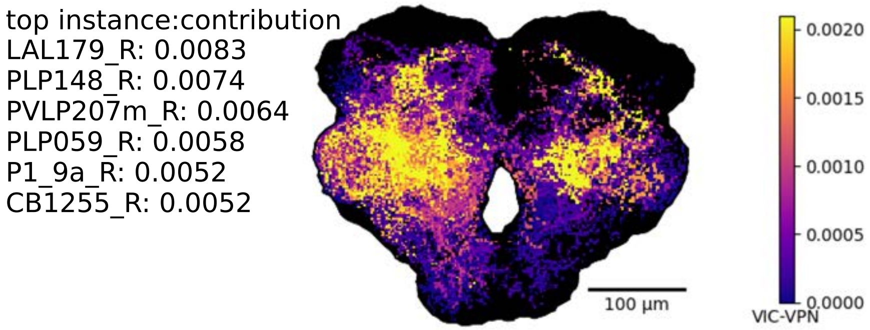


2-hop, cell count : 6691

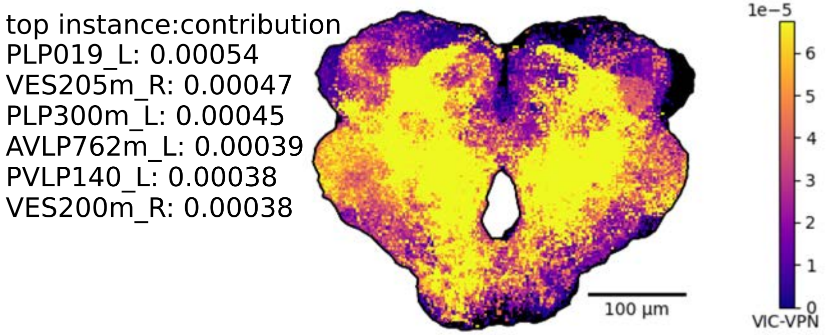

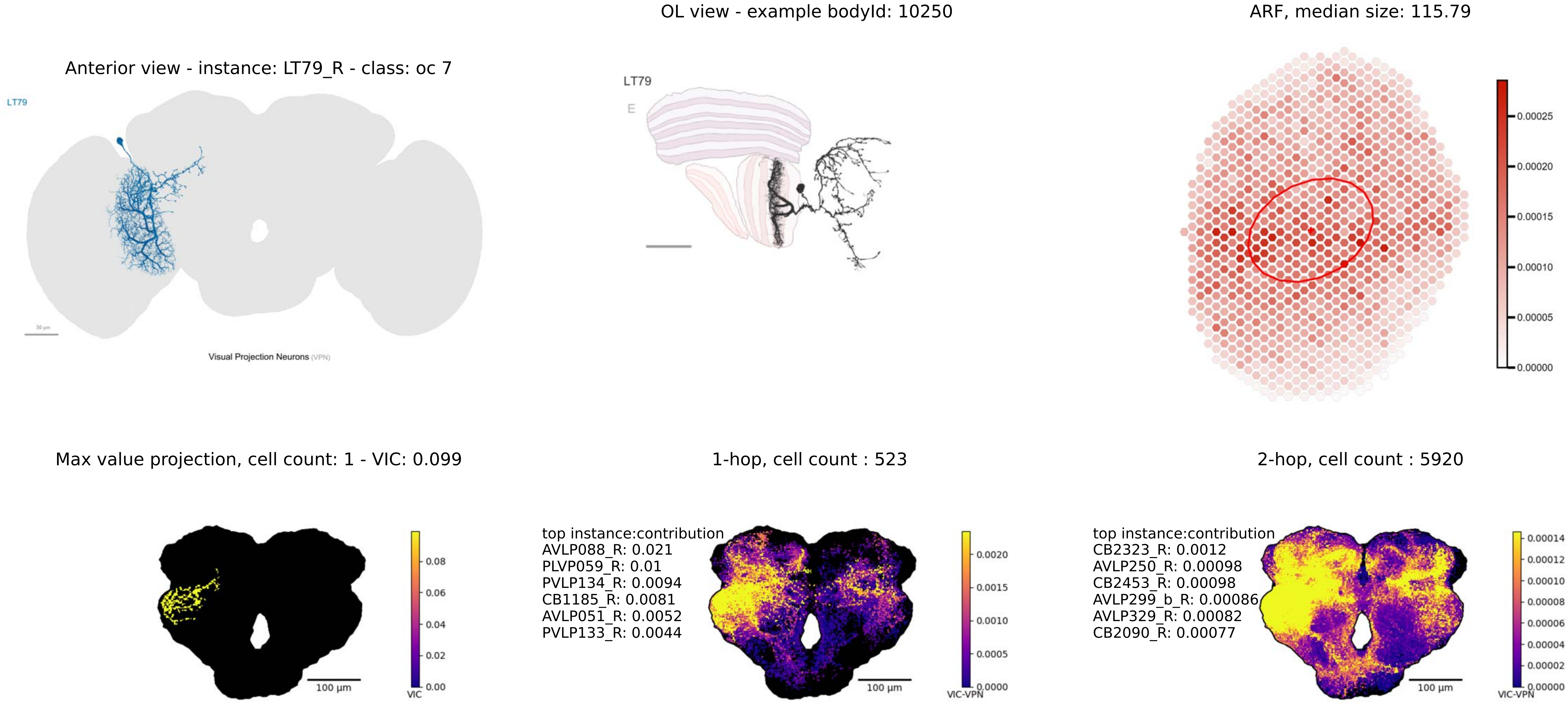

Anterior view - instance: LT79_R - class: oc 7
LT79
Visual Projection Neurons (VPN)
OL view - example bodyId: 10250
LT79
E
ARF, median size: 115.79
0.00025
0.00020
0.00015
0.00010
0.00005
0.00000
Max value projection, cell count: 1 - VIC: 0.099
100 μm
VIC
0.08
0.06
0.04
0.02
0.00
1-hop, cell count : 523
top instance:contribution
AVLP088_R: 0.021
PLVP059_R: 0.01
PVLP134_R: 0.0094
CB1185_R: 0.0081
AVLP051_R: 0.0052
PVLP133_R: 0.0044
100 μm
VIC-VPN
0.0020
0.0015
0.0010
0.0005
0.0000
2-hop, cell count : 5920
top instance:contribution
CB2323_R: 0.0012
AVLP250_R: 0.00098
CB2453_R: 0.00098
AVLP299_b_R: 0.00086
AVLP329_R: 0.00082
CB2090_R: 0.00077
100 μm
VIC-VPN
0.00014
0.00012
0.00010
0.00008
0.00006
0.00004
0.00002
0.00000

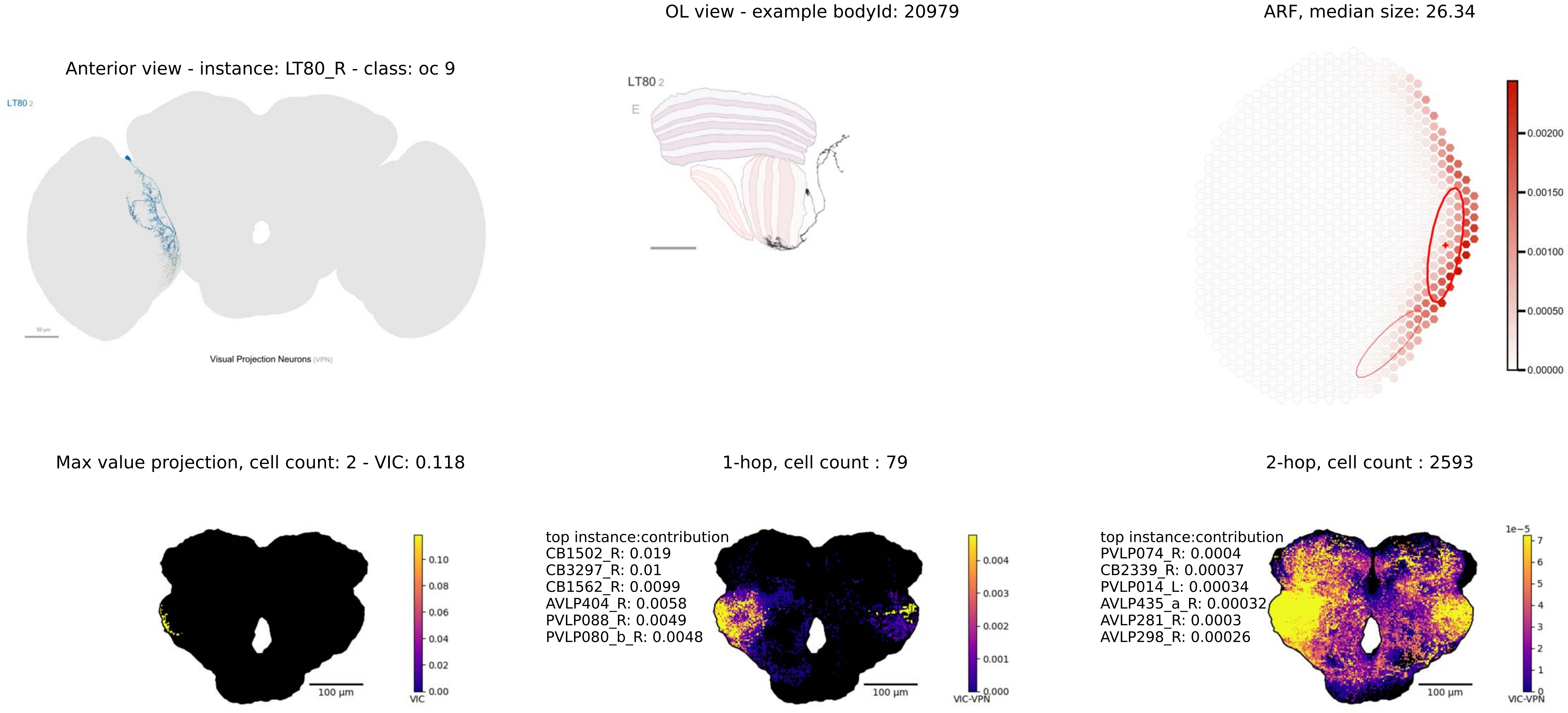
OL view - example bodyId: 20979
ARF, median size: 26.34
Anterior view - instance: LT80_R - class: oc 9
LT80 2
Visual Projection Neurons (VPN)
LT80 2
E
0.00200
0.00150
0.00100
0.00050
0.00000
Max value projection, cell count: 2 - VIC: 0.118
1-hop, cell count : 79
2-hop, cell count : 2593
0.10
0.08
0.06
0.04
0.02
0.00
VIC
100 µm
top instance:contribution
CB1502_R: 0.019
CB3297_R: 0.01
CB1562_R: 0.0099
AVLP404_R: 0.0058
PVLP088_R: 0.0049
PVLP080_b_R: 0.0048
0.004
0.003
0.002
0.001
0.000
VIC-VPN
100 µm
top instance:contribution
PVLP074_R: 0.0004
CB2339_R: 0.00037
PVLP014_L: 0.00034
AVLP435_a_R: 0.00032
AVLP281_R: 0.0003
AVLP298_R: 0.00026
1e−5
7
6
5
4
3
2
1
0
VIC-VPN
100 µm

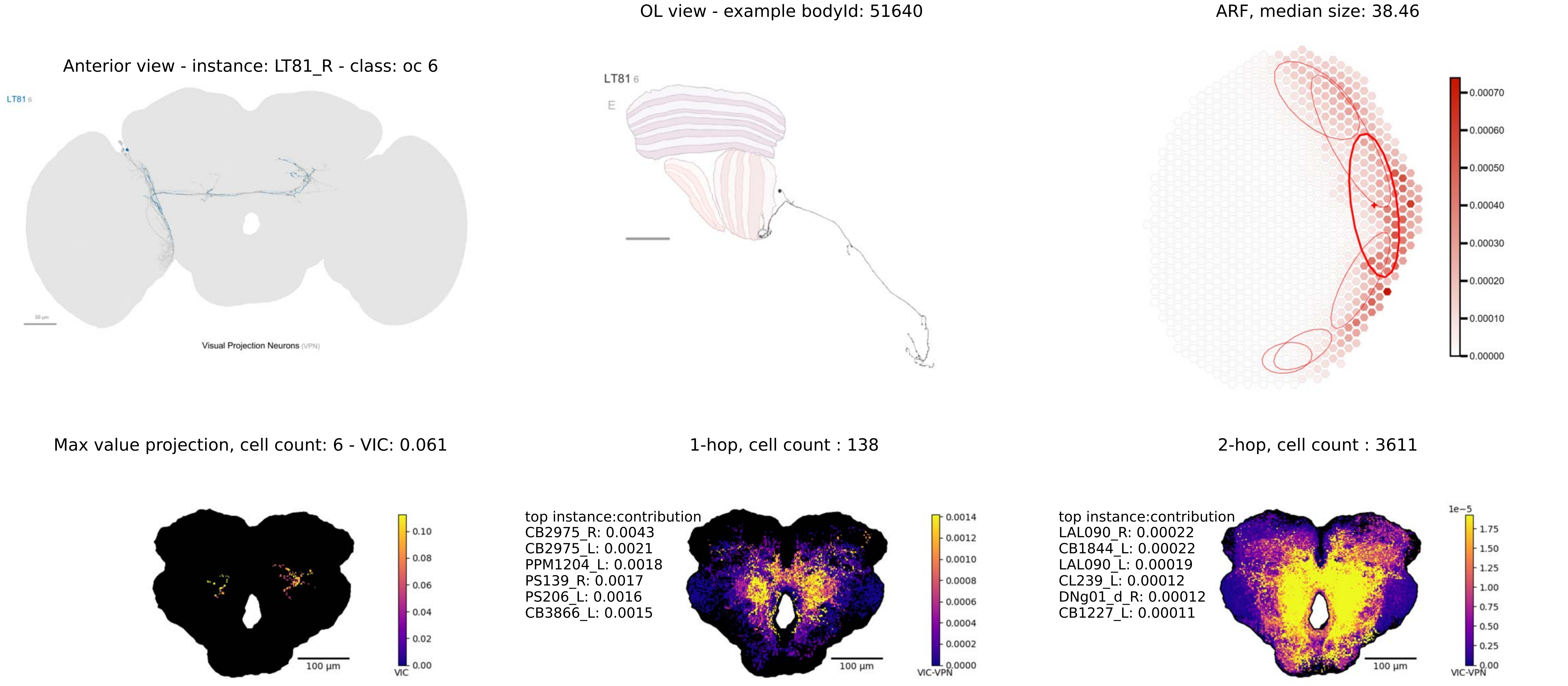
OL view - example bodyId: 51640
ARF, median size: 38.46
Anterior view - instance: LT81_R - class: oc 6
LT81 6
Visual Projection Neurons (VPN)
LT81 6
E
0.00070
0.00060
0.00050
0.00040
0.00030
0.00020
0.00010
0.00000
Max value projection, cell count: 6 - VIC: 0.061
1-hop, cell count : 138
2-hop, cell count : 3611
100 µm
0.10
0.08
0.06
0.04
0.02
0.00
VIC
top instance:contribution
CB2975_R: 0.0043
CB2975_L: 0.0021
PPM1204_L: 0.0018
PS139_R: 0.0017
PS206_L: 0.0016
CB3866_L: 0.0015
100 µm
0.0014
0.0012
0.0010
0.0008
0.0006
0.0004
0.0002
0.0000
VIC-VPN
top instance:contribution
LAL090_R: 0.00022
CB1844_L: 0.00022
LAL090_L: 0.00019
CL239_L: 0.00012
DNg01_d_R: 0.00012
CB1227_L: 0.00011
100 µm
1e−5
1.75
1.50
1.25
1.00
0.75
0.50
0.25
0.00
VIC-VPN

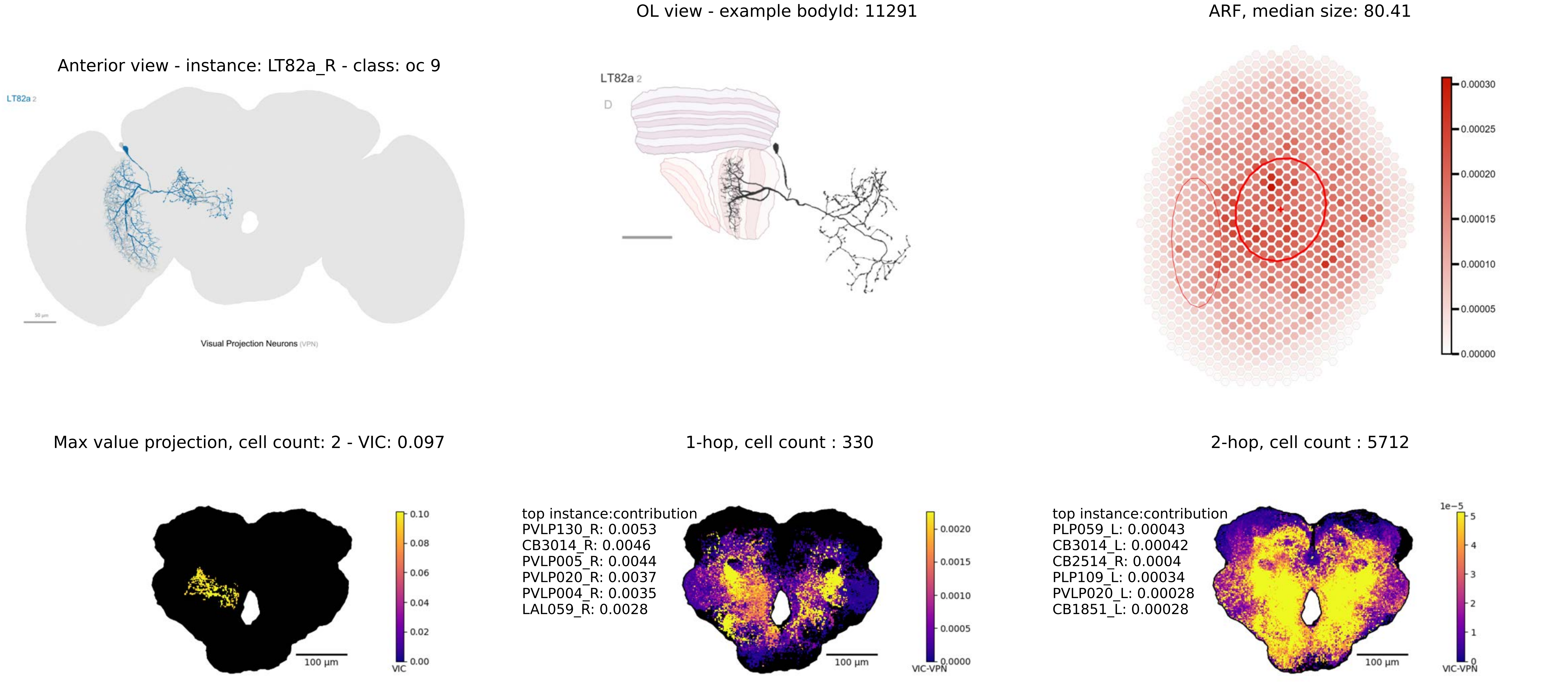
OL view - example bodyId: 11291
ARF, median size: 80.41
Anterior view - instance: LT82a_R - class: oc 9
LT82a 2
Visual Projection Neurons (VPN)
LT82a 2
D
0.00030
0.00025
0.00020
0.00015
0.00010
0.00005
0.00000
Max value projection, cell count: 2 - VIC: 0.097
1-hop, cell count : 330
2-hop, cell count : 5712
0.10
0.08
0.06
0.04
0.02
0.00
VIC
100 µm
top instance:contribution
PVLP130_R: 0.0053
CB3014_R: 0.0046
PVLP005_R: 0.0044
PVLP020_R: 0.0037
PVLP004_R: 0.0035
LAL059_R: 0.0028
0.0020
0.0015
0.0010
0.0005
0.0000
VIC-VPN
100 µm
top instance:contribution
PLP059_L: 0.00043
CB3014_L: 0.00042
CB2514_R: 0.0004
PLP109_L: 0.00034
PVLP020_L: 0.00028
CB1851_L: 0.00028
1e−5
5
4
3
2
1
0
VIC-VPN
100 µm

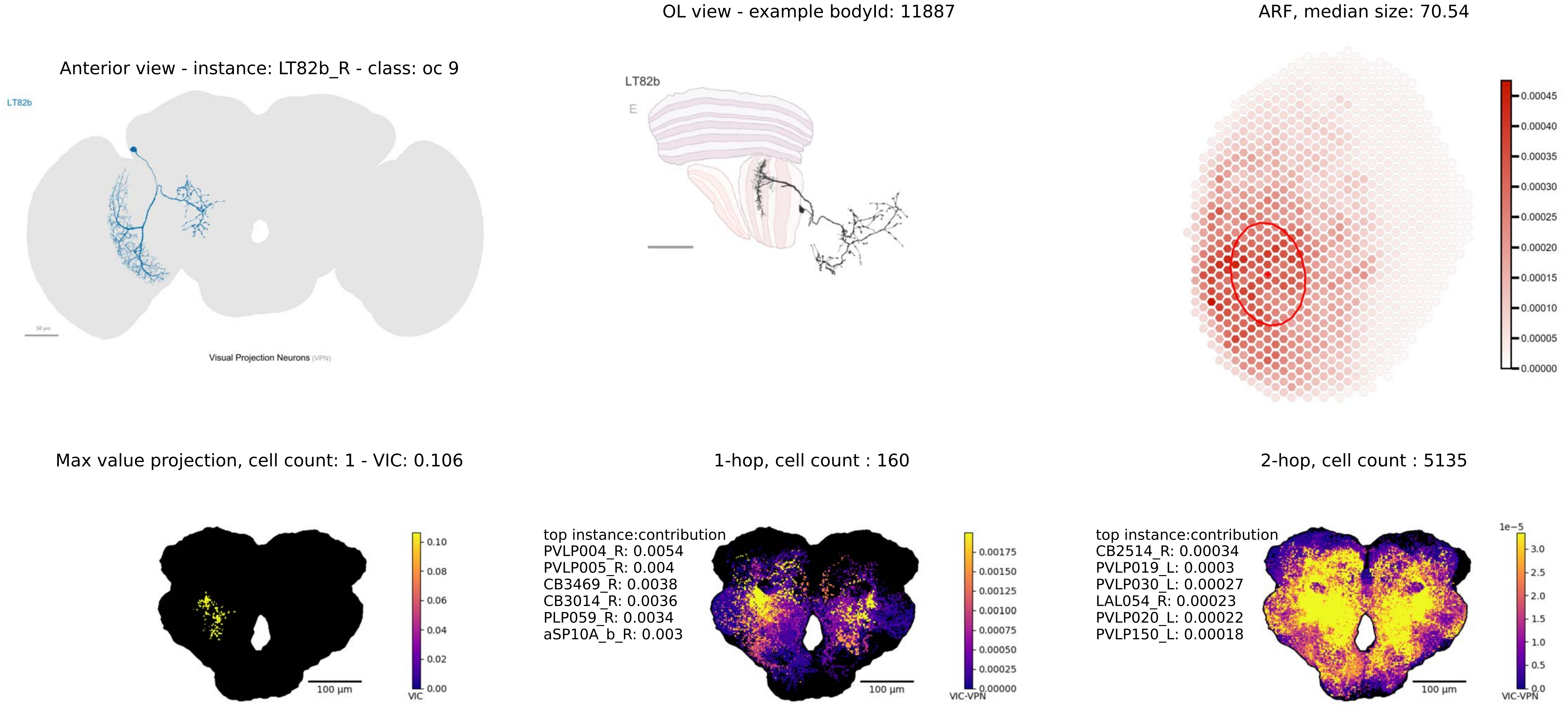

OL view - example bodyId: 11887
ARF, median size: 70.54
Anterior view - instance: LT82b_R - class: oc 9
LT82b
Visual Projection Neurons (VPN)
LT82b
E
Max value projection, cell count: 1 - VIC: 0.106
1-hop, cell count : 160
2-hop, cell count : 5135
100 µm
VIC
top instance:contribution
PVLP004_R: 0.0054
PVLP005_R: 0.004
CB3469_R: 0.0038
CB3014_R: 0.0036
PLP059_R: 0.0034
aSP10A_b_R: 0.003
VIC-VPN
top instance:contribution
CB2514_R: 0.00034
PVLP019_L: 0.0003
PVLP030_L: 0.00027
LAL054_R: 0.00023
PVLP020_L: 0.00022
PVLP150_L: 0.00018
1e−5

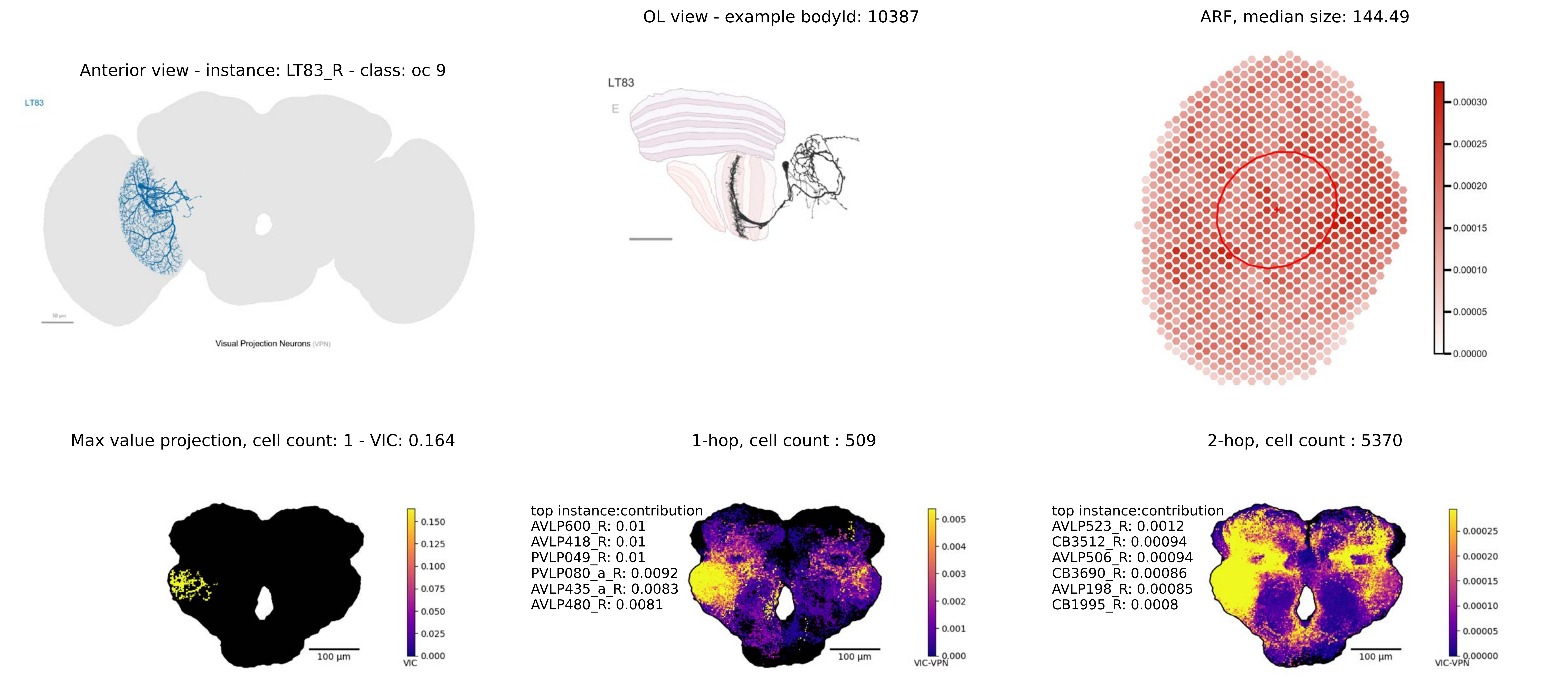
Anterior view - instance: LT83_R - class: oc 9
LT83
Visual Projection Neurons (VPN)
OL view - example bodyId: 10387
LT83
E
ARF, median size: 144.49
0.00030
0.00025
0.00020
0.00015
0.00010
0.00005
0.00000
Max value projection, cell count: 1 - VIC: 0.164
100 μm
0.150
0.125
0.100
0.075
0.050
0.025
0.000
VIC
1-hop, cell count : 509
top instance:contribution
AVLP600_R: 0.01
AVLP418_R: 0.01
PVLP049_R: 0.01
PVLP080_a_R: 0.0092
AVLP435_a_R: 0.0083
AVLP480_R: 0.0081
100 μm
0.005
0.004
0.003
0.002
0.001
0.000
VIC-VPN
2-hop, cell count : 5370
top instance:contribution
AVLP523_R: 0.0012
CB3512_R: 0.00094
AVLP506_R: 0.00094
CB3690_R: 0.00086
AVLP198_R: 0.00085
CB1995_R: 0.0008
100 μm
0.00025
0.00020
0.00015
0.00010
0.00005
0.00000
VIC-VPN

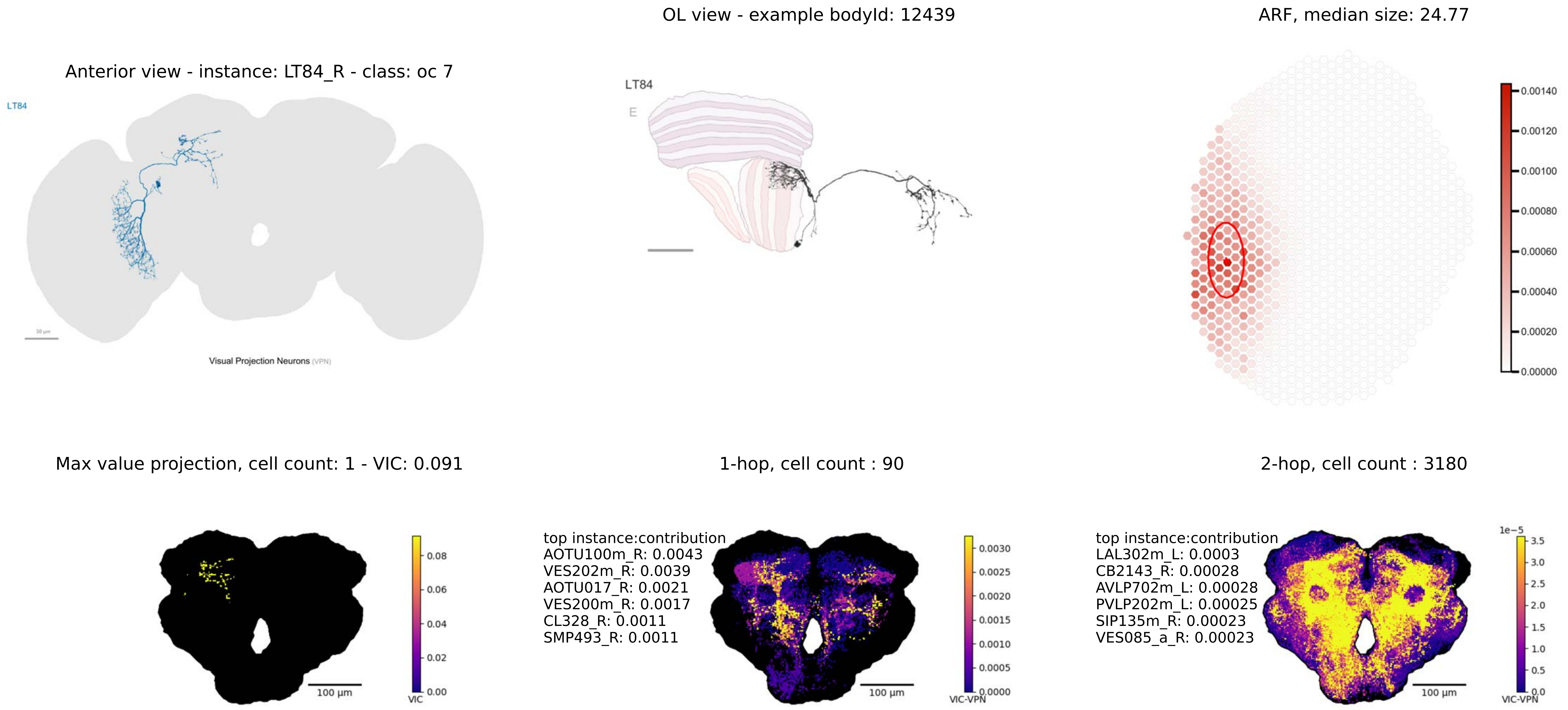
OL view - example bodyId: 12439
ARF, median size: 24.77
Anterior view - instance: LT84_R - class: oc 7
LT84
Visual Projection Neurons (VPN)
LT84
E
0.00140
0.00120
0.00100
0.00080
0.00060
0.00040
0.00020
0.00000
Max value projection, cell count: 1 - VIC: 0.091
1-hop, cell count : 90
2-hop, cell count : 3180
100 µm
VIC
0.08
0.06
0.04
0.02
0.00
top instance:contribution
AOTU100m_R: 0.0043
VES202m_R: 0.0039
AOTU017_R: 0.0021
VES200m_R: 0.0017
CL328_R: 0.0011
SMP493_R: 0.0011
100 µm
VIC-VPN
0.0030
0.0025
0.0020
0.0015
0.0010
0.0005
0.0000
top instance:contribution
LAL302m_L: 0.0003
CB2143_R: 0.00028
AVLP702m_L: 0.00028
PVLP202m_L: 0.00025
SIP135m_R: 0.00023
VES085_a_R: 0.00023
100 µm
1e−5
3.5
3.0
2.5
2.0
1.5
1.0
0.5
0.0
VIC-VPN

Anterior view - instance: LT85_R - class: oc 8

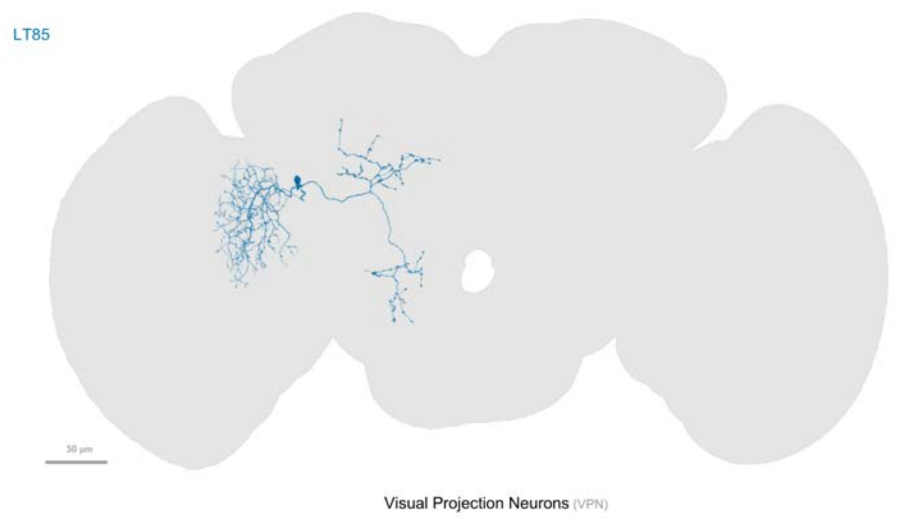


OL view - example bodyId: 19122

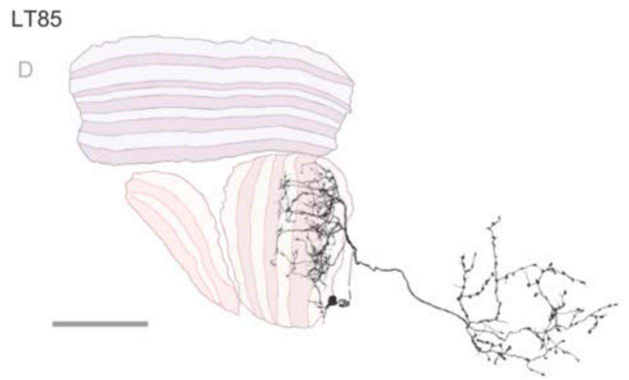


ARF, median size: 64.14

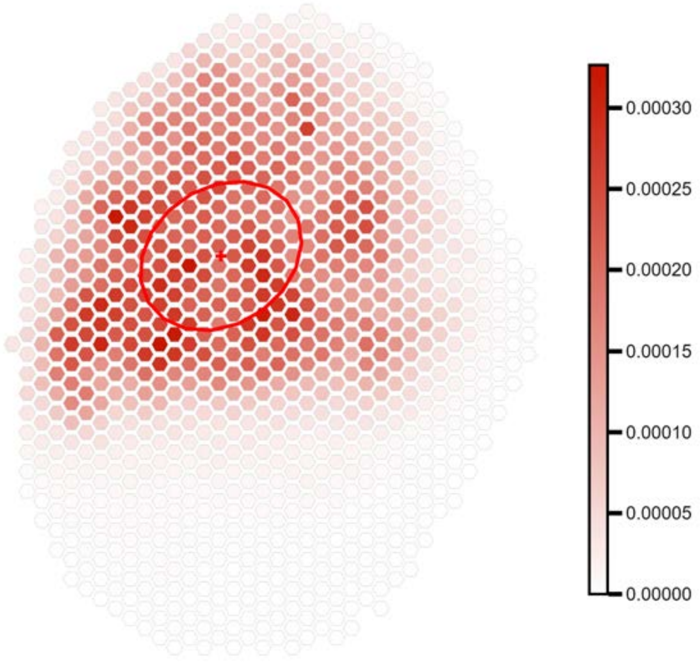


Max value projection, cell count: 1 - VIC: 0.077

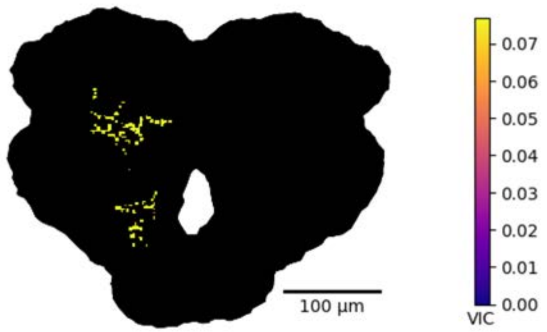


1-hop, cell count : 110

top instance:contribution
VES013_R: 0.0037
VES012_R: 0.0031
MBON20_R: 0.0012
GNG594_L: 0.0011
CB1985_R: 0.0011
DNp08_R: 0.0011

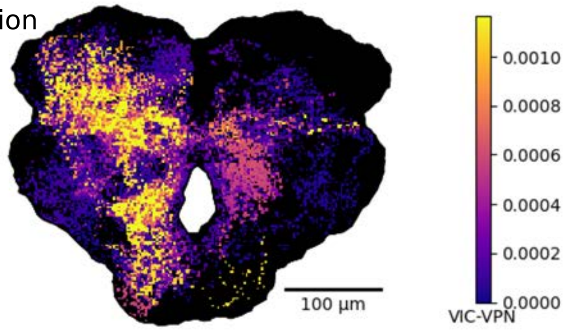


2-hop, cell count : 4536

top instance:contribution
PS305_R: 0.0003
DNge083_R: 0.0002
VES078_R: 0.00019
CL003_R: 0.00015
CB1853_R: 0.00013
PS173_R: 0.00013

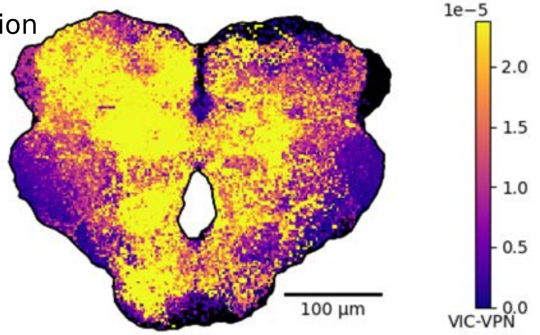

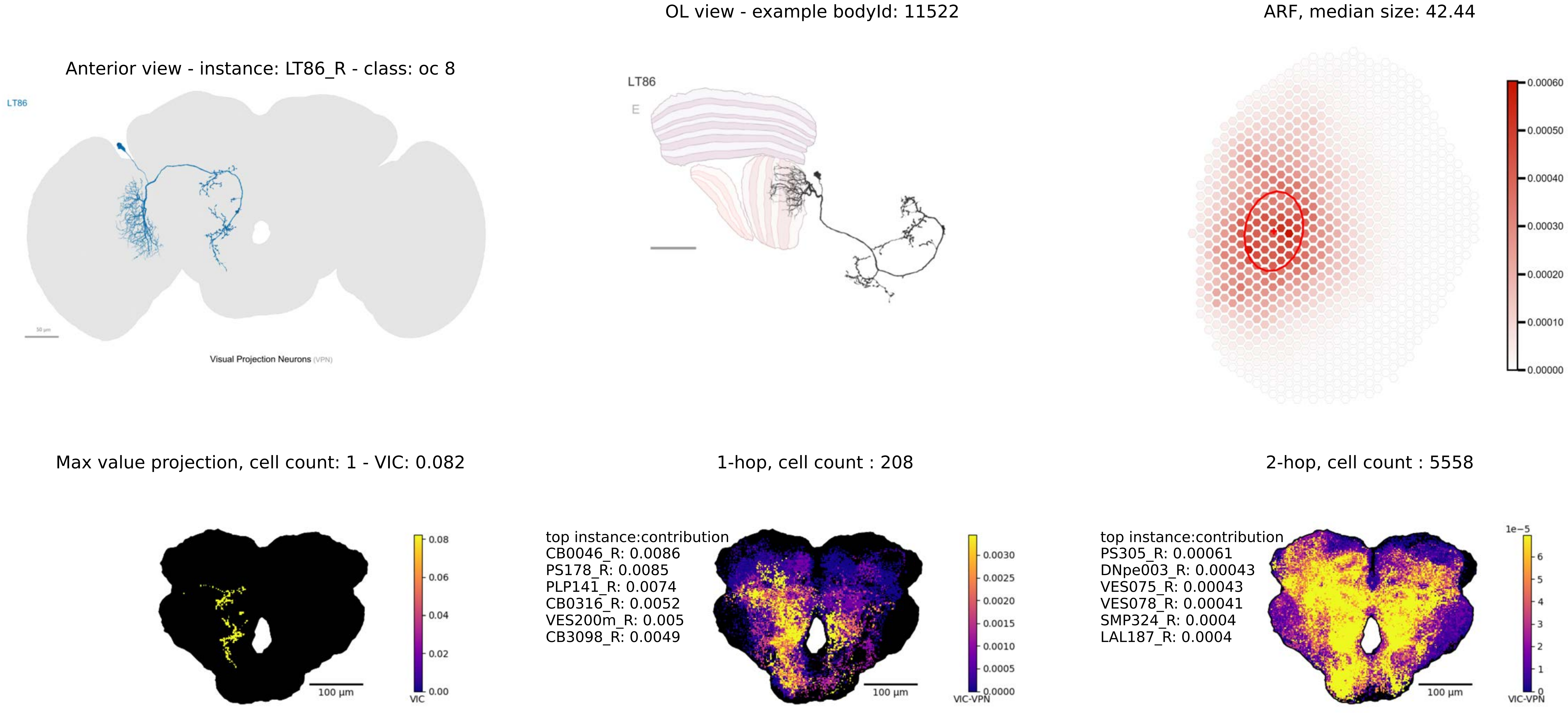

OL view - example bodyId: 11522
ARF, median size: 42.44
Anterior view - instance: LT86_R - class: oc 8
LT86
Visual Projection Neurons (VPN)
LT86
E
0.00060
0.00050
0.00040
0.00030
0.00020
0.00010
0.00000
Max value projection, cell count: 1 - VIC: 0.082
1-hop, cell count : 208
2-hop, cell count : 5558
0.08
0.06
0.04
0.02
0.00
VIC
100 µm
top instance:contribution
CB0046_R: 0.0086
PS178_R: 0.0085
PLP141_R: 0.0074
CB0316_R: 0.0052
VES200m_R: 0.005
CB3098_R: 0.0049
0.0030
0.0025
0.0020
0.0015
0.0010
0.0005
0.0000
VIC-VPN
100 µm
top instance:contribution
PS305_R: 0.00061
DNpe003_R: 0.00043
VES075_R: 0.00043
VES078_R: 0.00041
SMP324_R: 0.0004
LAL187_R: 0.0004
1e−5
6
5
4
3
2
1
0
VIC-VPN
100 µm

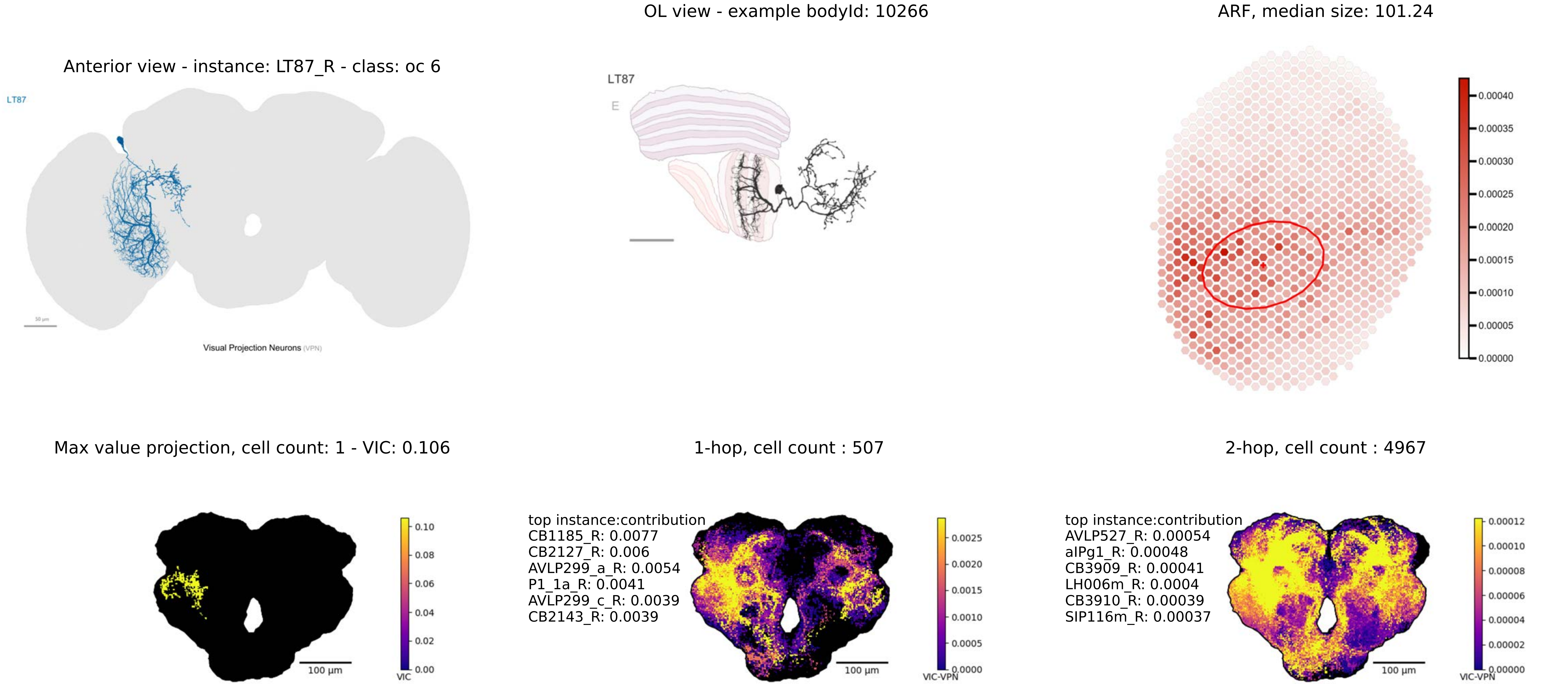
Anterior view - instance: LT87_R - class: oc 6
LT87
Visual Projection Neurons (VPN)
OL view - example bodyId: 10266
LT87
E
ARF, median size: 101.24
0.00040
0.00035
0.00030
0.00025
0.00020
0.00015
0.00010
0.00005
0.00000
Max value projection, cell count: 1 - VIC: 0.106
100 µm
0.10
0.08
0.06
0.04
0.02
0.00
VIC
1-hop, cell count : 507
top instance:contribution
CB1185_R: 0.0077
CB2127_R: 0.006
AVLP299_a_R: 0.0054
P1_1a_R: 0.0041
AVLP299_c_R: 0.0039
CB2143_R: 0.0039
100 µm
0.0025
0.0020
0.0015
0.0010
0.0005
0.0000
VIC-VPN
2-hop, cell count : 4967
top instance:contribution
AVLP527_R: 0.00054
aIPg1_R: 0.00048
CB3909_R: 0.00041
LH006m_R: 0.0004
CB3910_R: 0.00039
SIP116m_R: 0.00037
100 µm
0.00012
0.00010
0.00008
0.00006
0.00004
0.00002
0.00000
VIC-VPN

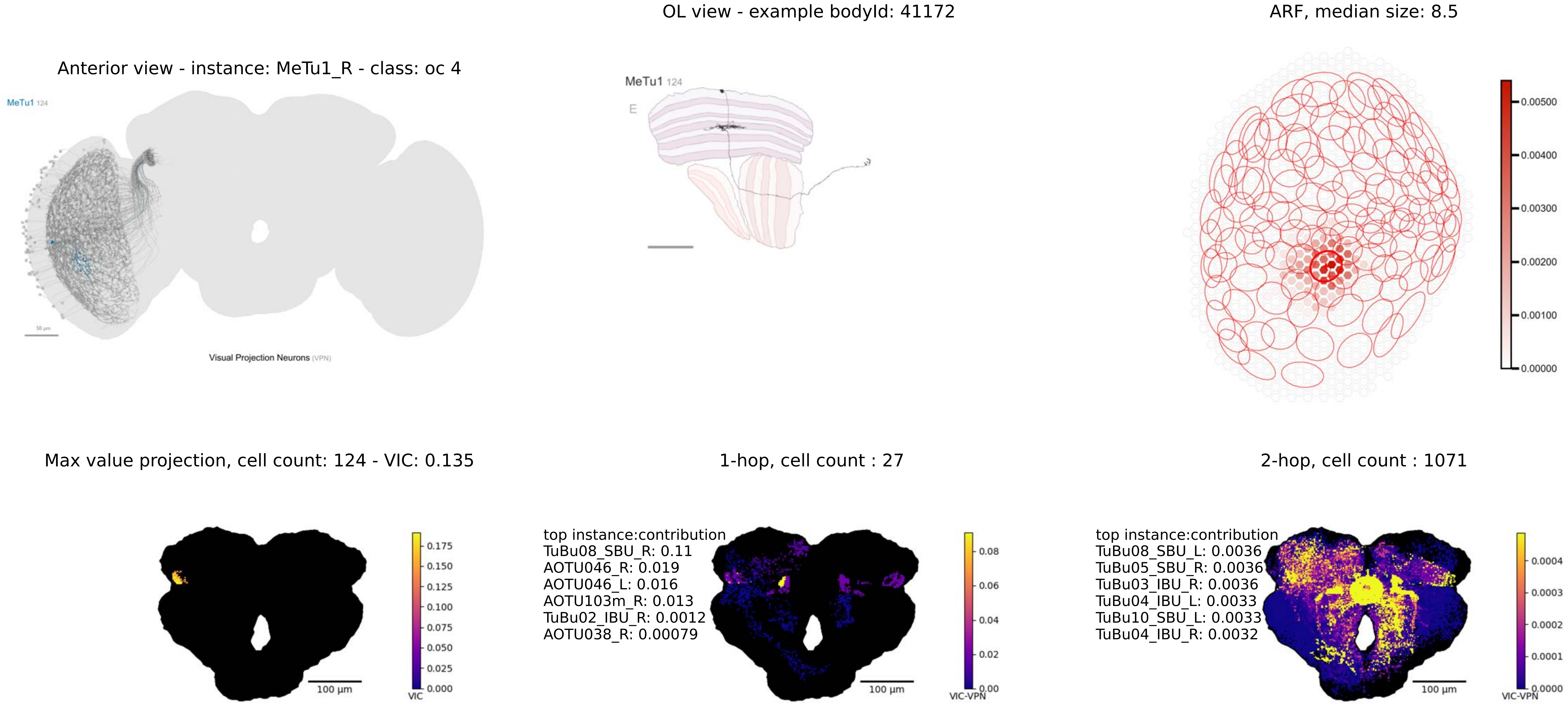

OL view - example bodyId: 41172
ARF, median size: 8.5
Anterior view - instance: MeTu1_R - class: oc 4
MeTu1 124
Visual Projection Neurons (VPN)
MeTu1 124
E
0.00500
0.00400
0.00300
0.00200
0.00100
0.00000
Max value projection, cell count: 124 - VIC: 0.135
1-hop, cell count : 27
2-hop, cell count : 1071
0.175
0.150
0.125
0.100
0.075
0.050
0.025
0.000
VIC
100 µm
top instance:contribution
TuBu08_SBU_R: 0.11
AOTU046_R: 0.019
AOTU046_L: 0.016
AOTU103m_R: 0.013
TuBu02_IBU_R: 0.0012
AOTU038_R: 0.00079
0.08
0.06
0.04
0.02
0.00
VIC-VPN
100 µm
top instance:contribution
TuBu08_SBU_L: 0.0036
TuBu05_SBU_R: 0.0036
TuBu03_IBU_R: 0.0036
TuBu04_IBU_L: 0.0033
TuBu10_SBU_L: 0.0033
TuBu04_IBU_R: 0.0032
0.0004
0.0003
0.0002
0.0001
0.0000
VIC-VPN
100 µm

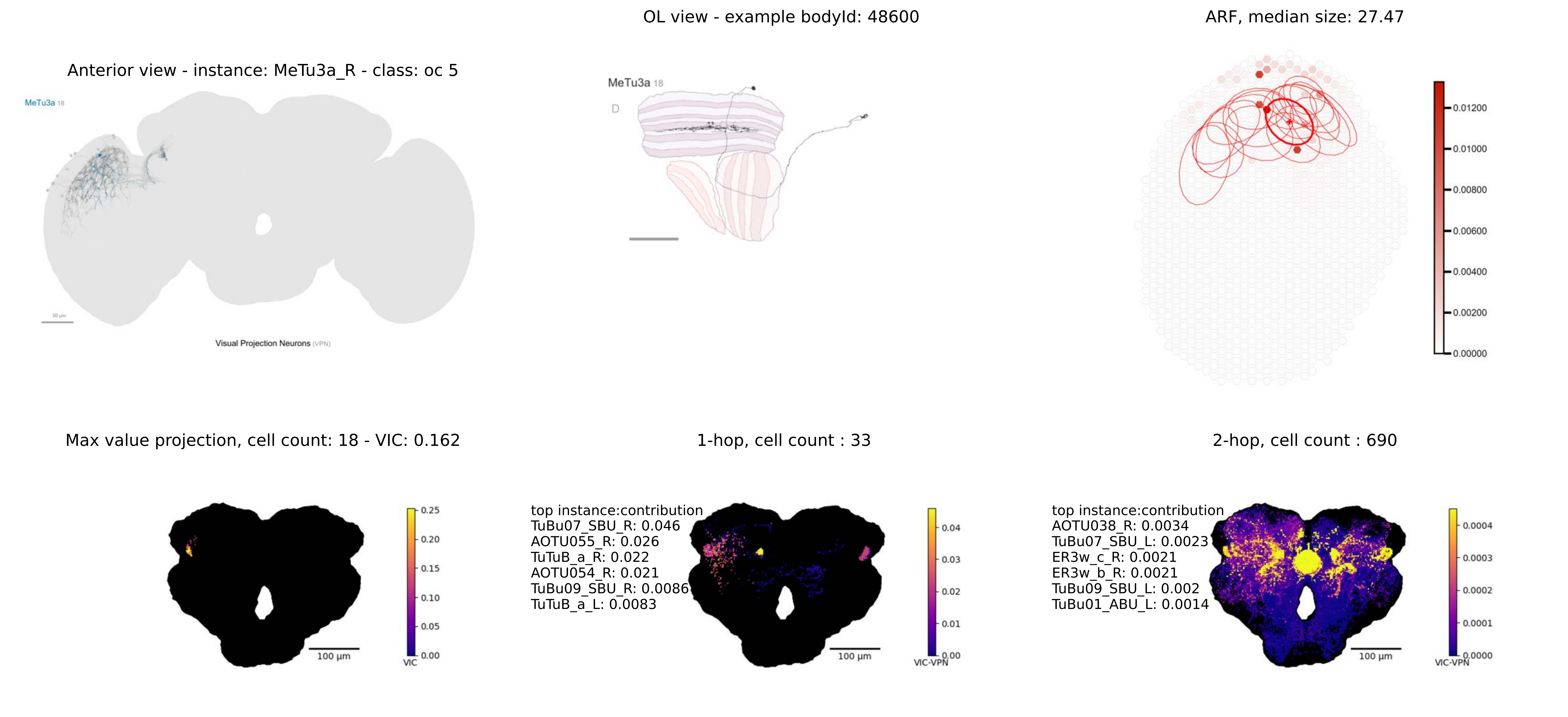

OL view - example bodyId: 48600
ARF, median size: 27.47
Anterior view - instance: MeTu3a_R - class: oc 5
MeTu3a 18
Visual Projection Neurons (VPN)
MeTu3a 18
D
0.01200
0.01000
0.00800
0.00600
0.00400
0.00200
0.00000
Max value projection, cell count: 18 - VIC: 0.162
1-hop, cell count : 33
2-hop, cell count : 690
0.25
0.20
0.15
0.10
0.05
0.00
VIC
100 µm
top instance:contribution
TuBu07_SBU_R: 0.046
AOTU055_R: 0.026
TuTuB_a_R: 0.022
AOTU054_R: 0.021
TuBu09_SBU_R: 0.0086
TuTuB_a_L: 0.0083
0.04
0.03
0.02
0.01
0.00
VIC-VPN
100 µm
top instance:contribution
AOTU038_R: 0.0034
TuBu07_SBU_L: 0.0023
ER3w_c_R: 0.0021
ER3w_b_R: 0.0021
TuBu09_SBU_L: 0.002
TuBu01_ABU_L: 0.0014
0.0004
0.0003
0.0002
0.0001
0.0000
VIC-VPN
100 µm

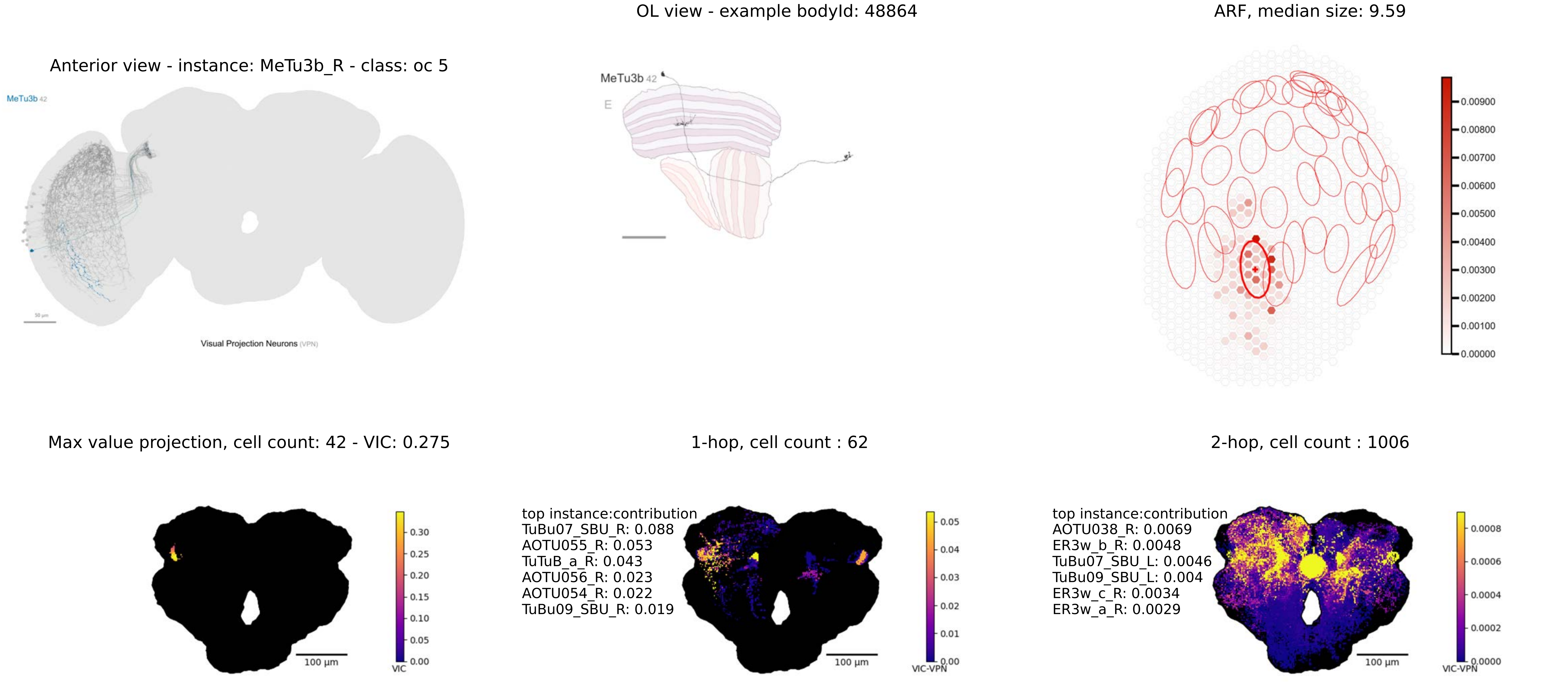

OL view - example bodyId: 48864
ARF, median size: 9.59
Anterior view - instance: MeTu3b_R - class: oc 5
MeTu3b 42
Visual Projection Neurons (VPN)
MeTu3b 42
E
0.00900
0.00800
0.00700
0.00600
0.00500
0.00400
0.00300
0.00200
0.00100
0.00000
Max value projection, cell count: 42 - VIC: 0.275
1-hop, cell count : 62
2-hop, cell count : 1006
100 µm
VIC
0.30
0.25
0.20
0.15
0.10
0.05
0.00
top instance:contribution
TuBu07_SBU_R: 0.088
AOTU055_R: 0.053
TuTuB_a_R: 0.043
AOTU056_R: 0.023
AOTU054_R: 0.022
TuBu09_SBU_R: 0.019
100 µm
VIC-VPN
0.05
0.04
0.03
0.02
0.01
0.00
top instance:contribution
AOTU038_R: 0.0069
ER3w_b_R: 0.0048
TuBu07_SBU_L: 0.0046
TuBu09_SBU_L: 0.004
ER3w_c_R: 0.0034
ER3w_a_R: 0.0029
100 µm
VIC-VPN
0.0008
0.0006
0.0004
0.0002
0.0000

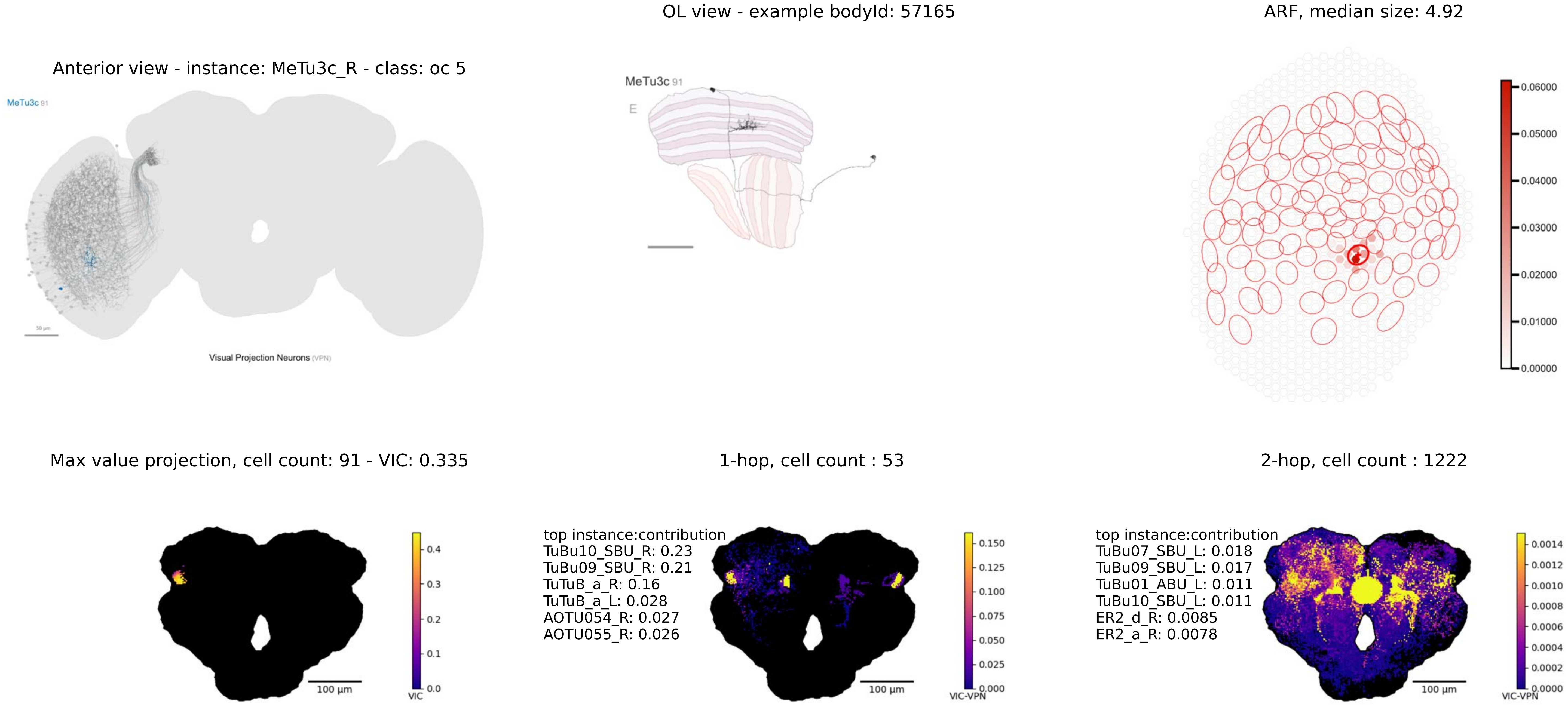

OL view - example bodyId: 57165
ARF, median size: 4.92
Anterior view - instance: MeTu3c_R - class: oc 5
MeTu3c 91
Visual Projection Neurons (VPN)
MeTu3c 91
E
Max value projection, cell count: 91 - VIC: 0.335
1-hop, cell count : 53
2-hop, cell count : 1222
top instance:contribution
TuBu10_SBU_R: 0.23
TuBu09_SBU_R: 0.21
TuTuB_a_R: 0.16
TuTuB_a_L: 0.028
AOTU054_R: 0.027
AOTU055_R: 0.026
top instance:contribution
TuBu07_SBU_L: 0.018
TuBu09_SBU_L: 0.017
TuBu01_ABU_L: 0.011
TuBu10_SBU_L: 0.011
ER2_d_R: 0.0085
ER2_a_R: 0.0078
100 µm
VIC
VIC-VPN

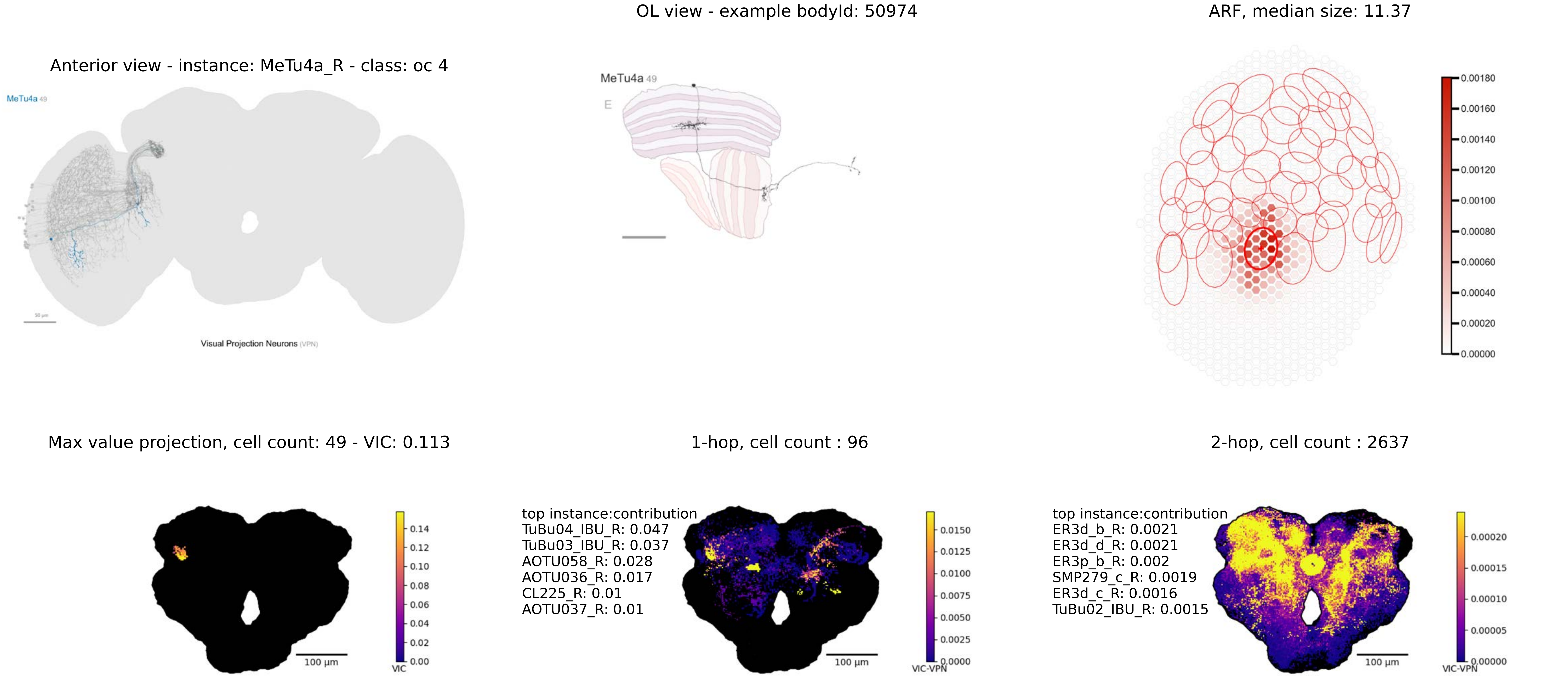

OL view - example bodyId: 50974
ARF, median size: 11.37
Anterior view - instance: MeTu4a_R - class: oc 4
MeTu4a 49
Visual Projection Neurons (VPN)
MeTu4a 49
E
0.00180
0.00160
0.00140
0.00120
0.00100
0.00080
0.00060
0.00040
0.00020
0.00000
Max value projection, cell count: 49 - VIC: 0.113
1-hop, cell count : 96
2-hop, cell count : 2637
100 µm
VIC
0.14
0.12
0.10
0.08
0.06
0.04
0.02
0.00
top instance:contribution
TuBu04_IBU_R: 0.047
TuBu03_IBU_R: 0.037
AOTU058_R: 0.028
AOTU036_R: 0.017
CL225_R: 0.01
AOTU037_R: 0.01
100 µm
VIC-VPN
0.0150
0.0125
0.0100
0.0075
0.0050
0.0025
0.0000
top instance:contribution
ER3d_b_R: 0.0021
ER3d_d_R: 0.0021
ER3p_b_R: 0.002
SMP279_c_R: 0.0019
ER3d_c_R: 0.0016
TuBu02_IBU_R: 0.0015
100 µm
VIC-VPN
0.00020
0.00015
0.00010
0.00005
0.00000

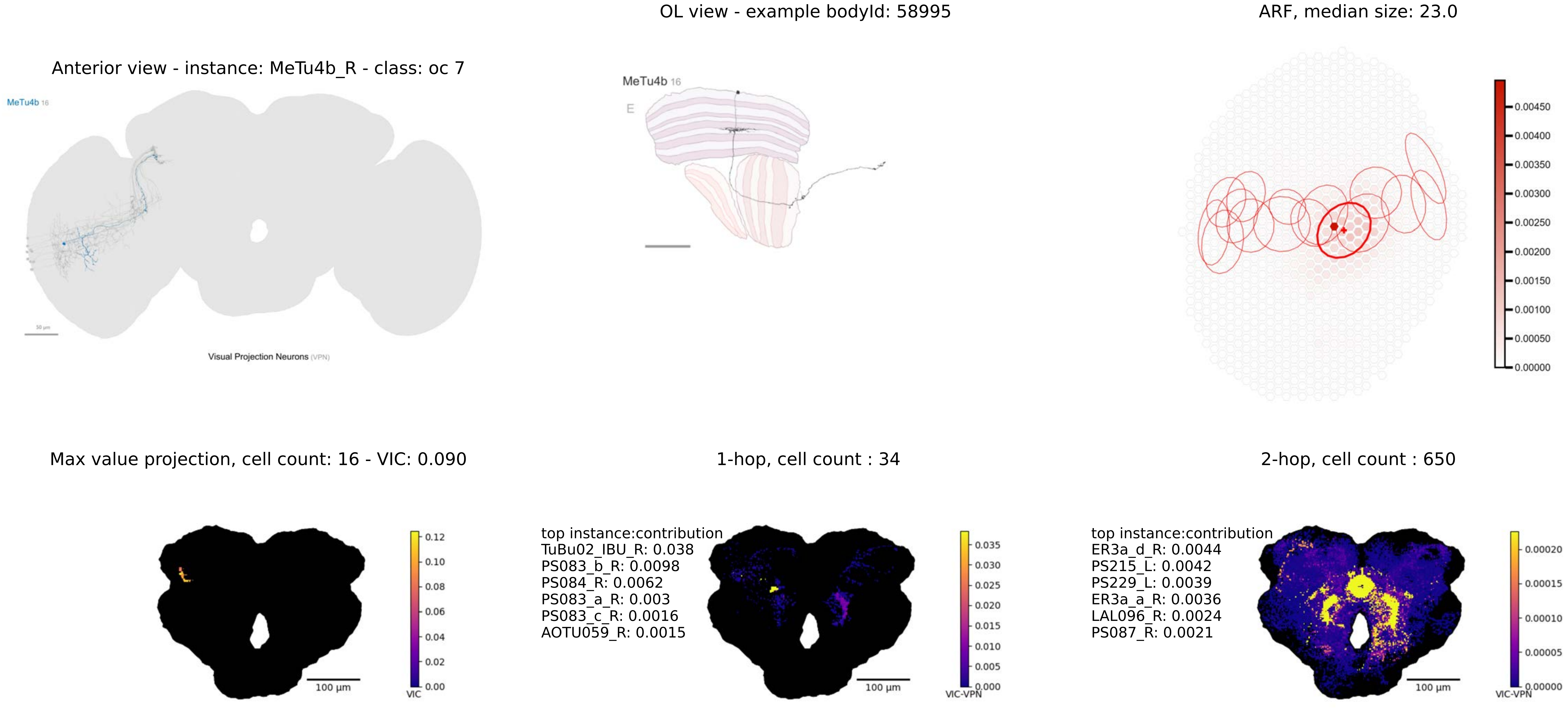

OL view - example bodyId: 58995
ARF, median size: 23.0
Anterior view - instance: MeTu4b_R - class: oc 7
MeTu4b 16
Visual Projection Neurons (VPN)
MeTu4b 16
E
0.00450
0.00400
0.00350
0.00300
0.00250
0.00200
0.00150
0.00100
0.00050
0.00000
Max value projection, cell count: 16 - VIC: 0.090
1-hop, cell count : 34
2-hop, cell count : 650
0.12
0.10
0.08
0.06
0.04
0.02
0.00
VIC
100 μm
top instance:contribution
TuBu02_IBU_R: 0.038
PS083_b_R: 0.0098
PS084_R: 0.0062
PS083_a_R: 0.003
PS083_c_R: 0.0016
AOTU059_R: 0.0015
0.035
0.030
0.025
0.020
0.015
0.010
0.005
0.000
VIC-VPN
100 μm
top instance:contribution
ER3a_d_R: 0.0044
PS215_L: 0.0042
PS229_L: 0.0039
ER3a_a_R: 0.0036
LAL096_R: 0.0024
PS087_R: 0.0021
0.00020
0.00015
0.00010
0.00005
0.00000
VIC-VPN
100 μm

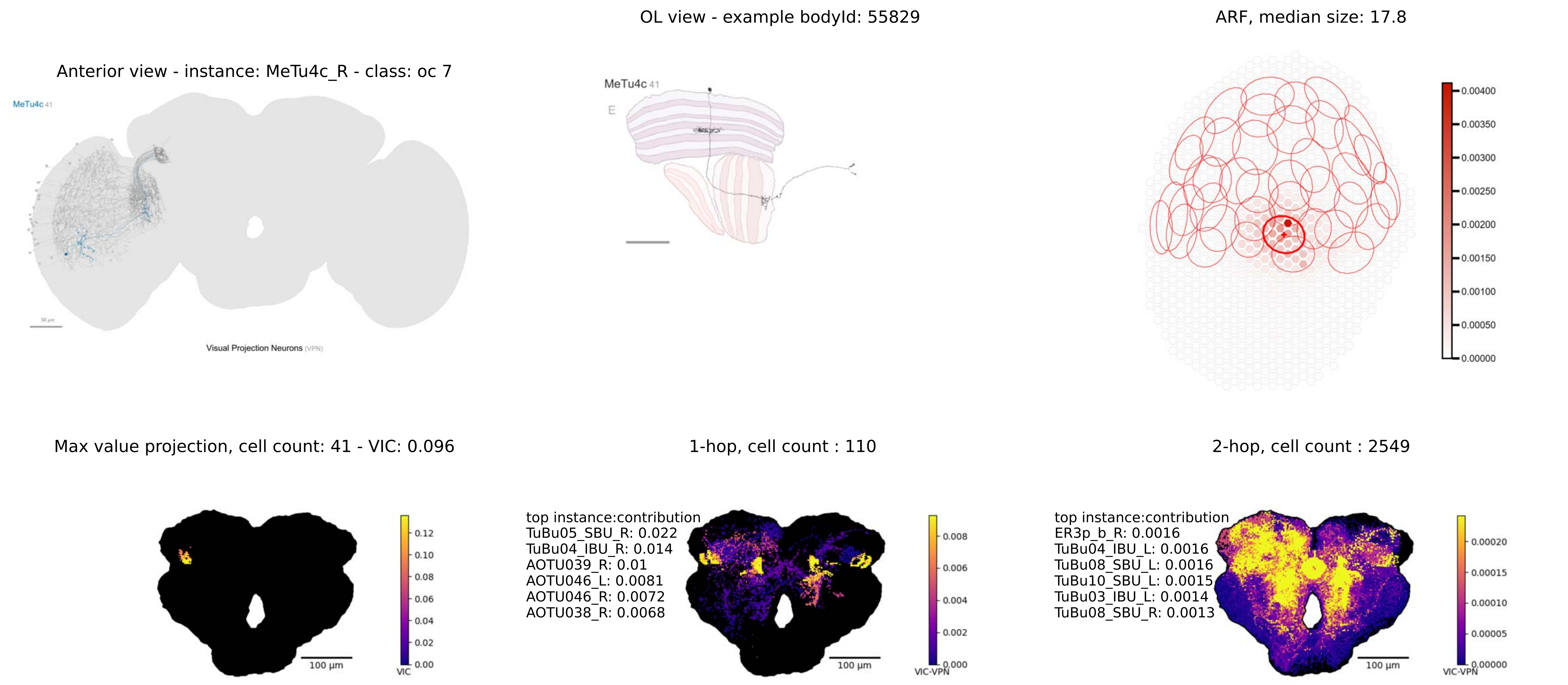
Anterior view - instance: MeTu4c_R - class: oc 7
MeTu4c 41
Visual Projection Neurons (VPN)
OL view - example bodyId: 55829
MeTu4c 41
E
ARF, median size: 17.8
0.00400
0.00350
0.00300
0.00250
0.00200
0.00150
0.00100
0.00050
0.00000
Max value projection, cell count: 41 - VIC: 0.096
0.12
0.10
0.08
0.06
0.04
0.02
0.00
VIC
100 µm
1-hop, cell count : 110
top instance:contribution
TuBu05_SBU_R: 0.022
TuBu04_IBU_R: 0.014
AOTU039_R: 0.01
AOTU046_L: 0.0081
AOTU046_R: 0.0072
AOTU038_R: 0.0068
0.008
0.006
0.004
0.002
0.000
VIC-VPN
100 µm
2-hop, cell count : 2549
top instance:contribution
ER3p_b_R: 0.0016
TuBu04_IBU_L: 0.0016
TuBu08_SBU_L: 0.0016
TuBu10_SBU_L: 0.0015
TuBu03_IBU_L: 0.0014
TuBu08_SBU_R: 0.0013
0.00020
0.00015
0.00010
0.00005
0.00000
VIC-VPN
100 µm

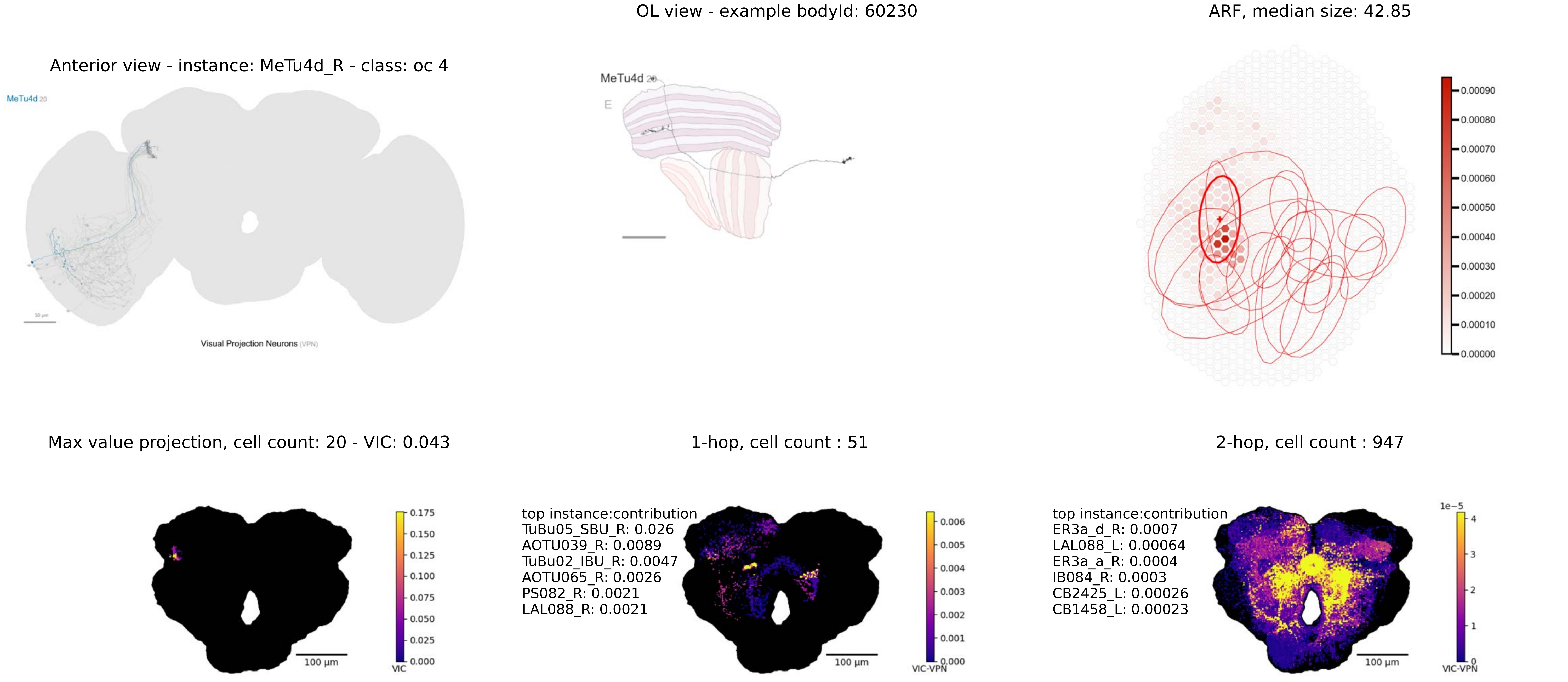
OL view - example bodyId: 60230
ARF, median size: 42.85
Anterior view - instance: MeTu4d_R - class: oc 4
MeTu4d 20
MeTu4d 20
E
Visual Projection Neurons (VPN)
0.00090
0.00080
0.00070
0.00060
0.00050
0.00040
0.00030
0.00020
0.00010
0.00000
Max value projection, cell count: 20 - VIC: 0.043
1-hop, cell count : 51
2-hop, cell count : 947
0.175
0.150
0.125
0.100
0.075
0.050
0.025
0.000
VIC
100 µm
top instance:contribution
TuBu05_SBU_R: 0.026
AOTU039_R: 0.0089
TuBu02_IBU_R: 0.0047
AOTU065_R: 0.0026
PS082_R: 0.0021
LAL088_R: 0.0021
0.006
0.005
0.004
0.003
0.002
0.001
0.000
VIC-VPN
100 µm
top instance:contribution
ER3a_d_R: 0.0007
LAL088_L: 0.00064
ER3a_a_R: 0.0004
IB084_R: 0.0003
CB2425_L: 0.00026
CB1458_L: 0.00023
1e−5
4
3
2
1
0
VIC-VPN
100 µm

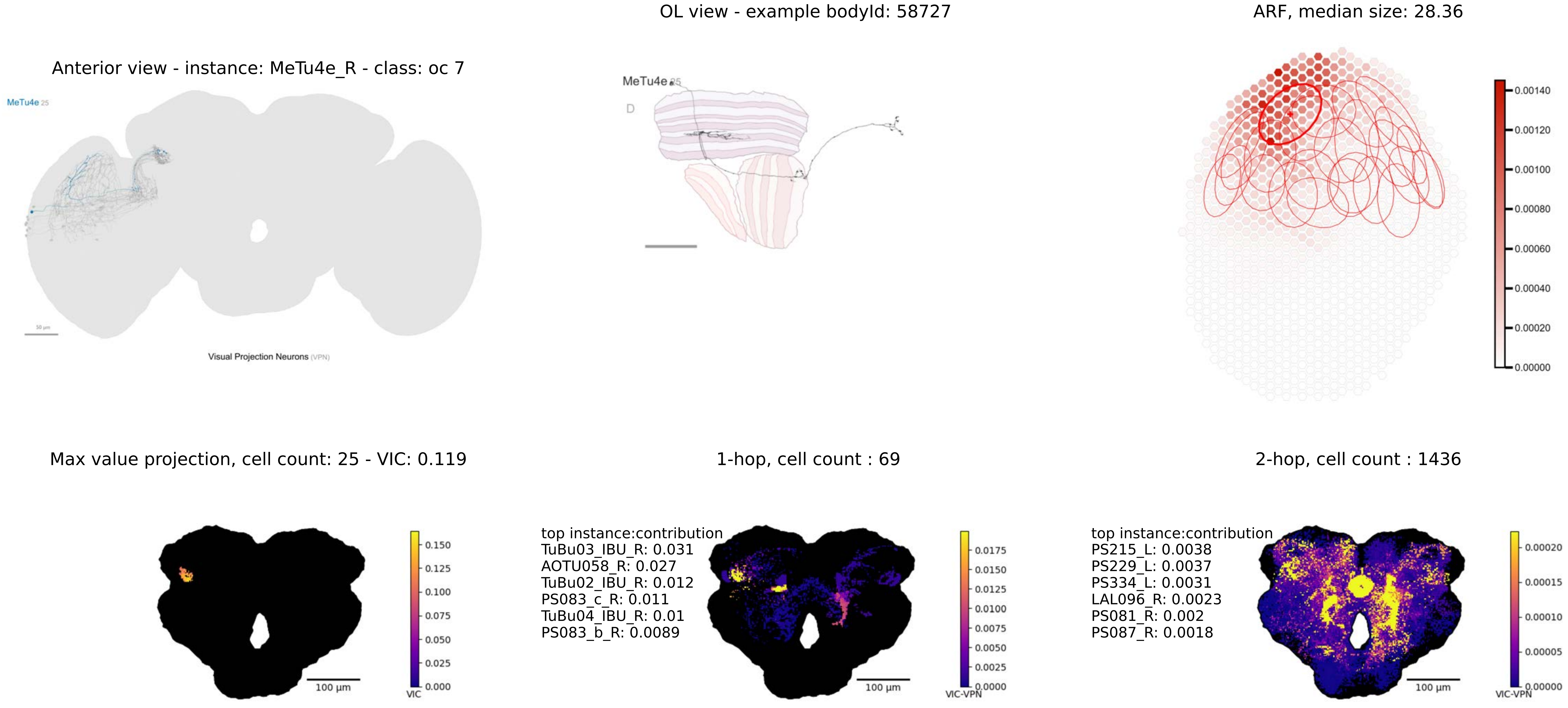

Anterior view - instance: MeTu4e_R - class: oc 7
MeTu4e 25
Visual Projection Neurons (VPN)
OL view - example bodyId: 58727
MeTu4e 25
D
ARF, median size: 28.36
0.00140
0.00120
0.00100
0.00080
0.00060
0.00040
0.00020
0.00000
Max value projection, cell count: 25 - VIC: 0.119
100 µm
0.150
0.125
0.100
0.075
0.050
0.025
0.000
VIC
1-hop, cell count : 69
top instance:contribution
TuBu03_IBU_R: 0.031
AOTU058_R: 0.027
TuBu02_IBU_R: 0.012
PS083_c_R: 0.011
TuBu04_IBU_R: 0.01
PS083_b_R: 0.0089
100 µm
0.0175
0.0150
0.0125
0.0100
0.0075
0.0050
0.0025
0.0000
VIC-VPN
2-hop, cell count : 1436
top instance:contribution
PS215_L: 0.0038
PS229_L: 0.0037
PS334_L: 0.0031
LAL096_R: 0.0023
PS081_R: 0.002
PS087_R: 0.0018
100 µm
0.00020
0.00015
0.00010
0.00005
0.00000
VIC-VPN

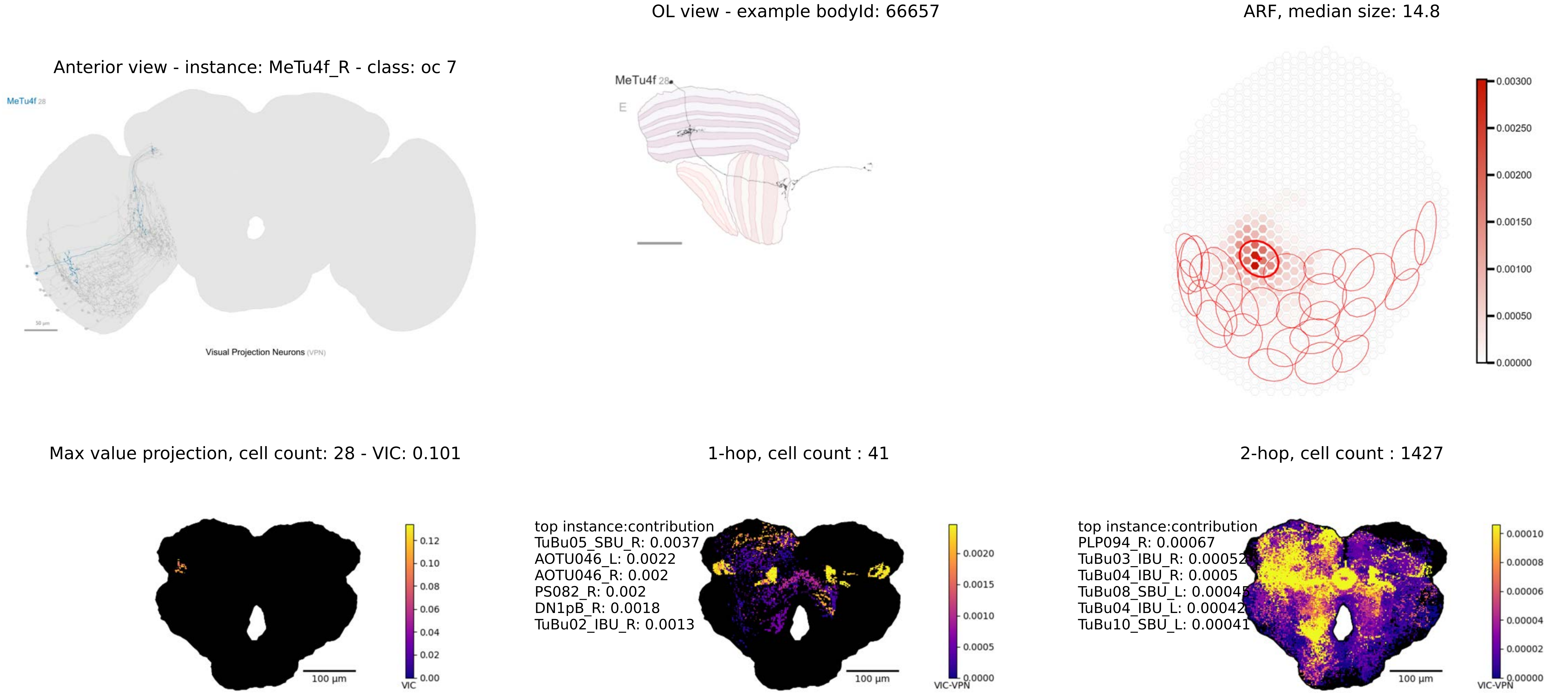

OL view - example bodyId: 66657
ARF, median size: 14.8
Anterior view - instance: MeTu4f_R - class: oc 7
MeTu4f 28
Visual Projection Neurons (VPN)
MeTu4f 28
E
0.00300
0.00250
0.00200
0.00150
0.00100
0.00050
0.00000
Max value projection, cell count: 28 - VIC: 0.101
1-hop, cell count : 41
2-hop, cell count : 1427
100 µm
0.12
0.10
0.08
0.06
0.04
0.02
0.00
VIC
top instance:contribution
TuBu05_SBU_R: 0.0037
AOTU046_L: 0.0022
AOTU046_R: 0.002
PS082_R: 0.002
DN1pB_R: 0.0018
TuBu02_IBU_R: 0.0013
100 µm
0.0020
0.0015
0.0010
0.0005
0.0000
VIC-VPN
top instance:contribution
PLP094_R: 0.00067
TuBu03_IBU_R: 0.00052
TuBu04_IBU_R: 0.0005
TuBu08_SBU_L: 0.00045
TuBu04_IBU_L: 0.00042
TuBu10_SBU_L: 0.00041
100 µm
0.00010
0.00008
0.00006
0.00004
0.00002
0.00000
VIC-VPN

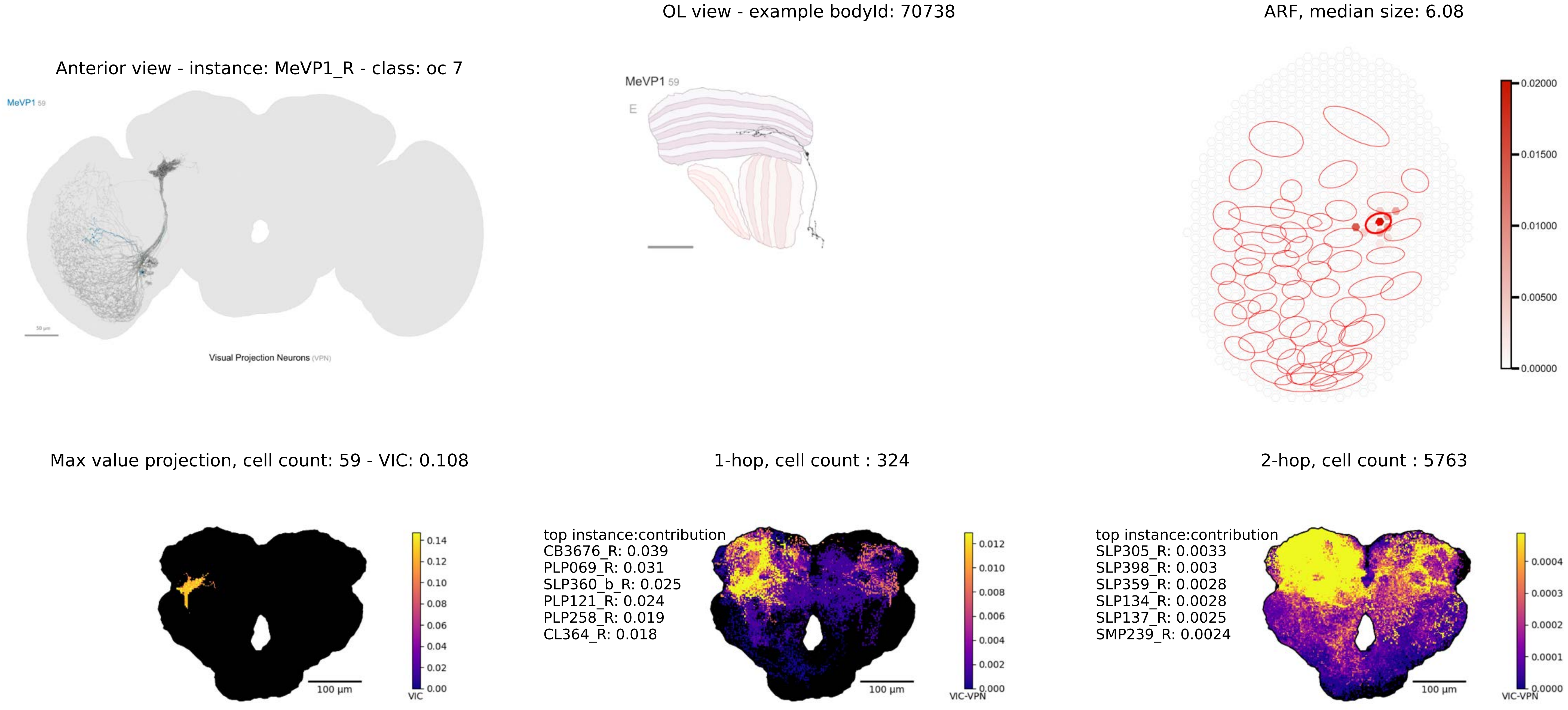
OL view - example bodyId: 70738
ARF, median size: 6.08
Anterior view - instance: MeVP1_R - class: oc 7
MeVP1 59
Visual Projection Neurons (VPN)
MeVP1 59
E
0.02000
0.01500
0.01000
0.00500
0.00000
Max value projection, cell count: 59 - VIC: 0.108
1-hop, cell count : 324
2-hop, cell count : 5763
0.14
0.12
0.10
0.08
0.06
0.04
0.02
0.00
VIC
100 µm
top instance:contribution
CB3676_R: 0.039
PLP069_R: 0.031
SLP360_b_R: 0.025
PLP121_R: 0.024
PLP258_R: 0.019
CL364_R: 0.018
0.012
0.010
0.008
0.006
0.004
0.002
0.000
VIC-VPN
100 µm
top instance:contribution
SLP305_R: 0.0033
SLP398_R: 0.003
SLP359_R: 0.0028
SLP134_R: 0.0028
SLP137_R: 0.0025
SMP239_R: 0.0024
0.0004
0.0003
0.0002
0.0001
0.0000
VIC-VPN
100 µm

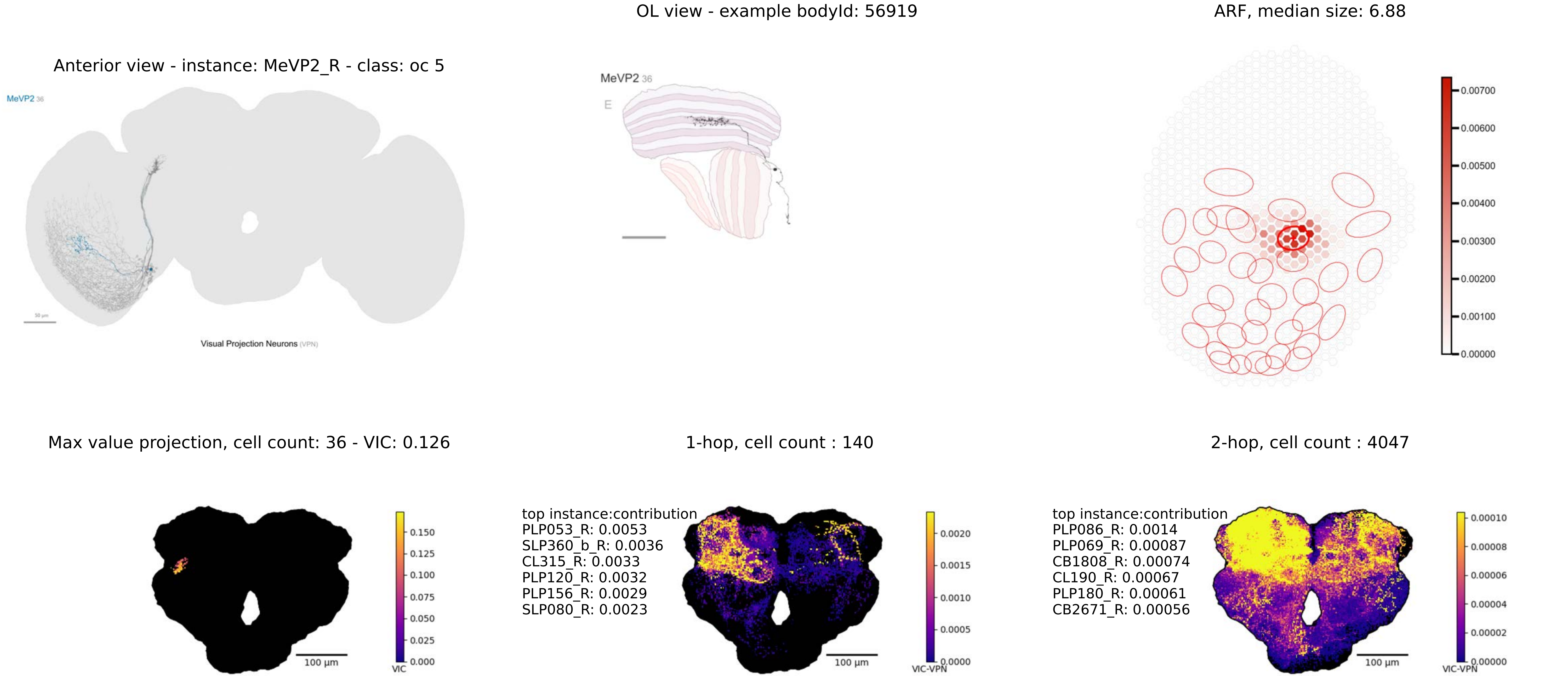

Anterior view - instance: MeVP2_R - class: oc 5
MeVP2 36
50 µm
Visual Projection Neurons (VPN)
OL view - example bodyId: 56919
MeVP2 36
E
ARF, median size: 6.88
0.00700
0.00600
0.00500
0.00400
0.00300
0.00200
0.00100
0.00000
Max value projection, cell count: 36 - VIC: 0.126
100 µm
0.150
0.125
0.100
0.075
0.050
0.025
0.000
VIC
1-hop, cell count : 140
top instance:contribution
PLP053_R: 0.0053
SLP360_b_R: 0.0036
CL315_R: 0.0033
PLP120_R: 0.0032
PLP156_R: 0.0029
SLP080_R: 0.0023
100 µm
0.0020
0.0015
0.0010
0.0005
0.0000
VIC-VPN
2-hop, cell count : 4047
top instance:contribution
PLP086_R: 0.0014
PLP069_R: 0.00087
CB1808_R: 0.00074
CL190_R: 0.00067
PLP180_R: 0.00061
CB2671_R: 0.00056
100 µm
0.00010
0.00008
0.00006
0.00004
0.00002
0.00000
VIC-VPN

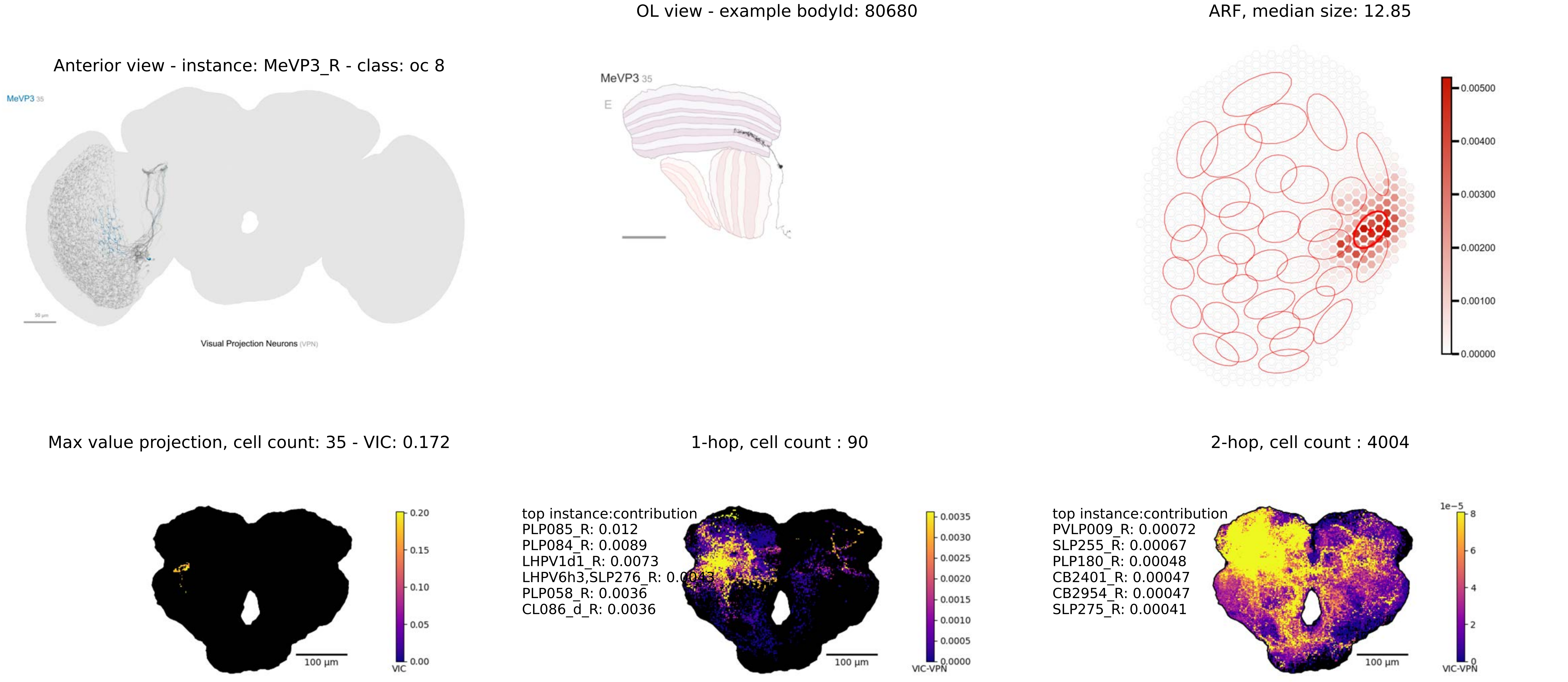
OL view - example bodyId: 80680
ARF, median size: 12.85
Anterior view - instance: MeVP3_R - class: oc 8
MeVP3 35
50 µm
Visual Projection Neurons (VPN)
MeVP3 35
E
0.00500
0.00400
0.00300
0.00200
0.00100
0.00000
Max value projection, cell count: 35 - VIC: 0.172
1-hop, cell count : 90
2-hop, cell count : 4004
0.20
0.15
0.10
0.05
0.00
VIC
100 µm
top instance:contribution
PLP085_R: 0.012
PLP084_R: 0.0089
LHPV1d1_R: 0.0073
LHPV6h3,SLP276_R: 0.0043
PLP058_R: 0.0036
CL086_d_R: 0.0036
100 µm
0.0035
0.0030
0.0025
0.0020
0.0015
0.0010
0.0005
0.0000
VIC-VPN
top instance:contribution
PVLP009_R: 0.00072
SLP255_R: 0.00067
PLP180_R: 0.00048
CB2401_R: 0.00047
CB2954_R: 0.00047
SLP275_R: 0.00041
100 µm
1e−5
8
6
4
2
0
VIC-VPN

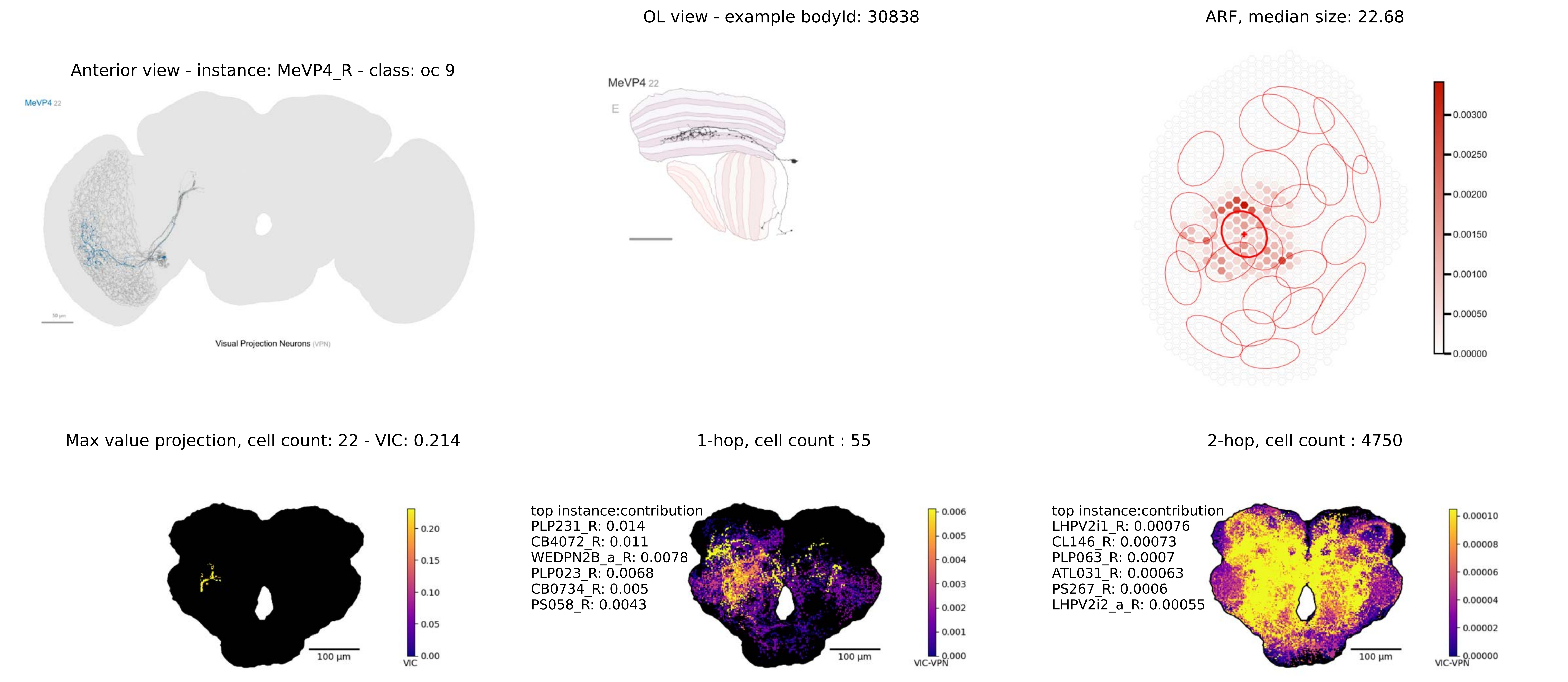

OL view - example bodyId: 30838
ARF, median size: 22.68
Anterior view - instance: MeVP4_R - class: oc 9
MeVP4 22
Visual Projection Neurons (VPN)
MeVP4 22
E
0.00300
0.00250
0.00200
0.00150
0.00100
0.00050
0.00000
Max value projection, cell count: 22 - VIC: 0.214
1-hop, cell count : 55
2-hop, cell count : 4750
100 µm
VIC
0.20
0.15
0.10
0.05
0.00
top instance:contribution
PLP231_R: 0.014
CB4072_R: 0.011
WEDPN2B_a_R: 0.0078
PLP023_R: 0.0068
CB0734_R: 0.005
PS058_R: 0.0043
VIC-VPN
0.006
0.005
0.004
0.003
0.002
0.001
0.000
top instance:contribution
LHPV2i1_R: 0.00076
CL146_R: 0.00073
PLP063_R: 0.0007
ATL031_R: 0.00063
PS267_R: 0.0006
LHPV2i2_a_R: 0.00055
0.00010
0.00008
0.00006
0.00004
0.00002
0.00000

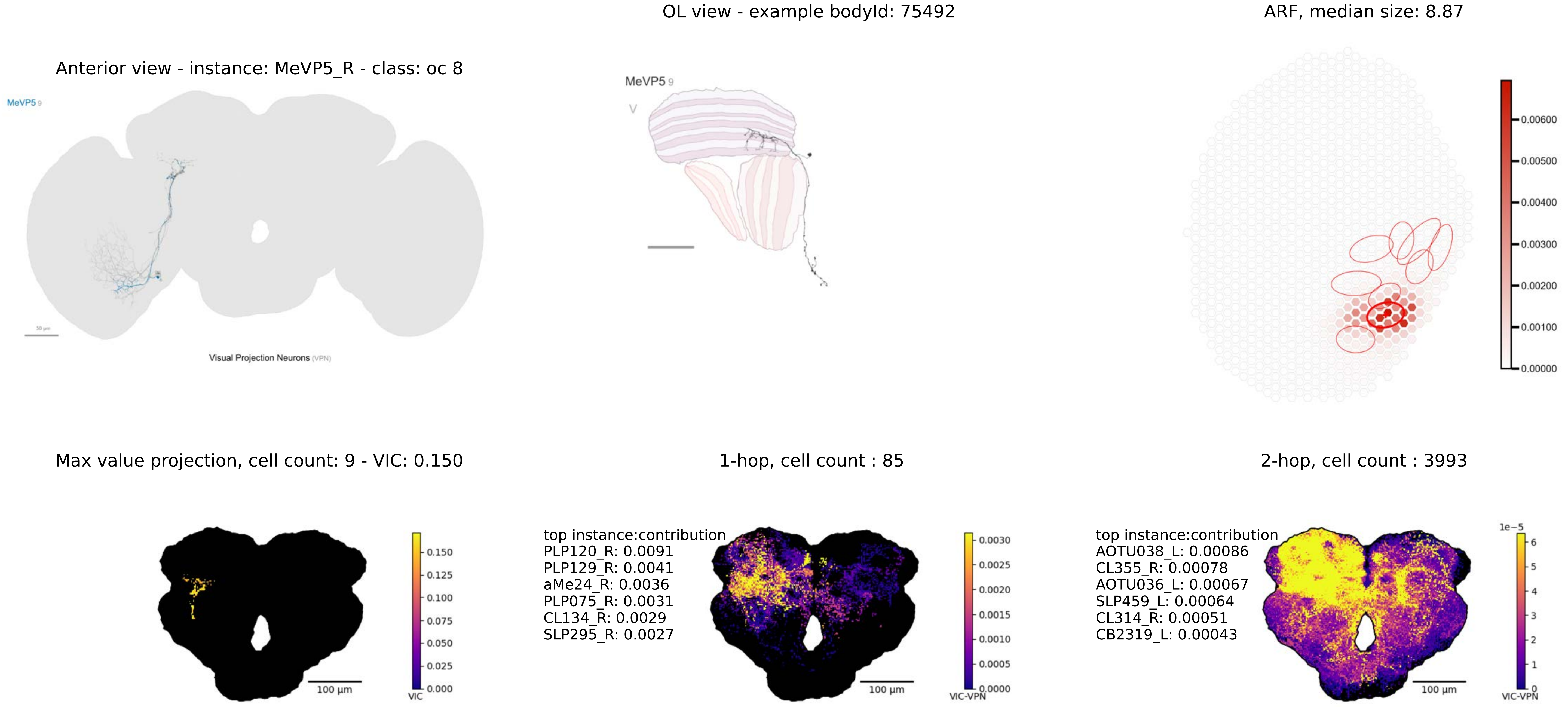

OL view - example bodyId: 75492
ARF, median size: 8.87
Anterior view - instance: MeVP5_R - class: oc 8
MeVP5 9
Visual Projection Neurons (VPN)
0.00600
0.00500
0.00400
0.00300
0.00200
0.00100
0.00000
Max value projection, cell count: 9 - VIC: 0.150
1-hop, cell count : 85
2-hop, cell count : 3993
0.150
0.125
0.100
0.075
0.050
0.025
0.000
VIC
100 µm
top instance:contribution
PLP120_R: 0.0091
PLP129_R: 0.0041
aMe24_R: 0.0036
PLP075_R: 0.0031
CL134_R: 0.0029
SLP295_R: 0.0027
0.0030
0.0025
0.0020
0.0015
0.0010
0.0005
0.0000
VIC-VPN
top instance:contribution
AOTU038_L: 0.00086
CL355_R: 0.00078
AOTU036_L: 0.00067
SLP459_L: 0.00064
CL314_R: 0.00051
CB2319_L: 0.00043
1e−5
6
5
4
3
2
1
0
VIC-VPN

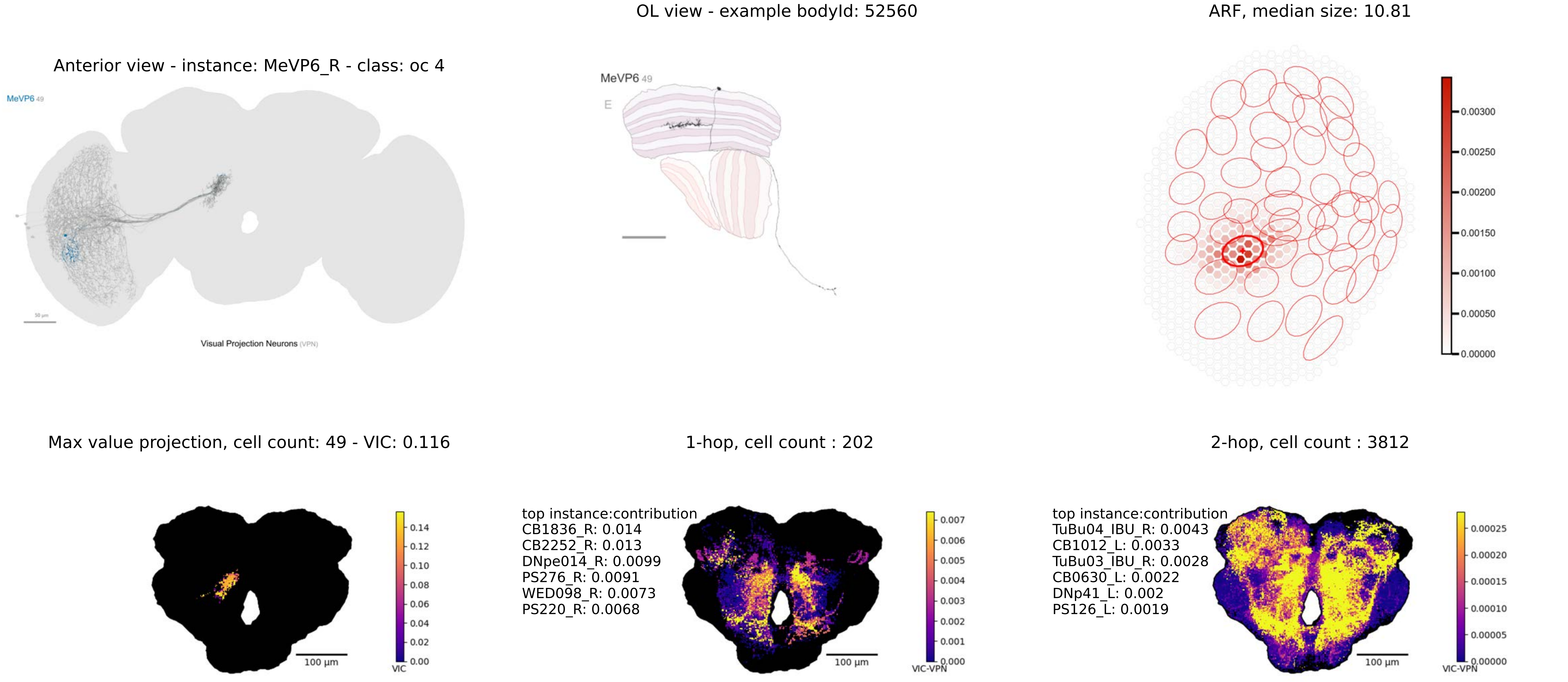

OL view - example bodyId: 52560
ARF, median size: 10.81
Anterior view - instance: MeVP6_R - class: oc 4
MeVP6 49
Visual Projection Neurons (VPN)
MeVP6 49
E
0.00300
0.00250
0.00200
0.00150
0.00100
0.00050
0.00000
Max value projection, cell count: 49 - VIC: 0.116
1-hop, cell count : 202
2-hop, cell count : 3812
0.14
0.12
0.10
0.08
0.06
0.04
0.02
0.00
VIC
100 µm
top instance:contribution
CB1836_R: 0.014
CB2252_R: 0.013
DNpe014_R: 0.0099
PS276_R: 0.0091
WED098_R: 0.0073
PS220_R: 0.0068
100 µm
0.007
0.006
0.005
0.004
0.003
0.002
0.001
0.000
VIC-VPN
top instance:contribution
TuBu04_lBU_R: 0.0043
CB1012_L: 0.0033
TuBu03_lBU_R: 0.0028
CB0630_L: 0.0022
DNp41_L: 0.002
PS126_L: 0.0019
100 µm
0.00025
0.00020
0.00015
0.00010
0.00005
0.00000
VIC-VPN

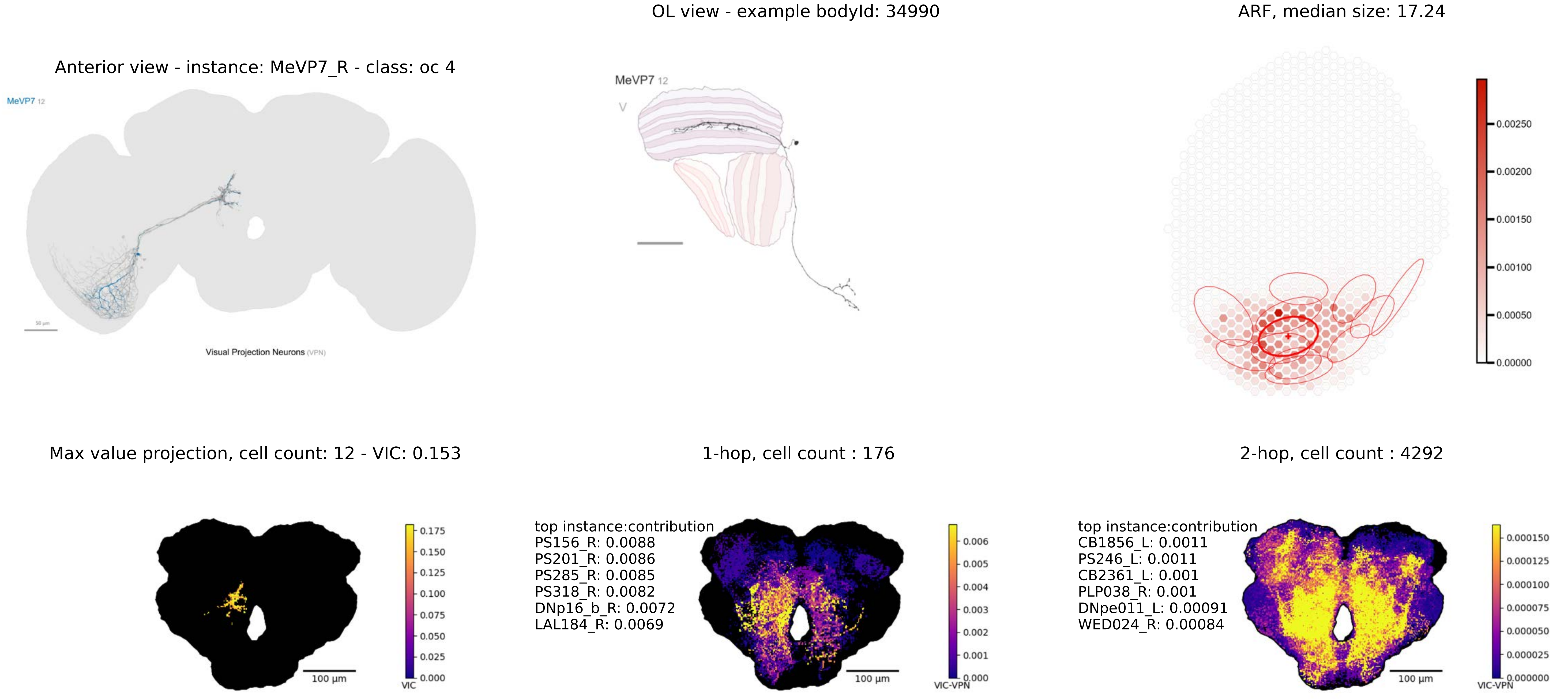
OL view - example bodyId: 34990
ARF, median size: 17.24
Anterior view - instance: MeVP7_R - class: oc 4
MeVP7 12
Visual Projection Neurons (VPN)
MeVP7 12
0.00250
0.00200
0.00150
0.00100
0.00050
0.00000
Max value projection, cell count: 12 - VIC: 0.153
1-hop, cell count : 176
2-hop, cell count : 4292
0.175
0.150
0.125
0.100
0.075
0.050
0.025
0.000
VIC
100 µm
top instance:contribution
PS156_R: 0.0088
PS201_R: 0.0086
PS285_R: 0.0085
PS318_R: 0.0082
DNp16_b_R: 0.0072
LAL184_R: 0.0069
0.006
0.005
0.004
0.003
0.002
0.001
0.000
VIC-VPN
100 µm
top instance:contribution
CB1856_L: 0.0011
PS246_L: 0.0011
CB2361_L: 0.001
PLP038_R: 0.001
DNpe011_L: 0.00091
WED024_R: 0.00084
0.000150
0.000125
0.000100
0.000075
0.000050
0.000025
0.000000
VIC-VPN
100 µm

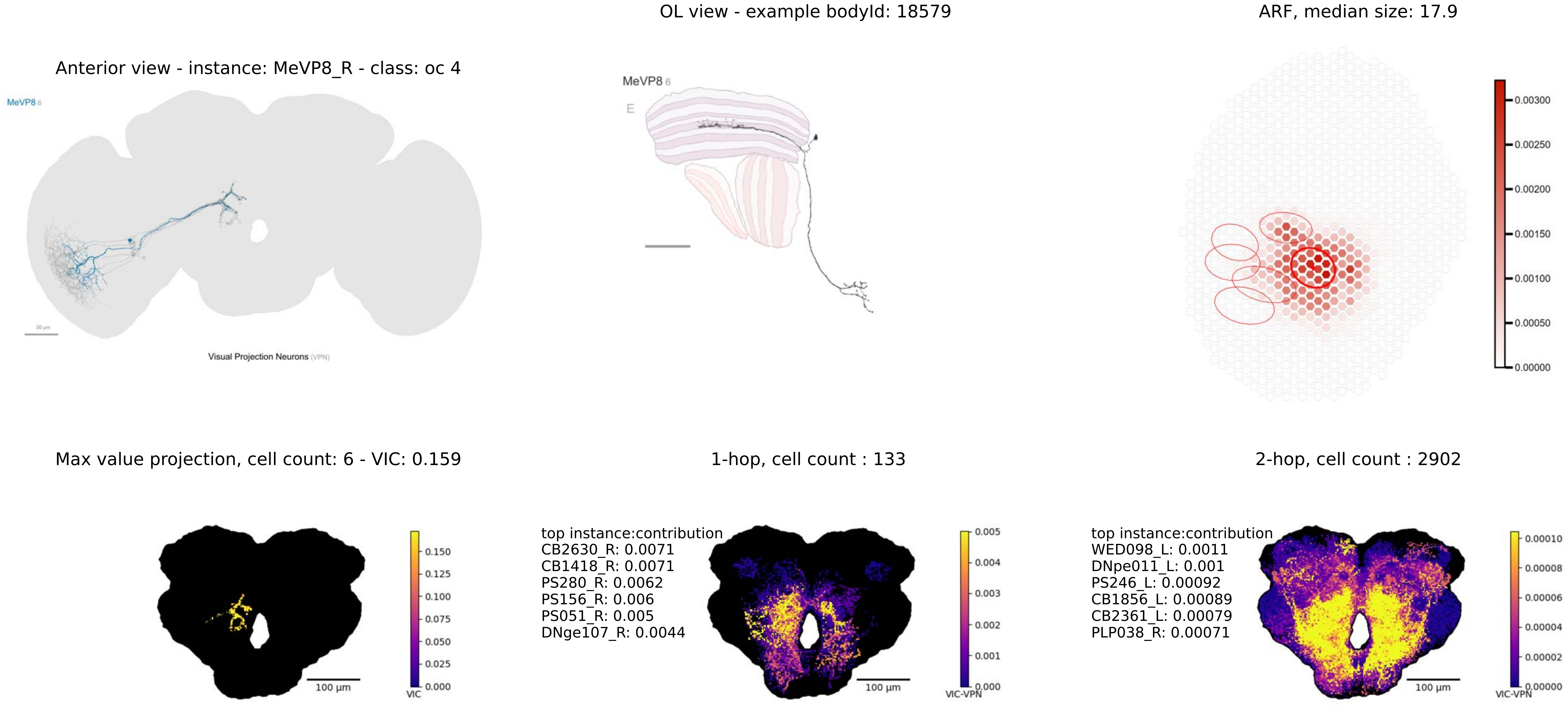
OL view - example bodyId: 18579
ARF, median size: 17.9
Anterior view - instance: MeVP8_R - class: oc 4
MeVP8 6
Visual Projection Neurons (VPN)
MeVP8 6
E
0.00300
0.00250
0.00200
0.00150
0.00100
0.00050
0.00000
Max value projection, cell count: 6 - VIC: 0.159
1-hop, cell count : 133
2-hop, cell count : 2902
100 µm
VIC
0.150
0.125
0.100
0.075
0.050
0.025
0.000
top instance:contribution
CB2630_R: 0.0071
CB1418_R: 0.0071
PS280_R: 0.0062
PS156_R: 0.006
PS051_R: 0.005
DNge107_R: 0.0044
100 µm
VIC-VPN
0.005
0.004
0.003
0.002
0.001
0.000
top instance:contribution
WED098_L: 0.0011
DNpe011_L: 0.001
PS246_L: 0.00092
CB1856_L: 0.00089
CB2361_L: 0.00079
PLP038_R: 0.00071
100 µm
VIC-VPN
0.00010
0.00008
0.00006
0.00004
0.00002
0.00000

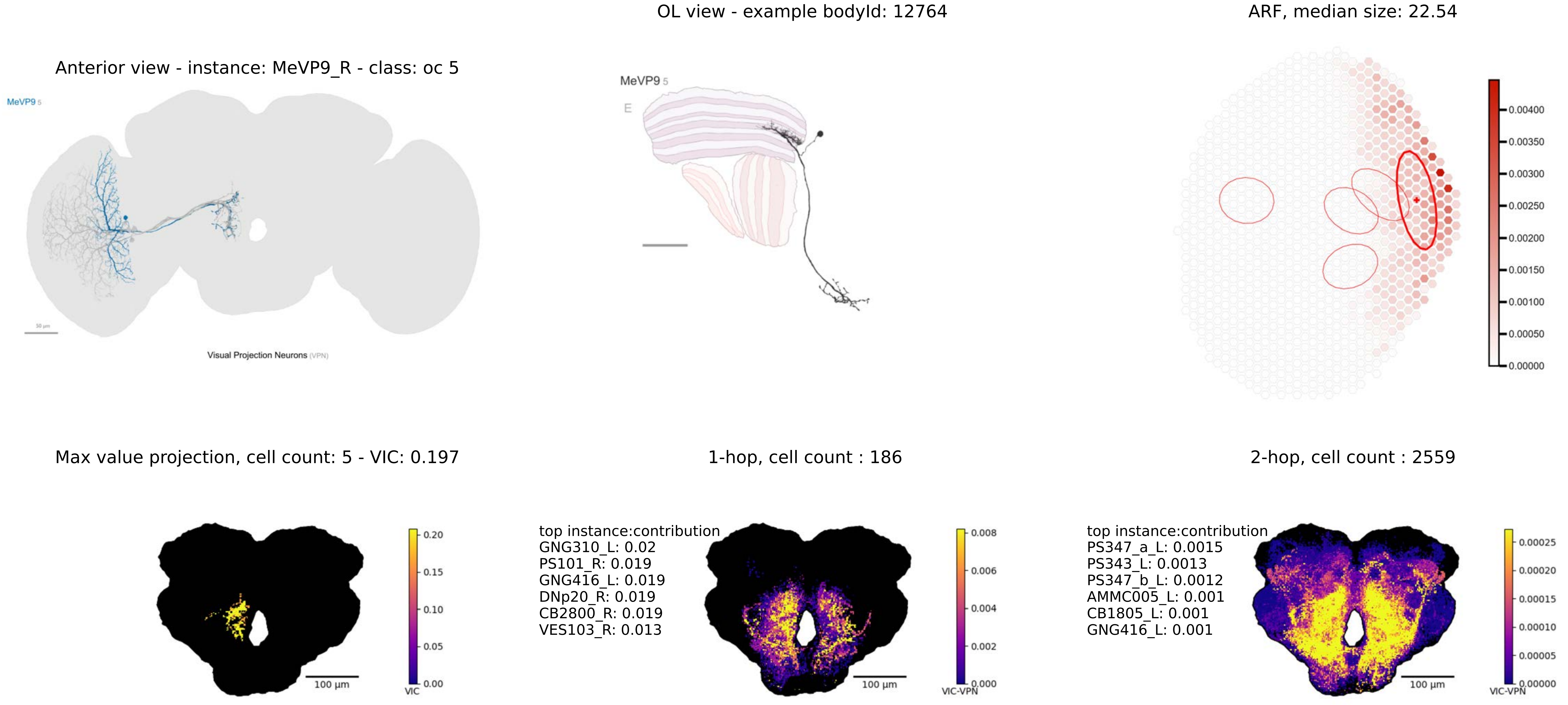
OL view - example bodyId: 12764
ARF, median size: 22.54
Anterior view - instance: MeVP9_R - class: oc 5
MeVP9 5
Visual Projection Neurons (VPN)
50 µm
MeVP9 5
E
0.00400
0.00350
0.00300
0.00250
0.00200
0.00150
0.00100
0.00050
0.00000
Max value projection, cell count: 5 - VIC: 0.197
1-hop, cell count : 186
2-hop, cell count : 2559
100 µm
0.20
0.15
0.10
0.05
0.00
VIC
top instance:contribution
GNG310_L: 0.02
PS101_R: 0.019
GNG416_L: 0.019
DNp20_R: 0.019
CB2800_R: 0.019
VES103_R: 0.013
100 µm
0.008
0.006
0.004
0.002
0.000
VIC-VPN
top instance:contribution
PS347_a_L: 0.0015
PS343_L: 0.0013
PS347_b_L: 0.0012
AMMC005_L: 0.001
CB1805_L: 0.001
GNG416_L: 0.001
100 µm
0.00025
0.00020
0.00015
0.00010
0.00005
0.00000
VIC-VPN

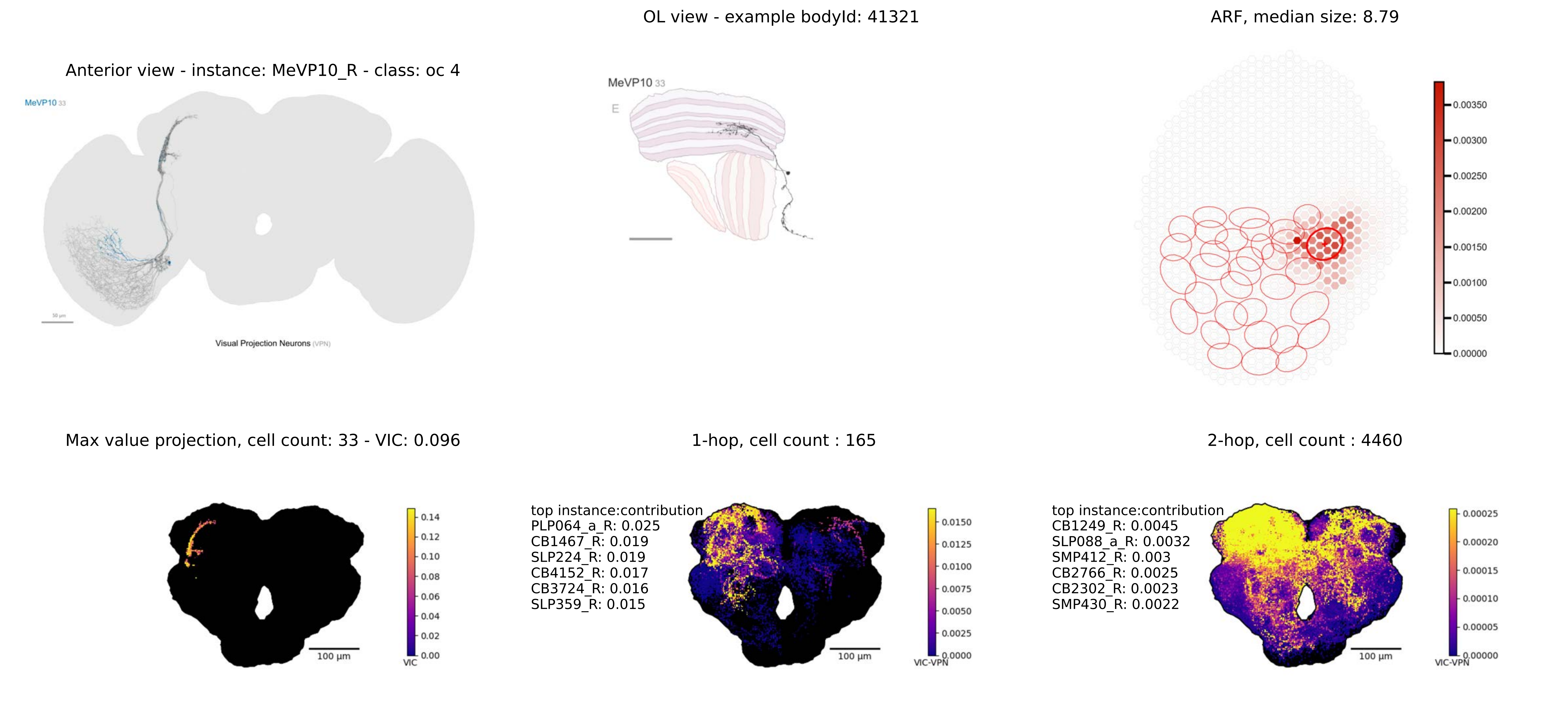
Anterior view - instance: MeVP10_R - class: oc 4
MeVP10 33
Visual Projection Neurons (VPN)
OL view - example bodyId: 41321
MeVP10 33
E
ARF, median size: 8.79
0.00350
0.00300
0.00250
0.00200
0.00150
0.00100
0.00050
0.00000
Max value projection, cell count: 33 - VIC: 0.096
0.14
0.12
0.10
0.08
0.06
0.04
0.02
0.00
VIC
100 µm
1-hop, cell count : 165
top instance:contribution
PLP064_a_R: 0.025
CB1467_R: 0.019
SLP224_R: 0.019
CB4152_R: 0.017
CB3724_R: 0.016
SLP359_R: 0.015
0.0150
0.0125
0.0100
0.0075
0.0050
0.0025
0.0000
VIC-VPN
100 µm
2-hop, cell count : 4460
top instance:contribution
CB1249_R: 0.0045
SLP088_a_R: 0.0032
SMP412_R: 0.003
CB2766_R: 0.0025
CB2302_R: 0.0023
SMP430_R: 0.0022
0.00025
0.00020
0.00015
0.00010
0.00005
0.00000
VIC-VPN
100 µm

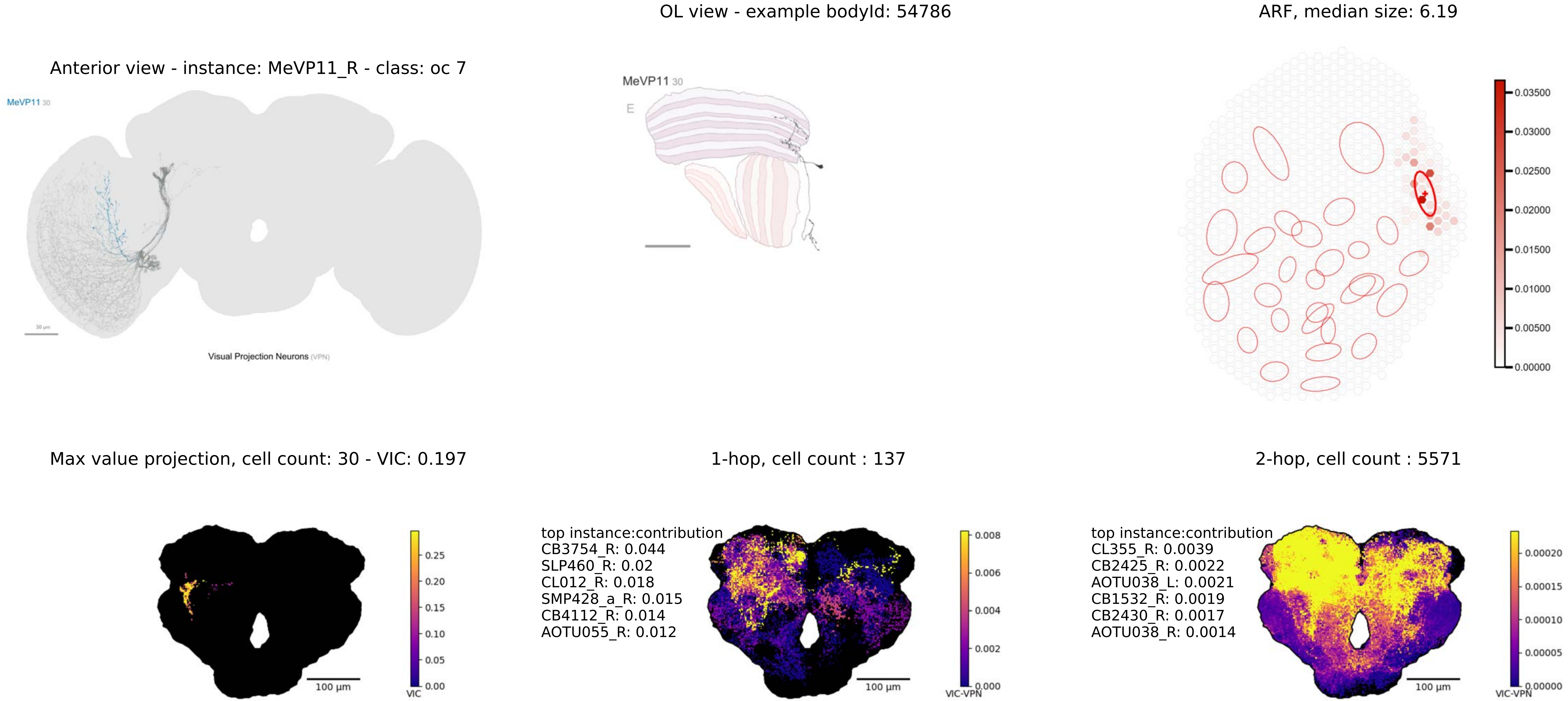
OL view - example bodyId: 54786
ARF, median size: 6.19
Anterior view - instance: MeVP11_R - class: oc 7
MeVP11 30
Visual Projection Neurons (VPN)
MeVP11 30
E
0.03500
0.03000
0.02500
0.02000
0.01500
0.01000
0.00500
0.00000
Max value projection, cell count: 30 - VIC: 0.197
1-hop, cell count : 137
2-hop, cell count : 5571
100 µm
0.25
0.20
0.15
0.10
0.05
0.00
VIC
top instance:contribution
CB3754_R: 0.044
SLP460_R: 0.02
CL012_R: 0.018
SMP428_a_R: 0.015
CB4112_R: 0.014
AOTU055_R: 0.012
100 µm
0.008
0.006
0.004
0.002
0.000
VIC-VPN
top instance:contribution
CL355_R: 0.0039
CB2425_R: 0.0022
AOTU038_L: 0.0021
CB1532_R: 0.0019
CB2430_R: 0.0017
AOTU038_R: 0.0014
100 µm
0.00020
0.00015
0.00010
0.00005
0.00000
VIC-VPN

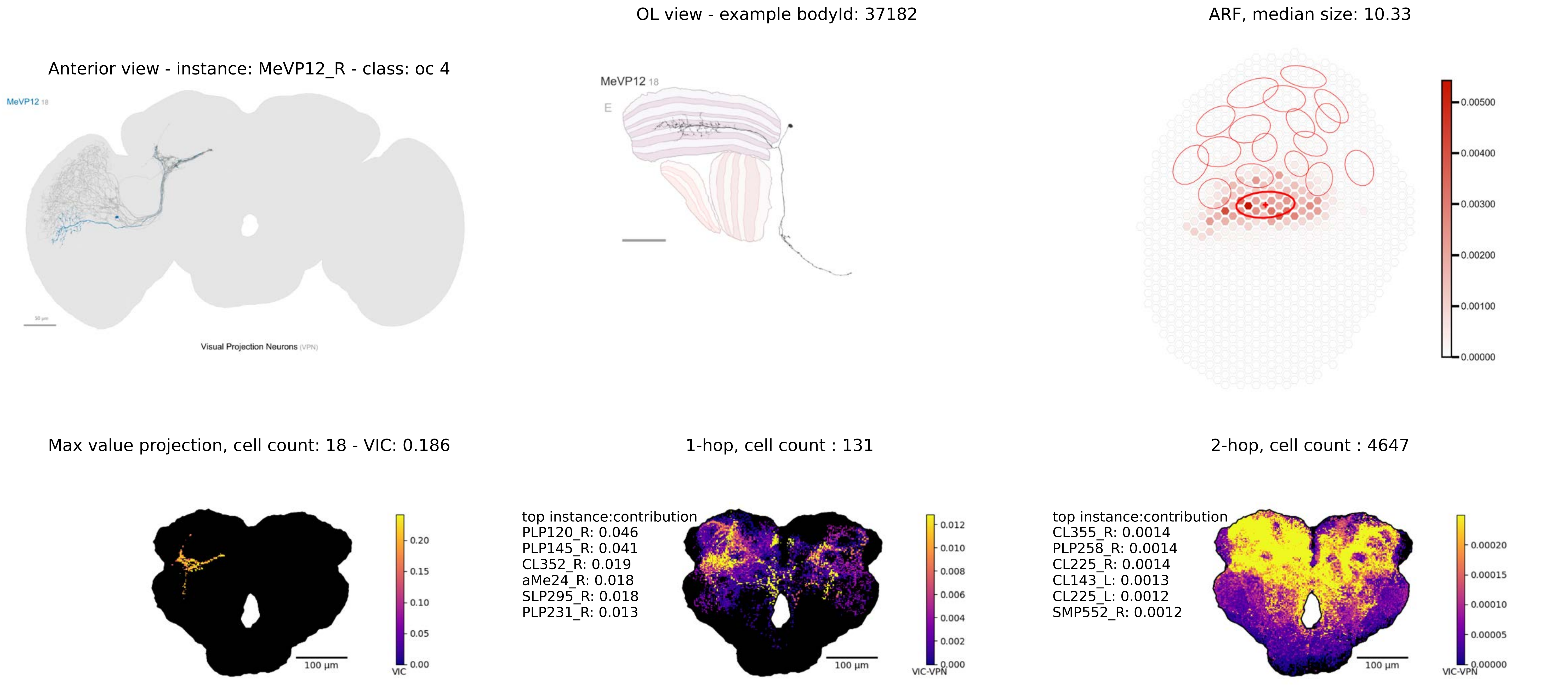
OL view - example bodyId: 37182
ARF, median size: 10.33
Anterior view - instance: MeVP12_R - class: oc 4
MeVP12 18
Visual Projection Neurons (VPN)
MeVP12 18
E
0.00500
0.00400
0.00300
0.00200
0.00100
0.00000
Max value projection, cell count: 18 - VIC: 0.186
1-hop, cell count : 131
2-hop, cell count : 4647
top instance:contribution
PLP120_R: 0.046
PLP145_R: 0.041
CL352_R: 0.019
aMe24_R: 0.018
SLP295_R: 0.018
PLP231_R: 0.013
top instance:contribution
CL355_R: 0.0014
PLP258_R: 0.0014
CL225_R: 0.0014
CL143_L: 0.0013
CL225_L: 0.0012
SMP552_R: 0.0012
100 µm
0.20
0.15
0.10
0.05
0.00
VIC
100 µm
0.012
0.010
0.008
0.006
0.004
0.002
0.000
VIC-VPN
100 µm
0.00020
0.00015
0.00010
0.00005
0.00000
VIC-VPN

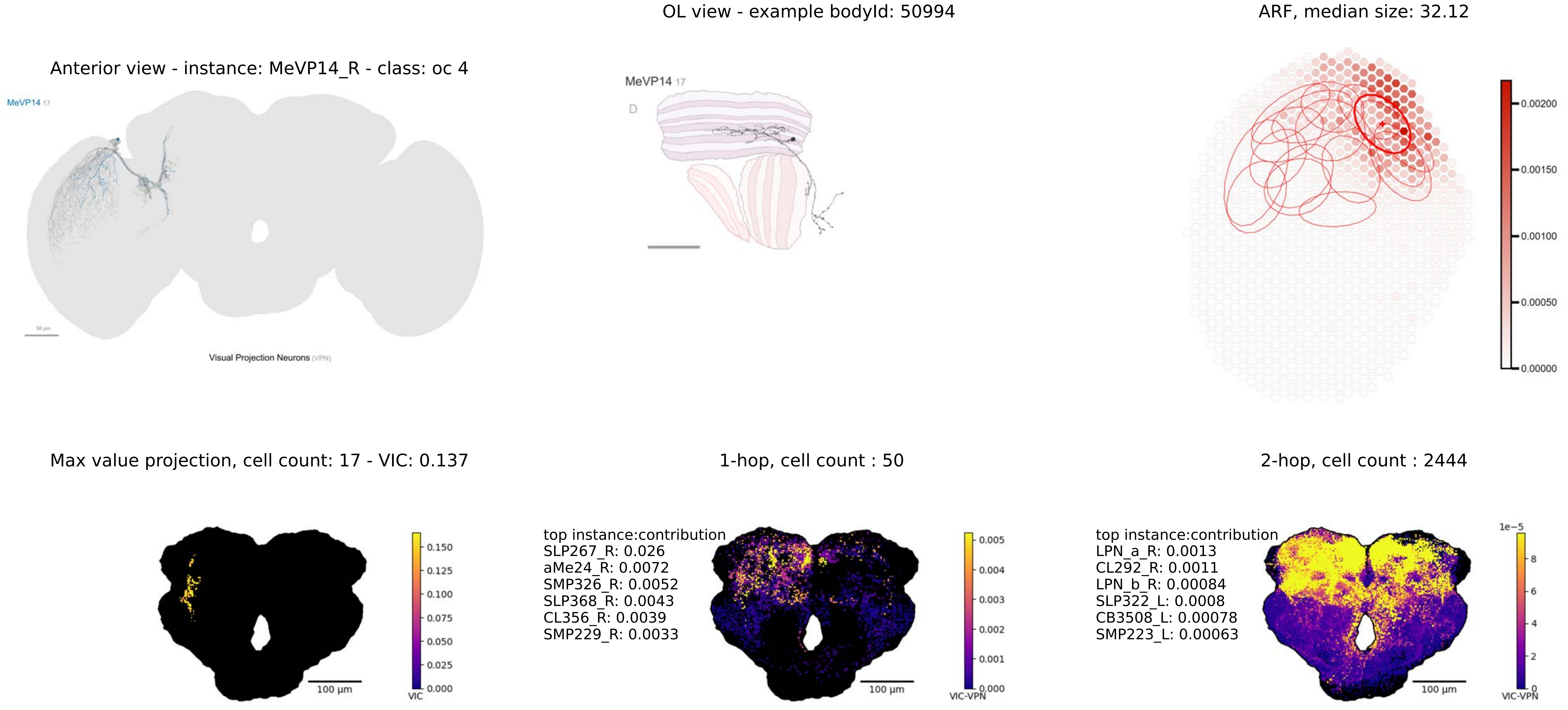
OL view - example bodyId: 50994
ARF, median size: 32.12
Anterior view - instance: MeVP14_R - class: oc 4
MeVP14 17
Visual Projection Neurons (VPN)
MeVP14 17
D
0.00200
0.00150
0.00100
0.00050
0.00000
Max value projection, cell count: 17 - VIC: 0.137
1-hop, cell count : 50
2-hop, cell count : 2444
0.150
0.125
0.100
0.075
0.050
0.025
0.000
VIC
100 µm
top instance:contribution
SLP267_R: 0.026
aMe24_R: 0.0072
SMP326_R: 0.0052
SLP368_R: 0.0043
CL356_R: 0.0039
SMP229_R: 0.0033
0.005
0.004
0.003
0.002
0.001
0.000
VIC-VPN
100 µm
top instance:contribution
LPN_a_R: 0.0013
CL292_R: 0.0011
LPN_b_R: 0.00084
SLP322_L: 0.0008
CB3508_L: 0.00078
SMP223_L: 0.00063
1e−5
8
6
4
2
0
VIC-VPN
100 µm

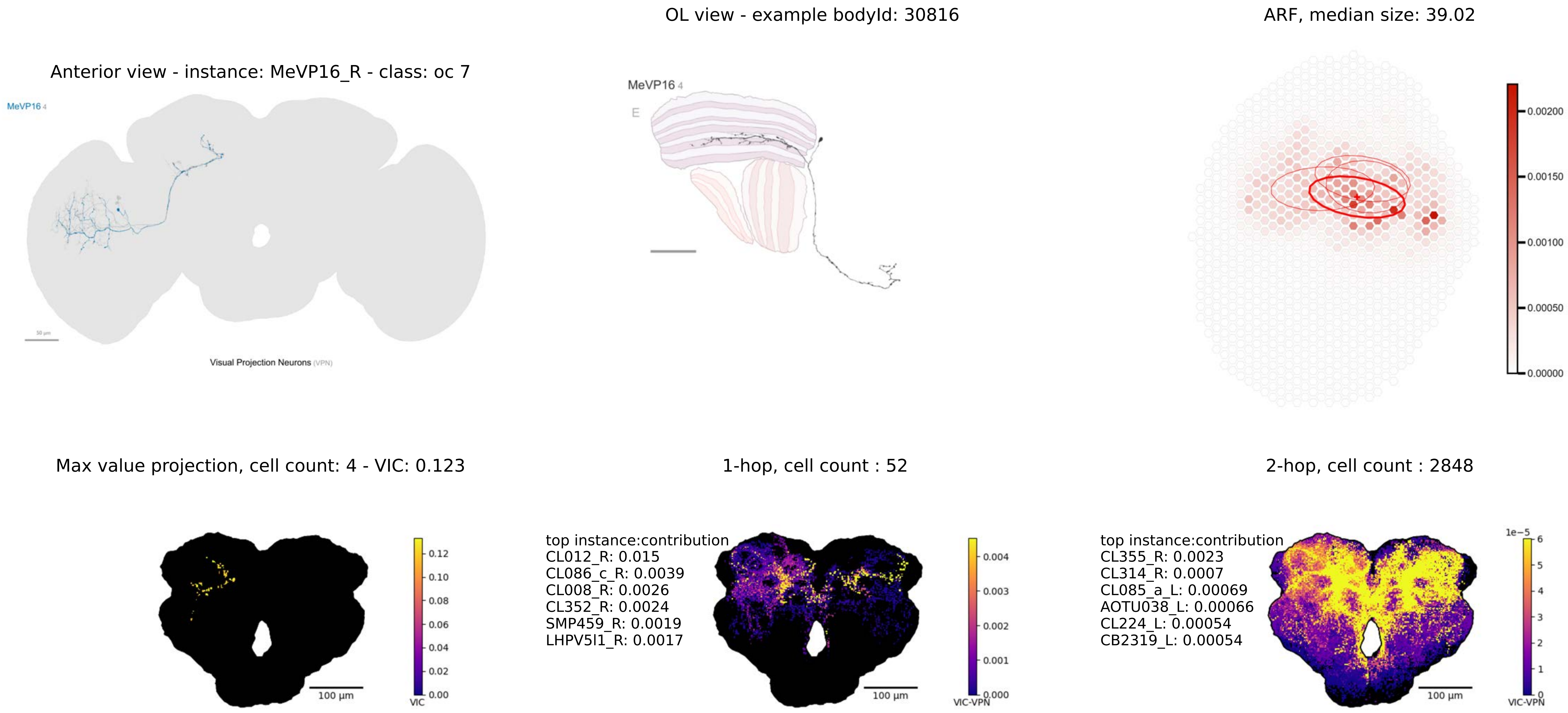
OL view - example bodyId: 30816
ARF, median size: 39.02
Anterior view - instance: MeVP16_R - class: oc 7
MeVP16 4
Visual Projection Neurons (VPN)
MeVP16 4
E
0.00200
0.00150
0.00100
0.00050
0.00000
Max value projection, cell count: 4 - VIC: 0.123
1-hop, cell count : 52
2-hop, cell count : 2848
0.12
0.10
0.08
0.06
0.04
0.02
0.00
VIC
100 µm
top instance:contribution
CL012_R: 0.015
CL086_c_R: 0.0039
CL008_R: 0.0026
CL352_R: 0.0024
SMP459_R: 0.0019
LHPV5l1_R: 0.0017
0.004
0.003
0.002
0.001
0.000
VIC-VPN
100 µm
top instance:contribution
CL355_R: 0.0023
CL314_R: 0.0007
CL085_a_L: 0.00069
AOTU038_L: 0.00066
CL224_L: 0.00054
CB2319_L: 0.00054
1e−5
6
5
4
3
2
1
0
VIC-VPN
100 µm

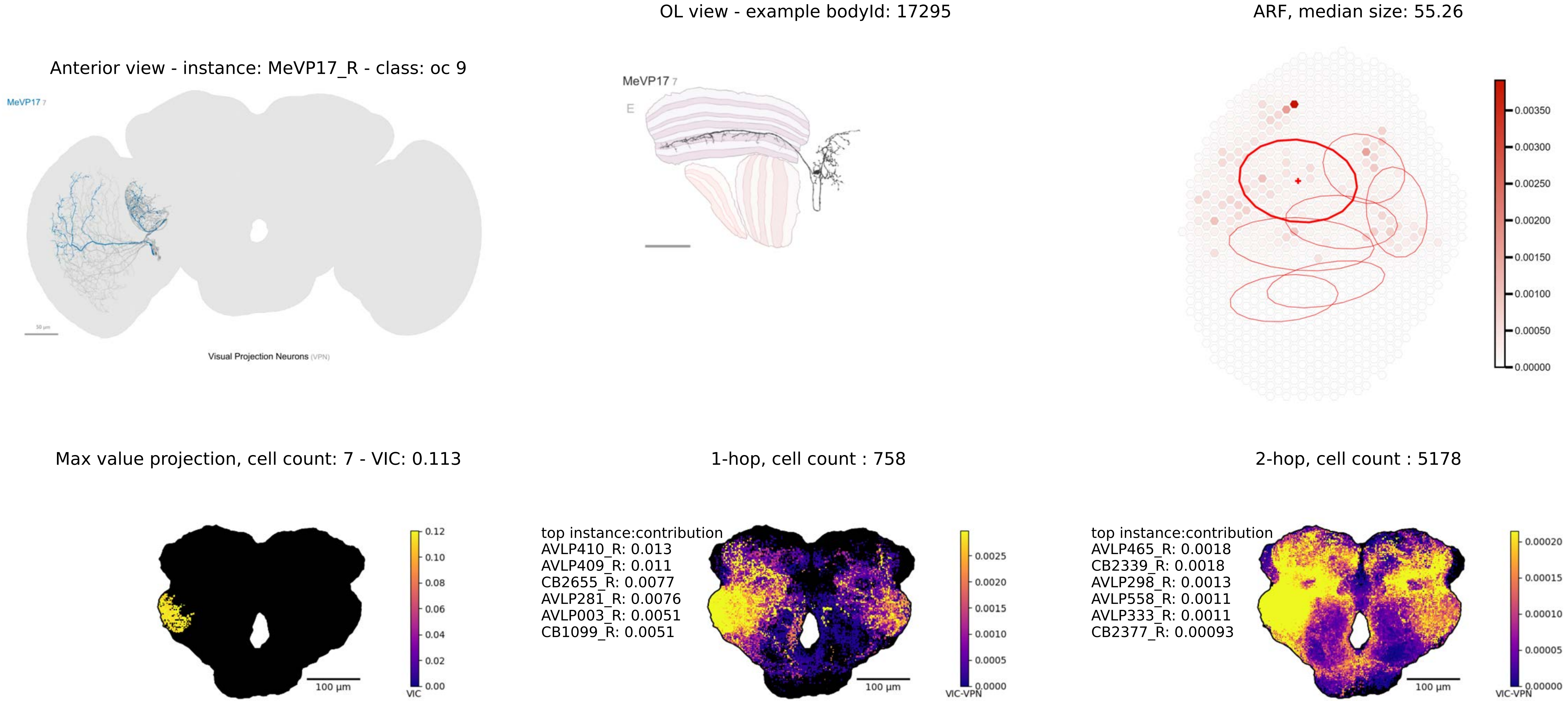
OL view - example bodyId: 17295
ARF, median size: 55.26
Anterior view - instance: MeVP17_R - class: oc 9
MeVP17 7
Visual Projection Neurons (VPN)
MeVP17 7
E
0.00350
0.00300
0.00250
0.00200
0.00150
0.00100
0.00050
0.00000
Max value projection, cell count: 7 - VIC: 0.113
1-hop, cell count : 758
2-hop, cell count : 5178
0.12
0.10
0.08
0.06
0.04
0.02
0.00
VIC
100 µm
top instance:contribution
AVLP410_R: 0.013
AVLP409_R: 0.011
CB2655_R: 0.0077
AVLP281_R: 0.0076
AVLP003_R: 0.0051
CB1099_R: 0.0051
0.0025
0.0020
0.0015
0.0010
0.0005
0.0000
VIC-VPN
100 µm
top instance:contribution
AVLP465_R: 0.0018
CB2339_R: 0.0018
AVLP298_R: 0.0013
AVLP558_R: 0.0011
AVLP333_R: 0.0011
CB2377_R: 0.00093
0.00020
0.00015
0.00010
0.00005
0.00000
VIC-VPN
100 µm

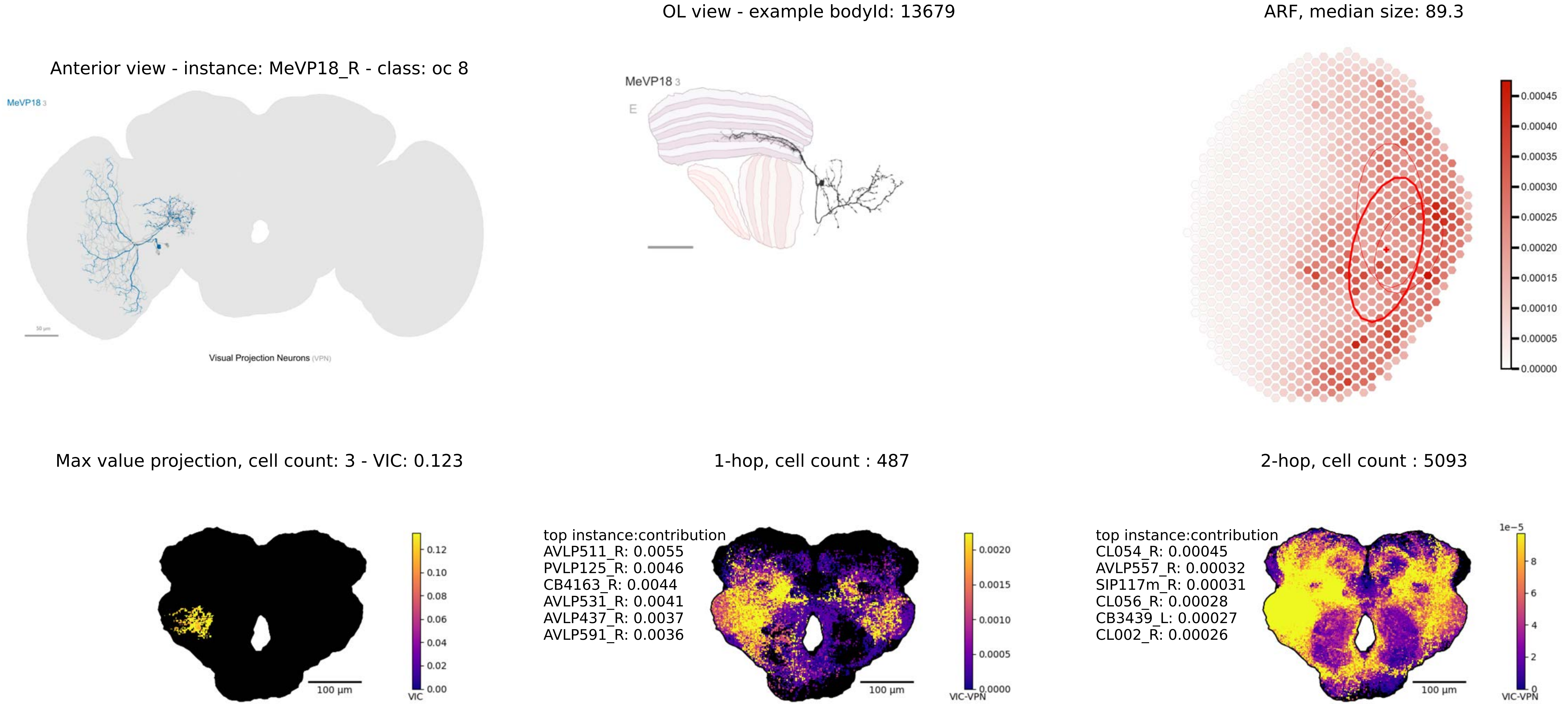

OL view - example bodyId: 13679
ARF, median size: 89.3
Anterior view - instance: MeVP18_R - class: oc 8
MeVP18 3
Visual Projection Neurons (VPN)
MeVP18 3
E
0.00045
0.00040
0.00035
0.00030
0.00025
0.00020
0.00015
0.00010
0.00005
0.00000
Max value projection, cell count: 3 - VIC: 0.123
1-hop, cell count : 487
2-hop, cell count : 5093
0.12
0.10
0.08
0.06
0.04
0.02
0.00
VIC
100 µm
top instance:contribution
AVLP511_R: 0.0055
PVLP125_R: 0.0046
CB4163_R: 0.0044
AVLP531_R: 0.0041
AVLP437_R: 0.0037
AVLP591_R: 0.0036
0.0020
0.0015
0.0010
0.0005
0.0000
VIC-VPN
100 µm
top instance:contribution
CL054_R: 0.00045
AVLP557_R: 0.00032
SIP117m_R: 0.00031
CL056_R: 0.00028
CB3439_L: 0.00027
CL002_R: 0.00026
1e−5
8
6
4
2
0
VIC-VPN
100 µm

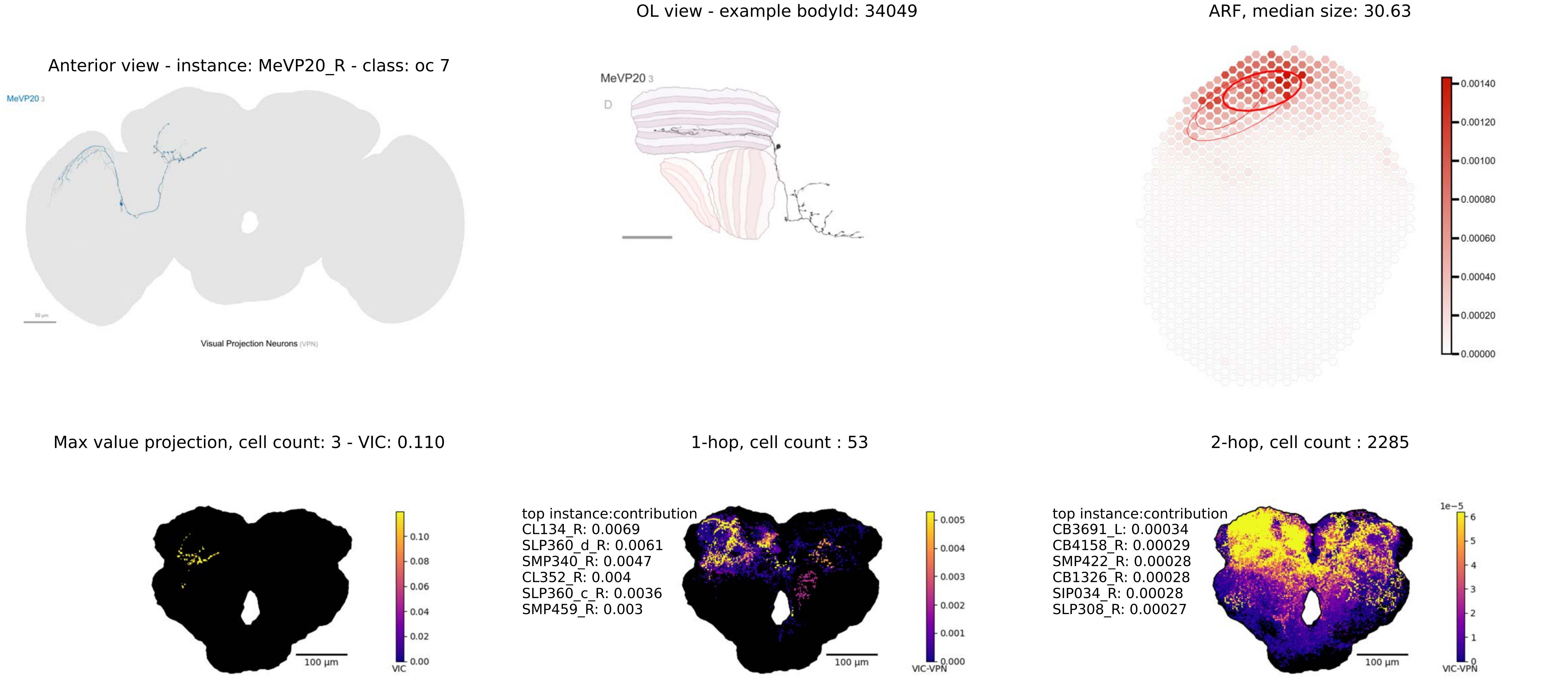

Anterior view - instance: MeVP20_R - class: oc 7
MeVP20 3
Visual Projection Neurons (VPN)
OL view - example bodyId: 34049
MeVP20 3
D
ARF, median size: 30.63
0.00140
0.00120
0.00100
0.00080
0.00060
0.00040
0.00020
0.00000
Max value projection, cell count: 3 - VIC: 0.110
0.10
0.08
0.06
0.04
0.02
0.00
VIC
100 µm
1-hop, cell count : 53
top instance:contribution
CL134_R: 0.0069
SLP360_d_R: 0.0061
SMP340_R: 0.0047
CL352_R: 0.004
SLP360_c_R: 0.0036
SMP459_R: 0.003
0.005
0.004
0.003
0.002
0.001
0.000
VIC-VPN
100 µm
2-hop, cell count : 2285
top instance:contribution
CB3691_L: 0.00034
CB4158_R: 0.00029
SMP422_R: 0.00028
CB1326_R: 0.00028
SIP034_R: 0.00028
SLP308_R: 0.00027
1e−5
6
5
4
3
2
1
0
VIC-VPN
100 µm

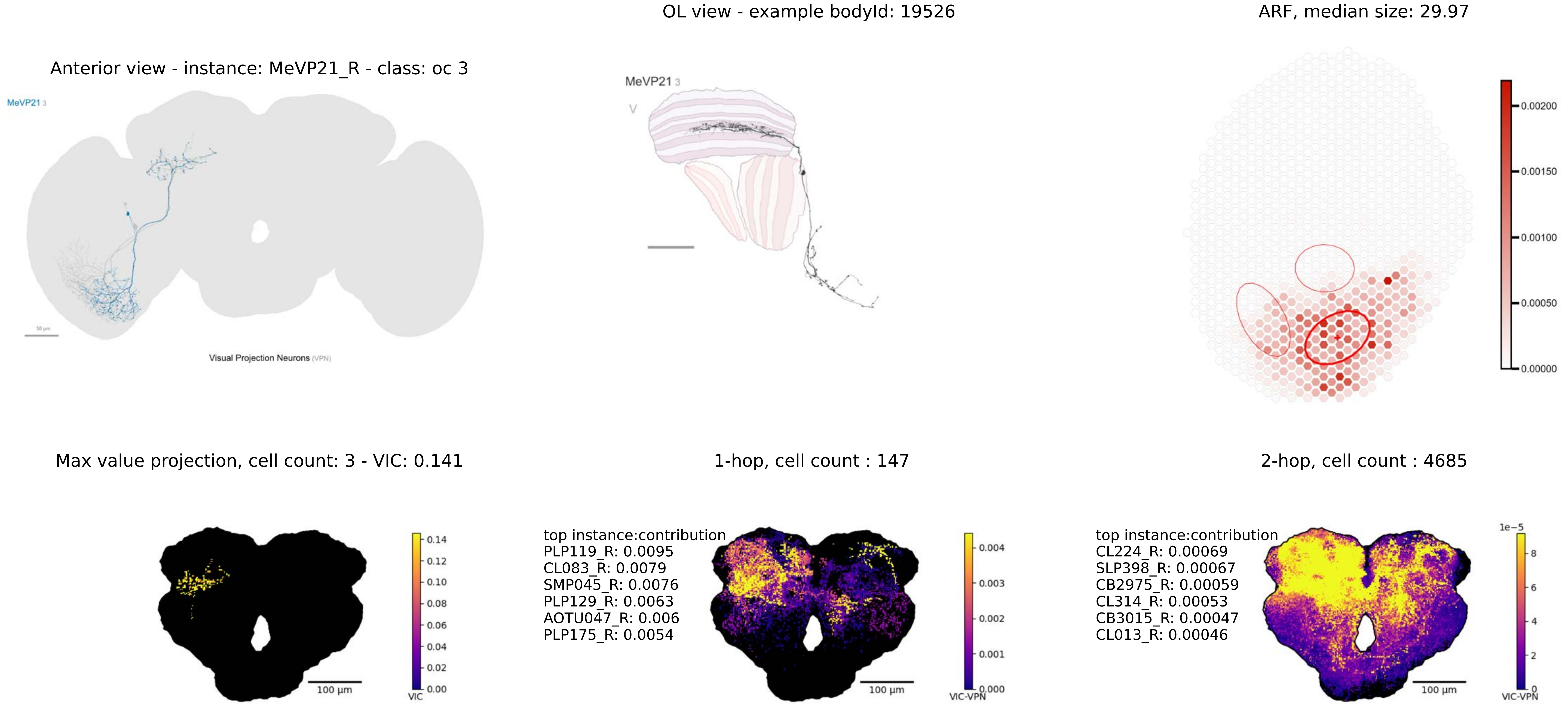

OL view - example bodyId: 19526
ARF, median size: 29.97
Anterior view - instance: MeVP21_R - class: oc 3
MeVP21 3
Visual Projection Neurons (VPN)
MeVP21 3
V
0.00200
0.00150
0.00100
0.00050
0.00000
Max value projection, cell count: 3 - VIC: 0.141
1-hop, cell count : 147
2-hop, cell count : 4685
0.14
0.12
0.10
0.08
0.06
0.04
0.02
0.00
VIC
100 μm
top instance:contribution
PLP119_R: 0.0095
CL083_R: 0.0079
SMP045_R: 0.0076
PLP129_R: 0.0063
AOTU047_R: 0.006
PLP175_R: 0.0054
100 μm
0.004
0.003
0.002
0.001
0.000
VIC-VPN
top instance:contribution
CL224_R: 0.00069
SLP398_R: 0.00067
CB2975_R: 0.00059
CL314_R: 0.00053
CB3015_R: 0.00047
CL013_R: 0.00046
100 μm
1e−5
8
6
4
2
0
VIC-VPN

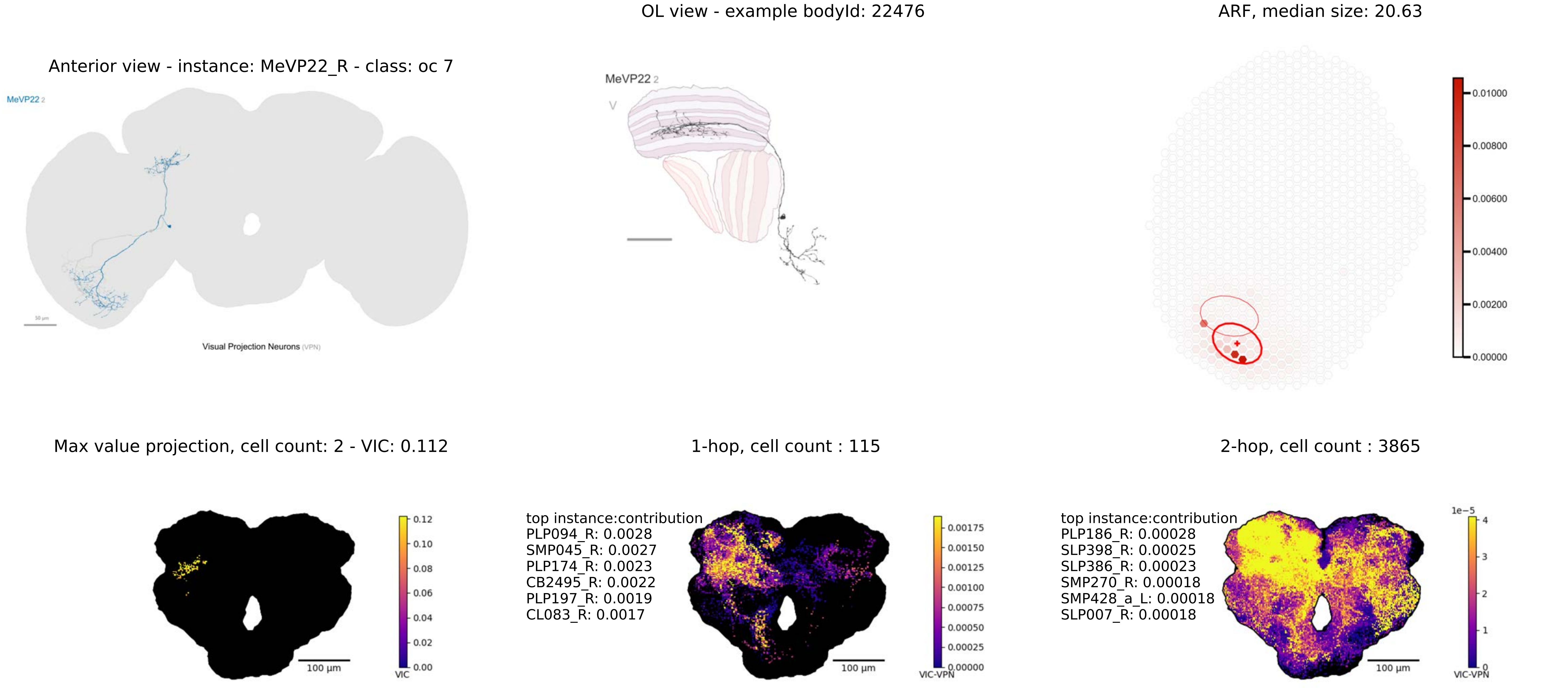
OL view - example bodyId: 22476
ARF, median size: 20.63
Anterior view - instance: MeVP22_R - class: oc 7
MeVP22 2
Visual Projection Neurons (VPN)
MeVP22 2
0.01000
0.00800
0.00600
0.00400
0.00200
0.00000
Max value projection, cell count: 2 - VIC: 0.112
1-hop, cell count : 115
2-hop, cell count : 3865
0.12
0.10
0.08
0.06
0.04
0.02
0.00
VIC
100 µm
top instance:contribution
PLP094_R: 0.0028
SMP045_R: 0.0027
PLP174_R: 0.0023
CB2495_R: 0.0022
PLP197_R: 0.0019
CL083_R: 0.0017
0.00175
0.00150
0.00125
0.00100
0.00075
0.00050
0.00025
0.00000
VIC-VPN
100 µm
top instance:contribution
PLP186_R: 0.00028
SLP398_R: 0.00025
SLP386_R: 0.00023
SMP270_R: 0.00018
SMP428_a_L: 0.00018
SLP007_R: 0.00018
1e−5
4
3
2
1
0
VIC-VPN
100 µm

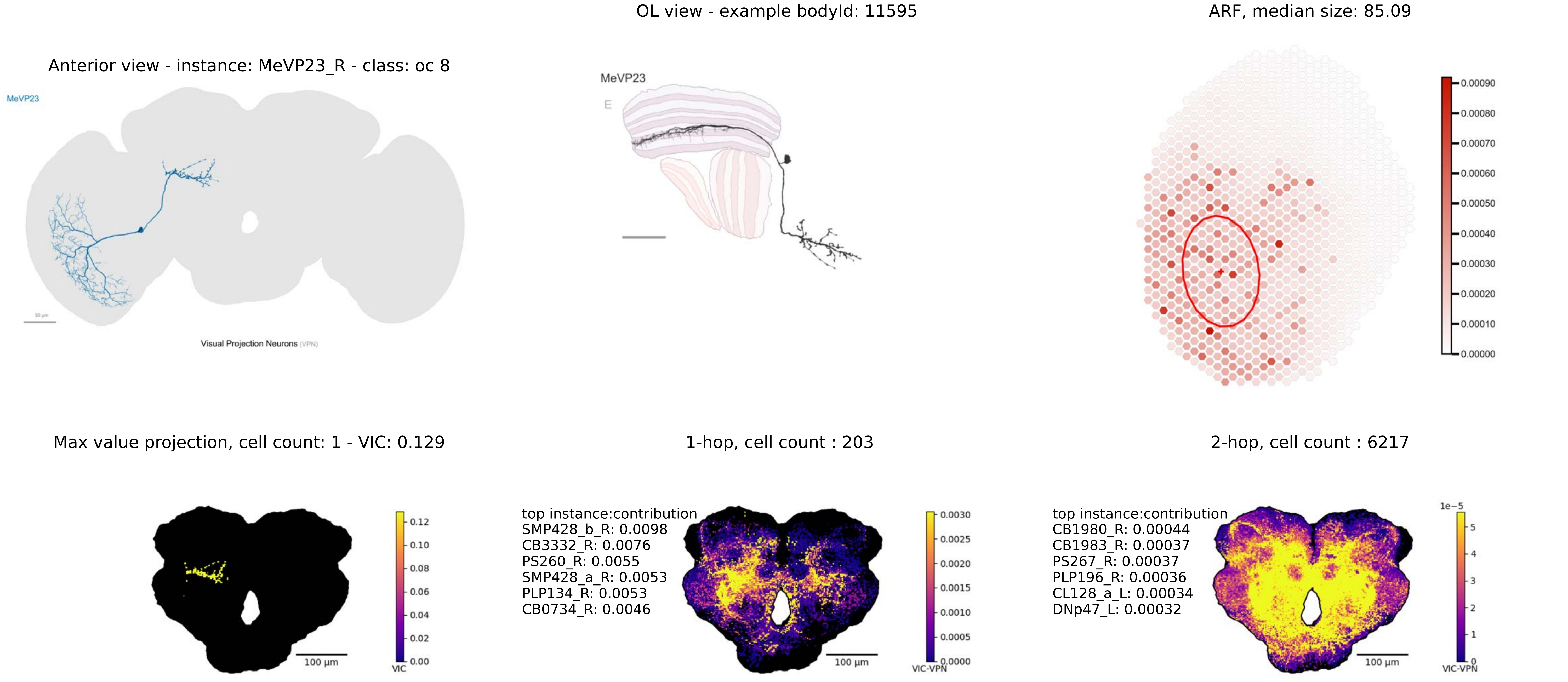
Anterior view - instance: MeVP23_R - class: oc 8
MeVP23
50 µm
Visual Projection Neurons (VPN)
OL view - example bodyId: 11595
MeVP23
E
ARF, median size: 85.09
0.00090
0.00080
0.00070
0.00060
0.00050
0.00040
0.00030
0.00020
0.00010
0.00000
Max value projection, cell count: 1 - VIC: 0.129
100 µm
0.12
0.10
0.08
0.06
0.04
0.02
0.00
VIC
1-hop, cell count : 203
top instance:contribution
SMP428_b_R: 0.0098
CB3332_R: 0.0076
PS260_R: 0.0055
SMP428_a_R: 0.0053
PLP134_R: 0.0053
CB0734_R: 0.0046
100 µm
0.0030
0.0025
0.0020
0.0015
0.0010
0.0005
0.0000
VIC-VPN
2-hop, cell count : 6217
top instance:contribution
CB1980_R: 0.00044
CB1983_R: 0.00037
PS267_R: 0.00037
PLP196_R: 0.00036
CL128_a_L: 0.00034
DNp47_L: 0.00032
100 µm
1e−5
5
4
3
2
1
0
VIC-VPN

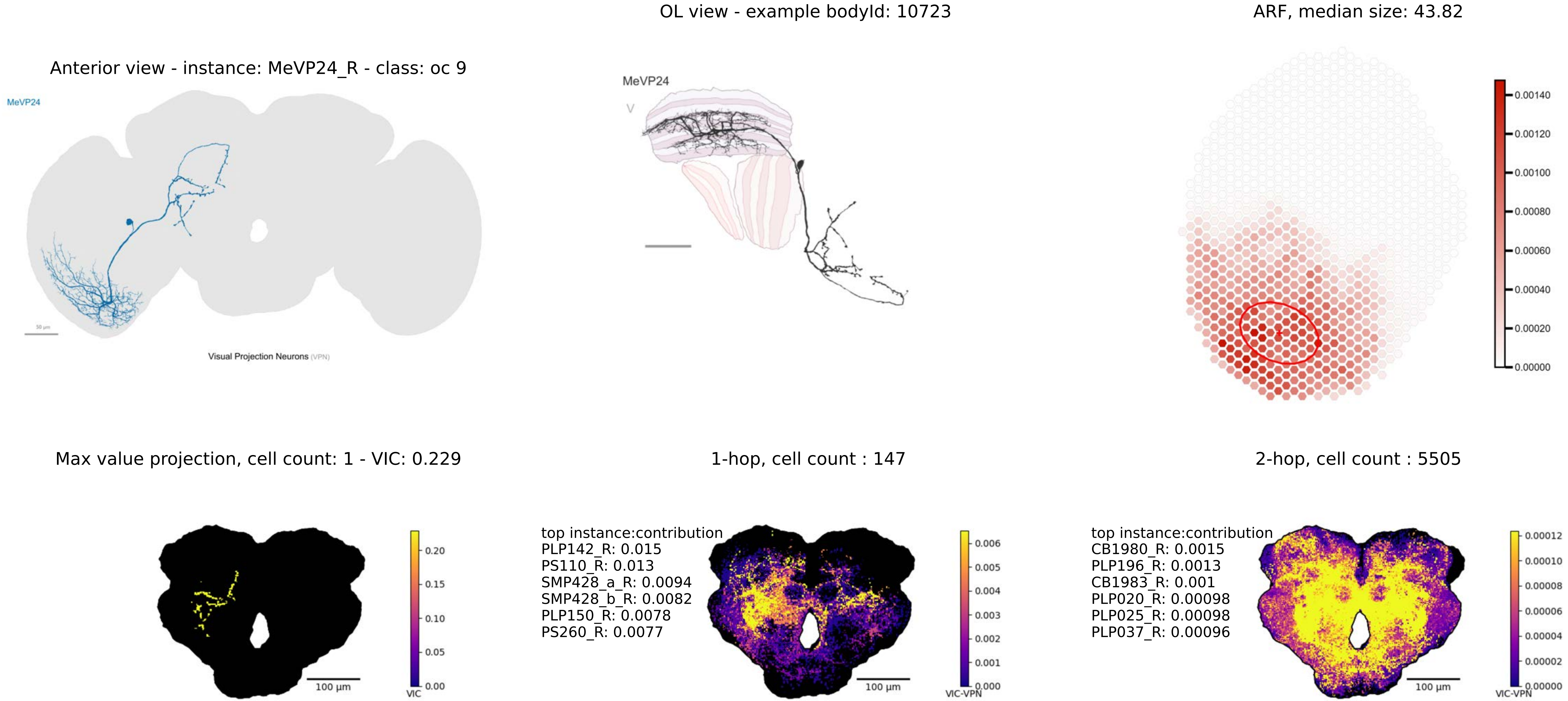
OL view - example bodyId: 10723
ARF, median size: 43.82
Anterior view - instance: MeVP24_R - class: oc 9
MeVP24
Visual Projection Neurons (VPN)
MeVP24
V
0.00140
0.00120
0.00100
0.00080
0.00060
0.00040
0.00020
0.00000
Max value projection, cell count: 1 - VIC: 0.229
1-hop, cell count : 147
2-hop, cell count : 5505
100 µm
VIC
0.20
0.15
0.10
0.05
0.00
top instance:contribution
PLP142_R: 0.015
PS110_R: 0.013
SMP428_a_R: 0.0094
SMP428_b_R: 0.0082
PLP150_R: 0.0078
PS260_R: 0.0077
VIC-VPN
0.006
0.005
0.004
0.003
0.002
0.001
0.000
top instance:contribution
CB1980_R: 0.0015
PLP196_R: 0.0013
CB1983_R: 0.001
PLP020_R: 0.00098
PLP025_R: 0.00098
PLP037_R: 0.00096
0.00012
0.00010
0.00008
0.00006
0.00004
0.00002
0.00000

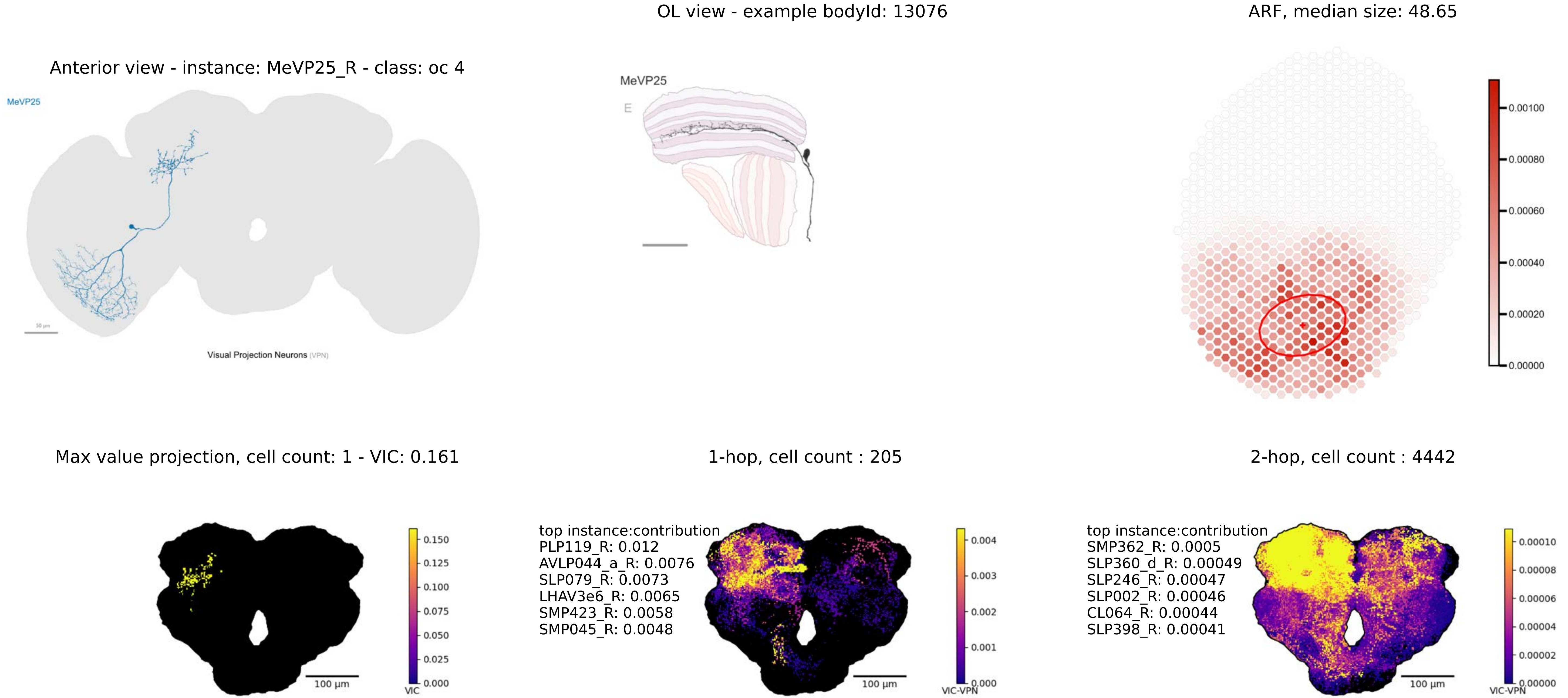
OL view - example bodyId: 13076
ARF, median size: 48.65
Anterior view - instance: MeVP25_R - class: oc 4
MeVP25
Visual Projection Neurons (VPN)
MeVP25
E
0.00100
0.00080
0.00060
0.00040
0.00020
0.00000
Max value projection, cell count: 1 - VIC: 0.161
1-hop, cell count : 205
2-hop, cell count : 4442
100 µm
0.150
0.125
0.100
0.075
0.050
0.025
0.000
VIC
top instance:contribution
PLP119_R: 0.012
AVLP044_a_R: 0.0076
SLP079_R: 0.0073
LHAV3e6_R: 0.0065
SMP423_R: 0.0058
SMP045_R: 0.0048
100 µm
0.004
0.003
0.002
0.001
0.000
VIC-VPN
top instance:contribution
SMP362_R: 0.0005
SLP360_d_R: 0.00049
SLP246_R: 0.00047
SLP002_R: 0.00046
CL064_R: 0.00044
SLP398_R: 0.00041
100 µm
0.00010
0.00008
0.00006
0.00004
0.00002
0.00000
VIC-VPN

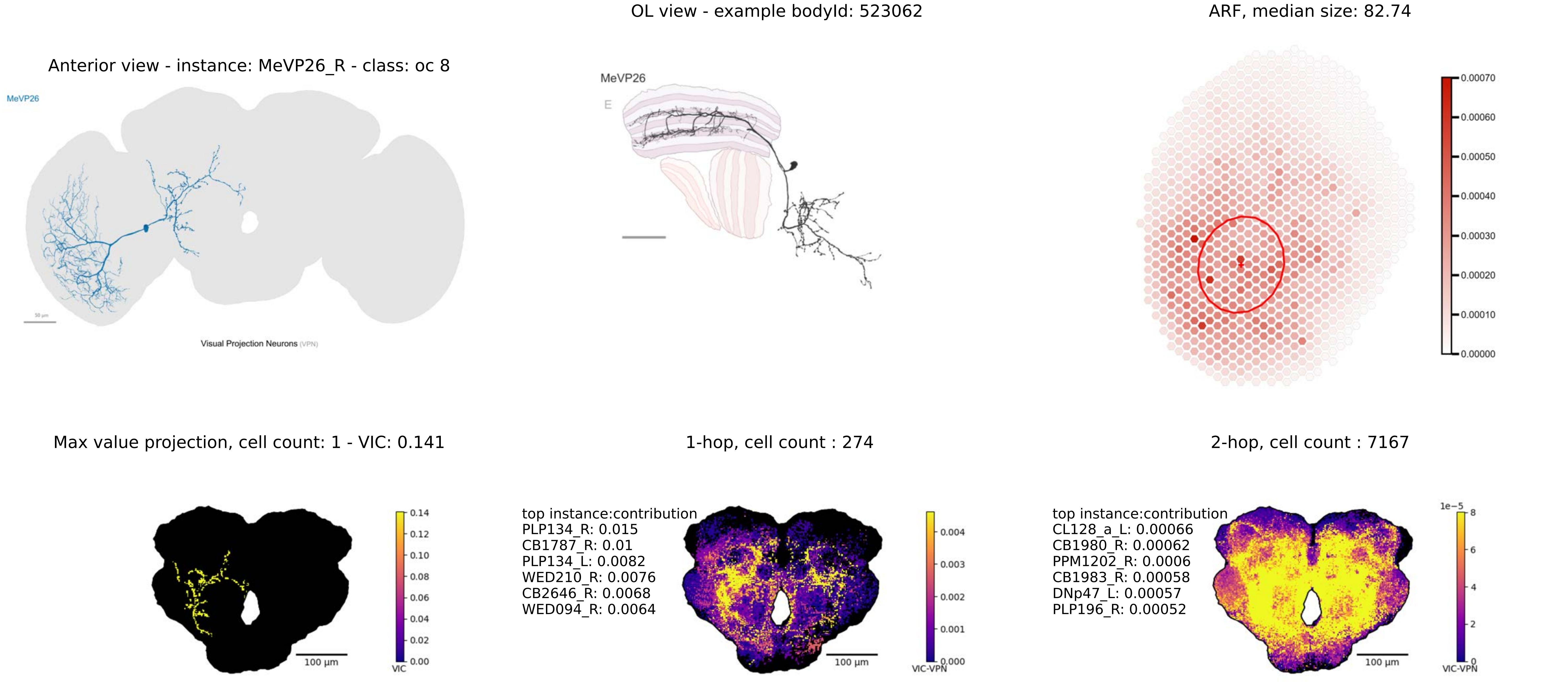
OL view - example bodyId: 523062
ARF, median size: 82.74
Anterior view - instance: MeVP26_R - class: oc 8
MeVP26
Visual Projection Neurons (VPN)
MeVP26
E
0.00070
0.00060
0.00050
0.00040
0.00030
0.00020
0.00010
0.00000
Max value projection, cell count: 1 - VIC: 0.141
1-hop, cell count : 274
2-hop, cell count : 7167
100 μm
VIC
0.14
0.12
0.10
0.08
0.06
0.04
0.02
0.00
top instance:contribution
PLP134_R: 0.015
CB1787_R: 0.01
PLP134_L: 0.0082
WED210_R: 0.0076
CB2646_R: 0.0068
WED094_R: 0.0064
100 μm
VIC-VPN
0.004
0.003
0.002
0.001
0.000
top instance:contribution
CL128_a_L: 0.00066
CB1980_R: 0.00062
PPM1202_R: 0.0006
CB1983_R: 0.00058
DNp47_L: 0.00057
PLP196_R: 0.00052
100 μm
1e−5
8
6
4
2
0
VIC-VPN

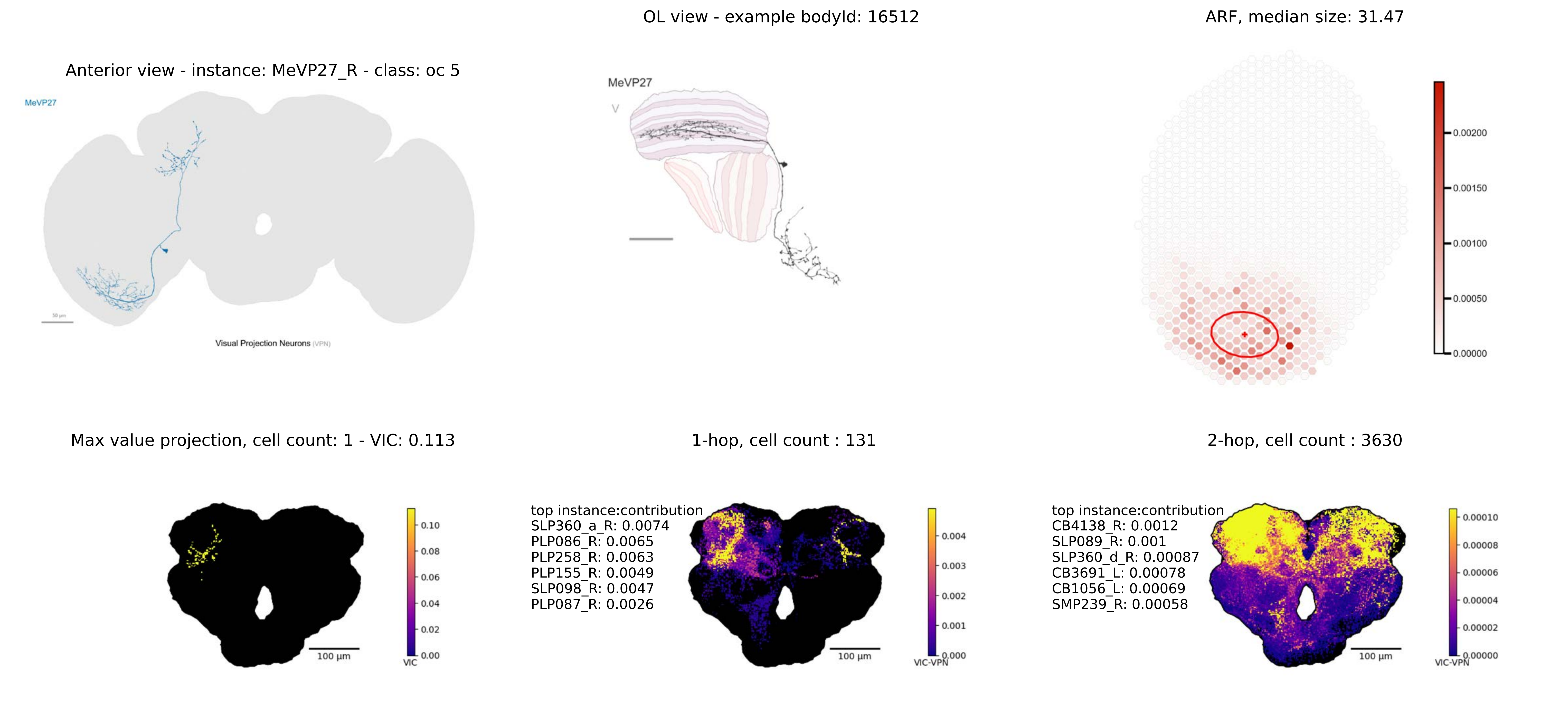

Anterior view - instance: MeVP27_R - class: oc 5
MeVP27
Visual Projection Neurons (VPN)
OL view - example bodyId: 16512
MeVP27
ARF, median size: 31.47
0.00200
0.00150
0.00100
0.00050
0.00000
Max value projection, cell count: 1 - VIC: 0.113
0.10
0.08
0.06
0.04
0.02
0.00
VIC
100 μm
1-hop, cell count : 131
top instance:contribution
SLP360_a_R: 0.0074
PLP086_R: 0.0065
PLP258_R: 0.0063
PLP155_R: 0.0049
SLP098_R: 0.0047
PLP087_R: 0.0026
0.004
0.003
0.002
0.001
0.000
VIC-VPN
100 μm
2-hop, cell count : 3630
top instance:contribution
CB4138_R: 0.0012
SLP089_R: 0.001
SLP360_d_R: 0.00087
CB3691_L: 0.00078
CB1056_L: 0.00069
SMP239_R: 0.00058
0.00010
0.00008
0.00006
0.00004
0.00002
0.00000
VIC-VPN
100 μm

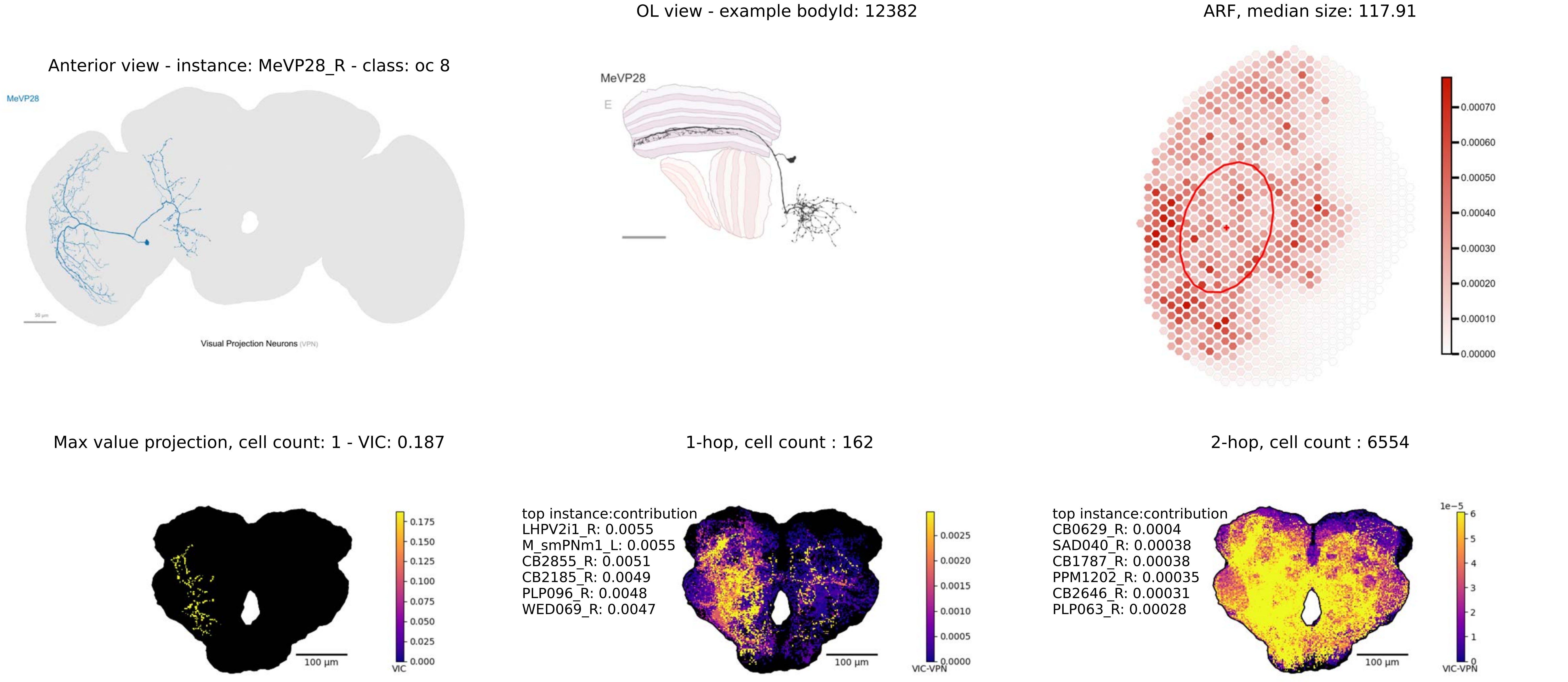

OL view - example bodyId: 12382
ARF, median size: 117.91
Anterior view - instance: MeVP28_R - class: oc 8
MeVP28
Visual Projection Neurons (VPN)
MeVP28
E
0.00070
0.00060
0.00050
0.00040
0.00030
0.00020
0.00010
0.00000
Max value projection, cell count: 1 - VIC: 0.187
1-hop, cell count : 162
2-hop, cell count : 6554
0.175
0.150
0.125
0.100
0.075
0.050
0.025
0.000
VIC
100 µm
top instance:contribution
LHPV2i1_R: 0.0055
M_smPNm1_L: 0.0055
CB2855_R: 0.0051
CB2185_R: 0.0049
PLP096_R: 0.0048
WED069_R: 0.0047
0.0025
0.0020
0.0015
0.0010
0.0005
0.0000
VIC-VPN
100 µm
top instance:contribution
CB0629_R: 0.0004
SAD040_R: 0.00038
CB1787_R: 0.00038
PPM1202_R: 0.00035
CB2646_R: 0.00031
PLP063_R: 0.00028
1e−5
6
5
4
3
2
1
0
VIC-VPN
100 µm

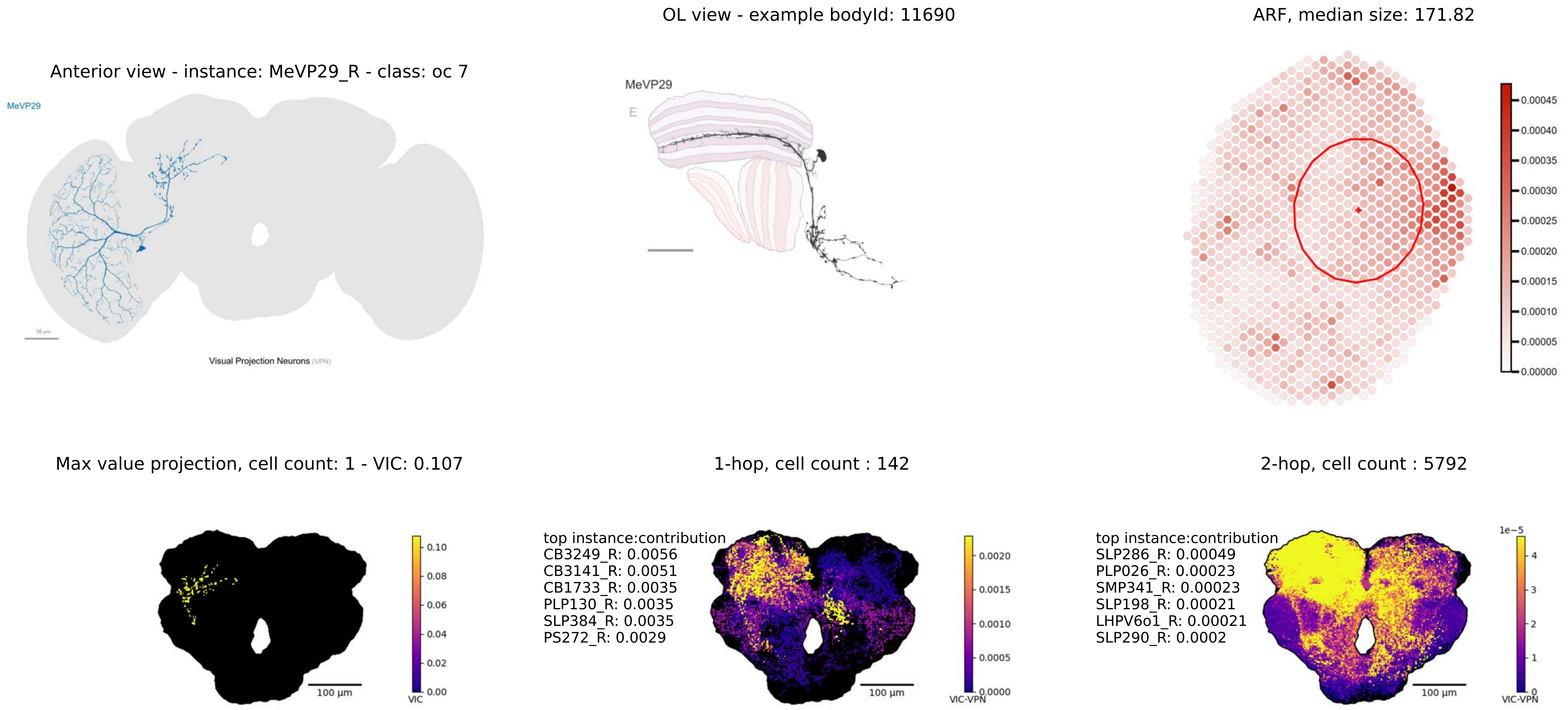
OL view - example bodyId: 11690
ARF, median size: 171.82
Anterior view - instance: MeVP29_R - class: oc 7
MeVP29
Visual Projection Neurons (VPN)
MeVP29
E
0.00045
0.00040
0.00035
0.00030
0.00025
0.00020
0.00015
0.00010
0.00005
0.00000
Max value projection, cell count: 1 - VIC: 0.107
1-hop, cell count : 142
2-hop, cell count : 5792
0.10
0.08
0.06
0.04
0.02
0.00
VIC
100 µm
top instance:contribution
CB3249_R: 0.0056
CB3141_R: 0.0051
CB1733_R: 0.0035
PLP130_R: 0.0035
SLP384_R: 0.0035
PS272_R: 0.0029
0.0020
0.0015
0.0010
0.0005
0.0000
VIC-VPN
100 µm
top instance:contribution
SLP286_R: 0.00049
PLP026_R: 0.00023
SMP341_R: 0.00023
SLP198_R: 0.00021
LHPV6o1_R: 0.00021
SLP290_R: 0.0002
1e−5
4
3
2
1
0
VIC-VPN
100 µm

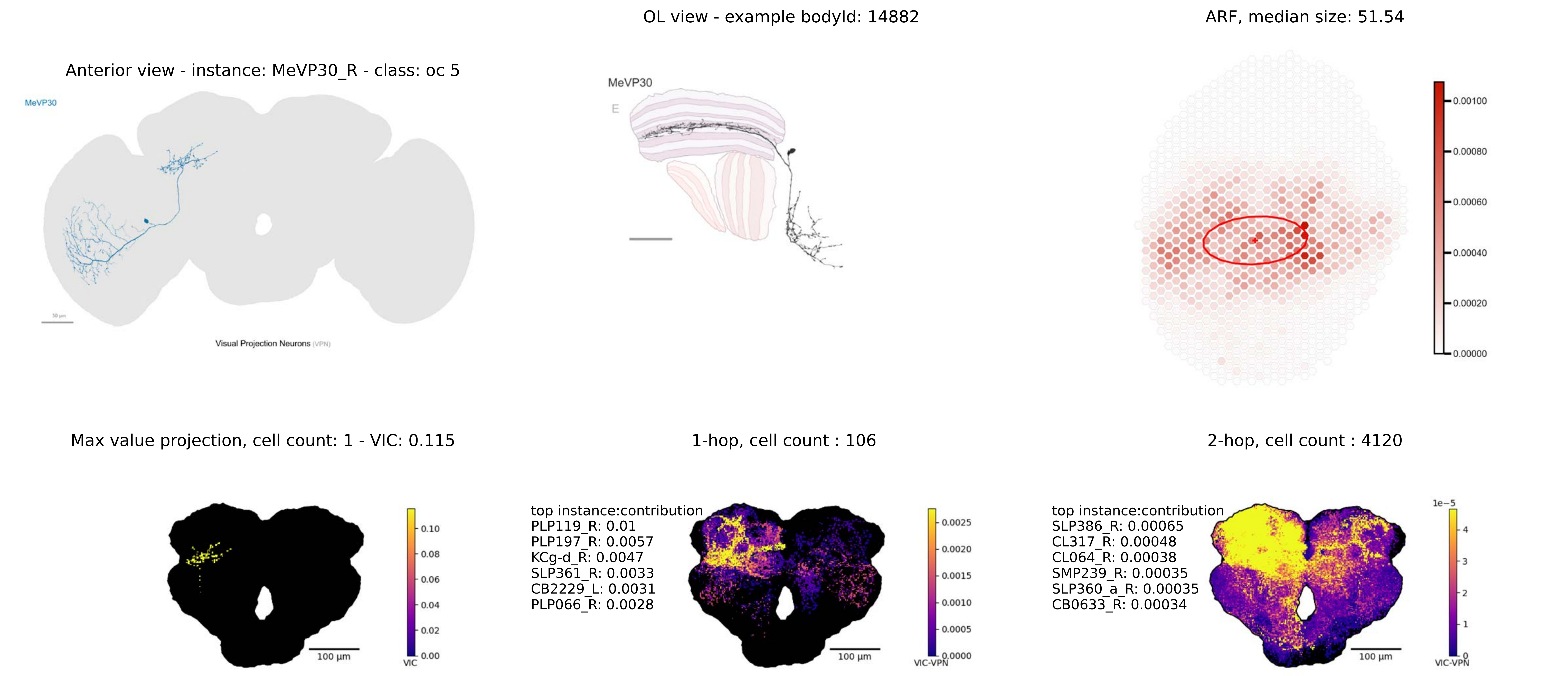
OL view - example bodyId: 14882
ARF, median size: 51.54
Anterior view - instance: MeVP30_R - class: oc 5
MeVP30
Visual Projection Neurons (VPN)
MeVP30
E
0.00100
0.00080
0.00060
0.00040
0.00020
0.00000
Max value projection, cell count: 1 - VIC: 0.115
1-hop, cell count : 106
2-hop, cell count : 4120
0.10
0.08
0.06
0.04
0.02
0.00
VIC
100 µm
top instance:contribution
PLP119_R: 0.01
PLP197_R: 0.0057
KCg-d_R: 0.0047
SLP361_R: 0.0033
CB2229_L: 0.0031
PLP066_R: 0.0028
0.0025
0.0020
0.0015
0.0010
0.0005
0.0000
VIC-VPN
100 µm
top instance:contribution
SLP386_R: 0.00065
CL317_R: 0.00048
CL064_R: 0.00038
SMP239_R: 0.00035
SLP360_a_R: 0.00035
CB0633_R: 0.00034
1e−5
4
3
2
1
0
VIC-VPN
100 µm

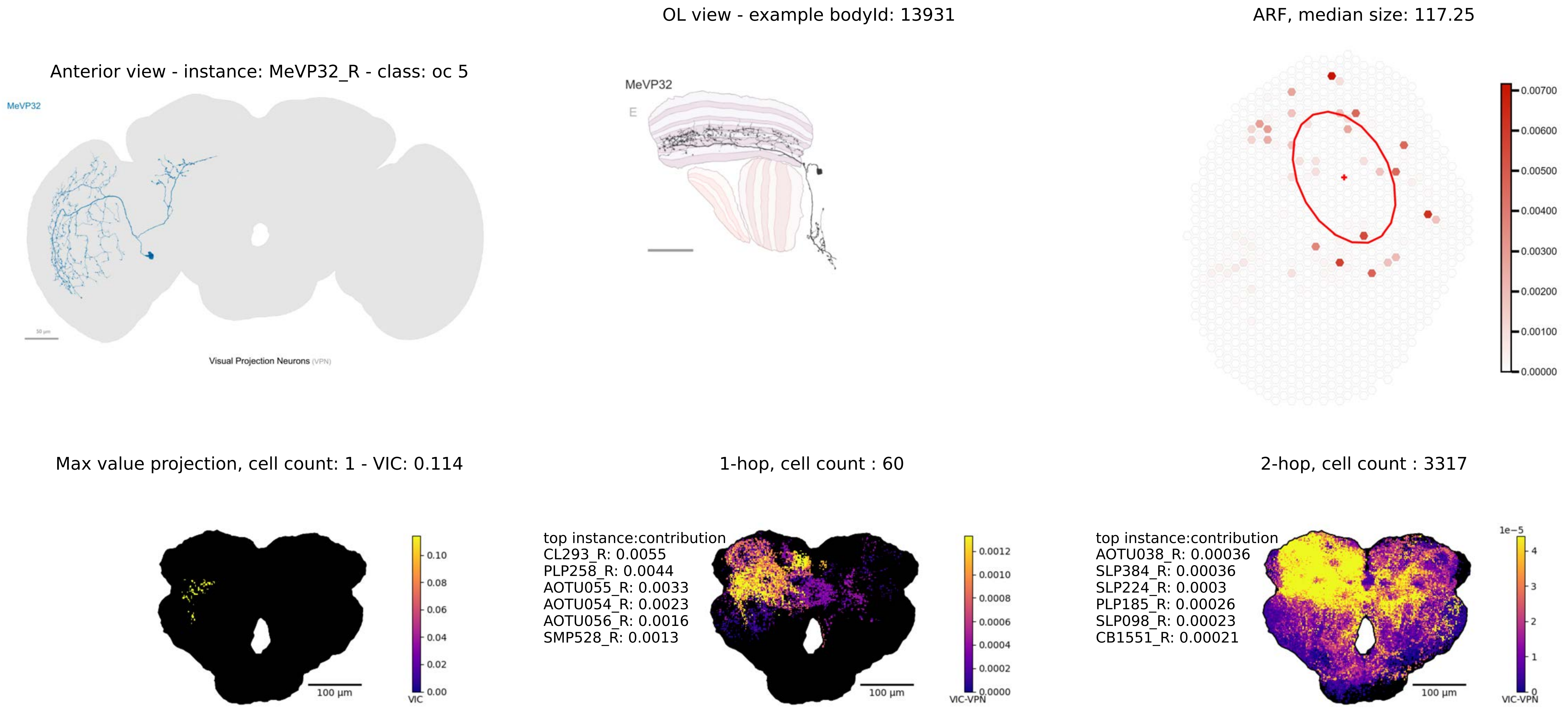

OL view - example bodyId: 13931
ARF, median size: 117.25
Anterior view - instance: MeVP32_R - class: oc 5
MeVP32
Visual Projection Neurons (VPN)
MeVP32
E
0.00700
0.00600
0.00500
0.00400
0.00300
0.00200
0.00100
0.00000
Max value projection, cell count: 1 - VIC: 0.114
1-hop, cell count : 60
2-hop, cell count : 3317
top instance:contribution
CL293_R: 0.0055
PLP258_R: 0.0044
AOTU055_R: 0.0033
AOTU054_R: 0.0023
AOTU056_R: 0.0016
SMP528_R: 0.0013
top instance:contribution
AOTU038_R: 0.00036
SLP384_R: 0.00036
SLP224_R: 0.0003
PLP185_R: 0.00026
SLP098_R: 0.00023
CB1551_R: 0.00021
0.10
0.08
0.06
0.04
0.02
0.00
VIC
100 µm
0.0012
0.0010
0.0008
0.0006
0.0004
0.0002
0.0000
VIC-VPN
100 µm
1e−5
4
3
2
1
0
VIC-VPN
100 µm

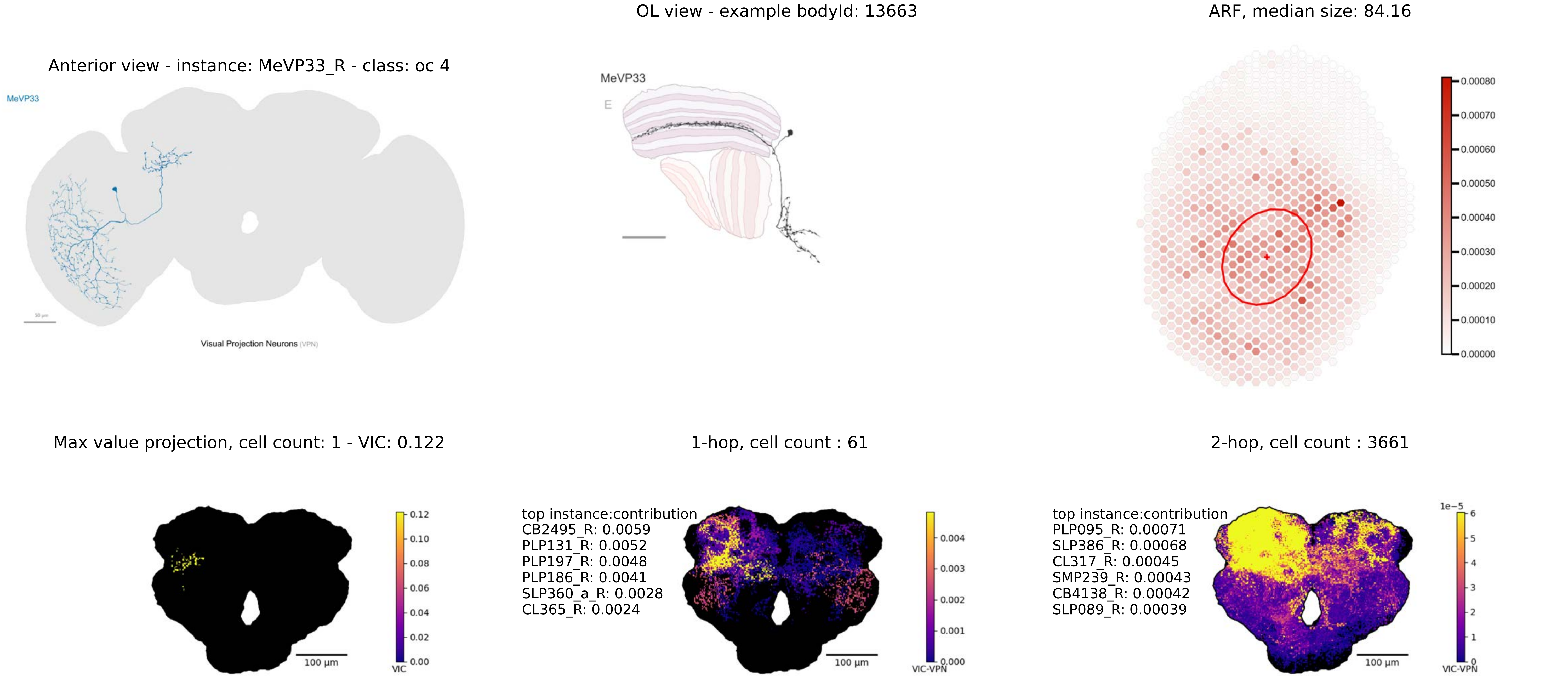
OL view - example bodyId: 13663
ARF, median size: 84.16
Anterior view - instance: MeVP33_R - class: oc 4
MeVP33
Visual Projection Neurons (VPN)
MeVP33
E
0.00080
0.00070
0.00060
0.00050
0.00040
0.00030
0.00020
0.00010
0.00000
Max value projection, cell count: 1 - VIC: 0.122
1-hop, cell count : 61
2-hop, cell count : 3661
0.12
0.10
0.08
0.06
0.04
0.02
0.00
VIC
100 µm
top instance:contribution
CB2495_R: 0.0059
PLP131_R: 0.0052
PLP197_R: 0.0048
PLP186_R: 0.0041
SLP360_a_R: 0.0028
CL365_R: 0.0024
0.004
0.003
0.002
0.001
0.000
VIC-VPN
100 µm
top instance:contribution
PLP095_R: 0.00071
SLP386_R: 0.00068
CL317_R: 0.00045
SMP239_R: 0.00043
CB4138_R: 0.00042
SLP089_R: 0.00039
1e−5
6
5
4
3
2
1
0
VIC-VPN
100 µm

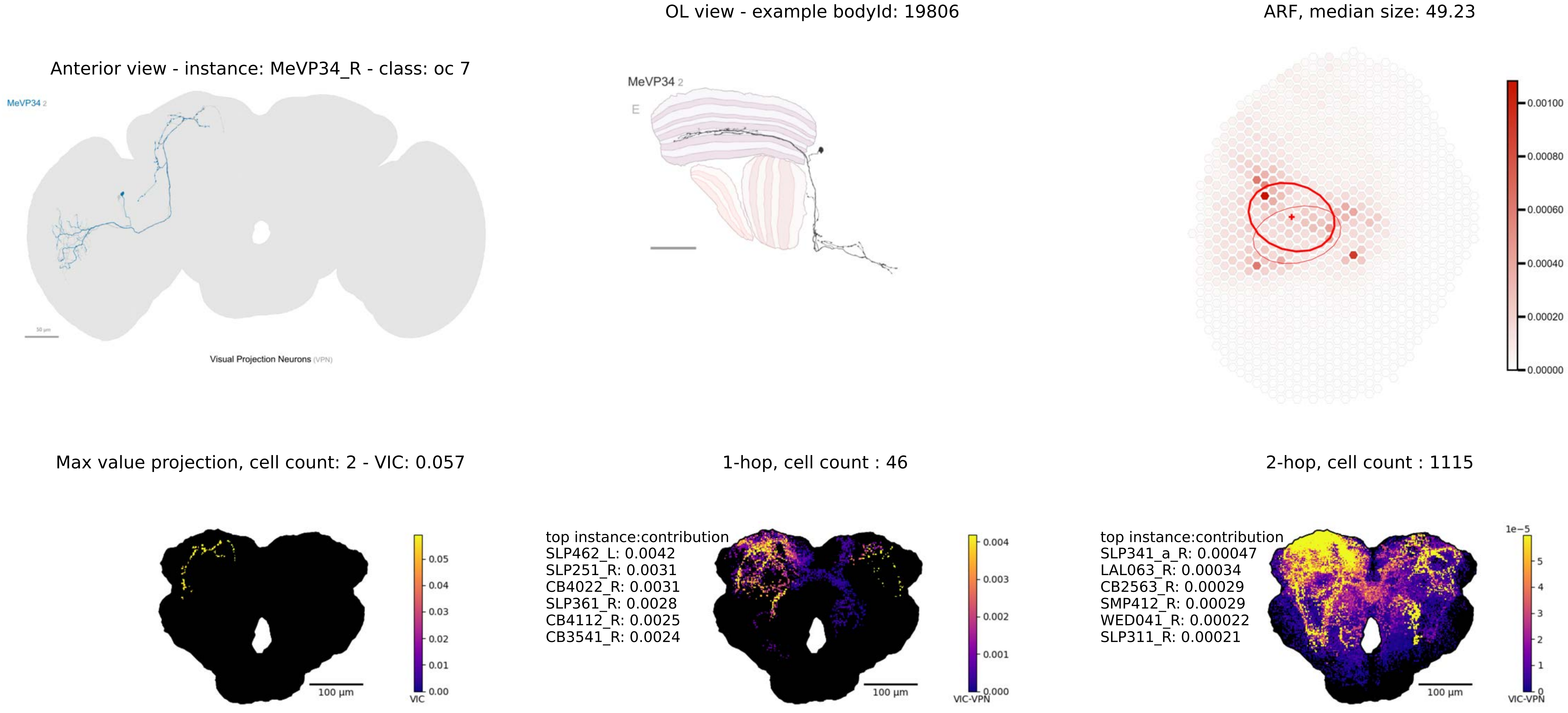
OL view - example bodyId: 19806
ARF, median size: 49.23
Anterior view - instance: MeVP34_R - class: oc 7
MeVP34 2
Visual Projection Neurons (VPN)
MeVP34 2
E
0.00100
0.00080
0.00060
0.00040
0.00020
0.00000
Max value projection, cell count: 2 - VIC: 0.057
1-hop, cell count : 46
2-hop, cell count : 1115
0.05
0.04
0.03
0.02
0.01
0.00
VIC
100 μm
top instance:contribution
SLP462_L: 0.0042
SLP251_R: 0.0031
CB4022_R: 0.0031
SLP361_R: 0.0028
CB4112_R: 0.0025
CB3541_R: 0.0024
0.004
0.003
0.002
0.001
0.000
VIC-VPN
100 μm
top instance:contribution
SLP341_a_R: 0.00047
LAL063_R: 0.00034
CB2563_R: 0.00029
SMP412_R: 0.00029
WED041_R: 0.00022
SLP311_R: 0.00021
1e−5
5
4
3
2
1
0
VIC-VPN
100 μm

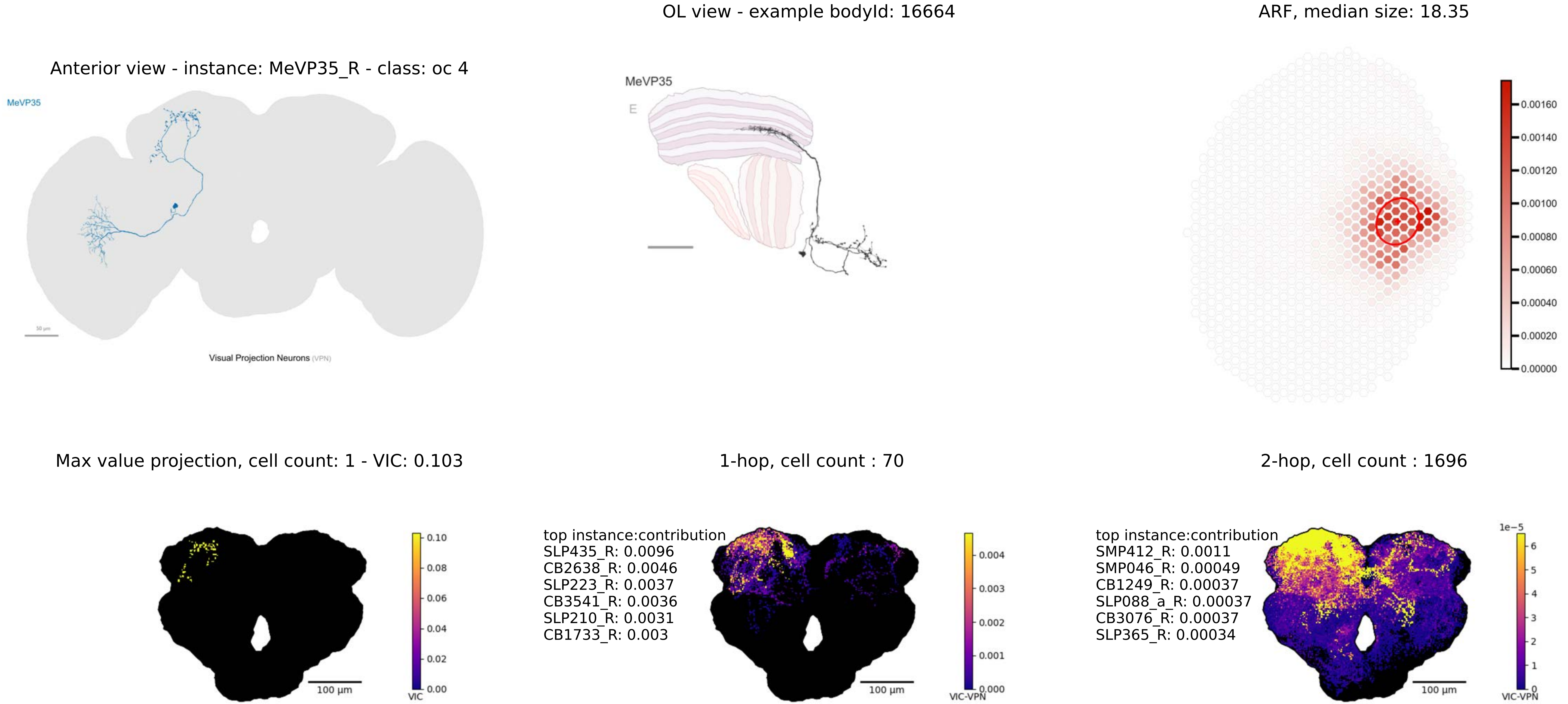
OL view - example bodyId: 16664
ARF, median size: 18.35
Anterior view - instance: MeVP35_R - class: oc 4
MeVP35
50 μm
Visual Projection Neurons (VPN)
MeVP35
E
0.00160
0.00140
0.00120
0.00100
0.00080
0.00060
0.00040
0.00020
0.00000
Max value projection, cell count: 1 - VIC: 0.103
1-hop, cell count : 70
2-hop, cell count : 1696
0.10
0.08
0.06
0.04
0.02
0.00
VIC
100 μm
top instance:contribution
SLP435_R: 0.0096
CB2638_R: 0.0046
SLP223_R: 0.0037
CB3541_R: 0.0036
SLP210_R: 0.0031
CB1733_R: 0.003
0.004
0.003
0.002
0.001
0.000
VIC-VPN
100 μm
top instance:contribution
SMP412_R: 0.0011
SMP046_R: 0.00049
CB1249_R: 0.00037
SLP088_a_R: 0.00037
CB3076_R: 0.00037
SLP365_R: 0.00034
1e−5
6
5
4
3
2
1
0
VIC-VPN
100 μm

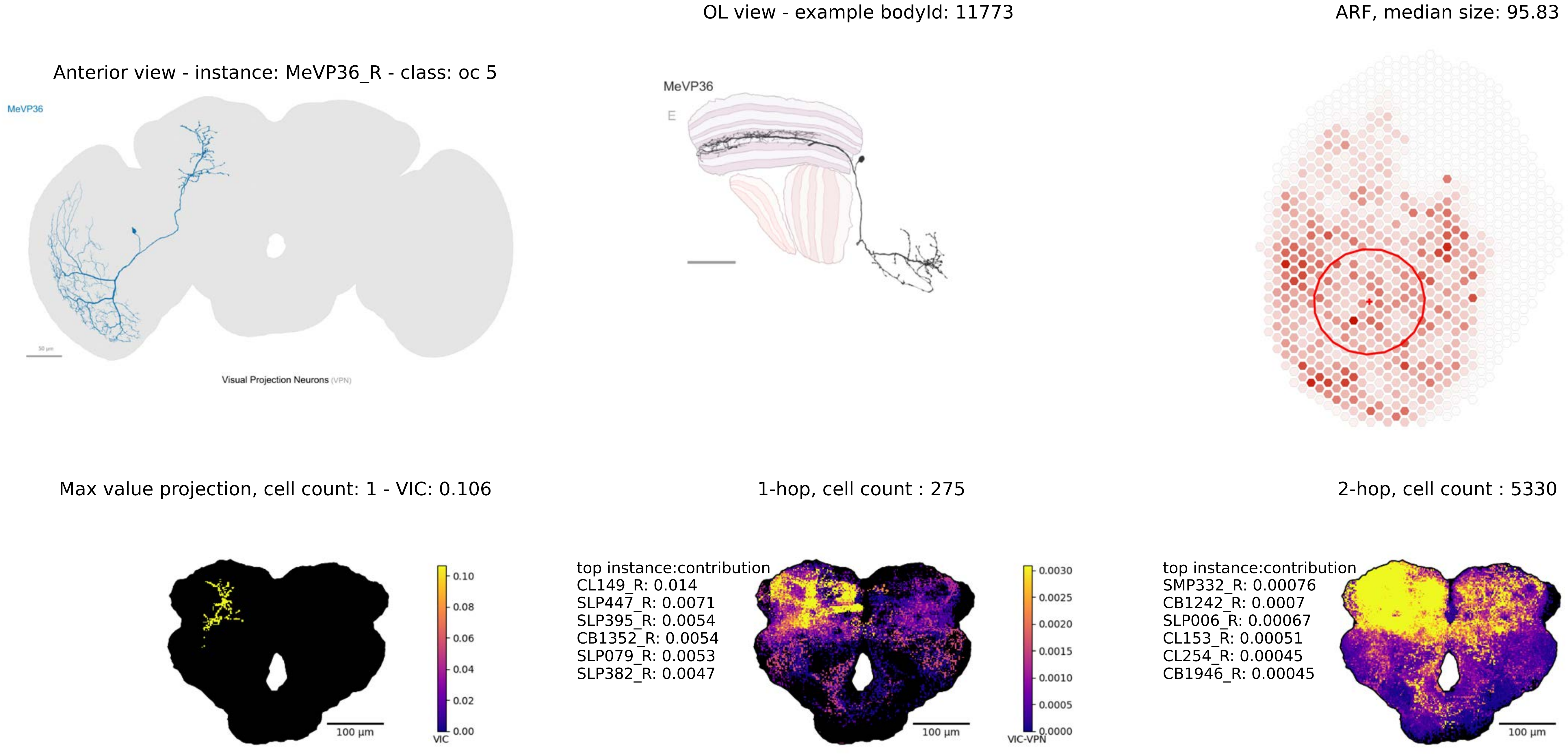

OL view - example bodyId: 11773
ARF, median size: 95.83
Anterior view - instance: MeVP36_R - class: oc 5
MeVP36
Visual Projection Neurons (VPN)
MeVP36
E
0.00070
0.00060
0.00050
0.00040
0.00030
0.00020
0.00010
0.00000
Max value projection, cell count: 1 - VIC: 0.106
1-hop, cell count : 275
2-hop, cell count : 5330
0.10
0.08
0.06
0.04
0.02
0.00
VIC
100 µm
top instance:contribution
CL149_R: 0.014
SLP447_R: 0.0071
SLP395_R: 0.0054
CB1352_R: 0.0054
SLP079_R: 0.0053
SLP382_R: 0.0047
0.0030
0.0025
0.0020
0.0015
0.0010
0.0005
0.0000
VIC-VPN
100 µm
top instance:contribution
SMP332_R: 0.00076
CB1242_R: 0.0007
SLP006_R: 0.00067
CL153_R: 0.00051
CL254_R: 0.00045
CB1946_R: 0.00045
1e−5
8
6
4
2
0
VIC-VPN
100 µm

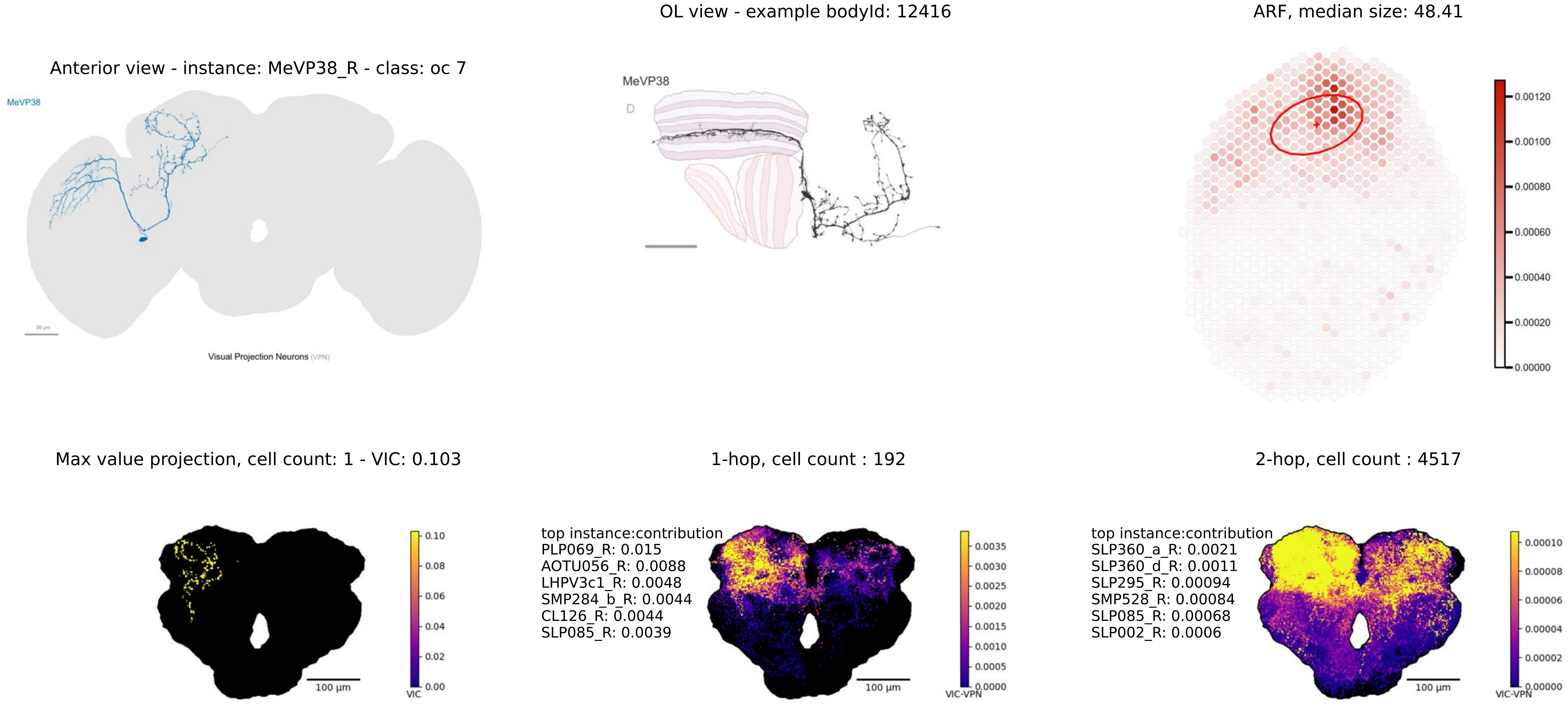

OL view - example bodyId: 12416
ARF, median size: 48.41
Anterior view - instance: MeVP38_R - class: oc 7
MeVP38
Visual Projection Neurons (VPN)
MeVP38
D
0.00120
0.00100
0.00080
0.00060
0.00040
0.00020
0.00000
Max value projection, cell count: 1 - VIC: 0.103
1-hop, cell count : 192
2-hop, cell count : 4517
0.10
0.08
0.06
0.04
0.02
0.00
VIC
100 µm
top instance:contribution
PLP069_R: 0.015
AOTU056_R: 0.0088
LHPV3c1_R: 0.0048
SMP284_b_R: 0.0044
CL126_R: 0.0044
SLP085_R: 0.0039
0.0035
0.0030
0.0025
0.0020
0.0015
0.0010
0.0005
0.0000
VIC-VPN
100 µm
top instance:contribution
SLP360_a_R: 0.0021
SLP360_d_R: 0.0011
SLP295_R: 0.00094
SMP528_R: 0.00084
SLP085_R: 0.00068
SLP002_R: 0.0006
0.00010
0.00008
0.00006
0.00004
0.00002
0.00000
VIC-VPN
100 µm

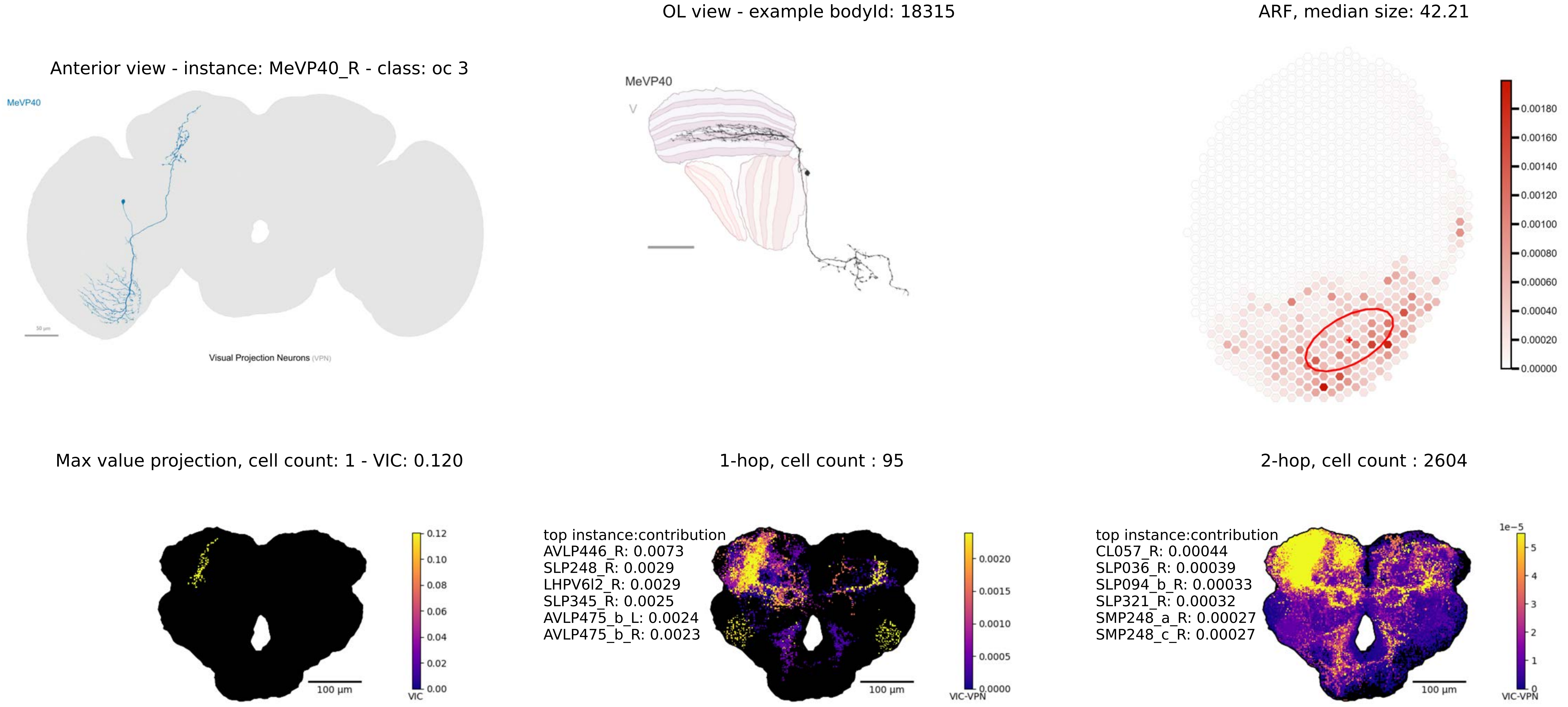

OL view - example bodyId: 18315
ARF, median size: 42.21
Anterior view - instance: MeVP40_R - class: oc 3
MeVP40
Visual Projection Neurons (VPN)
MeVP40
Max value projection, cell count: 1 - VIC: 0.120
1-hop, cell count : 95
2-hop, cell count : 2604
top instance:contribution
AVLP446_R: 0.0073
SLP248_R: 0.0029
LHPV6l2_R: 0.0029
SLP345_R: 0.0025
AVLP475_b_L: 0.0024
AVLP475_b_R: 0.0023
top instance:contribution
CL057_R: 0.00044
SLP036_R: 0.00039
SLP094_b_R: 0.00033
SLP321_R: 0.00032
SMP248_a_R: 0.00027
SMP248_c_R: 0.00027
100 µm
VIC
VIC-VPN

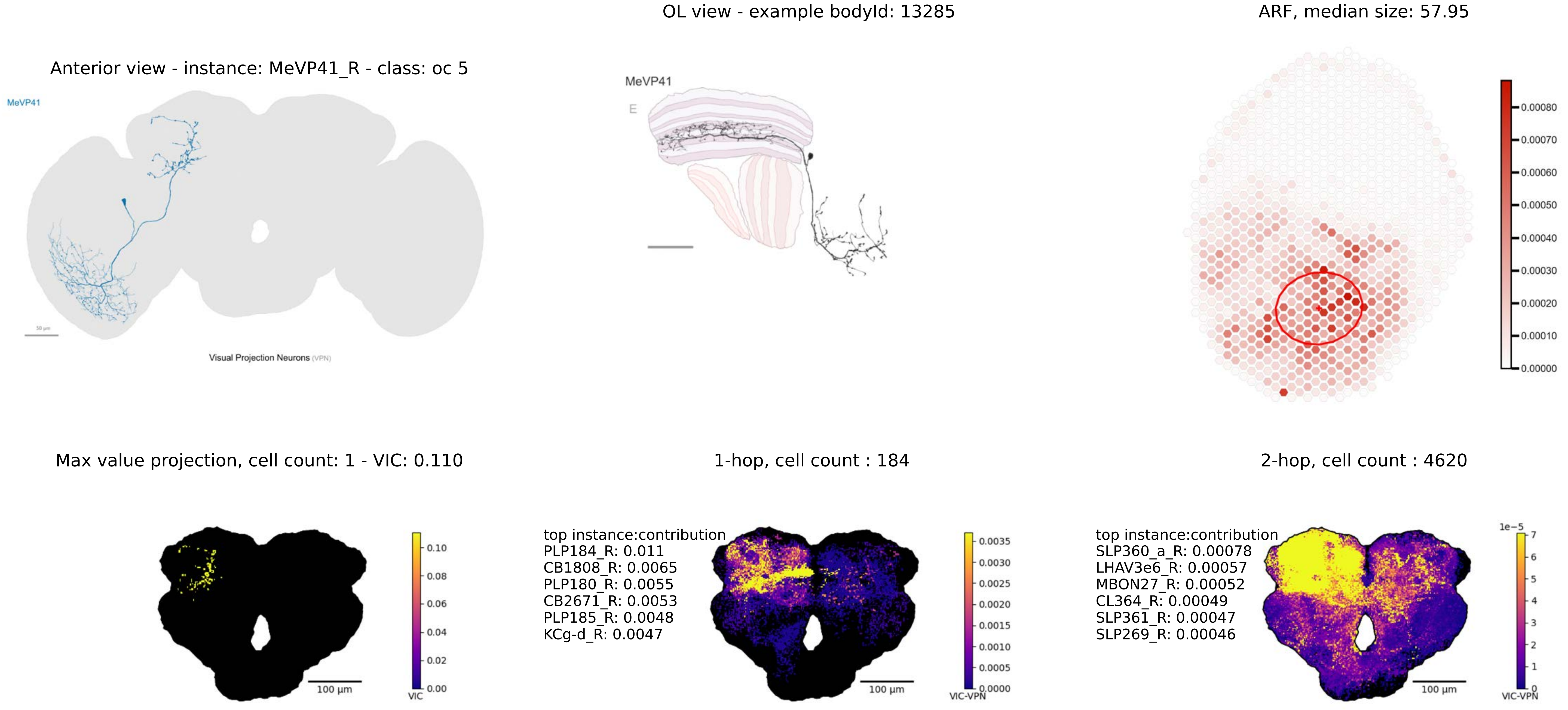

Anterior view - instance: MeVP41_R - class: oc 5
MeVP41
Visual Projection Neurons (VPN)
OL view - example bodyId: 13285
MeVP41
E
ARF, median size: 57.95
0.00080
0.00070
0.00060
0.00050
0.00040
0.00030
0.00020
0.00010
0.00000
Max value projection, cell count: 1 - VIC: 0.110
100 µm
0.10
0.08
0.06
0.04
0.02
0.00
VIC
1-hop, cell count : 184
top instance:contribution
PLP184_R: 0.011
CB1808_R: 0.0065
PLP180_R: 0.0055
CB2671_R: 0.0053
PLP185_R: 0.0048
KCg-d_R: 0.0047
100 µm
0.0035
0.0030
0.0025
0.0020
0.0015
0.0010
0.0005
0.0000
VIC-VPN
2-hop, cell count : 4620
top instance:contribution
SLP360_a_R: 0.00078
LHAV3e6_R: 0.00057
MBON27_R: 0.00052
CL364_R: 0.00049
SLP361_R: 0.00047
SLP269_R: 0.00046
100 µm
1e−5
7
6
5
4
3
2
1
0
VIC-VPN

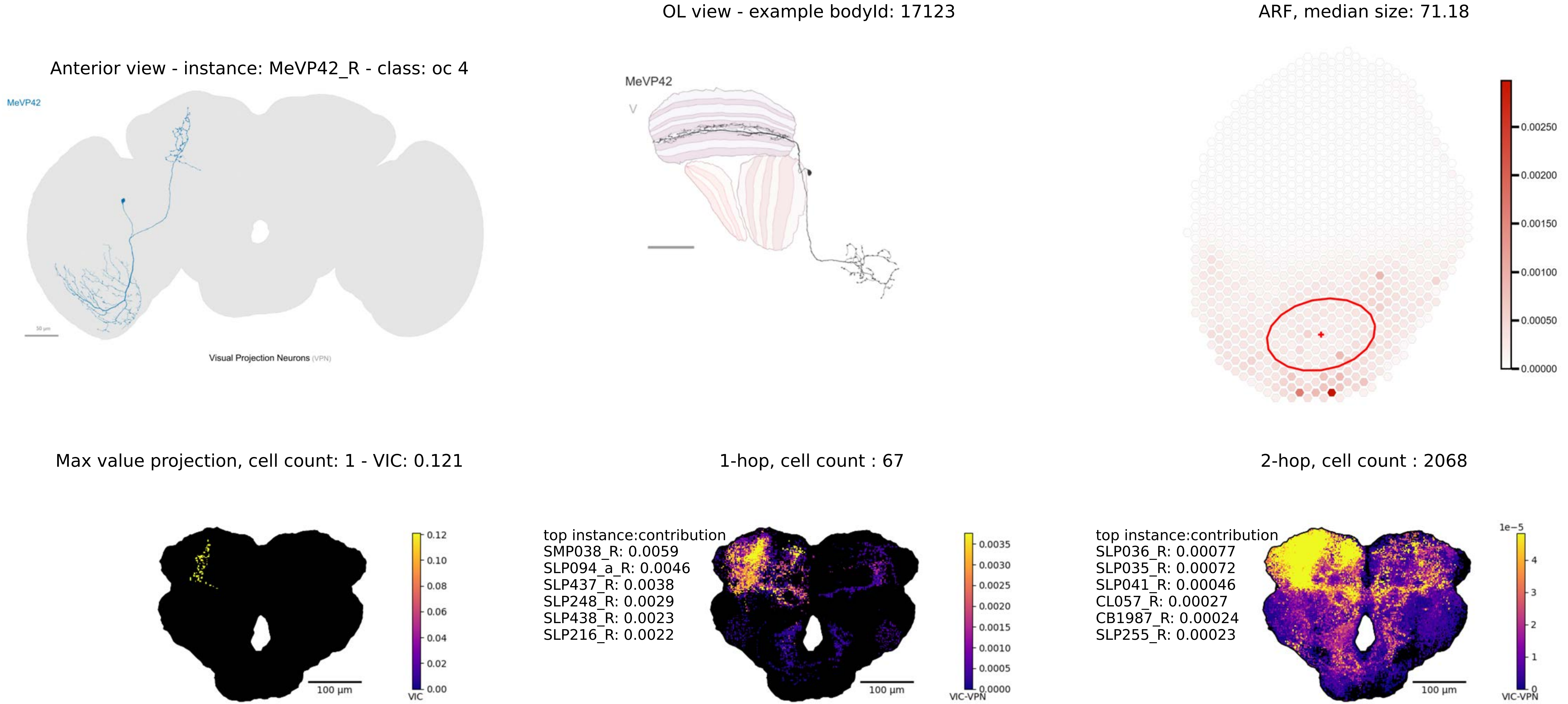
OL view - example bodyId: 17123
ARF, median size: 71.18
Anterior view - instance: MeVP42_R - class: oc 4
MeVP42
Visual Projection Neurons (VPN)
MeVP42
0.00250
0.00200
0.00150
0.00100
0.00050
0.00000
Max value projection, cell count: 1 - VIC: 0.121
1-hop, cell count : 67
2-hop, cell count : 2068
100 µm
VIC
0.12
0.10
0.08
0.06
0.04
0.02
0.00
top instance:contribution
SMP038_R: 0.0059
SLP094_a_R: 0.0046
SLP437_R: 0.0038
SLP248_R: 0.0029
SLP438_R: 0.0023
SLP216_R: 0.0022
100 µm
VIC-VPN
0.0035
0.0030
0.0025
0.0020
0.0015
0.0010
0.0005
0.0000
top instance:contribution
SLP036_R: 0.00077
SLP035_R: 0.00072
SLP041_R: 0.00046
CL057_R: 0.00027
CB1987_R: 0.00024
SLP255_R: 0.00023
100 µm
1e−5
4
3
2
1
0
VIC-VPN

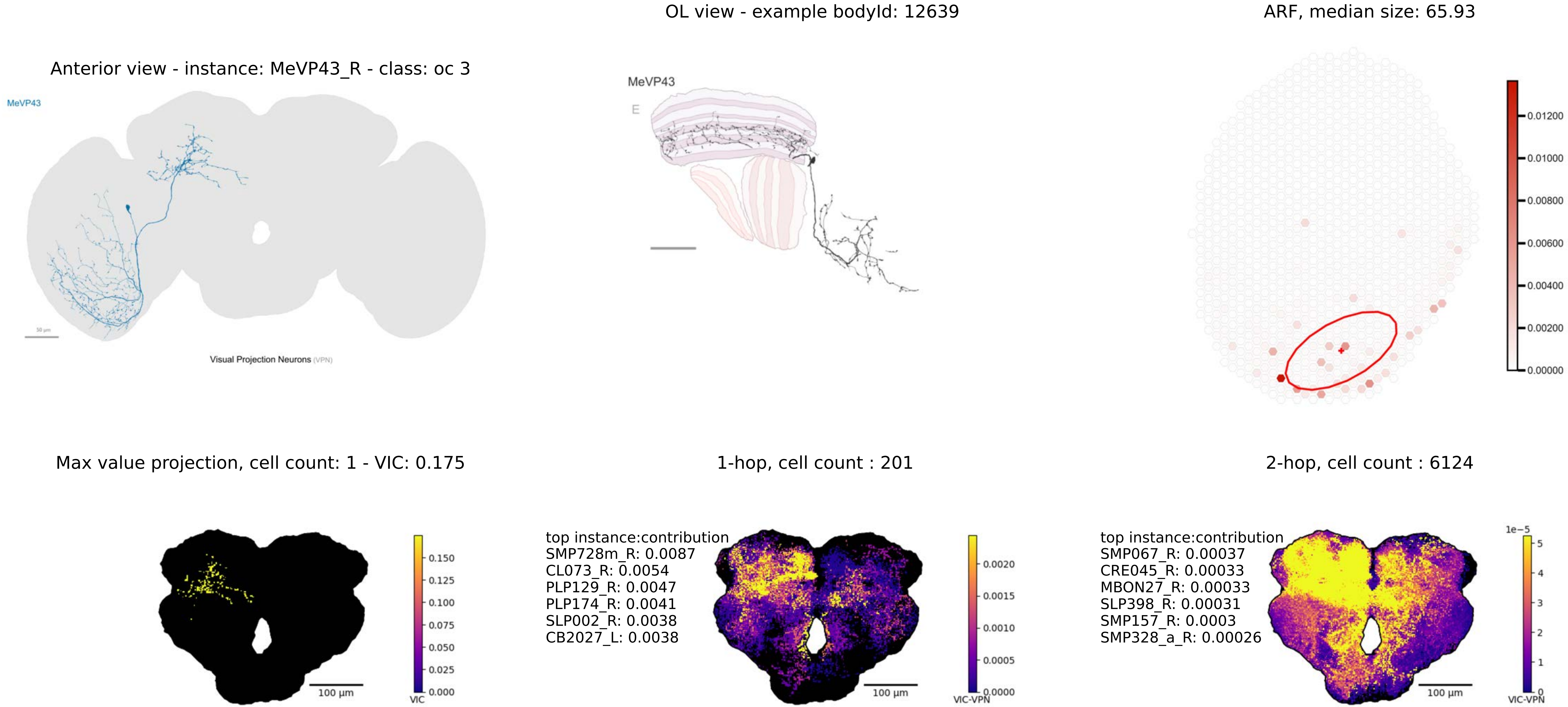
OL view - example bodyId: 12639
ARF, median size: 65.93
Anterior view - instance: MeVP43_R - class: oc 3
MeVP43
Visual Projection Neurons (VPN)
MeVP43
E
0.01200
0.01000
0.00800
0.00600
0.00400
0.00200
0.00000
Max value projection, cell count: 1 - VIC: 0.175
1-hop, cell count : 201
2-hop, cell count : 6124
0.150
0.125
0.100
0.075
0.050
0.025
0.000
VIC
100 µm
top instance:contribution
SMP728m_R: 0.0087
CL073_R: 0.0054
PLP129_R: 0.0047
PLP174_R: 0.0041
SLP002_R: 0.0038
CB2027_L: 0.0038
0.0020
0.0015
0.0010
0.0005
0.0000
VIC-VPN
100 µm
top instance:contribution
SMP067_R: 0.00037
CRE045_R: 0.00033
MBON27_R: 0.00033
SLP398_R: 0.00031
SMP157_R: 0.0003
SMP328_a_R: 0.00026
1e−5
5
4
3
2
1
0
VIC-VPN
100 µm

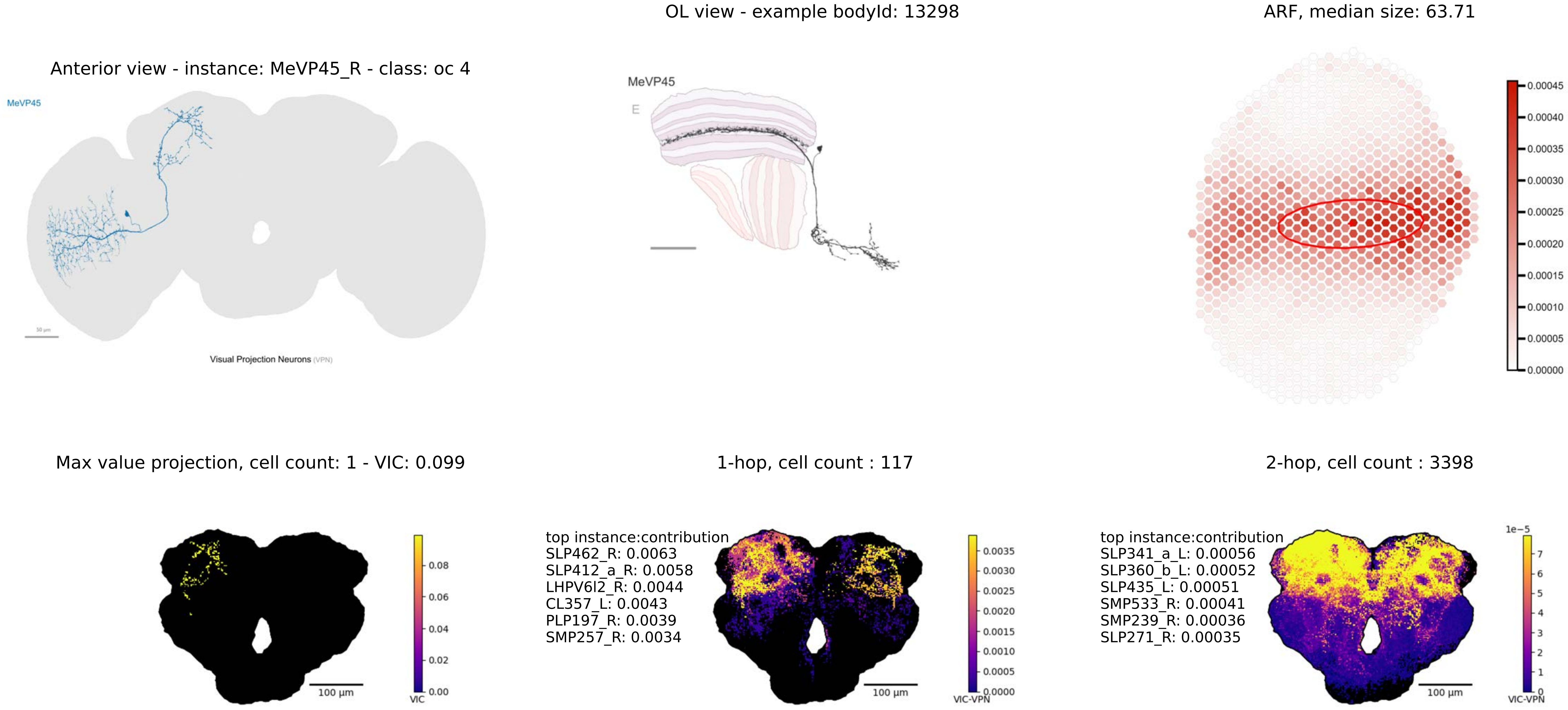
OL view - example bodyId: 13298
ARF, median size: 63.71
Anterior view - instance: MeVP45_R - class: oc 4
MeVP45
Visual Projection Neurons (VPN)
MeVP45
E
0.00045
0.00040
0.00035
0.00030
0.00025
0.00020
0.00015
0.00010
0.00005
0.00000
Max value projection, cell count: 1 - VIC: 0.099
1-hop, cell count : 117
2-hop, cell count : 3398
100 µm
0.08
0.06
0.04
0.02
0.00
VIC
top instance:contribution
SLP462_R: 0.0063
SLP412_a_R: 0.0058
LHPV6l2_R: 0.0044
CL357_L: 0.0043
PLP197_R: 0.0039
SMP257_R: 0.0034
100 µm
0.0035
0.0030
0.0025
0.0020
0.0015
0.0010
0.0005
0.0000
VIC-VPN
top instance:contribution
SLP341_a_L: 0.00056
SLP360_b_L: 0.00052
SLP435_L: 0.00051
SMP533_R: 0.00041
SMP239_R: 0.00036
SLP271_R: 0.00035
100 µm
1e−5
7
6
5
4
3
2
1
0
VIC-VPN

Anterior view - instance: MeVP46_R - class: oc 7

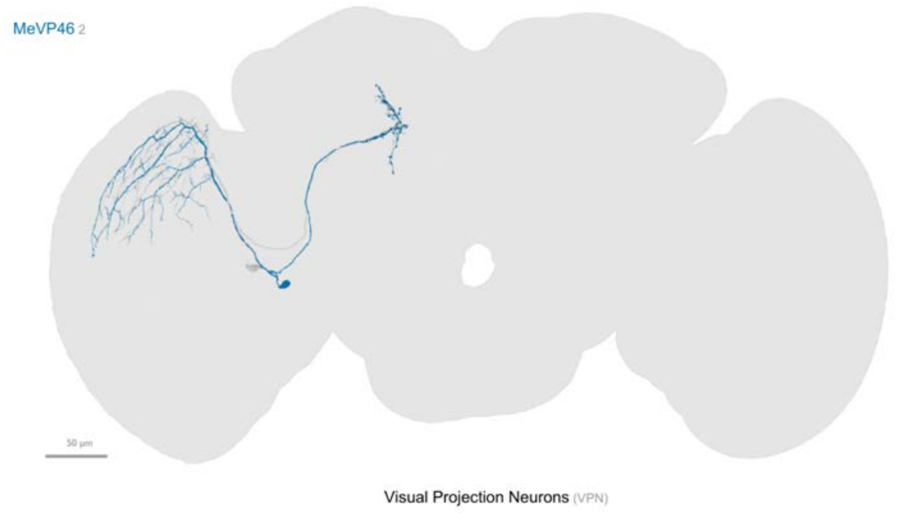


OL view - example bodyId: 14142

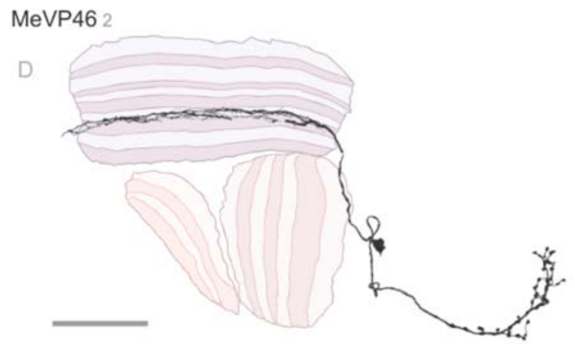


ARF, median size: 51.6

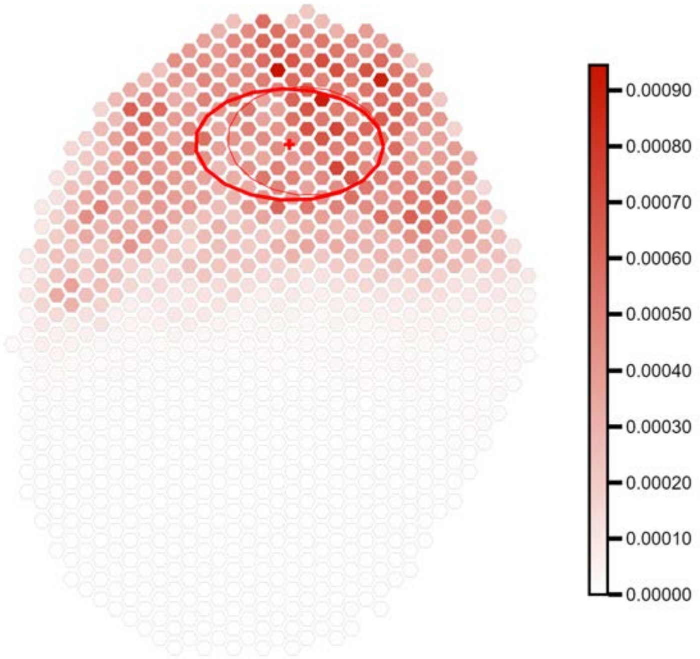


Max value projection, cell count: 2 - VIC: 0.147

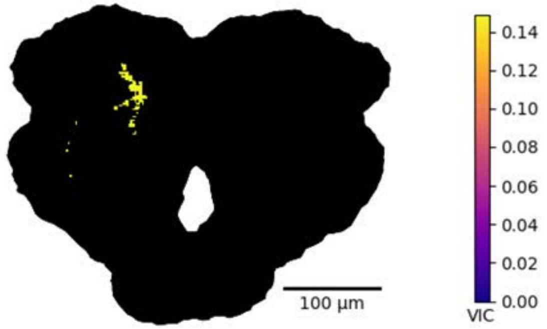


1-hop, cell count : 96

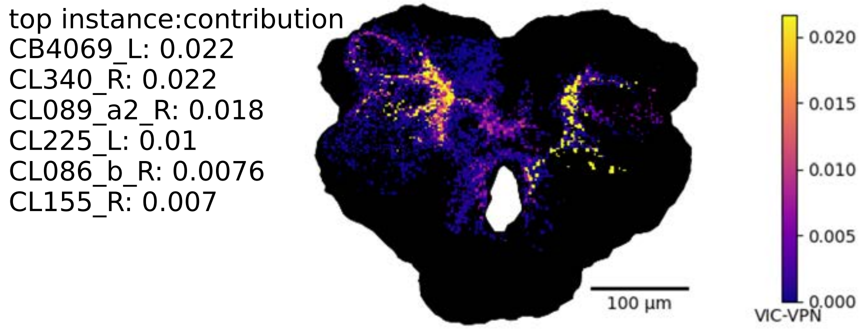


2-hop, cell count : 1965

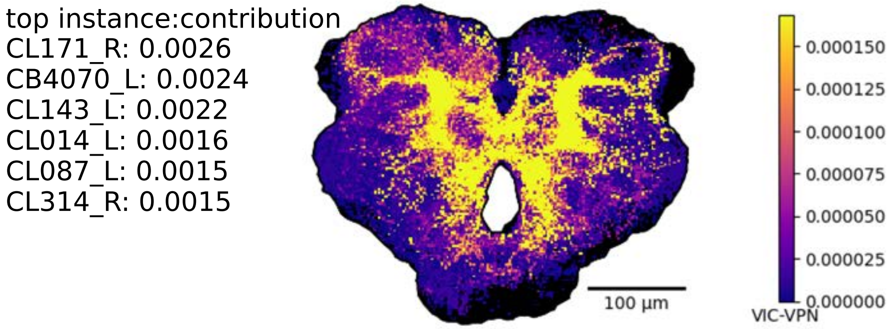

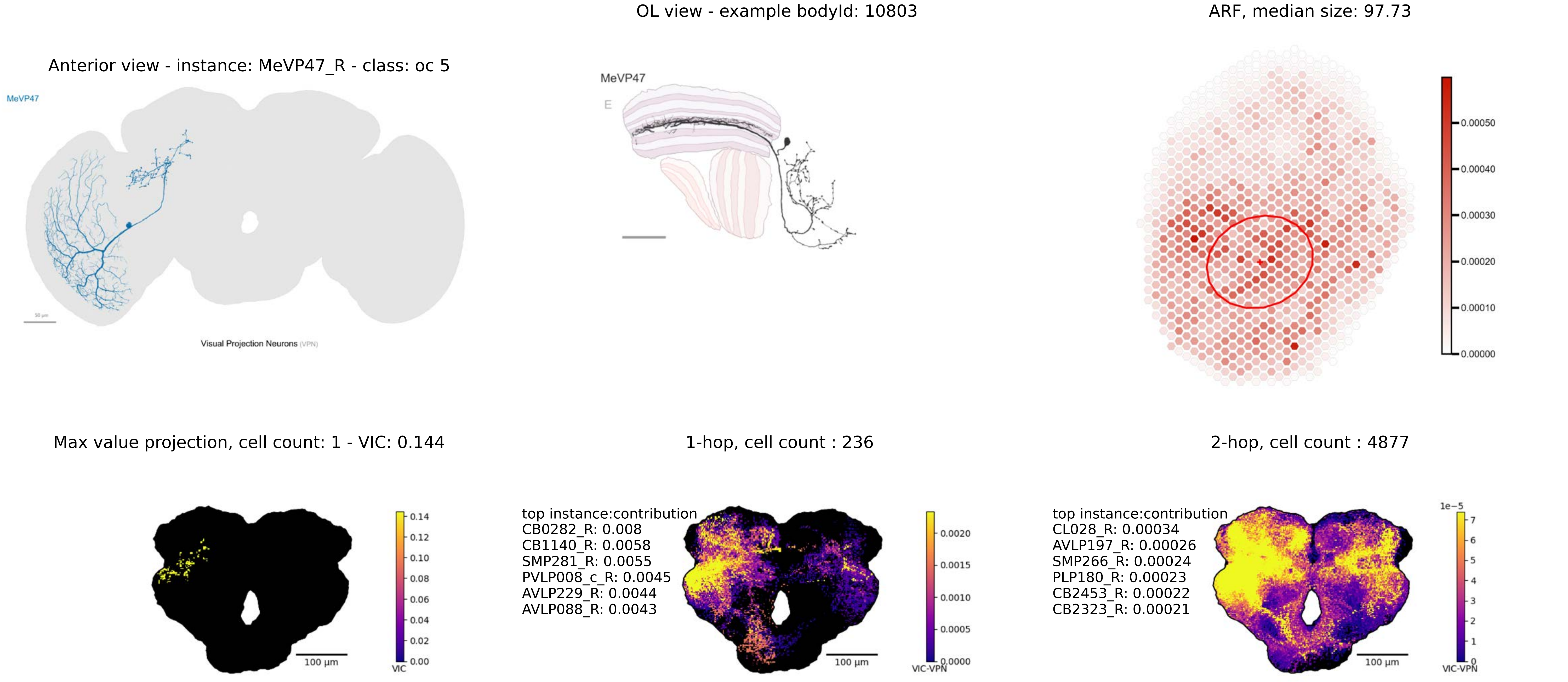
OL view - example bodyId: 10803
ARF, median size: 97.73
Anterior view - instance: MeVP47_R - class: oc 5
MeVP47
Visual Projection Neurons (VPN)
MeVP47
E
0.00050
0.00040
0.00030
0.00020
0.00010
0.00000
Max value projection, cell count: 1 - VIC: 0.144
1-hop, cell count : 236
2-hop, cell count : 4877
0.14
0.12
0.10
0.08
0.06
0.04
0.02
0.00
VIC
100 µm
top instance:contribution
CB0282_R: 0.008
CB1140_R: 0.0058
SMP281_R: 0.0055
PVLP008_c_R: 0.0045
AVLP229_R: 0.0044
AVLP088_R: 0.0043
0.0020
0.0015
0.0010
0.0005
0.0000
VIC-VPN
100 µm
top instance:contribution
CL028_R: 0.00034
AVLP197_R: 0.00026
SMP266_R: 0.00024
PLP180_R: 0.00023
CB2453_R: 0.00022
CB2323_R: 0.00021
1e−5
7
6
5
4
3
2
1
0
VIC-VPN
100 µm

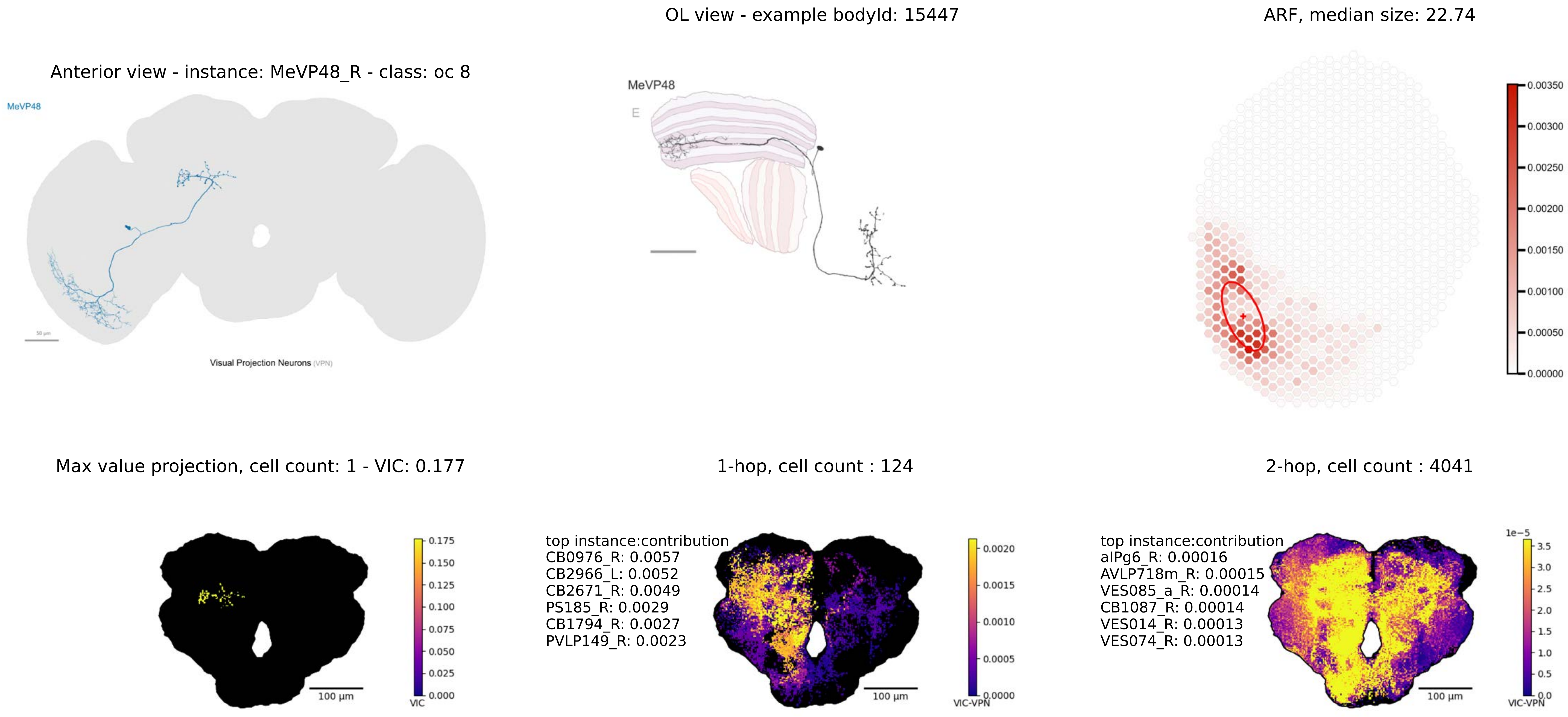
OL view - example bodyId: 15447
ARF, median size: 22.74
Anterior view - instance: MeVP48_R - class: oc 8
MeVP48
Visual Projection Neurons (VPN)
MeVP48
E
0.00350
0.00300
0.00250
0.00200
0.00150
0.00100
0.00050
0.00000
Max value projection, cell count: 1 - VIC: 0.177
1-hop, cell count : 124
2-hop, cell count : 4041
0.175
0.150
0.125
0.100
0.075
0.050
0.025
0.000
VIC
100 µm
top instance:contribution
CB0976_R: 0.0057
CB2966_L: 0.0052
CB2671_R: 0.0049
PS185_R: 0.0029
CB1794_R: 0.0027
PVLP149_R: 0.0023
0.0020
0.0015
0.0010
0.0005
0.0000
VIC-VPN
100 µm
top instance:contribution
aIPg6_R: 0.00016
AVLP718m_R: 0.00015
VES085_a_R: 0.00014
CB1087_R: 0.00014
VES014_R: 0.00013
VES074_R: 0.00013
1e−5
3.5
3.0
2.5
2.0
1.5
1.0
0.5
0.0
VIC-VPN
100 µm

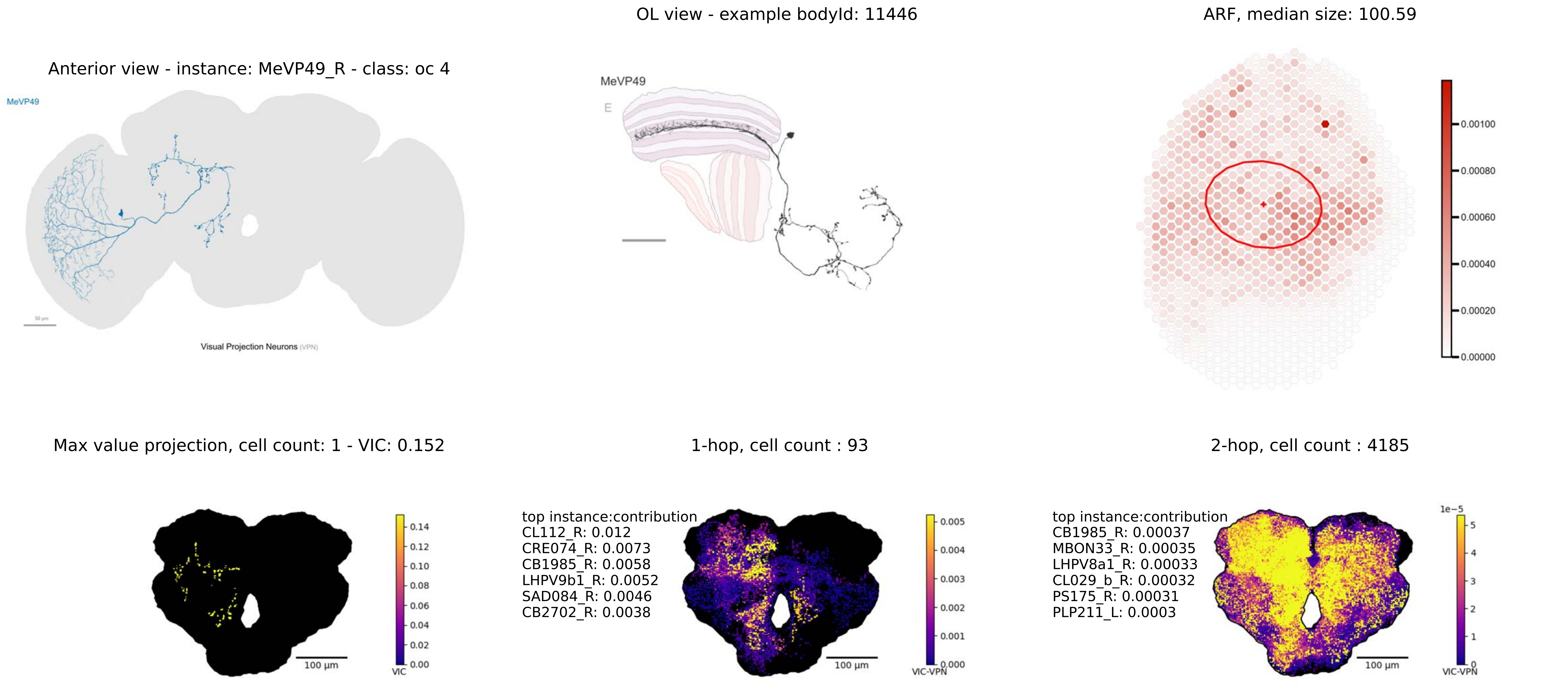

OL view - example bodyId: 11446
ARF, median size: 100.59
Anterior view - instance: MeVP49_R - class: oc 4
MeVP49
Visual Projection Neurons (VPN)
MeVP49
E
0.00100
0.00080
0.00060
0.00040
0.00020
0.00000
Max value projection, cell count: 1 - VIC: 0.152
1-hop, cell count : 93
2-hop, cell count : 4185
0.14
0.12
0.10
0.08
0.06
0.04
0.02
0.00
VIC
100 µm
top instance:contribution
CL112_R: 0.012
CRE074_R: 0.0073
CB1985_R: 0.0058
LHPV9b1_R: 0.0052
SAD084_R: 0.0046
CB2702_R: 0.0038
0.005
0.004
0.003
0.002
0.001
0.000
VIC-VPN
100 µm
top instance:contribution
CB1985_R: 0.00037
MBON33_R: 0.00035
LHPV8a1_R: 0.00033
CL029_b_R: 0.00032
PS175_R: 0.00031
PLP211_L: 0.0003
1e−5
5
4
3
2
1
0
VIC-VPN
100 µm

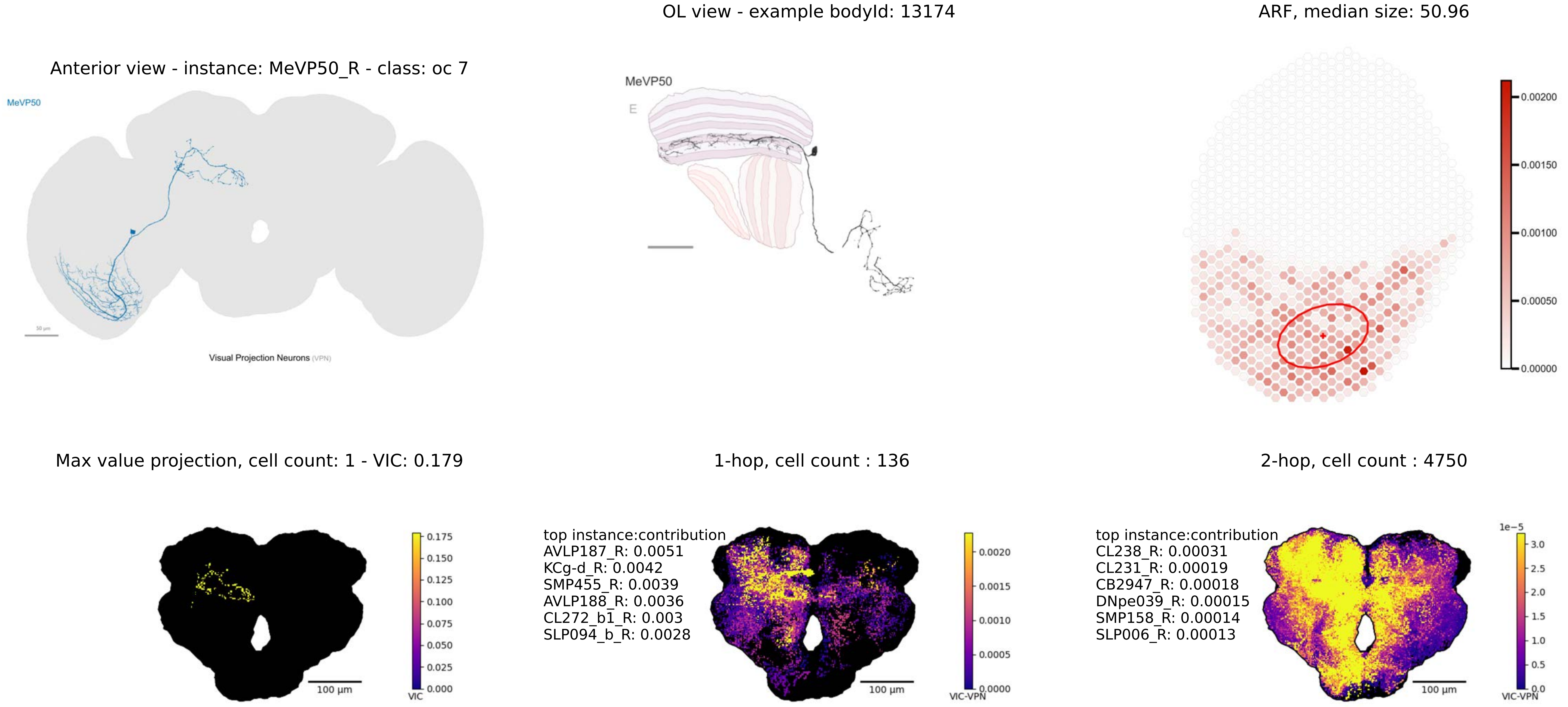

OL view - example bodyId: 13174
ARF, median size: 50.96
Anterior view - instance: MeVP50_R - class: oc 7
MeVP50
Visual Projection Neurons (VPN)
MeVP50
E
0.00200
0.00150
0.00100
0.00050
0.00000
Max value projection, cell count: 1 - VIC: 0.179
1-hop, cell count : 136
2-hop, cell count : 4750
0.175
0.150
0.125
0.100
0.075
0.050
0.025
0.000
VIC
100 µm
top instance:contribution
AVLP187_R: 0.0051
KCg-d_R: 0.0042
SMP455_R: 0.0039
AVLP188_R: 0.0036
CL272_b1_R: 0.003
SLP094_b_R: 0.0028
0.0020
0.0015
0.0010
0.0005
0.0000
VIC-VPN
100 µm
top instance:contribution
CL238_R: 0.00031
CL231_R: 0.00019
CB2947_R: 0.00018
DNpe039_R: 0.00015
SMP158_R: 0.00014
SLP006_R: 0.00013
1e−5
3.0
2.5
2.0
1.5
1.0
0.5
0.0
VIC-VPN
100 µm

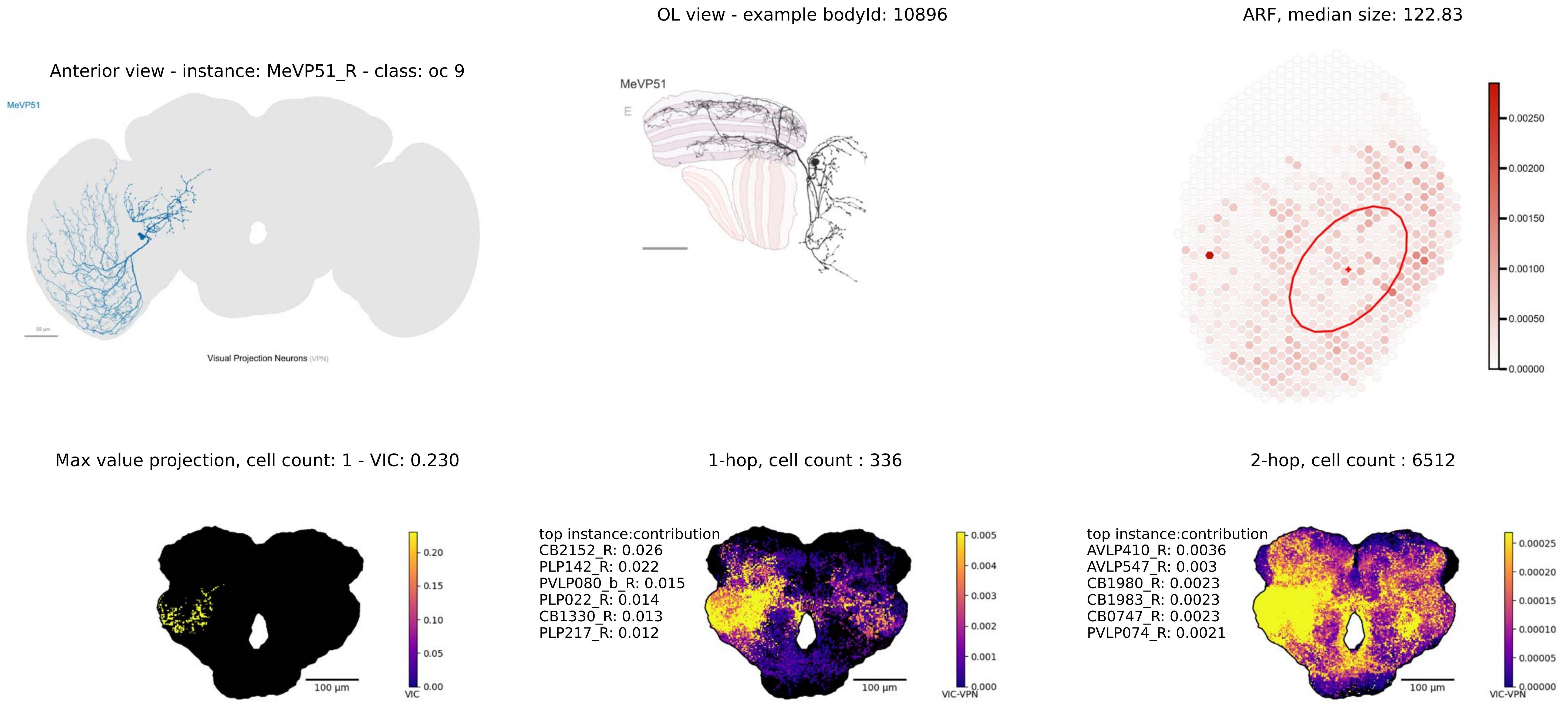
Anterior view - instance: MeVP51_R - class: oc 9
MeVP51
Visual Projection Neurons (VPN)
OL view - example bodyId: 10896
MeVP51
E
ARF, median size: 122.83
0.00250
0.00200
0.00150
0.00100
0.00050
0.00000
Max value projection, cell count: 1 - VIC: 0.230
100 µm
0.20
0.15
0.10
0.05
0.00
VIC
1-hop, cell count : 336
top instance:contribution
CB2152_R: 0.026
PLP142_R: 0.022
PVLP080_b_R: 0.015
PLP022_R: 0.014
CB1330_R: 0.013
PLP217_R: 0.012
100 µm
0.005
0.004
0.003
0.002
0.001
0.000
VIC-VPN
2-hop, cell count : 6512
top instance:contribution
AVLP410_R: 0.0036
AVLP547_R: 0.003
CB1980_R: 0.0023
CB1983_R: 0.0023
CB0747_R: 0.0023
PVLP074_R: 0.0021
100 µm
0.00025
0.00020
0.00015
0.00010
0.00005
0.00000
VIC-VPN

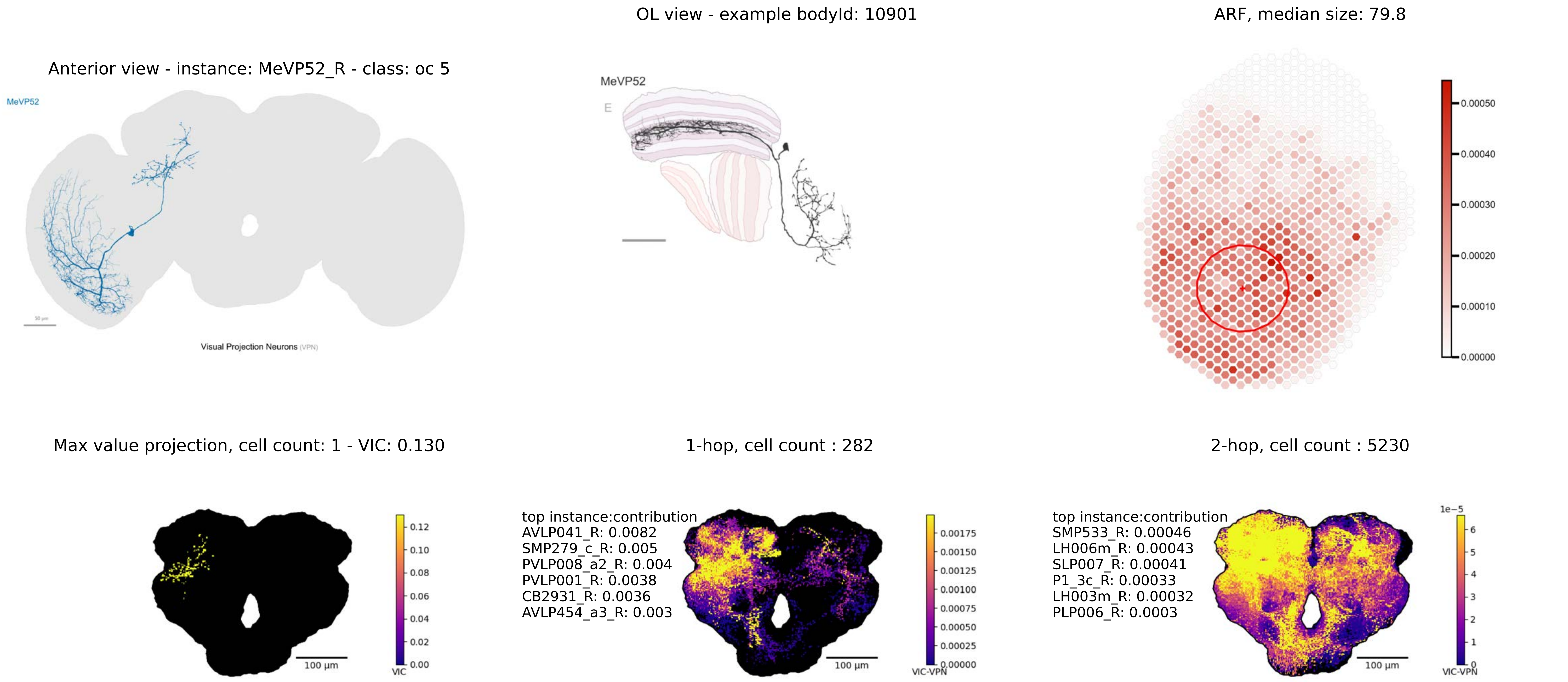

OL view - example bodyId: 10901
ARF, median size: 79.8
Anterior view - instance: MeVP52_R - class: oc 5
MeVP52
Visual Projection Neurons (VPN)
MeVP52
E
0.00050
0.00040
0.00030
0.00020
0.00010
0.00000
Max value projection, cell count: 1 - VIC: 0.130
1-hop, cell count : 282
2-hop, cell count : 5230
0.12
0.10
0.08
0.06
0.04
0.02
0.00
VIC
100 µm
top instance:contribution
AVLP041_R: 0.0082
SMP279_c_R: 0.005
PVLP008_a2_R: 0.004
PVLP001_R: 0.0038
CB2931_R: 0.0036
AVLP454_a3_R: 0.003
0.00175
0.00150
0.00125
0.00100
0.00075
0.00050
0.00025
0.00000
VIC-VPN
100 µm
top instance:contribution
SMP533_R: 0.00046
LH006m_R: 0.00043
SLP007_R: 0.00041
P1_3c_R: 0.00033
LH003m_R: 0.00032
PLP006_R: 0.0003
1e−5
6
5
4
3
2
1
0
VIC-VPN
100 µm

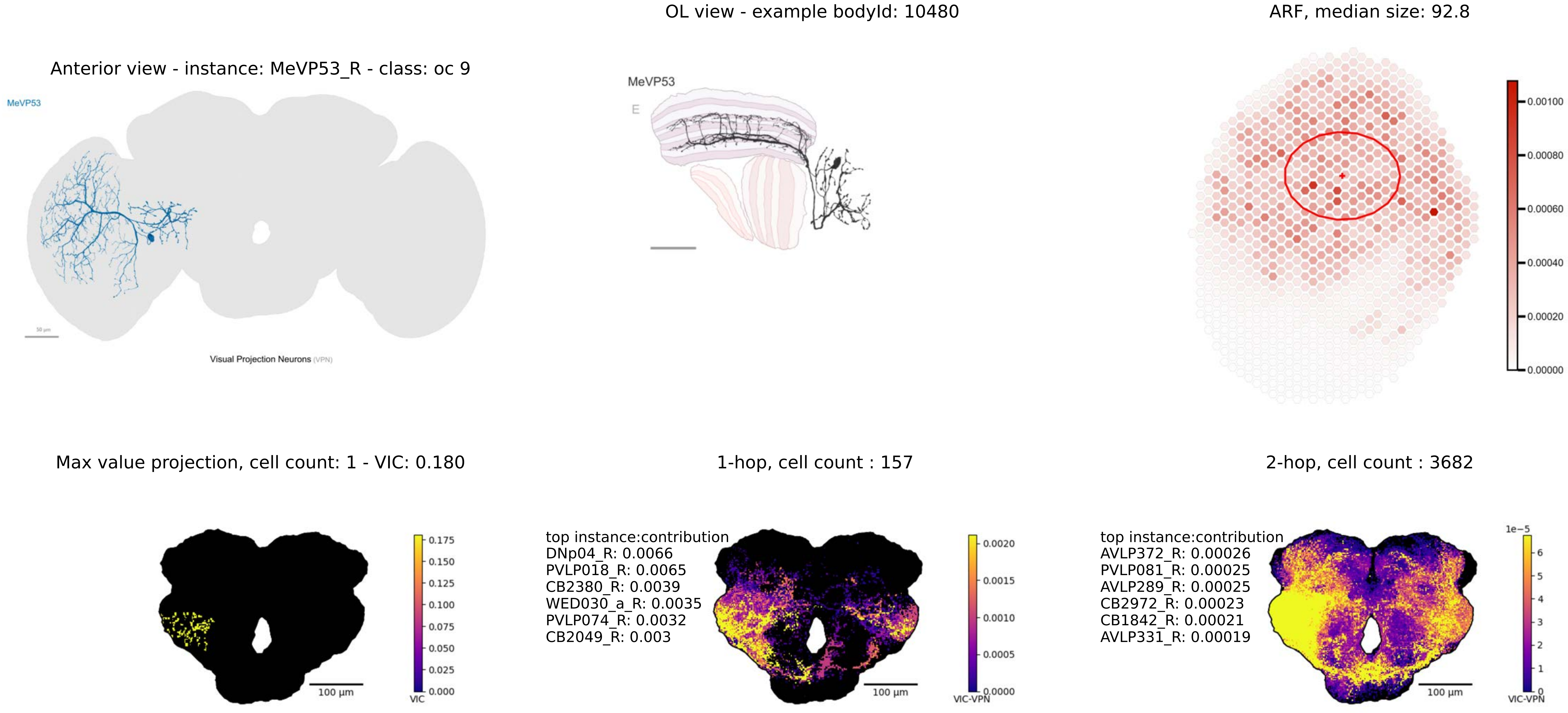

OL view - example bodyId: 10480
ARF, median size: 92.8
Anterior view - instance: MeVP53_R - class: oc 9
MeVP53
MeVP53
E
Visual Projection Neurons (VPN)
0.00100
0.00080
0.00060
0.00040
0.00020
0.00000
Max value projection, cell count: 1 - VIC: 0.180
1-hop, cell count : 157
2-hop, cell count : 3682
0.175
0.150
0.125
0.100
0.075
0.050
0.025
0.000
VIC
100 μm
top instance:contribution
DNp04_R: 0.0066
PVLP018_R: 0.0065
CB2380_R: 0.0039
WED030_a_R: 0.0035
PVLP074_R: 0.0032
CB2049_R: 0.003
0.0020
0.0015
0.0010
0.0005
0.0000
VIC-VPN
100 μm
top instance:contribution
AVLP372_R: 0.00026
PVLP081_R: 0.00025
AVLP289_R: 0.00025
CB2972_R: 0.00023
CB1842_R: 0.00021
AVLP331_R: 0.00019
1e−5
6
5
4
3
2
1
0
VIC-VPN
100 μm

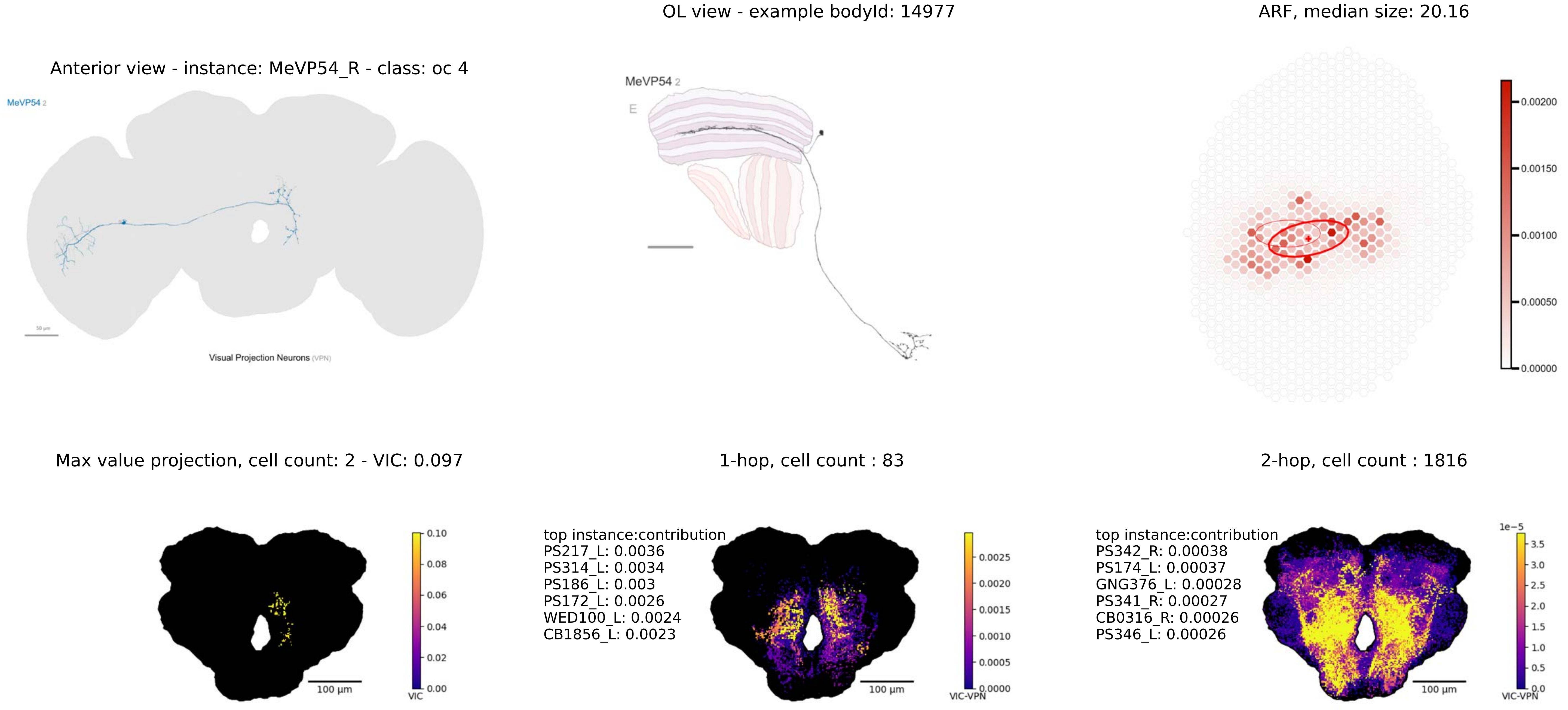

OL view - example bodyId: 14977
ARF, median size: 20.16
Anterior view - instance: MeVP54_R - class: oc 4
MeVP54 2
Visual Projection Neurons (VPN)
MeVP54 2
E
0.00200
0.00150
0.00100
0.00050
0.00000
Max value projection, cell count: 2 - VIC: 0.097
1-hop, cell count : 83
2-hop, cell count : 1816
0.10
0.08
0.06
0.04
0.02
0.00
VIC
100 µm
top instance:contribution
PS217_L: 0.0036
PS314_L: 0.0034
PS186_L: 0.003
PS172_L: 0.0026
WED100_L: 0.0024
CB1856_L: 0.0023
0.0025
0.0020
0.0015
0.0010
0.0005
0.0000
VIC-VPN
100 µm
top instance:contribution
PS342_R: 0.00038
PS174_L: 0.00037
GNG376_L: 0.00028
PS341_R: 0.00027
CB0316_R: 0.00026
PS346_L: 0.00026
1e−5
3.5
3.0
2.5
2.0
1.5
1.0
0.5
0.0
VIC-VPN
100 µm

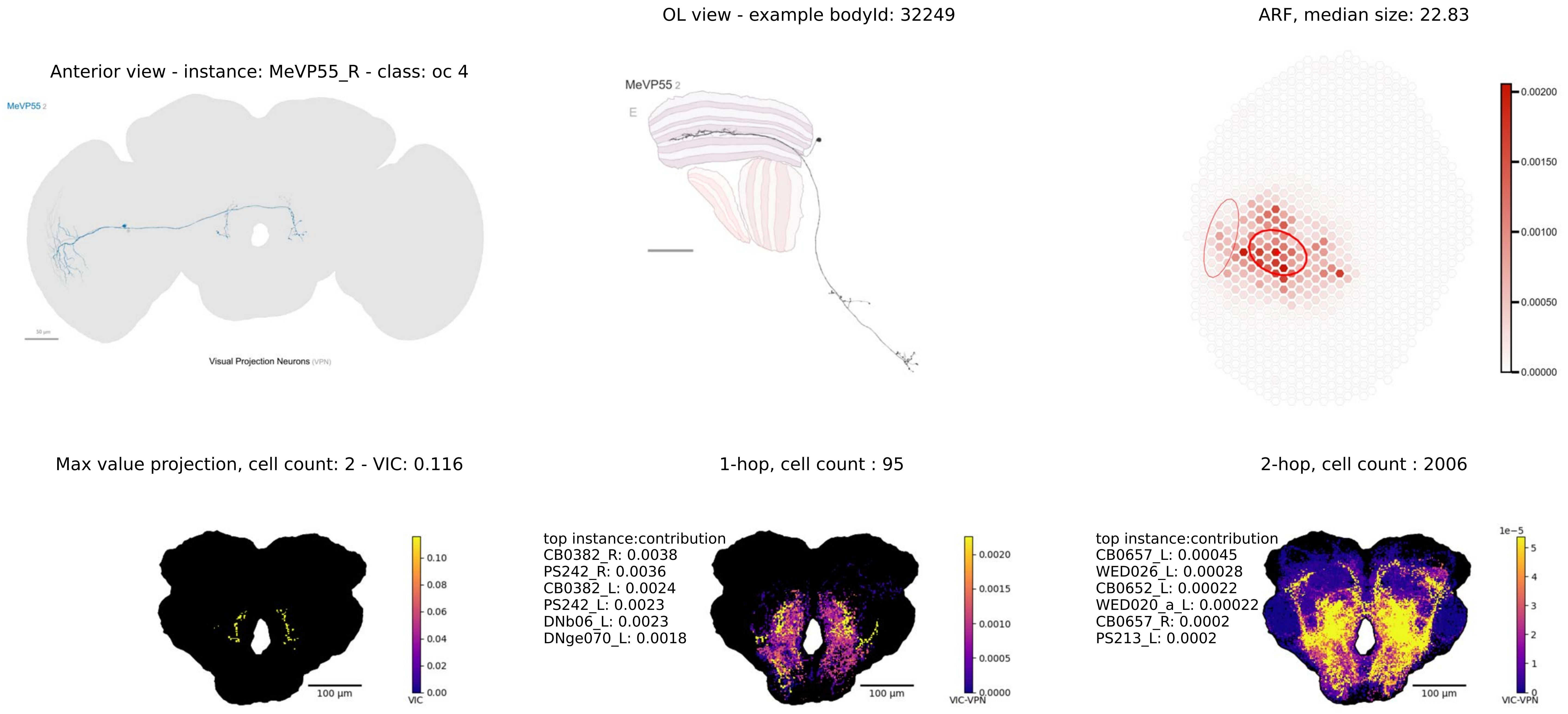
Anterior view - instance: MeVP55_R - class: oc 4
MeVP55 2
Visual Projection Neurons (VPN)
OL view - example bodyId: 32249
MeVP55 2
E
ARF, median size: 22.83
0.00200
0.00150
0.00100
0.00050
0.00000
Max value projection, cell count: 2 - VIC: 0.116
100 μm
0.10
0.08
0.06
0.04
0.02
0.00
VIC
1-hop, cell count : 95
top instance:contribution
CB0382_R: 0.0038
PS242_R: 0.0036
CB0382_L: 0.0024
PS242_L: 0.0023
DNb06_L: 0.0023
DNge070_L: 0.0018
100 μm
0.0020
0.0015
0.0010
0.0005
0.0000
VIC-VPN
2-hop, cell count : 2006
top instance:contribution
CB0657_L: 0.00045
WED026_L: 0.00028
CB0652_L: 0.00022
WED020_a_L: 0.00022
CB0657_R: 0.0002
PS213_L: 0.0002
100 μm
1e−5
5
4
3
2
1
0
VIC-VPN

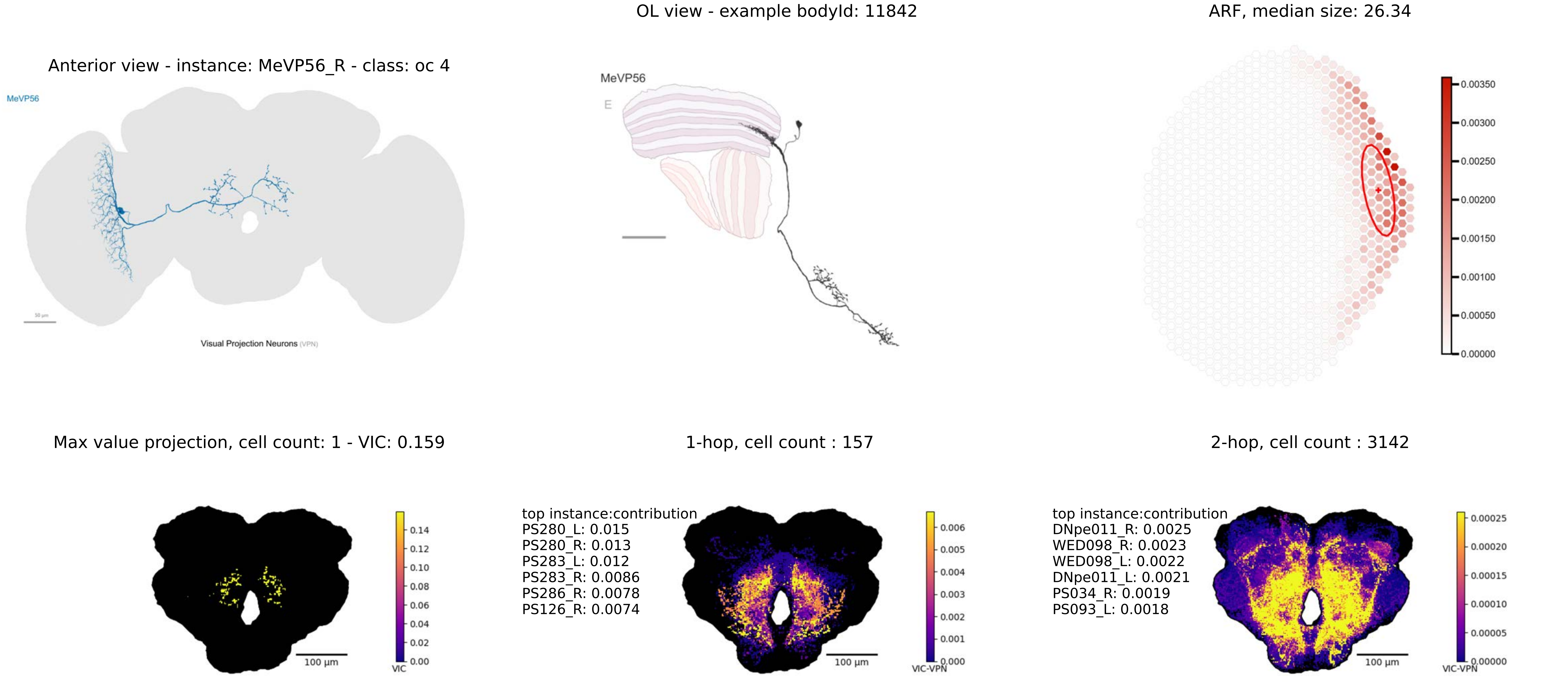

OL view - example bodyId: 11842
ARF, median size: 26.34
Anterior view - instance: MeVP56_R - class: oc 4
MeVP56
Visual Projection Neurons (VPN)
MeVP56
E
0.00350
0.00300
0.00250
0.00200
0.00150
0.00100
0.00050
0.00000
Max value projection, cell count: 1 - VIC: 0.159
1-hop, cell count : 157
2-hop, cell count : 3142
0.14
0.12
0.10
0.08
0.06
0.04
0.02
0.00
VIC
100 µm
top instance:contribution
PS280_L: 0.015
PS280_R: 0.013
PS283_L: 0.012
PS283_R: 0.0086
PS286_R: 0.0078
PS126_R: 0.0074
0.006
0.005
0.004
0.003
0.002
0.001
0.000
VIC-VPN
100 µm
top instance:contribution
DNpe011_R: 0.0025
WED098_R: 0.0023
WED098_L: 0.0022
DNpe011_L: 0.0021
PS034_R: 0.0019
PS093_L: 0.0018
0.00025
0.00020
0.00015
0.00010
0.00005
0.00000
VIC-VPN
100 µm

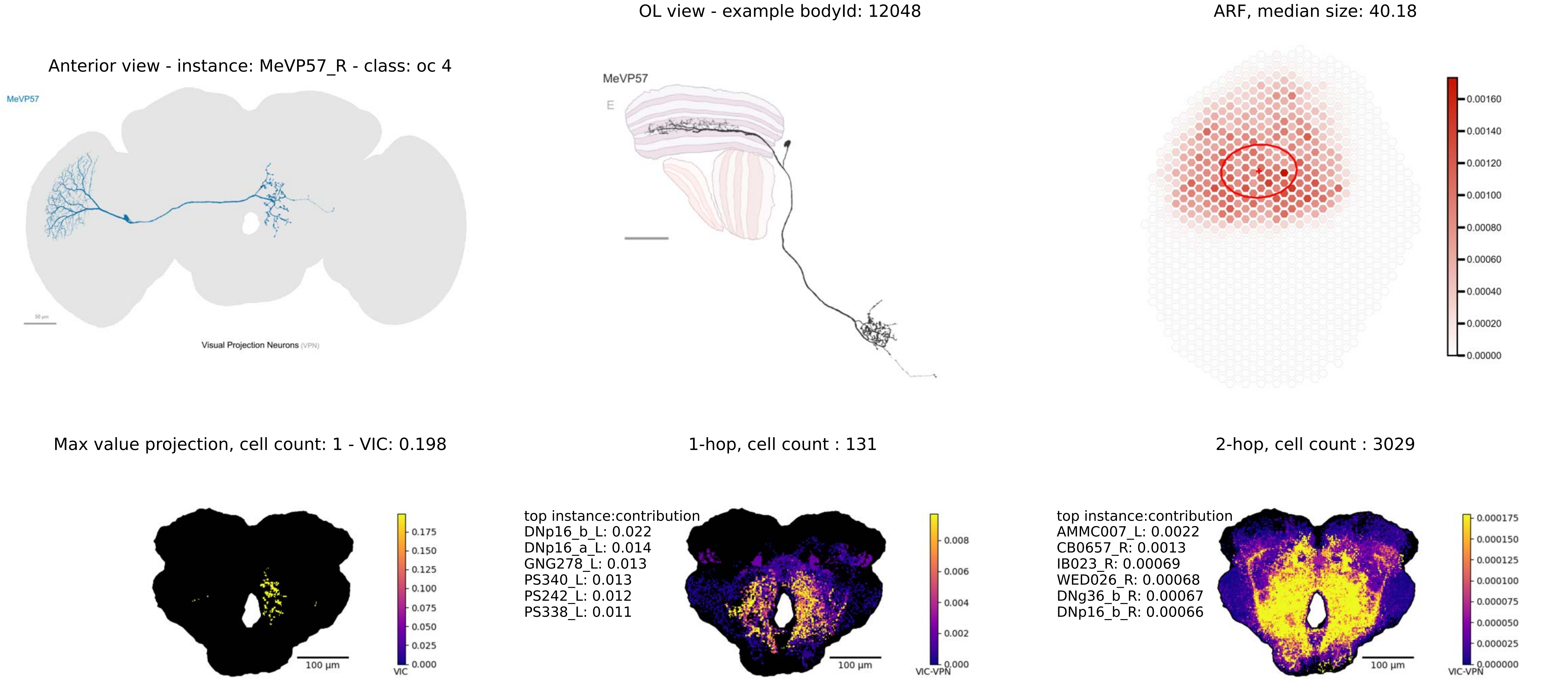
Anterior view - instance: MeVP57_R - class: oc 4
MeVP57
Visual Projection Neurons (VPN)
OL view - example bodyId: 12048
MeVP57
E
ARF, median size: 40.18
0.00160
0.00140
0.00120
0.00100
0.00080
0.00060
0.00040
0.00020
0.00000
Max value projection, cell count: 1 - VIC: 0.198
100 µm
0.175
0.150
0.125
0.100
0.075
0.050
0.025
0.000
VIC
1-hop, cell count : 131
top instance:contribution
DNp16_b_L: 0.022
DNp16_a_L: 0.014
GNG278_L: 0.013
PS340_L: 0.013
PS242_L: 0.012
PS338_L: 0.011
100 µm
0.008
0.006
0.004
0.002
0.000
VIC-VPN
2-hop, cell count : 3029
top instance:contribution
AMMC007_L: 0.0022
CB0657_R: 0.0013
IB023_R: 0.00069
WED026_R: 0.00068
DNg36_b_R: 0.00067
DNp16_b_R: 0.00066
100 µm
0.000175
0.000150
0.000125
0.000100
0.000075
0.000050
0.000025
0.000000
VIC-VPN

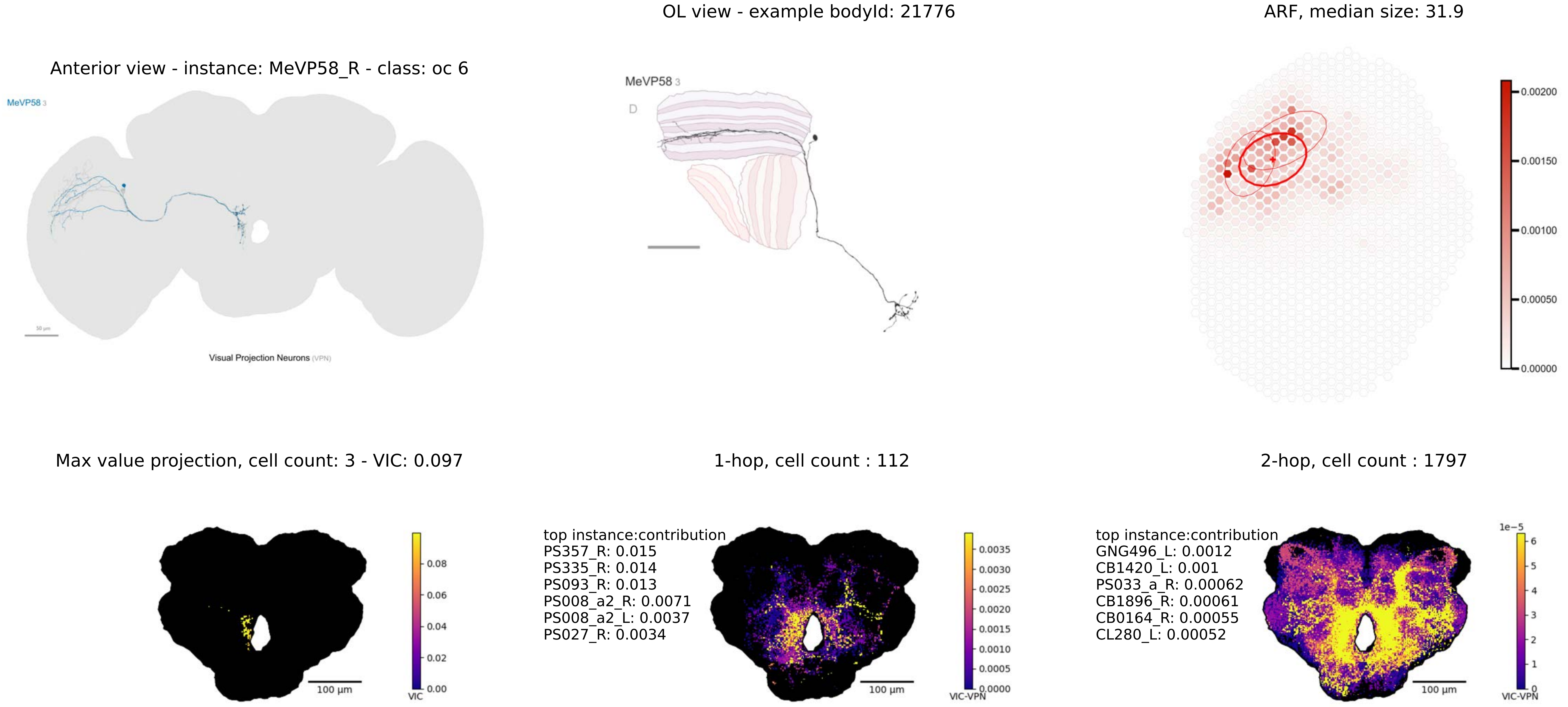

Anterior view - instance: MeVP58_R - class: oc 6
MeVP58 3
Visual Projection Neurons (VPN)
OL view - example bodyId: 21776
MeVP58 3
D
ARF, median size: 31.9
0.00200
0.00150
0.00100
0.00050
0.00000
Max value projection, cell count: 3 - VIC: 0.097
100 µm
0.08
0.06
0.04
0.02
0.00
VIC
1-hop, cell count : 112
top instance:contribution
PS357_R: 0.015
PS335_R: 0.014
PS093_R: 0.013
PS008_a2_R: 0.0071
PS008_a2_L: 0.0037
PS027_R: 0.0034
100 µm
0.0035
0.0030
0.0025
0.0020
0.0015
0.0010
0.0005
0.0000
VIC-VPN
2-hop, cell count : 1797
top instance:contribution
GNG496_L: 0.0012
CB1420_L: 0.001
PS033_a_R: 0.00062
CB1896_R: 0.00061
CB0164_R: 0.00055
CL280_L: 0.00052
100 µm
1e−5
6
5
4
3
2
1
0
VIC-VPN

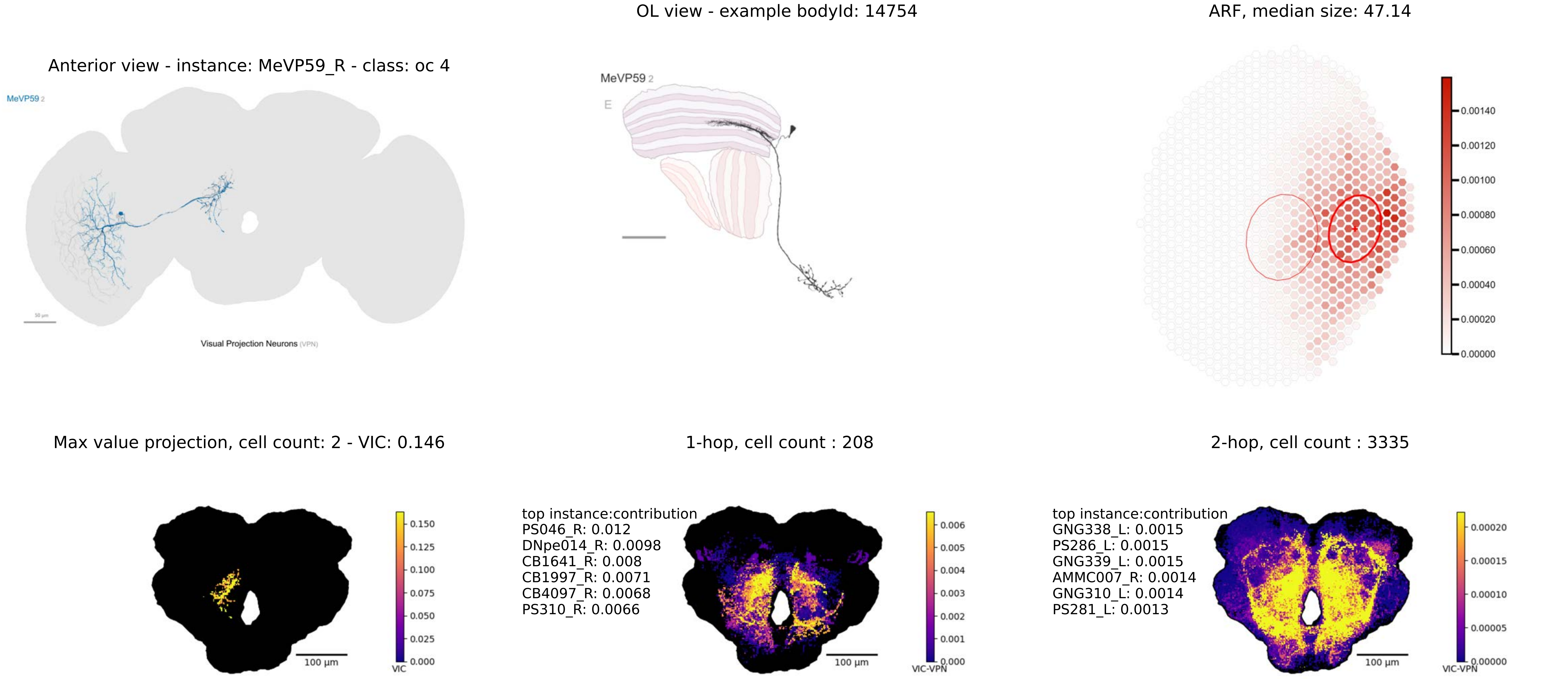

Anterior view - instance: MeVP59_R - class: oc 4
MeVP59 2
Visual Projection Neurons (VPN)
OL view - example bodyId: 14754
MeVP59 2
E
ARF, median size: 47.14
0.00140
0.00120
0.00100
0.00080
0.00060
0.00040
0.00020
0.00000
Max value projection, cell count: 2 - VIC: 0.146
0.150
0.125
0.100
0.075
0.050
0.025
0.000
VIC
100 µm
1-hop, cell count : 208
top instance:contribution
PS046_R: 0.012
DNpe014_R: 0.0098
CB1641_R: 0.008
CB1997_R: 0.0071
CB4097_R: 0.0068
PS310_R: 0.0066
0.006
0.005
0.004
0.003
0.002
0.001
0.000
VIC-VPN
100 µm
2-hop, cell count : 3335
top instance:contribution
GNG338_L: 0.0015
PS286_L: 0.0015
GNG339_L: 0.0015
AMMC007_R: 0.0014
GNG310_L: 0.0014
PS281_L: 0.0013
0.00020
0.00015
0.00010
0.00005
0.00000
VIC-VPN
100 µm

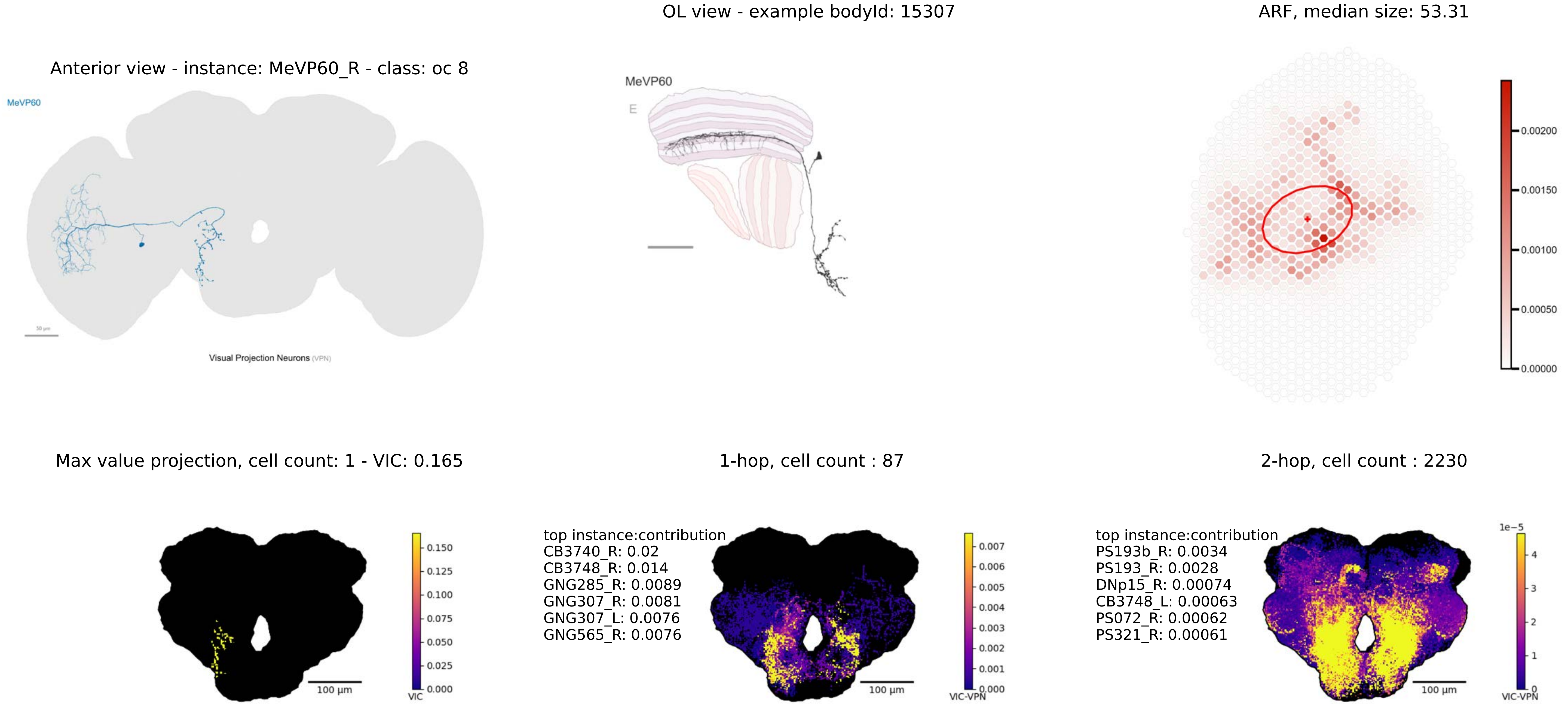
OL view - example bodyId: 15307
ARF, median size: 53.31
Anterior view - instance: MeVP60_R - class: oc 8
MeVP60
Visual Projection Neurons (VPN)
MeVP60
E
0.00200
0.00150
0.00100
0.00050
0.00000
Max value projection, cell count: 1 - VIC: 0.165
1-hop, cell count : 87
2-hop, cell count : 2230
100 µm
VIC
0.150
0.125
0.100
0.075
0.050
0.025
0.000
top instance:contribution
CB3740_R: 0.02
CB3748_R: 0.014
GNG285_R: 0.0089
GNG307_R: 0.0081
GNG307_L: 0.0076
GNG565_R: 0.0076
VIC-VPN
0.007
0.006
0.005
0.004
0.003
0.002
0.001
0.000
top instance:contribution
PS193b_R: 0.0034
PS193_R: 0.0028
DNp15_R: 0.00074
CB3748_L: 0.00063
PS072_R: 0.00062
PS321_R: 0.00061
1e−5
4
3
2
1
0
VIC-VPN

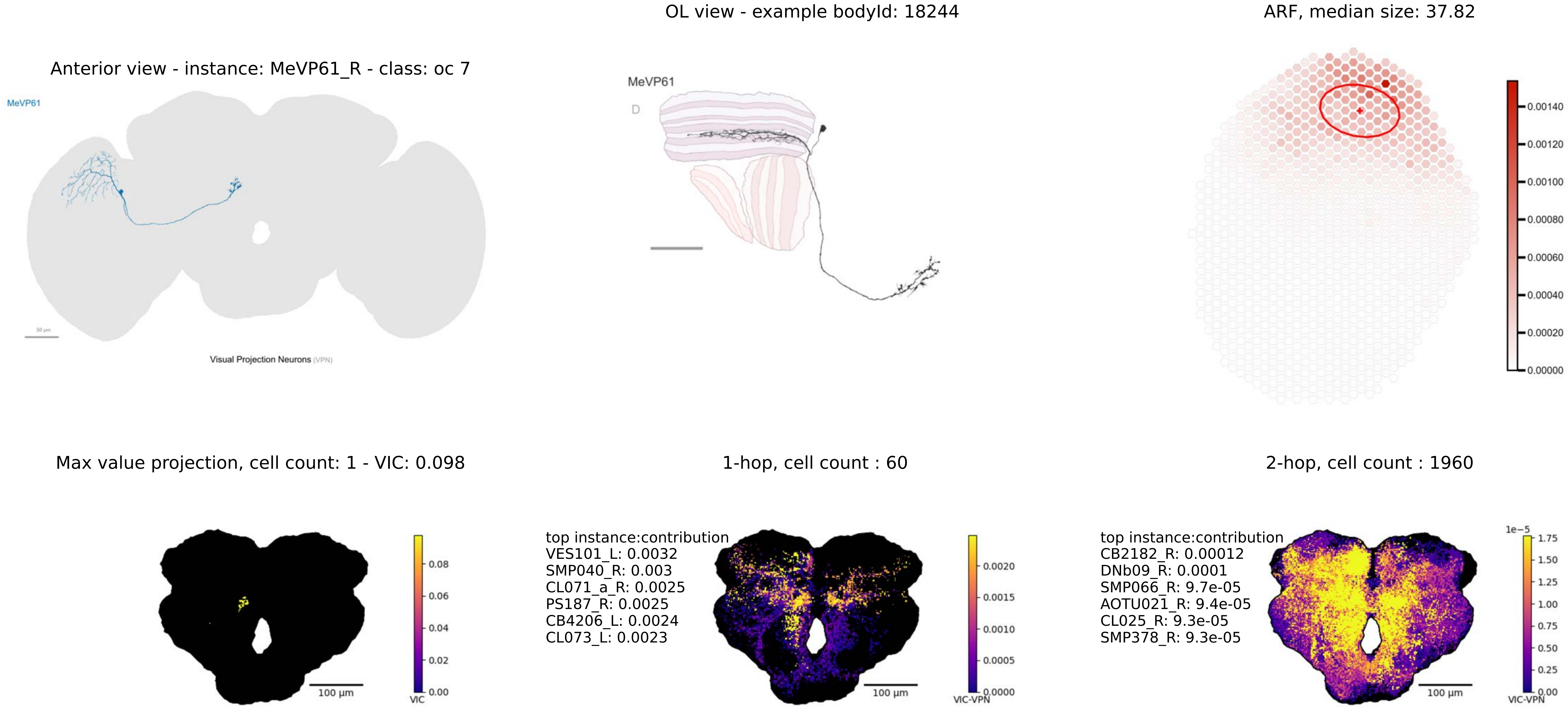

Anterior view - instance: MeVP61_R - class: oc 7
MeVP61
Visual Projection Neurons (VPN)
OL view - example bodyId: 18244
MeVP61
D
ARF, median size: 37.82
0.00140
0.00120
0.00100
0.00080
0.00060
0.00040
0.00020
0.00000
Max value projection, cell count: 1 - VIC: 0.098
0.08
0.06
0.04
0.02
0.00
VIC
100 µm
1-hop, cell count : 60
top instance:contribution
VES101_L: 0.0032
SMP040_R: 0.003
CL071_a_R: 0.0025
PS187_R: 0.0025
CB4206_L: 0.0024
CL073_L: 0.0023
0.0020
0.0015
0.0010
0.0005
0.0000
VIC-VPN
100 µm
2-hop, cell count : 1960
top instance:contribution
CB2182_R: 0.00012
DNb09_R: 0.0001
SMP066_R: 9.7e-05
AOTU021_R: 9.4e-05
CL025_R: 9.3e-05
SMP378_R: 9.3e-05
1e−5
1.75
1.50
1.25
1.00
0.75
0.50
0.25
0.00
VIC-VPN
100 µm

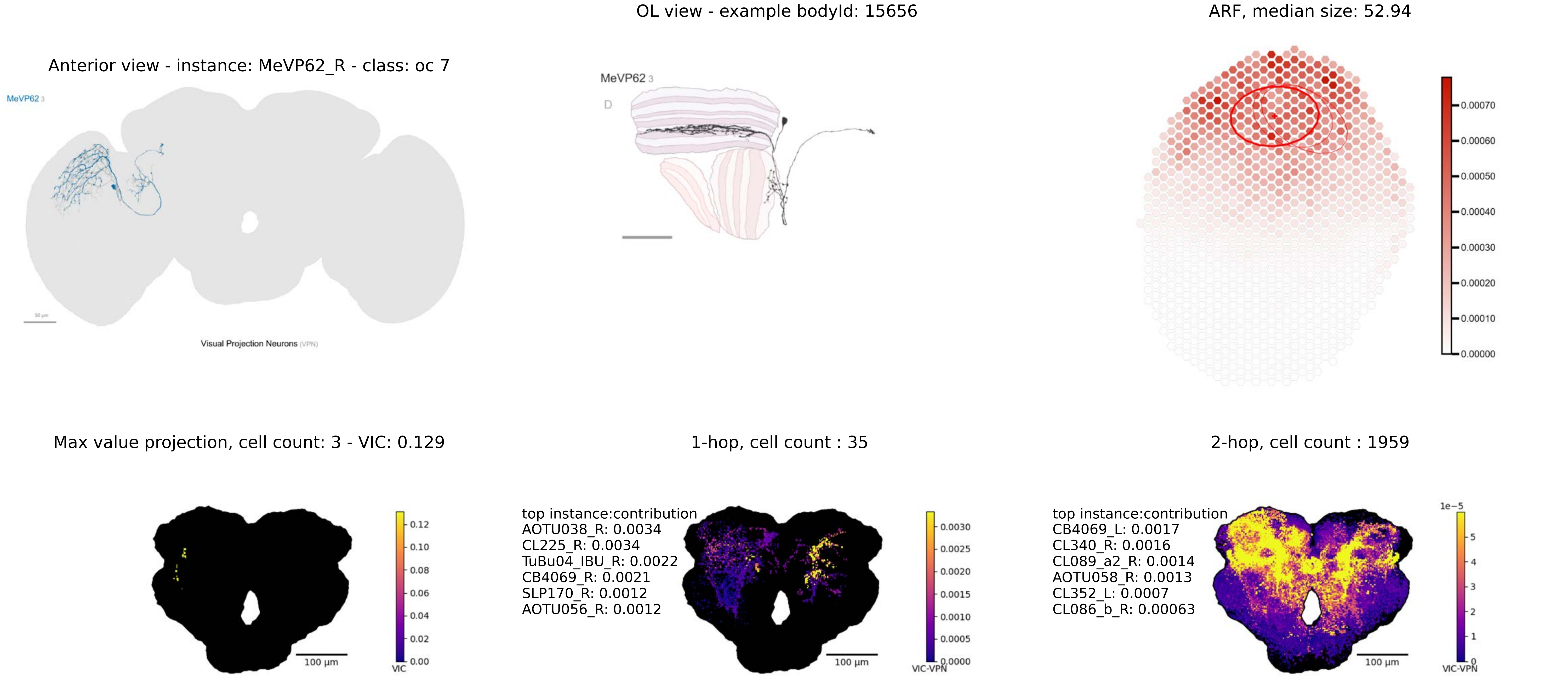

Anterior view - instance: MeVP62_R - class: oc 7
MeVP62 3
50 µm
Visual Projection Neurons (VPN)
OL view - example bodyId: 15656
MeVP62 3
D
ARF, median size: 52.94
0.00070
0.00060
0.00050
0.00040
0.00030
0.00020
0.00010
0.00000
Max value projection, cell count: 3 - VIC: 0.129
100 µm
0.12
0.10
0.08
0.06
0.04
0.02
0.00
VIC
1-hop, cell count : 35
top instance:contribution
AOTU038_R: 0.0034
CL225_R: 0.0034
TuBu04_IBU_R: 0.0022
CB4069_R: 0.0021
SLP170_R: 0.0012
AOTU056_R: 0.0012
100 µm
0.0030
0.0025
0.0020
0.0015
0.0010
0.0005
0.0000
VIC-VPN
2-hop, cell count : 1959
top instance:contribution
CB4069_L: 0.0017
CL340_R: 0.0016
CL089_a2_R: 0.0014
AOTU058_R: 0.0013
CL352_L: 0.0007
CL086_b_R: 0.00063
100 µm
1e−5
5
4
3
2
1
0
VIC-VPN

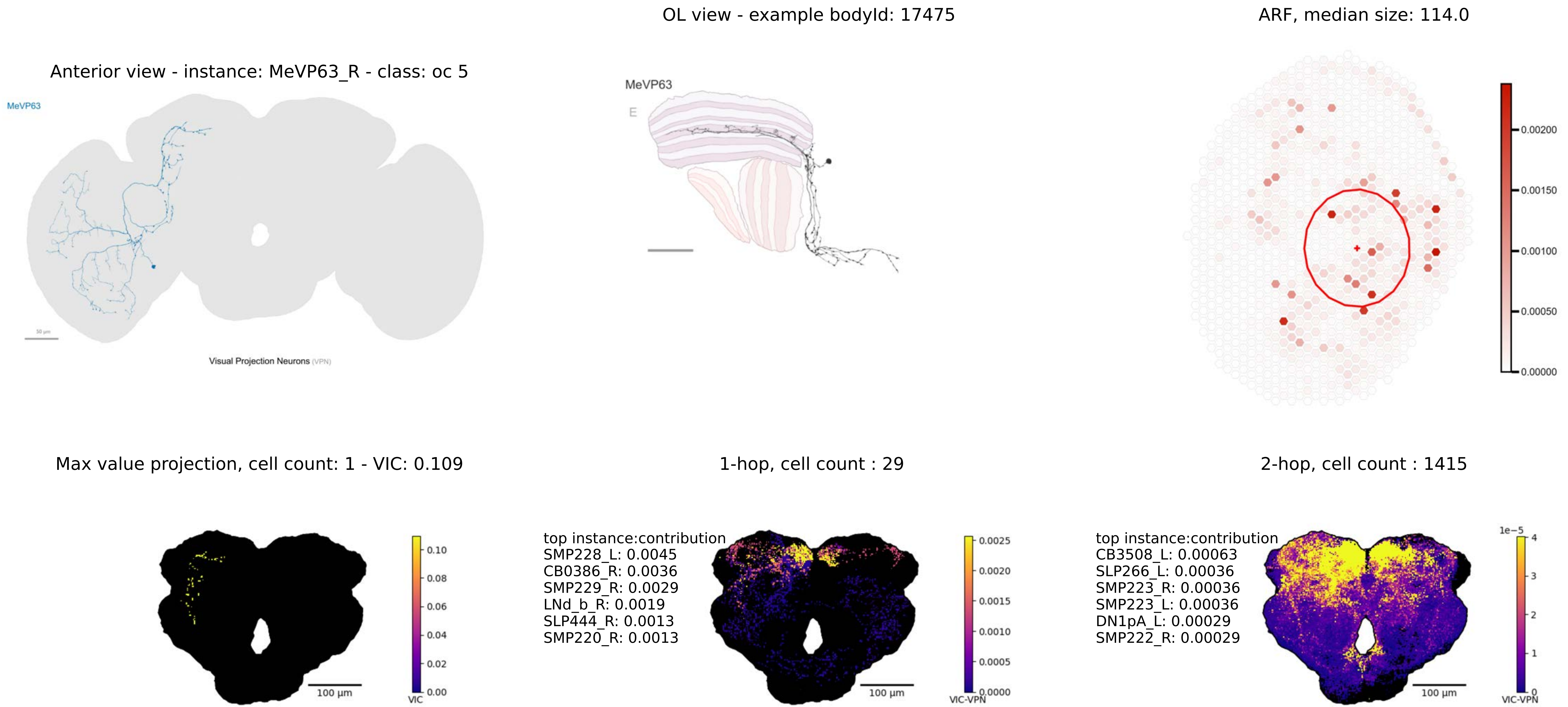

OL view - example bodyId: 17475
ARF, median size: 114.0
Anterior view - instance: MeVP63_R - class: oc 5
MeVP63
Visual Projection Neurons (VPN)
MeVP63
E
0.00200
0.00150
0.00100
0.00050
0.00000
Max value projection, cell count: 1 - VIC: 0.109
1-hop, cell count : 29
2-hop, cell count : 1415
0.10
0.08
0.06
0.04
0.02
0.00
VIC
100 µm
top instance:contribution
SMP228_L: 0.0045
CB0386_R: 0.0036
SMP229_R: 0.0029
LNd_b_R: 0.0019
SLP444_R: 0.0013
SMP220_R: 0.0013
0.0025
0.0020
0.0015
0.0010
0.0005
0.0000
VIC-VPN
100 µm
top instance:contribution
CB3508_L: 0.00063
SLP266_L: 0.00036
SMP223_R: 0.00036
SMP223_L: 0.00036
DN1pA_L: 0.00029
SMP222_R: 0.00029
1e−5
4
3
2
1
0
VIC-VPN
100 µm

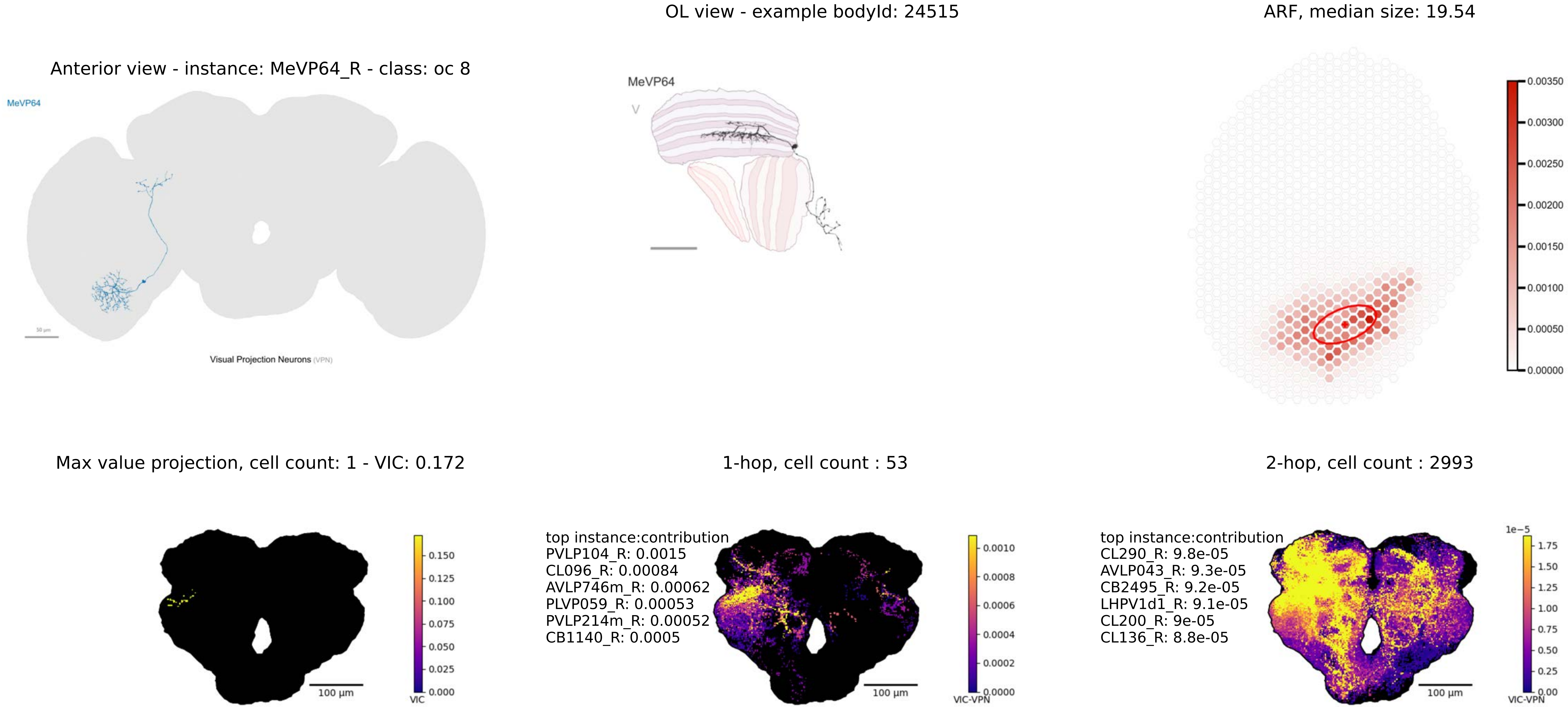

OL view - example bodyId: 24515
ARF, median size: 19.54
Anterior view - instance: MeVP64_R - class: oc 8
MeVP64
50 μm
Visual Projection Neurons (VPN)
MeVP64
V
0.00350
0.00300
0.00250
0.00200
0.00150
0.00100
0.00050
0.00000
Max value projection, cell count: 1 - VIC: 0.172
1-hop, cell count : 53
2-hop, cell count : 2993
0.150
0.125
0.100
0.075
0.050
0.025
0.000
VIC
100 μm
top instance:contribution
PVLP104_R: 0.0015
CL096_R: 0.00084
AVLP746m_R: 0.00062
PLVP059_R: 0.00053
PVLP214m_R: 0.00052
CB1140_R: 0.0005
0.0010
0.0008
0.0006
0.0004
0.0002
0.0000
VIC-VPN
100 μm
top instance:contribution
CL290_R: 9.8e-05
AVLP043_R: 9.3e-05
CB2495_R: 9.2e-05
LHPV1d1_R: 9.1e-05
CL200_R: 9e-05
CL136_R: 8.8e-05
1e−5
1.75
1.50
1.25
1.00
0.75
0.50
0.25
0.00
VIC-VPN
100 μm

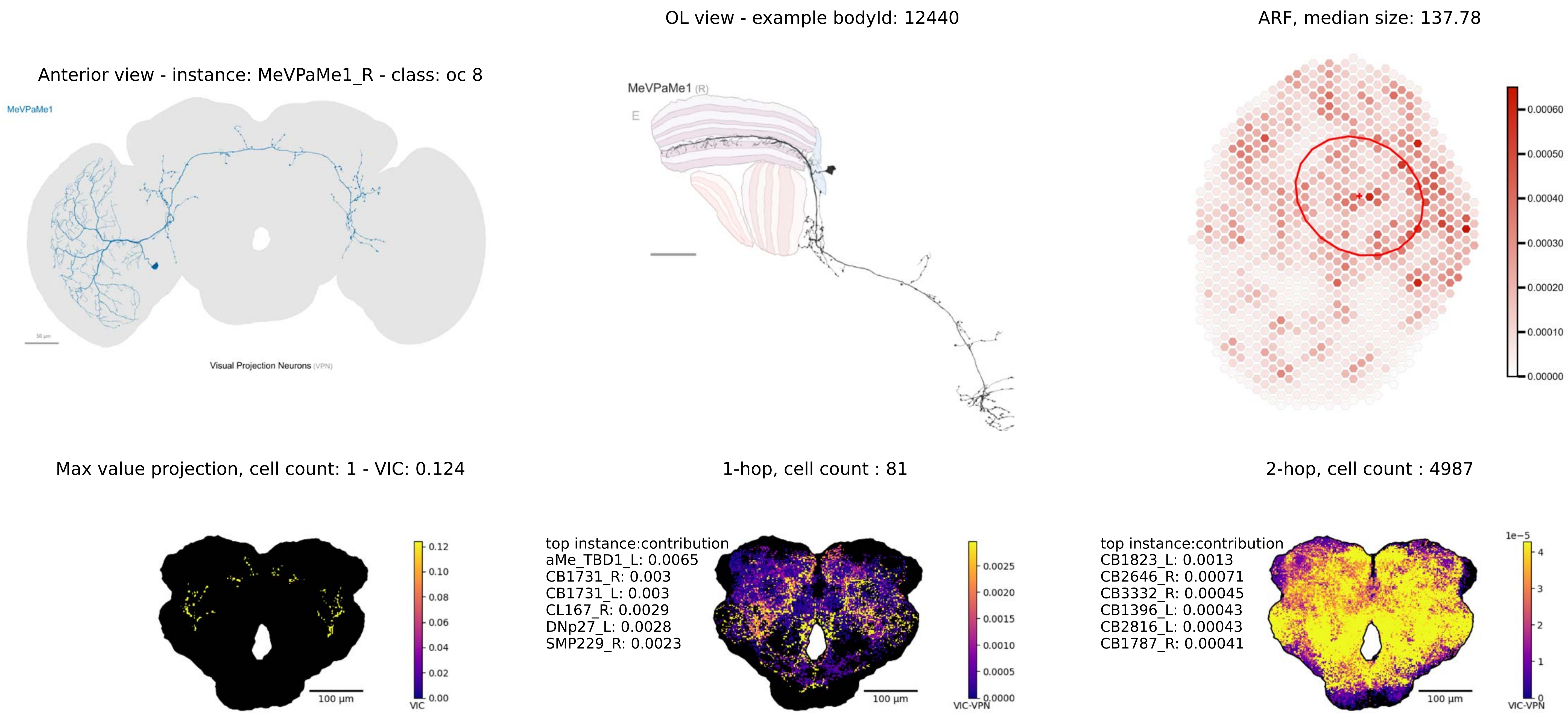
OL view - example bodyId: 12440
ARF, median size: 137.78
Anterior view - instance: MeVPaMe1_R - class: oc 8
MeVPaMe1
Visual Projection Neurons (VPN)
MeVPaMe1 (R)
E
0.00060
0.00050
0.00040
0.00030
0.00020
0.00010
0.00000
Max value projection, cell count: 1 - VIC: 0.124
1-hop, cell count : 81
2-hop, cell count : 4987
0.12
0.10
0.08
0.06
0.04
0.02
0.00
VIC
100 µm
top instance:contribution
aMe_TBD1_L: 0.0065
CB1731_R: 0.003
CB1731_L: 0.003
CL167_R: 0.0029
DNp27_L: 0.0028
SMP229_R: 0.0023
0.0025
0.0020
0.0015
0.0010
0.0005
0.0000
VIC-VPN
100 µm
top instance:contribution
CB1823_L: 0.0013
CB2646_R: 0.00071
CB3332_R: 0.00045
CB1396_L: 0.00043
CB2816_L: 0.00043
CB1787_R: 0.00041
1e−5
4
3
2
1
0
VIC-VPN
100 µm

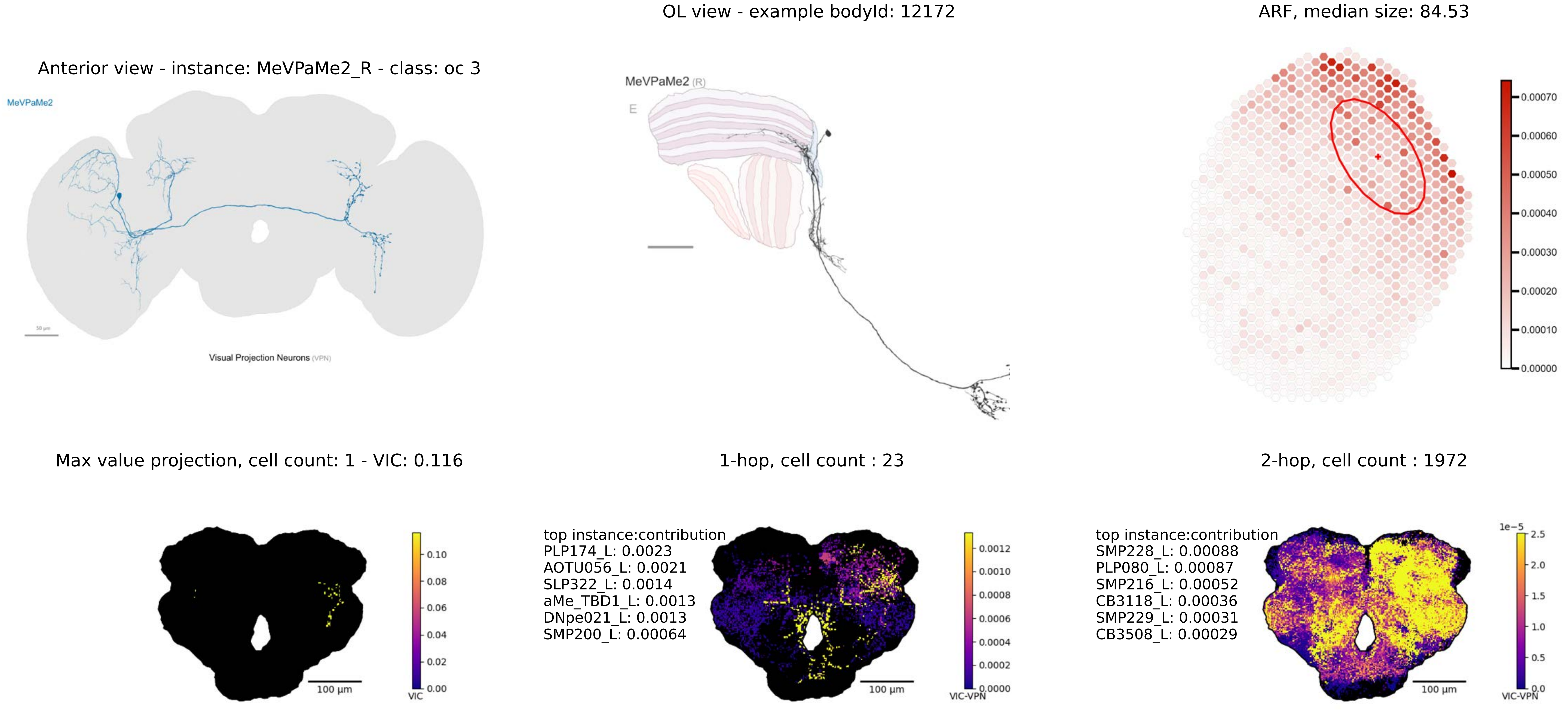
Anterior view - instance: MeVPaMe2_R - class: oc 3
MeVPaMe2
Visual Projection Neurons (VPN)
OL view - example bodyId: 12172
MeVPaMe2 (R)
E
ARF, median size: 84.53
0.00070
0.00060
0.00050
0.00040
0.00030
0.00020
0.00010
0.00000
Max value projection, cell count: 1 - VIC: 0.116
100 µm
0.10
0.08
0.06
0.04
0.02
0.00
VIC
1-hop, cell count : 23
top instance:contribution
PLP174_L: 0.0023
AOTU056_L: 0.0021
SLP322_L: 0.0014
aMe_TBD1_L: 0.0013
DNpe021_L: 0.0013
SMP200_L: 0.00064
100 µm
0.0012
0.0010
0.0008
0.0006
0.0004
0.0002
0.0000
VIC-VPN
2-hop, cell count : 1972
top instance:contribution
SMP228_L: 0.00088
PLP080_L: 0.00087
SMP216_L: 0.00052
CB3118_L: 0.00036
SMP229_L: 0.00031
CB3508_L: 0.00029
100 µm
1e−5
2.5
2.0
1.5
1.0
0.5
0.0
VIC-VPN

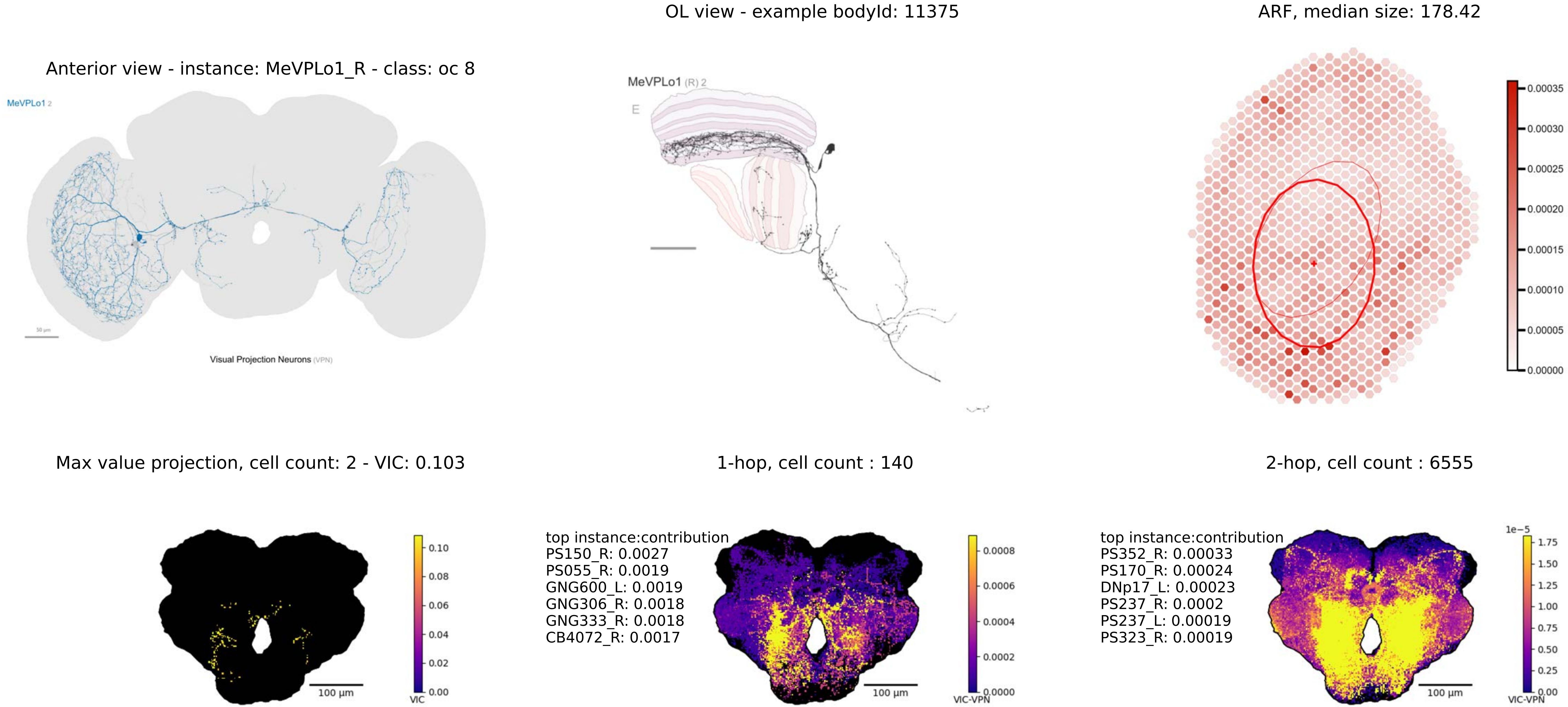

OL view - example bodyId: 11375
ARF, median size: 178.42
Anterior view - instance: MeVPLo1_R - class: oc 8
MeVPLo1 2
Visual Projection Neurons (VPN)
MeVPLo1 (R) 2
E
0.00035
0.00030
0.00025
0.00020
0.00015
0.00010
0.00005
0.00000
Max value projection, cell count: 2 - VIC: 0.103
1-hop, cell count : 140
2-hop, cell count : 6555
0.10
0.08
0.06
0.04
0.02
0.00
VIC
100 µm
top instance:contribution
PS150_R: 0.0027
PS055_R: 0.0019
GNG600_L: 0.0019
GNG306_R: 0.0018
GNG333_R: 0.0018
CB4072_R: 0.0017
0.0008
0.0006
0.0004
0.0002
0.0000
VIC-VPN
100 µm
top instance:contribution
PS352_R: 0.00033
PS170_R: 0.00024
DNp17_L: 0.00023
PS237_R: 0.0002
PS237_L: 0.00019
PS323_R: 0.00019
1e−5
1.75
1.50
1.25
1.00
0.75
0.50
0.25
0.00
VIC-VPN
100 µm

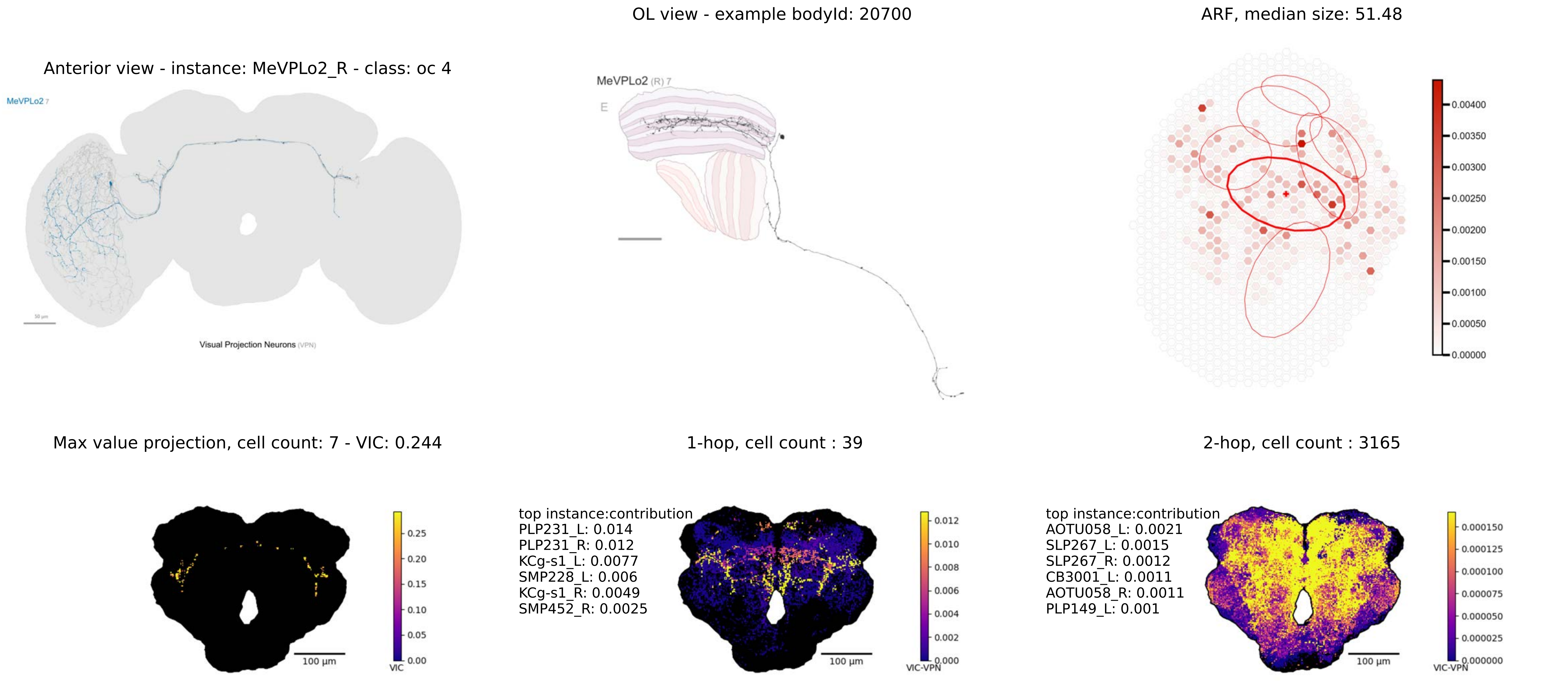

Anterior view - instance: MeVPLo2_R - class: oc 4
MeVPLo2 7
Visual Projection Neurons (VPN)
OL view - example bodyId: 20700
MeVPLo2 (R) 7
E
ARF, median size: 51.48
0.00400
0.00350
0.00300
0.00250
0.00200
0.00150
0.00100
0.00050
0.00000
Max value projection, cell count: 7 - VIC: 0.244
0.25
0.20
0.15
0.10
0.05
0.00
VIC
100 µm
1-hop, cell count : 39
top instance:contribution
PLP231_L: 0.014
PLP231_R: 0.012
KCg-s1_L: 0.0077
SMP228_L: 0.006
KCg-s1_R: 0.0049
SMP452_R: 0.0025
0.012
0.010
0.008
0.006
0.004
0.002
0.000
VIC-VPN
100 µm
2-hop, cell count : 3165
top instance:contribution
AOTU058_L: 0.0021
SLP267_L: 0.0015
SLP267_R: 0.0012
CB3001_L: 0.0011
AOTU058_R: 0.0011
PLP149_L: 0.001
0.000150
0.000125
0.000100
0.000075
0.000050
0.000025
0.000000
VIC-VPN
100 µm

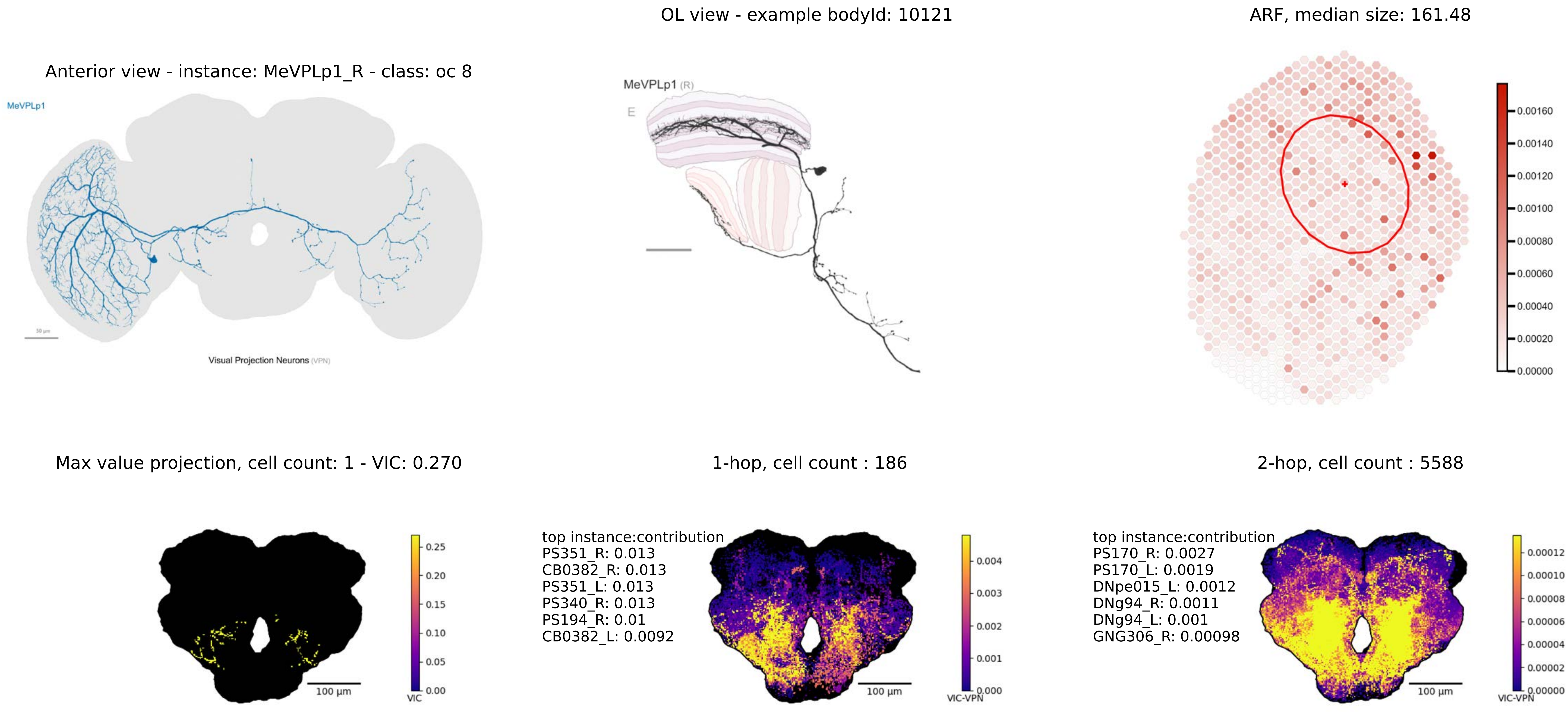

OL view - example bodyId: 10121
ARF, median size: 161.48
Anterior view - instance: MeVPLp1_R - class: oc 8
MeVPLp1
Visual Projection Neurons (VPN)
MeVPLp1 (R)
E
0.00160
0.00140
0.00120
0.00100
0.00080
0.00060
0.00040
0.00020
0.00000
Max value projection, cell count: 1 - VIC: 0.270
1-hop, cell count : 186
2-hop, cell count : 5588
100 µm
VIC
0.25
0.20
0.15
0.10
0.05
0.00
top instance:contribution
PS351_R: 0.013
CB0382_R: 0.013
PS351_L: 0.013
PS340_R: 0.013
PS194_R: 0.01
CB0382_L: 0.0092
VIC-VPN
0.004
0.003
0.002
0.001
0.000
top instance:contribution
PS170_R: 0.0027
PS170_L: 0.0019
DNpe015_L: 0.0012
DNg94_R: 0.0011
DNg94_L: 0.001
GNG306_R: 0.00098
0.00012
0.00010
0.00008
0.00006
0.00004
0.00002
0.00000

OL view - example bodyId: 11684

ARF, median size: 39.08

Anterior view - instance: MeVPLp2_R - class: oc 8

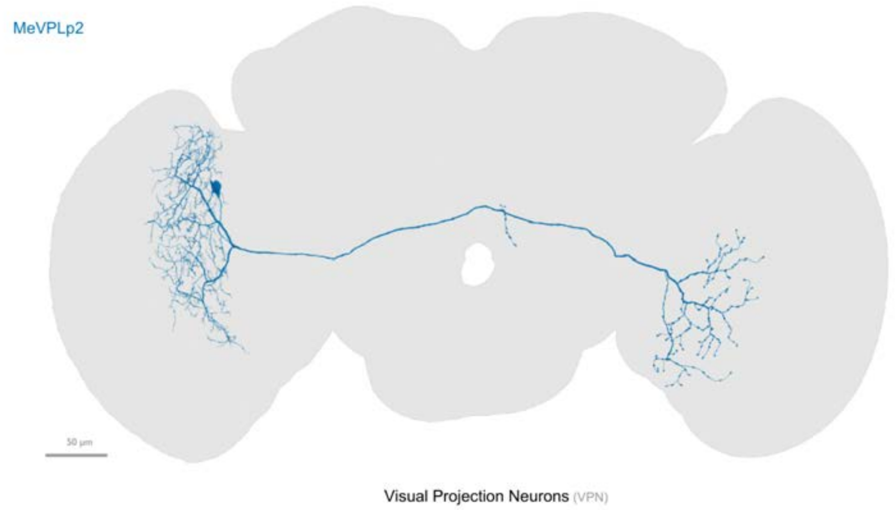


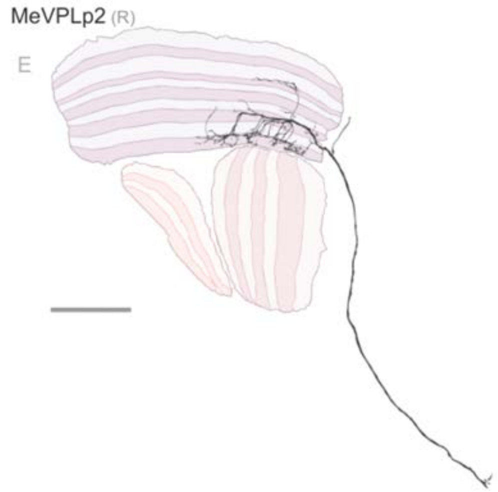


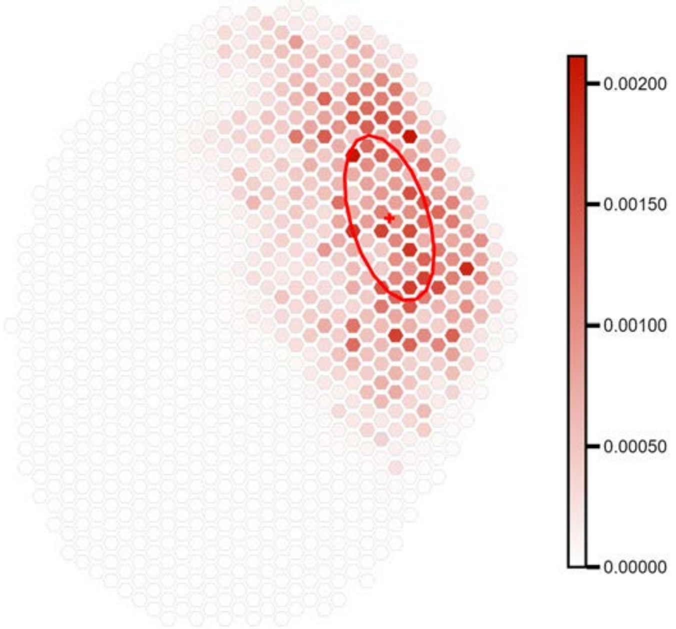


fewer than 100 output connections in central brain

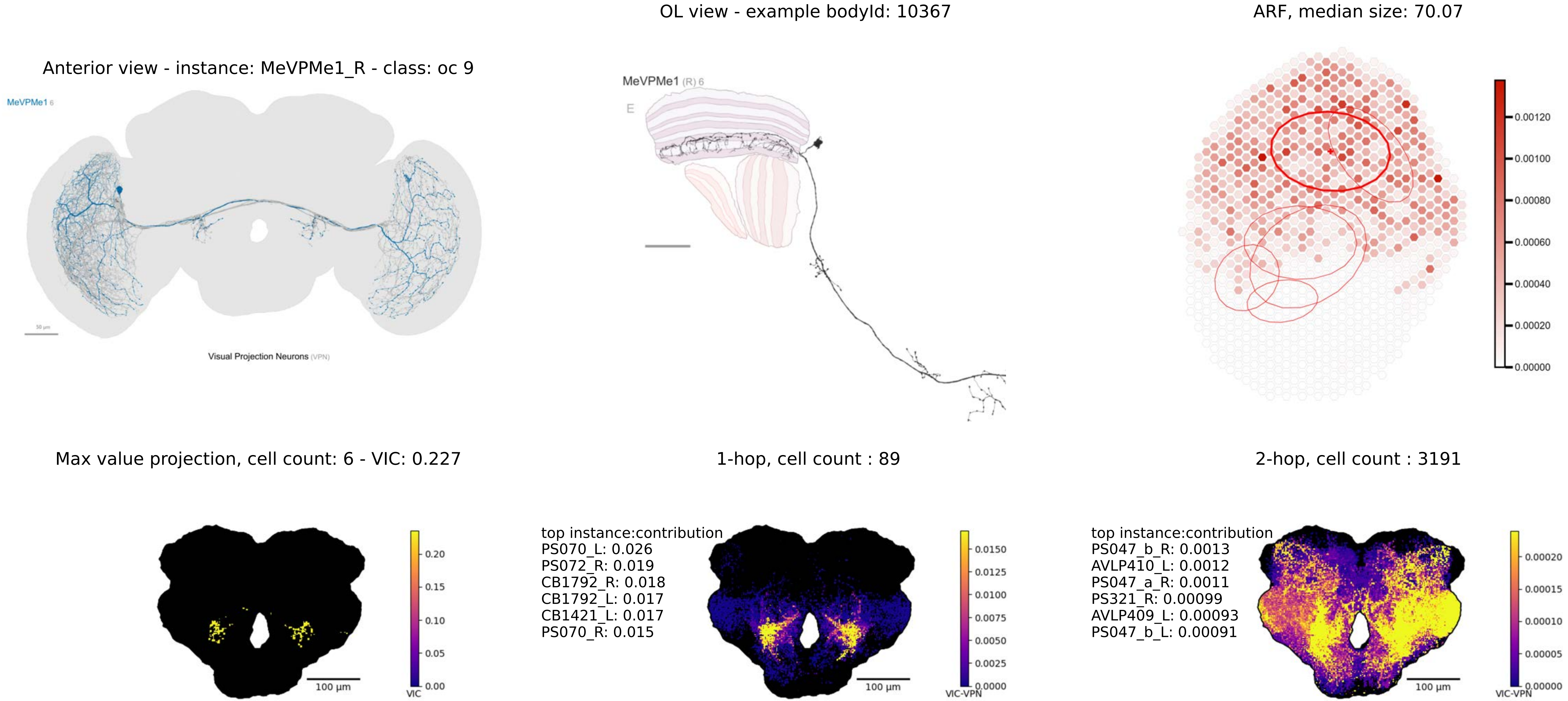
OL view - example bodyId: 10367
ARF, median size: 70.07
Anterior view - instance: MeVPMe1_R - class: oc 9
MeVPMe1 6
Visual Projection Neurons (VPN)
MeVPMe1 (R) 6
E
0.00120
0.00100
0.00080
0.00060
0.00040
0.00020
0.00000
Max value projection, cell count: 6 - VIC: 0.227
1-hop, cell count : 89
2-hop, cell count : 3191
0.20
0.15
0.10
0.05
0.00
VIC
100 µm
top instance:contribution
PS070_L: 0.026
PS072_R: 0.019
CB1792_R: 0.018
CB1792_L: 0.017
CB1421_L: 0.017
PS070_R: 0.015
0.0150
0.0125
0.0100
0.0075
0.0050
0.0025
0.0000
VIC-VPN
100 µm
top instance:contribution
PS047_b_R: 0.0013
AVLP410_L: 0.0012
PS047_a_R: 0.0011
PS321_R: 0.00099
AVLP409_L: 0.00093
PS047_b_L: 0.00091
0.00020
0.00015
0.00010
0.00005
0.00000
VIC-VPN
100 µm

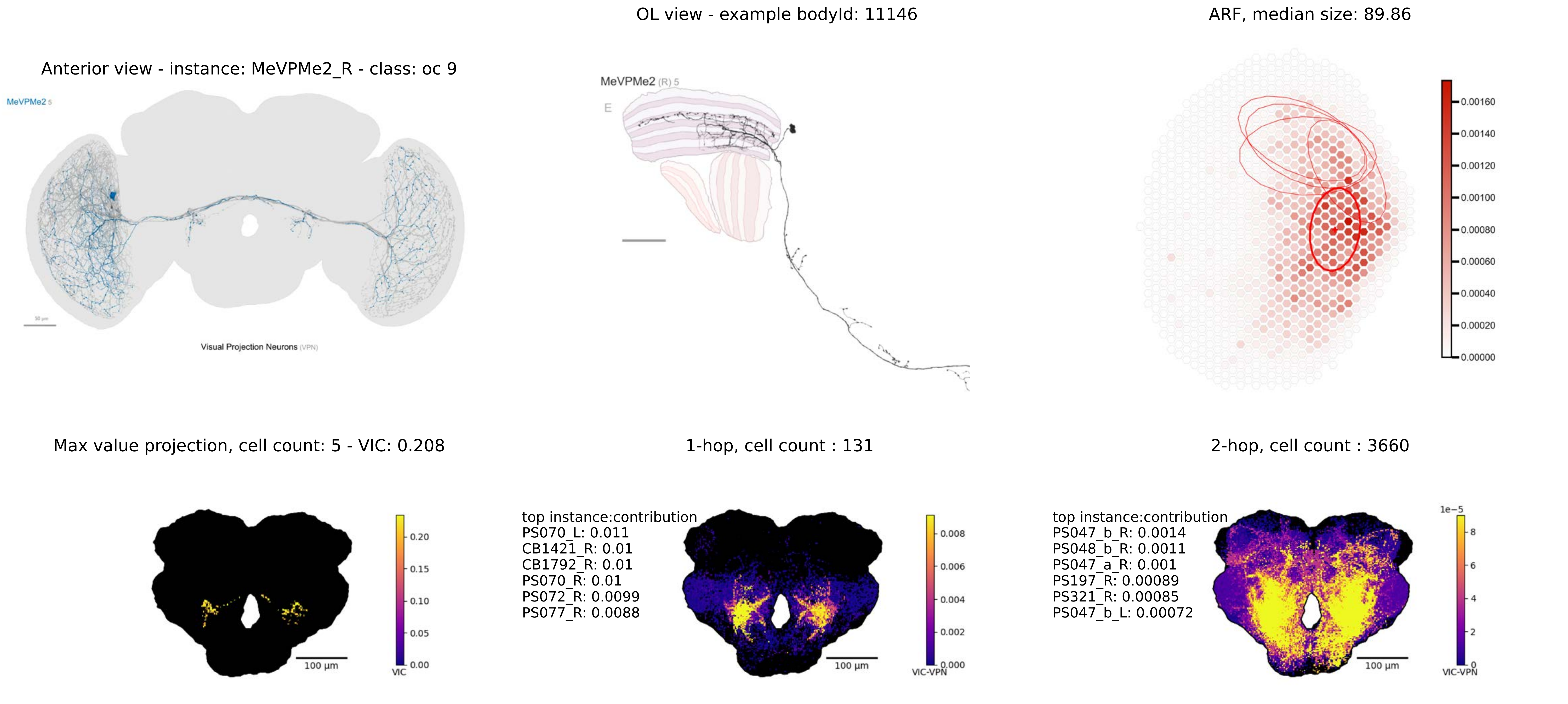

Anterior view - instance: MeVPMe2_R - class: oc 9
MeVPMe2 5
Visual Projection Neurons (VPN)
OL view - example bodyId: 11146
MeVPMe2 (R) 5
E
ARF, median size: 89.86
0.00160
0.00140
0.00120
0.00100
0.00080
0.00060
0.00040
0.00020
0.00000
Max value projection, cell count: 5 - VIC: 0.208
100 µm
VIC
0.20
0.15
0.10
0.05
0.00
1-hop, cell count : 131
top instance:contribution
PS070_L: 0.011
CB1421_R: 0.01
CB1792_R: 0.01
PS070_R: 0.01
PS072_R: 0.0099
PS077_R: 0.0088
100 µm
VIC-VPN
0.008
0.006
0.004
0.002
0.000
2-hop, cell count : 3660
top instance:contribution
PS047_b_R: 0.0014
PS048_b_R: 0.0011
PS047_a_R: 0.001
PS197_R: 0.00089
PS321_R: 0.00085
PS047_b_L: 0.00072
100 µm
1e−5
VIC-VPN
8
6
4
2
0

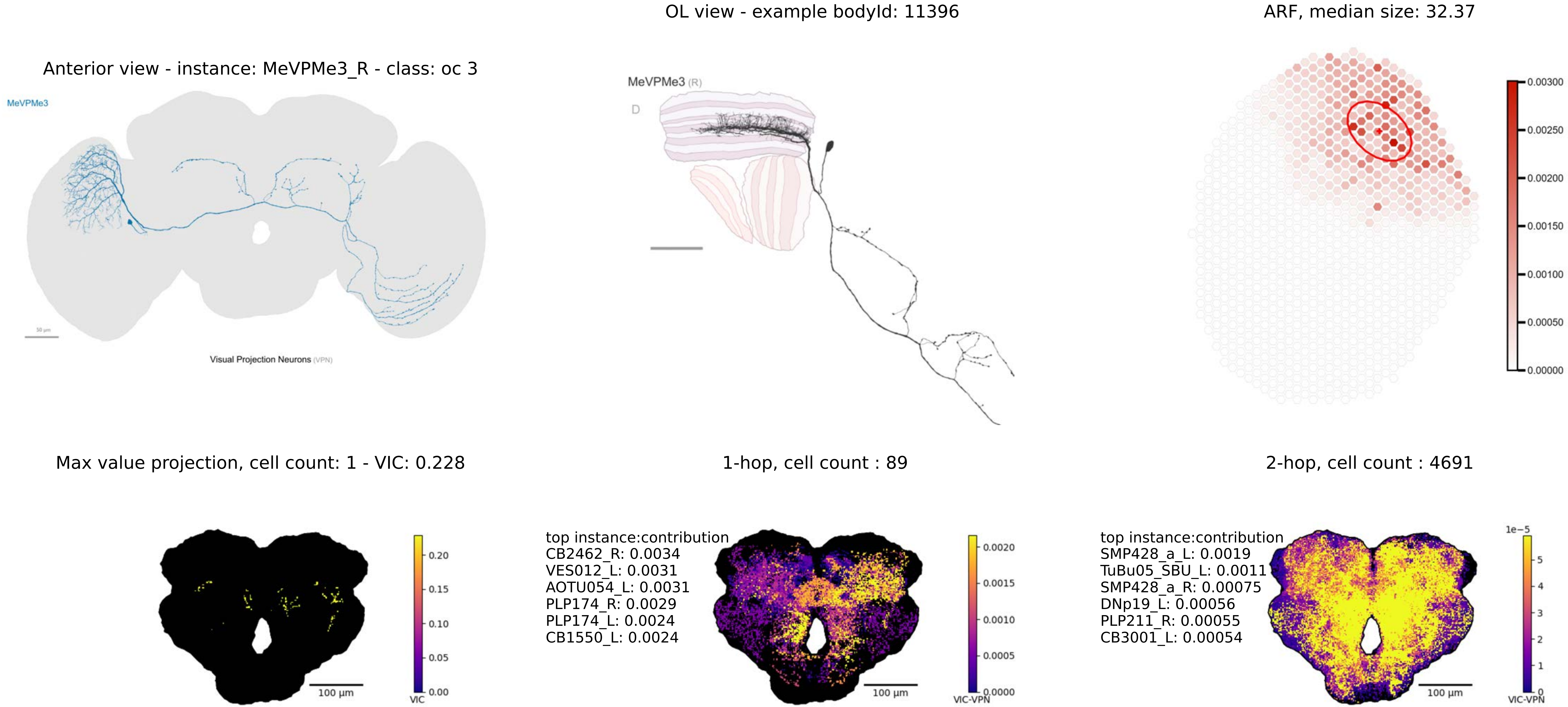

OL view - example bodyId: 11396
ARF, median size: 32.37
Anterior view - instance: MeVPMe3_R - class: oc 3
MeVPMe3
50 µm
Visual Projection Neurons (VPN)
MeVPMe3 (R)
D
0.00300
0.00250
0.00200
0.00150
0.00100
0.00050
0.00000
Max value projection, cell count: 1 - VIC: 0.228
1-hop, cell count : 89
2-hop, cell count : 4691
100 µm
VIC
0.20
0.15
0.10
0.05
0.00
top instance:contribution
CB2462_R: 0.0034
VES012_L: 0.0031
AOTU054_L: 0.0031
PLP174_R: 0.0029
PLP174_L: 0.0024
CB1550_L: 0.0024
0.0020
0.0015
0.0010
0.0005
0.0000
VIC-VPN
top instance:contribution
SMP428_a_L: 0.0019
TuBu05_SBU_L: 0.0011
SMP428_a_R: 0.00075
DNp19_L: 0.00056
PLP211_R: 0.00055
CB3001_L: 0.00054
1e−5
5
4
3
2
1
0
VIC-VPN

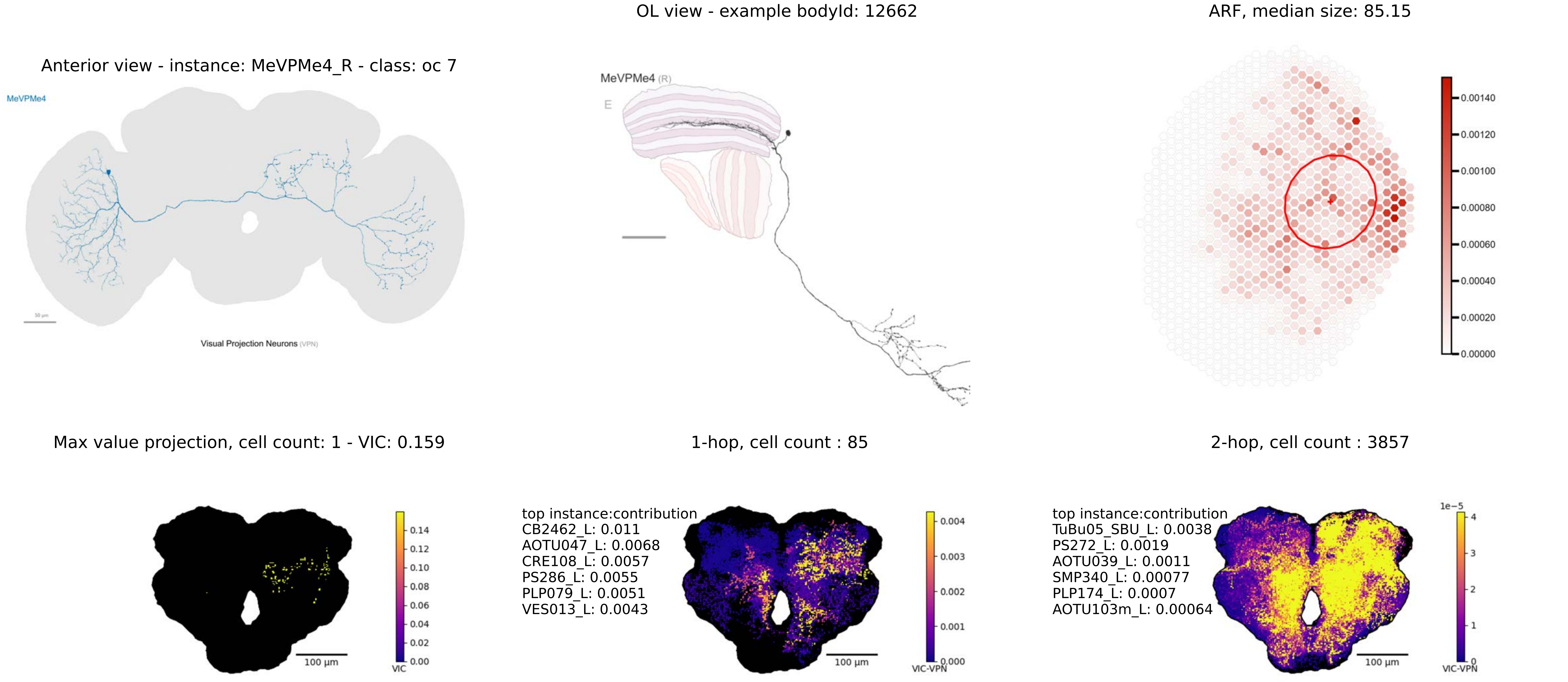
OL view - example bodyId: 12662
ARF, median size: 85.15
Anterior view - instance: MeVPMe4_R - class: oc 7
MeVPMe4
Visual Projection Neurons (VPN)
MeVPMe4 (R)
E
0.00140
0.00120
0.00100
0.00080
0.00060
0.00040
0.00020
0.00000
Max value projection, cell count: 1 - VIC: 0.159
1-hop, cell count : 85
2-hop, cell count : 3857
0.14
0.12
0.10
0.08
0.06
0.04
0.02
0.00
VIC
100 µm
top instance:contribution
CB2462_L: 0.011
AOTU047_L: 0.0068
CRE108_L: 0.0057
PS286_L: 0.0055
PLP079_L: 0.0051
VES013_L: 0.0043
0.004
0.003
0.002
0.001
0.000
VIC-VPN
100 µm
top instance:contribution
TuBu05_SBU_L: 0.0038
PS272_L: 0.0019
AOTU039_L: 0.0011
SMP340_L: 0.00077
PLP174_L: 0.0007
AOTU103m_L: 0.00064
1e−5
4
3
2
1
0
VIC-VPN
100 µm

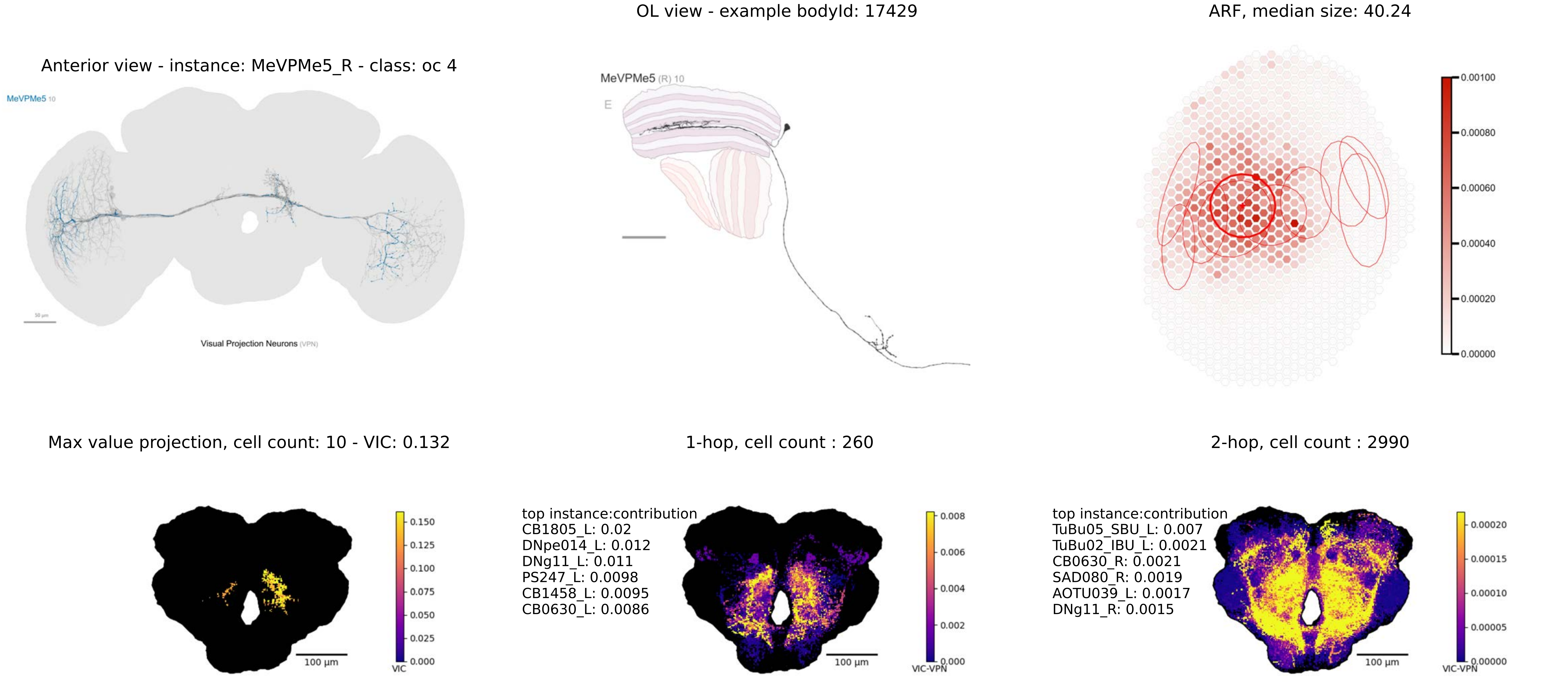
OL view - example bodyId: 17429
ARF, median size: 40.24
Anterior view - instance: MeVPMe5_R - class: oc 4
MeVPMe5 10
Visual Projection Neurons (VPN)
MeVPMe5 (R) 10
E
0.00100
0.00080
0.00060
0.00040
0.00020
0.00000
Max value projection, cell count: 10 - VIC: 0.132
1-hop, cell count : 260
2-hop, cell count : 2990
100 µm
VIC
0.150
0.125
0.100
0.075
0.050
0.025
0.000
top instance:contribution
CB1805_L: 0.02
DNpe014_L: 0.012
DNg11_L: 0.011
PS247_L: 0.0098
CB1458_L: 0.0095
CB0630_L: 0.0086
VIC-VPN
0.008
0.006
0.004
0.002
0.000
top instance:contribution
TuBu05_SBU_L: 0.007
TuBu02_IBU_L: 0.0021
CB0630_R: 0.0021
SAD080_R: 0.0019
AOTU039_L: 0.0017
DNg11_R: 0.0015
0.00020
0.00015
0.00010
0.00005
0.00000

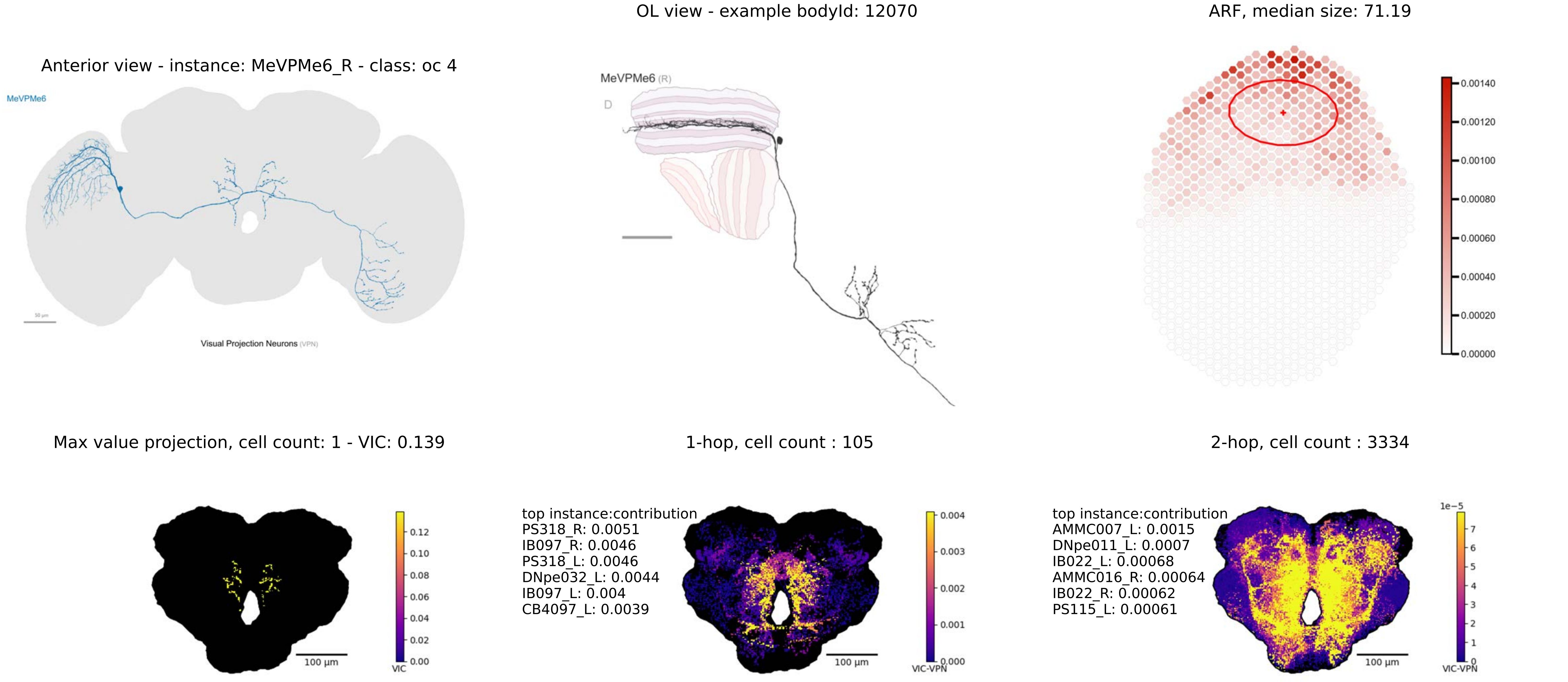

Anterior view - instance: MeVPMe6_R - class: oc 4
MeVPMe6
Visual Projection Neurons (VPN)
OL view - example bodyId: 12070
MeVPMe6 (R)
D
ARF, median size: 71.19
0.00140
0.00120
0.00100
0.00080
0.00060
0.00040
0.00020
0.00000
Max value projection, cell count: 1 - VIC: 0.139
0.12
0.10
0.08
0.06
0.04
0.02
0.00
VIC
100 μm
1-hop, cell count : 105
top instance:contribution
PS318_R: 0.0051
IB097_R: 0.0046
PS318_L: 0.0046
DNpe032_L: 0.0044
IB097_L: 0.004
CB4097_L: 0.0039
0.004
0.003
0.002
0.001
0.000
VIC-VPN
100 μm
2-hop, cell count : 3334
top instance:contribution
AMMC007_L: 0.0015
DNpe011_L: 0.0007
IB022_L: 0.00068
AMMC016_R: 0.00064
IB022_R: 0.00062
PS115_L: 0.00061
1e−5
7
6
5
4
3
2
1
0
VIC-VPN
100 μm

Anterior view - instance: MeVPMe7_R - class: oc 5

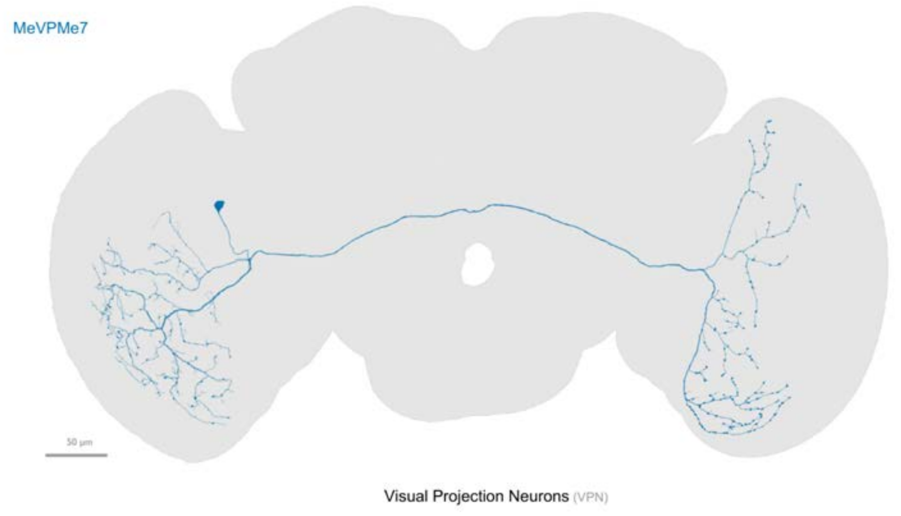


fewer than 100 output connections in central brain

OL view - example bodyId: 11669

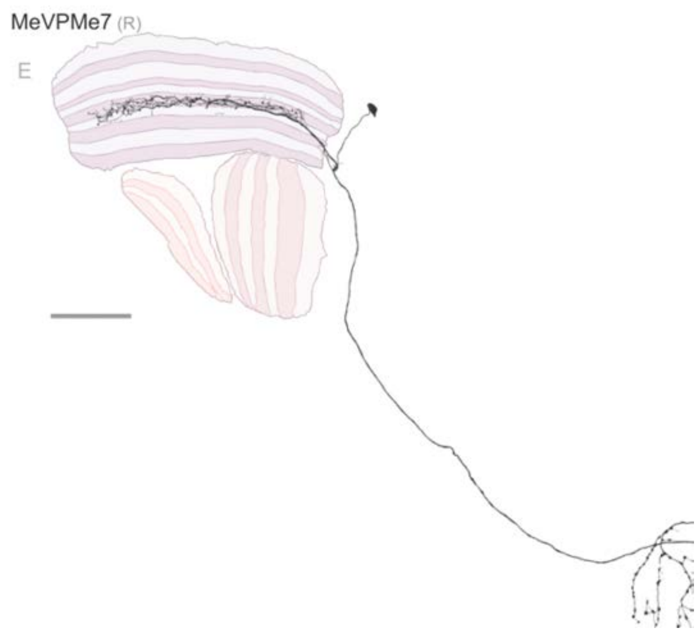


ARF, median size: 81.22

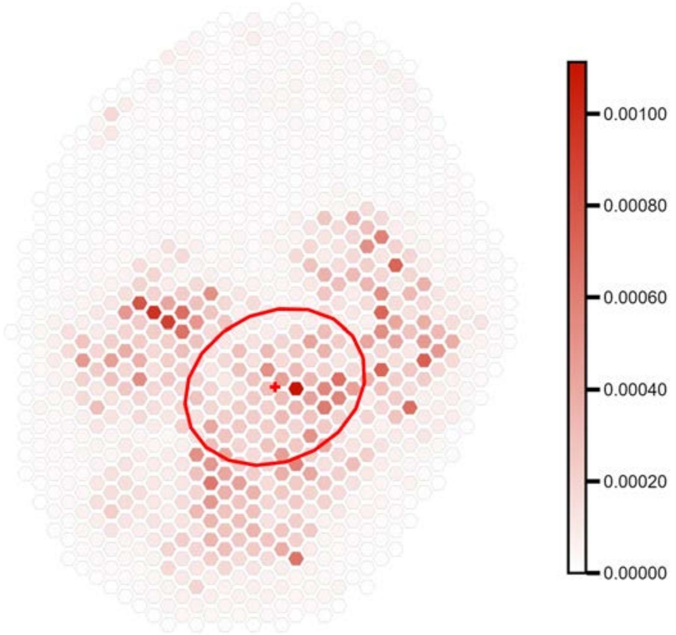

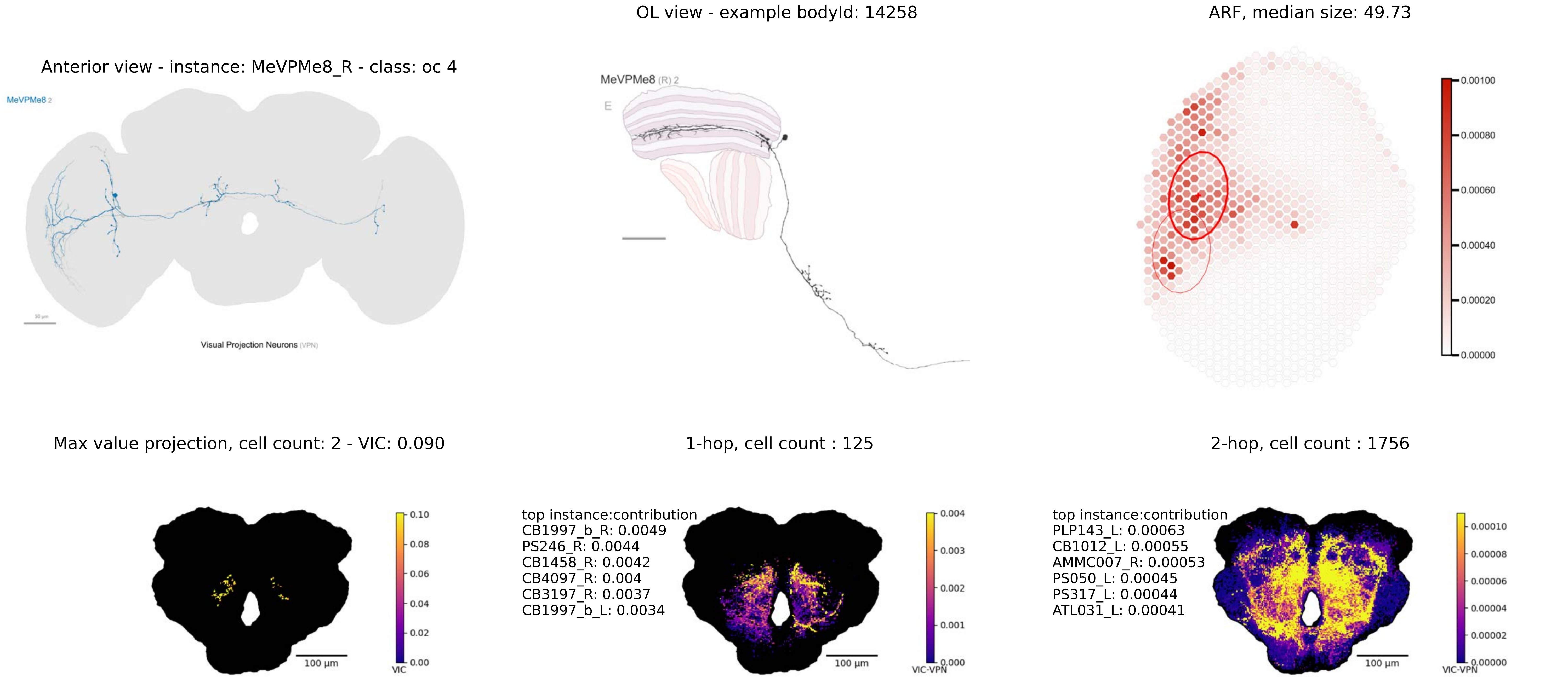

OL view - example bodyId: 14258
ARF, median size: 49.73
Anterior view - instance: MeVPMe8_R - class: oc 4
MeVPMe8 2
Visual Projection Neurons (VPN)
MeVPMe8 (R) 2
E
0.00100
0.00080
0.00060
0.00040
0.00020
0.00000
Max value projection, cell count: 2 - VIC: 0.090
1-hop, cell count : 125
2-hop, cell count : 1756
100 µm
VIC
0.10
0.08
0.06
0.04
0.02
0.00
top instance:contribution
CB1997_b_R: 0.0049
PS246_R: 0.0044
CB1458_R: 0.0042
CB4097_R: 0.004
CB3197_R: 0.0037
CB1997_b_L: 0.0034
VIC-VPN
0.004
0.003
0.002
0.001
0.000
top instance:contribution
PLP143_L: 0.00063
CB1012_L: 0.00055
AMMC007_R: 0.00053
PS050_L: 0.00045
PS317_L: 0.00044
ATL031_L: 0.00041
0.00010
0.00008
0.00006
0.00004
0.00002
0.00000

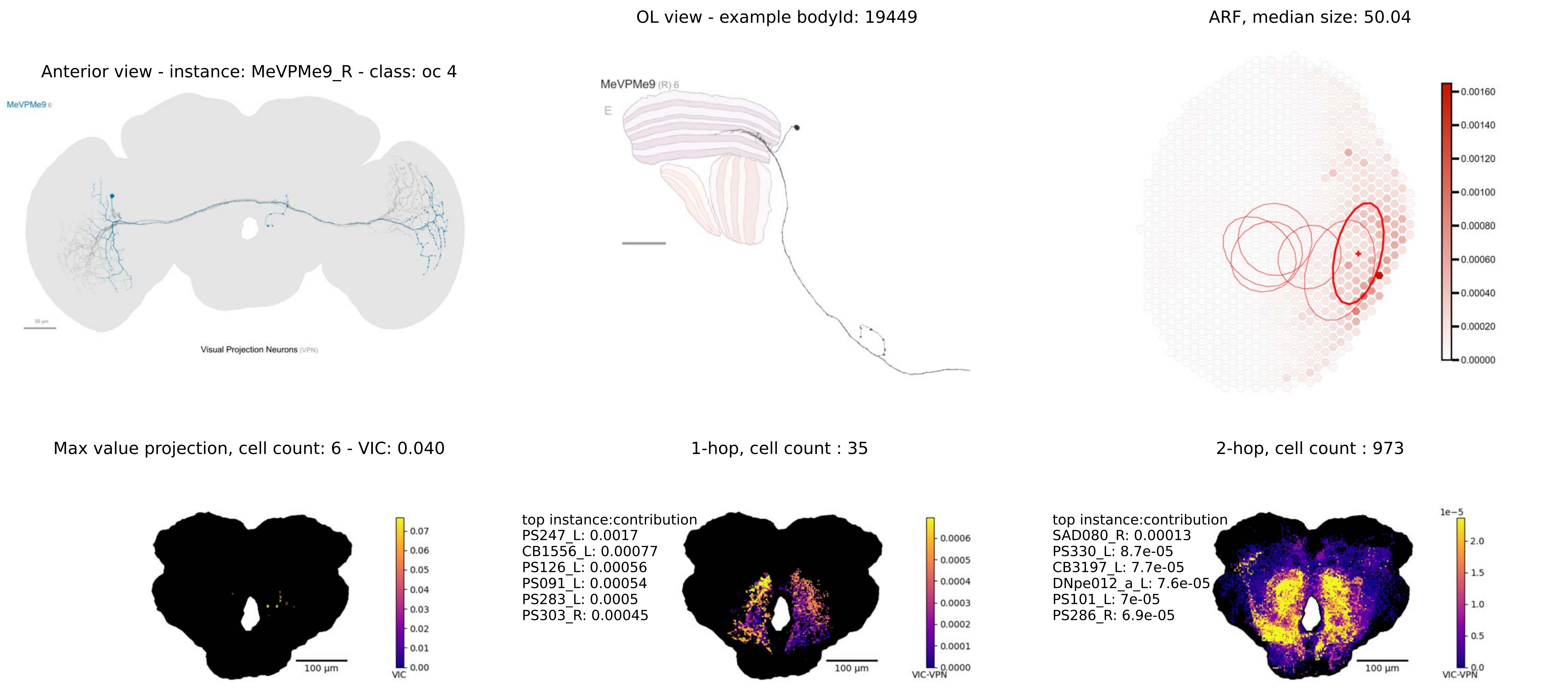
Anterior view - instance: MeVPMe9_R - class: oc 4
MeVPMe9 6
50 µm
Visual Projection Neurons (VPN)
OL view - example bodyId: 19449
MeVPMe9 (R) 6
E
ARF, median size: 50.04
0.00160
0.00140
0.00120
0.00100
0.00080
0.00060
0.00040
0.00020
0.00000
Max value projection, cell count: 6 - VIC: 0.040
100 µm
0.07
0.06
0.05
0.04
0.03
0.02
0.01
0.00
VIC
1-hop, cell count : 35
top instance:contribution
PS247_L: 0.0017
CB1556_L: 0.00077
PS126_L: 0.00056
PS091_L: 0.00054
PS283_L: 0.0005
PS303_R: 0.00045
100 µm
0.0006
0.0005
0.0004
0.0003
0.0002
0.0001
0.0000
VIC-VPN
2-hop, cell count : 973
top instance:contribution
SAD080_R: 0.00013
PS330_L: 8.7e-05
CB3197_L: 7.7e-05
DNpe012_a_L: 7.6e-05
PS101_L: 7e-05
PS286_R: 6.9e-05
100 µm
1e−5
2.0
1.5
1.0
0.5
0.0
VIC-VPN

Anterior view - instance: MeVPMe11_R - class: oc 3

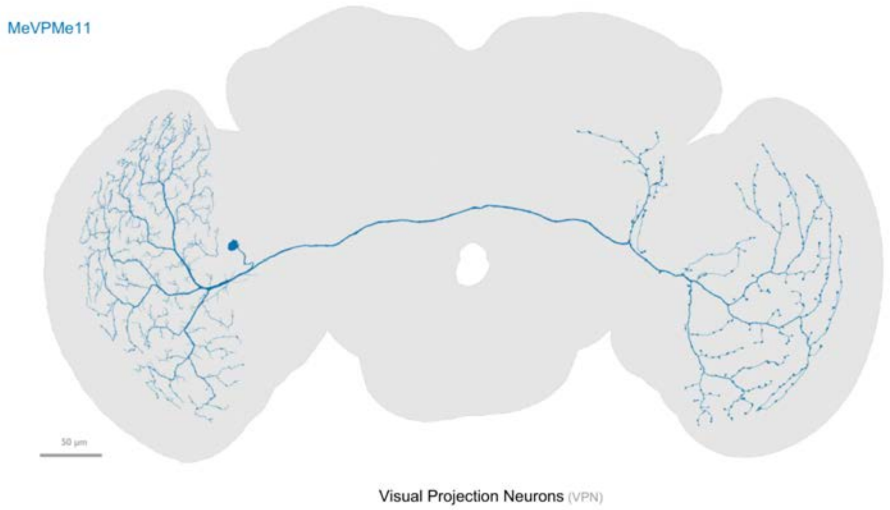


fewer than 100 output connections in central brain

OL view - example bodyId: 11202

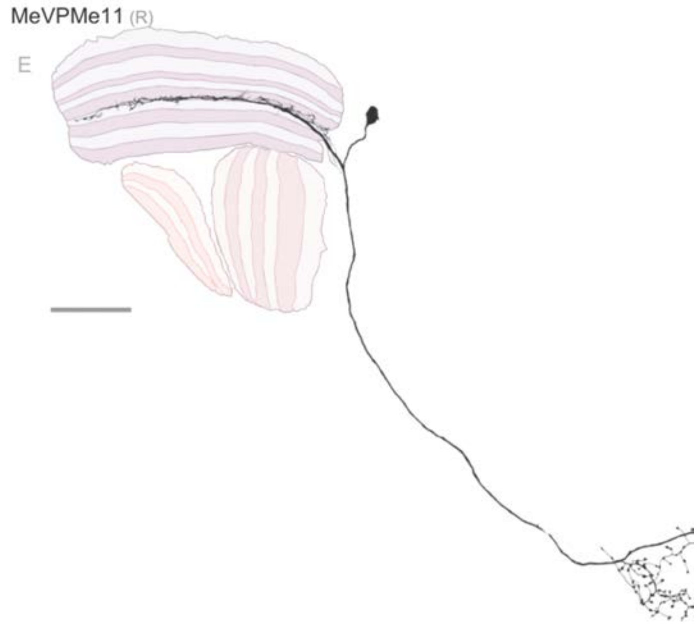


ARF, median size: 97.47

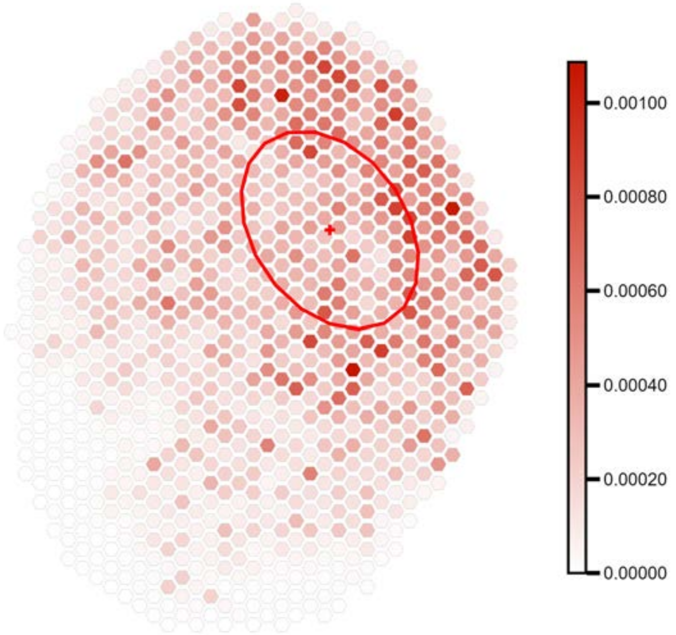

Anterior view - instance: MeVPMe12_R - class: oc 7

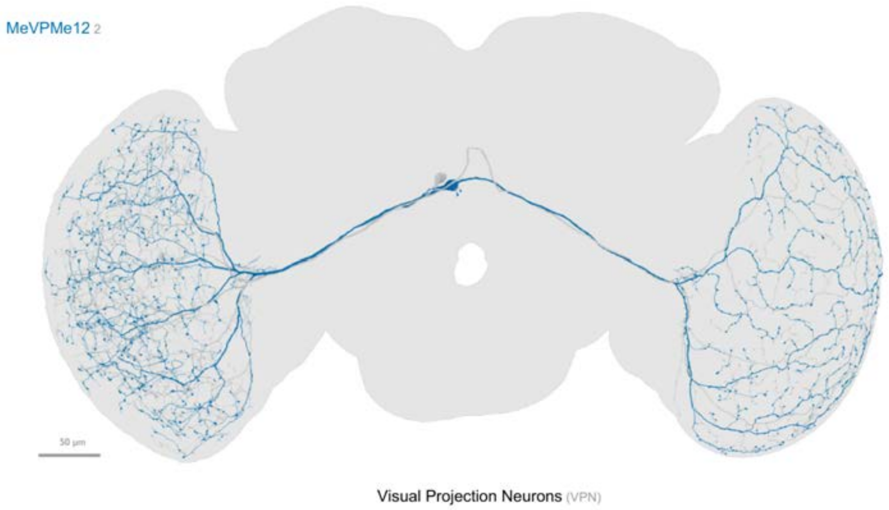


fewer than 100 output connections in central brain

OL view - example bodyId: 10358

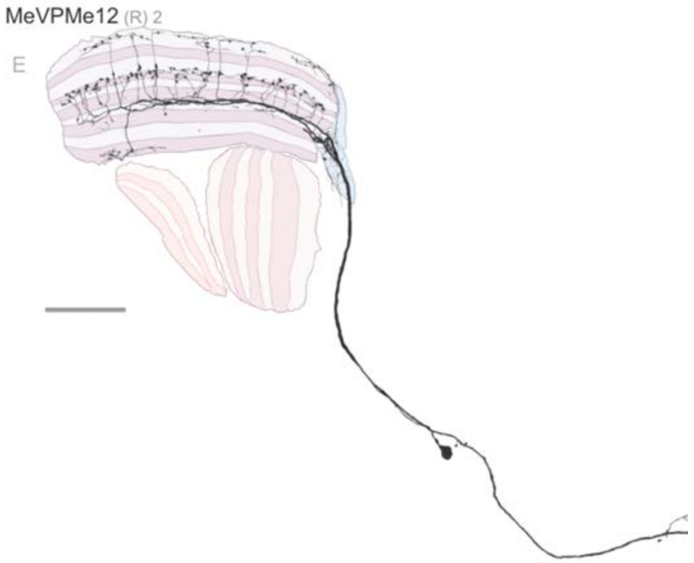


ARF, median size: 145.2

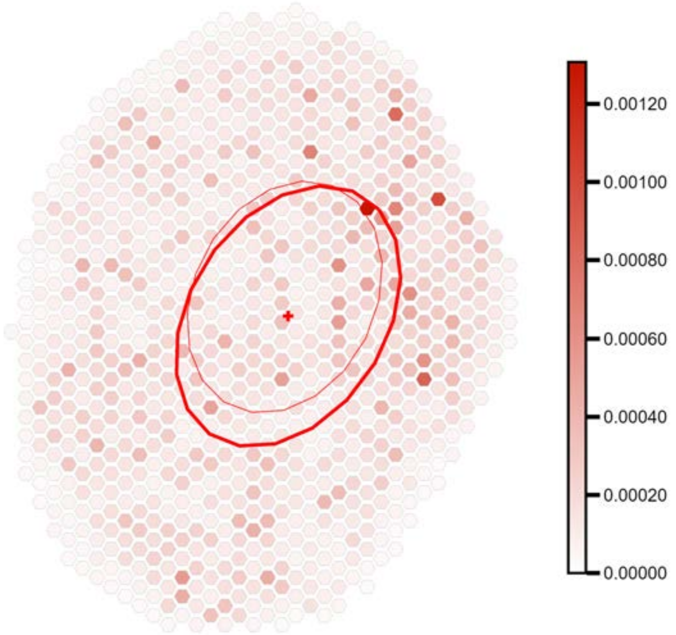

Anterior view - instance: MeVPMe13_R - class: oc 5

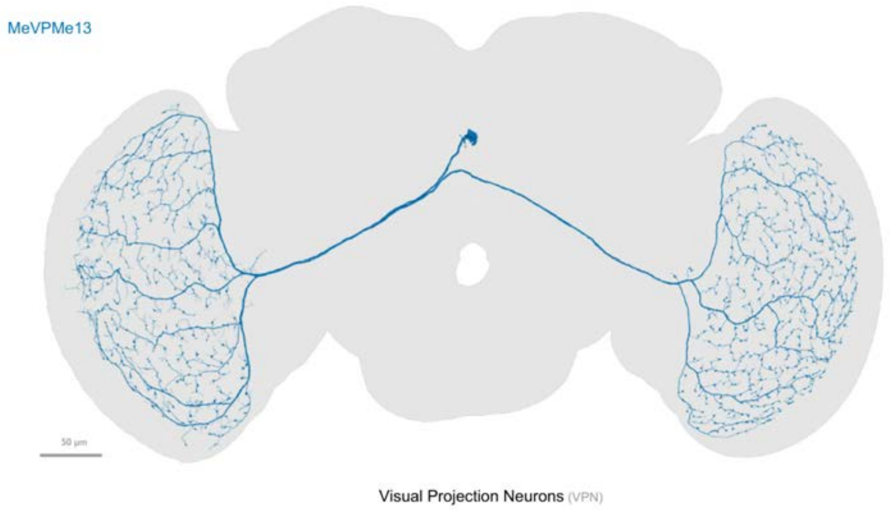


fewer than 100 output connections in central brain

OL view - example bodyId: 10275

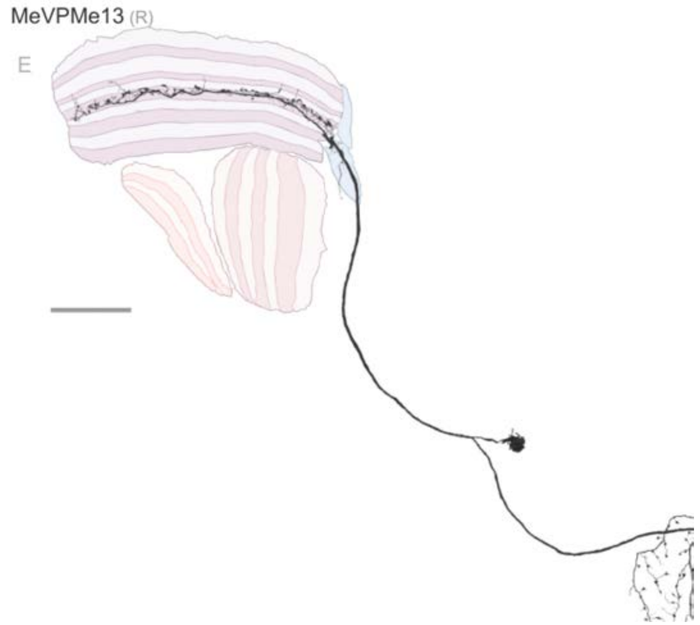


ARF, median size: 142.19

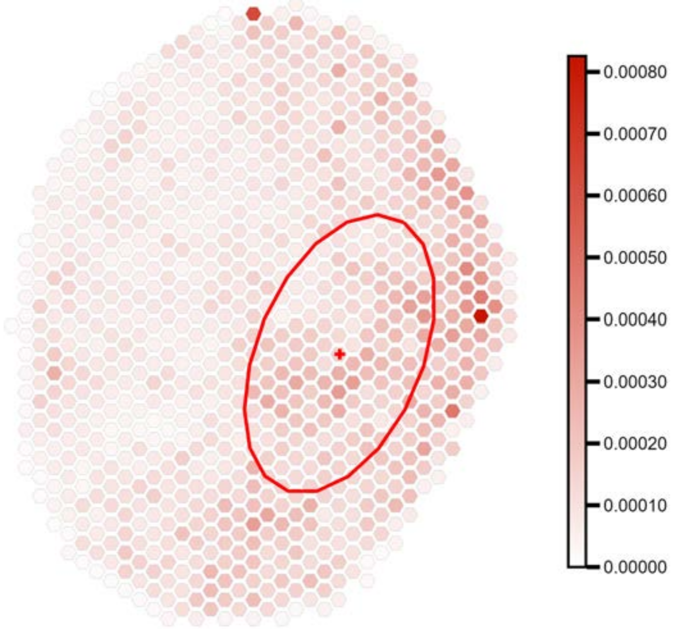

Anterior view - instance: MeVPOL1_R - class: oc 8

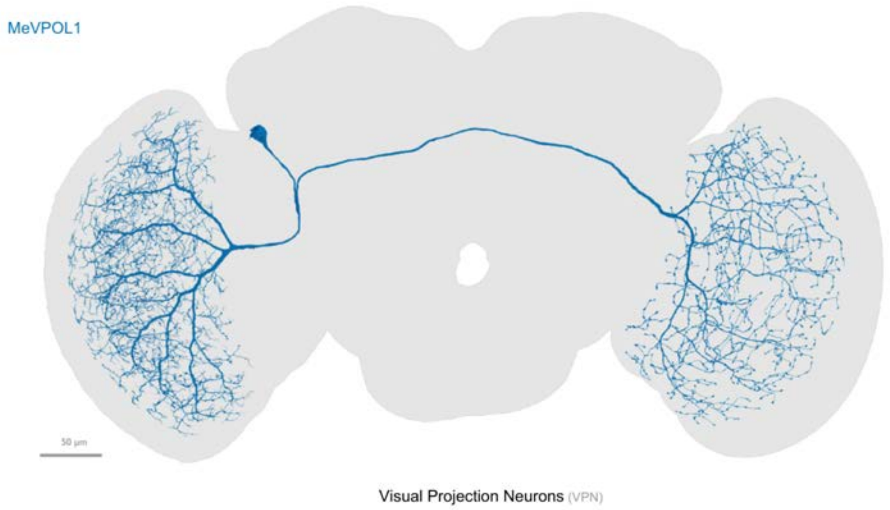


fewer than 100 output connections in central brain

OL view - example bodyId: 10330

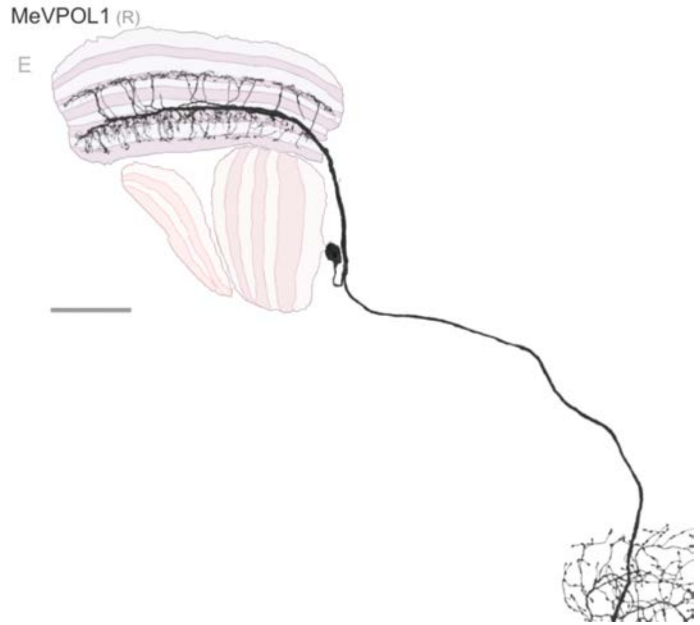


ARF, median size: 117.98

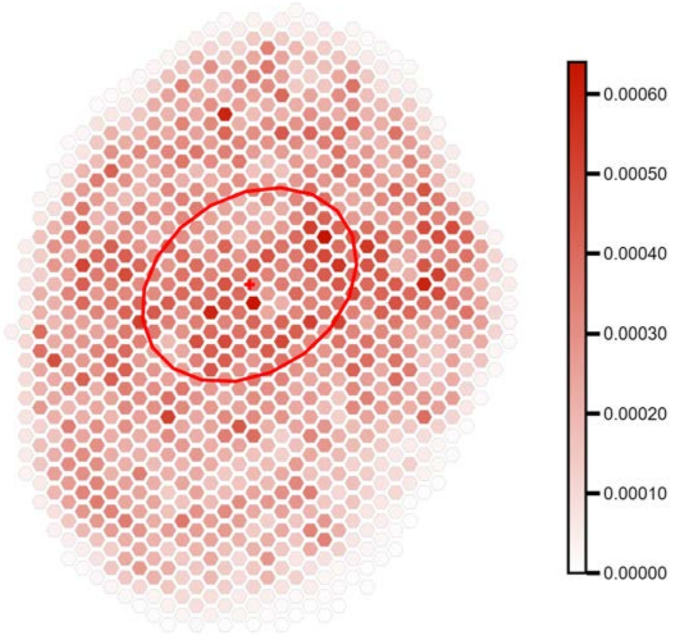

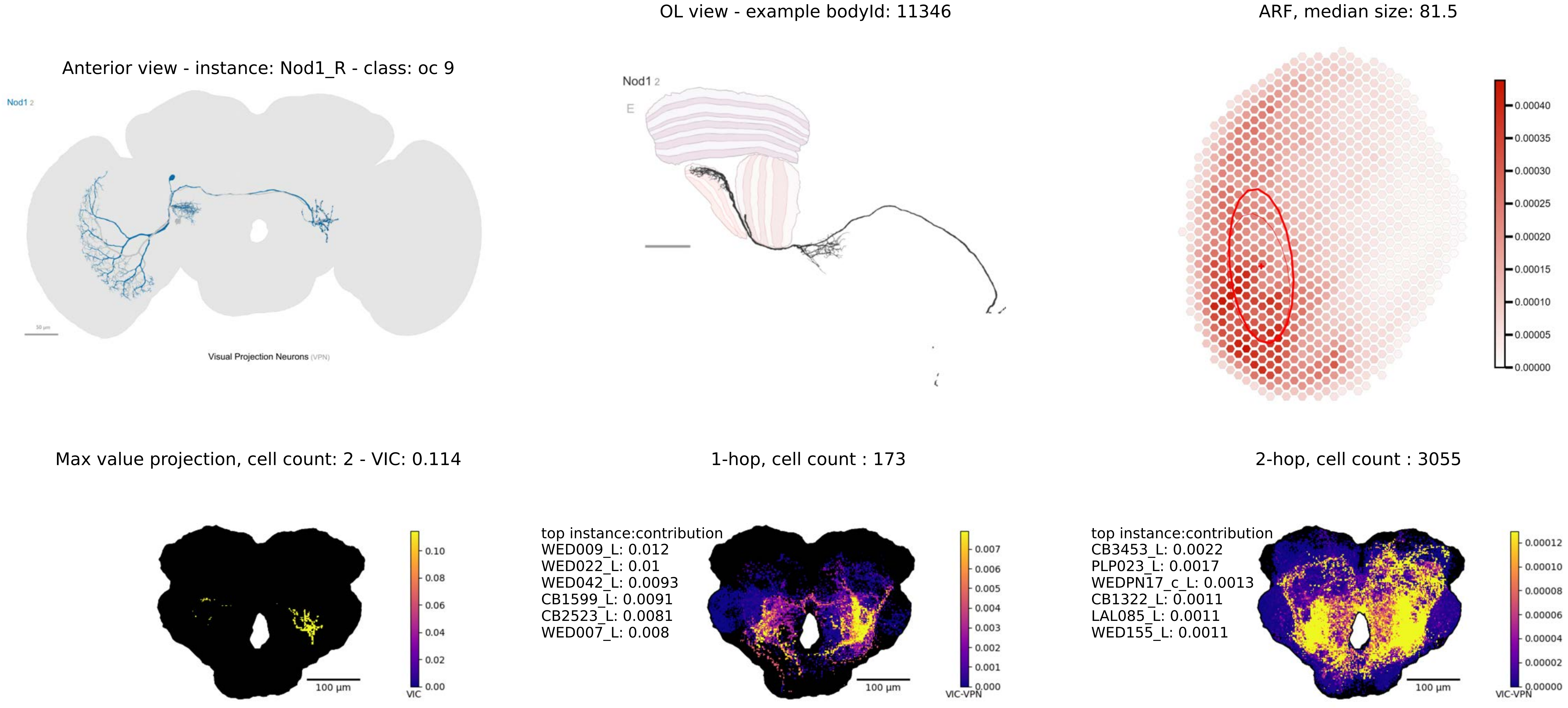
Anterior view - instance: Nod1_R - class: oc 9
Nod1 2
Visual Projection Neurons (VPN)
OL view - example bodyId: 11346
Nod1 2
E
ARF, median size: 81.5
0.00040
0.00035
0.00030
0.00025
0.00020
0.00015
0.00010
0.00005
0.00000
Max value projection, cell count: 2 - VIC: 0.114
100 µm
VIC
0.10
0.08
0.06
0.04
0.02
0.00
1-hop, cell count : 173
top instance:contribution
WED009_L: 0.012
WED022_L: 0.01
WED042_L: 0.0093
CB1599_L: 0.0091
CB2523_L: 0.0081
WED007_L: 0.008
100 µm
VIC-VPN
0.007
0.006
0.005
0.004
0.003
0.002
0.001
0.000
2-hop, cell count : 3055
top instance:contribution
CB3453_L: 0.0022
PLP023_L: 0.0017
WEDPN17_c_L: 0.0013
CB1322_L: 0.0011
LAL085_L: 0.0011
WED155_L: 0.0011
100 µm
VIC-VPN
0.00012
0.00010
0.00008
0.00006
0.00004
0.00002
0.00000

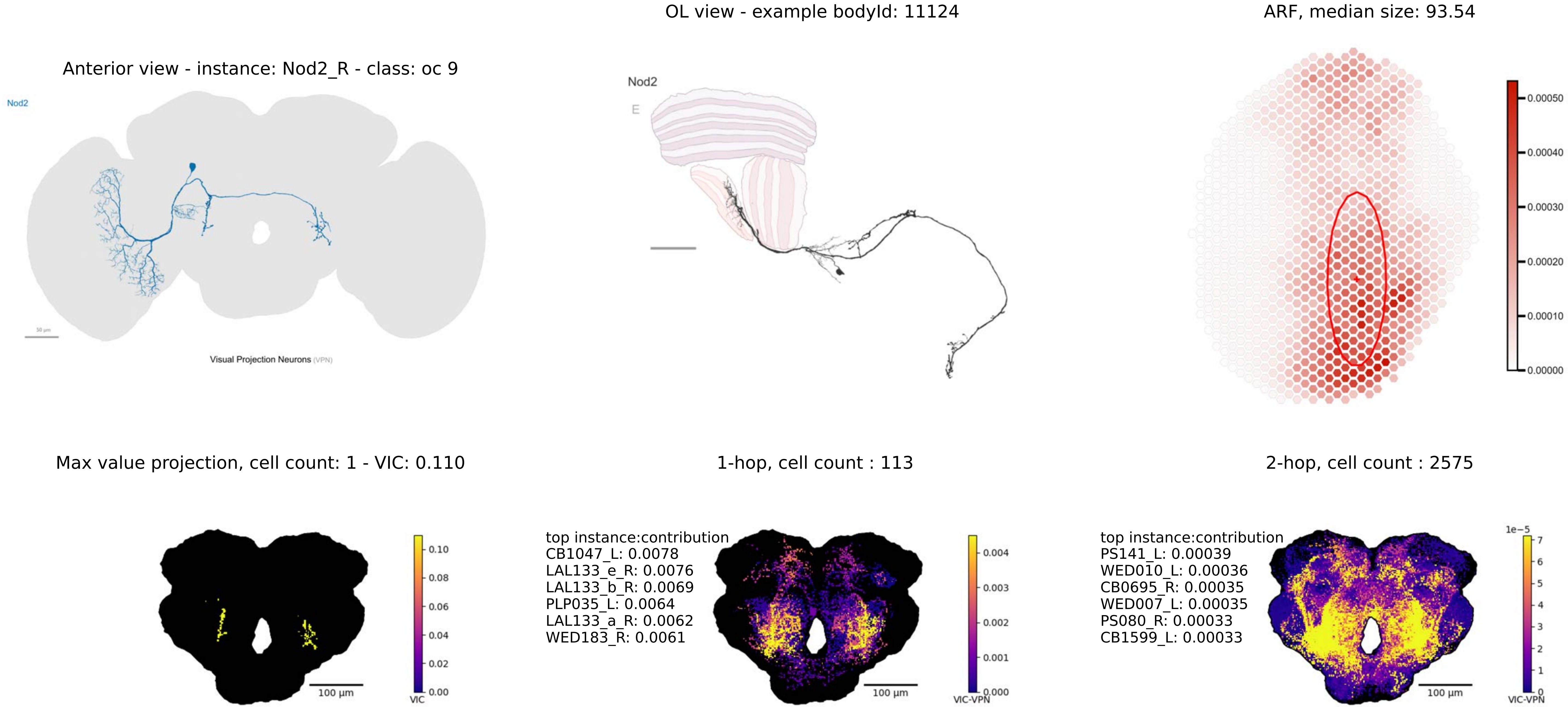
OL view - example bodyId: 11124
ARF, median size: 93.54
Anterior view - instance: Nod2_R - class: oc 9
Nod2
Visual Projection Neurons (VPN)
Nod2
E
0.00050
0.00040
0.00030
0.00020
0.00010
0.00000
Max value projection, cell count: 1 - VIC: 0.110
1-hop, cell count : 113
2-hop, cell count : 2575
100 µm
0.10
0.08
0.06
0.04
0.02
0.00
VIC
top instance:contribution
CB1047_L: 0.0078
LAL133_e_R: 0.0076
LAL133_b_R: 0.0069
PLP035_L: 0.0064
LAL133_a_R: 0.0062
WED183_R: 0.0061
100 µm
0.004
0.003
0.002
0.001
0.000
VIC-VPN
top instance:contribution
PS141_L: 0.00039
WED010_L: 0.00036
CB0695_R: 0.00035
WED007_L: 0.00035
PS080_R: 0.00033
CB1599_L: 0.00033
100 µm
1e−5
7
6
5
4
3
2
1
0
VIC-VPN

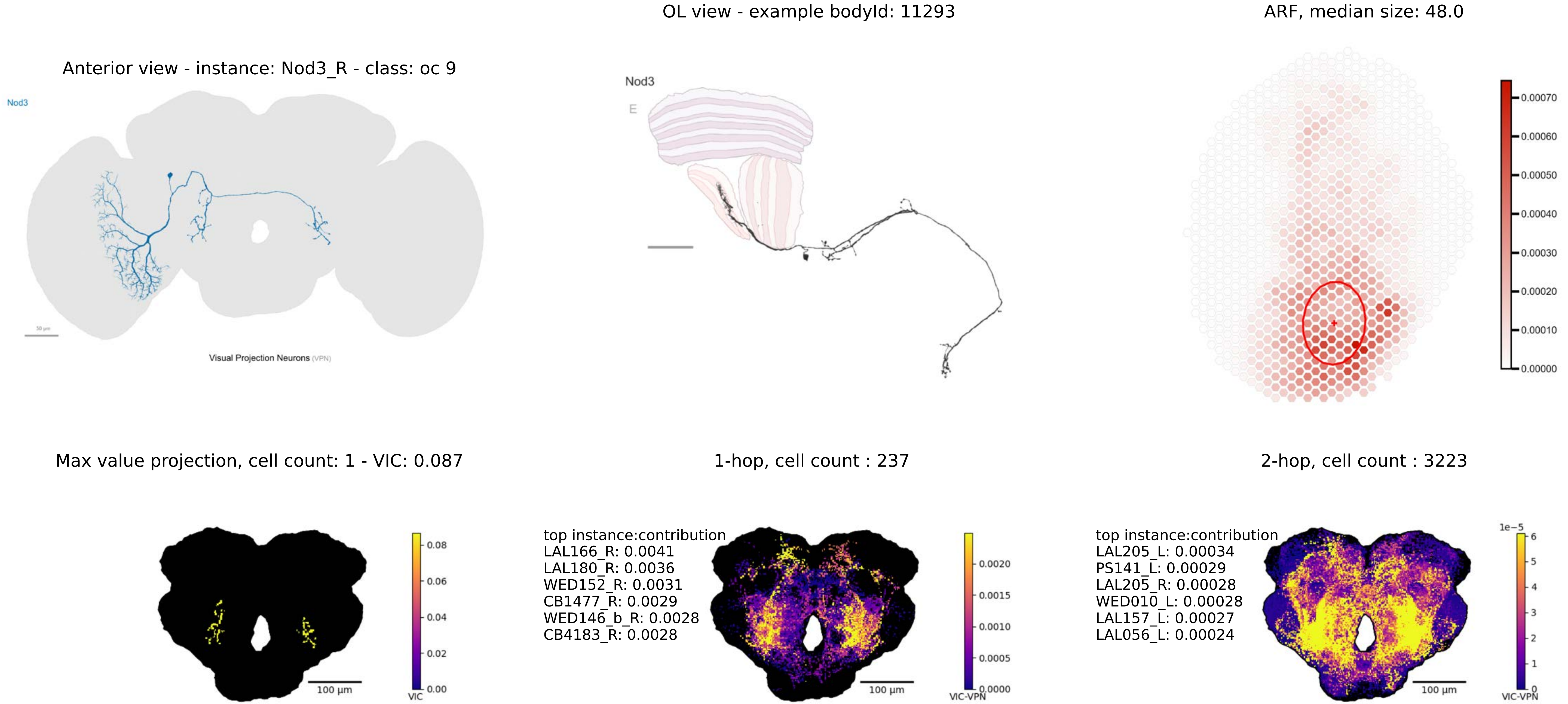

OL view - example bodyId: 11293
ARF, median size: 48.0
Anterior view - instance: Nod3_R - class: oc 9
Nod3
Visual Projection Neurons (VPN)
Nod3
E
0.00070
0.00060
0.00050
0.00040
0.00030
0.00020
0.00010
0.00000
Max value projection, cell count: 1 - VIC: 0.087
1-hop, cell count : 237
2-hop, cell count : 3223
0.08
0.06
0.04
0.02
0.00
VIC
100 µm
top instance:contribution
LAL166_R: 0.0041
LAL180_R: 0.0036
WED152_R: 0.0031
CB1477_R: 0.0029
WED146_b_R: 0.0028
CB4183_R: 0.0028
0.0020
0.0015
0.0010
0.0005
0.0000
VIC-VPN
100 µm
top instance:contribution
LAL205_L: 0.00034
PS141_L: 0.00029
LAL205_R: 0.00028
WED010_L: 0.00028
LAL157_L: 0.00027
LAL056_L: 0.00024
1e−5
6
5
4
3
2
1
0
VIC-VPN
100 µm

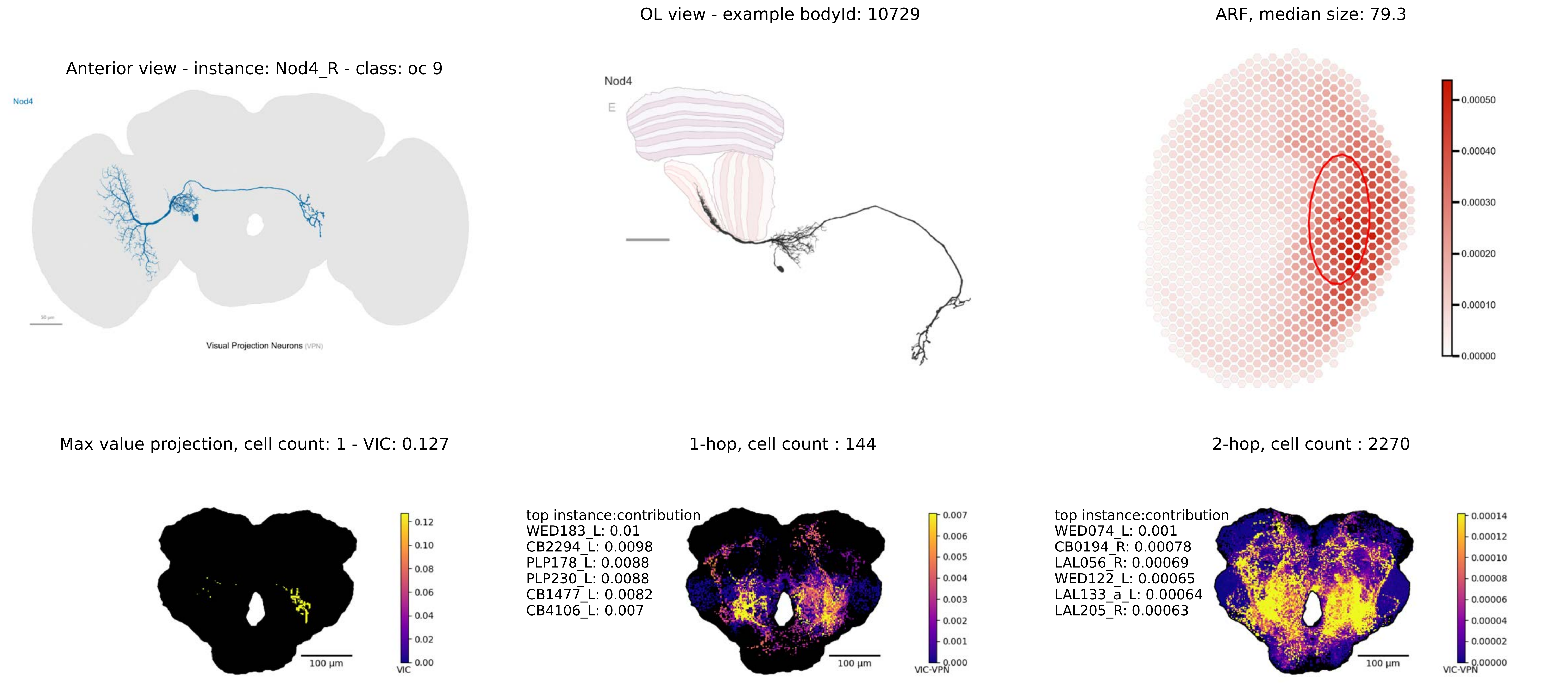

OL view - example bodyId: 10729
ARF, median size: 79.3
Anterior view - instance: Nod4_R - class: oc 9
Nod4
Visual Projection Neurons (VPN)
Nod4
E
0.00050
0.00040
0.00030
0.00020
0.00010
0.00000
Max value projection, cell count: 1 - VIC: 0.127
1-hop, cell count : 144
2-hop, cell count : 2270
100 µm
VIC
0.12
0.10
0.08
0.06
0.04
0.02
0.00
top instance:contribution
WED183_L: 0.01
CB2294_L: 0.0098
PLP178_L: 0.0088
PLP230_L: 0.0088
CB1477_L: 0.0082
CB4106_L: 0.007
VIC-VPN
0.007
0.006
0.005
0.004
0.003
0.002
0.001
0.000
top instance:contribution
WED074_L: 0.001
CB0194_R: 0.00078
LAL056_R: 0.00069
WED122_L: 0.00065
LAL133_a_L: 0.00064
LAL205_R: 0.00063
0.00014
0.00012
0.00010
0.00008
0.00006
0.00004
0.00002
0.00000

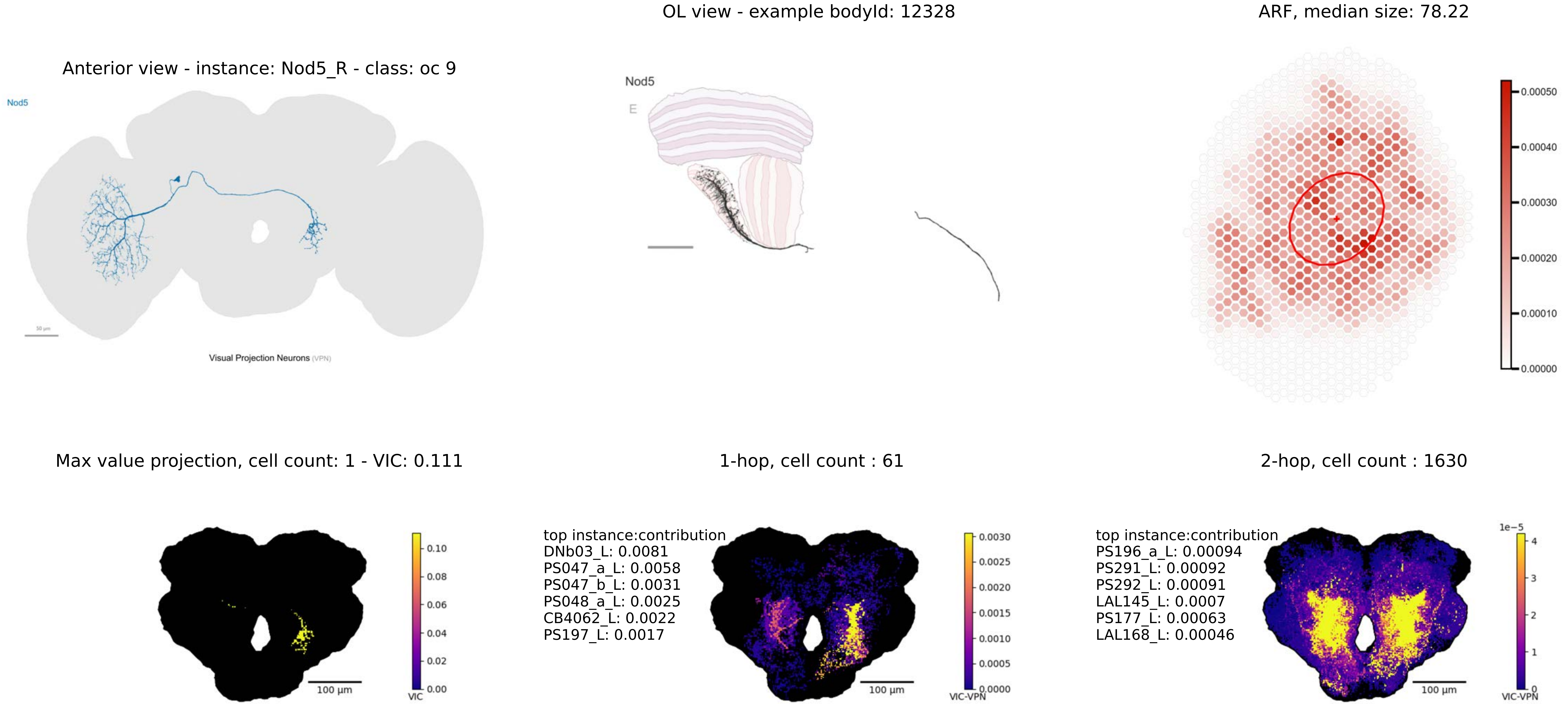
OL view - example bodyId: 12328
ARF, median size: 78.22
Anterior view - instance: Nod5_R - class: oc 9
Nod5
50 μm
Visual Projection Neurons (VPN)
Nod5
E
0.00050
0.00040
0.00030
0.00020
0.00010
0.00000
Max value projection, cell count: 1 - VIC: 0.111
1-hop, cell count : 61
2-hop, cell count : 1630
100 μm
VIC
0.10
0.08
0.06
0.04
0.02
0.00
top instance:contribution
DNb03_L: 0.0081
PS047_a_L: 0.0058
PS047_b_L: 0.0031
PS048_a_L: 0.0025
CB4062_L: 0.0022
PS197_L: 0.0017
VIC-VPN
0.0030
0.0025
0.0020
0.0015
0.0010
0.0005
0.0000
top instance:contribution
PS196_a_L: 0.00094
PS291_L: 0.00092
PS292_L: 0.00091
LAL145_L: 0.0007
PS177_L: 0.00063
LAL168_L: 0.00046
1e−5
4
3
2
1
0
VIC-VPN

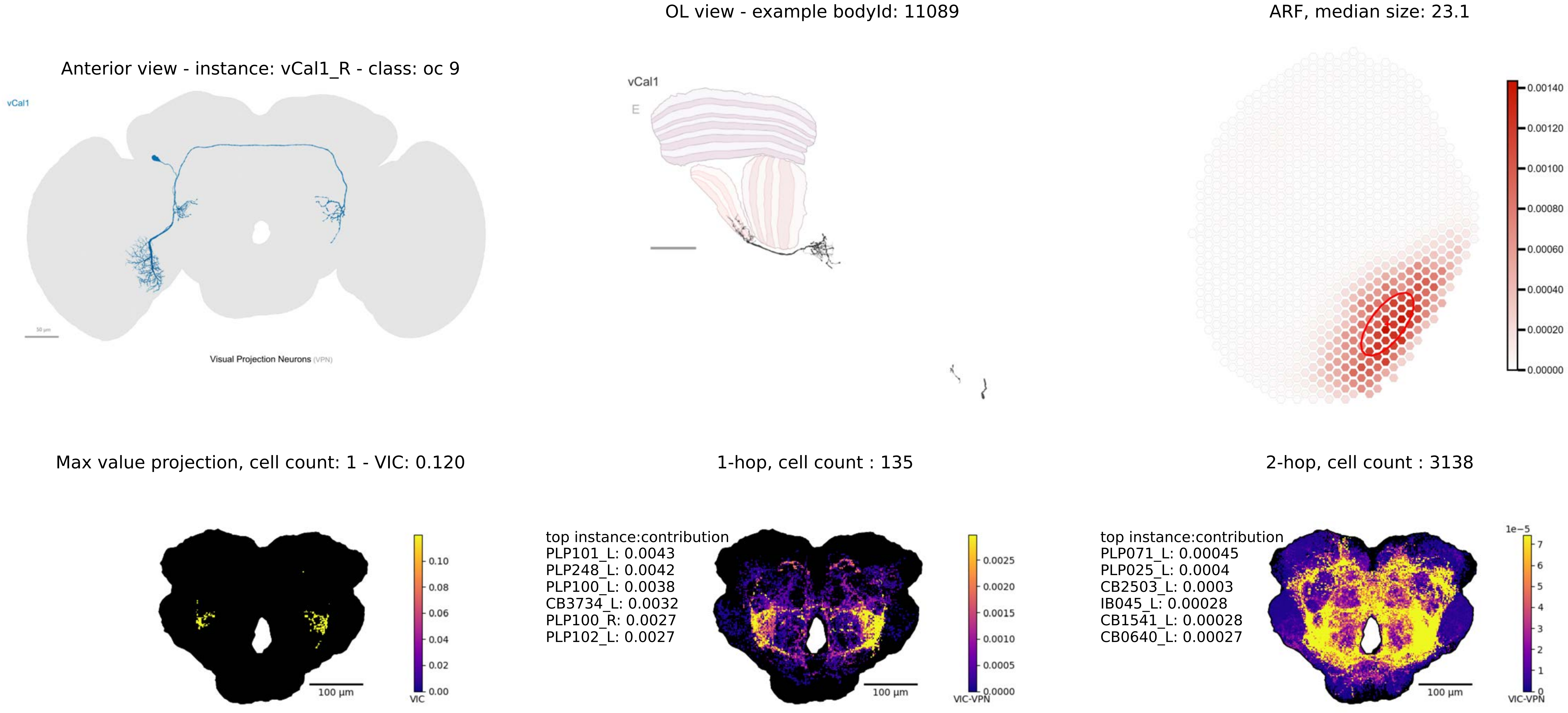

OL view - example bodyId: 11089
ARF, median size: 23.1
Anterior view - instance: vCal1_R - class: oc 9
vCal1
Visual Projection Neurons (VPN)
vCal1
E
0.00140
0.00120
0.00100
0.00080
0.00060
0.00040
0.00020
0.00000
Max value projection, cell count: 1 - VIC: 0.120
1-hop, cell count : 135
2-hop, cell count : 3138
100 µm
0.10
0.08
0.06
0.04
0.02
0.00
VIC
top instance:contribution
PLP101_L: 0.0043
PLP248_L: 0.0042
PLP100_L: 0.0038
CB3734_L: 0.0032
PLP100_R: 0.0027
PLP102_L: 0.0027
100 µm
0.0025
0.0020
0.0015
0.0010
0.0005
0.0000
VIC-VPN
top instance:contribution
PLP071_L: 0.00045
PLP025_L: 0.0004
CB2503_L: 0.0003
IB045_L: 0.00028
CB1541_L: 0.00028
CB0640_L: 0.00027
100 µm
1e−5
7
6
5
4
3
2
1
0
VIC-VPN

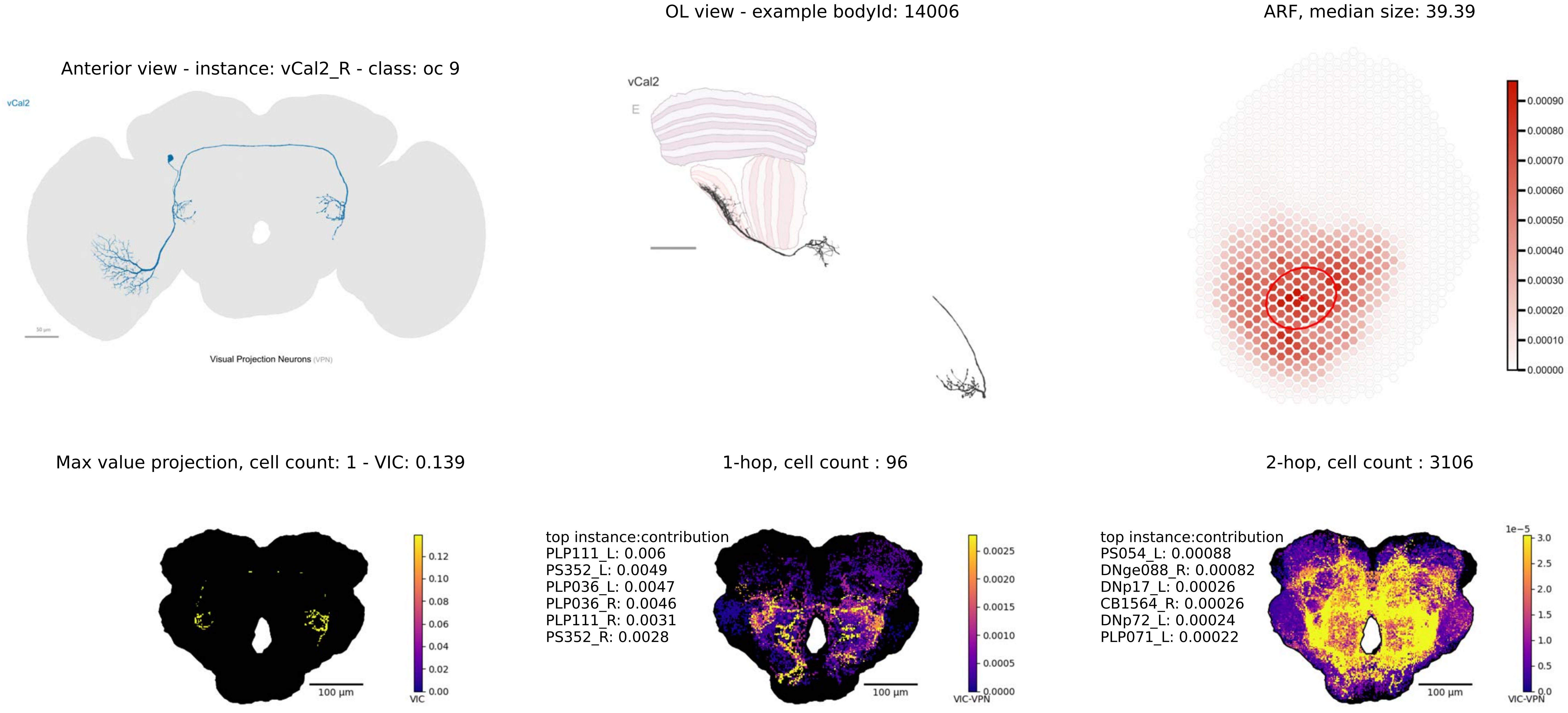
OL view - example bodyId: 14006
ARF, median size: 39.39
Anterior view - instance: vCal2_R - class: oc 9
vCal2
Visual Projection Neurons (VPN)
vCal2
E
Max value projection, cell count: 1 - VIC: 0.139
1-hop, cell count : 96
2-hop, cell count : 3106
top instance:contribution
PLP111_L: 0.006
PS352_L: 0.0049
PLP036_L: 0.0047
PLP036_R: 0.0046
PLP111_R: 0.0031
PS352_R: 0.0028
top instance:contribution
PS054_L: 0.00088
DNge088_R: 0.00082
DNp17_L: 0.00026
CB1564_R: 0.00026
DNp72_L: 0.00024
PLP071_L: 0.00022
100 µm
VIC
VIC-VPN
1e−5

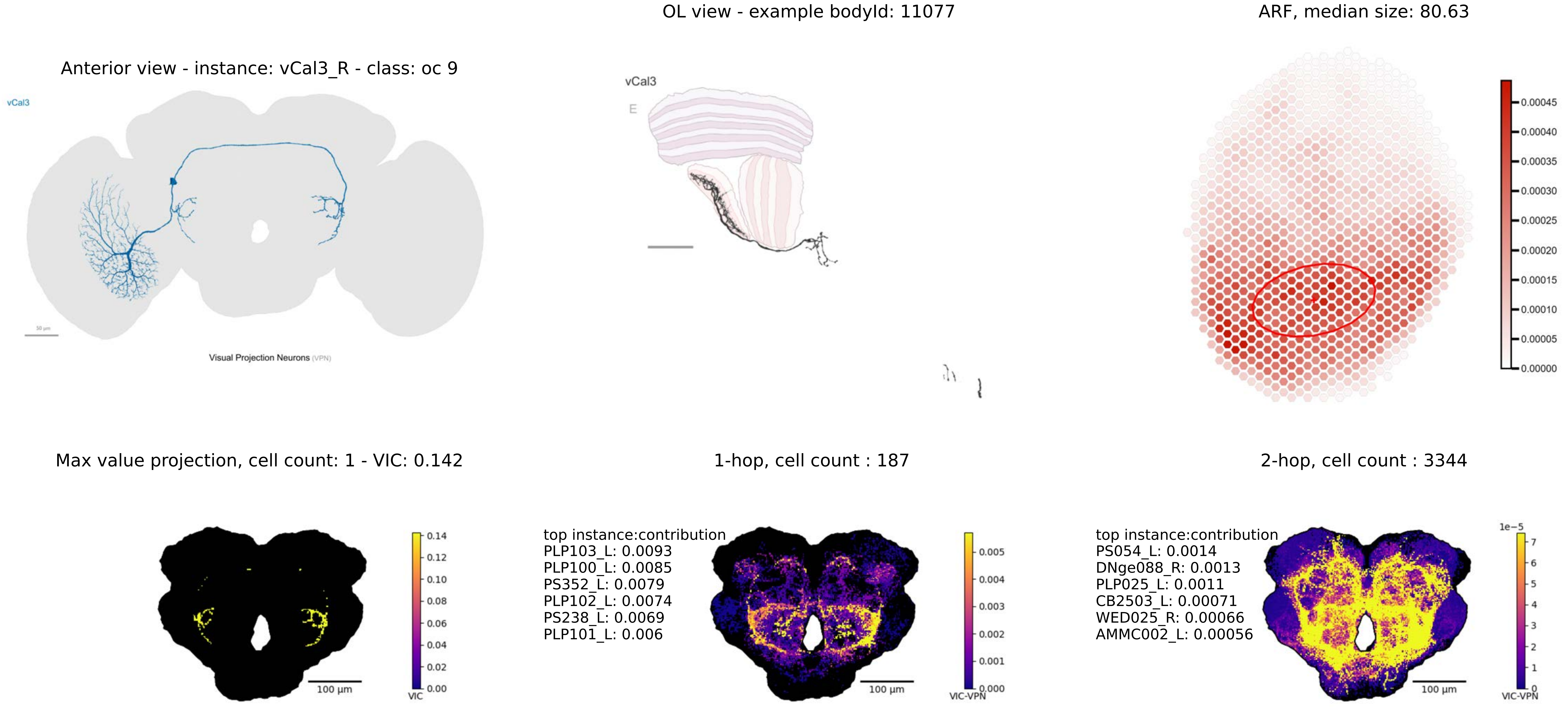
OL view - example bodyId: 11077
ARF, median size: 80.63
Anterior view - instance: vCal3_R - class: oc 9
vCal3
Visual Projection Neurons (VPN)
vCal3
E
0.00045
0.00040
0.00035
0.00030
0.00025
0.00020
0.00015
0.00010
0.00005
0.00000
Max value projection, cell count: 1 - VIC: 0.142
1-hop, cell count : 187
2-hop, cell count : 3344
100 µm
0.14
0.12
0.10
0.08
0.06
0.04
0.02
0.00
VIC
top instance:contribution
PLP103_L: 0.0093
PLP100_L: 0.0085
PS352_L: 0.0079
PLP102_L: 0.0074
PS238_L: 0.0069
PLP101_L: 0.006
100 µm
0.005
0.004
0.003
0.002
0.001
0.000
VIC-VPN
top instance:contribution
PS054_L: 0.0014
DNge088_R: 0.0013
PLP025_L: 0.0011
CB2503_L: 0.00071
WED025_R: 0.00066
AMMC002_L: 0.00056
100 µm
1e−5
7
6
5
4
3
2
1
0
VIC-VPN

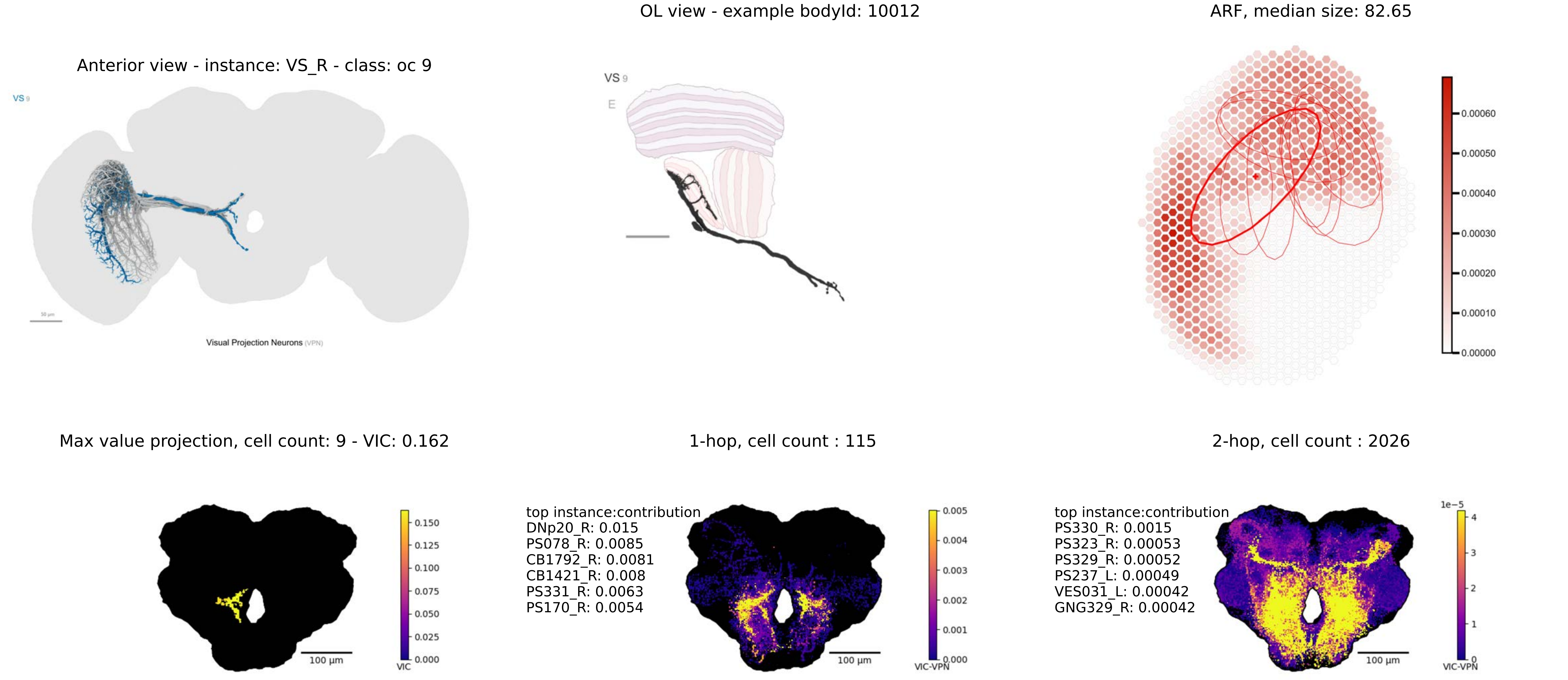

OL view - example bodyId: 10012
ARF, median size: 82.65
Anterior view - instance: VS_R - class: oc 9
VS 9
E
Visual Projection Neurons (VPN)
0.00060
0.00050
0.00040
0.00030
0.00020
0.00010
0.00000
Max value projection, cell count: 9 - VIC: 0.162
1-hop, cell count : 115
2-hop, cell count : 2026
0.150
0.125
0.100
0.075
0.050
0.025
0.000
VIC
100 µm
top instance:contribution
DNp20_R: 0.015
PS078_R: 0.0085
CB1792_R: 0.0081
CB1421_R: 0.008
PS331_R: 0.0063
PS170_R: 0.0054
0.005
0.004
0.003
0.002
0.001
0.000
VIC-VPN
100 µm
top instance:contribution
PS330_R: 0.0015
PS323_R: 0.00053
PS329_R: 0.00052
PS237_L: 0.00049
VES031_L: 0.00042
GNG329_R: 0.00042
1e−5
4
3
2
1
0
VIC-VPN
100 µm

Anterior view - instance: VSm_R - class: oc 9

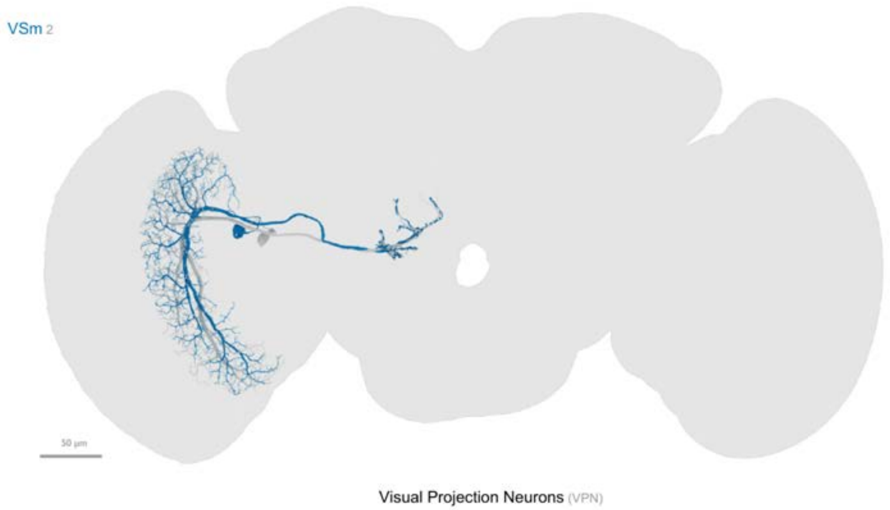


OL view - example bodyId: 11134

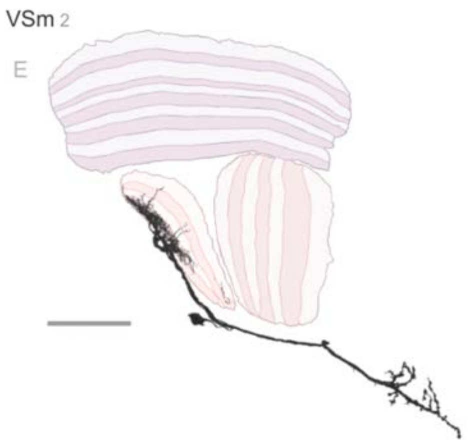


ARF, median size: 102.21

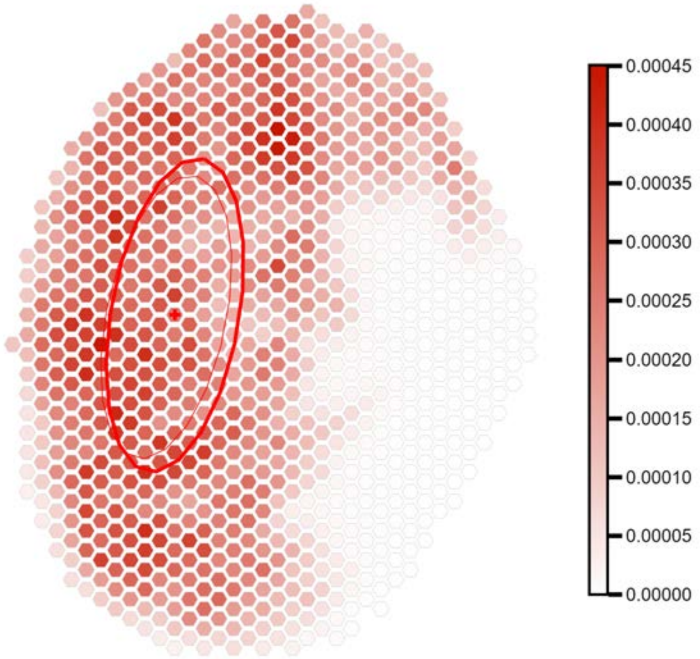


Max value projection, cell count: 2 - VIC: 0.150

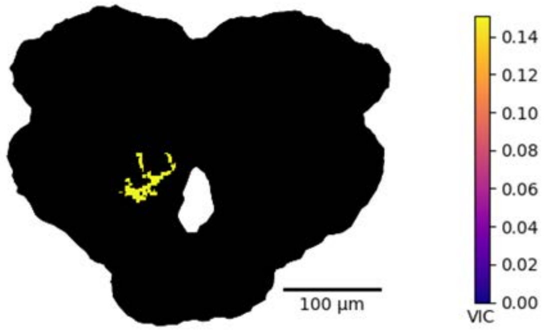


1-hop, cell count : 113

top instance:contribution
DNg46_R: 0.04
PS237_R: 0.023
PS213_R: 0.023
PS078_R: 0.022
DNge097_R: 0.016
CB4066_R: 0.013

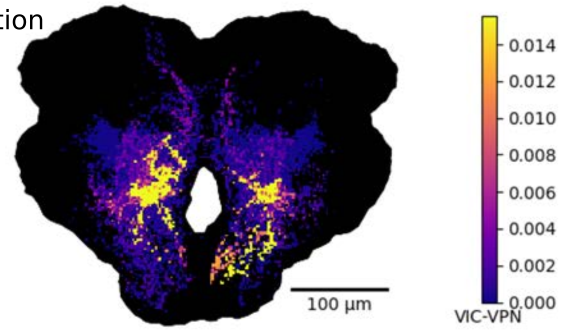


2-hop, cell count : 2459

top instance:contribution
DNp72_L: 0.0058
GNG520_L: 0.0055
GNG520_R: 0.005
PS352_L: 0.0032
DNp17_L: 0.0029
DNg58_L: 0.0029

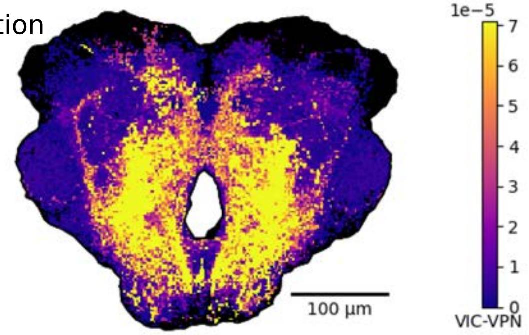

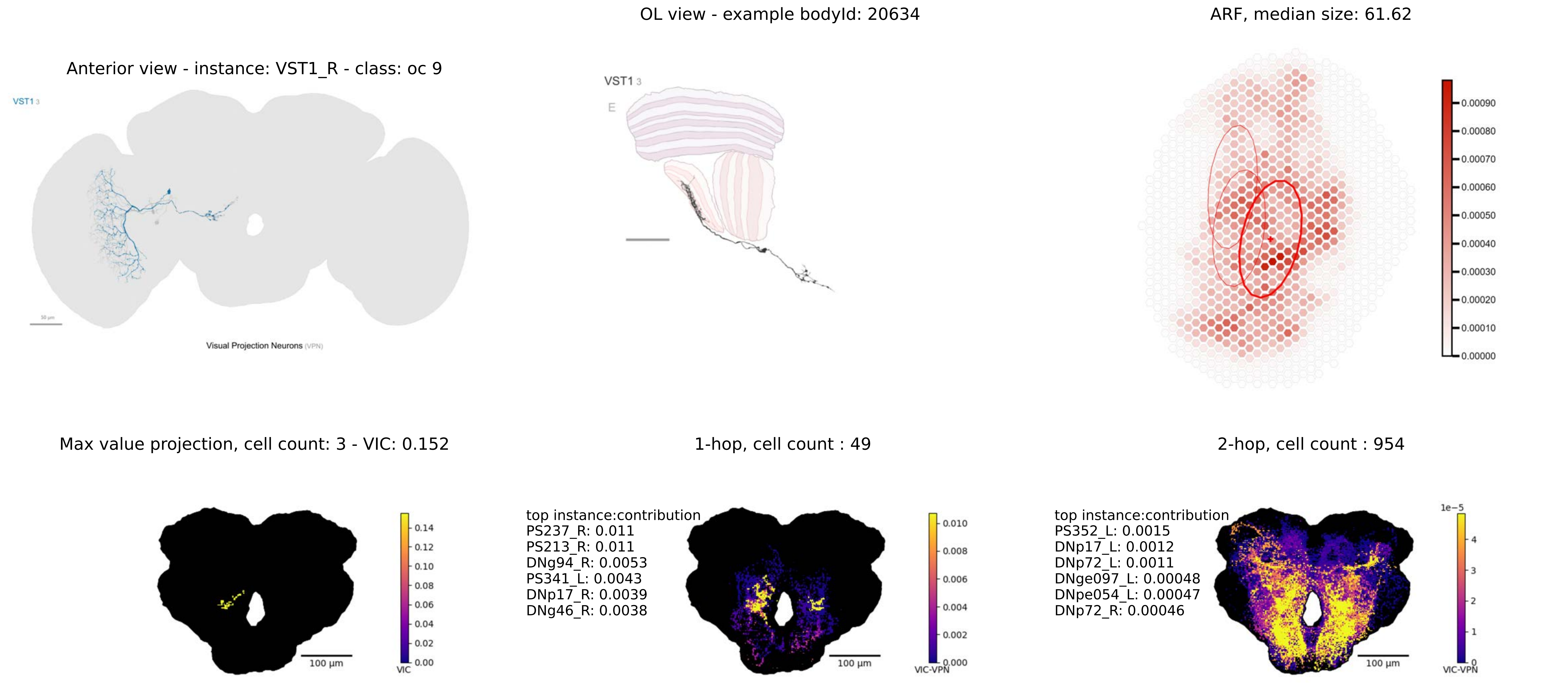

OL view - example bodyId: 20634
ARF, median size: 61.62
Anterior view - instance: VST1_R - class: oc 9
VST1 3
Visual Projection Neurons (VPN)
VST1 3
E
0.00090
0.00080
0.00070
0.00060
0.00050
0.00040
0.00030
0.00020
0.00010
0.00000
Max value projection, cell count: 3 - VIC: 0.152
1-hop, cell count : 49
2-hop, cell count : 954
0.14
0.12
0.10
0.08
0.06
0.04
0.02
0.00
VIC
100 µm
top instance:contribution
PS237_R: 0.011
PS213_R: 0.011
DNg94_R: 0.0053
PS341_L: 0.0043
DNp17_R: 0.0039
DNg46_R: 0.0038
0.010
0.008
0.006
0.004
0.002
0.000
VIC-VPN
100 µm
top instance:contribution
PS352_L: 0.0015
DNp17_L: 0.0012
DNp72_L: 0.0011
DNge097_L: 0.00048
DNpe054_L: 0.00047
DNp72_R: 0.00046
1e−5
4
3
2
1
0
VIC-VPN
100 µm

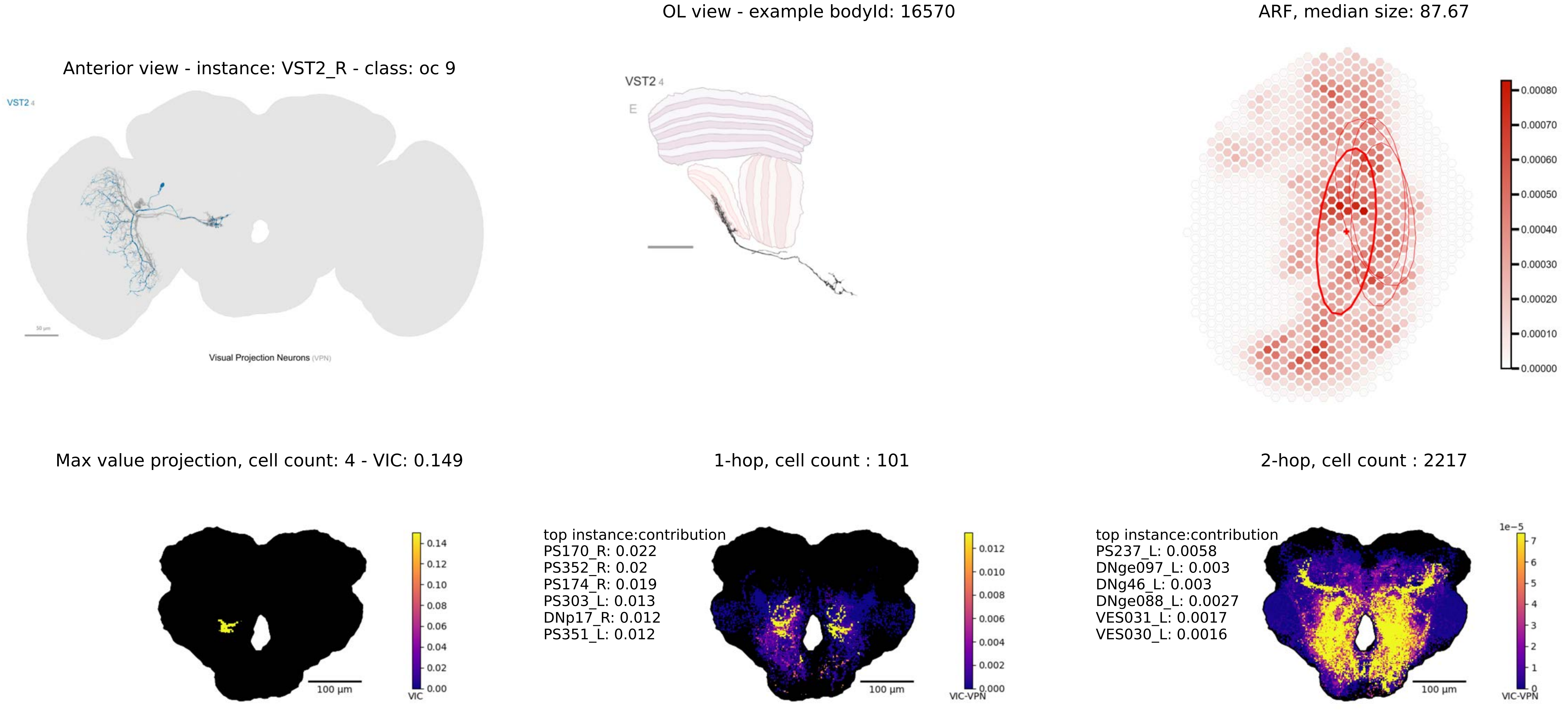
OL view - example bodyId: 16570
ARF, median size: 87.67
Anterior view - instance: VST2_R - class: oc 9
VST2 4
Visual Projection Neurons (VPN)
VST2 4
E
0.00080
0.00070
0.00060
0.00050
0.00040
0.00030
0.00020
0.00010
0.00000
Max value projection, cell count: 4 - VIC: 0.149
1-hop, cell count : 101
2-hop, cell count : 2217
100 µm
VIC
0.14
0.12
0.10
0.08
0.06
0.04
0.02
0.00
top instance:contribution
PS170_R: 0.022
PS352_R: 0.02
PS174_R: 0.019
PS303_L: 0.013
DNp17_R: 0.012
PS351_L: 0.012
100 µm
VIC-VPN
0.012
0.010
0.008
0.006
0.004
0.002
0.000
top instance:contribution
PS237_L: 0.0058
DNge097_L: 0.003
DNg46_L: 0.003
DNge088_L: 0.0027
VES031_L: 0.0017
VES030_L: 0.0016
100 µm
1e−5
7
6
5
4
3
2
1
0
VIC-VPN
